# To enrich or not to enrich: Enhancing (glyco)peptide ionization using the CaptiveSpray nanoBooster™

**DOI:** 10.1101/597922

**Authors:** Kathirvel Alagesan, Daniel Kolarich

**Author notes:** Corresponding Author: Dr. Kathirvel Alagesan, Institute for Glycomics, Building G26, Griffith University, Gold Coast campus, Southport, 4222 QLD, Australia, &.

## Abstract

The CaptiveSpray source ensures a stable spray and excellent nano ESI performance facilitated by a vortex gas that sweeps around the emitter spray tip to support liquid desolvation and focus the Taylor cone. Enriching the vortex gas with dopant solvents provides tremendous opportunities to increase ionization efficiency, in particular for hydrophilic compounds such as glycopeptides. How this CaptiveSpray nanobooster benefits their analysis, however, has to date not been systematically studied.

We evaluated various dopant solvents such as (i) acetone (ii) acetonitrile (iii) methanol (iv) ethanol and (v) isopropanol for their ability to enhance glycopeptide ionization. Using a synthetic IgG2 glycopeptide as a standard, acetonitrile provided a five-fold increase in signal intensities and resulted in an overall charge state increase compared to conventional CaptiveSpray ionization. This trend remained the same when tryptic IgG (glyco)peptides were analyzed and allowed highly sensitive detection of glycopeptides even without any enrichment. While acetone dopant gas enhanced glycopeptide ionization by doubling glycopeptide signal intensities, all other tested solvents resulted either in ion suppression or adduct formation. This is in agreement with and can be explained by their individual physio-chemical properties of the solvents. Finally, by omitting glycopeptide enrichment steps, we established a bias-free human Immunoglobulin G (IgG) subclass specific glycosylation profile applying the optimized CaptiveSpray nanoBooster nano-LC-ESI-MS/MS analysis conditions.

## INTRODUCTION

Glycosylation is one of the most complex yet universal posttranslational modifications (PTMs) that significantly enhances the functional diversity of proteins and influences their biological activity ^1^. Understanding the relationship between glycoprotein structure, glycosylation site location within a protein sequence and the function glycans fulfil on individual glycoproteins requires highly sensitive and selective methods that enable scientists to collect detailed information on primary structure aspects such as peptide sequence, glycan composition/structure and sites of glycosylation ^2^. Mass Spectrometry (MS) has emerged as the tool of choice to identify, characterize or quantify complex glycoproteins/proteins due to its universal applicability, sensitivity and high throughput capabilities ^3^. Its analytical capacities are further significantly increased by combining MS with orthogonal separation techniques such as ion-mobility MS ^4^, capillary electrophoresis ^5^ or nano-scale liquid chromatography (nano-LC) ^6^. The low flow rates used in nano-LC provide several advantages for glycoproteomics applications, in particular with respect to the ionization efficiency of hydrophilic compounds such as glycopeptides ^7 8^. The reduced flow rate and small orifice diameter significantly reduces the size of initially produced analyte droplets in nano-LC-ESI, thereby requiring less evaporation/fission cycles prior MS detection ^9–11^. Besides the impact of the droplet size on ionization, the charge state distribution of peptides and proteins also depends on the sample solvent and/or sheath gas ^12–13^.

However, during the ESI process, hydrophobic molecules tend to be more efficiently ionized compared to hydrophilic ones ^14^. Subsequently, glycopeptide signal strengths are significantly lower compared to their unmodified counterparts, mostly due to the presence of the hydrophilic glycan moiety ^7^. Glycan microheterogeneity is an additional factor hampering glycopeptide analyses as it reduces their global abundance in the entire peptide pool of a proteolytic digest ^15^. Consequently, glycopeptide enrichment has become a common sample preparation step in most glycoproteomic experiments to facilitate their detection and identification ^16–18^.

Despite increased sensitivity nano-LC-ESI ionization sources frequently suffer from issues such as unstable spray fostering the development of various variations of LC-emitter geometries to make nano-spray ESI plug and play like. The CaptiveSpray LC source emitter initially developed by Michrome and then further by Bruker represents one such development where a vortex gas (e.g. lab air) sweeps around the LC emitter spray, thereby concentrating and focusing the Taylor cone spray into the MS source. The CaptiveSpray emitter also allows for the gas that flows coaxially around the emitter to be enriched with a specific dopant solvent (known as CaptiveSpray nanoBooster™). The CaptiveSpray nanoBooster™ setup using acetonitrile (ACN) as a dopant has been described to significantly improve glycopeptide ionisation and thus facilitate their detection and analysis (personal communication with Dr. Kristina Marx, Bruker). Depending upon the used dopant, either charge stripping or supercharging of peptides and glycopeptides can be achieved during the ionization process. Dopant enriched nitrogen gas combined with sheatless CE-ESI-MS has been shown to improve sensitivity of glycopeptides, however requiring prior glycopeptide enrichment ^19^. However, to date a systematic evaluation on the influence of dopant solvent on CaptiveSpray nanoBooster™ ionization when omitting the glycopeptide enrichment step in nano-LC ESI MS/MS analysis is still lacking.

In the present study, we used a synthetic IgG2 subclass glycopeptide carrying a biantennary sialyated *N*-glycan to systematically investigate the ionization behavior using the CaptiveSpray nanoBooster™ in combination with various MS compatible solvents. We also investigated the influence the amount of organic LC eluent has on the ionization efficiency of glycopeptides during a nano-LC ESI MS/MS experiment and finally established the human IgG subclass glycosylation profile without any glycopeptide enrichment using the optimized CaptiveSpray nanobooster™ conditions.

## MATERIALS AND METHODS

### Materials

If not otherwise stated, all materials were purchased in the highest possible quality from Sigma-Aldrich (St. Louis, MO, USA). Trypsin (sequencing grade) was obtained from Roche Diagnostic GmbH (Mannheim, Germany). Water was used after purification with a Milli Q-8 direct system (Merck KGaA, Darmstadt, Germany). Human Immunoglobulin G (IgG) was obtained from BioreclamationIVT (New York, USA). The amino acid numbering applied for all proteins analyzed in this study is based on the respective UniProtKB entries.

### Glycopeptide synthesis

Solid Phase Glycopeptide Synthesis (SPGPS) was performed manually using 5-mL and 10-mL disposable polypropylene syringes with a bottom filter. All peptides and glycopeptides were synthesized by SPGPS using previously reported fluorenylmethoxycarbonyl (Fmoc) protocols ^7, 20–21^.

### In-solution protease digestion

10 µg of protein were reduced with 1 μL of 100 mM dithiothreitol (DTT) (in H_2_O) (99°C, 5 min) and then subsequently alkylated with 1 μL 500 mM iodoacetamide (IAA) solution (in H_2_O) at room temperature for 60 min in dark. Prior trypsin digestion, the samples were subjected to chloroform-methanol precipitation as described earlier ^22^. The protein pellet was resolubilized in 50 μL of 25 mM ammoniumbicarbonate and trypsin was added in a 1:30 ratio (enzyme:substrate). After overnight incubation at 37°C the resulting glycopeptide/peptide mixtures were dried in the speedvac without additional heating. The samples were stored at -25°C until further experiments.

### LC-MS Analysis Parameters

Nano-LC-ESI-MS analysis was carried out on an Ultimate 3000 RSLC-nano LC system (Dionex/Thermo Scientific, Sunnyvale, CA) coupled to an amaZon speed ETD ion trap mass spectrometer (IT-MS) equipped with the CaptiveSpray nanoBooster™ (both Bruker, Bremen, Germany). The dried glycopeptide sample was reconstituted in 200 µL of 0.1 % Trifluoroacetic acid (TFA). For each LC analysis glycopeptides corresponding to 150 ng (3 µL) were injected using the µL-pickup injection option.

In nano-LC mode peptides were concentrated on a C18 precolumn (Acclaim PepMap100™, Thermo, 100 µm x 20 mm, 5 µm particle size) and separated by reversed phase chromatography on a C18 analytical column (Acclaim PepMap™, Thermo, 75 µm x 15 cm, 3 µm particle size). The samples were loaded in 99 % loading buffer (0.1 % TFA) for 5 min on the precolumn at a flow rate of 5 µL/min before the captured peptides were subjected to reversed phase nanoLC at a flowrate of 400 nL/min on a column equilibrated in 95 % buffer A [0.1 % formic acid (FA)]. The gradient conditions were as follows: increase of buffer B (90 % acetonitrile containing 0.1 % formic acid) from 5 % to 45 % (6-36 min), further increase to 70 % B (36-38 min), followed by a steeper increase to 90 % B (40-42 min). The column was held at 90 % B for 10 min (42-52 min) before it was equilibrated in starting conditions for 13 min. The amaZon ETD speed ion trap was set-up to perform CID on the three most intense signals in every MS scan. An *m/z* range from 400-1600 was used for data dependent precursor scanning. The MS data was recorded using the instrument’s “enhanced resolution mode”. MS/MS data was acquired in “ultra-mode” over an *m/z* range from 100-2000. A detailed parameter setting is provided in the Supplementary Table S1 following MIRAGE ^23^ and MIPAE ^24^ recommendations.

Data analysis was performed using ProteinScape 4.0 (Bruker, Germany) and MASCOT 2.6 (MatrixScience, United Kingdom) using the following search parameters: Cysteine as carbamidomethyl was set as fixed modification, and oxidation (Met) were set as variable modifications. Up to two missed cleavages were allowed. Peptide tolerance was set at ±0.5 Da for MS and at ±0.5 Da for MS/MS. The data were searched against the SwissProt protein database (taxonomy restriction: *Homo sapiens*, SwissProt 2011_08; 531,473 sequences; 188,463,640 residues) and protein identification results are provided in Supplementary File F2.

## RESULTS AND DISCUSSION

### Rationale and study design

Glycoprotein-focused glycoproteomics aims to acquire comprehensive data on protein specific glycosylation micro and macro-heterogeneity ^7, 15^. For this purpose, glycopeptide enrichment is frequently applied to improve glycopeptide detection in the background of non-modified peptides in complex sample matrices. Hydrophilic Interaction Chromatography (HILIC) has extensively been applied for this purpose due to its low bias towards different glycan types. However, a recent systematic evaluation of the mobile phase effect on glycopeptide enrichment indicated that zwitterionic (ZIC)-HILIC glycopeptide efficiency primarily relied upon the used solvent ^25^. Thus, if the analytical conditions allow the detection of glycopeptides from complex samples without any enrichment steps, any enrichment-derived bias would be significantly reduced. Since glycopeptide ionization efficiency is one major limiting step for successful glycopeptide detection, we investigated how CaptiveSpray nanoBooster™ ionization can be employed for glycoproteomics when omitting the glycopeptide enrichment step. Using the CaptiveSpray nanoBooster™ system also enabled us to investigate how different dopant solvents added to the nitrogen vortex spray such as acetone, acetonitrile (ACN), methanol (MeOH), ethanol (EtOH), and isopropanol (IPA) influence glycopeptide ionization efficiency for subsequent enrichment-free glycoproteomics *(Supplementary Figure S1)*.

### Glycopeptide ionization enhancement depends upon the dopant solvent

First, we used a synthetic *N*-glycopeptide to investigate the effect various solvents have on glycopeptide ionization using CaptiveSpray nanoBooster™ system. A synthetic *N*-glycopeptide corresponding to the tryptic peptide sequence derived from IgG2 carrying a disialylated, bi-antennary *N-*glycan was analyzed using offline injection in a sample solvent containing 30 % ACN+0.1 % FA. The ion trap’s SPS target mass was set at *m/z* 1350 and any increase or decrease in the signal intensities was expressed in relation to the signal intensity that was observed without using the nanoBooster™ option. In addition to glycopeptide signal intensity, parameters such as background noise and adduct formation were considered to determine the most suitable CaptiveSpray nanoBooster™ ionization dopant solvent. The synthetic IgG2 *N*-glycopeptide was detected as doubly, triply and quadruply charged signals, but the individual charge state signal intensities were highly dopant solvent dependent (Figure 1). Using the CaptiveSpray ESI-MS ionization without any desolvation solvent or just nitrogen as a vortex gas, mainly triply charged signals were detected for the IgG2 synthetic *N*-glycopeptide (Figure 1). MeOH, EtOH and IPA as dopants all resulted in a significant charge stripping and thus ion intensity increase of the doubly charged precursor (Figure 1). Interestingly, potassium adduct formation for the triply charged ions (but not for the doubly charged ones) was observed when using these OH-group containing dopant organic solvents. In contrast, the use of acetone and ACN shifted the detected signals towards higher charge states of up to 25 % quadruply charged signals for ACN (Figure 1b). In ESI the charge state distribution of peptides and proteins is known to depend on the sample solvent and/or sheath gas ^9, 26^. In agreement with previous observations, we found that enhanced glycopeptide ionization and increased charge states were dependent on the physio-chemical properties of the solvents such as dielectric constant and surface tension. ACN exhibits the highest dielectric constant and surface tension of the tested solvents clearly supporting the notion that its physio-chemical properties are the major factors responsible for ACN’s ionization supportive features (Supplementary Table S3 & Figure 1).

**Figure 1:**
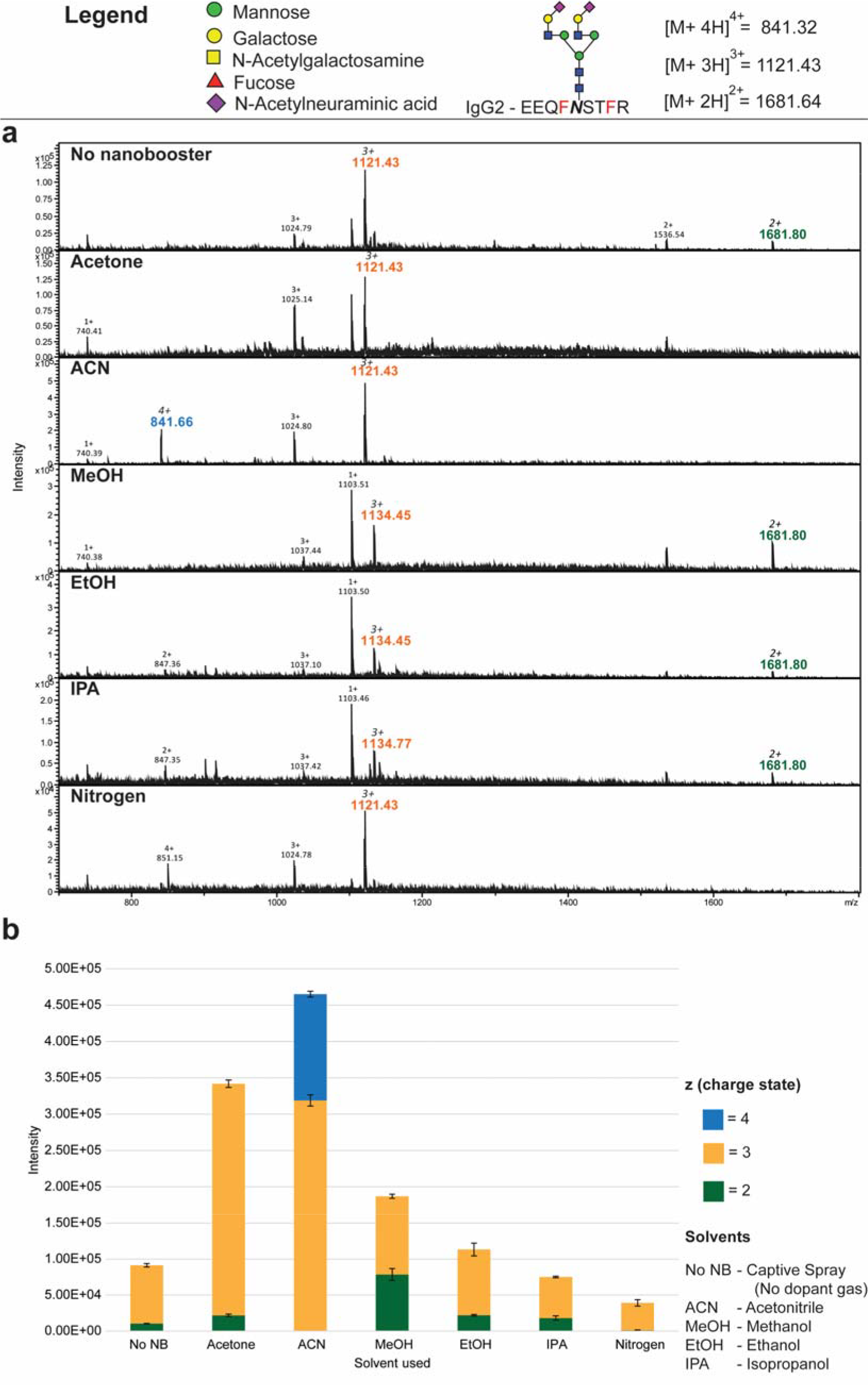
Influence different dopant solvents have on the signal intensity and charge state distribution of a synthetic IgG2 N-glycopeptide. **(a)** Representative summed spectra corresponding to 167 fmol of glycopeptide were obtained using different dopant solvents (spectra summed over 20 sec, 500 fmol/µL of synthetic *N*-glycopeptide, offline injection at 1 µL/min flowrate). When using MeOH, EtOH and IPA as dopant solvents the signal of a low molecular weight contaminant (*m/z* 1103.51) is significantly enhanced. In addition, cation adduct formation (*m/z* 1134.45) of the triply charged signals is drastically increased, further complicating glycopeptide analyses. Both, Acetone and ACN dopant solvents resulted in better signal intensities for the analyte and clearly reduced background noise. (**b**) Summary of the dopant solvent influence on CaptiveSpray nanoBooster™ ionization for an IgG2 synthetic *N*-glycopeptide. Acetone and ACN were the most suitable dopant solvents for increasing glycopeptide signal intensity. The signal intensities were summed up for all detected charge states from summed MS spectra (20 sec = 167 fmol of glycopeptide). Mean and standard deviation have been determined from triplicates. (*Quantitation analysis – See Supplementary Table S2 and For CID MSMS Spectra see supplementary Figure S2*).

The influence of the tested dopant solvents on glycopeptide ionization efficiency was determined by comparing the sum of the absolute signal intensities across all charge states. From the tested solvents ACN (5.0-fold), Acetone (3.74-fold), MeOH (2.05-fold) and EtOH (1.24-fold) enhanced glycopeptide signal intensities whereas IPA and nitrogen actually resulted in a decreased signal intensity (by 0.82-fold and 0.43-fold respectively, Figure 1b). Despite MeOH and EtOH having a higher dielectric constant in comparison to acetone, these solvents had an opposite effect on glycopeptide ionization efficiency. This could possibly be explained by the aprotic and protic properties of the solvents. The polar-protic solvents MeOH and EtOH are likely to participate in hydrogen bonding with the glycopeptide analyte molecules, thereby reducing their ionization efficiency. Even though glycopeptide ionization was marginally enhanced by MeOH and EtOH, their use as dopant solvents came with additional disadvantages such as increased adduct formation, a higher level of background noise especially in the lower mass region and an increased ionization of singly charged contaminant peaks (Figure 1a). All these factors made MeOH and EtOH as well as IPA unsuitable dopant solvents for glycoproteomics analyses using the CaptiveSpray nanoBooster™ ionization device. These results also clearly supported the hypothesis that the observed enhanced glycopeptide ionization was mainly dependent on the physio-chemical properties of the dopant solvent. In summary, using Acetone and ACN as dopant solvent resulted in higher signal intensities as well as lower background noise, hence these two solvents were selected for the further analyses.

### To enrich or not to enrich-CaptiveSpray nanobooster enhances glycopeptide ionization

In a recent study we demonstrated that the presence of reducing and alkylation agents significantly interfered with ZIC-HILIC based glycopeptide enrichment. More importantly, glycopeptide enrichment efficiency was strongly dependent on the used organic phase ^25^. Motivated by our dopant solvent-dependent glycopeptide ionization enhancement results, we evaluated the feasibility of CaptiveSpray nanoBooster™ ionization for unbiased (=enrichment free) glycoproteomics. First, we used our synthetic glycopeptide to evaluate any influence the organic solvent content had on ionization enhancement efficiency using three different ACN concentrations (10 %, 30 %, and 50 % ACN containing 0.1 % FA). Next, we used human IgG as a model protein and analyzed the tryptic (glyco)peptides without any prior enrichment using Reverse phase (RP)-CaptiveSpray nanoBooster™ nano-LC ESI-MS/MS analysis.

Independent of the sample organic solvent concentration both dopant solvents (ACN and acetone) enhanced glycopeptide ionization under offline and online MS conditions (Supplementary Figure S3; Figure 2 and Supplementary Table S4 and S5). Glycopeptides were generally detected in higher charge states when ACN was used as the dopant solvent. When injected in a solvent containing 10 % ACN with 0.1% FA the synthetic glycopeptide was mainly detected as a triply charged ion (~ 95%) when acetone was used as dopant solvent. In the presence of ACN as the dopant solvent, however, the triply charged signal decreased to 25% while the quadruply charged ions became the most intense signal (Supplementary Figure S3-A). Interestingly, these quadruply charged signals were observed only when ACN was used a dopant solvent. Also, the ratio of triply charged signal to that of quadruply charged signal was found to be dependent on the ACN concentration in the infusion solvent (Supplementary Figure S3-B-C). These data clearly indicated in addition to dopant solvent, the charge state enhancement of glycopeptides is also dependent on the organic solvent composition of the injection solution.

**Figure 2:**
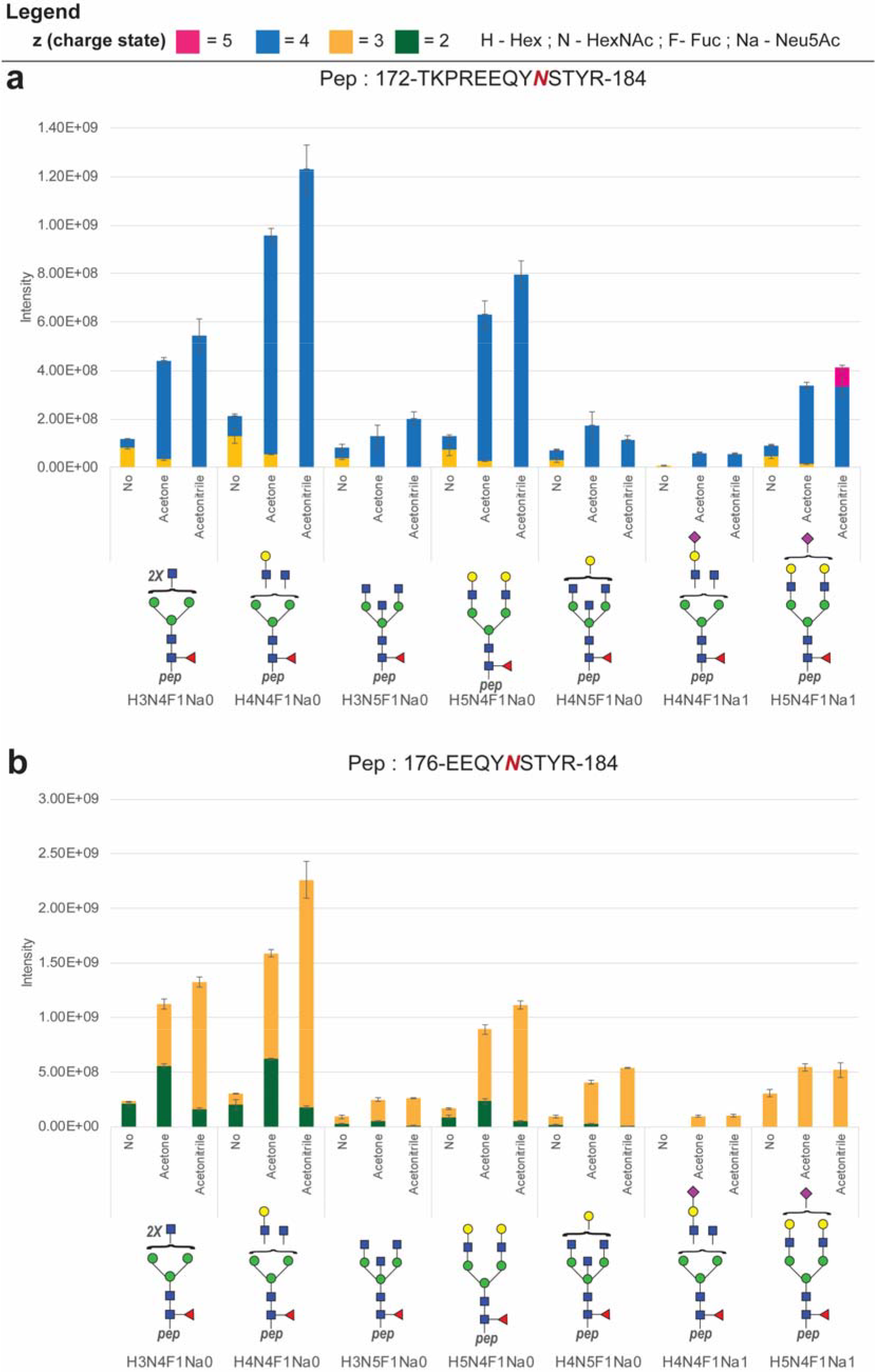
Influence of acetone and acetonitrile enriched dopant gas on tryptic Fc N-glycopeptides from Human IgG. **(a)** corresponding to the peptide backbone 172-184 TKPREEQYNSTYR and **(b)** 176-184 EEQYNSTYR in comparison to default CaptiveSpray nanoLC-ESI-MS/MS analysis. Error bars represent the standard deviation (n=3). The absolute abundances were determined using the area under the curve of extracted ion chromatograms (EIC’s) produced from all glycoforms and for each charge state signal detected for each single glycopeptide signal (*Quantitation results – See Supplementary file F1-Table S5-2*)

A similar effect was observed for the individual standard glycoprotein digests of IgG when analyzed without any glycopeptide enrichment (Figure 2). As reported earlier, IgG2 derived glycopeptides provided the most abundant signals followed by IgG1 and IgG4 ^25, 27^. However, the number of reported glycopeptides was higher than the previous study employing HILIC enrichment ^25^. Thus, is it evident that, use of CaptiveSpray nanoBooster™ is beneficial in glycoproteomics analysis reducing the sample preparation steps.

As observed for the synthetic IgG-2 glycopeptide, CaptiveSpray nanoBooster™ nano-LC-ESI-MS/MS analysis of human IgG exhibited a similar charge state enhancement when using ACN as dopant solvent. About 50 % of the ions for the IgG1 glycopeptide ^172^TKPREEQYNSTYR^184^ carrying the H5N4F1Na1 glycan were detected as [M + 4H]^4+^ when analyzed in a conventional CaptiveSpray setup (Figure 2A). This value increased to 95 % when acetone was used as dopant gas. When acetonitrile was used as dopant gas, about 19 % of the ions were detected as [M + 5H]^5+^ ions, which where otherwise not detected at all (Figure 2-A). A similar trend of charge state enhancement was observed for all the glycopeptides detected when using acetonitrile as a dopant solvent. Both dopant solvents acetone and acetonitrile provided better coverage of neutral and sialylated glycopeptides and similar glycosylation profiles if not better compared to the conventional setup. However, due to their higher intensities and supercharging properties, acetonitrile was selected for further experiments to establish human IgG subclass specific glycosylation profiles. IgG subclass glycovariants are known to orchestrate either pro- or anti-inflammatory effector pathways initiated via differential binding to Fcγ-receptors. In general, agalactosylated IgG antibodies promote inflammation, whereas galactosylation and terminal sialylation suppress inflammation ^28–30^. Therefore, we ventured to establish a subclass specific IgG glycosylation profile using commercially available IgG samples. The IgG samples were subjected to protease digestion after reduction and alkylation and analyzed by CaptiveSpray nanoBooster™ nano-LC-ESI-MS/MS using acetonitrile as dopant solvent. Overall, ten glycan compositions present on seven different peptide backbones were determined by CaptiveSpray nanoBooster™-RP nano-LC-ESI-MS/MS analysis. In total, 43 *N*-glycopeptides derived from IgG1-4 subclasses were identified and relatively quantified in addition to six previously reported *O*-glycopeptides derived from IgG3 (Supplementary Table S6). Next to the expected IgG3 glycopeptides, IgG2 subclass specific glycopeptides (*named as IgG3 variant in Figure 3a*) were present in the IgG3 sample due to a polymorphic variant frequent in populations of European ancestry ^31^. The majority of the identified glycoforms are bi-anntennary and core fucosylated *N*-glycans. Glycoforms lacking the core fucose were only detected as very low abundant signals of insufficient intensity for appropriate quantitation. A singly galactosylated, core fucosylated *N*-glycan was the most abundant glycoform in IgG1 and IgG2, while the doubly galactosylated, core fucosylated *N*-glycan was the most abundant glycoform in IgG3 (Figure 3). IgG4 carried a comparable heterogenous glycosylation with the non-galactosylated, core fucosylated *N*-glycan being slightly more abundant. While the IgG3 was the most homogenously glycosylated immunoglobulin exhibiting three glycoforms, the IgG3 variant was detected in five different glycoforms. IgG3 *N*-glycans did not exhibit any detectable levels of sialic acid, however the major *O*-glycoforms detected in the hinge region were found to be sialylated as in agreement with previous reports ^32^. On up to 20% of *N*-glycans one NeuAc residue was detected on all other IgGs (Figure 3-a/b).

**Figure 3:**
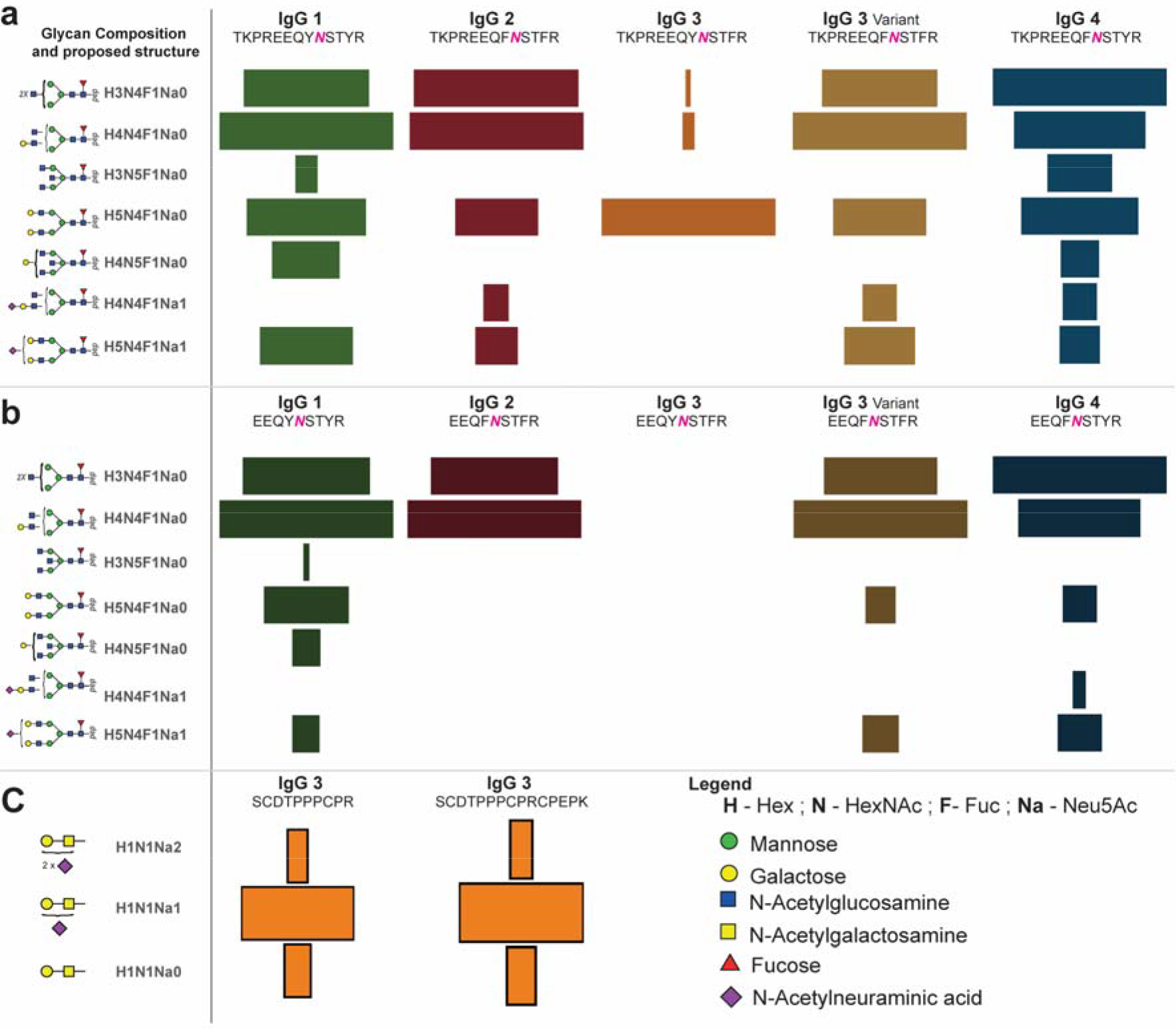
Relative quantitative glycoprofiles of the individual IgG 1-4 subclasses. Glycopeptides were analyzed using CaptiveSpray nanoBooster™ ionization using acetonitrile as dopant solvent. **(a)** *N*-glycoprofile with of glycopeptide containing one missed cleavage and **(b)** No-miss cleavage of glycopeptide. **(c)** *O*-glycoprofile determined for the IgG3 glycopeptide with and without a missed cleavage (*for detailed quantitation results – see supplementary file F1 – Table S61 to S6-4*).

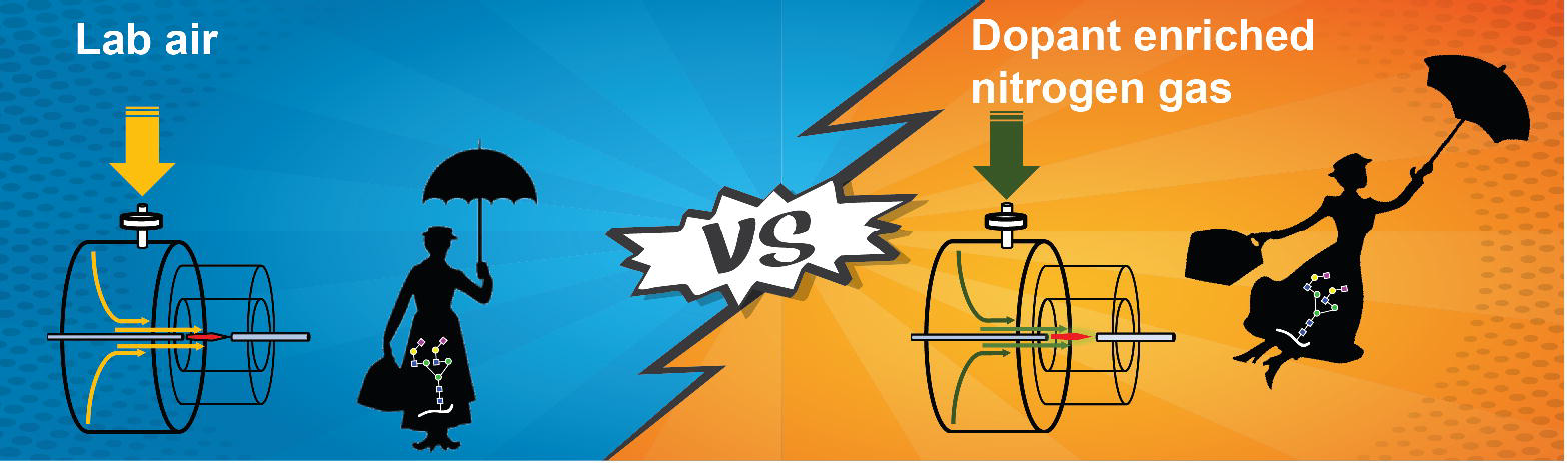

## CONCLUSIONS

The availability of a synthetic *N*-glycopeptide standard provided an opportunity to systematically evaluate glycopeptide analysis methods and fostered the development of a glycoproteomics approach that is employing a novel ionization interface, the CaptiveSpray nanoBooster™. The opportunity to use ACN as dopant solvent during the ionization process provides clear advantages for glycopeptide analyses by enhancing the detectable charge states and ionization efficiency of hydrophilic compounds such as glycopeptides. This allows for the elimination of specific enrichment steps. The ability to “supercharge” glycopeptide ion precursors was found to depend on the used dopant solvent and can provide an important advantage for ETD-fragmentation analyses of glycopeptides. In summary, we demonstrated that glycopeptide ionization step plays a crucial role in glycoproteomics and that this can be significantly enhanced using dopant solvents in vortex spray of ionization sources such as the CaptiveSpray nanoBooster™.

## ACKNOWLEDGEMENTS

We thank the Beilstein-Institut for supporting KA with a PhD scholarship and the Max Planck Society for financial support. DK is the recipient of an Australian Research Council Future Fellowship (project number FT160100344) funded by the Australian Government.

