## Supplementary Table for "To enrich or not to enrich: Enhancing (glyco)peptide ionization using the CaptiveSpray nanoBooster™"

*Corresponding Author: Dr. Kathirvel Alagesan

T +61 7 5552 7026

F +61 7 5552 9040

 &

### Supplementary Figures

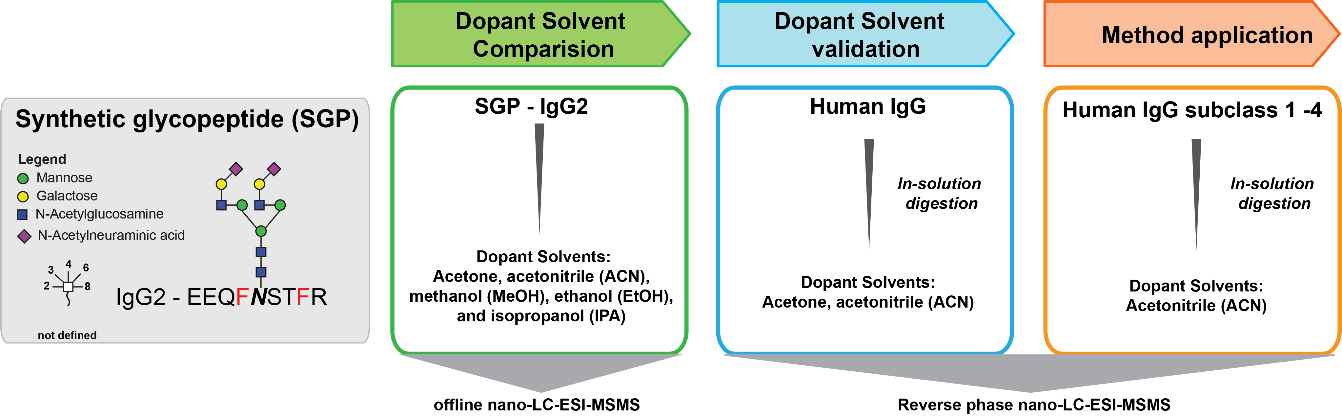

Supplementary Figure S1: Experimental design – systematic evaluation of glycopeptide ionization behavior in CaptiveSpray nanoBooster™ using various dopant solvents [(i) Acetone (ii) Acetonitrile (iii) Methanol (iv) Ethanol (v) Isopropanol] for unbiased glycoproteomics using synthetic IgG 2 subclass tryptic glycopeptide carrying a biantennary N-glycan. The optimal solvent determined was then used to establish the IgG subclass specific glycosylation profile without any prior enrichment step using CaptiveSpray nanoBooster™ RP-nLC-ESI-MSMS.

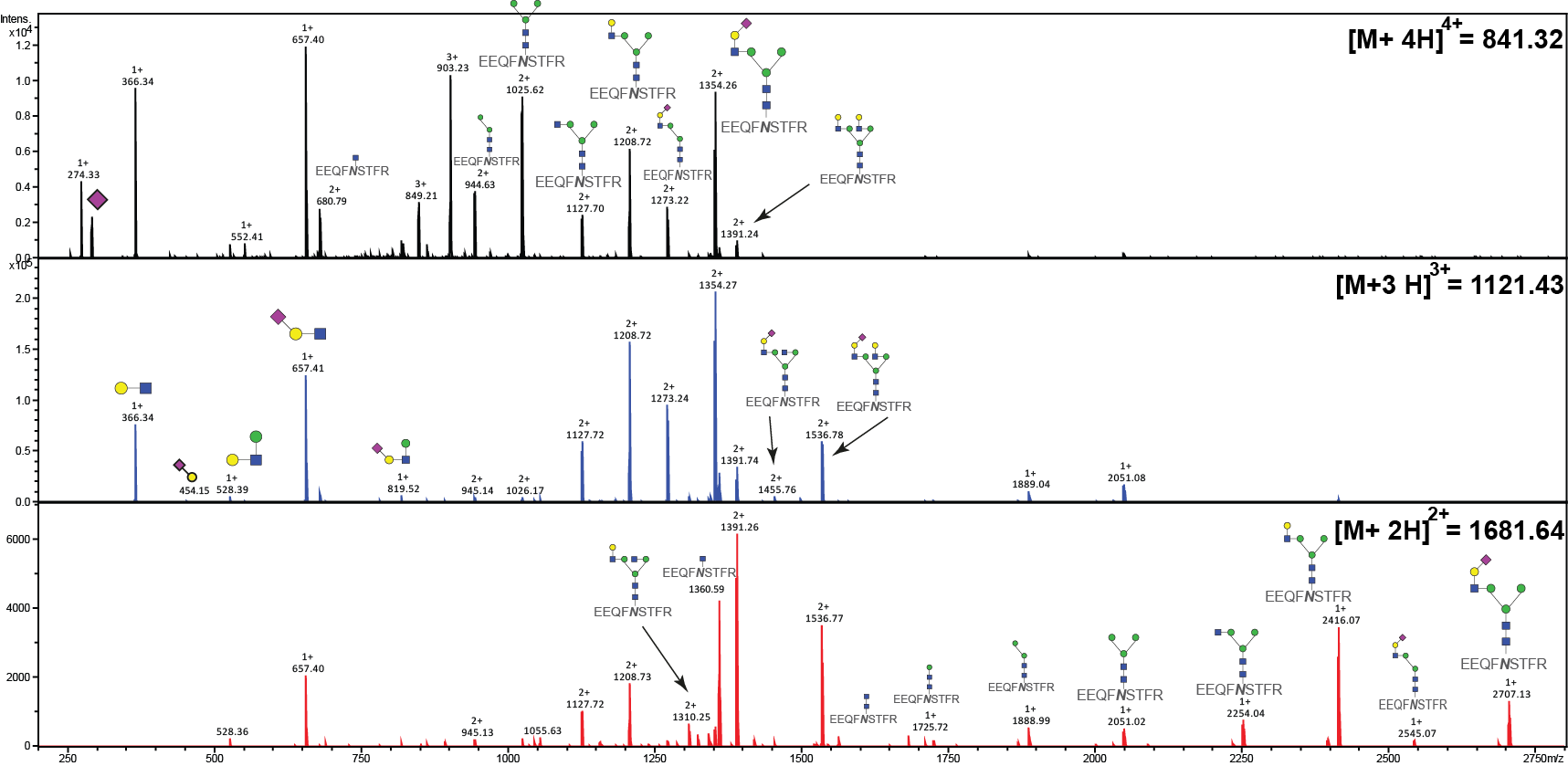

Supplementary Figure S2: CID tandem MS spectrum of a synthetic glycopeptide corresponding to tryptic peptide sequence derived from human IgG 2 carrying a biantennary N-glycan.

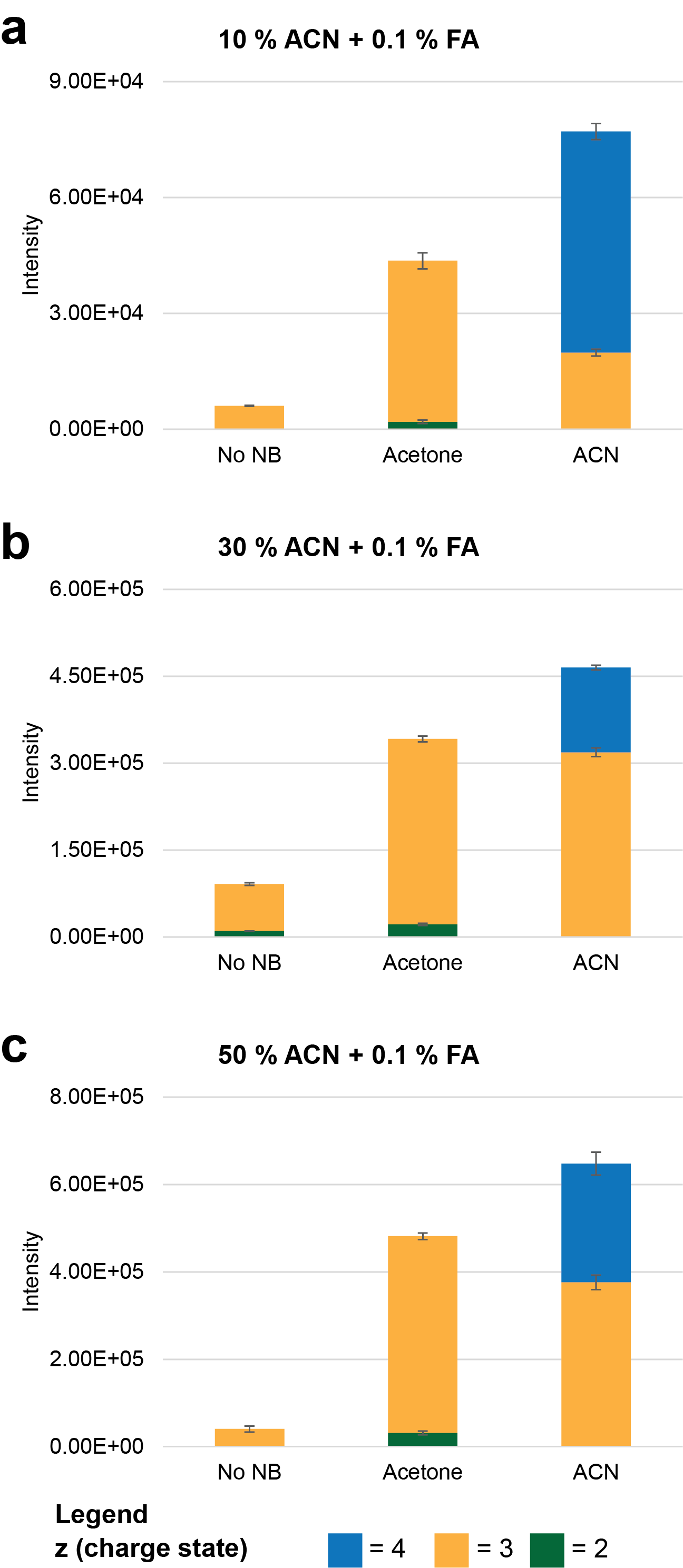

*Supplementary Figure S3: Effect of percent organic solvent composition on glycopeptide ionization using CaptiveSpray nanoBooster™ system*. Representative summed spectra obtained using different dopant solvents (spectra summed over 20 sec, 500 fmol/µL of synthetic N-glycopeptide, offline injection at 1 µL/min flowrate) at various percentage organic solvent composition (A) 10% ACN + 0.1 % FA (B) 30% ACN + 0.1 % FA (C) 50% ACN + 0.1 % FA. Independent of organic solvent composition in which the analyte is present both Acetone and ACN enhanced glycopeptide ionization.

### Supplementary Tables

**Supplementary Table S1:** Ion trap settings applied in this study, following the MIRAGE guidelines ([www.beilstein-mirage.org](http://www.beilstein-mirage.org))

| **MS Settings - General** |  |
| --- | --- |
| ESI probe | CaptiveSpray™ |
| Capillary voltage | 1.3 kV |
| SPS | *m/z* 900 for LC-MS analysis |
| Compound stability | 100% |
| Trap Drive Level | 100% |
| Spectra averaging | 5 |
| Dry gas temperature | 150°C |
| Dry Gas flow | 3 L/min |
| Maximum accumulation time | 200 ms |
| Ion mode | positive |
| **MS-Scan** |  |
| MS Scan mode | Enhanced scan |
| ICC target | 200000 |
| Mass detection range | *m/z* 400-1600 |
| **MS2** |  |
| MS scan mode | ultrascan |
| SPS MS(n) | automatic |
| MS(n) spectra averages | 5 |
| MS(n) ICC target | 40000 |
| Preferred charge state | ≥Doubly, |
| Active exclusion | Off |
| Mass detection range | *m/z* 100-2000 |
| Isolation width | 3 Da |
| Exclude singly charged ions | on |
| No. of precursor ions | 3 |
| SmartFrag | Enhanced |
| SmartFrag Start amplitude | 30% |
| SmartFrag End amplitude | 120% |
| Fragmentation width | 5 *m/z* |

***Supplementary Table S3:*** Physiochemical properties of the used solvents

| **Solvent** | **Solvent Polarity** | **Dielectric Constant** | **Surface tension @ 20°C in mN/m** |
| --- | --- | --- | --- |
| Acetone | Polar - Aprotic | 21 | 25.20 |
| ACN | Polar - Aprotic | 37.5 | 29.1 |
| MeOH | Polar - Protic | 32.6 | 22.70 |
| EtOH | Polar - Protic | 24.3 | 22.10 |
| IPA | Polar - Protic | 18 | 23.00 |
| Water | Polar - Protic | 78.5 | 72.80 |
| Nitrogen |  | ~1 |  |

Surface tension values were obtained from http://www.surface-tension.de/ and dielectric constants information retrieved from http://chem.libretexts.org/Core/Organic_Chemistry/Fundamentals/
Intermolecular_Forces/Polar_Protic_and_Aprotic_Solvents on 30/03/2019

**Quatitation results available in separate excel sheet (SUPPLEMENTARY FILE-F1)**

***Supplementary Table S2:* Quantitation results related to the figure 1-b in the main text**

***Supplementary Table S4:* Quantitation results for the** Figure S3 in the supplementary file

***Supplementary Table S5-1 & 2 :* Quantitation results for the** Figure 2 in the main text

***Supplementary Table S6-1 to 4:* Quantitation results for the** Figure 3 in the main text

**ProteinScape results (SUPPLEMENTARY FILE-F2)**
