## supplementary File F2 for "To enrich or not to enrich: Enhancing (glyco)peptide ionization using the CaptiveSpray nanoBooster™"

### Protein Report
### IonTrap_Human_Swiss_CID_2019-03-05 17:55:08

BRUKER

#### Project Info

**Name:** Alagesan
**Note:** Nanobooster
**Date:** Mar 5, 2019

#### Sample Info & Protocols

**Name:** Nanobooster - IgG
**Date:** Mar 5, 2019

#### Search Result Info

**Search Result:** IonTrap_Human_Swiss_CID_2019-03-05 17:55:08
**Location:** /Alagesan/Nanobooster - IgG/Ka_IgG Trypsin No Enrich No NB_BA5_01_8724.mgf
**Search Method:** IonTrap_Human_Swiss_CID
**Version:** 1.2
**Search Engine(s):** Mascot, 2.6.0
**Ident. Compound(s):** 0/2206
**Database(s):** SwissProt, SwissProt_2017_04.fasta
**Note:**

#### Results

##### Protein 1: Immunoglobulin gamma-1 heavy chain OS=Homo sapiens PE=1 SV=1

| | | | |
|---|---|---|---|
| **Accession:** | IGG1_HUMAN | **Score:** | 1074.4 |
| **Database:** | SwissProt | **Seq. Coverage [%]:** | 38.8 |
| **MW [kDa] / pI:** | 49.3 / 8.9 | **No. of Peptides:** | 15 |
| **Modification(s):** | Carbamidomethyl, Oxidation | | |

| 10 | 20 | 30 | 40 | 50 | 60 | 70 | 80 | 90 | 100 |
|---|---|---|---|---|---|---|---|---|---|
| QVQLVQSGGG | VVQPGRSLRL | SCAASGFTFS | RYTIHWVRQA | PGKGLEWVAV | MSYBGBBKHY | ADSVNGRFTI | SRNDSKNTLY | LNMNSLRPED | TAVYYCARIR |
| **110** | **120** | **130** | **140** | **150** | **160** | **170** | **180** | **190** | **200** |
| DTAMFFAHWG | QGTLVTVSSA | STKGPSVFPL | APSSKSTSGG | TAALGCLVKD | YFPEPVTVSW | NSGALTSGVH | TFPAVLQSSG | LYSLSSVVTV | PSSSLGTQTY |
| **210** | **220** | **230** | **240** | **250** | **260** | **270** | **280** | **290** | **300** |
| ICNVNHKPSN | TKVDKKVEPK | SCDKTHTCPP | CPAPELLGGP | SVFLFPPKPK | DTLMISRTPE | VTCVVVDVSH | EDPEVKFNWY | VDGVEVHNAK | TKPREEQYNS |
| **310** | **320** | **330** | **340** | **350** | **360** | **370** | **380** | **390** | **400** |
| TYRVVSVLTV | LHQDWLNGKE | YKCKVSNKAL | PAPIEKTISK | AKGQPREPQV | YTLPPSRDEL | TKNQVSLTCL | VKGFYPSDIA | VEWESNGQPE | NNYKTTPPVL |
| **410** | **420** | **430** | **440** | **450** | | | | | |
| DSDGSFFLYS | KLTVDKSRWQ | QGNVFSCSVM | HEALHNHYTQ | KSLSLSPGK | | | | | |

MS/MS Peptide Matches

| Cmpd. | m/z meas. | z | Δ m/z [ppm] | Δ m/z [Da] | Rt [min] | Score | No. of Cmpds. | Site [%] | P | Sequence | Modification | Range | Type |
|---|---|---|---|---|---|---|---|---|---|---|---|---|---|
| 771 | 593.894 | 2 | 112.89 | 0.0670 | 27.02 | 57.09 | 12 | 0.0 | 0 | K.GPSVFPLAPSSK.S | | 124-135 | CID |
| 678 | 661.43 | 2 | 132.08 | 0.0873 | 24.95 | 132.32 | 17 | 0.0 | 0 | K.STSGGTAALGCLVK.D | Carbamidomethyl: 11 | 136-149 | CID |
| 1000 | 834.754 | 4 | 405.09 | 0.3380 | 31.85 | 81.3 | 1 | 0.0 | 1 | K.SCDKTHTCPPCPAPELLGGPSVFLFPPKPK.D | Carbamidomethyl: 2, 8, 11 | 221-250 | CID |
| 1059 | 949.265 | 3 | 464.75 | 0.4410 | 32.99 | 89.33 | 34 | 0.0 | 0 | K.THTCPPCPAPELLGGPSVFLFPPKPK.D | Carbamidomethyl: 4, 7 | 225-250 | CID |
| 524 | 418.263 | 2 | 101.03 | 0.0423 | 21.98 | 47.69 | 6 | 0.0 | 0 | K.DTLMISR.T | | 251-257 | CID |
| 417 | 426.262 | 2 | 102.76 | 0.0438 | 19.69 | 53.68 | 3 | 0.0 | 0 | K.DTLMISR.T | Oxidation: 4 | 251-257 | CID |

| Cmpd. | m/z meas. | z | Δ m/z [ppm] | Δ m/z [Da] | Rt [min] | Score | No. of Cmpds. | Site [%] | P | Sequence | Modification | Range | Type |
|---|---|---|---|---|---|---|---|---|---|---|---|---|---|
| 805 | 839.46 | 2 | 65.97 | 0.0554 | 27.66 | 117.4 | 11 | 0.0 | 0 | K.FNWYVDGVEVHNAK.T | | 277-290 | CID |
| 1114 | 904.608 | 2 | 111.79 | 0.1011 | 34.20 | 90.66 | 13 | 0.0 | 0 | R.VVSVLTVLHQDWLNGK.E | | 304-319 | CID |
| 492 | 419.83 | 2 | 178.01 | 0.0747 | 21.37 | 30.25 | 2 | 0.0 | 0 | K.ALPAPIEK.T | | 329-336 | CID |
| 604 | 643.898 | 2 | 89.15 | 0.0574 | 23.55 | 40.06 | 9 | 0.0 | 0 | R.EPQVYTLPPSR.D | | 347-357 | CID |
| 647 | 937.042 | 2 | 56.86 | 0.0533 | 24.35 | 56.09 | 9 | 0.0 | 1 | R.EPQVYTLPPSRDELTK.N | | 347-362 | CID |
| 756 | 581.394 | 2 | 129.97 | 0.0756 | 26.72 | 69.41 | 15 | 0.0 | 0 | K.NQVSLTCLVK.G | Carbamidomethyl: 7 | 363-372 | CID |
| 1011 | 937.574 | 2 | 116.75 | 0.1095 | 32.07 | 106.79 | 27 | 0.0 | 0 | K.TTPPVLDSDGSFFLYSK.L | | 395-411 | CID |
| 166 | 409.798 | 2 | 141.16 | 0.0578 | 13.55 | 44.26 | 3 | 0.0 | 1 | K.LTVDKSR.W | | 412-418 | CID |
| 726 | 934.552 | 3 | 133.53 | 0.1248 | 26.09 | 58.09 | 12 | 0.0 | 0 | R.WQQGNVFSCSVMHEALHNHYTQK.S | Carbamidomethyl: 9 | 419-441 | CID |

#### Protein 2: Immunoglobulin heavy constant gamma 2 OS=Homo sapiens GN=IGHG2 PE=1 SV=2

**Accession:** IGHG2_HUMAN
**Database:** SwissProt
**MW [kDa] / pI:** 35.9 / 7.7
**Modification(s):** Carbamidomethyl, Oxidation

**Score:** 1016.8
**Seq. Coverage [%]:** 48.8
**No. of Peptides:** 16

| 10 | 20 | 30 | 40 | 50 | 60 | 70 | 80 | 90 | 100 |
|---|---|---|---|---|---|---|---|---|---|
| ASTKGPSVFP | LAPCSRSTSE | STAALGCLVK | DYFPEPVTVS | WNSGALTSGV | HTFPAVLQSS | GLYSLSSVVT | VPSSNFGTQT | YTCNVDHKPS | NTKVDKTVER |
| **110** | **120** | **130** | **140** | **150** | **160** | **170** | **180** | **190** | **200** |
| KCCVECPPCP | APPVAGPSVF | LFPPKPKDTL | MISRTPEVTC | VVVDVSHEDP | EVQFNWYVDG | VEVHNAKTKP | REEQFNSTFR | VVSVLTVVHQ | DWLNGKEYKC |
| **210** | **220** | **230** | **240** | **250** | **260** | **270** | **280** | **290** | **300** |
| KVSNKGLPAP | IEKTISKTKG | QPREPQVYTL | PPSREEMTKN | QVSLTCLVKG | FYPSDISVEW | ESNGQPENNY | KTTPPMLDSD | GSFFLYSKLT | VDKSRWQQGN |
| **310** | **320** | **330** | | | | | | | |
| VFSCSVMHEA | LHNHYTQKSL | SLSPGK | | | | | | | |

MS/MS Peptide Matches

| Cmpd. | m/z meas. | z | Δ m/z [ppm] | Δ m/z [Da] | Rt [min] | Score | No. of Cmpds. | Site [%] | P | Sequence | Modification | Range | Type |
|---|---|---|---|---|---|---|---|---|---|---|---|---|---|
| 818 | 644.407 | 2 | 120.52 | 0.0777 | 27.99 | 74.43 | 22 | 0.0 | 0 | K.GPSVFPLAPCSR.S | Carbamidomethyl: 10 | 5-16 | CID |
| 689 | 712.414 | 2 | 77.91 | 0.0555 | 25.26 | 85.41 | 16 | 0.0 | 0 | R.STSESTAALGCLVK.D | Carbamidomethyl: 11 | 17-30 | CID |
| 117 | 423.783 | 2 | 107.10 | 0.0454 | 12.21 | 23.77 | 1 | 0.0 | 1 | K.VDKTVER.K | | 94-100 | CID |
| 941 | 1013.183 | 3 | 341.55 | 0.3459 | 30.53 | 83.88 | 4 | 0.0 | 1 | R.KCCVECPPCPAPPVAGPSVFLFPPKPK.D | Carbamidomethyl: 2, 3, 6, 9 | 101-127 | CID |
| 1031 | 970.542 | 3 | 415.67 | 0.4033 | 32.46 | 91.42 | 12 | 0.0 | 0 | K.CCVECPPCPAPPVAGPSVFLFPPKPK.D | Carbamidomethyl: 1, 2, 5, 8 | 102-127 | CID |
| 524 | 418.263 | 2 | 101.03 | 0.0423 | 21.98 | 47.69 | 6 | 0.0 | 0 | K.DTLMISR.T | | 128-134 | CID |
| 417 | 426.262 | 2 | 102.76 | 0.0438 | 19.69 | 53.68 | 3 | 0.0 | 0 | K.DTLMISR.T | Oxidation: 4 | 128-134 | CID |
| 1010 | 898.095 | 2 | 664.00 | 0.5959 | 32.04 | 77.96 | 3 | 0.0 | 0 | R.VVSVLTVVHQDWLNGK.E | | 181-196 | CID |
| 942 | 738.785 | 3 | 67.10 | 0.0496 | 30.57 | 94.33 | 6 | 0.0 | 1 | R.VVSVLTVVHQDWLNGKEYK.C | | 181-199 | CID |
| 604 | 643.898 | 2 | 89.15 | 0.0574 | 23.55 | 40.06 | 9 | 0.0 | 0 | R.EPQVYTLPPSR.E | | 224-234 | CID |
| 631 | 953.01 | 2 | 36.98 | 0.0352 | 24.06 | 76.6 | 24 | 0.0 | 1 | R.EPQVYTLPPSREEMTK.N | | 224-239 | CID |
| 756 | 581.394 | 2 | 129.97 | 0.0756 | 26.72 | 69.41 | 15 | 0.0 | 0 | K.NQVSLTCLVK.G | Carbamidomethyl: 7 | 240-249 | CID |
| 1023 | 953.549 | 2 | 103.22 | 0.0984 | 32.33 | 78.77 | 25 | 0.0 | 0 | K.TTPPMLDSDGSFFLYSK.L | | 272-288 | CID |
| 950 | 961.502 | 2 | 56.12 | 0.0540 | 30.69 | 17.04 | 1 | 0.0 | 0 | K.TTPPMLDSDGSFFLYSK.L | Oxidation: 5 | 272-288 | CID |
| 166 | 409.798 | 2 | 141.16 | 0.0578 | 13.55 | 44.26 | 3 | 0.0 | 1 | K.LTVDKSR.W | | 289-295 | CID |
| 726 | 934.552 | 3 | 133.53 | 0.1248 | 26.09 | 58.09 | 12 | 0.0 | 0 | R.WQQGNVFSCSVMHEALHNHYTQK.S | Carbamidomethyl: 9 | 296-318 | CID |

#### Protein 3: Immunoglobulin heavy constant gamma 3 OS=Homo sapiens GN=IGHG3 PE=1 SV=2

**Accession:** IGHG3_HUMAN
**Database:** SwissProt
**MW [kDa] / pI:** 41.3 / 8.2
**Modification(s):** Carbamidomethyl, Oxidation

**Score:** 941.3
**Seq. Coverage [%]:** 38.2
**No. of Peptides:** 14

BRUKER

| 10 | 20 | 30 | 40 | 50 | 60 | 70 | 80 | 90 | 100 |
|---|---|---|---|---|---|---|---|---|---|
| ASTKGPSVFP | LAPCSRSTSG | GTAALGCLVK | DYFPEPVTVS | WNSGALTSGV | HTFPAVLQSS | GLYSLSSVVT | VPSSSLGTQT | YTCNVNHKPS | NTKVDKRVEL |
| **110** | **120** | **130** | **140** | **150** | **160** | **170** | **180** | **190** | **200** |
| KTPLGDTTHT | CPRCPEPKSC | DTPPPCPRCP | EPKSCDTPPP | CPRCPEPKSC | DTPPPCPRCP | APELLGGPSV | FLFPPKPKDT | LMISRTPEVT | CVVVDVSHED |
| **210** | **220** | **230** | **240** | **250** | **260** | **270** | **280** | **290** | **300** |
| PEVQFKWYVD | GVEVHNAKTK | PREEQYNSTF | RVVSVLTVLH | QDWLNGKEYK | CKVSNKALPA | PIEKTISKTK | GQPREPQVYT | LPPSREEMTK | NQVSLTCLVK |
| **310** | **320** | **330** | **340** | **350** | **360** | **370** | **380** | | |
| GFYPSDIAVE | WESSGQPENN | YNTTPPMLDS | DGSFFLYSKL | TVDKSRWQQG | NIFSCSVMHE | ALHNRFTQKS | LSLSPGK | | |

**MS/MS Peptide Matches**

| Cmpd. | m/z meas. | z | Δ m/z [ppm] | Δ m/z [Da] | Rt [min] | Score | No. of Cmpds. | Site [%] | P | Sequence | Modification | Range | Type |
|---|---|---|---|---|---|---|---|---|---|---|---|---|---|
| 818 | 644.407 | 2 | 120.52 | 0.0777 | 27.99 | 74.43 | 22 | 0.0 | 0 | K.GPSVFPLAPCSR.S | Carbamidomethyl: 10 | 5-16 | CID |
| 678 | 661.43 | 2 | 132.08 | 0.0873 | 24.95 | 132.32 | 17 | 0.0 | 0 | R.STSGGTAALGCLVK.D | Carbamidomethyl: 11 | 17-30 | CID |
| 300 | 678.332 | 2 | 14.37 | 0.0097 | 16.89 | 88.5 | 2 | 0.0 | 0 | K.TPLGDTTHTCPR.C | Carbamidomethyl: 10 | 102-113 | CID |
| 256 | 593.803 | 2 | 84.86 | 0.0504 | 15.77 | 67.38 | 12 | 0.0 | 0 | K.SCDTPPPCPR.C | Carbamidomethyl: 2, 8 | 119-128 | CID |
| 1104 | 717.754 | 3 | 37.32 | 0.0268 | 34.04 | 76.08 | 2 | 0.0 | 0 | R.CPAPELLGGPSVFLFPPKPK.D | Carbamidomethyl: 1 | 159-178 | CID |
| 524 | 418.263 | 2 | 101.03 | 0.0423 | 21.98 | 47.69 | 6 | 0.0 | 0 | K.DTLMISR.T | | 179-185 | CID |
| 417 | 426.262 | 2 | 102.76 | 0.0438 | 19.69 | 53.68 | 3 | 0.0 | 0 | K.DTLMISR.T | Oxidation: 4 | 179-185 | CID |
| 610 | 472.968 | 3 | 140.13 | 0.0663 | 23.68 | 49.98 | 3 | 0.0 | 0 | K.WYVDGVEVHNAK.T | | 207-218 | CID |
| 1114 | 904.608 | 2 | 111.79 | 0.1011 | 34.20 | 90.66 | 13 | 0.0 | 0 | R.VVSVLTVLHQDWLNGK.E | | 232-247 | CID |
| 492 | 419.83 | 2 | 178.01 | 0.0747 | 21.37 | 30.25 | 2 | 0.0 | 0 | K.ALPAPIEK.T | | 257-264 | CID |
| 604 | 643.898 | 2 | 89.15 | 0.0574 | 23.55 | 40.06 | 9 | 0.0 | 0 | R.EPQVYTLPPSR.E | | 275-285 | CID |
| 631 | 953.01 | 2 | 36.98 | 0.0352 | 24.06 | 76.6 | 24 | 0.0 | 1 | R.EPQVYTLPPSREEMTK.N | | 275-290 | CID |
| 756 | 581.394 | 2 | 129.97 | 0.0756 | 26.72 | 69.41 | 15 | 0.0 | 0 | K.NQVSLTCLVK.G | Carbamidomethyl: 7 | 291-300 | CID |
| 166 | 409.798 | 2 | 141.16 | 0.0578 | 13.55 | 44.26 | 3 | 0.0 | 1 | K.LTVDKSR.W | | 340-346 | CID |

#### Protein 4: Immunoglobulin kappa constant OS=Homo sapiens GN=IGKC PE=1 SV=2

| | | | |
|---|---|---|---|
| **Accession:** | IGKC_HUMAN | **Score:** | 624.3 |
| **Database:** | SwissProt | **Seq. Coverage [%]:** | 74.8 |
| **MW [kDa] / pI:** | 11.8 / 6.1 | **No. of Peptides:** | 8 |
| **Modification(s):** | Carbamidomethyl | | |

10 20 30 40 50 60 70 80 90 100
RTVAAPSVFI FPPSDEQLKS GTASVVCLLN NFYPREAKVQ WKVDNALQSG NSQESVTEQD SKDSTYSLSS TLTLSKADYE KHKVYACEVT HQGLSSPVTK
110
SFNRGEC

**MS/MS Peptide Matches**

| Cmpd. | m/z meas. | z | Δ m/z [ppm] | Δ m/z [Da] | Rt [min] | Score | No. of Cmpds. | Site [%] | P | Sequence | Modification | Range | Type |
|---|---|---|---|---|---|---|---|---|---|---|---|---|---|
| 959 | 1052.092 | 2 | 498.62 | 0.5243 | 30.85 | 45.56 | 6 | 0.0 | 1 | -.RTVAAPSVFIFPPSDEQLK.S | | 1-19 | CID |
| 1086 | 973.639 | 2 | 125.20 | 0.1219 | 33.62 | 98.44 | 46 | 0.0 | 0 | R.TVAAPSVFIFPPSDEQLK.S | | 2-19 | CID |
| 1161 | 899.557 | 2 | 117.57 | 0.1057 | 35.05 | 65.38 | 28 | 0.0 | 0 | K.SGTASVVCLLNNFYPR.E | Carbamidomethyl: 8 | 20-35 | CID |
| 797 | 751.954 | 2 | 94.62 | 0.0711 | 27.53 | 100.05 | 11 | 0.0 | 0 | K.DSTYSLSSTLTLSK.A | | 63-76 | CID |
| 94 | 445.819 | 2 | 217.70 | 0.0970 | 11.28 | 26.65 | 5 | 0.0 | 1 | K.ADYEKHK.V | | 77-83 | CID |
| 405 | 714.404 | 3 | 54.43 | 0.0389 | 19.29 | 127.78 | 3 | 0.0 | 1 | K.HKVYACEVTHQGLSSPVTK.S | Carbamidomethyl: 6 | 82-100 | CID |
| 506 | 626.059 | 3 | 125.42 | 0.0785 | 21.64 | 117.71 | 9 | 0.0 | 0 | K.VYACEVTHQGLSSPVTK.S | Carbamidomethyl: 4 | 84-100 | CID |
| 181 | 435.256 | 2 | 169.70 | 0.0738 | 13.92 | 42.72 | 3 | 0.0 | 1 | K.SFNRGEC.- | Carbamidomethyl: 7 | 101-107 | CID |

#### Protein 5: Immunoglobulin heavy constant gamma 4 OS=Homo sapiens GN=IGHG4 PE=1 SV=1

| | | | |
|---|---|---|---|
| **Accession:** | IGHG4_HUMAN | **Score:** | 619.4 |
| **Database:** | SwissProt | **Seq. Coverage [%]:** | 39.8 |
| **MW [kDa] / pI:** | 35.9 / 7.2 | **No. of Peptides:** | 10 |
| **Modification(s):** | Carbamidomethyl, Oxidation | | |

| 10 | 20 | 30 | 40 | 50 | 60 | 70 | 80 | 90 | 100 |
|---|---|---|---|---|---|---|---|---|---|
| ASTKGPSVFP | LAPCSRSTSE | STAALGCLVK | DYFPEPVTVS | WNSGALTSGV | HTFPAVLQSS | GLYSLSSVVT | VPSSSLGTKT | YTCNVDHKPS | NTKVDKRVES |
| **110** | **120** | **130** | **140** | **150** | **160** | **170** | **180** | **190** | **200** |
| KYGPPCPSCP | APEFLGGPSV | FLFPPKPKDT | LMISRTPEVT | CVVVDVSQED | PEVQFNWYVD | GVEVHNAKTK | PREEQFNSTY | RVVSVLTVLH | QDWLNGKEYK |
| **210** | **220** | **230** | **240** | **250** | **260** | **270** | **280** | **290** | **300** |
| CKVSNKGLPS | SIEKTISKAK | GQPREPQVYT | LPPSQEEMTK | NQVSLTCLVK | GFYPSDIAVE | WESNGQPENN | YKTTPPVLDS | DGSFFLYSRL | TVDKSRWQEG |
| **310** | **320** | **330** | | | | | | | |
| NVFSCSVMHE | ALHNHYTQKS | LSLSLGK | | | | | | | |

MS/MS Peptide Matches

| Cmpd. | m/z meas. | z | Δ m/z [ppm] | Δ m/z [Da] | Rt [min] | Score | No. of Cmpds. | Site [%] | P | Sequence | Modification | Range | Type |
|---|---|---|---|---|---|---|---|---|---|---|---|---|---|
| 818 | 644.407 | 2 | 120.52 | 0.0777 | 27.99 | 74.43 | 22 | 0.0 | 0 | K.GPSVFPLAPCSR.S | Carbamidomethyl: 10 | 5-16 | CID |
| 689 | 712.414 | 2 | 77.91 | 0.0555 | 25.26 | 85.41 | 16 | 0.0 | 0 | R.STSESTAALGCLVK.D | Carbamidomethyl: 11 | 17-30 | CID |
| 178 | 832.942 | 2 | 64.25 | 0.0535 | 13.86 | 46.31 | 8 | 0.0 | 0 | K.TYTCNVDHKPSNTK.V | Carbamidomethyl: 4 | 80-93 | CID |
| 1140 | 982.217 | 3 | 400.36 | 0.3931 | 34.67 | 54.97 | 6 | 0.0 | 0 | K.YGPPCPSCPAPEFLGGPSVFLFPPKPK.D | Carbamidomethyl: 5, 8 | 102-128 | CID |
| 524 | 418.263 | 2 | 101.03 | 0.0423 | 21.98 | 47.69 | 6 | 0.0 | 0 | K.DTLMISR.T | | 129-135 | CID |
| 417 | 426.262 | 2 | 102.76 | 0.0438 | 19.69 | 53.68 | 3 | 0.0 | 0 | K.DTLMISR.T | Oxidation: 4 | 129-135 | CID |
| 1114 | 904.608 | 2 | 111.79 | 0.1011 | 34.20 | 90.66 | 13 | 0.0 | 0 | R.VVSVLTVLHQDWLNGK.E | | 182-197 | CID |
| 756 | 581.394 | 2 | 129.97 | 0.0756 | 26.72 | 69.41 | 15 | 0.0 | 0 | K.NQVSLTCLVK.G | Carbamidomethyl: 7 | 241-250 | CID |
| 166 | 409.798 | 2 | 141.16 | 0.0578 | 13.55 | 44.26 | 3 | 0.0 | 1 | R.LTVDKSR.W | | 290-296 | CID |
| 732 | 934.817 | 3 | 66.08 | 0.0618 | 26.19 | 52.54 | 8 | 0.0 | 0 | R.WQEGNVFSCSVMHEALHNHYTQK.S | Carbamidomethyl: 9 | 297-319 | CID |

#### Protein 6: Immunoglobulin lambda constant 2 OS=Homo sapiens GN=IGLC2 PE=1 SV=1

**Accession:** IGLC2_HUMAN
**Database:** SwissProt
**MW [kDa] / pI:** 11.3 / 6.9

**Score:** 570.2
**Seq. Coverage [%]:** 85.8
**No. of Peptides:** 7

**Modification(s):** Carbamidomethyl

```
         10         20         30         40         50         60         70         80         90        100
GQPKAAPSVT LFPPSSEELQ ANKATLVCLI SDFYPGAVTV AWKADSSPVK AGVETTTPSK QSNNKYAASS YLSLTPEQWK SHRSYSCQVT HEGSTVEKTV
        110
APTECS
```

**MS/MS Peptide Matches**

| Cmpd. | m/z meas. | z | Δ m/z [ppm] | Δ m/z [Da] | Rt [min] | Score | No. of Cmpds. | Site [%] | P | Sequence | Modification | Range | Type |
|---|---|---|---|---|---|---|---|---|---|---|---|---|---|
| 902 | 993.578 | 2 | 65.87 | 0.0654 | 29.67 | 116.47 | 28 | 0.0 | 0 | K.AAPSVTLFPPSSEELQANK.A | | 5-23 | CID |
| 1396 | 1106.605 | 2 | 475.05 | 0.5254 | 39.87 | 85.8 | 4 | 0.0 | 0 | K.ATLVCLISDFYPGAVTVAWK.A | Carbamidomethyl: 5 | 24-43 | CID |
| 336 | 558.995 | 3 | 69.16 | 0.0387 | 17.69 | 29.5 | 1 | 0.0 | 1 | K.ADSSPVKAGVETTTPSK.Q | | 44-60 | CID |
| 198 | 495.827 | 2 | 137.68 | 0.0683 | 14.33 | 74.39 | 19 | 0.0 | 0 | K.AGVETTTPSK.Q | | 51-60 | CID |
| 164 | 781.434 | 2 | 50.60 | 0.0395 | 13.49 | 52.91 | 2 | 0.0 | 1 | K.AGVETTTPSKQSNNK.Y | | 51-65 | CID |
| 931 | 872.57 | 2 | 156.97 | 0.1369 | 30.31 | 92.88 | 17 | 0.0 | 0 | K.YAASSYLSLTPEQWK.S | | 66-80 | CID |
| 294 | 856.468 | 2 | 98.98 | 0.0848 | 16.72 | 118.24 | 10 | 0.0 | 0 | R.SYSCQVTHEGSTVEK.T | Carbamidomethyl: 4 | 84-98 | CID |

#### Protein 7: Immunoglobulin kappa light chain OS=Homo sapiens PE=1 SV=1

**Accession:** IGK_HUMAN
**Database:** SwissProt
**MW [kDa] / pI:** 23.4 / 6.9
**Modification(s):** Carbamidomethyl

**Score:** 527.4
**Seq. Coverage [%]:** 33.6
**No. of Peptides:** 7

| 10 | 20 | 30 | 40 | 50 | 60 | 70 | 80 | 90 | 100 |
|---|---|---|---|---|---|---|---|---|---|
| DIQMTQSPST | LSASVGDRVT | ITCRASQSIN | TWLAWYQQKP | GKAPKLLMYK | ASSLESGVPS | RFIGSGSGTE | FTLTISSLQP | DDFATYYCQQ | YNSDSKMFGQ |
| **110** | **120** | **130** | **140** | **150** | **160** | **170** | **180** | **190** | **200** |
| GTKVEVKGTV | AAPSVFIFPP | SDEQLKSGTA | SVVCLLNNFY | PREAKVQWKV | DNALQSGNSQ | ESVTEQDSKD | STYSLSSTLT | LSKADYEKHK | VYACEVTHQG |
| **210** | **220** | | | | | | | | |
| LSSPVTKSFN | RGEC | | | | | | | | |

**MS/MS Peptide Matches**

| Cmpd. | m/z meas. | z | Δ m/z [ppm] | Δ m/z [Da] | Rt [min] | Score | No. of Cmpds. | Site [%] | P | Sequence | Modification | Range | Type |
|---|---|---|---|---|---|---|---|---|---|---|---|---|---|
| 369 | 545.291 | 2 | 19.48 | 0.0106 | 18.35 | 47.14 | 3 | 0.0 | 0 | K.ASSLESGVPSR.F | | 51-61 | CID |
| 1161 | 899.557 | 2 | 117.57 | 0.1057 | 35.05 | 65.38 | 28 | 0.0 | 0 | K.SGTASVVCLLNNFYPR.E | Carbamidomethyl: 8 | 127-142 | CID |
| 797 | 751.954 | 2 | 94.62 | 0.0711 | 27.53 | 100.05 | 11 | 0.0 | 0 | K.DSTYSLSSTLTLSK.A | | 170-183 | CID |
| 94 | 445.819 | 2 | 217.70 | 0.0970 | 11.28 | 26.65 | 5 | 0.0 | 1 | K.ADYEKHK.V | | 184-190 | CID |
| 405 | 714.404 | 3 | 54.43 | 0.0389 | 19.29 | 127.78 | 3 | 0.0 | 1 | K.HKVYACEVTHQGLSSPVTK.S | Carbamidomethyl: 6 | 189-207 | CID |
| 506 | 626.059 | 3 | 125.42 | 0.0785 | 21.64 | 117.71 | 9 | 0.0 | 0 | K.VYACEVTHQGLSSPVTK.S | Carbamidomethyl: 4 | 191-207 | CID |
| 181 | 435.256 | 2 | 169.70 | 0.0738 | 13.92 | 42.72 | 3 | 0.0 | 1 | K.SFNRGEC.- | Carbamidomethyl: 7 | 208-214 | CID |

#### Protein 8: Immunoglobulin lambda-1 light chain OS=Homo sapiens PE=1 SV=1

| | | | |
|---|---|---|---|
| **Accession:** | IGL1_HUMAN | **Score:** | 515.6 |
| **Database:** | SwissProt | **Seq. Coverage [%]:** | 44.0 |
| **MW [kDa] / pI:** | 22.8 / 6.3 | **No. of Peptides:** | 7 |
| **Modification(s):** | Carbamidomethyl | | |

| 10 | 20 | 30 | 40 | 50 | 60 | 70 | 80 | 90 | 100 |
|---|---|---|---|---|---|---|---|---|---|
| QSALTQPPSA | SGSLGQSVTI | SCTGTSSDVG | GYNYVSWYQQ | HAGKAPKVII | YEVNKRPSGV | PDRFSGSKSG | NTASLTVSGL | QAEDEADYYC | SSYEGSDNFV |
| 110 | 120 | 130 | 140 | 150 | 160 | 170 | 180 | 190 | 200 |
| FGTGTKVTVL | GQPKANPTVT | LFPPSSEELQ | ANKATLVCLI | SDFYPGAVTV | AWKADGSPVK | AGVETTKPSK | QSNNKYAASS | YLSLTPEQWK | SHRSYSCQVT |
| 210 | 220 | | | | | | | | |
| HEGSTVEKTV | APTECS | | | | | | | | |

MS/MS Peptide Matches

| Cmpd. | m/z meas. | z | Δ m/z [ppm] | Δ m/z [Da] | Rt [min] | Score | No. of Cmpds. | Site [%] | P | Sequence | Modification | Range | Type |
|---|---|---|---|---|---|---|---|---|---|---|---|---|---|
| 165 | 442.304 | 2 | 142.78 | 0.0631 | 13.50 | 36.48 | 3 | 0.0 | 0 | K.RPSGVPDR.F | | 56-63 | CID |
| 414 | 421.321 | 2 | 143.08 | 0.0603 | 19.60 | 41.41 | 3 | 0.0 | 0 | K.VTVLGQPK.A | | 107-114 | CID |
| 897 | 1022.08 | 2 | 55.48 | 0.0567 | 29.56 | 79.87 | 22 | 0.0 | 0 | K.ANPTVTLFPPSSEELQANK.A | | 115-133 | CID |
| 1396 | 1106.605 | 2 | 475.05 | 0.5254 | 39.87 | 85.8 | 4 | 0.0 | 0 | K.ATLVCLISDFYPGAVTVAWK.A | Carbamidomethyl: 5 | 134-153 | CID |
| 135 | 509.364 | 2 | 160.25 | 0.0816 | 12.73 | 60.87 | 8 | 0.0 | 0 | K.AGVETTKPSK.Q | | 161-170 | CID |
| 931 | 872.57 | 2 | 156.97 | 0.1369 | 30.31 | 92.88 | 17 | 0.0 | 0 | K.YAASSYLSLTPEQWK.S | | 176-190 | CID |
| 294 | 856.468 | 2 | 98.98 | 0.0848 | 16.72 | 118.24 | 10 | 0.0 | 0 | R.SYSCQVTHEGSTVEK.T | Carbamidomethyl: 4 | 194-208 | CID |

#### Protein 9: Immunoglobulin kappa variable 2D-28 OS=Homo sapiens GN=IGKV2D-28 PE=1 SV=2

**Accession:** KVD28_HUMAN
**Database:** SwissProt
**MW [kDa] / pI:** 12.9 / 5.6

**Score:** 124.1
**Seq. Coverage [%]:** 16.7
**No. of Peptides:** 2

| 10 | 20 | 30 | 40 | 50 | 60 | 70 | 80 | 90 | 100 |
|---|---|---|---|---|---|---|---|---|---|
| MRLPAQLLGL | LMLWVSGSSG | DIVMTQSPLS | LPVTPGEPAS | ISCRSSQSLL | HSNGYNYLDW | YLQKPGQSPQ | LLIYLGSNRA | SGVPDRFSGS | GSGTDFTLKI |
| 110 | 120 | 130 | | | | | | | |
| SRVEAEDVGV | YYCMQALQTP | | | | | | | | |

MS/MS Peptide Matches

| Cmpd. | m/z meas. | z | Δ m/z [ppm] | Δ m/z [Da] | Rt [min] | Score | No. of Cmpds. | Site [%] | P | Sequence | Modification | Range | Type |
|---|---|---|---|---|---|---|---|---|---|---|---|---|---|
| 671 | 662.715 | 3 | 87.60 | 0.0580 | 24.80 | 30.68 | 1 | 0.0 | 1 | R.ASGVPDRFSGSGSGTDFTLK.I | | 80-99 | CID |
| 623 | 652.352 | 2 | 61.51 | 0.0401 | 23.91 | 93.42 | 4 | 0.0 | 0 | R.FSGSGSGTDFTLK.I | | 87-99 | CID |

#### Protein 10: Immunoglobulin lambda variable 3-10 OS=Homo sapiens GN=IGLV3-10 PE=3 SV=2

**Accession:** LV310_HUMAN
**Database:** SwissProt
**MW [kDa] / pI:** 12.4 / 4.7
**Modification(s):** Carbamidomethyl

**Score:** 119.7
**Seq. Coverage [%]:** 15.7
**No. of Peptides:** 2

| 10 | 20 | 30 | 40 | 50 | 60 | 70 | 80 | 90 | 100 |
|---|---|---|---|---|---|---|---|---|---|
| MAWTPLLLPL | LTFCTVSEAS | YELTQPPSVS | VSPGQTARIT | CSGDALPKKY | AYWYQQKSGQ | APVLVIYEDS | KRPSGIPERF | SGSSSGTMAT | LTISGAQVED |
| **110** | **120** | | | | | | | | |
| EADYYCYSTD | SSGNH | | | | | | | | |

MS/MS Peptide Matches

| Cmpd. | m/z meas. | z | Δ m/z [ppm] | Δ m/z [Da] | Rt [min] | Score | No. of Cmpds. | Site [%] | P | Sequence | Modification | Range | Type |
|---|---|---|---|---|---|---|---|---|---|---|---|---|---|
| 401 | 531.312 | 2 | 82.02 | 0.0436 | 19.03 | 77.1 | 3 | 0.0 | 0 | R.ITCSGDALPK.K | Carbamidomethyl: 3 | 39-48 | CID |
| 227 | 456.324 | 2 | 147.93 | 0.0675 | 15.06 | 42.58 | 3 | 0.0 | 0 | K.RPSGIPER.F | | 72-79 | CID |

#### Protein 11: Immunoglobulin heavy variable 3-7 OS=Homo sapiens GN=IGHV3-7 PE=1 SV=2

**Accession:** HV307_HUMAN
**Database:** SwissProt
**MW [kDa] / pI:** 12.9 / 6.2
**Modification(s):** Carbamidomethyl

**Score:** 109.5
**Seq. Coverage [%]:** 15.4
**No. of Peptides:** 2

| 10 | 20 | 30 | 40 | 50 | 60 | 70 | 80 | 90 | 100 |
|---|---|---|---|---|---|---|---|---|---|
| MELGLSWVFL | VAILEGVQCE | VQLVESGGGL | VQPGGSLRLS | CAASGFTFSS | YWMSWVRQAP | GKGLEWVANI | KQDGSEKYYV | DSVKGRFTIS | RDNAKNSLYL |
| 110 | 120 | | | | | | | | |
| QMNSLRAEDT | AVYYCAR | | | | | | | | |

MS/MS Peptide Matches

| Cmpd. | m/z meas. | z | Δ m/z [ppm] | Δ m/z [Da] | Rt [min] | Score | No. of Cmpds. | Site [%] | P | Sequence | Modification | Range | Type |
|---|---|---|---|---|---|---|---|---|---|---|---|---|---|
| 400 | 437.288 | 2 | 152.63 | 0.0667 | 19.02 | 23.44 | 2 | 0.0 | 0 | K.YYVDSVK.G | | 78-84 | CID |
| 481 | 659.862 | 2 | 108.74 | 0.0717 | 21.11 | 86.09 | 6 | 0.0 | 0 | R.AEDTAVYYCAR.- | Carbamidomethyl: 9 | 107-117 | CID |

#### Protein 12: Immunoglobulin kappa variable 3-11 OS=Homo sapiens GN=IGKV3-11 PE=1 SV=1

**Accession:** KV311_HUMAN
**Database:** SwissProt
**MW [kDa] / pI:** 12.6 / 4.9

**Score:** 70.7
**Seq. Coverage [%]:** 7.8
**No. of Peptides:** 1

| 10 | 20 | 30 | 40 | 50 | 60 | 70 | 80 | 90 | 100 |
|---|---|---|---|---|---|---|---|---|---|
| MEAPAQLLFL | LLLWLPDTTG | EIVLTQSPAT | LSLSPGERAT | LSCRASQSVS | SYLAWYQQKP | GQAPRLLIYD | ASNRATGIPA | RFSGSGSGTD | FTLTISSLEP |
| 110 | 120 | | | | | | | | |
| EDFAVYYCQQ | RSNWP | | | | | | | | |

MS/MS Peptide Matches

| Cmpd. | m/z meas. | z | Δ m/z [ppm] | Δ m/z [Da] | Rt [min] | Score | No. of Cmpds. | Site [%] | P | Sequence | Modification | Range | Type |
|---|---|---|---|---|---|---|---|---|---|---|---|---|---|
| 643 | 532.829 | 2 | 72.49 | 0.0386 | 24.26 | 70.74 | 3 | 0.0 | 0 | R.LLIYDASNR.A | | 66-74 | CID |

#### Protein 13: Immunoglobulin kappa variable 3-20 OS=Homo sapiens GN=IGKV3-20 PE=1 SV=2

| | | | |
|---|---|---|---|
| **Accession:** | KV320_HUMAN | **Score:** | 69.4 |
| **Database:** | SwissProt | **Seq. Coverage [%]:** | 7.8 |
| **MW [kDa] / pI:** | 12.5 / 4.9 | **No. of Peptides:** | 1 |

```
        10         20         30         40         50         60         70         80         90        100
METPAQLLFL LLLWLPDTTG EIVLTQSPGT LSLSPGERAT LSCRASQSVS SSYLAWYQQK PGQAPRLLIY GASSRATGIP DRFSGSGSGT DFTLTISRLE
       110        120
PEDFAVYYCQ QYGSSP
```

MS/MS Peptide Matches

| Cmpd. | m/z meas. | z | Δ m/z [ppm] | Δ m/z [Da] | Rt [min] | Score | No. of Cmpds. | Site [%] | P | Sequence | Modification | Range | Type |
|---|---|---|---|---|---|---|---|---|---|---|---|---|---|
| 618 | 490.308 | 2 | 52.64 | 0.0258 | 23.81 | 69.39 | 3 | 0.0 | 0 | R.LLIYGASSR.A | | 67-75 | CID |

#### Protein 14: Immunoglobulin kappa variable 3-15 OS=Homo sapiens GN=IGKV3-15 PE=1 SV=2

| | | | |
|---|---|---|---|
| **Accession:** | KV315_HUMAN | **Score:** | 67.4 |
| **Database:** | SwissProt | **Seq. Coverage [%]:** | 7.8 |
| **MW [kDa] / pI:** | 12.5 / 5.1 | **No. of Peptides:** | 1 |

```
        10         20         30         40         50         60         70         80         90        100
MEAPAQLLFL LLLWLPDTTG EIVMTQSPAT LSVSPGERAT LSCRASQSVS SNLAWYQQKP GQAPRLLIYG ASTRATGIPA RFSGSGSGTE FTLTISSLQS
       110        120
EDFAVYYCQQ YNNWP
```

MS/MS Peptide Matches

| Cmpd. | m/z meas. | z | Δ m/z [ppm] | Δ m/z [Da] | Rt [min] | Score | No. of Cmpds. | Site [%] | P | Sequence | Modification | Range | Type |
|---|---|---|---|---|---|---|---|---|---|---|---|---|---|
| 641 | 497.354 | 2 | 128.66 | 0.0640 | 24.22 | 67.4 | 3 | 0.0 | 0 | R.LLIYGASTR.A | | 66-74 | CID |

#### Protein 15: Immunoglobulin kappa variable 4-1 OS=Homo sapiens GN=IGKV4-1 PE=1 SV=1

**Accession:** KV401_HUMAN
**Database:** SwissProt
**MW [kDa] / pI:** 13.4 / 5.1
**Score:** 58.1
**Seq. Coverage [%]:** 7.4
**No. of Peptides:** 1

| 10 | 20 | 30 | 40 | 50 | 60 | 70 | 80 | 90 | 100 |
|---|---|---|---|---|---|---|---|---|---|
| MVLQTQVFIS | LLLWISGAYG | DIVMTQSPDS | LAVSLGERAT | INCKSSQSVL | YSSNNKNYLA | WYQQKPGQPP | KLLIYWASTR | ESGVPDRFSG | SGSGTDFTLT |
| **110** | **120** | **130** | | | | | | | |
| ISSLQAEDVA | VYYCQQYYST | P | | | | | | | |

MS/MS Peptide Matches

| Cmpd. | m/z meas. | z | Δ m/z [ppm] | Δ m/z [Da] | Rt [min] | Score | No. of Cmpds. | Site [%] | P | Sequence | Modification | Range | Type |
|---|---|---|---|---|---|---|---|---|---|---|---|---|---|
| 938 | 561.89 | 2 | 126.48 | 0.0711 | 30.49 | 58.12 | 1 | 0.0 | 0 | K.LLIYWASTR.E | | 72-80 | CID |

#### Project Info

| | | | |
|---|---|---|---|
| **Name:** | Alagesan | **Date:** | Mar 5, 2019 |
| **Note:** | Nanobooster | | |

#### Sample Info & Protocols

| | |
|---|---|
| **Name:** | Nanobooster - IgG |
| **Date:** | Mar 5, 2019 |

#### Search Result Info

| | | | |
|---|---|---|---|
| **Search Result:** | IonTrap_Human_Swiss_CID_2019-03-05 17:55:08 | | |
| **Location:** | /Alagesan/Nanobooster - IgG/Ka_IgG Trypsin No Enrich No NB_BA5_01_8729.mgf | | |
| **Search Method:** | IonTrap_Human_Swiss_CID | **Version:** | 1.2 |
| **Search Engine(s):** | Mascot, 2.6.0 | **Ident. Compound(s):** | 0/2351 |
| **Database(s):** | SwissProt, SwissProt_2017_04.fasta | | |
| **Note:** | | | |

#### Results

##### Protein 1: Immunoglobulin heavy constant gamma 1 OS=Homo sapiens GN=IGHG1 PE=1 SV=1

**Accession:** IGHG1_HUMAN
**Database:** SwissProt
**MW [kDa] / pI:** 36.1 / 8.5
**Modification(s):** Carbamidomethyl, Oxidation
**Score:** 1270.0
**Seq. Coverage [%]:** 66.1
**No. of Peptides:** 18

```
        10         20         30         40         50         60         70         80         90        100
ASTKGPSVFP LAPSSKSTSG GTAALGCLVK DYFPEPVTVS WNSGALTSGV HTFPAVLQSS GLYSLSSVVT VPSSSLGTQT YICNVNHKPS NTKVDKKVEP
       110        120        130        140        150        160        170        180        190        200
KSCDKTHTCP PCPAPELLGG PSVFLFPPKP KDTLMISRTP EVTCVVVDVS HEDPEVKFNW YVDGVEVHNA KTKPREEQYN STYRVVSVLT VLHQDWLNGK
       210        220        230        240        250        260        270        280        290        300
EYKCKVSNKA LPAPIEKTIS KAKGQPREPQ VYTLPPSRDE LTKNQVSLTC LVKGFYPSDI AVEWESNGQP ENNYKTTPPV LDSDGSFFLY SKLTVDKSRW
       310        320        330        340
QQGNVFSCSV MHEALHNHYT QKSLSLSPGK
```

MS/MS Peptide Matches

| Cmpd. | m/z meas. | z | Δ m/z [ppm] | Δ m/z [Da] | Rt [min] | Score | No. of Cmpds. | Site [%] | P | Sequence | Modification | Range | Type |
|---|---|---|---|---|---|---|---|---|---|---|---|---|---|
| 805 | 593.884 | 2 | 96.05 | 0.0570 | 27.43 | 57.75 | 11 | 0.0 | 0 | K.GPSVFPLAPSSK.S | | 5-16 | CID |
| 697 | 661.417 | 2 | 112.42 | 0.0743 | 25.12 | 110.24 | 12 | 0.0 | 0 | K.STSGGTAALGCLVK.D | Carbamidomethyl: 11 | 17-30 | CID |
| 1031 | 834.746 | 4 | 395.50 | 0.3300 | 31.94 | 57.73 | 3 | 0.0 | 1 | K.SCDKTHTCPPCPAPELLGGPSVFLFPPKPK.D | Carbamidomethyl: 2, 8, 11 | 102-131 | CID |
| 1082 | 949.263 | 3 | 462.65 | 0.4390 | 32.94 | 86.49 | 12 | 0.0 | 0 | K.THTCPPCPAPELLGGPSVFLFPPKPK.D | Carbamidomethyl: 4, 7 | 106-131 | CID |
| 555 | 418.247 | 2 | 62.78 | 0.0263 | 22.46 | 47.79 | 7 | 0.0 | 0 | K.DTLMISR.T | | 132-138 | CID |
| 428 | 426.285 | 2 | 156.72 | 0.0668 | 19.86 | 55.77 | 6 | 0.0 | 0 | K.DTLMISR.T | Oxidation: 4 | 132-138 | CID |
| 792 | 713.756 | 3 | 105.57 | 0.0753 | 27.17 | 90.89 | 1 | 0.0 | 0 | R.TPEVTCVVVDVSHEDPEVK.F | Carbamidomethyl: 6 | 139-157 | CID |
| 790 | 839.495 | 2 | 107.66 | 0.0904 | 27.15 | 98.08 | 9 | 0.0 | 0 | K.FNWYVDGVEVHNAK.T | | 158-171 | CID |

| Cmpd. | m/z meas. | z | Δ m/z [ppm] | Δ m/z [Da] | Rt [min] | Score | No. of Cmpds. | Site [%] | P | Sequence | Modification | Range | Type |
|---|---|---|---|---|---|---|---|---|---|---|---|---|---|
| 1148 | 905.098 | 2 | 653.52 | 0.5911 | 34.32 | 107.99 | 7 | 0.0 | 0 | R.VVSVLTVLHQDWLNGK.E | | 185-200 | CID |
| 1091 | 743.807 | 3 | 537.65 | 0.3997 | 33.09 | 15.14 | 1 | 0.0 | 1 | R.VVSVLTVLHQDWLNGKEYK.C | | 185-203 | CID |
| 506 | 419.828 | 2 | 173.25 | 0.0727 | 21.48 | 20.1 | 2 | 0.0 | 0 | K.ALPAPIEK.T | | 210-217 | CID |
| 609 | 643.883 | 2 | 65.85 | 0.0424 | 23.44 | 55.2 | 8 | 0.0 | 0 | R.EPQVYTLPPSR.D | | 228-238 | CID |
| 644 | 937.014 | 2 | 26.98 | 0.0253 | 24.11 | 61.99 | 11 | 0.0 | 1 | R.EPQVYTLPPSRDELTK.N | | 228-243 | CID |
| 766 | 581.387 | 2 | 117.92 | 0.0686 | 26.61 | 82.75 | 13 | 0.0 | 0 | K.NQVSLTCLVK.G | Carbamidomethyl: 7 | 244-253 | CID |
| 1445 | 1272.578 | 2 | 6.82 | 0.0087 | 40.31 | 118.89 | 40 | 0.0 | 0 | K.GFYPSDIAVEWESNGQPENNYK.T | | 254-275 | CID |
| 1059 | 937.551 | 2 | 92.22 | 0.0865 | 32.50 | 105.96 | 15 | 0.0 | 0 | K.TTPPVLDSDGSFFLYSK.L | | 276-292 | CID |
| 158 | 409.811 | 2 | 172.89 | 0.0708 | 13.68 | 48.73 | 4 | 0.0 | 1 | K.LTVDKSR.W | | 293-299 | CID |
| 735 | 934.806 | 3 | 405.36 | 0.3788 | 25.99 | 48.47 | 9 | 0.0 | 0 | R.WQQGNVFSCSVMHEALHNHYTQK.S | Carbamidomethyl: 9 | 300-322 | CID |

#### Protein 2: Immunoglobulin heavy constant gamma 2 OS=Homo sapiens GN=IGHG2 PE=1 SV=2

| | | | |
|---|---|---|---|
| **Accession:** | IGHG2_HUMAN | **Score:** | 1006.6 |
| **Database:** | SwissProt | **Seq. Coverage [%]:** | 52.5 |
| **MW [kDa] / pI:** | 35.9 / 7.7 | **No. of Peptides:** | 17 |
| **Modification(s):** | Carbamidomethyl, Oxidation | | |

```
        10         20         30         40         50         60         70         80         90        100
ASTKGPSVFP LAPCSRSTSE STAALGCLVK DYFPEPVTVS WNSGALTSGV HTFPAVLQSS GLYSLSSVVT VPSSNFGTQT YTCNVDHKPS NTKVDKTVER
       110        120        130        140        150        160        170        180        190        200
KCCVECPPCP APPVAGPSVF LFPPKPKDTL MISRTPEVTC VVVDVSHEDP EVQFNWYVDG VEVHNAKTKP REEQFNSTFR VVSVLTVVHQ DWLNGKEYKC
       210        220        230        240        250        260        270        280        290        300
KVSNKGLPAP IEKTISKTKG QPREPQVYTL PPSREEMTKN QVSLTCLVKG FYPSDISVEW ESNGQPENNY KTTPPMLDSD GSFFLYSKLT VDKSRWQQGN
       310        320        330
VFSCSVMHEA LHNHYTQKSL SLSPGK
```

MS/MS Peptide Matches

| Cmpd. | m/z meas. | z | Δ m/z [ppm] | Δ m/z [Da] | Rt [min] | Score | No. of Cmpds. | Site [%] | P | Sequence | Modification | Range | Type |
|---|---|---|---|---|---|---|---|---|---|---|---|---|---|
| 846 | 644.394 | 2 | 100.34 | 0.0647 | 28.27 | 65.13 | 15 | 0.0 | 0 | K.GPSVFPLAPCSR.S | Carbamidomethyl: 10 | 5-16 | CID |
| 716 | 712.439 | 2 | 113.01 | 0.0805 | 25.56 | 82.04 | 11 | 0.0 | 0 | R.STSESTAALGCLVK.D | Carbamidomethyl: 11 | 17-30 | CID |
| 113 | 423.817 | 2 | 187.34 | 0.0794 | 12.30 | 37.57 | 4 | 0.0 | 1 | K.VDKTVER.K | | 94-100 | CID |
| 966 | 1013.195 | 3 | 353.40 | 0.3579 | 30.71 | 69.6 | 2 | 0.0 | 1 | R.KCCVECPPCPAPPVAGPSVFLFPPKPK.D | Carbamidomethyl: 2, 3, 6, 9 | 101-127 | CID |
| 1041 | 970.541 | 3 | 414.64 | 0.4023 | 32.16 | 79.5 | 9 | 0.0 | 0 | K.CCVECPPCPAPPVAGPSVFLFPPKPK.D | Carbamidomethyl: 1, 2, 5, 8 | 102-127 | CID |
| 555 | 418.247 | 2 | 62.78 | 0.0263 | 22.46 | 47.79 | 7 | 0.0 | 0 | K.DTLMISR.T | | 128-134 | CID |
| 428 | 426.285 | 2 | 156.72 | 0.0668 | 19.86 | 55.77 | 6 | 0.0 | 0 | K.DTLMISR.T | Oxidation: 4 | 128-134 | CID |
| 1044 | 599.077 | 3 | 682.40 | 0.4085 | 32.20 | 35.92 | 6 | 0.0 | 0 | R.VVSVLTVVHQDWLNGK.E | | 181-196 | CID |
| 965 | 738.78 | 3 | 60.34 | 0.0446 | 30.67 | 85.43 | 2 | 0.0 | 1 | R.VVSVLTVVHQDWLNGKEYK.C | | 181-199 | CID |
| 479 | 626.762 | 2 | -166.89 | -0.1046 | 20.88 | 16.53 | 1 | 0.0 | 1 | K.VSNKGLPAPIEK.T | | 202-213 | CID |
| 609 | 643.883 | 2 | 65.85 | 0.0424 | 23.44 | 55.2 | 8 | 0.0 | 0 | R.EPQVYTLPPSR.E | | 224-234 | CID |
| 602 | 953.01 | 2 | 36.98 | 0.0352 | 23.33 | 82.3 | 19 | 0.0 | 1 | R.EPQVYTLPPSREEMTK.N | | 224-239 | CID |
| 766 | 581.387 | 2 | 117.92 | 0.0686 | 26.61 | 82.75 | 13 | 0.0 | 0 | K.NQVSLTCLVK.G | Carbamidomethyl: 7 | 240-249 | CID |
| 1054 | 953.537 | 2 | 90.63 | 0.0864 | 32.40 | 90.29 | 11 | 0.0 | 0 | K.TTPPMLDSDGSFFLYSK.L | | 272-288 | CID |
| 964 | 961.492 | 2 | 45.72 | 0.0440 | 30.67 | 23.53 | 1 | 0.0 | 0 | K.TTPPMLDSDGSFFLYSK.L | Oxidation: 5 | 272-288 | CID |
| 158 | 409.811 | 2 | 172.89 | 0.0708 | 13.68 | 48.73 | 4 | 0.0 | 1 | K.LTVDKSR.W | | 289-295 | CID |
| 735 | 934.806 | 3 | 405.36 | 0.3788 | 25.99 | 48.47 | 9 | 0.0 | 0 | R.WQQGNVFSCSVMHEALHNHYTQK.S | Carbamidomethyl: 9 | 296-318 | CID |

#### Protein 3: Immunoglobulin kappa constant OS=Homo sapiens GN=IGKC PE=1 SV=2

**Accession:** IGKC_HUMAN
**Database:** SwissProt
**MW [kDa] / pI:** 11.8 / 6.1
**Modification(s):** Carbamidomethyl

**Score:** 979.0
**Seq. Coverage [%]:** 93.5
**No. of Peptides:** 11

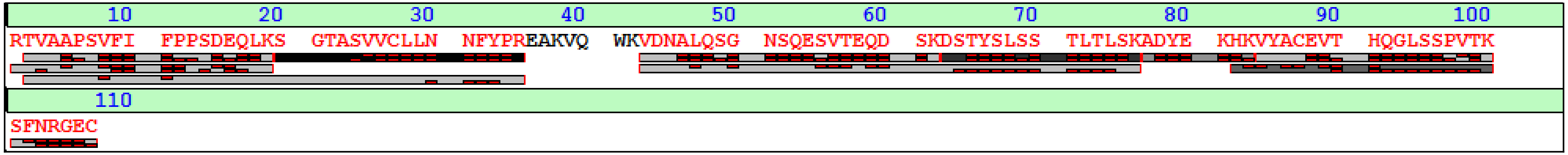


MS/MS Peptide Matches

| Cmpd. | m/z meas. | z | Δ m/z [ppm] | Δ m/z [Da] | Rt [min] | Score | No. of Cmpds. | Site [%] | P | Sequence | Modification | Range | Type |
|---|---|---|---|---|---|---|---|---|---|---|---|---|---|
| 998 | 1051.616 | 2 | 45.96 | 0.0483 | 31.27 | 42.13 | 6 | 0.0 | 1 | -.RTVAAPSVFIFPPSDEQLK.S | | 1-19 | CID |
| 1109 | 973.653 | 2 | 139.58 | 0.1359 | 33.47 | 107.3 | 30 | 0.0 | 0 | R.TVAAPSVFIFPPSDEQLK.S | | 2-19 | CID |
| 1422 | 1242.641 | 3 | 269.42 | 0.3347 | 39.83 | 21.39 | 1 | 0.0 | 1 | R.TVAAPSVFIFPPSDEQLKSGTASVVCLLNNFYP R.E | Carbamidomethyl: 26 | 2-35 | CID |
| 1190 | 899.571 | 2 | 133.13 | 0.1197 | 35.19 | 70.54 | 23 | 0.0 | 0 | K.SGTASVVCLLNNFYPR.E | Carbamidomethyl: 8 | 20-35 | CID |
| 2340 | 1068.502 | 2 | 13.10 | 0.0140 | 58.65 | 150.79 | 30 | 0.0 | 0 | K.VDNALQSGNSQESVTEQDSK.D | | 43-62 | CID |
| 2294 | 1207.195 | 3 | -38.35 | -0.0463 | 57.44 | 159.02 | 37 | 0.0 | 1 | K.VDNALQSGNSQESVTEQDSKDSTYSLSSTLTLS K.A | | 43-76 | CID |
| 836 | 751.948 | 2 | 86.64 | 0.0651 | 27.99 | 94.08 | 6 | 0.0 | 0 | K.DSTYSLSSTLTLSK.A | | 63-76 | CID |
| 89 | 445.785 | 2 | 141.42 | 0.0630 | 11.33 | 34.83 | 5 | 0.0 | 1 | K.ADYEKHK.V | | 77-83 | CID |
| 418 | 714.388 | 3 | 32.03 | 0.0229 | 19.62 | 137.08 | 8 | 0.0 | 1 | K.HKVYACEVTHQGLSSPVTK.S | Carbamidomethyl: 6 | 82-100 | CID |
| 510 | 938.499 | 2 | 33.99 | 0.0319 | 21.57 | 116.38 | 5 | 0.0 | 0 | K.VYACEVTHQGLSSPVTK.S | Carbamidomethyl: 4 | 84-100 | CID |
| 171 | 435.221 | 2 | 89.27 | 0.0388 | 13.99 | 45.47 | 7 | 0.0 | 1 | K.SFNRGEC.- | Carbamidomethyl: 7 | 101-107 | CID |

#### Protein 4: Immunoglobulin heavy constant gamma 3 OS=Homo sapiens GN=IGHG3 PE=1 SV=2

**Accession:** IGHG3_HUMAN
**Database:** SwissProt
**MW [kDa] / pI:** 41.3 / 8.2
**Modification(s):** Carbamidomethyl, Oxidation

**Score:** 954.3
**Seq. Coverage [%]:** 39.0
**No. of Peptides:** 15

| 10 | 20 | 30 | 40 | 50 | 60 | 70 | 80 | 90 | 100 |
|---|---|---|---|---|---|---|---|---|---|
| ASTKGPSVFP | LAPCSRSTSG | GTAALGCLVK | DYFPEPVTVS | WNSGALTSGV | HTFPAVLQSS | GLYSLSSVVT | VPSSSLGTQT | YTCNVNHKPS | NTKVDKRVEL |
| **110** | **120** | **130** | **140** | **150** | **160** | **170** | **180** | **190** | **200** |
| KTPLGDTTHT | CPRCPEPKSC | DTPPPCPRCP | EPKSCDTPPP | CPRCPEPKSC | DTPPPCPRCP | APELLGGPSV | FLFPPKPKDT | LMISRTPEVT | CVVVDVSHED |
| **210** | **220** | **230** | **240** | **250** | **260** | **270** | **280** | **290** | **300** |
| PEVQFKWYVD | GVEVHNAKTK | PREEQYNSTF | RVVSVLTVLH | QDWLNGKEYK | CKVSNKALPA | PIEKTISKTK | GQPREPQVYT | LPPSREEMTK | NQVSLTCLVK |
| **310** | **320** | **330** | **340** | **350** | **360** | **370** | **380** | | |
| GFYPSDIAVE | WESSGQPENN | YNTTPPMLDS | DGSFFLYSKL | TVDKSRWQQG | NIFSCSVMHE | ALHNRFTQKS | LSLSPGK | | |

MS/MS Peptide Matches

| Cmpd. | m/z meas. | z | Δ m/z [ppm] | Δ m/z [Da] | Rt [min] | Score | No. of Cmpds. | Site [%] | P | Sequence | Modification | Range | Type |
|---|---|---|---|---|---|---|---|---|---|---|---|---|---|
| 846 | 644.394 | 2 | 100.34 | 0.0647 | 28.27 | 65.13 | 15 | 0.0 | 0 | K.GPSVFPLAPCSR.S | Carbamidomethyl: 10 | 5-16 | CID |
| 697 | 661.417 | 2 | 112.42 | 0.0743 | 25.12 | 110.24 | 12 | 0.0 | 0 | R.STSGGTAALGCLVK.D | Carbamidomethyl: 11 | 17-30 | CID |
| 299 | 452.602 | 3 | 113.60 | 0.0514 | 16.95 | 60.4 | 3 | 0.0 | 0 | K.TPLGDTTHTCPR.C | Carbamidomethyl: 10 | 102-113 | CID |
| 253 | 593.792 | 2 | 66.33 | 0.0394 | 15.93 | 61.97 | 11 | 0.0 | 0 | K.SCDTPPPCPR.C | Carbamidomethyl: 2, 8 | 119-128 | CID |
| 1136 | 717.809 | 3 | 113.95 | 0.0818 | 34.10 | 90.41 | 2 | 0.0 | 0 | R.CPAPELLGGPSVFLFPPKPK.D | Carbamidomethyl: 1 | 159-178 | CID |
| 555 | 418.247 | 2 | 62.78 | 0.0263 | 22.46 | 47.79 | 7 | 0.0 | 0 | K.DTLMISR.T | | 179-185 | CID |
| 428 | 426.285 | 2 | 156.72 | 0.0668 | 19.86 | 55.77 | 6 | 0.0 | 0 | K.DTLMISR.T | Oxidation: 4 | 179-185 | CID |
| 627 | 472.932 | 3 | 64.01 | 0.0303 | 23.81 | 50.36 | 3 | 0.0 | 0 | K.WYVDGVEVHNAK.T | | 207-218 | CID |
| 1148 | 905.098 | 2 | 653.52 | 0.5911 | 34.32 | 107.99 | 7 | 0.0 | 0 | R.VVSVLTVLHQDWLNGK.E | | 232-247 | CID |
| 1091 | 743.807 | 3 | 537.65 | 0.3997 | 33.09 | 15.14 | 1 | 0.0 | 1 | R.VVSVLTVLHQDWLNGKEYK.C | | 232-250 | CID |
| 506 | 419.828 | 2 | 173.25 | 0.0727 | 21.48 | 20.1 | 2 | 0.0 | 0 | K.ALPAPIEK.T | | 257-264 | CID |
| 609 | 643.883 | 2 | 65.85 | 0.0424 | 23.44 | 55.2 | 8 | 0.0 | 0 | R.EPQVYTLPPSR.E | | 275-285 | CID |
| 602 | 953.01 | 2 | 36.98 | 0.0352 | 23.33 | 82.3 | 19 | 0.0 | 1 | R.EPQVYTLPPSREEMTK.N | | 275-290 | CID |
| 766 | 581.387 | 2 | 117.92 | 0.0686 | 26.61 | 82.75 | 13 | 0.0 | 0 | K.NQVSLTCLVK.G | Carbamidomethyl: 7 | 291-300 | CID |
| 158 | 409.811 | 2 | 172.89 | 0.0708 | 13.68 | 48.73 | 4 | 0.0 | 1 | K.LTVDKSR.W | | 340-346 | CID |

BRUKER

#### Protein 5: Immunoglobulin kappa light chain OS=Homo sapiens PE=1 SV=1

| | | | |
|---|---|---|---|
| **Accession:** | IGK_HUMAN | **Score:** | 895.0 |
| **Database:** | SwissProt | **Seq. Coverage [%]:** | 43.0 |
| **MW [kDa] / pI:** | 23.4 / 6.9 | **No. of Peptides:** | 9 |
| **Modification(s):** | Carbamidomethyl | | |

| 10 | 20 | 30 | 40 | 50 | 60 | 70 | 80 | 90 | 100 |
|---|---|---|---|---|---|---|---|---|---|
| DIQMTQSPST | LSASVGDRVT | ITCRASQSIN | TWLAWYQQKP | GKAPKLLMYK | ASSLESGVPS | RFIGSGSGTE | FTLTISSLQP | DDFATYYCQQ | YNSDSKMFGQ |
| **110** | **120** | **130** | **140** | **150** | **160** | **170** | **180** | **190** | **200** |
| GTKVEVKGTV | AAPSVFIFPP | SDEQLKSGTA | SVVCLLNNFY | PREAKVQWKV | DNALQSGNSQ | ESVTEQDSKD | STYSLSSTLT | LSKADYEKHK | VYACEVTHQG |
| **210** | **220** | | | | | | | | |
| LSSPVTKSFN | RGEC | | | | | | | | |

##### MS/MS Peptide Matches

| Cmpd. | m/z meas. | z | Δ m/z [ppm] | Δ m/z [Da] | Rt [min] | Score | No. of Cmpds. | Site [%] | P | Sequence | Modification | Range | Type |
|---|---|---|---|---|---|---|---|---|---|---|---|---|---|
| 377 | 545.324 | 2 | 80.00 | 0.0436 | 18.59 | 86.77 | 2 | 0.0 | 0 | K.ASSLESGVPSR.F | | 51-61 | CID |
| 1190 | 899.571 | 2 | 133.13 | 0.1197 | 35.19 | 70.54 | 23 | 0.0 | 0 | K.SGTASVVCLLNNFYPR.E | Carbamidomethyl: 8 | 127-142 | CID |
| 2340 | 1068.502 | 2 | 13.10 | 0.0140 | 58.65 | 150.79 | 30 | 0.0 | 0 | K.VDNALQSGNSQESVTEQDSK.D | | 150-169 | CID |
| 2294 | 1207.195 | 3 | -38.35 | -0.0463 | 57.44 | 159.02 | 37 | 0.0 | 1 | K.VDNALQSGNSQESVTEQDSKDSTYSLSSTLTLSK.A | | 150-183 | CID |
| 836 | 751.948 | 2 | 86.64 | 0.0651 | 27.99 | 94.08 | 6 | 0.0 | 0 | K.DSTYSLSSTLTLSK.A | | 170-183 | CID |
| 89 | 445.785 | 2 | 141.42 | 0.0630 | 11.33 | 34.83 | 5 | 0.0 | 1 | K.ADYEKHK.V | | 184-190 | CID |
| 418 | 714.388 | 3 | 32.03 | 0.0229 | 19.62 | 137.08 | 8 | 0.0 | 1 | K.HKVYACEVTHQGLSSPVTK.S | Carbamidomethyl: 6 | 189-207 | CID |
| 510 | 938.499 | 2 | 33.99 | 0.0319 | 21.57 | 116.38 | 5 | 0.0 | 0 | K.VYACEVTHQGLSSPVTK.S | Carbamidomethyl: 4 | 191-207 | CID |
| 171 | 435.221 | 2 | 89.27 | 0.0388 | 13.99 | 45.47 | 7 | 0.0 | 1 | K.SFNRGEC.- | Carbamidomethyl: 7 | 208-214 | CID |

#### Protein 6: Immunoglobulin heavy constant gamma 4 OS=Homo sapiens GN=IGHG4 PE=1 SV=1

**Accession:** IGHG4_HUMAN
**Database:** SwissProt
**MW [kDa] / pI:** 35.9 / 7.2
**Modification(s):** Carbamidomethyl, Oxidation

**Score:** 746.2
**Seq. Coverage [%]:** 47.4
**No. of Peptides:** 11

| 10 | 20 | 30 | 40 | 50 | 60 | 70 | 80 | 90 | 100 |
|---|---|---|---|---|---|---|---|---|---|
| ASTKGPSVFP | LAPCSRSTSE | STAALGCLVK | DYFPEPVTVS | WNSGALTSGV | HTFPAVLQSS | GLYSLSSVVT | VPSSSLGTKT | YTCNVDHKPS | NTKVDKRVES |
| **110** | **120** | **130** | **140** | **150** | **160** | **170** | **180** | **190** | **200** |
| KYGPPCPSCP | APEFLGGPSV | FLFPPKPKDT | LMISRTPEVT | CVVVDVSQED | PEVQFNWYVD | GVEVHNAKTK | PREEQFNSTY | RVVSVLTVLH | QDWLNGKEYK |
| **210** | **220** | **230** | **240** | **250** | **260** | **270** | **280** | **290** | **300** |
| CKVSNKGLPS | SIEKTISKAK | GQPREPQVYT | LPPSQEEMTK | NQVSLTCLVK | GFYPSDIAVE | WESNGQPENN | YKTTPPVLDS | DGSFFLYSRL | TVDKSRWQEG |
| **310** | **320** | **330** | | | | | | | |
| NVFSCSVMHE | ALHNHYTQKS | LSLSLGK | | | | | | | |

##### MS/MS Peptide Matches

| Cmpd. | m/z meas. | z | Δ m/z [ppm] | Δ m/z [Da] | Rt [min] | Score | No. of Cmpds. | Site [%] | P | Sequence | Modification | Range | Type |
|---|---|---|---|---|---|---|---|---|---|---|---|---|---|
| 846 | 644.394 | 2 | 100.34 | 0.0647 | 28.27 | 65.13 | 15 | 0.0 | 0 | K.GPSVFPLAPCSR.S | Carbamidomethyl: 10 | 5-16 | CID |
| 716 | 712.439 | 2 | 113.01 | 0.0805 | 25.56 | 82.04 | 11 | 0.0 | 0 | R.STSESTAALGCLVK.D | Carbamidomethyl: 11 | 17-30 | CID |
| 170 | 555.676 | 3 | 146.24 | 0.0813 | 13.98 | 43.99 | 7 | 0.0 | 0 | K.TYTCNVDHKPSNTK.V | Carbamidomethyl: 4 | 80-93 | CID |
| 1170 | 982.217 | 3 | 400.36 | 0.3931 | 34.76 | 46.21 | 6 | 0.0 | 0 | K.YGPPCPSCPAPEFLGGPSVFLFPPKPK.D | Carbamidomethyl: 5, 8 | 102-128 | CID |
| 555 | 418.247 | 2 | 62.78 | 0.0263 | 22.46 | 47.79 | 7 | 0.0 | 0 | K.DTLMISR.T | | 129-135 | CID |
| 428 | 426.285 | 2 | 156.72 | 0.0668 | 19.86 | 55.77 | 6 | 0.0 | 0 | K.DTLMISR.T | Oxidation: 4 | 129-135 | CID |
| 1148 | 905.098 | 2 | 653.52 | 0.5911 | 34.32 | 107.99 | 7 | 0.0 | 0 | R.VVSVLTVLHQDWLNGK.E | | 182-197 | CID |
| 1091 | 743.807 | 3 | 537.65 | 0.3997 | 33.09 | 15.14 | 1 | 0.0 | 1 | R.VVSVLTVLHQDWLNGKEYK.C | | 182-200 | CID |
| 766 | 581.387 | 2 | 117.92 | 0.0686 | 26.61 | 82.75 | 13 | 0.0 | 0 | K.NQVSLTCLVK.G | Carbamidomethyl: 7 | 241-250 | CID |

| Cmpd. | m/z meas. | z | Δ m/z [ppm] | Δ m/z [Da] | Rt [min] | Score | No. of Cmpds. | Site [%] | P | Sequence | Modification | Range | Type |
|---|---|---|---|---|---|---|---|---|---|---|---|---|---|
| 1445 | 1272.578 | 2 | 6.82 | 0.0087 | 40.31 | 118.89 | 40 | 0.0 | 0 | K.GFYPSDIAVEWESNGQPENNYK.T | | 251-272 | CID |
| 158 | 409.811 | 2 | 172.89 | 0.0708 | 13.68 | 48.73 | 4 | 0.0 | 1 | R.LTVDKSR.W | | 290-296 | CID |

#### Protein 7: Immunoglobulin lambda-1 light chain OS=Homo sapiens PE=1 SV=1

**Accession:** IGL1_HUMAN
**Database:** SwissProt
**MW [kDa] / pI:** 22.8 / 6.3
**Modification(s):** Carbamidomethyl

**Score:** 588.9
**Seq. Coverage [%]:** 50.0
**No. of Peptides:** 9

```
        10         20         30         40         50         60         70         80         90        100
QSALTQPPSA SGSLGQSVTI SCTGTSSDVG GYNYVSWYQQ HAGKAPKVII YEVNKRPSGV PDRFSGSKSG NTASLTVSGL QAEDEADYYC SSYEGSDNFV
       110        120        130        140        150        160        170        180        190        200
FGTGTKVTVL GQPKANPTVT LFPPSSEELQ ANKATLVCLI SDFYPGAVTV AWKADGSPVK AGVETTKPSK QSNNKYAASS YLSLTPEQWK SHRSYSCQVT
       210        220
HEGSTVEKTV APTECS
```

MS/MS Peptide Matches

| Cmpd. | m/z meas. | z | Δ m/z [ppm] | Δ m/z [Da] | Rt [min] | Score | No. of Cmpds. | Site [%] | P | Sequence | Modification | Range | Type |
|---|---|---|---|---|---|---|---|---|---|---|---|---|---|
| 159 | 442.296 | 2 | 124.69 | 0.0551 | 13.69 | 24.91 | 3 | 0.0 | 0 | K.RPSGVPDR.F | | 56-63 | CID |
| 312 | 463.977 | 3 | 138.75 | 0.0644 | 17.22 | 34.33 | 2 | 0.0 | 1 | K.RPSGVPDRFSGSK.S | | 56-68 | CID |
| 426 | 421.318 | 2 | 135.95 | 0.0573 | 19.82 | 43.36 | 3 | 0.0 | 0 | K.VTVLGQPK.A | | 107-114 | CID |
| 889 | 1022.064 | 2 | 39.83 | 0.0407 | 29.07 | 77.38 | 12 | 0.0 | 0 | K.ANPTVTLFPPSSEELQANK.A | | 115-133 | CID |
| 1430 | 1106.134 | 2 | 49.22 | 0.0544 | 40.01 | 98.45 | 4 | 0.0 | 0 | K.ATLVCLISDFYPGAVTVAWK.A | Carbamidomethyl: 5 | 134-153 | CID |
| 136 | 509.336 | 2 | 105.27 | 0.0536 | 13.02 | 74.9 | 7 | 0.0 | 0 | K.AGVETTKPSK.Q | | 161-170 | CID |
| 960 | 872.522 | 2 | 101.95 | 0.0889 | 30.58 | 94.61 | 16 | 0.0 | 0 | K.YAASSYLSLTPEQWK.S | | 176-190 | CID |

| Cmpd. | m/z meas. | z | Δ m/z [ppm] | Δ m/z [Da] | Rt [min] | Score | No. of Cmpds. | Site [%] | P | Sequence | Modification | Range | Type |
|---|---|---|---|---|---|---|---|---|---|---|---|---|---|
| 282 | 856.511 | 2 | 149.19 | 0.1278 | 16.64 | 105.15 | 10 | 0.0 | 0 | R.SYSCQVTHEGSTVEK.T | Carbamidomethyl: 4 | 194-208 | CID |
| 489 | 853.065 | 3 | 24.36 | 0.0208 | 21.15 | 35.84 | 2 | 0.0 | 1 | R.SYSCQVTHEGSTVEKTVAPTECS.- | Carbamidomethyl: 4, 22 | 194-216 | CID |

#### Protein 8: Immunoglobulin lambda constant 3 OS=Homo sapiens GN=IGLC3 PE=1 SV=1

**Accession:** IGLC3_HUMAN
**Database:** SwissProt
**MW [kDa] / pI:** 11.3 / 6.9
**Modification(s):** Carbamidomethyl

**Score:** 578.7
**Seq. Coverage [%]:** 93.4
**No. of Peptides:** 8

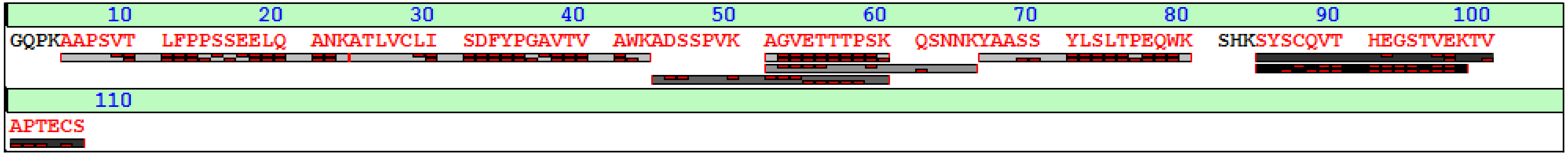


MS/MS Peptide Matches

| Cmpd. | m/z meas. | z | Δ m/z [ppm] | Δ m/z [Da] | Rt [min] | Score | No. of Cmpds. | Site [%] | P | Sequence | Modification | Range | Type |
|---|---|---|---|---|---|---|---|---|---|---|---|---|---|
| 880 | 993.557 | 2 | 44.73 | 0.0444 | 28.92 | 91.55 | 19 | 0.0 | 0 | K.AAPSVTLFPPSSEELQANK.A | | 5-23 | CID |
| 1430 | 1106.134 | 2 | 49.22 | 0.0544 | 40.01 | 98.45 | 4 | 0.0 | 0 | K.ATLVCLISDFYPGAVTVAWK.A | Carbamidomethyl: 5 | 24-43 | CID |
| 339 | 559.025 | 3 | 122.83 | 0.0687 | 17.82 | 29.9 | 2 | 0.0 | 1 | K.ADSSPVKAGVETTTPSK.Q | | 44-60 | CID |
| 207 | 495.794 | 2 | 71.11 | 0.0353 | 14.85 | 79.07 | 15 | 0.0 | 0 | K.AGVETTTPSK.Q | | 51-60 | CID |
| 153 | 521.332 | 3 | 127.77 | 0.0666 | 13.51 | 44.12 | 3 | 0.0 | 1 | K.AGVETTTPSKQSNNK.Y | | 51-65 | CID |
| 960 | 872.522 | 2 | 101.95 | 0.0889 | 30.58 | 94.61 | 16 | 0.0 | 0 | K.YAASSYLSLTPEQWK.S | | 66-80 | CID |
| 282 | 856.511 | 2 | 149.19 | 0.1278 | 16.64 | 105.15 | 10 | 0.0 | 0 | K.SYSCQVTHEGSTVEK.T | Carbamidomethyl: 4 | 84-98 | CID |
| 489 | 853.065 | 3 | 24.36 | 0.0208 | 21.15 | 35.84 | 2 | 0.0 | 1 | K.SYSCQVTHEGSTVEKTVAPTECS.- | Carbamidomethyl: 4, 22 | 84-106 | CID |

#### Protein 9: Alpha-1-antitrypsin OS=Homo sapiens GN=SERPINA1 PE=1 SV=3

| | | | |
|---|---|---|---|
| **Accession:** | A1AT_HUMAN | **Score:** | 233.8 |
| **Database:** | SwissProt | **Seq. Coverage [%]:** | 19.9 |
| **MW [kDa] / pI:** | 46.7 / 5.4 | **No. of Peptides:** | 4 |

| | | | | | | | | | |
|---|---|---|---|---|---|---|---|---|---|
| 10 | 20 | 30 | 40 | 50 | 60 | 70 | 80 | 90 | 100 |
| MPSSVSWGIL | LLAGLCCLVP | VSLAEDPQGD | AAQKTDTSHH | DQDHPTFNKI | TPNLAEFAFS | LYRQLAHQSN | STNIFFSPVS | IATAFAMLSL | GTKADTHDEI |
| 110 | 120 | 130 | 140 | 150 | 160 | 170 | 180 | 190 | 200 |
| LEGLNFNLTE | IPEAQIHEGF | QELLRTLNQP | DSQLQLTTGN | GLFLSEGLKL | VDKFLEDVKK | LYHSEAFTVN | FGDTEEAKKQ | INDYVEKGTQ | GKIVDLVKEL |
| 210 | 220 | 230 | 240 | 250 | 260 | 270 | 280 | 290 | 300 |
| DRDTVFALVN | YIFFKGKWER | PFEVKDTEEE | DFHVDQVTTV | KVPMMKRLGM | FNIQHCKKLS | SWVLLMKYLG | NATAIFFLPD | EGKLQHLENE | LTHDIITKFL |
| 310 | 320 | 330 | 340 | 350 | 360 | 370 | 380 | 390 | 400 |
| ENEDRRSASL | HLPKLSITGT | YDLKSVLGQL | GITKVFSNGA | DLSGVTEEAP | LKLSKAVHKA | VLTIDEKGTE | AAGAMFLEAI | PMSIPPEVKF | NKPFVFLMIE |
| 410 | 420 | | | | | | | | |
| QNTKSPLFMG | KVVNPTQK | | | | | | | | |

##### MS/MS Peptide Matches

| Cmpd. | m/z meas. | z | Δ m/z [ppm] | Δ m/z [Da] | Rt [min] | Score | No. of Cmpds. | Site [%] | P | Sequence | Modification | Range | Type |
|---|---|---|---|---|---|---|---|---|---|---|---|---|---|
| 1515 | 1045.632 | 2 | 77.00 | 0.0805 | 41.87 | 74.58 | 3 | 0.0 | 1 | K.ELDRDTVFALVNYIFFK.G | | 199-215 | CID |
| 502 | 425.96 | 3 | 141.67 | 0.0603 | 21.40 | 25.19 | 4 | 0.0 | 1 | K.GKWERPFEVK.D | | 216-225 | CID |
| 2285 | 917.959 | 2 | 538.35 | 0.4939 | 57.22 | 63.9 | 4 | 0.0 | 0 | K.VFSNGADLSGVTEEAPLK.L | | 335-352 | CID |
| 1312 | 1043.932 | 3 | 369.15 | 0.3852 | 37.64 | 54.66 | 1 | 0.0 | 1 | K.AVLTIDEKGTEAAGAMFLEAIPMSIPPEVK.F | | 360-389 | CID |

#### Protein 10: Immunoglobulin heavy variable 3-33 OS=Homo sapiens GN=IGHV3-33 PE=1 SV=2

| | | | |
|---|---|---|---|
| **Accession:** | HV333_HUMAN | **Score:** | 101.3 |
| **Database:** | SwissProt | **Seq. Coverage [%]:** | 21.4 |
| **MW [kDa] / pI:** | 13.1 / 9.1 | **No. of Peptides:** | 3 |
| **Modification(s):** | Carbamidomethyl | | |

```
         10         20         30         40         50         60         70         80         90        100
MEFGLSWVFL VALLRGVQCQ VQLVESGGGV VQPGRSLRLS CAASGFTFSS YGMHWVRQAP GKGLEWVAVI WYDGSNKYYA DSVKGRFTIS RDNSKNTLYL
        110        120
QMNSLRAEDT AVYYCAR
```

MS/MS Peptide Matches

| Cmpd. | m/z meas. | z | Δ m/z [ppm] | Δ m/z [Da] | Rt [min] | Score | No. of Cmpds. | Site [%] | P | Sequence | Modification | Range | Type |
|---|---|---|---|---|---|---|---|---|---|---|---|---|---|
| 271 | 423.255 | 2 | 116.69 | 0.0494 | 16.39 | 24.6 | 3 | 0.0 | 0 | K.YYADSVK.G | | 78-84 | CID |
| 360 | 418.804 | 2 | 151.66 | 0.0635 | 18.20 | 19.77 | 2 | 0.0 | 1 | K.GRFTISR.D | | 85-91 | CID |
| 456 | 659.846 | 2 | 84.49 | 0.0557 | 20.39 | 56.97 | 4 | 0.0 | 0 | R.AEDTAVYYCAR.- | Carbamidomethyl: 9 | 107-117 | CID |

#### Protein 11: Serum albumin OS=Homo sapiens GN=ALB PE=1 SV=2

| | | | |
|---|---|---|---|
| **Accession:** | ALBU_HUMAN | **Score:** | 99.0 |
| **Database:** | SwissProt | **Seq. Coverage [%]:** | 7.4 |
| **MW [kDa] / pI:** | 69.3 / 5.9 | **No. of Peptides:** | 3 |
| **Modification(s):** | Carbamidomethyl | | |

| 10 | 20 | 30 | 40 | 50 | 60 | 70 | 80 | 90 | 100 |
|---|---|---|---|---|---|---|---|---|---|
| MKWVTFISLL | FLFSSAYSRG | VFRRDAHKSE | VAHRFKDLGE | ENFKALVLIA | FAQYLQQCPF | EDHVKLVNEV | TEFAKTCVAD | ESAENCDKSL | HTLFGDKLCT |
| 110 | 120 | 130 | 140 | 150 | 160 | 170 | 180 | 190 | 200 |
| VATLRETYGE | MADCCAKQEP | ERNECFLQHK | DDNPNLPRLV | RPEVDVMCTA | FHDNEETFLK | KYLYEIARRH | PYFYAPELLF | FAKRYKAAFT | ECCQAADKAA |
| 210 | 220 | 230 | 240 | 250 | 260 | 270 | 280 | 290 | 300 |
| CLLPKLDELR | DEGKASSAKQ | RLKCASLQKF | GERAFKAWAV | ARLSQRFPKA | EFAEVSKLVT | DLTKVHTECC | HGDLLECADD | RADLAKYICE | NQDSISSKLK |
| 310 | 320 | 330 | 340 | 350 | 360 | 370 | 380 | 390 | 400 |
| ECCEKPLLEK | SHCIAEVEND | EMPADLPSLA | ADFVESKDVC | KNYAEAKDVF | LGMFLYEYAR | RHPDYSVVLL | LRLAKTYETT | LEKCCAAADP | HECYAKVFDE |
| 410 | 420 | 430 | 440 | 450 | 460 | 470 | 480 | 490 | 500 |
| FKPLVEEPQN | LIKQNCELFE | QLGEYKFQNA | LLVRYTKKVP | QVSTPTLVEV | SRNLGKVGSK | CCKHPEAKRM | PCAEDYLSVV | LNQLCVLHEK | TPVSDRVTKC |
| 510 | 520 | 530 | 540 | 550 | 560 | 570 | 580 | 590 | 600 |
| CTESLVNRRP | CFSALEVDET | YVPKEFNAET | FTFHADICTL | SEKERQIKKQ | TALVELVKHK | PKATKEQLKA | VMDDFAAFVE | KCCKADDKET | CFAEEGKKLV |
| 610 | | | | | | | | | |
| AASQAALGL | | | | | | | | | |

MS/MS Peptide Matches

| Cmpd. | m/z meas. | z | Δ m/z [ppm] | Δ m/z [Da] | Rt [min] | Score | No. of Cmpds. | Site [%] | P | Sequence | Modification | Range | Type |
|---|---|---|---|---|---|---|---|---|---|---|---|---|---|
| 1410 | 831.199 | 3 | 520.59 | 0.4325 | 39.56 | 27.87 | 1 | 0.0 | 0 | K.ALVLIAFAQYLQQCPFEDHVK.L | Carbamidomethyl: 14 | 45-65 | CID |
| 370 | 516.304 | 3 | 65.01 | 0.0336 | 18.45 | 18.87 | 2 | 0.0 | 1 | K.LKECCEKPLLEK.S | Carbamidomethyl: 4, 5 | 299-310 | CID |
| 872 | 490.017 | 3 | 131.57 | 0.0645 | 28.77 | 52.27 | 1 | 0.0 | 1 | R.RHPDYSVVLLLR.L | | 361-372 | CID |

#### Protein 12: Immunoglobulin lambda variable 3-10 OS=Homo sapiens GN=IGLV3-10 PE=3 SV=2

**Accession:** LV310_HUMAN
**Database:** SwissProt
**MW [kDa] / pI:** 12.4 / 4.7
**Modification(s):** Carbamidomethyl

**Score:** 93.9
**Seq. Coverage [%]:** 15.7
**No. of Peptides:** 2

| 10 | 20 | 30 | 40 | 50 | 60 | 70 | 80 | 90 | 100 |
|---|---|---|---|---|---|---|---|---|---|
| MAWTPLLLPL | LTFCTVSEAS | YELTQPPSVS | VSPGQTARIT | CSGDALPKKY | AYWYQQKSGQ | APVLVIYEDS | KRPSGIPERF | SGSSSGTMAT | LTISGAQVED |
| 110 | 120 | | | | | | | | |
| EADYYCYSTD | SSGNH | | | | | | | | |

MS/MS Peptide Matches

| Cmpd. | m/z meas. | z | Δ m/z [ppm] | Δ m/z [Da] | Rt [min] | Score | No. of Cmpds. | Site [%] | P | Sequence | Modification | Range | Type |
|---|---|---|---|---|---|---|---|---|---|---|---|---|---|
| 397 | 531.336 | 2 | 127.20 | 0.0676 | 19.01 | 64.17 | 3 | 0.0 | 0 | R.ITCSGDALPK.K | Carbamidomethyl: 3 | 39-48 | CID |
| 220 | 456.323 | 2 | 145.73 | 0.0665 | 15.19 | 29.72 | 3 | 0.0 | 0 | K.RPSGIPER.F | | 72-79 | CID |

#### Protein 13: Immunoglobulin kappa variable 3-11 OS=Homo sapiens GN=IGKV3-11 PE=1 SV=1

**Accession:** KV311_HUMAN
**Database:** SwissProt
**MW [kDa] / pI:** 12.6 / 4.9

**Score:** 67.1
**Seq. Coverage [%]:** 7.8
**No. of Peptides:** 1

| 10 | 20 | 30 | 40 | 50 | 60 | 70 | 80 | 90 | 100 |
|---|---|---|---|---|---|---|---|---|---|
| MEAPAQLLFL | LLLWLPDTTG | EIVLTQSPAT | LSLSPGERAT | LSCRASQSVS | SYLAWYQQKP | GQAPRLLIYD | ASNRATGIPA | RFSGSGSGTD | FTLTISSLEP |
| 110 | 120 | | | | | | | | |
| EDFAVYYCQQ | RSNWP | | | | | | | | |

MS/MS Peptide Matches

| Cmpd. | m/z meas. | z | Δ m/z [ppm] | Δ m/z [Da] | Rt [min] | Score | No. of Cmpds. | Site [%] | P | Sequence | Modification | Range | Type |
|---|---|---|---|---|---|---|---|---|---|---|---|---|---|
| 655 | 532.835 | 2 | 83.75 | 0.0446 | 24.31 | 67.12 | 3 | 0.0 | 0 | R.LLIYDASNR.A | | 66-74 | CID |

##### Protein 14: Immunoglobulin kappa variable 3-15 OS=Homo sapiens GN=IGKV3-15 PE=1 SV=2

| | | | |
|---|---|---|---|
| **Accession:** | KV315_HUMAN | **Score:** | 62.9 |
| **Database:** | SwissProt | **Seq. Coverage [%]:** | 7.8 |
| **MW [kDa] / pI:** | 12.5 / 5.1 | **No. of Peptides:** | 1 |

```
        10         20         30         40         50         60         70         80         90        100
MEAPAQLLFL LLLWLPDTTG EIVMTQSPAT LSVSPGERAT LSCRASQSVS SNLAWYQQKP GQAPRLLIYG ASTRATGIPA RFSGSGSGTE FTLTISSLQS
       110        120
EDFAVYYCQQ YNNWP
```

MS/MS Peptide Matches

| Cmpd. | m/z meas. | z | Δ m/z [ppm] | Δ m/z [Da] | Rt [min] | Score | No. of Cmpds. | Site [%] | P | Sequence | Modification | Range | Type |
|---|---|---|---|---|---|---|---|---|---|---|---|---|---|
| 663 | 497.365 | 2 | 150.78 | 0.0750 | 24.45 | 62.86 | 7 | 0.0 | 0 | R.LLIYGASTR.A | | 66-74 | CID |

##### Protein 15: Immunoglobulin kappa variable 2D-28 OS=Homo sapiens GN=IGKV2D-28 PE=1 SV=2

| | | | |
|---|---|---|---|
| **Accession:** | KVD28_HUMAN | **Score:** | 57.9 |
| **Database:** | SwissProt | **Seq. Coverage [%]:** | 16.7 |
| **MW [kDa] / pI:** | 12.9 / 5.6 | **No. of Peptides:** | 1 |

```
        10         20         30         40         50         60         70         80         90        100
MRLPAQLLGL LMLWVSGSSG DIVMTQSPLS LPVTPGEPAS ISCRSSQSLL HSNGYNYLDW YLQKPGQSPQ LLIYLGSNRA SGVPDRFSGS GSGTDFTLKI
       110        120        130
SRVEAEDVGV YYCMQALQTP
```

MS/MS Peptide Matches

| Cmpd. | m/z meas. | z | Δ m/z [ppm] | Δ m/z [Da] | Rt [min] | Score | No. of Cmpds. | Site [%] | P | Sequence | Modification | Range | Type |
|---|---|---|---|---|---|---|---|---|---|---|---|---|---|
| 684 | 662.709 | 3 | 78.54 | 0.0520 | 24.86 | 57.9 | 2 | 0.0 | 1 | R.ASGVPDRFSGSGSGTDFTLK.I | | 80-99 | CID |

#### Protein 16: Immunoglobulin kappa variable 3-20 OS=Homo sapiens GN=IGKV3-20 PE=1 SV=2

**Accession:** KV320_HUMAN
**Database:** SwissProt
**MW [kDa] / pI:** 12.5 / 4.9

**Score:** 55.2
**Seq. Coverage [%]:** 7.8
**No. of Peptides:** 1

| 10 | 20 | 30 | 40 | 50 | 60 | 70 | 80 | 90 | 100 |
|---|---|---|---|---|---|---|---|---|---|
| METPAQLLFL | LLLWLPDTTG | EIVLTQSPGT | LSLSPGERAT | LSCRASQSVS | SSYLAWYQQK | PGQAPRLLIY | GASSRATGIP | DRFSGSGSGT | DFTLTISRLE |
| **110** | **120** | | | | | | | | |
| PEDFAVYYCQ | QYGSSP | | | | | | | | |

MS/MS Peptide Matches

| Cmpd. | m/z meas. | z | Δ m/z [ppm] | Δ m/z [Da] | Rt [min] | Score | No. of Cmpds. | Site [%] | P | Sequence | Modification | Range | Type |
|---|---|---|---|---|---|---|---|---|---|---|---|---|---|
| 645 | 490.353 | 2 | 144.42 | 0.0708 | 24.11 | 55.19 | 1 | 0.0 | 0 | R.LLIYGASSR.A | | 67-75 | CID |

#### Protein 17: Immunoglobulin heavy variable 5-51 OS=Homo sapiens GN=IGHV5-51 PE=3 SV=1

**Accession:** HV551_HUMAN
**Database:** SwissProt
**MW [kDa] / pI:** 12.7 / 8.4
**Modification(s):** Carbamidomethyl

**Score:** 49.4
**Seq. Coverage [%]:** 9.4
**No. of Peptides:** 1

| 10 | 20 | 30 | 40 | 50 | 60 | 70 | 80 | 90 | 100 |
|---|---|---|---|---|---|---|---|---|---|
| MGSTAILALL | LAVLQGVCAE | VQLVQSGAEV | KKPGESLKIS | CKGSGYSFTS | YWIGWVRQMP | GKGLEWMGII | YPGDSDTRYS | PSFQGQVTIS | ADKSISTAYL |
| 110 | 120 | | | | | | | | |
| QWSSLKASDT | AMYYCAR | | | | | | | | |

MS/MS Peptide Matches

| Cmpd. | m/z meas. | z | Δ m/z [ppm] | Δ m/z [Da] | Rt [min] | Score | No. of Cmpds. | Site [%] | P | Sequence | Modification | Range | Type |
|---|---|---|---|---|---|---|---|---|---|---|---|---|---|
| 473 | 654.69 | 2 | -123.72 | -0.0810 | 20.75 | 49.35 | 2 | 0.0 | 0 | K.ASDTAMYYCAR.- | Carbamidomethyl: 9 | 107-117 | CID |

BRUKER

#### Project Info

| | | | |
|---|---|---|---|
| **Name:** | Alagesan | **Date:** | Mar 5, 2019 |
| **Note:** | Nanobooster | | |

#### Sample Info & Protocols

| | |
|---|---|
| **Name:** | Nanobooster - IgG |
| **Date:** | Mar 5, 2019 |

#### Search Result Info

| | | | |
|---|---|---|---|
| **Search Result:** | IonTrap_Human_Swiss_CID_2019-03-05 17:55:08 | | |
| **Location:** | /Alagesan/Nanobooster - IgG/Ka_IgG Trypsin No Enrich No NB_BA5_01_8734.mgf | | |
| **Search Method:** | IonTrap_Human_Swiss_CID | **Version:** | 1.2 |
| **Search Engine(s):** | Mascot, 2.6.0 | **Ident. Compound(s):** | 0/2308 |
| **Database(s):** | SwissProt, SwissProt_2017_04.fasta | | |
| **Note:** | | | |

#### Results

##### Protein 1: Immunoglobulin heavy constant gamma 1 OS=Homo sapiens GN=IGHG1 PE=1 SV=1

| | | | |
|---|---|---|---|
| **Accession:** | IGHG1_HUMAN | **Score:** | 1184.1 |
| **Database:** | SwissProt | **Seq. Coverage [%]:** | 65.2 |
| **MW [kDa] / pI:** | 36.1 / 8.5 | **No. of Peptides:** | 17 |
| **Modification(s):** | Carbamidomethyl, Oxidation | | |

| 10 | 20 | 30 | 40 | 50 | 60 | 70 | 80 | 90 | 100 |
|---|---|---|---|---|---|---|---|---|---|
| ASTKGPSVFP | LAPSSKSTSG | GTAALGCLVK | DYFPEPVTVS | WNSGALTSGV | HTFPAVLQSS | GLYSLSSVVT | VPSSSLGTQT | YICNVNHKPS | NTKVDKKVEP |
| **110** | **120** | **130** | **140** | **150** | **160** | **170** | **180** | **190** | **200** |
| KSCDKTHTCP | PCPAPELLGG | PSVFLFPPKP | KDTLMISRTP | EVTCVVVDVS | HEDPEVKFNW | YVDGVEVHNA | KTKPREEQYN | STYRVVSVLT | VLHQDWLNGK |
| **210** | **220** | **230** | **240** | **250** | **260** | **270** | **280** | **290** | **300** |
| EYKCKVSNKA | LPAPIEKTIS | KAKGQPREPQ | VYTLPPSRDE | LTKNQVSLTC | LVKGFYPSDI | AVEWESNGQP | ENNYKTTPPV | LDSDGSFFLY | SKLTVDKSRW |
| **310** | **320** | **330** | **340** | | | | | | |
| QQGNVFSCSV | MHEALHNHYT | QKSLSLSPGK | | | | | | | |

**MS/MS Peptide Matches**

| Cmpd. | m/z meas. | z | Δ m/z [ppm] | Δ m/z [Da] | Rt [min] | Score | No. of Cmpds. | Site [%] | P | Sequence | Modification | Range | Type |
|---|---|---|---|---|---|---|---|---|---|---|---|---|---|
| 805 | 593.888 | 2 | 102.79 | 0.0610 | 27.51 | 52.97 | 8 | 0.0 | 0 | K.GPSVFPLAPSSK.S | | 5-16 | CID |
| 694 | 661.419 | 2 | 115.44 | 0.0763 | 25.16 | 132.45 | 11 | 0.0 | 0 | K.STSGGTAALGCLVK.D | Carbamidomethyl: 11 | 17-30 | CID |
| 1032 | 834.765 | 4 | 418.27 | 0.3490 | 32.01 | 41.46 | 2 | 0.0 | 1 | K.SCDKTHTCPPCPAPELLGGPSVFLFPPKPK.D | Carbamidomethyl: 2, 8, 11 | 102-131 | CID |
| 1068 | 949.222 | 3 | 419.43 | 0.3980 | 32.78 | 76.34 | 9 | 0.0 | 0 | K.THTCPPCPAPELLGGPSVFLFPPKPK.D | Carbamidomethyl: 4, 7 | 106-131 | CID |
| 529 | 418.264 | 2 | 103.43 | 0.0433 | 21.94 | 47.56 | 7 | 0.0 | 0 | K.DTLMISR.T | | 132-138 | CID |
| 428 | 426.274 | 2 | 130.91 | 0.0558 | 19.84 | 41.03 | 3 | 0.0 | 0 | K.DTLMISR.T | Oxidation: 4 | 132-138 | CID |
| 794 | 713.737 | 3 | 78.94 | 0.0563 | 27.27 | 84.85 | 3 | 0.0 | 0 | R.TPEVTCVVVDVSHEDPEVK.F | Carbamidomethyl: 6 | 139-157 | CID |
| 785 | 839.479 | 2 | 88.60 | 0.0744 | 27.10 | 112.53 | 10 | 0.0 | 0 | K.FNWYVDGVEVHNAK.T | | 158-171 | CID |

| Cmpd. | m/z meas. | z | Δ m/z [ppm] | Δ m/z [Da] | Rt [min] | Score | No. of Cmpds. | Site [%] | P | Sequence | Modification | Range | Type |
|---|---|---|---|---|---|---|---|---|---|---|---|---|---|
| 1137 | 603.386 | 3 | 75.66 | 0.0457 | 34.26 | 82.54 | 10 | 0.0 | 0 | R.VVSVLTVLHQDWLNGK.E | | 185-200 | CID |
| 507 | 419.814 | 2 | 139.90 | 0.0587 | 21.51 | 21.91 | 2 | 0.0 | 0 | K.ALPAPIEK.T | | 210-217 | CID |
| 610 | 643.86 | 2 | 30.13 | 0.0194 | 23.49 | 40.08 | 6 | 0.0 | 0 | R.EPQVYTLPPSR.D | | 228-238 | CID |
| 643 | 937.03 | 2 | 44.05 | 0.0413 | 24.15 | 50.07 | 10 | 0.0 | 1 | R.EPQVYTLPPSRDELTK.N | | 228-243 | CID |
| 772 | 581.383 | 2 | 111.04 | 0.0646 | 26.85 | 80.28 | 14 | 0.0 | 0 | K.NQVSLTCLVK.G | Carbamidomethyl: 7 | 244-253 | CID |
| 969 | 1273.035 | 2 | 365.93 | 0.4657 | 30.78 | 114.44 | 37 | 0.0 | 0 | K.GFYPSDIAVEWESNGQPENNYK.T | | 254-275 | CID |
| 1055 | 937.556 | 2 | 97.55 | 0.0915 | 32.49 | 71.95 | 17 | 0.0 | 0 | K.TTPPVLDSDGSFFLYSK.L | | 276-292 | CID |
| 157 | 409.816 | 2 | 185.09 | 0.0758 | 13.70 | 49.06 | 4 | 0.0 | 1 | K.LTVDKSR.W | | 293-299 | CID |
| 739 | 934.834 | 3 | 435.32 | 0.4068 | 26.16 | 84.54 | 7 | 0.0 | 0 | R.WQQGNVFSCSVMHEALHNHYTQK.S | Carbamidomethyl: 9 | 300-322 | CID |

#### Protein 2: Immunoglobulin heavy constant gamma 2 OS=Homo sapiens GN=IGHG2 PE=1 SV=2

| | | | |
|---|---|---|---|
| **Accession:** | IGHG2_HUMAN | **Score:** | 1033.1 |
| **Database:** | SwissProt | **Seq. Coverage [%]:** | 48.8 |
| **MW [kDa] / pI:** | 35.9 / 7.7 | **No. of Peptides:** | 16 |
| **Modification(s):** | Carbamidomethyl, Oxidation | | |

| 10 | 20 | 30 | 40 | 50 | 60 | 70 | 80 | 90 | 100 |
|---|---|---|---|---|---|---|---|---|---|
| ASTKGPSVFP | LAPCSRSTSE | STAALGCLVK | DYFPEPVTVS | WNSGALTSGV | HTFPAVLQSS | GLYSLSSVVT | VPSSNFGTQT | YTCNVDHKPS | NTKVDKTVER |
| **110** | **120** | **130** | **140** | **150** | **160** | **170** | **180** | **190** | **200** |
| KCCVECPPCP | APPVAGPSVF | LFPPKPKDTL | MISRTPEVTC | VVVDVSHEDP | EVQFNWYVDG | VEVHNAKTKP | REEQFNSTFR | VVSVLTVVHQ | DWLNGKEYKC |
| **210** | **220** | **230** | **240** | **250** | **260** | **270** | **280** | **290** | **300** |
| KVSNKGLPAP | IEKTISKTKG | QPREPQVYTL | PPSREEMTKN | QVSLTCLVKG | FYPSDISVEW | ESNGQPENNY | KTTPPMLDSD | GSFFLYSKLT | VDKSRWQQGN |
| **310** | **320** | **330** | | | | | | | |
| VFSCSVMHEA | LHNHYTQKSL | SLSPGK | | | | | | | |

MS/MS Peptide Matches

| Cmpd. | m/z meas. | z | Δ m/z [ppm] | Δ m/z [Da] | Rt [min] | Score | No. of Cmpds. | Site [%] | P | Sequence | Modification | Range | Type |
|---|---|---|---|---|---|---|---|---|---|---|---|---|---|
| 840 | 644.4 | 2 | 109.65 | 0.0707 | 28.24 | 59.86 | 11 | 0.0 | 0 | K.GPSVFPLAPCSR.S | Carbamidomethyl: 10 | 5-16 | CID |
| 703 | 712.396 | 2 | 52.64 | 0.0375 | 25.41 | 83.86 | 11 | 0.0 | 0 | R.STSESTAALGCLVK.D | Carbamidomethyl: 11 | 17-30 | CID |
| 112 | 423.812 | 2 | 175.54 | 0.0744 | 12.31 | 44.5 | 3 | 0.0 | 1 | K.VDKTVER.K | | 94-100 | CID |
| 968 | 1013.183 | 3 | 341.55 | 0.3459 | 30.74 | 83.5 | 2 | 0.0 | 1 | R.KCCVECPPCPAPPVAGPSVFLFPPKPK.D | Carbamidomethyl: 2, 3, 6, 9 | 101-127 | CID |
| 1036 | 970.519 | 3 | 391.96 | 0.3803 | 32.07 | 71.23 | 9 | 0.0 | 0 | K.CCVECPPCPAPPVAGPSVFLFPPKPK.D | Carbamidomethyl: 1, 2, 5, 8 | 102-127 | CID |
| 529 | 418.264 | 2 | 103.43 | 0.0433 | 21.94 | 47.56 | 7 | 0.0 | 0 | K.DTLMISR.T | | 128-134 | CID |
| 428 | 426.274 | 2 | 130.91 | 0.0558 | 19.84 | 41.03 | 3 | 0.0 | 0 | K.DTLMISR.T | Oxidation: 4 | 128-134 | CID |
| 1035 | 898.07 | 2 | 636.15 | 0.5709 | 32.06 | 53.04 | 3 | 0.0 | 0 | R.VVSVLTVVHQDWLNGK.E | | 181-196 | CID |
| 966 | 738.808 | 3 | 98.24 | 0.0726 | 30.72 | 76.61 | 3 | 0.0 | 1 | R.VVSVLTVVHQDWLNGKEYK.C | | 181-199 | CID |
| 610 | 643.86 | 2 | 30.13 | 0.0194 | 23.49 | 40.08 | 6 | 0.0 | 0 | R.EPQVYTLPPSR.E | | 224-234 | CID |
| 618 | 952.997 | 2 | 23.34 | 0.0222 | 23.70 | 96.77 | 18 | 0.0 | 1 | R.EPQVYTLPPSREEMTK.N | | 224-239 | CID |
| 772 | 581.383 | 2 | 111.04 | 0.0646 | 26.85 | 80.28 | 14 | 0.0 | 0 | K.NQVSLTCLVK.G | Carbamidomethyl: 7 | 240-249 | CID |
| 1049 | 953.465 | 2 | 15.12 | 0.0144 | 32.33 | 102.49 | 9 | 0.0 | 0 | K.TTPPMLDSDGSFFLYSK.L | | 272-288 | CID |
| 965 | 961.495 | 2 | 48.84 | 0.0470 | 30.68 | 18.74 | 3 | 0.0 | 0 | K.TTPPMLDSDGSFFLYSK.L | Oxidation: 5 | 272-288 | CID |
| 157 | 409.816 | 2 | 185.09 | 0.0758 | 13.70 | 49.06 | 4 | 0.0 | 1 | K.LTVDKSR.W | | 289-295 | CID |
| 739 | 934.834 | 3 | 435.32 | 0.4068 | 26.16 | 84.54 | 7 | 0.0 | 0 | R.WQQGNVFSCSVMHEALHNHYTQK.S | Carbamidomethyl: 9 | 296-318 | CID |

#### Protein 3: Immunoglobulin kappa constant OS=Homo sapiens GN=IGKC PE=1 SV=2

**Accession:** IGKC_HUMAN
**Database:** SwissProt
**MW [kDa] / pI:** 11.8 / 6.1
**Modification(s):** Carbamidomethyl

**Score:** 932.6
**Seq. Coverage [%]:** 93.5
**No. of Peptides:** 10

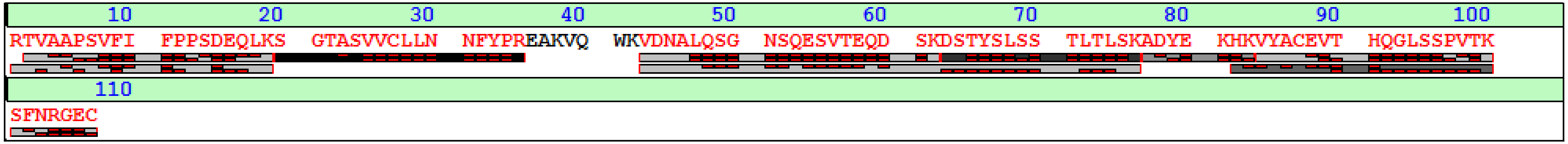


MS/MS Peptide Matches

| Cmpd. | m/z meas. | z | Δ m/z [ppm] | Δ m/z [Da] | Rt [min] | Score | No. of Cmpds. | Site [%] | P | Sequence | Modification | Range | Type |
|---|---|---|---|---|---|---|---|---|---|---|---|---|---|
| 985 | 1051.613 | 2 | 43.11 | 0.0453 | 31.04 | 42.51 | 3 | 0.0 | 1 | -.RTVAAPSVFIFPPSDEQLK.S | | 1-19 | CID |
| 1103 | 973.675 | 2 | 162.18 | 0.1579 | 33.49 | 86.97 | 30 | 0.0 | 0 | R.TVAAPSVFIFPPSDEQLK.S | | 2-19 | CID |
| 1184 | 899.556 | 2 | 116.46 | 0.1047 | 35.25 | 68.37 | 25 | 0.0 | 0 | K.SGTASVVCLLNNFYPR.E | Carbamidomethyl: 8 | 20-35 | CID |
| 435 | 1068.53 | 2 | 39.31 | 0.0420 | 19.98 | 169.85 | 49 | 0.0 | 0 | K.VDNALQSGNSQESVTEQDSK.D | | 43-62 | CID |
| 813 | 1207.559 | 3 | 263.17 | 0.3177 | 27.64 | 155.08 | 46 | 0.0 | 1 | K.VDNALQSGNSQESVTEQDSKDSTYSLSSTLTLS K.A | | 43-76 | CID |
| 831 | 751.956 | 2 | 97.28 | 0.0731 | 27.99 | 86.19 | 5 | 0.0 | 0 | K.DSTYSLSSTLTLSK.A | | 63-76 | CID |
| 90 | 445.801 | 2 | 177.32 | 0.0790 | 11.34 | 31.68 | 6 | 0.0 | 1 | K.ADYEKHK.V | | 77-83 | CID |
| 403 | 714.403 | 3 | 53.03 | 0.0379 | 19.34 | 123.98 | 7 | 0.0 | 1 | K.HKVYACEVTHQGLSSPVTK.S | Carbamidomethyl: 6 | 82-100 | CID |
| 521 | 938.507 | 2 | 42.52 | 0.0399 | 21.79 | 113.57 | 6 | 0.0 | 0 | K.VYACEVTHQGLSSPVTK.S | Carbamidomethyl: 4 | 84-100 | CID |
| 168 | 435.244 | 2 | 142.12 | 0.0618 | 13.97 | 54.37 | 7 | 0.0 | 1 | K.SFNRGEC.- | Carbamidomethyl: 7 | 101-107 | CID |

#### Protein 4: Immunoglobulin heavy constant gamma 3 OS=Homo sapiens GN=IGHG3 PE=1 SV=2

**Accession:** IGHG3_HUMAN
**Database:** SwissProt
**MW [kDa] / pI:** 41.3 / 8.2
**Modification(s):** Carbamidomethyl, Oxidation

**Score:** 883.5
**Seq. Coverage [%]:** 35.0
**No. of Peptides:** 13

| 10 | 20 | 30 | 40 | 50 | 60 | 70 | 80 | 90 | 100 |
|---|---|---|---|---|---|---|---|---|---|
| ASTKGPSVFP | LAPCSRSTSG | GTAALGCLVK | DYFPEPVTVS | WNSGALTSGV | HTFPAVLQSS | GLYSLSSVVT | VPSSSLGTQT | YTCNVNHKPS | NTKVDKRVEL |
| **110** | **120** | **130** | **140** | **150** | **160** | **170** | **180** | **190** | **200** |
| KTPLGDTTHT | CPRCPEPKSC | DTPPPCPRCP | EPKSCDTPPP | CPRCPEPKSC | DTPPPCPRCP | APELLGGPSV | FLFPPKPKDT | LMISRTPEVT | CVVVDVSHED |
| **210** | **220** | **230** | **240** | **250** | **260** | **270** | **280** | **290** | **300** |
| PEVQFKWYVD | GVEVHNAKTK | PREEQYNSTF | RVVSVLTVLH | QDWLNGKEYK | CKVSNKALPA | PIEKTISKTK | GQPREPQVYT | LPPSREEMTK | NQVSLTCLVK |
| **310** | **320** | **330** | **340** | **350** | **360** | **370** | **380** | | |
| GFYPSDIAVE | WESSGQPENN | YNTTPPMLDS | DGSFFLYSKL | TVDKSRWQQG | NIFSCSVMHE | ALHNRFTQKS | LSLSPGK | | |

**MS/MS Peptide Matches**

| Cmpd. | m/z meas. | z | Δ m/z [ppm] | Δ m/z [Da] | Rt [min] | Score | No. of Cmpds. | Site [%] | P | Sequence | Modification | Range | Type |
|---|---|---|---|---|---|---|---|---|---|---|---|---|---|
| 840 | 644.4 | 2 | 109.65 | 0.0707 | 28.24 | 59.86 | 11 | 0.0 | 0 | K.GPSVFPLAPCSR.S | Carbamidomethyl: 10 | 5-16 | CID |
| 694 | 661.419 | 2 | 115.44 | 0.0763 | 25.16 | 132.45 | 11 | 0.0 | 0 | R.STSGGTAALGCLVK.D | Carbamidomethyl: 11 | 17-30 | CID |
| 295 | 452.58 | 3 | 64.98 | 0.0294 | 16.98 | 68.04 | 3 | 0.0 | 0 | K.TPLGDTTHTCPR.C | Carbamidomethyl: 10 | 102-113 | CID |
| 259 | 593.787 | 2 | 57.91 | 0.0344 | 16.14 | 68.47 | 12 | 0.0 | 0 | K.SCDTPPPCPR.C | Carbamidomethyl: 2, 8 | 119-128 | CID |
| 1128 | 717.794 | 3 | 93.05 | 0.0668 | 34.09 | 95.43 | 3 | 0.0 | 0 | R.CPAPELLGGPSVFLFPPKPK.D | Carbamidomethyl: 1 | 159-178 | CID |
| 529 | 418.264 | 2 | 103.43 | 0.0433 | 21.94 | 47.56 | 7 | 0.0 | 0 | K.DTLMISR.T | | 179-185 | CID |
| 428 | 426.274 | 2 | 130.91 | 0.0558 | 19.84 | 41.03 | 3 | 0.0 | 0 | K.DTLMISR.T | Oxidation: 4 | 179-185 | CID |
| 1137 | 603.386 | 3 | 75.66 | 0.0457 | 34.26 | 82.54 | 10 | 0.0 | 0 | R.VVSVLTVLHQDWLNGK.E | | 232-247 | CID |
| 507 | 419.814 | 2 | 139.90 | 0.0587 | 21.51 | 21.91 | 2 | 0.0 | 0 | K.ALPAPIEK.T | | 257-264 | CID |
| 610 | 643.86 | 2 | 30.13 | 0.0194 | 23.49 | 40.08 | 6 | 0.0 | 0 | R.EPQVYTLPPSR.E | | 275-285 | CID |
| 618 | 952.997 | 2 | 23.34 | 0.0222 | 23.70 | 96.77 | 18 | 0.0 | 1 | R.EPQVYTLPPSREEMTK.N | | 275-290 | CID |
| 772 | 581.383 | 2 | 111.04 | 0.0646 | 26.85 | 80.28 | 14 | 0.0 | 0 | K.NQVSLTCLVK.G | Carbamidomethyl: 7 | 291-300 | CID |
| 157 | 409.816 | 2 | 185.09 | 0.0758 | 13.70 | 49.06 | 4 | 0.0 | 1 | K.LTVDKSR.W | | 340-346 | CID |

#### Protein 5: Immunoglobulin kappa light chain OS=Homo sapiens PE=1 SV=1

| | | | |
|---|---|---|---|
| **Accession:** | IGK_HUMAN | **Score:** | 865.0 |
| **Database:** | SwissProt | **Seq. Coverage [%]:** | 43.0 |
| **MW [kDa] / pI:** | 23.4 / 6.9 | **No. of Peptides:** | 9 |
| **Modification(s):** | Carbamidomethyl | | |

| 10 | 20 | 30 | 40 | 50 | 60 | 70 | 80 | 90 | 100 |
|---|---|---|---|---|---|---|---|---|---|
| DIQMTQSPST | LSASVGDRVT | ITCRASQSIN | TWLAWYQQKP | GKAPKLLMYK | ASSLESGVPS | RFIGSGSGTE | FTLTISSLQP | DDFATYYCQQ | YNSDSKMFGQ |
| **110** | **120** | **130** | **140** | **150** | **160** | **170** | **180** | **190** | **200** |
| GTKVEVKGTV | AAPSVFIFPP | SDEQLKSGTA | SVVCLLNNFY | PREAKVQWKV | DNALQSGNSQ | ESVTEQDSKD | STYSLSSTLT | LSKADYEKHK | VYACEVTHQG |
| **210** | **220** | | | | | | | | |
| LSSPVTKSFN | RGEC | | | | | | | | |

**MS/MS Peptide Matches**

| Cmpd. | m/z meas. | z | Δ m/z [ppm] | Δ m/z [Da] | Rt [min] | Score | No. of Cmpds. | Site [%] | P | Sequence | Modification | Range | Type |
|---|---|---|---|---|---|---|---|---|---|---|---|---|---|
| 375 | 545.316 | 2 | 65.33 | 0.0356 | 18.67 | 61.87 | 3 | 0.0 | 0 | K.ASSLESGVPSR.F | | 51-61 | CID |
| 1184 | 899.556 | 2 | 116.46 | 0.1047 | 35.25 | 68.37 | 25 | 0.0 | 0 | K.SGTASVVCLLNNFYPR.E | Carbamidomethyl: 8 | 127-142 | CID |
| 435 | 1068.53 | 2 | 39.31 | 0.0420 | 19.98 | 169.85 | 49 | 0.0 | 0 | K.VDNALQSGNSQESVTEQDSK.D | | 150-169 | CID |
| 813 | 1207.559 | 3 | 263.17 | 0.3177 | 27.64 | 155.08 | 46 | 0.0 | 1 | K.VDNALQSGNSQESVTEQDSKDSTYSLSSTLTLSK.A | | 150-183 | CID |
| 831 | 751.956 | 2 | 97.28 | 0.0731 | 27.99 | 86.19 | 5 | 0.0 | 0 | K.DSTYSLSSTLTLSK.A | | 170-183 | CID |
| 90 | 445.801 | 2 | 177.32 | 0.0790 | 11.34 | 31.68 | 6 | 0.0 | 1 | K.ADYEKHK.V | | 184-190 | CID |
| 403 | 714.403 | 3 | 53.03 | 0.0379 | 19.34 | 123.98 | 7 | 0.0 | 1 | K.HKVYACEVTHQGLSSPVTK.S | Carbamidomethyl: 6 | 189-207 | CID |
| 521 | 938.507 | 2 | 42.52 | 0.0399 | 21.79 | 113.57 | 6 | 0.0 | 0 | K.VYACEVTHQGLSSPVTK.S | Carbamidomethyl: 4 | 191-207 | CID |
| 168 | 435.244 | 2 | 142.12 | 0.0618 | 13.97 | 54.37 | 7 | 0.0 | 1 | K.SFNRGEC.- | Carbamidomethyl: 7 | 208-214 | CID |

#### Protein 6: Immunoglobulin heavy constant gamma 4 OS=Homo sapiens GN=IGHG4 PE=1 SV=1

| | | | |
|---|---|---|---|
| **Accession:** | IGHG4_HUMAN | **Score:** | 748.8 |
| **Database:** | SwissProt | **Seq. Coverage [%]:** | 51.4 |
| **MW [kDa] / pI:** | 35.9 / 7.2 | **No. of Peptides:** | 11 |
| **Modification(s):** | Carbamidomethyl, Oxidation | | |

```
        10         20         30         40         50         60         70         80         90        100
ASTKGPSVFP LAPCSRSTSE STAALGCLVK DYFPEPVTVS WNSGALTSGV HTFPAVLQSS GLYSLSSVVT VPSSSLGTKT YTCNVDHKPS NTKVDKRVES
       110        120        130        140        150        160        170        180        190        200
KYGPPCPSCP APEFLGGPSV FLFPPKPKDT LMISRTPEVT CVVVDVSQED PEVQFNWYVD GVEVHNAKTK PREEQFNSTY RVVSVLTVLH QDWLNGKEYK
       210        220        230        240        250        260        270        280        290        300
CKVSNKGLPS SIEKTISKAK GQPREPQVYT LPPSQEEMTK NQVSLTCLVK GFYPSDIAVE WESNGQPENN YKTTPPVLDS DGSFFLYSRL TVDKSRWQEG
       310        320        330
NVFSCSVMHE ALHNHYTQKS LSLSLGK
```

**MS/MS Peptide Matches**

| Cmpd. | m/z meas. | z | Δ m/z [ppm] | Δ m/z [Da] | Rt [min] | Score | No. of Cmpds. | Site [%] | P | Sequence | Modification | Range | Type |
|---|---|---|---|---|---|---|---|---|---|---|---|---|---|
| 840 | 644.4 | 2 | 109.65 | 0.0707 | 28.24 | 59.86 | 11 | 0.0 | 0 | K.GPSVFPLAPCSR.S | Carbamidomethyl: 10 | 5-16 | CID |
| 703 | 712.396 | 2 | 52.64 | 0.0375 | 25.41 | 83.86 | 11 | 0.0 | 0 | R.STSESTAALGCLVK.D | Carbamidomethyl: 11 | 17-30 | CID |
| 175 | 555.652 | 3 | 103.04 | 0.0573 | 14.16 | 43.75 | 7 | 0.0 | 0 | K.TYTCNVDHKPSNTK.V | Carbamidomethyl: 4 | 80-93 | CID |
| 1167 | 981.903 | 3 | 80.54 | 0.0791 | 34.88 | 79.09 | 3 | 0.0 | 0 | K.YGPPCPSCPAPEFLGGPSVFLFPPKPK.D | Carbamidomethyl: 5, 8 | 102-128 | CID |
| 529 | 418.264 | 2 | 103.43 | 0.0433 | 21.94 | 47.56 | 7 | 0.0 | 0 | K.DTLMISR.T | | 129-135 | CID |
| 428 | 426.274 | 2 | 130.91 | 0.0558 | 19.84 | 41.03 | 3 | 0.0 | 0 | K.DTLMISR.T | Oxidation: 4 | 129-135 | CID |
| 1137 | 603.386 | 3 | 75.66 | 0.0457 | 34.26 | 82.54 | 10 | 0.0 | 0 | R.VVSVLTVLHQDWLNGK.E | | 182-197 | CID |
| 689 | 939.491 | 2 | 572.46 | 0.5375 | 25.02 | 18.48 | 2 | 0.0 | 0 | R.EPQVYTLPPSQEEMTK.N | | 225-240 | CID |
| 772 | 581.383 | 2 | 111.04 | 0.0646 | 26.85 | 80.28 | 14 | 0.0 | 0 | K.NQVSLTCLVK.G | Carbamidomethyl: 7 | 241-250 | CID |

| Cmpd. | m/z meas. | z | Δ m/z [ppm] | Δ m/z [Da] | Rt [min] | Score | No. of Cmpds. | Site [%] | P | Sequence | Modification | Range | Type |
|---|---|---|---|---|---|---|---|---|---|---|---|---|---|
| 969 | 1273.035 | 2 | 365.93 | 0.4657 | 30.78 | 114.44 | 37 | 0.0 | 0 | K.GFYPSDIAVEWESNGQPENNYK.T | | 251-272 | CID |
| 157 | 409.816 | 2 | 185.09 | 0.0758 | 13.70 | 49.06 | 4 | 0.0 | 1 | R.LTVDKSR.W | | 290-296 | CID |

#### Protein 7: Immunoglobulin lambda constant 3 OS=Homo sapiens GN=IGLC3 PE=1 SV=1

**Accession:** IGLC3_HUMAN
**Database:** SwissProt
**MW [kDa] / pI:** 11.3 / 6.9
**Modification(s):** Carbamidomethyl

**Score:** 597.9
**Seq. Coverage [%]:** 93.4
**No. of Peptides:** 8

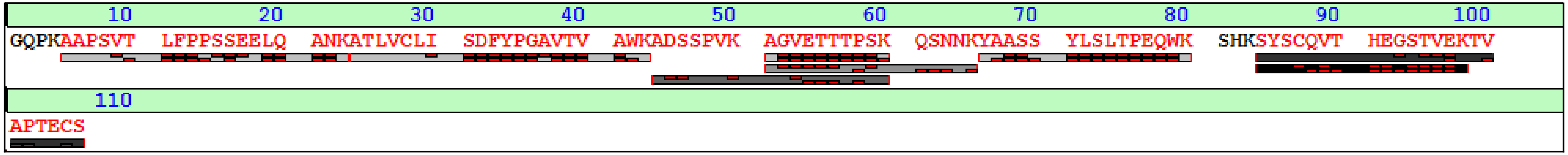


MS/MS Peptide Matches

| Cmpd. | m/z meas. | z | Δ m/z [ppm] | Δ m/z [Da] | Rt [min] | Score | No. of Cmpds. | Site [%] | P | Sequence | Modification | Range | Type |
|---|---|---|---|---|---|---|---|---|---|---|---|---|---|
| 873 | 993.515 | 2 | 2.45 | 0.0024 | 28.86 | 90.77 | 20 | 0.0 | 0 | K.AAPSVTLFPPSSEELQANK.A | | 5-23 | CID |
| 1435 | 1106.632 | 2 | 499.46 | 0.5524 | 40.21 | 73.78 | 3 | 0.0 | 0 | K.ATLVCLISDFYPGAVTVAWK.A | Carbamidomethyl: 5 | 24-43 | CID |
| 332 | 558.986 | 3 | 53.06 | 0.0297 | 17.81 | 37.98 | 1 | 0.0 | 1 | K.ADSSPVKAGVETTTPSK.Q | | 44-60 | CID |
| 198 | 495.764 | 2 | 10.60 | 0.0053 | 14.67 | 72.59 | 15 | 0.0 | 0 | K.AGVETTTPSK.Q | | 51-60 | CID |
| 149 | 521.325 | 3 | 114.34 | 0.0596 | 13.48 | 59.19 | 4 | 0.0 | 1 | K.AGVETTTPSKQSNNK.Y | | 51-65 | CID |
| 943 | 872.52 | 2 | 99.66 | 0.0869 | 30.21 | 86.96 | 12 | 0.0 | 0 | K.YAASSYLSLTPEQWK.S | | 66-80 | CID |
| 314 | 856.375 | 2 | -9.61 | -0.0082 | 17.40 | 118.26 | 7 | 0.0 | 0 | K.SYSCQVTHEGSTVEK.T | Carbamidomethyl: 4 | 84-98 | CID |
| 489 | 853.077 | 3 | 38.43 | 0.0328 | 21.15 | 58.35 | 3 | 0.0 | 1 | K.SYSCQVTHEGSTVEKTVAPTECS.- | Carbamidomethyl: 4, 22 | 84-106 | CID |

#### Protein 8: Immunoglobulin lambda-1 light chain OS=Homo sapiens PE=1 SV=1

**Accession:** IGL1_HUMAN
**Database:** SwissProt
**MW [kDa] / pI:** 22.8 / 6.3
**Modification(s):** Carbamidomethyl

**Score:** 584.4
**Seq. Coverage [%]:** 50.0
**No. of Peptides:** 9

| 10 | 20 | 30 | 40 | 50 | 60 | 70 | 80 | 90 | 100 |
|---|---|---|---|---|---|---|---|---|---|
| QSALTQPPSA | SGSLGQSVTI | SCTGTSSDVG | GYNYVSWYQQ | HAGKAPKVII | YEVNKRPSGV | PDRFSGSKSG | NTASLTVSGL | QAEDEADYYC | SSYEGSDNFV |
| 110 | 120 | 130 | 140 | 150 | 160 | 170 | 180 | 190 | 200 |
| FGTGTKVTVL | GQPKANPTVT | LFPPSSEELQ | ANKATLVCLI | SDFYPGAVTV | AWKADGSPVK | AGVETTKPSK | QSNNKYAASS | YLSLTPEQWK | SHRSYSCQVT |
| 210 | 220 | | | | | | | | |
| HEGSTVEKTV | APTECS | | | | | | | | |

##### MS/MS Peptide Matches

| Cmpd. | m/z meas. | z | Δ m/z [ppm] | Δ m/z [Da] | Rt [min] | Score | No. of Cmpds. | Site [%] | P | Sequence | Modification | Range | Type |
|---|---|---|---|---|---|---|---|---|---|---|---|---|---|
| 158 | 442.315 | 2 | 167.65 | 0.0741 | 13.71 | 26.85 | 3 | 0.0 | 0 | K.RPSGVPDR.F | | 56-63 | CID |
| 312 | 463.976 | 3 | 136.60 | 0.0634 | 17.34 | 25.13 | 3 | 0.0 | 1 | K.RPSGVPDRFSGSK.S | | 56-68 | CID |
| 425 | 421.314 | 2 | 126.46 | 0.0533 | 19.79 | 47.86 | 3 | 0.0 | 0 | K.VTVLGQPK.A | | 107-114 | CID |
| 903 | 1022.039 | 2 | 15.37 | 0.0157 | 29.43 | 74.38 | 13 | 0.0 | 0 | K.ANPTVTLFPPSSEELQANK.A | | 115-133 | CID |
| 1435 | 1106.632 | 2 | 499.46 | 0.5524 | 40.21 | 73.78 | 3 | 0.0 | 0 | K.ATLVCLISDFYPGAVTVAWK.A | Carbamidomethyl: 5 | 134-153 | CID |
| 129 | 509.357 | 2 | 146.50 | 0.0746 | 12.82 | 72.87 | 6 | 0.0 | 0 | K.AGVETTKPSK.Q | | 161-170 | CID |
| 943 | 872.52 | 2 | 99.66 | 0.0869 | 30.21 | 86.96 | 12 | 0.0 | 0 | K.YAASSYLSLTPEQWK.S | | 176-190 | CID |
| 314 | 856.375 | 2 | -9.61 | -0.0082 | 17.40 | 118.26 | 7 | 0.0 | 0 | R.SYSCQVTHEGSTVEK.T | Carbamidomethyl: 4 | 194-208 | CID |
| 489 | 853.077 | 3 | 38.43 | 0.0328 | 21.15 | 58.35 | 3 | 0.0 | 1 | R.SYSCQVTHEGSTVEKTVAPTECS.- | Carbamidomethyl: 4, 22 | 194-216 | CID |

#### Protein 9: Immunoglobulin kappa variable 2D-28 OS=Homo sapiens GN=IGKV2D-28 PE=1 SV=2

| | | | |
|---|---|---|---|
| **Accession:** | KVD28_HUMAN | **Score:** | 148.8 |
| **Database:** | SwissProt | **Seq. Coverage [%]:** | 16.7 |
| **MW [kDa] / pI:** | 12.9 / 5.6 | **No. of Peptides:** | 2 |

```
       10         20         30         40         50         60         70         80         90        100
MRLPAQLLGL LMLWVSGSSG DIVMTQSPLS LPVTPGEPAS ISCRSSQSLL HSNGYNYLDW YLQKPGQSPQ LLIYLGSNRA SGVPDRFSGS GSGTDFTLKI
      110        120        130
SRVEAEDVGV YYCMQALQTP
```

MS/MS Peptide Matches

| Cmpd. | m/z meas. | z | Δ m/z [ppm] | Δ m/z [Da] | Rt [min] | Score | No. of Cmpds. | Site [%] | P | Sequence | Modification | Range | Type |
|---|---|---|---|---|---|---|---|---|---|---|---|---|---|
| 680 | 662.673 | 3 | 24.21 | 0.0160 | 24.87 | 51.66 | 3 | 0.0 | 1 | R.ASGVPDRFSGSGSGTDFTLK.I | | 80-99 | CID |
| 632 | 652.353 | 2 | 63.05 | 0.0411 | 23.95 | 97.11 | 3 | 0.0 | 0 | R.FSGSGSGTDFTLK.I | | 87-99 | CID |

#### Protein 10: Immunoglobulin heavy variable 3-7 OS=Homo sapiens GN=IGHV3-7 PE=1 SV=2

| | | | |
|---|---|---|---|
| **Accession:** | HV307_HUMAN | **Score:** | 135.9 |
| **Database:** | SwissProt | **Seq. Coverage [%]:** | 21.4 |
| **MW [kDa] / pI:** | 12.9 / 6.2 | **No. of Peptides:** | 3 |
| **Modification(s):** | Carbamidomethyl | | |

```
       10         20         30         40         50         60         70         80         90        100
MELGLSWVFL VAILEGVQCE VQLVESGGGL VQPGGSLRLS CAASGFTFSS YWMSWVRQAP GKGLEWVANI KQDGSEKYYV DSVKGRFTIS RDNAKNSLYL
      110        120
QMNSLRAEDT AVYYCAR
```

MS/MS Peptide Matches

| Cmpd. | m/z meas. | z | Δ m/z [ppm] | Δ m/z [Da] | Rt [min] | Score | No. of Cmpds. | Site [%] | P | Sequence | Modification | Range | Type |
|---|---|---|---|---|---|---|---|---|---|---|---|---|---|
| 393 | 437.273 | 2 | 118.32 | 0.0517 | 19.06 | 35.52 | 2 | 0.0 | 0 | K.YYVDSVK.G | | 78-84 | CID |
| 355 | 418.78 | 2 | 94.35 | 0.0395 | 18.24 | 21.02 | 2 | 0.0 | 1 | K.GRFTISR.D | | 85-91 | CID |
| 457 | 659.853 | 2 | 95.10 | 0.0627 | 20.39 | 79.38 | 4 | 0.0 | 0 | R.AEDTAVYYCAR.- | Carbamidomethyl: 9 | 107-117 | CID |

#### Protein 11: Immunoglobulin lambda variable 3-10 OS=Homo sapiens GN=IGLV3-10 PE=3 SV=2

**Accession:** LV310_HUMAN
**Database:** SwissProt
**MW [kDa] / pI:** 12.4 / 4.7
**Modification(s):** Carbamidomethyl

**Score:** 106.0
**Seq. Coverage [%]:** 15.7
**No. of Peptides:** 2

| 10 | 20 | 30 | 40 | 50 | 60 | 70 | 80 | 90 | 100 |
|---|---|---|---|---|---|---|---|---|---|
| MAWTPLLLPL | LTFCTVSEAS | YELTQPPSVS | VSPGQTARIT | CSGDALPKKY | AYWYQQKSGQ | APVLVIYEDS | KRPSGIPERF | SGSSSGTMAT | LTISGAQVED |
| 110 | 120 | | | | | | | | |
| EADYYCYSTD | SSGNH | | | | | | | | |

MS/MS Peptide Matches

| Cmpd. | m/z meas. | z | Δ m/z [ppm] | Δ m/z [Da] | Rt [min] | Score | No. of Cmpds. | Site [%] | P | Sequence | Modification | Range | Type |
|---|---|---|---|---|---|---|---|---|---|---|---|---|---|
| 394 | 531.308 | 2 | 74.49 | 0.0396 | 19.07 | 63.66 | 2 | 0.0 | 0 | R.ITCSGDALPK.K | Carbamidomethyl: 3 | 39-48 | CID |
| 220 | 456.327 | 2 | 154.50 | 0.0705 | 15.23 | 42.36 | 3 | 0.0 | 0 | K.RPSGIPER.F | | 72-79 | CID |

#### Protein 12: Alpha-1-antitrypsin OS=Homo sapiens GN=SERPINA1 PE=1 SV=3

**Accession:** A1AT_HUMAN
**Database:** SwissProt
**MW [kDa] / pI:** 46.7 / 5.4
**Modification(s):** Carbamidomethyl

**Score:** 98.0
**Seq. Coverage [%]:** 9.1
**No. of Peptides:** 3

| 10 | 20 | 30 | 40 | 50 | 60 | 70 | 80 | 90 | 100 |
|---|---|---|---|---|---|---|---|---|---|
| MPSSVSWGIL | LLAGLCCLVP | VSLAEDPQGD | AAQKTDTSHH | DQDHPTFNKI | TPNLAEFAFS | LYRQLAHQSN | STNIFFSPVS | IATAFAMLSL | GTKADTHDEI |
| 110 | 120 | 130 | 140 | 150 | 160 | 170 | 180 | 190 | 200 |
| LEGLNFNLTE | IPEAQIHEGF | QELLRTLNQP | DSQLQLTTGN | GLFLSEGLKL | VDKFLEDVKK | LYHSEAFTVN | FGDTEEAKKQ | INDYVEKGTQ | GKIVDLVKEL |
| 210 | 220 | 230 | 240 | 250 | 260 | 270 | 280 | 290 | 300 |
| DRDTVFALVN | YIFFKGKWER | PFEVKDTEEE | DFHVDQVTTV | KVPMMKRLGM | FNIQHCKKLS | SWVLLMKYLG | NATAIFFLPD | EGKLQHLENE | LTHDIITKFL |
| 310 | 320 | 330 | 340 | 350 | 360 | 370 | 380 | 390 | 400 |
| ENEDRRSASL | HLPKLSITGT | YDLKSVLGQL | GITKVFSNGA | DLSGVTEEAP | LKLSKAVHKA | VLTIDEKGTE | AAGAMFLEAI | PMSIPPEVKF | NKPFVFLMIE |
| 410 | 420 | | | | | | | | |
| QNTKSPLFMG | KVVNPTQK | | | | | | | | |

**MS/MS Peptide Matches**

| Cmpd. | m/z meas. | z | Δ m/z [ppm] | Δ m/z [Da] | Rt [min] | Score | No. of Cmpds. | Site [%] | P | Sequence | Modification | Range | Type |
|---|---|---|---|---|---|---|---|---|---|---|---|---|---|
| 1510 | 1045.575 | 2 | 22.49 | 0.0235 | 41.87 | 58.58 | 5 | 0.0 | 1 | K.ELDRDTVFALVNYIFFK.G | | 199-215 | CID |
| 492 | 425.916 | 3 | 38.36 | 0.0163 | 21.20 | 24.39 | 3 | 0.0 | 1 | K.GKWERPFEVK.D | | 216-225 | CID |
| 556 | 468.658 | 3 | 182.10 | 0.0853 | 22.47 | 15.04 | 1 | 0.0 | 1 | K.RLGMFNIQHCK.K | Carbamidomethyl: 10 | 247-257 | CID |

##### Protein 13: Serum albumin OS=Homo sapiens GN=ALB PE=1 SV=2

**Accession:** ALBU_HUMAN
**Database:** SwissProt
**MW [kDa] / pI:** 69.3 / 5.9
**Modification(s):** Carbamidomethyl

**Score:** 85.8
**Seq. Coverage [%]:** 4.1
**No. of Peptides:** 2

| | | | | | | | | | |
|---|---|---|---|---|---|---|---|---|---|
| 10 | 20 | 30 | 40 | 50 | 60 | 70 | 80 | 90 | 100 |
| MKWVTFISLL | FLFSSAYSRG | VFRRDAHKSE | VAHRFKDLGE | ENFKALVLIA | FAQYLQQCPF | EDHVKLVNEV | TEFAKTCVAD | ESAENCDKSL | HTLFGDKLCT |
| 110 | 120 | 130 | 140 | 150 | 160 | 170 | 180 | 190 | 200 |
| VATLRETYGE | MADCCAKQEP | ERNECFLQHK | DDNPNLPRLV | RPEVDVMCTA | FHDNEETFLK | KYLYEIARRH | PYFYAPELLF | FAKRYKAAFT | ECCQAADKAA |
| 210 | 220 | 230 | 240 | 250 | 260 | 270 | 280 | 290 | 300 |
| CLLPKLDELR | DEGKASSAKQ | RLKCASLQKF | GERAFKAWAV | ARLSQRFPKA | EFAEVSKLVT | DLTKVHTECC | HGDLLECADD | RADLAKYICE | NQDSISSKLK |
| 310 | 320 | 330 | 340 | 350 | 360 | 370 | 380 | 390 | 400 |
| ECCEKPLLEK | SHCIAEVEND | EMPADLPSLA | ADFVESKDVC | KNYAEAKDVF | LGMFLYEYAR | RHPDYSVVLL | LRLAKTYETT | LEKCCAADP | HECYAKVFDE |
| 410 | 420 | 430 | 440 | 450 | 460 | 470 | 480 | 490 | 500 |
| FKPLVEEPQN | LIKQNCELFE | QLGEYKFQNA | LLVRYTKKVP | QVSTPTLVEV | SRNLGKVGSK | CCKHPEAKRM | PCAEDYLSVV | LNQLCVLHEK | TPVSDRVTKC |
| 510 | 520 | 530 | 540 | 550 | 560 | 570 | 580 | 590 | 600 |
| CTESLVNRRP | CFSALEVDET | YVPKEFNAET | FTFHADICTL | SEKERQIKKQ | TALVELVKHK | PKATKEQLKA | VMDDFAAFVE | KCCKADDKET | CFAEEGKKLV |
| 610 | | | | | | | | | |
| AASQAALGL | | | | | | | | | |

**MS/MS Peptide Matches**

| Cmpd. | m/z meas. | z | Δ m/z [ppm] | Δ m/z [Da] | Rt [min] | Score | No. of Cmpds. | Site [%] | P | Sequence | Modification | Range | Type |
|---|---|---|---|---|---|---|---|---|---|---|---|---|---|
| 365 | 516.349 | 3 | 152.17 | 0.0786 | 18.46 | 23.34 | 1 | 0.0 | 1 | K.LKECCEKPLLEK.S | Carbamidomethyl: 4, 5 | 299-310 | CID |
| 1392 | 812.452 | 2 | 67.18 | 0.0546 | 39.23 | 62.44 | 1 | 0.0 | 0 | K.DVFLGMFLYEYAR.R | | 348-360 | CID |

#### Protein 14: Immunoglobulin kappa variable 3-20 OS=Homo sapiens GN=IGKV3-20 PE=1 SV=2

| | | | |
|---|---|---|---|
| **Accession:** | KV320_HUMAN | **Score:** | 69.5 |
| **Database:** | SwissProt | **Seq. Coverage [%]:** | 7.8 |
| **MW [kDa] / pI:** | 12.5 / 4.9 | **No. of Peptides:** | 1 |

| 10 | 20 | 30 | 40 | 50 | 60 | 70 | 80 | 90 | 100 |
|---|---|---|---|---|---|---|---|---|---|
| METPAQLLFL | LLLWLPDTTG | EIVLTQSPGT | LSLSPGERAT | LSCRASQSVS | SSYLAWYQQK | PGQAPRLLIY | GASSRATGIP | DRFSGSGSGT | DFTLTISRLE |
| 110 | 120 | | | | | | | | |
| PEDFAVYYCQ | QYGSSP | | | | | | | | |

MS/MS Peptide Matches

| Cmpd. | m/z meas. | z | Δ m/z [ppm] | Δ m/z [Da] | Rt [min] | Score | No. of Cmpds. | Site [%] | P | Sequence | Modification | Range | Type |
|---|---|---|---|---|---|---|---|---|---|---|---|---|---|
| 638 | 490.321 | 2 | 79.16 | 0.0388 | 24.05 | 69.47 | 3 | 0.0 | 0 | R.LLIYGASSR.A | | 67-75 | CID |

#### Protein 15: Immunoglobulin kappa variable 3-15 OS=Homo sapiens GN=IGKV3-15 PE=1 SV=2

**Accession:** KV315_HUMAN
**Database:** SwissProt
**MW [kDa] / pI:** 12.5 / 5.1

**Score:** 67.5
**Seq. Coverage [%]:** 7.8
**No. of Peptides:** 1

| 10 | 20 | 30 | 40 | 50 | 60 | 70 | 80 | 90 | 100 |
|---|---|---|---|---|---|---|---|---|---|
| MEAPAQLLFL | LLLWLPDTTG | EIVMTQSPAT | LSVSPGERAT | LSCRASQSVS | SNLAWYQQKP | GQAPRLLIYG | ASTRATGIPA | RFSGSGSGTE | FTLTISSLQS |
| 110 | 120 | | | | | | | | |
| EDFAVYYCQQ | YNNWP | | | | | | | | |

MS/MS Peptide Matches

| Cmpd. | m/z meas. | z | Δ m/z [ppm] | Δ m/z [Da] | Rt [min] | Score | No. of Cmpds. | Site [%] | P | Sequence | Modification | Range | Type |
|---|---|---|---|---|---|---|---|---|---|---|---|---|---|
| 655 | 497.335 | 2 | 90.46 | 0.0450 | 24.39 | 67.52 | 5 | 0.0 | 0 | R.LLIYGASTR.A | | 66-74 | CID |

#### Protein 16: Immunoglobulin kappa variable 3-11 OS=Homo sapiens GN=IGKV3-11 PE=1 SV=1

**Accession:** KV311_HUMAN
**Database:** SwissProt
**MW [kDa] / pI:** 12.6 / 4.9

**Score:** 63.4
**Seq. Coverage [%]:** 7.8
**No. of Peptides:** 1

| 10 | 20 | 30 | 40 | 50 | 60 | 70 | 80 | 90 | 100 |
|---|---|---|---|---|---|---|---|---|---|
| MEAPAQLLFL | LLLWLPDTTG | EIVLTQSPAT | LSLSPGERAT | LSCRASQSVS | SYLAWYQQKP | GQAPRLLIYD | ASNRATGIPA | RFSGSGSGTD | FTLTISSLEP |
| 110 | 120 | | | | | | | | |
| EDFAVYYCQQ | RSNWP | | | | | | | | |

MS/MS Peptide Matches

| Cmpd. | m/z meas. | z | Δ m/z [ppm] | Δ m/z [Da] | Rt [min] | Score | No. of Cmpds. | Site [%] | P | Sequence | Modification | Range | Type |
|---|---|---|---|---|---|---|---|---|---|---|---|---|---|
| 652 | 532.831 | 2 | 76.24 | 0.0406 | 24.31 | 63.39 | 4 | 0.0 | 0 | R.LLIYDASNR.A | | 66-74 | CID |

#### Protein 17: Trypsin-1 OS=Homo sapiens GN=PRSS1 PE=1 SV=1

| | | | |
|---|---|---|---|
| **Accession:** | TRY1_HUMAN | **Score:** | 42.9 |
| **Database:** | SwissProt | **Seq. Coverage [%]:** | 3.2 |
| **MW [kDa] / pI:** | 26.5 / 6.1 | **No. of Peptides:** | 1 |

| 10 | 20 | 30 | 40 | 50 | 60 | 70 | 80 | 90 | 100 |
|---|---|---|---|---|---|---|---|---|---|
| MNPLLILTFV | AAALAAPFDD | DDKIVGGYNC | EENSVPYQVS | LNSGYHFCGG | SLINEQWVVS | AGHCYKSRIQ | VRLGEHNIEV | LEGNEQFINA | AKIIRHPQYD |
| 110 | 120 | 130 | 140 | 150 | 160 | 170 | 180 | 190 | 200 |
| RKTLNNDIML | IKLSSRAVIN | ARVSTISLPT | APPATGTKCL | ISGWGNTASS | GADYPDELQC | LDAPVLSQAK | CEASYPGKIT | SNMFCVGFLE | GGKDSCQGDS |
| 210 | 220 | 230 | 240 | 250 | | | | | |
| GGPVVCNGQL | QGVVSWGDGC | AQKNKPGVYT | KVYNYVKWIK | NTIAANS | | | | | |

MS/MS Peptide Matches

| Cmpd. | m/z meas. | z | Δ m/z [ppm] | Δ m/z [Da] | Rt [min] | Score | No. of Cmpds. | Site [%] | P | Sequence | Modification | Range | Type |
|---|---|---|---|---|---|---|---|---|---|---|---|---|---|
| 143 | 453.828 | 2 | 159.10 | 0.0722 | 13.32 | 42.89 | 4 | 0.0 | 0 | K.NKPGVYTK.V | | 224-231 | CID |

##### Protein 18: Immunoglobulin heavy variable 5-51 OS=Homo sapiens GN=IGHV5-51 PE=3 SV=1

| | | | |
|---|---|---|---|
| **Accession:** | HV551_HUMAN | **Score:** | 36.2 |
| **Database:** | SwissProt | **Seq. Coverage [%]:** | 9.4 |
| **MW [kDa] / pI:** | 12.7 / 8.4 | **No. of Peptides:** | 1 |
| **Modification(s):** | Carbamidomethyl | | |

```
        10         20         30         40         50         60         70         80         90        100
MGSTAILALL LAVLQGVCAE VQLVQSGAEV KKPGESLKIS CKGSGYSFTS YWIGWVRQMP GKGLEWMGII YPGDSDTRYS PSFQGQVTIS ADKSISTAYL
       110        120
QWSSLKASDT AMYYCAR
```

**MS/MS Peptide Matches**

| Cmpd. | m/z meas. | z | Δ m/z [ppm] | Δ m/z [Da] | Rt [min] | Score | No. of Cmpds. | Site [%] | P | Sequence | Modification | Range | Type |
|---|---|---|---|---|---|---|---|---|---|---|---|---|---|
| 469 | 654.695 | 2 | -116.08 | -0.0760 | 20.65 | 36.16 | 3 | 0.0 | 0 | K.ASDTAMYYCAR.- | Carbamidomethyl: 9 | 107-117 | CID |

#### Project Info

| | | | |
|---|---|---|---|
| **Name:** | Alagesan | **Date:** | Mar 5, 2019 |
| **Note:** | Nanobooster | | |

#### Sample Info & Protocols

**Name:** Nanobooster - IgG
**Date:** Mar 5, 2019

#### Search Result Info

| | | | |
|---|---|---|---|
| **Search Result:** | IonTrap_Human_Swiss_CID_2019-03-05 17:55:08 | | |
| **Location:** | /Alagesan/Nanobooster - IgG/Ka_IgG Trypsin NoEnrich NB Ace_BA5_01_8756.mgf | | |
| **Search Method:** | IonTrap_Human_Swiss_CID | **Version:** | 1.2 |
| **Search Engine(s):** | Mascot, 2.6.0 | **Ident. Compound(s):** | 0/2070 |
| **Database(s):** | SwissProt, SwissProt_2017_04.fasta | | |
| **Note:** | | | |

#### Results

##### Protein 1: Immunoglobulin kappa constant OS=Homo sapiens GN=IGKC PE=1 SV=2

**Accession:** IGKC_HUMAN
**Database:** SwissProt
**MW [kDa] / pI:** 11.8 / 6.1
**Modification(s):** Carbamidomethyl

**Score:** 1051.3
**Seq. Coverage [%]:** 97.2
**No. of Peptides:** 11

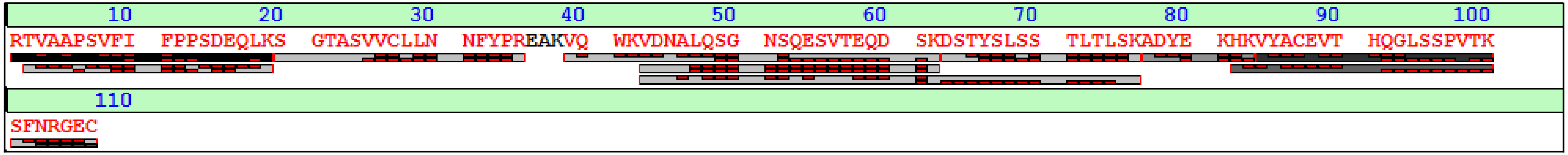


MS/MS Peptide Matches

| Cmpd. | m/z meas. | z | Δ m/z [ppm] | Δ m/z [Da] | Rt [min] | Score | No. of Cmpds. | Site [%] | P | Sequence | Modification | Range | Type |
|---|---|---|---|---|---|---|---|---|---|---|---|---|---|
| 911 | 701.434 | 3 | 75.75 | 0.0531 | 31.45 | 59.48 | 13 | 0.0 | 1 | -.RTVAAPSVFIFPPSDEQLK.S | | 1-19 | CID |
| 1003 | 973.986 | 2 | 481.64 | 0.4689 | 33.47 | 98.64 | 44 | 0.0 | 0 | R.TVAAPSVFIFPPSDEQLK.S | | 2-19 | CID |
| 1088 | 899.554 | 2 | 114.23 | 0.1027 | 35.37 | 76.76 | 32 | 0.0 | 0 | K.SGTASVVCLLNNFYPR.E | Carbamidomethyl: 8 | 20-35 | CID |
| 547 | 893.069 | 3 | -28.94 | -0.0258 | 23.71 | 129.07 | 3 | 0.0 | 1 | K.VQWKVDNALQSGNSQESVTEQDSK.D | | 39-62 | CID |
| 332 | 1068.665 | 2 | 165.65 | 0.1770 | 19.23 | 169.91 | 75 | 0.0 | 0 | K.VDNALQSGNSQESVTEQDSK.D | | 43-62 | CID |
| 688 | 1207.567 | 3 | 269.79 | 0.3257 | 26.98 | 180.82 | 28 | 0.0 | 1 | K.VDNALQSGNSQESVTEQDSKDSTYSLSSTLTLSK.A | | 43-76 | CID |
| 723 | 751.954 | 2 | 94.62 | 0.0711 | 27.69 | 89.2 | 8 | 0.0 | 0 | K.DSTYSLSSTLTLSK.A | | 63-76 | CID |
| 7 | 445.783 | 2 | 136.93 | 0.0610 | 11.37 | 24.83 | 2 | 0.0 | 1 | K.ADYEKHK.V | | 77-83 | CID |
| 337 | 536.112 | 4 | 161.08 | 0.0863 | 19.36 | 64.25 | 10 | 0.0 | 1 | K.HKVYACEVTHQGLSSPVTK.S | Carbamidomethyl: 6 | 82-100 | CID |
| 450 | 626.25 | 3 | 430.54 | 0.2695 | 21.62 | 114.13 | 9 | 0.0 | 0 | K.VYACEVTHQGLSSPVTK.S | Carbamidomethyl: 4 | 84-100 | CID |
| 94 | 435.186 | 2 | 8.84 | 0.0038 | 13.89 | 44.18 | 8 | 0.0 | 1 | K.SFNRGEC.- | Carbamidomethyl: 7 | 101-107 | CID |

#### Protein 2: Immunoglobulin heavy constant gamma 1 OS=Homo sapiens GN=IGHG1 PE=1 SV=1

| | | | |
|---|---|---|---|
| **Accession:** | IGHG1_HUMAN | **Score:** | 1041.1 |
| **Database:** | SwissProt | **Seq. Coverage [%]:** | 54.8 |
| **MW [kDa] / pI:** | 36.1 / 8.5 | **No. of Peptides:** | 15 |
| **Modification(s):** | Carbamidomethyl, Oxidation | | |

| 10 | 20 | 30 | 40 | 50 | 60 | 70 | 80 | 90 | 100 |
|---|---|---|---|---|---|---|---|---|---|
| ASTKGPSVFP | LAPSSKSTSG | GTAALGCLVK | DYFPEPVTVS | WNSGALTSGV | HTFPAVLQSS | GLYSLSSVVT | VPSSSLGTQT | YICNVNHKPS | NTKVDKKVEP |
| 110 | 120 | 130 | 140 | 150 | 160 | 170 | 180 | 190 | 200 |
| KSCDKTHTCP | PCPAPELLGG | PSVFLFPPKP | KDTLMISRTP | EVTCVVVDVS | HEDPEVKFNW | YVDGVEVHNA | KTKPREEQYN | STYRVVSVLT | VLHQDWLNGK |
| 210 | 220 | 230 | 240 | 250 | 260 | 270 | 280 | 290 | 300 |
| EYKCKVSNKA | LPAPIEKTIS | KAKGQPREPQ | VYTLPPSRDE | LTKNQVSLTC | LVKGFYPSDI | AVEWESNGQP | ENNYKTTPPV | LDSDGSFFLY | SKLTVDKSRW |
| 310 | 320 | 330 | 340 | | | | | | |
| QQGNVFSCSV | MHEALHNHYT | QKSLSLPGK | | | | | | | |

##### MS/MS Peptide Matches

| Cmpd. | m/z meas. | z | Δ m/z [ppm] | Δ m/z [Da] | Rt [min] | Score | No. of Cmpds. | Site [%] | P | Sequence | Modification | Range | Type |
|---|---|---|---|---|---|---|---|---|---|---|---|---|---|
| 715 | 593.857 | 2 | 50.58 | 0.0300 | 27.55 | 57.92 | 9 | 0.0 | 0 | K.GPSVFPLAPSSK.S | | 5-16 | CID |
| 630 | 661.384 | 2 | 62.52 | 0.0413 | 25.52 | 113.07 | 16 | 0.0 | 0 | K.STSGGTAALGCLVK.D | Carbamidomethyl: 11 | 17-30 | CID |
| 984 | 712.231 | 4 | 507.34 | 0.3612 | 33.01 | 58.03 | 12 | 0.0 | 0 | K.THTCPPCPAPELLGGPSVFLFPPKPK.D | Carbamidomethyl: 4, 7 | 106-131 | CID |
| 457 | 418.231 | 2 | 24.52 | 0.0103 | 21.76 | 47.88 | 12 | 0.0 | 0 | K.DTLMISR.T | | 132-138 | CID |
| 360 | 426.264 | 2 | 107.45 | 0.0458 | 19.84 | 44.05 | 4 | 0.0 | 0 | K.DTLMISR.T | Oxidation: 4 | 132-138 | CID |
| 700 | 713.684 | 3 | 4.68 | 0.0033 | 27.25 | 93.61 | 3 | 0.0 | 0 | R.TPEVTCVVVDVSHEDPEVK.F | Carbamidomethyl: 6 | 139-157 | CID |
| 1296 | 950.409 | 4 | 211.17 | 0.2007 | 39.64 | 49.37 | 9 | 0.0 | 1 | R.TPEVTCVVVDVSHEDPEVKFNWYVDGVEVHNAK.T | Carbamidomethyl: 6 | 139-171 | CID |
| 712 | 560.002 | 3 | 112.79 | 0.0632 | 27.48 | 72.62 | 8 | 0.0 | 0 | K.FNWYVDGVEVHNAK.T | | 158-171 | CID |

| Cmpd. | m/z meas. | z | Δ m/z [ppm] | Δ m/z [Da] | Rt [min] | Score | No. of Cmpds. | Site [%] | P | Sequence | Modification | Range | Type |
|---|---|---|---|---|---|---|---|---|---|---|---|---|---|
| 1049 | 603.42 | 3 | 132.02 | 0.0797 | 34.48 | 56.7 | 8 | 0.0 | 0 | R.VVSVLTVLHQDWLNGK.E | | 185-200 | CID |
| 445 | 419.797 | 2 | 99.40 | 0.0417 | 21.48 | 27.25 | 5 | 0.0 | 0 | K.ALPAPIEK.T | | 210-217 | CID |
| 538 | 643.836 | 2 | -7.15 | -0.0046 | 23.52 | 63.66 | 11 | 0.0 | 0 | R.EPQVYTLPPSR.D | | 228-238 | CID |
| 576 | 625.073 | 3 | 124.95 | 0.0781 | 24.31 | 30.89 | 9 | 0.0 | 1 | R.EPQVYTLPPSRDELTK.N | | 228-243 | CID |
| 680 | 581.357 | 2 | 66.32 | 0.0386 | 26.76 | 82.12 | 12 | 0.0 | 0 | K.NQVSLTCLVK.G | Carbamidomethyl: 7 | 244-253 | CID |
| 880 | 1273.041 | 2 | 370.65 | 0.4717 | 30.87 | 132.13 | 40 | 0.0 | 0 | K.GFYPSDIAVEWESNGQPENNYK.T | | 254-275 | CID |
| 948 | 937.873 | 2 | 435.70 | 0.4085 | 32.23 | 111.78 | 16 | 0.0 | 0 | K.TTPPVLDSDGSFFLYSK.L | | 276-292 | CID |

#### Protein 3: Immunoglobulin kappa light chain OS=Homo sapiens PE=1 SV=1

**Accession:** IGK_HUMAN
**Database:** SwissProt
**MW [kDa] / pI:** 23.4 / 6.9
**Modification(s):** Carbamidomethyl

**Score:** 1013.7
**Seq. Coverage [%]:** 48.1
**No. of Peptides:** 10

```
10         20         30         40         50         60         70         80         90         100
DIQMTQSPST LSASVGDRVT ITCRASQSIN TWLAWYQQKP GKAPKLLMYK ASSLESGVPS RFIGSGSGTE FTLTISSLQP DDFATYYCQQ YNSDSKMFGQ
110        120        130        140        150        160        170        180        190        200
GTKVEVKGTV AAPSVFIFPP SDEQLKSGTA SVVCLLNNFY PREAKVQWKV DNALQSGNSQ ESVTEQDSKD STYSLSSTLT LSKADYEKHK VYACEVTHQG
210        220
LSSPVTKSFN RGEC
```

##### MS/MS Peptide Matches

| Cmpd. | m/z meas. | z | Δ m/z [ppm] | Δ m/z [Da] | Rt [min] | Score | No. of Cmpds. | Site [%] | P | Sequence | Modification | Range | Type |
|---|---|---|---|---|---|---|---|---|---|---|---|---|---|
| 607 | 946.914 | 2 | -42.83 | -0.0406 | 24.96 | 120.56 | 1 | 0.0 | 0 | -.DIQMTQSPSTLSASVGDR.V | | 1-18 | CID |
| 1088 | 899.554 | 2 | 114.23 | 0.1027 | 35.37 | 76.76 | 32 | 0.0 | 0 | K.SGTASVVCLLNNFYPR.E | Carbamidomethyl: 8 | 127-142 | CID |

| Cmpd. | m/z meas. | z | Δ m/z [ppm] | Δ m/z [Da] | Rt [min] | Score | No. of Cmpds. | Site [%] | P | Sequence | Modification | Range | Type |
|---|---|---|---|---|---|---|---|---|---|---|---|---|---|
| 547 | 893.069 | 3 | -28.94 | -0.0258 | 23.71 | 129.07 | 3 | 0.0 | 1 | K.VQWKVDNALQSGNSQESVTEQDSK.D | | 146-169 | CID |
| 332 | 1068.665 | 2 | 165.65 | 0.1770 | 19.23 | 169.91 | 75 | 0.0 | 0 | K.VDNALQSGNSQESVTEQDSK.D | | 150-169 | CID |
| 688 | 1207.567 | 3 | 269.79 | 0.3257 | 26.98 | 180.82 | 28 | 0.0 | 1 | K.VDNALQSGNSQESVTEQDSKDSTYSLSSTLTLSK.A | | 150-183 | CID |
| 723 | 751.954 | 2 | 94.62 | 0.0711 | 27.69 | 89.2 | 8 | 0.0 | 0 | K.DSTYSLSSTLTLSK.A | | 170-183 | CID |
| 7 | 445.783 | 2 | 136.93 | 0.0610 | 11.37 | 24.83 | 2 | 0.0 | 1 | K.ADYEKHK.V | | 184-190 | CID |
| 337 | 536.112 | 4 | 161.08 | 0.0863 | 19.36 | 64.25 | 10 | 0.0 | 1 | K.HKVYACEVTHQGLSSPVTK.S | Carbamidomethyl: 6 | 189-207 | CID |
| 450 | 626.25 | 3 | 430.54 | 0.2695 | 21.62 | 114.13 | 9 | 0.0 | 0 | K.VYACEVTHQGLSSPVTK.S | Carbamidomethyl: 4 | 191-207 | CID |
| 94 | 435.186 | 2 | 8.84 | 0.0038 | 13.89 | 44.18 | 8 | 0.0 | 1 | K.SFNRGEC.- | Carbamidomethyl: 7 | 208-214 | CID |

#### Protein 4: Immunoglobulin heavy constant gamma 2 OS=Homo sapiens GN=IGHG2 PE=1 SV=2

**Accession:** IGHG2_HUMAN
**Database:** SwissProt
**MW [kDa] / pI:** 35.9 / 7.7
**Modification(s):** Carbamidomethyl, Oxidation

**Score:** 823.2
**Seq. Coverage [%]:** 50.0
**No. of Peptides:** 15

| 10 | 20 | 30 | 40 | 50 | 60 | 70 | 80 | 90 | 100 |
|---|---|---|---|---|---|---|---|---|---|
| ASTKGPSVFP | LAPCSRSTSE | STAALGCLVK | DYFPEPVTVS | WNSGALTSGV | HTFPAVLQSS | GLYSLSSVVT | VPSSNFGTQT | YTCNVDHKPS | NTKVDKTVER |
| **110** | **120** | **130** | **140** | **150** | **160** | **170** | **180** | **190** | **200** |
| KCCVECPPCP | APPVAGPSVF | LFPPKPKDTL | MISRTPEVTC | VVVDVSHEDP | EVQFNWYVDG | VEVHNAKTKP | REEQFNSTFR | VVSVLTVVHQ | DWLNGKEYKC |
| **210** | **220** | **230** | **240** | **250** | **260** | **270** | **280** | **290** | **300** |
| KVSNKGLPAP | IEKTISKTKG | QPREPQVYTL | PPSREEMTKN | QVSLTCLVKG | FYPSDISVEW | ESNGQPENNY | KTTPPMLDSD | GSFFLYSKLT | VDKSRWQQGN |
| **310** | **320** | **330** | | | | | | | |
| VFSCSVMHEA | LHNHYTQKSL | SLSPGK | | | | | | | |

MS/MS Peptide Matches

| Cmpd. | m/z meas. | z | Δ m/z [ppm] | Δ m/z [Da] | Rt [min] | Score | No. of Cmpds. | Site [%] | P | Sequence | Modification | Range | Type |
|---|---|---|---|---|---|---|---|---|---|---|---|---|---|
| 746 | 644.392 | 2 | 97.24 | 0.0627 | 28.14 | 60.96 | 13 | 0.0 | 0 | K.GPSVFPLAPCSR.S | Carbamidomethyl: 10 | 5-16 | CID |
| 642 | 712.408 | 2 | 69.49 | 0.0495 | 25.83 | 89.0 | 16 | 0.0 | 0 | R.STSESTAALGCLVK.D | Carbamidomethyl: 11 | 17-30 | CID |
| 875 | 760.229 | 4 | 459.79 | 0.3494 | 30.78 | 42.15 | 4 | 0.0 | 1 | R.KCCVECPPCPAPPVAGPSVFLFPPKPK.D | Carbamidomethyl: 2, 3, 6, 9 | 101-127 | CID |
| 951 | 970.486 | 3 | 357.94 | 0.3473 | 32.31 | 81.32 | 9 | 0.0 | 0 | K.CCVECPPCPAPPVAGPSVFLFPPKPK.D | Carbamidomethyl: 1, 2, 5, 8 | 102-127 | CID |
| 457 | 418.231 | 2 | 24.52 | 0.0103 | 21.76 | 47.88 | 12 | 0.0 | 0 | K.DTLMISR.T | | 128-134 | CID |
| 360 | 426.264 | 2 | 107.45 | 0.0458 | 19.84 | 44.05 | 4 | 0.0 | 0 | K.DTLMISR.T | Oxidation: 4 | 128-134 | CID |
| 1296 | 950.409 | 4 | 220.74 | 0.2097 | 39.64 | 49.37 | 9 | 0.0 | 0 | R.TPEVTCVVVDVSHEDPEVQFNWYVDGVEVHNAK.T | Carbamidomethyl: 6 | 135-167 | CID |
| 943 | 599.078 | 3 | 684.08 | 0.4095 | 32.10 | 30.59 | 3 | 0.0 | 0 | R.VVSVLTVVHQDWLNGK.E | | 181-196 | CID |
| 878 | 554.378 | 4 | 134.60 | 0.0746 | 30.83 | 17.12 | 1 | 0.0 | 1 | R.VVSVLTVVHQDWLNGKEYK.C | | 181-199 | CID |
| 455 | 412.737 | 2 | -25.33 | -0.0105 | 21.72 | 25.92 | 1 | 0.0 | 0 | K.GLPAPIEK.T | | 206-213 | CID |
| 538 | 643.836 | 2 | -7.15 | -0.0046 | 23.52 | 63.66 | 11 | 0.0 | 0 | R.EPQVYTLPPSR.E | | 224-234 | CID |
| 508 | 635.664 | 3 | 18.46 | 0.0117 | 22.79 | 42.89 | 27 | 0.0 | 1 | R.EPQVYTLPPSREEMTK.N | | 224-239 | CID |
| 680 | 581.357 | 2 | 66.32 | 0.0386 | 26.76 | 82.12 | 12 | 0.0 | 0 | K.NQVSLTCLVK.G | Carbamidomethyl: 7 | 240-249 | CID |
| 967 | 953.658 | 2 | 217.54 | 0.2074 | 32.65 | 90.37 | 10 | 0.0 | 0 | K.TTPPMLDSDGSFFLYSK.L | | 272-288 | CID |
| 876 | 961.457 | 2 | 9.32 | 0.0090 | 30.78 | 55.79 | 3 | 0.0 | 0 | K.TTPPMLDSDGSFFLYSK.L | Oxidation: 5 | 272-288 | CID |

#### Protein 5: Immunoglobulin heavy constant gamma 3 OS=Homo sapiens GN=IGHG3 PE=1 SV=2

**Accession:** IGHG3_HUMAN
**Database:** SwissProt
**MW [kDa] / pI:** 41.3 / 8.2
**Modification(s):** Carbamidomethyl, Oxidation

**Score:** 767.3
**Seq. Coverage [%]:** 36.3
**No. of Peptides:** 13

| 10 | 20 | 30 | 40 | 50 | 60 | 70 | 80 | 90 | 100 |
|---|---|---|---|---|---|---|---|---|---|
| ASTKGPSVFP | LAPCSRSTSG | GTAALGCLVK | DYFPEPVTVS | WNSGALTSGV | HTFPAVLQSS | GLYSLSSVVT | VPSSSLGTQT | YTCNVNHKPS | NTKVDKRVEL |
| 110 | 120 | 130 | 140 | 150 | 160 | 170 | 180 | 190 | 200 |
| KTPLGDTTHT | CPRCPEPKSC | DTPPPCPRCP | EPKSCDTPPP | CPRCPEPKSC | DTPPPCPRCP | APELLGGPSV | FLFPPKPKDT | LMISRTPEVT | CVVVDVSHED |
| 210 | 220 | 230 | 240 | 250 | 260 | 270 | 280 | 290 | 300 |
| PEVQFKWYVD | GVEVHNAKTK | PREEQYNSTF | RVVSVLTVLH | QDWLNGKEYK | CKVSNKALPA | PIEKTISKTK | GQPREPQVYT | LPPSREEMTK | NQVSLTCLVK |
| 310 | 320 | 330 | 340 | 350 | 360 | 370 | 380 | | |
| GFYPSDIAVE | WESSGQPENN | YNTTPPMLDS | DGSFFLYSKL | TVDKSRWQQG | NIFSCSVMHE | ALHNRFTQKS | LSLSPGK | | |

MS/MS Peptide Matches

| Cmpd. | m/z meas. | z | Δ m/z [ppm] | Δ m/z [Da] | Rt [min] | Score | No. of Cmpds. | Site [%] | P | Sequence | Modification | Range | Type |
|---|---|---|---|---|---|---|---|---|---|---|---|---|---|
| 746 | 644.392 | 2 | 97.24 | 0.0627 | 28.14 | 60.96 | 13 | 0.0 | 0 | K.GPSVFPLAPCSR.S | Carbamidomethyl: 10 | 5-16 | CID |
| 630 | 661.384 | 2 | 62.52 | 0.0413 | 25.52 | 113.07 | 16 | 0.0 | 0 | R.STSGGTAALGCLVK.D | Carbamidomethyl: 11 | 17-30 | CID |
| 227 | 452.544 | 3 | -14.57 | -0.0066 | 16.91 | 51.36 | 4 | 0.0 | 0 | K.TPLGDTTHTCPR.C | Carbamidomethyl: 10 | 102-113 | CID |
| 199 | 593.76 | 2 | 12.44 | 0.0074 | 16.25 | 63.01 | 12 | 0.0 | 0 | K.SCDTPPPCPR.C | Carbamidomethyl: 2, 8 | 119-128 | CID |
| 1045 | 717.796 | 3 | 95.84 | 0.0688 | 34.38 | 76.03 | 3 | 0.0 | 0 | R.CPAPELLGGPSVFLFPPKPK.D | Carbamidomethyl: 1 | 159-178 | CID |
| 457 | 418.231 | 2 | 24.52 | 0.0103 | 21.76 | 47.88 | 12 | 0.0 | 0 | K.DTLMISR.T | | 179-185 | CID |
| 360 | 426.264 | 2 | 107.45 | 0.0458 | 19.84 | 44.05 | 4 | 0.0 | 0 | K.DTLMISR.T | Oxidation: 4 | 179-185 | CID |
| 550 | 472.97 | 3 | 144.36 | 0.0683 | 23.78 | 38.34 | 3 | 0.0 | 0 | K.WYVDGVEVHNAK.T | | 207-218 | CID |
| 1049 | 603.42 | 3 | 132.02 | 0.0797 | 34.48 | 56.7 | 8 | 0.0 | 0 | R.VVSVLTVLHQDWLNGK.E | | 232-247 | CID |
| 445 | 419.797 | 2 | 99.40 | 0.0417 | 21.48 | 27.25 | 5 | 0.0 | 0 | K.ALPAPIEK.T | | 257-264 | CID |
| 538 | 643.836 | 2 | -7.15 | -0.0046 | 23.52 | 63.66 | 11 | 0.0 | 0 | R.EPQVYTLPPSR.E | | 275-285 | CID |
| 508 | 635.664 | 3 | 18.46 | 0.0117 | 22.79 | 42.89 | 27 | 0.0 | 1 | R.EPQVYTLPPSREEMTK.N | | 275-290 | CID |
| 680 | 581.357 | 2 | 66.32 | 0.0386 | 26.76 | 82.12 | 12 | 0.0 | 0 | K.NQVSLTCLVK.G | Carbamidomethyl: 7 | 291-300 | CID |

#### Protein 6: Immunoglobulin heavy constant gamma 4 OS=Homo sapiens GN=IGHG4 PE=1 SV=1

| | | | |
|---|---|---|---|
| **Accession:** | IGHG4_HUMAN | **Score:** | 725.2 |
| **Database:** | SwissProt | **Seq. Coverage [%]:** | 49.8 |
| **MW [kDa] / pI:** | 35.9 / 7.2 | **No. of Peptides:** | 12 |
| **Modification(s):** | Carbamidomethyl, Oxidation | | |

| 10 | 20 | 30 | 40 | 50 | 60 | 70 | 80 | 90 | 100 |
|---|---|---|---|---|---|---|---|---|---|
| ASTKGPSVFP | LAPCSRSTSE | STAALGCLVK | DYFPEPVTVS | WNSGALTSGV | HTFPAVLQSS | GLYSLSSVVT | VPSSSLGTKT | YTCNVDHKPS | NTKVDKRVES |
| **110** | **120** | **130** | **140** | **150** | **160** | **170** | **180** | **190** | **200** |
| KYGPPCPSCP | APEFLGGPSV | FLFPPKPKDT | LMISRTPEVT | CVVVDVSQED | PEVQFNWYVD | GVEVHNAKTK | PREEQFNSTY | RVVSVLTVLH | QDWLNGKEYK |
| **210** | **220** | **230** | **240** | **250** | **260** | **270** | **280** | **290** | **300** |
| CKVSNKGLPS | SIEKTISKAK | GQPREPQVYT | LPPSQEEMTK | NQVSLTCLVK | GFYPSDIAVE | WESNGQPENN | YKTTPPVLDS | DGSFFLYSRL | TVDKSRWQEG |
| **310** | **320** | **330** | | | | | | | |
| NVFSCSVMHE | ALHNHYTQKS | LSLSLGK | | | | | | | |

**MS/MS Peptide Matches**

| Cmpd. | m/z meas. | z | Δ m/z [ppm] | Δ m/z [Da] | Rt [min] | Score | No. of Cmpds. | Site [%] | P | Sequence | Modification | Range | Type |
|---|---|---|---|---|---|---|---|---|---|---|---|---|---|
| 746 | 644.392 | 2 | 97.24 | 0.0627 | 28.14 | 60.96 | 13 | 0.0 | 0 | K.GPSVFPLAPCSR.S | Carbamidomethyl: 10 | 5-16 | CID |
| 642 | 712.408 | 2 | 69.49 | 0.0495 | 25.83 | 89.0 | 16 | 0.0 | 0 | R.STSESTAALGCLVK.D | Carbamidomethyl: 11 | 17-30 | CID |
| 96 | 416.993 | 4 | 108.21 | 0.0451 | 13.95 | 25.92 | 3 | 0.0 | 0 | K.TYTCNVDHKPSNTK.V | Carbamidomethyl: 4 | 80-93 | CID |
| 1074 | 982.209 | 3 | 392.21 | 0.3851 | 35.03 | 63.32 | 5 | 0.0 | 0 | K.YGPPCPSCPAPEFLGGPSVFLFPPKPK.D | Carbamidomethyl: 5, 8 | 102-128 | CID |
| 457 | 418.231 | 2 | 24.52 | 0.0103 | 21.76 | 47.88 | 12 | 0.0 | 0 | K.DTLMISR.T | | 129-135 | CID |
| 360 | 426.264 | 2 | 107.45 | 0.0458 | 19.84 | 44.05 | 4 | 0.0 | 0 | K.DTLMISR.T | Oxidation: 4 | 129-135 | CID |
| 1049 | 603.42 | 3 | 132.02 | 0.0797 | 34.48 | 56.7 | 8 | 0.0 | 0 | R.VVSVLTVLHQDWLNGK.E | | 182-197 | CID |
| 376 | 415.816 | 2 | 195.94 | 0.0815 | 20.10 | 17.46 | 2 | 0.0 | 0 | K.GLPSSIEK.T | | 207-214 | CID |
| 596 | 938.947 | 2 | -6.91 | -0.0065 | 24.72 | 28.16 | 1 | 0.0 | 0 | R.EPQVYTLPPSQEEMTK.N | | 225-240 | CID |

| Cmpd. | m/z meas. | z | Δ m/z [ppm] | Δ m/z [Da] | Rt [min] | Score | No. of Cmpds. | Site [%] | P | Sequence | Modification | Range | Type |
|---|---|---|---|---|---|---|---|---|---|---|---|---|---|
| 680 | 581.357 | 2 | 66.32 | 0.0386 | 26.76 | 82.12 | 12 | 0.0 | 0 | K.NQVSLTCLVK.G | Carbamidomethyl: 7 | 241-250 | CID |
| 880 | 1273.041 | 2 | 370.65 | 0.4717 | 30.87 | 132.13 | 40 | 0.0 | 0 | K.GFYPSDIAVEWESNGQPENNYK.T | | 251-272 | CID |
| 960 | 951.501 | 2 | 35.08 | 0.0334 | 32.50 | 77.47 | 1 | 0.0 | 0 | K.TTPPVLDSDGSFFLYSR.L | | 273-289 | CID |

#### Protein 7: Immunoglobulin lambda constant 3 OS=Homo sapiens GN=IGLC3 PE=1 SV=1

**Accession:** IGLC3_HUMAN
**Database:** SwissProt
**MW [kDa] / pI:** 11.3 / 6.9
**Modification(s):** Carbamidomethyl

**Score:** 619.7
**Seq. Coverage [%]:** 93.4
**No. of Peptides:** 8

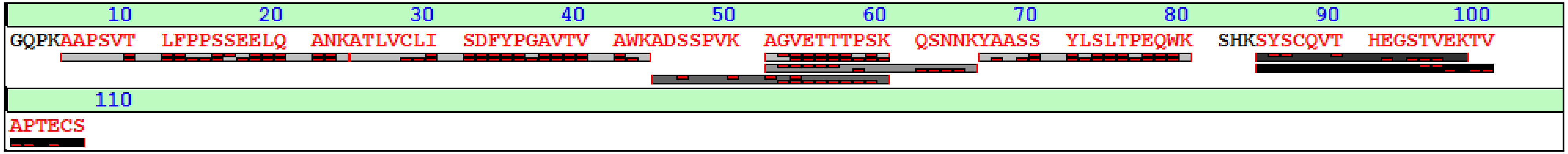


MS/MS Peptide Matches

| Cmpd. | m/z meas. | z | Δ m/z [ppm] | Δ m/z [Da] | Rt [min] | Score | No. of Cmpds. | Site [%] | P | Sequence | Modification | Range | Type |
|---|---|---|---|---|---|---|---|---|---|---|---|---|---|
| 829 | 993.571 | 2 | 58.82 | 0.0584 | 29.81 | 83.63 | 19 | 0.0 | 0 | K.AAPSVTLFPPSSEELQANK.A | | 5-23 | CID |
| 1325 | 1106.118 | 2 | 34.76 | 0.0384 | 40.29 | 107.09 | 3 | 0.0 | 0 | K.ATLVCLISDFYPGAVTVAWK.A | Carbamidomethyl: 5 | 24-43 | CID |
| 267 | 558.987 | 3 | 54.85 | 0.0307 | 17.76 | 65.13 | 2 | 0.0 | 1 | K.ADSSPVKAGVETTTPSK.Q | | 44-60 | CID |
| 124 | 495.75 | 2 | -17.64 | -0.0087 | 14.57 | 79.13 | 17 | 0.0 | 0 | K.AGVETTTPSK.Q | | 51-60 | CID |
| 74 | 521.321 | 3 | 106.67 | 0.0556 | 13.42 | 70.9 | 3 | 0.0 | 1 | K.AGVETTTPSKQSNNK.Y | | 51-65 | CID |
| 863 | 872.589 | 2 | 178.75 | 0.1559 | 30.47 | 95.15 | 18 | 0.0 | 0 | K.YAASSYLSLTPEQWK.S | | 66-80 | CID |
| 222 | 571.336 | 3 | 136.69 | 0.0781 | 16.77 | 61.95 | 15 | 0.0 | 0 | K.SYSCQVTHEGSTVEK.T | Carbamidomethyl: 4 | 84-98 | CID |
| 428 | 853.078 | 3 | 39.60 | 0.0338 | 21.11 | 56.72 | 3 | 0.0 | 1 | K.SYSCQVTHEGSTVEKTVAPTECS.- | Carbamidomethyl: 4, 22 | 84-106 | CID |

##### Protein 8: Immunoglobulin lambda-like polypeptide 5 OS=Homo sapiens GN=IGLL5 PE=2 SV=2

| | | | |
|---|---|---|---|
| **Accession:** | IGLL5_HUMAN | **Score:** | 467.1 |
| **Database:** | SwissProt | **Seq. Coverage [%]:** | 43.9 |
| **MW [kDa] / pI:** | 23.0 / 9.1 | **No. of Peptides:** | 7 |
| **Modification(s):** | Carbamidomethyl | | |

| 10 | 20 | 30 | 40 | 50 | 60 | 70 | 80 | 90 | 100 |
|---|---|---|---|---|---|---|---|---|---|
| MRPKTGQVGC | ETPEELGPGP | RQRWPLLLLG | LAMVAHGLLR | PMVAPQSGDP | DPGASVGSSR | SSLRSLWGRL | LLQPSPQRAD | PRCWPRGFWS | EPQSLCYVFG |
| **110** | **120** | **130** | **140** | **150** | **160** | **170** | **180** | **190** | **200** |
| TGTKVTVLGQ | PKANPTVTLF | PPSSEELQAN | KATLVCLISD | FYPGAVTVAW | KADGSPVKAG | VETTKPSKQS | NNKYAASSYL | SLTPEQWKSH | RSYSCQVTHE |
| **210** | **220** | | | | | | | | |
| GSTVEKTVAP | TECS | | | | | | | | |

**MS/MS Peptide Matches**

| Cmpd. | m/z meas. | z | Δ m/z [ppm] | Δ m/z [Da] | Rt [min] | Score | No. of Cmpds. | Site [%] | P | Sequence | Modification | Range | Type |
|---|---|---|---|---|---|---|---|---|---|---|---|---|---|
| 601 | 526.353 | 2 | 69.23 | 0.0364 | 24.84 | 17.41 | 1 | 0.0 | 0 | R.LLLQPSPQR.A | | 70-78 | CID |
| 356 | 421.301 | 2 | 95.60 | 0.0403 | 19.76 | 53.67 | 4 | 0.0 | 0 | K.VTVLGQPK.A | | 105-112 | CID |
| 794 | 1022.064 | 2 | 39.83 | 0.0407 | 29.08 | 75.08 | 12 | 0.0 | 0 | K.ANPTVTLFPPSSEELQANK.A | | 113-131 | CID |
| 1325 | 1106.118 | 2 | 34.76 | 0.0384 | 40.29 | 107.09 | 3 | 0.0 | 0 | K.ATLVCLISDFYPGAVTVAWK.A | Carbamidomethyl: 5 | 132-151 | CID |
| 863 | 872.589 | 2 | 178.75 | 0.1559 | 30.47 | 95.15 | 18 | 0.0 | 0 | K.YAASSYLSLTPEQWK.S | | 174-188 | CID |
| 222 | 571.336 | 3 | 136.69 | 0.0781 | 16.77 | 61.95 | 15 | 0.0 | 0 | R.SYSCQVTHEGSTVEK.T | Carbamidomethyl: 4 | 192-206 | CID |
| 428 | 853.078 | 3 | 39.60 | 0.0338 | 21.11 | 56.72 | 3 | 0.0 | 1 | R.SYSCQVTHEGSTVEKTVAPTECS.- | Carbamidomethyl: 4, 22 | 192-214 | CID |

#### Protein 9: Immunoglobulin kappa variable 2D-28 OS=Homo sapiens GN=IGKV2D-28 PE=1 SV=2

| | | | |
|---|---|---|---|
| **Accession:** | KVD28_HUMAN | **Score:** | 167.2 |
| **Database:** | SwissProt | **Seq. Coverage [%]:** | 16.7 |
| **MW [kDa] / pI:** | 12.9 / 5.6 | **No. of Peptides:** | 2 |

| 10 | 20 | 30 | 40 | 50 | 60 | 70 | 80 | 90 | 100 |
|---|---|---|---|---|---|---|---|---|---|
| MRLPAQLLGL | LMLWVSGSSG | DIVMTQSPLS | LPVTPGEPAS | ISCRSSQSLL | HSNGYNYLDW | YLQKPGQSPQ | LLIYLGSNRA | SGVPDRFSGS | GSGTDFTLKI |
| **110** | **120** | **130** | | | | | | | |
| SRVEAEDVGV | YYCMQALQTP | | | | | | | | |

##### MS/MS Peptide Matches

| Cmpd. | m/z meas. | z | Δ m/z [ppm] | Δ m/z [Da] | Rt [min] | Score | No. of Cmpds. | Site [%] | P | Sequence | Modification | Range | Type |
|---|---|---|---|---|---|---|---|---|---|---|---|---|---|
| 602 | 662.688 | 3 | 46.85 | 0.0310 | 24.87 | 73.23 | 3 | 0.0 | 1 | R.ASGVPDRFSGSGSGTDFTLK.I | | 80-99 | CID |
| 556 | 652.314 | 2 | 3.26 | 0.0021 | 23.89 | 93.96 | 3 | 0.0 | 0 | R.FSGSGSGTDFTLK.I | | 87-99 | CID |

#### Protein 10: Immunoglobulin heavy variable 3-7 OS=Homo sapiens GN=IGHV3-7 PE=1 SV=2

| | | | |
|---|---|---|---|
| **Accession:** | HV307_HUMAN | **Score:** | 140.6 |
| **Database:** | SwissProt | **Seq. Coverage [%]:** | 17.1 |
| **MW [kDa] / pI:** | 12.9 / 6.2 | **No. of Peptides:** | 2 |
| **Modification(s):** | Carbamidomethyl | | |

| 10 | 20 | 30 | 40 | 50 | 60 | 70 | 80 | 90 | 100 |
|---|---|---|---|---|---|---|---|---|---|
| MELGLSWVFL | VAILEGVQCE | VQLVESGGGL | VQPGGSLRLS | CAASGFTFSS | YWMSWVRQAP | GKGLEWVANI | KQDGSEKYYV | DSVKGRFTIS | RDNAKNSLYL |
| **110** | **120** | | | | | | | | |
| QMNSLRAEDT | AVYYCAR | | | | | | | | |

##### MS/MS Peptide Matches

| Cmpd. | m/z meas. | z | Δ m/z [ppm] | Δ m/z [Da] | Rt [min] | Score | No. of Cmpds. | Site [%] | P | Sequence | Modification | Range | Type |
|---|---|---|---|---|---|---|---|---|---|---|---|---|---|
| 791 | 515.329 | 2 | 75.65 | 0.0390 | 29.00 | 57.16 | 1 | 0.0 | 0 | K.GLEWVANIK.Q | | 63-71 | CID |
| 395 | 659.83 | 2 | 60.24 | 0.0397 | 20.46 | 83.46 | 3 | 0.0 | 0 | R.AEDTAVYYCAR.- | Carbamidomethyl: 9 | 107-117 | CID |

#### Protein 11: Serum albumin OS=Homo sapiens GN=ALB PE=1 SV=2

**Accession:** ALBU_HUMAN
**Database:** SwissProt
**MW [kDa] / pI:** 69.3 / 5.9
**Modification(s):** Carbamidomethyl

**Score:** 133.1
**Seq. Coverage [%]:** 6.6
**No. of Peptides:** 2

| 10 | 20 | 30 | 40 | 50 | 60 | 70 | 80 | 90 | 100 |
|---|---|---|---|---|---|---|---|---|---|
| MKWVTFISLL | FLFSSAYSRG | VFRRDAHKSE | VAHRFKDLGE | ENFKALVLIA | FAQYLQQCPF | EDHVKLVNEV | TEFAKTCVAD | ESAENCDKSL | HTLFGDKLCT |
| 110 | 120 | 130 | 140 | 150 | 160 | 170 | 180 | 190 | 200 |
| VATLRETYGE | MADCCAKQEP | ERNECFLQHK | DDNPNLPRLV | RPEVDVMCTA | FHDNEETFLK | KYLYEIARRH | PYFYAPELLF | FAKRYKAAFT | ECCQAADKAA |
| 210 | 220 | 230 | 240 | 250 | 260 | 270 | 280 | 290 | 300 |
| CLLPKLDELR | DEGKASSAKQ | RLKCASLQKF | GERAFKAWAV | ARLSQRFPKA | EFAEVSKLVT | DLTKVHTECC | HGDLLECADD | RADLAKYICE | NQDSISSKLK |
| 310 | 320 | 330 | 340 | 350 | 360 | 370 | 380 | 390 | 400 |
| ECCEKPLLEK | SHCIAEVEND | EMPADLPSLA | ADFVESKDVC | KNYAEAKDVF | LGMFLYEYAR | RHPDYSVVLL | LRLAKTYETT | LEKCCAADP | HECYAKVFDE |
| 410 | 420 | 430 | 440 | 450 | 460 | 470 | 480 | 490 | 500 |
| FKPLVEEPQN | LIKQNCELFE | QLGEYKFQNA | LLVRYTKKVP | QVSTPTLVEV | SRNLGKVGSK | CCKHPEAKRM | PCAEDYLSVV | LNQLCVLHEK | TPVSDRVTKC |
| 510 | 520 | 530 | 540 | 550 | 560 | 570 | 580 | 590 | 600 |
| CTESLVNRRP | CFSALEVDET | YVPKEFNAET | FTFHADICTL | SEKERQIKKQ | TALVELVKHK | PKATKEQLKA | VMDDFAAFVE | KCCKADDKET | CFAEEGKKLV |
| 610 | | | | | | | | | |
| AASQAALGL | | | | | | | | | |

MS/MS Peptide Matches

| Cmpd. | m/z meas. | z | Δ m/z [ppm] | Δ m/z [Da] | Rt [min] | Score | No. of Cmpds. | Site [%] | P | Sequence | Modification | Range | Type |
|---|---|---|---|---|---|---|---|---|---|---|---|---|---|
| 1161 | 992.515 | 3 | 398.46 | 0.3953 | 36.93 | 53.73 | 4 | 0.0 | 0 | K.SHCIAEVENDEMPADLPSLAADFVESK.D | Carbamidomethyl: 3 | 311-337 | CID |
| 1280 | 812.431 | 2 | 41.33 | 0.0336 | 39.31 | 79.37 | 3 | 0.0 | 0 | K.DVFLGMFLYEYAR.R | | 348-360 | CID |

#### Protein 12: Immunoglobulin kappa variable 3-11 OS=Homo sapiens GN=IGKV3-11 PE=1 SV=1

**Accession:** KV311_HUMAN
**Database:** SwissProt
**MW [kDa] / pI:** 12.6 / 4.9
**Modification(s):** Carbamidomethyl

**Score:** 112.1
**Seq. Coverage [%]:** 33.9
**No. of Peptides:** 2

```
         10         20         30         40         50         60         70         80         90        100
MEAPAQLLFL LLLWLPDTTG EIVLTQSPAT LSLSPGERAT LSCRASQSVS SYLAWYQQKP GQAPRLLIYD ASNRATGIPA RFSGSGSGTD FTLTISSLEP
        110        120
EDFAVYYCQQ RSNWP
```

MS/MS Peptide Matches

| Cmpd. | m/z meas. | z | Δ m/z [ppm] | Δ m/z [Da] | Rt [min] | Score | No. of Cmpds. | Site [%] | P | Sequence | Modification | Range | Type |
|---|---|---|---|---|---|---|---|---|---|---|---|---|---|
| 577 | 532.839 | 2 | 91.25 | 0.0486 | 24.32 | 71.27 | 4 | 0.0 | 0 | R.LLIYDASNR.A | | 66-74 | CID |
| 1132 | 1121.86 | 3 | 311.98 | 0.3499 | 36.34 | 40.86 | 3 | 0.0 | 0 | R.FSGSGSGTDFTLTISSLEPEDFAVYYCQQR.S | Carbamidomethyl: 27 | 82-111 | CID |

#### Protein 13: Immunoglobulin kappa variable 3-20 OS=Homo sapiens GN=IGKV3-20 PE=1 SV=2

**Accession:** KV320_HUMAN
**Database:** SwissProt
**MW [kDa] / pI:** 12.5 / 4.9

**Score:** 66.7
**Seq. Coverage [%]:** 7.8
**No. of Peptides:** 1

| 10 | 20 | 30 | 40 | 50 | 60 | 70 | 80 | 90 | 100 |
|---|---|---|---|---|---|---|---|---|---|
| METPAQLLFL | LLLWLPDTTG | EIVLTQSPGT | LSLSPGERAT | LSCRASQSVS | SSYLAWYQQK | PGQAPRLLIY | GASSRATGIP | DRFSGSGSGT | DFTLTISRLE |
| 110 | 120 | | | | | | | | |
| PEDFAVYYCQ | QYGSSP | | | | | | | | |

MS/MS Peptide Matches

| Cmpd. | m/z meas. | z | Δ m/z [ppm] | Δ m/z [Da] | Rt [min] | Score | No. of Cmpds. | Site [%] | P | Sequence | Modification | Range | Type |
|---|---|---|---|---|---|---|---|---|---|---|---|---|---|
| 560 | 490.328 | 2 | 93.43 | 0.0458 | 23.95 | 66.73 | 3 | 0.0 | 0 | R.LLIYGASSR.A | | 67-75 | CID |

#### Protein 14: Immunoglobulin kappa variable 3-15 OS=Homo sapiens GN=IGKV3-15 PE=1 SV=2

**Accession:** KV315_HUMAN
**Database:** SwissProt
**MW [kDa] / pI:** 12.5 / 5.1

**Score:** 66.6
**Seq. Coverage [%]:** 7.8
**No. of Peptides:** 1

| 10 | 20 | 30 | 40 | 50 | 60 | 70 | 80 | 90 | 100 |
|---|---|---|---|---|---|---|---|---|---|
| MEAPAQLLFL | LLLWLPDTTG | EIVMTQSPAT | LSVSPGERAT | LSCRASQSVS | SNLAWYQQKP | GQAPRLLIYG | ASTRATGIPA | RFSGSGSGTE | FTLTISSLQS |
| 110 | 120 | | | | | | | | |
| EDFAVYYCQQ | YNNWP | | | | | | | | |

MS/MS Peptide Matches

| Cmpd. | m/z meas. | z | Δ m/z [ppm] | Δ m/z [Da] | Rt [min] | Score | No. of Cmpds. | Site [%] | P | Sequence | Modification | Range | Type |
|---|---|---|---|---|---|---|---|---|---|---|---|---|---|
| 580 | 497.328 | 2 | 76.38 | 0.0380 | 24.39 | 66.58 | 3 | 0.0 | 0 | R.LLIYGASTR.A | | 66-74 | CID |

#### Protein 15: Immunoglobulin kappa variable 4-1 OS=Homo sapiens GN=IGKV4-1 PE=1 SV=1

**Accession:** KV401_HUMAN
**Database:** SwissProt
**MW [kDa] / pI:** 13.4 / 5.1

**Score:** 55.5
**Seq. Coverage [%]:** 12.4
**No. of Peptides:** 1

| 10 | 20 | 30 | 40 | 50 | 60 | 70 | 80 | 90 | 100 |
|---|---|---|---|---|---|---|---|---|---|
| MVLQTQVFIS | LLLWISGAYG | DIVMTQSPDS | LAVSLGERAT | INCKSSQSVL | YSSNNKNYLA | WYQQKPGQPP | KLLIYWASTR | ESGVPDRFSG | SGSGTDFTLT |
| 110 | 120 | 130 | | | | | | | |
| ISSLQAEDVA | VYYCQQYYST | P | | | | | | | |

MS/MS Peptide Matches

| Cmpd. | m/z meas. | z | Δ m/z [ppm] | Δ m/z [Da] | Rt [min] | Score | No. of Cmpds. | Site [%] | P | Sequence | Modification | Range | Type |
|---|---|---|---|---|---|---|---|---|---|---|---|---|---|
| 611 | 606.687 | 3 | 62.15 | 0.0377 | 25.02 | 55.47 | 2 | 0.0 | 0 | K.NYLAWYQQKPGQPPK.L | | 57-71 | CID |

#### Protein 16: Immunoglobulin heavy variable 3-73 OS=Homo sapiens GN=IGHV3-73 PE=3 SV=1

| | | | |
|---|---|---|---|
| **Accession:** | HV373_HUMAN | **Score:** | 51.4 |
| **Database:** | SwissProt | **Seq. Coverage [%]:** | 5.9 |
| **MW [kDa] / pI:** | 12.8 / 9.3 | **No. of Peptides:** | 1 |

| 10 | 20 | 30 | 40 | 50 | 60 | 70 | 80 | 90 | 100 |
|---|---|---|---|---|---|---|---|---|---|
| MEFGLSWVFL | VAILKGVQCE | VQLVESGGGL | VQPGGSLKLS | CAASGFTFSG | SAMHWVRQAS | GKGLEWVGRI | RSKANSYATA | YAASVKGRFT | ISRDDSKNTA |
| 110 | 120 | | | | | | | | |
| YLQMNSLKTE | DTAVYYCTR | | | | | | | | |

MS/MS Peptide Matches

| Cmpd. | m/z meas. | z | Δ m/z [ppm] | Δ m/z [Da] | Rt [min] | Score | No. of Cmpds. | Site [%] | P | Sequence | Modification | Range | Type |
|---|---|---|---|---|---|---|---|---|---|---|---|---|---|
| 584 | 408.77 | 2 | 118.00 | 0.0482 | 24.47 | 51.43 | 4 | 0.0 | 0 | K.GLEWVGR.I | | 63-69 | CID |

### Protein Report
#### IonTrap_Human_Swiss_CID_2019-03-05 17:55:08

##### Project Info

| | | | |
|---|---|---|---|
| **Name:** | Alagesan | **Date:** | Mar 5, 2019 |
| **Note:** | Nanobooster | | |

##### Sample Info & Protocols

| | |
|---|---|
| **Name:** | Nanobooster - IgG |
| **Date:** | Mar 5, 2019 |

##### Search Result Info

| | | | |
|---|---|---|---|
| **Search Result:** | IonTrap_Human_Swiss_CID_2019-03-05 17:55:08 | | |
| **Location:** | /Alagesan/Nanobooster - IgG/Ka_IgG Trypsin NoEnrich NB Ace_BA5_01_8763.mgf | | |
| **Search Method:** | IonTrap_Human_Swiss_CID | **Version:** | 1.2 |
| **Search Engine(s):** | Mascot, 2.6.0 | **Ident. Compound(s):** | 0/2071 |
| **Database(s):** | SwissProt, SwissProt_2017_04.fasta | | |
| **Note:** | | | |

#### Results

##### Protein 1: Immunoglobulin kappa light chain OS=Homo sapiens PE=1 SV=1

| | | | |
|---|---|---|---|
| **Accession:** | IGK_HUMAN | **Score:** | 987.3 |
| **Database:** | SwissProt | **Seq. Coverage [%]:** | 48.1 |
| **MW [kDa] / pI:** | 23.4 / 6.9 | **No. of Peptides:** | 10 |
| **Modification(s):** | Carbamidomethyl | | |

```
        10         20         30         40         50         60         70         80         90        100
DIQMTQSPST LSASVGDRVT ITCRASQSIN TWLAWYQQKP GKAPKLLMYK ASSLESGVPS RFIGSGSGTE FTLTISSLQP DDFATYYCQQ YNSDSKMFGQ
       110        120        130        140        150        160        170        180        190        200
GTKVEVKGTV AAPSVFIFPP SDEQLKSGTA SVVCLLNNFY PREAKVQWKV DNALQSGNSQ ESVTEQDSKD STYSLSSTLT LSKADYEKHK VYACEVTHQG
       210        220
LSSPVTKSFN RGEC
```

**MS/MS Peptide Matches**

| Cmpd. | m/z meas. | z | Δ m/z [ppm] | Δ m/z [Da] | Rt [min] | Score | No. of Cmpds. | Site [%] | P | Sequence | Modification | Range | Type |
|---|---|---|---|---|---|---|---|---|---|---|---|---|---|
| 605 | 946.928 | 2 | -28.04 | -0.0266 | 25.11 | 152.71 | 1 | 0.0 | 0 | -.DIQMTQSPSTLSASVGDR.V | | 1-18 | CID |
| 1105 | 899.625 | 2 | 193.17 | 0.1737 | 35.78 | 76.7 | 31 | 0.0 | 0 | K.SGTASVVCLLNNFYPR.E | Carbamidomethyl: 8 | 127-142 | CID |
| 549 | 893.364 | 3 | 301.37 | 0.2692 | 23.88 | 108.94 | 3 | 0.0 | 1 | K.VQWKVDNALQSGNSQESVTEQDSK.D | | 146-169 | CID |
| 326 | 1068.659 | 2 | 160.04 | 0.1710 | 19.11 | 165.53 | 70 | 0.0 | 0 | K.VDNALQSGNSQESVTEQDSK.D | | 150-169 | CID |
| 727 | 1207.545 | 3 | 251.57 | 0.3037 | 27.89 | 151.13 | 22 | 0.0 | 1 | K.VDNALQSGNSQESVTEQDSKDSTYSLSSTLTLS K.A | | 150-183 | CID |
| 738 | 751.951 | 2 | 90.63 | 0.0681 | 28.11 | 89.01 | 7 | 0.0 | 0 | K.DSTYSLSSTLTLSK.A | | 170-183 | CID |
| 10 | 445.766 | 2 | 98.79 | 0.0440 | 11.32 | 18.69 | 1 | 0.0 | 1 | K.ADYEKHK.V | | 184-190 | CID |
| 337 | 536.334 | 4 | 575.24 | 0.3083 | 19.42 | 58.21 | 7 | 0.0 | 1 | K.HKVYACEVTHQGLSSPVTK.S | Carbamidomethyl: 6 | 189-207 | CID |
| 448 | 626.028 | 3 | 75.89 | 0.0475 | 21.68 | 112.2 | 8 | 0.0 | 0 | K.VYACEVTHQGLSSPVTK.S | Carbamidomethyl: 4 | 191-207 | CID |

| Cmpd. | m/z meas. | z | Δ m/z [ppm] | Δ m/z [Da] | Rt [min] | Score | No. of Cmpds. | Site [%] | P | Sequence | Modification | Range | Type |
|---|---|---|---|---|---|---|---|---|---|---|---|---|---|
| 99 | 435.158 | 2 | -55.50 | -0.0242 | 13.99 | 54.22 | 9 | 0.0 | 1 | K.SFNRGEC.- | Carbamidomethyl: 7 | 208-214 | CID |

#### Protein 2: Immunoglobulin kappa constant OS=Homo sapiens GN=IGKC PE=1 SV=2

**Accession:** IGKC_HUMAN
**Database:** SwissProt
**MW [kDa] / pI:** 11.8 / 6.1
**Modification(s):** Carbamidomethyl
**Score:** 986.6
**Seq. Coverage [%]:** 97.2
**No. of Peptides:** 11

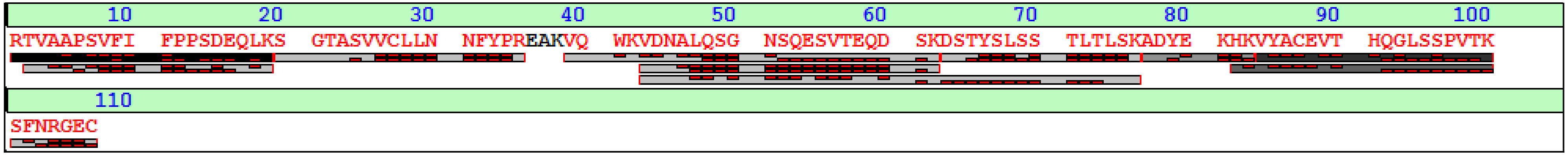


MS/MS Peptide Matches

| Cmpd. | m/z meas. | z | Δ m/z [ppm] | Δ m/z [Da] | Rt [min] | Score | No. of Cmpds. | Site [%] | P | Sequence | Modification | Range | Type |
|---|---|---|---|---|---|---|---|---|---|---|---|---|---|
| 898 | 701.422 | 3 | 58.64 | 0.0411 | 31.28 | 47.3 | 11 | 0.0 | 1 | -.RTVAAPSVFIFPPSDEQLK.S | | 1-19 | CID |
| 1027 | 973.658 | 2 | 144.72 | 0.1409 | 34.07 | 104.68 | 40 | 0.0 | 0 | R.TVAAPSVFIFPPSDEQLK.S | | 2-19 | CID |
| 1105 | 899.625 | 2 | 193.17 | 0.1737 | 35.78 | 76.7 | 31 | 0.0 | 0 | K.SGTASVVCLLNNFYPR.E | Carbamidomethyl: 8 | 20-35 | CID |
| 549 | 893.364 | 3 | 301.37 | 0.2692 | 23.88 | 108.94 | 3 | 0.0 | 1 | K.VQWKVDNALQSGNSQESVTEQDSK.D | | 39-62 | CID |
| 326 | 1068.659 | 2 | 160.04 | 0.1710 | 19.11 | 165.53 | 70 | 0.0 | 0 | K.VDNALQSGNSQESVTEQDSK.D | | 43-62 | CID |
| 727 | 1207.545 | 3 | 251.57 | 0.3037 | 27.89 | 151.13 | 22 | 0.0 | 1 | K.VDNALQSGNSQESVTEQDSKDSTYSLSSTLTLSK.A | | 43-76 | CID |
| 738 | 751.951 | 2 | 90.63 | 0.0681 | 28.11 | 89.01 | 7 | 0.0 | 0 | K.DSTYSLSSTLTLSK.A | | 63-76 | CID |
| 10 | 445.766 | 2 | 98.79 | 0.0440 | 11.32 | 18.69 | 1 | 0.0 | 1 | K.ADYEKHK.V | | 77-83 | CID |
| 337 | 536.334 | 4 | 575.24 | 0.3083 | 19.42 | 58.21 | 7 | 0.0 | 1 | K.HKVYACEVTHQGLSSPVTK.S | Carbamidomethyl: 6 | 82-100 | CID |
| 448 | 626.028 | 3 | 75.89 | 0.0475 | 21.68 | 112.2 | 8 | 0.0 | 0 | K.VYACEVTHQGLSSPVTK.S | Carbamidomethyl: 4 | 84-100 | CID |

| Cmpd. | m/z meas. | z | Δ m/z [ppm] | Δ m/z [Da] | Rt [min] | Score | No. of Cmpds. | Site [%] | P | Sequence | Modification | Range | Type |
|---|---|---|---|---|---|---|---|---|---|---|---|---|---|
| 99 | 435.158 | 2 | -55.50 | -0.0242 | 13.99 | 54.22 | 9 | 0.0 | 1 | K.SFNRGEC.- | Carbamidomethyl: 7 | 101-107 | CID |

#### Protein 3: Immunoglobulin heavy constant gamma 1 OS=Homo sapiens GN=IGHG1 PE=1 SV=1

**Accession:** IGHG1_HUMAN
**Database:** SwissProt
**MW [kDa] / pI:** 36.1 / 8.5
**Modification(s):** Carbamidomethyl, Oxidation

**Score:** 971.1
**Seq. Coverage [%]:** 54.8
**No. of Peptides:** 15

| 10 | 20 | 30 | 40 | 50 | 60 | 70 | 80 | 90 | 100 |
|---|---|---|---|---|---|---|---|---|---|
| ASTKGPSVFP | LAPSSKSTSG | GTAALGCLVK | DYFPEPVTVS | WNSGALTSGV | HTFPAVLQSS | GLYSLSSVVT | VPSSSLGTQT | YICNVNHKPS | NTKVDKKVEP |
| **110** | **120** | **130** | **140** | **150** | **160** | **170** | **180** | **190** | **200** |
| KSCDKTHTCP | PCPAPELLGG | PSVFLFPPKP | KDTLMISRTP | EVTCVVVDVS | HEDPEVKFNW | YVDGVEVHNA | KTKPREEQYN | STYRVVSVLT | VLHQDWLNGK |
| **210** | **220** | **230** | **240** | **250** | **260** | **270** | **280** | **290** | **300** |
| EYKCKVSNKA | LPAPIEKTIS | KAKGQPREPQ | VYTLPPSRDE | LTKNQVSLTC | LVKGFYPSDI | AVEWESNGQP | ENNYKTTPPV | LDSDGSFFLY | SKLTVDKSRW |
| **310** | **320** | **330** | **340** | | | | | | |
| QQGNVFSCSV | MHEALHNHYT | QKSLSLSPGK | | | | | | | |

MS/MS Peptide Matches

| Cmpd. | m/z meas. | z | Δ m/z [ppm] | Δ m/z [Da] | Rt [min] | Score | No. of Cmpds. | Site [%] | P | Sequence | Modification | Range | Type |
|---|---|---|---|---|---|---|---|---|---|---|---|---|---|
| 708 | 593.838 | 2 | 18.59 | 0.0110 | 27.57 | 52.15 | 14 | 0.0 | 0 | K.GPSVFPLAPSSK.S | | 5-16 | CID |
| 613 | 661.426 | 2 | 126.03 | 0.0833 | 25.27 | 112.76 | 18 | 0.0 | 0 | K.STSGGTAALGCLVK.D | Carbamidomethyl: 11 | 17-30 | CID |
| 1009 | 712.188 | 4 | 446.93 | 0.3182 | 33.61 | 42.31 | 5 | 0.0 | 0 | K.THTCPPCPAPELLGGPSVFLFPPKPK.D | Carbamidomethyl: 4, 7 | 106-131 | CID |
| 456 | 418.225 | 2 | 10.17 | 0.0043 | 21.87 | 47.58 | 9 | 0.0 | 0 | K.DTLMISR.T | | 132-138 | CID |
| 358 | 426.256 | 2 | 88.68 | 0.0378 | 19.90 | 47.25 | 4 | 0.0 | 0 | K.DTLMISR.T | Oxidation: 4 | 132-138 | CID |
| 704 | 713.719 | 3 | 53.72 | 0.0383 | 27.49 | 67.03 | 1 | 0.0 | 0 | R.TPEVTCVVVDVSHEDPEVK.F | Carbamidomethyl: 6 | 139-157 | CID |

| Cmpd. | m/z meas. | z | Δ m/z [ppm] | Δ m/z [Da] | Rt [min] | Score | No. of Cmpds. | Site [%] | P | Sequence | Modification | Range | Type |
|---|---|---|---|---|---|---|---|---|---|---|---|---|---|
| 1140 | 950.29 | 4 | 85.93 | 0.0817 | 36.58 | 44.3 | 7 | 0.0 | 1 | R.TPEVTCVVVDVSHEDPEVKFNWYVDGVEVHNAK.T | Carbamidomethyl: 6 | 139-171 | CID |
| 699 | 560.022 | 3 | 148.51 | 0.0832 | 27.40 | 72.64 | 14 | 0.0 | 0 | K.FNWYVDGVEVHNAK.T | | 158-171 | CID |
| 1057 | 603.404 | 3 | 105.50 | 0.0637 | 34.72 | 71.31 | 10 | 0.0 | 0 | R.VVSVLTVLHQDWLNGK.E | | 185-200 | CID |
| 446 | 419.792 | 2 | 87.48 | 0.0367 | 21.62 | 31.37 | 2 | 0.0 | 0 | K.ALPAPIEK.T | | 210-217 | CID |
| 544 | 643.819 | 2 | -33.55 | -0.0216 | 23.78 | 57.48 | 9 | 0.0 | 0 | R.EPQVYTLPPSR.D | | 228-238 | CID |
| 585 | 625.043 | 3 | 76.95 | 0.0481 | 24.65 | 26.23 | 8 | 0.0 | 1 | R.EPQVYTLPPSRDELTK.N | | 228-243 | CID |
| 682 | 581.368 | 2 | 85.24 | 0.0496 | 26.98 | 68.52 | 14 | 0.0 | 0 | K.NQVSLTCLVK.G | Carbamidomethyl: 7 | 244-253 | CID |
| 893 | 1273.046 | 2 | 374.58 | 0.4767 | 31.19 | 125.42 | 37 | 0.0 | 0 | K.GFYPSDIAVEWESNGQPENNYK.T | | 254-275 | CID |
| 956 | 937.694 | 2 | 244.76 | 0.2295 | 32.47 | 104.76 | 19 | 0.0 | 0 | K.TTPPVLDSDGSFFLYSK.L | | 276-292 | CID |

#### Protein 4: Immunoglobulin heavy constant gamma 3 OS=Homo sapiens GN=IGHG3 PE=1 SV=2

**Accession:** IGHG3_HUMAN
**Database:** SwissProt
**MW [kDa] / pI:** 41.3 / 8.2
**Modification(s):** Carbamidomethyl, Oxidation

**Score:** 848.6
**Seq. Coverage [%]:** 36.3
**No. of Peptides:** 13

| | | | | | | | | | |
|---|---|---|---|---|---|---|---|---|---|
| 10 | 20 | 30 | 40 | 50 | 60 | 70 | 80 | 90 | 100 |
| ASTKGPSVFP | LAPCSRSTSG | GTAALGCLVK | DYFPEPVTVS | WNSGALTSGV | HTFPAVLQSS | GLYSLSSVVT | VPSSSLGTQT | YTCNVNHKPS | NTKVDKRVEL |
| 110 | 120 | 130 | 140 | 150 | 160 | 170 | 180 | 190 | 200 |
| KTPLGDTTHT | CPRCPEPKSC | DTPPPCPRCP | EPKSCDTPPP | CPRCPEPKSC | DTPPPCPRCP | APELLGGPSV | FLFPPKPKDT | LMISRTPEVT | CVVVDVSHED |
| 210 | 220 | 230 | 240 | 250 | 260 | 270 | 280 | 290 | 300 |
| PEVQFKWYVD | GVEVHNAKTK | PREEQYNSTF | RVVSVLTVLH | QDWLNGKEYK | CKVSNKALPA | PIEKTISKTK | GQPREPQVYT | LPPSREEMTK | NQVSLTCLVK |
| 310 | 320 | 330 | 340 | 350 | 360 | 370 | 380 | | |
| GFYPSDIAVE | WESSGQPENN | YNTTPPMLDS | DGSFFLYSKL | TVDKSRWQQG | NIFSCSVMHE | ALHNRFTQKS | LSLSPGK | | |

MS/MS Peptide Matches

| Cmpd. | m/z meas. | z | Δ m/z [ppm] | Δ m/z [Da] | Rt [min] | Score | No. of Cmpds. | Site [%] | P | Sequence | Modification | Range | Type |
|---|---|---|---|---|---|---|---|---|---|---|---|---|---|
| 745 | 644.38 | 2 | 78.61 | 0.0507 | 28.23 | 69.32 | 14 | 0.0 | 0 | K.GPSVFPLAPCSR.S | Carbamidomethyl: 10 | 5-16 | CID |
| 613 | 661.426 | 2 | 126.03 | 0.0833 | 25.27 | 112.76 | 18 | 0.0 | 0 | R.STSGGTAALGCLVK.D | Carbamidomethyl: 11 | 17-30 | CID |
| 228 | 452.548 | 3 | -5.73 | -0.0026 | 17.01 | 72.7 | 3 | 0.0 | 0 | K.TPLGDTTHTCPR.C | Carbamidomethyl: 10 | 102-113 | CID |
| 192 | 593.786 | 2 | 56.23 | 0.0334 | 16.17 | 68.69 | 11 | 0.0 | 0 | K.SCDTPPPCPR.C | Carbamidomethyl: 2, 8 | 119-128 | CID |
| 1047 | 717.832 | 3 | 145.99 | 0.1048 | 34.52 | 100.35 | 3 | 0.0 | 0 | R.CPAPELLGGPSVFLFPPKPK.D | Carbamidomethyl: 1 | 159-178 | CID |
| 456 | 418.225 | 2 | 10.17 | 0.0043 | 21.87 | 47.58 | 9 | 0.0 | 0 | K.DTLMISR.T | | 179-185 | CID |
| 358 | 426.256 | 2 | 88.68 | 0.0378 | 19.90 | 47.25 | 4 | 0.0 | 0 | K.DTLMISR.T | Oxidation: 4 | 179-185 | CID |
| 546 | 472.942 | 3 | 85.15 | 0.0403 | 23.84 | 52.07 | 3 | 0.0 | 0 | K.WYVDGVEVHNAK.T | | 207-218 | CID |
| 1057 | 603.404 | 3 | 105.50 | 0.0637 | 34.72 | 71.31 | 10 | 0.0 | 0 | R.VVSVLTVLHQDWLNGK.E | | 232-247 | CID |
| 446 | 419.792 | 2 | 87.48 | 0.0367 | 21.62 | 31.37 | 2 | 0.0 | 0 | K.ALPAPIEK.T | | 257-264 | CID |
| 544 | 643.819 | 2 | -33.55 | -0.0216 | 23.78 | 57.48 | 9 | 0.0 | 0 | R.EPQVYTLPPSR.E | | 275-285 | CID |
| 539 | 635.94 | 3 | 452.66 | 0.2877 | 23.70 | 49.15 | 27 | 0.0 | 1 | R.EPQVYTLPPSREEMTK.N | | 275-290 | CID |
| 682 | 581.368 | 2 | 85.24 | 0.0496 | 26.98 | 68.52 | 14 | 0.0 | 0 | K.NQVSLTCLVK.G | Carbamidomethyl: 7 | 291-300 | CID |

#### Protein 5: Immunoglobulin heavy constant gamma 2 OS=Homo sapiens GN=IGHG2 PE=1 SV=2

**Accession:** IGHG2_HUMAN
**Database:** SwissProt
**MW [kDa] / pI:** 35.9 / 7.7
**Modification(s):** Carbamidomethyl, Oxidation

**Score:** 789.7
**Seq. Coverage [%]:** 47.5
**No. of Peptides:** 14

| 10 | 20 | 30 | 40 | 50 | 60 | 70 | 80 | 90 | 100 |
|---|---|---|---|---|---|---|---|---|---|
| ASTKGPSVFP | LAPCSRSTSE | STAALGCLVK | DYFPEPVTVS | WNSGALTSGV | HTFPAVLQSS | GLYSLSSVVT | VPSSNFGTQT | YTCNVDHKPS | NTKVDKTVER |
| **110** | **120** | **130** | **140** | **150** | **160** | **170** | **180** | **190** | **200** |
| KCCVECPPCP | APPVAGPSVF | LFPPKPKDTL | MISRTPEVTC | VVVDVSHEDP | EVQFNWYVDG | VEVHNAKTKP | REEQFNSTFR | VVSVLTVVHQ | DWLNGKEYKC |
| **210** | **220** | **230** | **240** | **250** | **260** | **270** | **280** | **290** | **300** |
| KVSNKGLPAP | IEKTISKTKG | QPREPQVYTL | PPSREEMTKN | QVSLTCLVKG | FYPSDISVEW | ESNGQPENNY | KTTPPMLDSD | GSFFLYSKLT | VDKSRWQQGN |
| **310** | **320** | **330** | | | | | | | |
| VFSCSVMHEA | LHNHYTQKSL | SLSPGK | | | | | | | |

##### MS/MS Peptide Matches

| Cmpd. | m/z meas. | z | Δ m/z [ppm] | Δ m/z [Da] | Rt [min] | Score | No. of Cmpds. | Site [%] | P | Sequence | Modification | Range | Type |
|---|---|---|---|---|---|---|---|---|---|---|---|---|---|
| 745 | 644.38 | 2 | 78.61 | 0.0507 | 28.23 | 69.32 | 14 | 0.0 | 0 | K.GPSVFPLAPCSR.S | Carbamidomethyl: 10 | 5-16 | CID |
| 640 | 712.423 | 2 | 90.55 | 0.0645 | 25.97 | 84.0 | 15 | 0.0 | 0 | R.STSESTAALGCLVK.D | Carbamidomethyl: 11 | 17-30 | CID |
| 886 | 760.215 | 4 | 441.36 | 0.3354 | 31.06 | 48.41 | 3 | 0.0 | 1 | R.KCCVECPPCPAPPVAGPSVFLFPPKPK.D | Carbamidomethyl: 2, 3, 6, 9 | 101-127 | CID |
| 964 | 970.476 | 3 | 347.64 | 0.3373 | 32.62 | 78.0 | 7 | 0.0 | 0 | K.CCVECPPCPAPPVAGPSVFLFPPKPK.D | Carbamidomethyl: 1, 2, 5, 8 | 102-127 | CID |
| 456 | 418.225 | 2 | 10.17 | 0.0043 | 21.87 | 47.58 | 9 | 0.0 | 0 | K.DTLMISR.T | | 128-134 | CID |
| 358 | 426.256 | 2 | 88.68 | 0.0378 | 19.90 | 47.25 | 4 | 0.0 | 0 | K.DTLMISR.T | Oxidation: 4 | 128-134 | CID |
| 1140 | 950.29 | 4 | 95.50 | 0.0907 | 36.58 | 44.3 | 7 | 0.0 | 0 | R.TPEVTCVVVDVSHEDPEVQFNWYVDGVEVHNAK.T | Carbamidomethyl: 6 | 135-167 | CID |
| 962 | 599.037 | 3 | 615.59 | 0.3685 | 32.56 | 59.03 | 5 | 0.0 | 0 | R.VVSVLTVVHQDWLNGK.E | | 181-196 | CID |
| 888 | 554.45 | 4 | 264.50 | 0.1466 | 31.07 | 21.12 | 1 | 0.0 | 1 | R.VVSVLTVVHQDWLNGKEYK.C | | 181-199 | CID |
| 544 | 643.819 | 2 | -33.55 | -0.0216 | 23.78 | 57.48 | 9 | 0.0 | 0 | R.EPQVYTLPPSR.E | | 224-234 | CID |
| 539 | 635.94 | 3 | 452.66 | 0.2877 | 23.70 | 49.15 | 27 | 0.0 | 1 | R.EPQVYTLPPSREEMTK.N | | 224-239 | CID |
| 682 | 581.368 | 2 | 85.24 | 0.0496 | 26.98 | 68.52 | 14 | 0.0 | 0 | K.NQVSLTCLVK.G | Carbamidomethyl: 7 | 240-249 | CID |
| 968 | 953.485 | 2 | 36.10 | 0.0344 | 32.71 | 80.66 | 13 | 0.0 | 0 | K.TTPPMLDSDGSFFLYSK.L | | 272-288 | CID |
| 887 | 961.468 | 2 | 20.76 | 0.0200 | 31.07 | 34.87 | 3 | 0.0 | 0 | K.TTPPMLDSDGSFFLYSK.L | Oxidation: 5 | 272-288 | CID |

BRUKER

#### Protein 6: Immunoglobulin heavy constant gamma 4 OS=Homo sapiens GN=IGHG4 PE=1 SV=1

| | | | |
|---|---|---|---|
| **Accession:** | IGHG4_HUMAN | **Score:** | 686.4 |
| **Database:** | SwissProt | **Seq. Coverage [%]:** | 44.6 |
| **MW [kDa] / pI:** | 35.9 / 7.2 | **No. of Peptides:** | 11 |
| **Modification(s):** | Carbamidomethyl, Oxidation | | |

| 10 | 20 | 30 | 40 | 50 | 60 | 70 | 80 | 90 | 100 |
|---|---|---|---|---|---|---|---|---|---|
| ASTKGPSVFP | LAPCSRSTSE | STAALGCLVK | DYFPEPVTVS | WNSGALTSGV | HTFPAVLQSS | GLYSLSSVVT | VPSSSLGTKT | YTCNVDHKPS | NTKVDKRVES |
| **110** | **120** | **130** | **140** | **150** | **160** | **170** | **180** | **190** | **200** |
| KYGPPCPSCP | APEFLGGPSV | FLFPPKPKDT | LMISRTPEVT | CVVVDVSQED | PEVQFNWYVD | GVEVHNAKTK | PREEQFNSTY | RVVSVLTVLH | QDWLNGKEYK |
| **210** | **220** | **230** | **240** | **250** | **260** | **270** | **280** | **290** | **300** |
| CKVSNKGLPS | SIEKTISKAK | GQPREPQVYT | LPPSQEEMTK | NQVSLTCLVK | GFYPSDIAVE | WESNGQPENN | YKTTPPVLDS | DGSFFLYSRL | TVDKSRWQEG |
| **310** | **320** | **330** | | | | | | | |
| NVFSCSVMHE | ALHNHYTQKS | LSLSLGK | | | | | | | |

##### MS/MS Peptide Matches

| Cmpd. | m/z meas. | z | Δ m/z [ppm] | Δ m/z [Da] | Rt [min] | Score | No. of Cmpds. | Site [%] | P | Sequence | Modification | Range | Type |
|---|---|---|---|---|---|---|---|---|---|---|---|---|---|
| 745 | 644.38 | 2 | 78.61 | 0.0507 | 28.23 | 69.32 | 14 | 0.0 | 0 | K.GPSVFPLAPCSR.S | Carbamidomethyl: 10 | 5-16 | CID |
| 640 | 712.423 | 2 | 90.55 | 0.0645 | 25.97 | 84.0 | 15 | 0.0 | 0 | R.STSESTAALGCLVK.D | Carbamidomethyl: 11 | 17-30 | CID |
| 95 | 417.13 | 4 | 436.79 | 0.1821 | 13.91 | 33.14 | 3 | 0.0 | 0 | K.TYTCNVDHKPSNTK.V | Carbamidomethyl: 4 | 80-93 | CID |
| 1091 | 982.246 | 3 | 429.89 | 0.4221 | 35.49 | 90.18 | 7 | 0.0 | 0 | K.YGPPCPSCPAPEFLGGPSVFLFPPKPK.D | Carbamidomethyl: 5, 8 | 102-128 | CID |
| 456 | 418.225 | 2 | 10.17 | 0.0043 | 21.87 | 47.58 | 9 | 0.0 | 0 | K.DTLMISR.T | | 129-135 | CID |
| 358 | 426.256 | 2 | 88.68 | 0.0378 | 19.90 | 47.25 | 4 | 0.0 | 0 | K.DTLMISR.T | Oxidation: 4 | 129-135 | CID |
| 1057 | 603.404 | 3 | 105.50 | 0.0637 | 34.72 | 71.31 | 10 | 0.0 | 0 | R.VVSVLTVLHQDWLNGK.E | | 182-197 | CID |
| 373 | 415.802 | 2 | 162.26 | 0.0675 | 20.16 | 27.18 | 2 | 0.0 | 0 | K.GLPSSIEK.T | | 207-214 | CID |
| 595 | 938.958 | 2 | 4.80 | 0.0045 | 24.87 | 22.53 | 2 | 0.0 | 0 | R.EPQVYTLPPSQEEMTK.N | | 225-240 | CID |

| Cmpd. | m/z meas. | z | Δ m/z [ppm] | Δ m/z [Da] | Rt [min] | Score | No. of Cmpds. | Site [%] | P | Sequence | Modification | Range | Type |
|---|---|---|---|---|---|---|---|---|---|---|---|---|---|
| 682 | 581.368 | 2 | 85.24 | 0.0496 | 26.98 | 68.52 | 14 | 0.0 | 0 | K.NQVSLTCLVK.G | Carbamidomethyl: 7 | 241-250 | CID |
| 893 | 1273.046 | 2 | 374.58 | 0.4767 | 31.19 | 125.42 | 37 | 0.0 | 0 | K.GFYPSDIAVEWESNGQPENNYK.T | | 251-272 | CID |

#### Protein 7: Immunoglobulin lambda constant 3 OS=Homo sapiens GN=IGLC3 PE=1 SV=1

**Accession:** IGLC3_HUMAN
**Database:** SwissProt
**MW [kDa] / pI:** 11.3 / 6.9
**Modification(s):** Carbamidomethyl

**Score:** 624.8
**Seq. Coverage [%]:** 93.4
**No. of Peptides:** 9

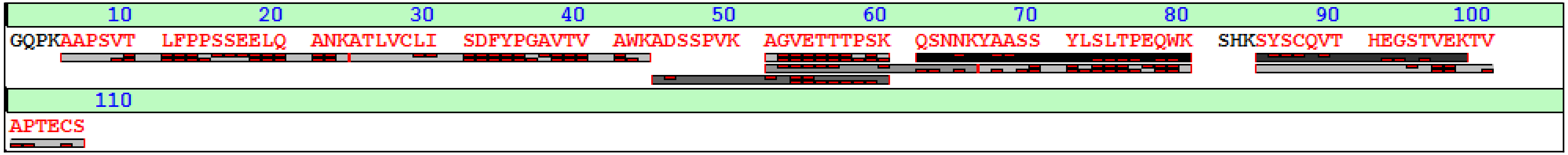


MS/MS Peptide Matches

| Cmpd. | m/z meas. | z | Δ m/z [ppm] | Δ m/z [Da] | Rt [min] | Score | No. of Cmpds. | Site [%] | P | Sequence | Modification | Range | Type |
|---|---|---|---|---|---|---|---|---|---|---|---|---|---|
| 831 | 993.583 | 2 | 70.90 | 0.0704 | 29.90 | 96.73 | 20 | 0.0 | 0 | K.AAPSVTLFPPSSEELQANK.A | | 5-23 | CID |
| 1335 | 1106.568 | 2 | 441.60 | 0.4884 | 40.62 | 98.23 | 3 | 0.0 | 0 | K.ATLVCLISDFYPGAVTVAWK.A | Carbamidomethyl: 5 | 24-43 | CID |
| 266 | 558.975 | 3 | 33.38 | 0.0187 | 17.82 | 69.71 | 2 | 0.0 | 1 | K.ADSSPVKAGVETTTPSK.Q | | 44-60 | CID |
| 145 | 495.787 | 2 | 56.99 | 0.0283 | 15.02 | 79.25 | 18 | 0.0 | 0 | K.AGVETTTPSK.Q | | 51-60 | CID |
| 83 | 521.313 | 3 | 91.32 | 0.0476 | 13.63 | 61.56 | 4 | 0.0 | 1 | K.AGVETTTPSKQSNNK.Y | | 51-65 | CID |
| 794 | 772.686 | 3 | 394.10 | 0.3044 | 29.18 | 19.29 | 1 | 0.0 | 1 | K.QSNNKYAASSYLSLTPEQWK.S | | 61-80 | CID |
| 853 | 872.463 | 2 | 34.33 | 0.0299 | 30.32 | 94.77 | 19 | 0.0 | 0 | K.YAASSYLSLTPEQWK.S | | 66-80 | CID |
| 247 | 571.285 | 3 | 47.41 | 0.0271 | 17.47 | 59.66 | 14 | 0.0 | 0 | K.SYSCQVTHEGSTVEK.T | Carbamidomethyl: 4 | 84-98 | CID |
| 426 | 853.057 | 3 | 14.98 | 0.0128 | 21.20 | 45.58 | 3 | 0.0 | 1 | K.SYSCQVTHEGSTVEKTVAPTECS.- | Carbamidomethyl: 4, 22 | 84-106 | CID |

#### Protein 8: Immunoglobulin lambda-like polypeptide 5 OS=Homo sapiens GN=IGLL5 PE=2 SV=2

| | | | |
|---|---|---|---|
| **Accession:** | IGLL5_HUMAN | **Score:** | 440.8 |
| **Database:** | SwissProt | **Seq. Coverage [%]:** | 42.1 |
| **MW [kDa] / pI:** | 23.0 / 9.1 | **No. of Peptides:** | 7 |
| **Modification(s):** | Carbamidomethyl | | |

| 10 | 20 | 30 | 40 | 50 | 60 | 70 | 80 | 90 | 100 |
|---|---|---|---|---|---|---|---|---|---|
| MRPKTGQVGC | ETPEELGPGP | RQRWPLLLLG | LAMVAHGLLR | PMVAPQSGDP | DPGASVGSSR | SSLRSLWGRL | LLQPSPQRAD | PRCWPRGFWS | EPQSLCYVFG |
| **110** | **120** | **130** | **140** | **150** | **160** | **170** | **180** | **190** | **200** |
| TGTKVTVLGQ | PKANPTVTLF | PPSSEELQAN | KATLVCLISD | FYPGAVTVAW | KADGSPVKAG | VETTKPSKQS | NNKYAASSYL | SLTPEQWKSH | RSYSCQVTHE |
| **210** | **220** | | | | | | | | |
| GSTVEKTVAP | TECS | | | | | | | | |

##### MS/MS Peptide Matches

| Cmpd. | m/z meas. | z | Δ m/z [ppm] | Δ m/z [Da] | Rt [min] | Score | No. of Cmpds. | Site [%] | P | Sequence | Modification | Range | Type |
|---|---|---|---|---|---|---|---|---|---|---|---|---|---|
| 359 | 421.321 | 2 | 143.08 | 0.0603 | 19.93 | 47.61 | 4 | 0.0 | 0 | K.VTVLGQPK.A | | 105-112 | CID |
| 818 | 1022.008 | 2 | -14.96 | -0.0153 | 29.66 | 75.63 | 13 | 0.0 | 0 | K.ANPTVTLFPPSSEELQANK.A | | 113-131 | CID |
| 1335 | 1106.568 | 2 | 441.60 | 0.4884 | 40.62 | 98.23 | 3 | 0.0 | 0 | K.ATLVCLISDFYPGAVTVAWK.A | Carbamidomethyl: 5 | 132-151 | CID |
| 794 | 772.686 | 3 | 394.10 | 0.3044 | 29.18 | 19.29 | 1 | 0.0 | 1 | K.QSNNKYAASSYLSLTPEQWK.S | | 169-188 | CID |
| 853 | 872.463 | 2 | 34.33 | 0.0299 | 30.32 | 94.77 | 19 | 0.0 | 0 | K.YAASSYLSLTPEQWK.S | | 174-188 | CID |
| 247 | 571.285 | 3 | 47.41 | 0.0271 | 17.47 | 59.66 | 14 | 0.0 | 0 | R.SYSCQVTHEGSTVEK.T | Carbamidomethyl: 4 | 192-206 | CID |
| 426 | 853.057 | 3 | 14.98 | 0.0128 | 21.20 | 45.58 | 3 | 0.0 | 1 | R.SYSCQVTHEGSTVEKTVAPTECS.- | Carbamidomethyl: 4, 22 | 192-214 | CID |

##### Protein 9: Immunoglobulin kappa variable 2D-28 OS=Homo sapiens GN=IGKV2D-28 PE=1 SV=2

| | | | |
|---|---|---|---|
| **Accession:** | KVD28_HUMAN | **Score:** | 196.8 |
| **Database:** | SwissProt | **Seq. Coverage [%]:** | 16.7 |
| **MW [kDa] / pI:** | 12.9 / 5.6 | **No. of Peptides:** | 2 |

```
        10         20         30         40         50         60         70         80         90        100
MRLPAQLLGL LMLWVSGSSG DIVMTQSPLS LPVTPGEPAS ISCRSSQSLL HSNGYNYLDW YLQKPGQSPQ LLIYLGSNRA SGVPDRFSGS GSGTDFTLKI
       110        120        130
SRVEAEDVGV YYCMQALQTP
```

MS/MS Peptide Matches

| Cmpd. | m/z meas. | z | Δ m/z [ppm] | Δ m/z [Da] | Rt [min] | Score | No. of Cmpds. | Site [%] | P | Sequence | Modification | Range | Type |
|---|---|---|---|---|---|---|---|---|---|---|---|---|---|
| 601 | 662.682 | 3 | 37.80 | 0.0250 | 25.03 | 80.96 | 3 | 0.0 | 1 | R.ASGVPDRFSGSGSGTDFTLK.I | | 80-99 | CID |
| 555 | 652.313 | 2 | 1.73 | 0.0011 | 24.02 | 115.8 | 3 | 0.0 | 0 | R.FSGSGSGTDFTLK.I | | 87-99 | CID |

##### Protein 10: Immunoglobulin heavy variable 3-30-3 OS=Homo sapiens GN=IGHV3-30-3 PE=3 SV=1

| | | | |
|---|---|---|---|
| **Accession:** | HVC33_HUMAN | **Score:** | 121.6 |
| **Database:** | SwissProt | **Seq. Coverage [%]:** | 15.4 |
| **MW [kDa] / pI:** | 13.0 / 9.1 | **No. of Peptides:** | 2 |
| **Modification(s):** | Carbamidomethyl | | |

```
        10         20         30         40         50         60         70         80         90        100
MEFGLSWVFL VALLRGVQCQ VQLVESGGGV VQPGRSLRLS CAASGFTFSS YAMHWVRQAP GKGLEWVAVI SYDGSNKYYA DSVKGRFTIS RDNSKNTLYL
       110        120
QMNSLRAEDT AVYYCAR
```

MS/MS Peptide Matches

| Cmpd. | m/z meas. | z | Δ m/z [ppm] | Δ m/z [Da] | Rt [min] | Score | No. of Cmpds. | Site [%] | P | Sequence | Modification | Range | Type |
|---|---|---|---|---|---|---|---|---|---|---|---|---|---|
| 211 | 423.264 | 2 | 137.95 | 0.0584 | 16.57 | 31.1 | 5 | 0.0 | 0 | K.YYADSVK.G | | 78-84 | CID |
| 398 | 659.819 | 2 | 43.57 | 0.0287 | 20.62 | 90.5 | 3 | 0.0 | 0 | R.AEDTAVYYCAR.- | Carbamidomethyl: 9 | 107-117 | CID |

#### Protein 11: Immunoglobulin heavy variable 1-69 OS=Homo sapiens GN=IGHV1-69 PE=1 SV=2

**Accession:** HV169_HUMAN
**Database:** SwissProt
**MW [kDa] / pI:** 12.7 / 9.3
**Modification(s):** Carbamidomethyl

**Score:** 80.9
**Seq. Coverage [%]:** 9.4
**No. of Peptides:** 1

```
        10         20         30         40         50         60         70         80         90        100
MDWTWRFLFV VAAATGVQSQ VQLVQSGAEV KKPGSSVKVS CKASGGTFSS YAISWVRQAP GQGLEWMGGI IPIFGTANYA QKFQGRVTIT ADKSTSTAYM
       110        120
ELSSLRSEDT AVYYCAR
```

MS/MS Peptide Matches

| Cmpd. | m/z meas. | z | Δ m/z [ppm] | Δ m/z [Da] | Rt [min] | Score | No. of Cmpds. | Site [%] | P | Sequence | Modification | Range | Type |
|---|---|---|---|---|---|---|---|---|---|---|---|---|---|
| 390 | 667.813 | 2 | 37.87 | 0.0253 | 20.46 | 80.85 | 2 | 0.0 | 0 | R.SEDTAVYYCAR.- | Carbamidomethyl: 9 | 107-117 | CID |

#### Protein 12: Serum albumin OS=Homo sapiens GN=ALB PE=1 SV=2

**Accession:** ALBU_HUMAN
**Database:** SwissProt
**MW [kDa] / pI:** 69.3 / 5.9

**Score:** 79.7
**Seq. Coverage [%]:** 2.1
**No. of Peptides:** 1

| 10 | 20 | 30 | 40 | 50 | 60 | 70 | 80 | 90 | 100 |
|---|---|---|---|---|---|---|---|---|---|
| MKWVTFISLL | FLFSSAYSRG | VFRRDAHKSE | VAHRFKDLGE | ENFKALVLIA | FAQYLQQCPF | EDHVKLVNEV | TEFAKTCVAD | ESAENCDKSL | HTLFGDKLCT |
| **110** | **120** | **130** | **140** | **150** | **160** | **170** | **180** | **190** | **200** |
| VATLRETYGE | MADCCAKQEP | ERNECFLQHK | DDNPNLPRLV | RPEVDVMCTA | FHDNEETFLK | KYLYEIARRH | PYFYAPELLF | FAKRYKAAFT | ECCQAADKAA |
| **210** | **220** | **230** | **240** | **250** | **260** | **270** | **280** | **290** | **300** |
| CLLPKLDELR | DEGKASSAKQ | RLKCASLQKF | GERAFKAWAV | ARLSQRFPKA | EFAEVSKLVT | DLTKVHTECC | HGDLLECADD | RADLAKYICE | NQDSISSKLK |
| **310** | **320** | **330** | **340** | **350** | **360** | **370** | **380** | **390** | **400** |
| ECCEKPLLEK | SHCIAEVEND | EMPADLPSLA | ADFVESKDVC | KNYAEAKDVF | LGMFLYEYAR | RHPDYSVVLL | LRLAKTYETT | LEKCCAADP | HECYAKVFDE |
| **410** | **420** | **430** | **440** | **450** | **460** | **470** | **480** | **490** | **500** |
| FKPLVEEPQN | LIKQNCELFE | QLGEYKFQNA | LLVRYTKKVP | QVSTPTLVEV | SRNLGKVGSK | CCKHPEAKRM | PCAEDYLSVV | LNQLCVLHEK | TPVSDRVTKC |
| **510** | **520** | **530** | **540** | **550** | **560** | **570** | **580** | **590** | **600** |
| CTESLVNRRP | CFSALEVDET | YVPKEFNAET | FTFHADICTL | SEKERQIKKQ | TALVELVKHK | PKATKEQLKA | VMDDFAAFVE | KCCKADDKET | CFAEEGKKLV |
| **610** | | | | | | | | | |
| AASQAALGL | | | | | | | | | |

MS/MS Peptide Matches

| Cmpd. | m/z meas. | z | Δ m/z [ppm] | Δ m/z [Da] | Rt [min] | Score | No. of Cmpds. | Site [%] | P | Sequence | Modification | Range | Type |
|---|---|---|---|---|---|---|---|---|---|---|---|---|---|
| 1297 | 812.471 | 2 | 90.56 | 0.0736 | 39.76 | 79.68 | 2 | 0.0 | 0 | K.DVFLGMFLYEYAR.R | | 348-360 | CID |

#### Protein 13: Immunoglobulin kappa variable 3-20 OS=Homo sapiens GN=IGKV3-20 PE=1 SV=2

| | | | |
|---|---|---|---|
| **Accession:** | KV320_HUMAN | **Score:** | 69.6 |
| **Database:** | SwissProt | **Seq. Coverage [%]:** | 7.8 |
| **MW [kDa] / pI:** | 12.5 / 4.9 | **No. of Peptides:** | 1 |

| 10 | 20 | 30 | 40 | 50 | 60 | 70 | 80 | 90 | 100 |
|---|---|---|---|---|---|---|---|---|---|
| METPAQLLFL | LLLWLPDTTG | EIVLTQSPGT | LSLSPGERAT | LSCRASQSVS | SSYLAWYQQK | PGQAPRLLIY | GASSRATGIP | DRFSGSGSGT | DFTLTISRLE |
| 110 | 120 | | | | | | | | |
| PEDFAVYYCQ | QYGSSP | | | | | | | | |

MS/MS Peptide Matches

| Cmpd. | m/z meas. | z | Δ m/z [ppm] | Δ m/z [Da] | Rt [min] | Score | No. of Cmpds. | Site [%] | P | Sequence | Modification | Range | Type |
|---|---|---|---|---|---|---|---|---|---|---|---|---|---|
| 564 | 490.326 | 2 | 89.35 | 0.0438 | 24.17 | 69.59 | 3 | 0.0 | 0 | R.LLIYGASSR.A | | 67-75 | CID |

#### Protein 14: Immunoglobulin heavy constant alpha 1 OS=Homo sapiens GN=IGHA1 PE=1 SV=2

**Accession:** IGHA1_HUMAN
**Database:** SwissProt
**MW [kDa] / pI:** 37.6 / 6.1
**Modification(s):** Carbamidomethyl

**Score:** 69.0
**Seq. Coverage [%]:** 5.9
**No. of Peptides:** 2

| 10 | 20 | 30 | 40 | 50 | 60 | 70 | 80 | 90 | 100 |
|---|---|---|---|---|---|---|---|---|---|
| ASPTSPKVFP | LSLCSTQPDG | NVVIACLVQG | FFPQEPLSVT | WSESGQGVTA | RNFPPSQDAS | GDLYTTSSQL | TLPATQCLAG | KSVTCHVKHY | TNPSQDVTVP |
| 110 | 120 | 130 | 140 | 150 | 160 | 170 | 180 | 190 | 200 |
| CPVPSTPPTP | SPSTPPTPSP | SCCHPRLSLH | RPALEDLLLG | SEANLTCTLT | GLRDASGVTF | TWTPSSGKSA | VQGPPERDLC | GCYSVSSVLP | GCAEPWNHGK |
| 210 | 220 | 230 | 240 | 250 | 260 | 270 | 280 | 290 | 300 |
| TFTCTAAYPE | SKTPLTATLS | KSGNTFRPEV | HLLPPPSEEL | ALNELVTLTC | LARGFSPKDV | LVRWLQGSQE | LPREKYLTWA | SRQEPSQGTT | TFAVTSILRV |
| 310 | 320 | 330 | 340 | 350 | 360 | | | | |
| AAEDWKKGDT | FSCMVGHEAL | PLAFTQKTID | RLAGKPTHVN | VSVVMAEVDG | TCY | | | | |

MS/MS Peptide Matches

| Cmpd. | m/z meas. | z | Δ m/z [ppm] | Δ m/z [Da] | Rt [min] | Score | No. of Cmpds. | Site [%] | P | Sequence | Modification | Range | Type |
|---|---|---|---|---|---|---|---|---|---|---|---|---|---|
| 403 | 688.323 | 2 | 13.72 | 0.0094 | 20.69 | 51.25 | 1 | 0.0 | 0 | K.TFTCTAAYPESK.T | Carbamidomethyl: 4 | 201-212 | CID |

| Cmpd. | m/z meas. | z | Δ m/z [ppm] | Δ m/z [Da] | Rt [min] | Score | No. of Cmpds. | Site [%] | P | Sequence | Modification | Range | Type |
|---|---|---|---|---|---|---|---|---|---|---|---|---|---|
| 518 | 577.321 | 2 | 30.04 | 0.0173 | 23.20 | 17.78 | 1 | 0.0 | 1 | R.EKYLTWASR.Q | | 274-282 | CID |

#### Protein 15: Immunoglobulin kappa variable 3-11 OS=Homo sapiens GN=IGKV3-11 PE=1 SV=1

**Accession:** KV311_HUMAN
**Database:** SwissProt
**MW [kDa] / pI:** 12.6 / 4.9
**Score:** 66.6
**Seq. Coverage [%]:** 7.8
**No. of Peptides:** 1

| 10 | 20 | 30 | 40 | 50 | 60 | 70 | 80 | 90 | 100 |
|---|---|---|---|---|---|---|---|---|---|
| MEAPAQLLFL | LLLWLPDTTG | EIVLTQSPAT | LSLSPGERAT | LSCRASQSVS | SYLAWYQQKP | GQAPRLLIYD | ASNRATGIPA | RFSGSGSGTD | FTLTISSLEP |
| **110** | **120** | | | | | | | | |
| EDFAVYYCQQ | RSNWP | | | | | | | | |

MS/MS Peptide Matches

| Cmpd. | m/z meas. | z | Δ m/z [ppm] | Δ m/z [Da] | Rt [min] | Score | No. of Cmpds. | Site [%] | P | Sequence | Modification | Range | Type |
|---|---|---|---|---|---|---|---|---|---|---|---|---|---|
| 572 | 532.826 | 2 | 66.85 | 0.0356 | 24.34 | 66.59 | 5 | 0.0 | 0 | R.LLIYDASNR.A | | 66-74 | CID |

#### Protein 16: Immunoglobulin kappa variable 3-15 OS=Homo sapiens GN=IGKV3-15 PE=1 SV=2

**Accession:** KV315_HUMAN
**Database:** SwissProt
**MW [kDa] / pI:** 12.5 / 5.1
**Score:** 65.3
**Seq. Coverage [%]:** 7.8
**No. of Peptides:** 1

| 10 | 20 | 30 | 40 | 50 | 60 | 70 | 80 | 90 | 100 |
|---|---|---|---|---|---|---|---|---|---|
| MEAPAQLLFL | LLLWLPDTTG | EIVMTQSPAT | LSVSPGERAT | LSCRASQSVS | SNLAWYQQKP | GQAPRLLIYG | ASTRATGIPA | RFSGSGSGTE | FTLTISSLQS |
| **110** | **120** | | | | | | | | |
| EDFAVYYCQQ | YNNWP | | | | | | | | |

MS/MS Peptide Matches

| Cmpd. | m/z meas. | z | Δ m/z [ppm] | Δ m/z [Da] | Rt [min] | Score | No. of Cmpds. | Site [%] | P | Sequence | Modification | Range | Type |
|---|---|---|---|---|---|---|---|---|---|---|---|---|---|
| 581 | 497.335 | 2 | 90.46 | 0.0450 | 24.54 | 65.31 | 3 | 0.0 | 0 | R.LLIYGASTR.A | | 66-74 | CID |

#### Protein 17: Immunoglobulin kappa variable 4-1 OS=Homo sapiens GN=IGKV4-1 PE=1 SV=1

| | | | |
|---|---|---|---|
| **Accession:** | KV401_HUMAN | **Score:** | 52.6 |
| **Database:** | SwissProt | **Seq. Coverage [%]:** | 12.4 |
| **MW [kDa] / pI:** | 13.4 / 5.1 | **No. of Peptides:** | 1 |

```
        10         20         30         40         50         60         70         80         90        100
MVLQTQVFIS LLLWISGAYG DIVMTQSPDS LAVSLGERAT INCKSSQSVL YSSNNKNYLA WYQQKPGQPP KLLIYWASTR ESGVPDRFSG SGSGTDFTLT
       110        120        130
ISSLQAEDVA VYYCQQYYST P
```

MS/MS Peptide Matches

| Cmpd. | m/z meas. | z | Δ m/z [ppm] | Δ m/z [Da] | Rt [min] | Score | No. of Cmpds. | Site [%] | P | Sequence | Modification | Range | Type |
|---|---|---|---|---|---|---|---|---|---|---|---|---|---|
| 608 | 606.68 | 3 | 50.62 | 0.0307 | 25.15 | 52.6 | 1 | 0.0 | 0 | K.NYLAWYQQKPGQPPK.L | | 57-71 | CID |

#### Protein 18: Immunoglobulin heavy variable 3-73 OS=Homo sapiens GN=IGHV3-73 PE=3 SV=1

| | | | |
|---|---|---|---|
| **Accession:** | HV373_HUMAN | **Score:** | 40.7 |
| **Database:** | SwissProt | **Seq. Coverage [%]:** | 5.9 |
| **MW [kDa] / pI:** | 12.8 / 9.3 | **No. of Peptides:** | 1 |

| 10 | 20 | 30 | 40 | 50 | 60 | 70 | 80 | 90 | 100 |
|---|---|---|---|---|---|---|---|---|---|
| MEFGLSWVFL | VAILKGVQCE | VQLVESGGGL | VQPGGSLKLS | CAASGFTFSG | SAMHWVRQAS | GKGLEWVGRI | RSKANSYATA | YAASVKGRFT | ISRDDSKNTA |
| 110 | 120 | | | | | | | | |
| YLQMNSLKTE | DTAVYYCTR | | | | | | | | |

MS/MS Peptide Matches

| Cmpd. | m/z meas. | z | Δ m/z [ppm] | Δ m/z [Da] | Rt [min] | Score | No. of Cmpds. | Site [%] | P | Sequence | Modification | Range | Type |
|---|---|---|---|---|---|---|---|---|---|---|---|---|---|
| 588 | 408.736 | 2 | 34.82 | 0.0142 | 24.70 | 40.71 | 3 | 0.0 | 0 | K.GLEWVGR.I | | 63-69 | CID |

#### Protein 19: Immunoglobulin lambda variable 5-45 OS=Homo sapiens GN=IGLV5-45 PE=3 SV=1

**Accession:** LV545_HUMAN
**Database:** SwissProt
**MW [kDa] / pI:** 13.2 / 6.7

**Score:** 40.6
**Seq. Coverage [%]:** 11.4
**No. of Peptides:** 1

| 10 | 20 | 30 | 40 | 50 | 60 | 70 | 80 | 90 | 100 |
|---|---|---|---|---|---|---|---|---|---|
| MAWTPLLLLF | LSHCTGSLSQ | AVLTQPSSLS | ASPGASASLT | CTLCSGINVG | TYRIYWYQQK | PGSPPQYLLR | YKSDSDKQQG | SGVPSRFSGS | KDASANAGIL |
| 110 | 120 | 130 | | | | | | | |
| LISGLQSEDE | ADYYCMIWHS | SAS | | | | | | | |

MS/MS Peptide Matches

| Cmpd. | m/z meas. | z | Δ m/z [ppm] | Δ m/z [Da] | Rt [min] | Score | No. of Cmpds. | Site [%] | P | Sequence | Modification | Range | Type |
|---|---|---|---|---|---|---|---|---|---|---|---|---|---|
| 65 | 483.298 | 3 | 139.53 | 0.0674 | 13.18 | 40.57 | 2 | 0.0 | 1 | K.SDSDKQQGSGVPSR.F | | 73-86 | CID |

BRUKER

#### Project Info

**Name:** Alagesan
**Note:** Nanobooster
**Date:** Mar 5, 2019

#### Sample Info & Protocols

**Name:** Nanobooster - IgG
**Date:** Mar 5, 2019

#### Search Result Info

**Search Result:** IonTrap_Human_Swiss_CID_2019-03-05 17:55:08
**Location:** /Alagesan/Nanobooster - IgG/Ka_IgG Trypsin NoEnrich NB Ace_BA5_01_8768.mgf
**Search Method:** IonTrap_Human_Swiss_CID
**Version:** 1.2
**Search Engine(s):** Mascot, 2.6.0
**Ident. Compound(s):** 0/2084
**Database(s):** SwissProt, SwissProt_2017_04.fasta
**Note:**

#### Results

##### Protein 1: Immunoglobulin kappa constant OS=Homo sapiens GN=IGKC PE=1 SV=2

**Accession:** IGKC_HUMAN
**Database:** SwissProt
**MW [kDa] / pI:** 11.8 / 6.1
**Modification(s):** Carbamidomethyl

**Score:** 1016.0
**Seq. Coverage [%]:** 97.2
**No. of Peptides:** 11

10 20 30 40 50 60 70 80 90 100
RTVAAPSVFI FPPSDEQLKS GTASVVCLLN NFYPREAKVQ WKVDNALQSG NSQESVTEQD SKDSTYSLSS TLTLSKADYE KHKVYACEVT HQGLSSPVTK

110
SFNRGEC

MS/MS Peptide Matches

| Cmpd. | m/z meas. | z | Δ m/z [ppm] | Δ m/z [Da] | Rt [min] | Score | No. of Cmpds. | Site [%] | P | Sequence | Modification | Range | Type |
|---|---|---|---|---|---|---|---|---|---|---|---|---|---|
| 920 | 701.441 | 3 | 85.73 | 0.0601 | 31.64 | 49.51 | 14 | 0.0 | 1 | -.RTVAAPSVFIFPPSDEQLK.S | | 1-19 | CID |
| 1027 | 973.667 | 2 | 153.96 | 0.1499 | 33.99 | 93.83 | 45 | 0.0 | 0 | R.TVAAPSVFIFPPSDEQLK.S | | 2-19 | CID |
| 1112 | 899.617 | 2 | 184.28 | 0.1657 | 35.81 | 77.55 | 29 | 0.0 | 0 | K.SGTASVVCLLNNFYPR.E | Carbamidomethyl: 8 | 20-35 | CID |
| 555 | 893.078 | 3 | -18.87 | -0.0168 | 23.83 | 128.05 | 3 | 0.0 | 1 | K.VQWKVDNALQSGNSQESVTEQDSK.D | | 39-62 | CID |
| 348 | 1068.541 | 2 | 49.60 | 0.0530 | 19.55 | 170.02 | 73 | 0.0 | 0 | K.VDNALQSGNSQESVTEQDSK.D | | 43-62 | CID |
| 718 | 1207.477 | 3 | 195.24 | 0.2357 | 27.56 | 153.68 | 21 | 0.0 | 1 | K.VDNALQSGNSQESVTEQDSKDSTYSLSSTLTLSK.A | | 43-76 | CID |
| 744 | 751.941 | 2 | 77.33 | 0.0581 | 28.07 | 96.92 | 7 | 0.0 | 0 | K.DSTYSLSSTLTLSK.A | | 63-76 | CID |
| 6 | 445.774 | 2 | 116.74 | 0.0520 | 11.31 | 15.82 | 2 | 0.0 | 1 | K.ADYEKHK.V | | 77-83 | CID |
| 338 | 536.321 | 4 | 550.99 | 0.2953 | 19.35 | 67.26 | 8 | 0.0 | 1 | K.HKVYACEVTHQGLSSPVTK.S | Carbamidomethyl: 6 | 82-100 | CID |
| 456 | 626.151 | 3 | 272.39 | 0.1705 | 21.74 | 115.13 | 11 | 0.0 | 0 | K.VYACEVTHQGLSSPVTK.S | Carbamidomethyl: 4 | 84-100 | CID |
| 108 | 435.178 | 2 | -9.54 | -0.0042 | 14.06 | 48.19 | 6 | 0.0 | 1 | K.SFNRGEC.- | Carbamidomethyl: 7 | 101-107 | CID |

##### Protein 2: Immunoglobulin kappa light chain OS=Homo sapiens PE=1 SV=1

| | | | |
|---|---|---|---|
| **Accession:** | IGK_HUMAN | **Score:** | 1015.0 |
| **Database:** | SwissProt | **Seq. Coverage [%]:** | 48.1 |
| **MW [kDa] / pI:** | 23.4 / 6.9 | **No. of Peptides:** | 10 |
| **Modification(s):** | Carbamidomethyl | | |

```
        10         20         30         40         50         60         70         80         90        100
DIQMTQSPST LSASVGDRVT ITCRASQSIN TWLAWYQQKP GKAPKLLMYK ASSLESGVPS RFIGSGSGTE FTLTISSLQP DDFATYYCQQ YNSDSKMFGQ
       110        120        130        140        150        160        170        180        190        200
GTKVEVKGTV AAPSVFIFPP SDEQLKSGTA SVVCLLNNFY PREAKVQWKV DNALQSGNSQ ESVTEQDSKD STYSLSSTLT LSKADYEKHK VYACEVTHQG
       210        220
LSSPVTKSFN RGEC
```

MS/MS Peptide Matches

| Cmpd. | m/z meas. | z | Δ m/z [ppm] | Δ m/z [Da] | Rt [min] | Score | No. of Cmpds. | Site [%] | P | Sequence | Modification | Range | Type |
|---|---|---|---|---|---|---|---|---|---|---|---|---|---|
| 619 | 946.898 | 2 | -59.72 | -0.0566 | 25.16 | 142.4 | 3 | 0.0 | 0 | -.DIQMTQSPSTLSASVGDR.V | | 1-18 | CID |
| 1112 | 899.617 | 2 | 184.28 | 0.1657 | 35.81 | 77.55 | 29 | 0.0 | 0 | K.SGTASVVCLLNNFYPR.E | Carbamidomethyl: 8 | 127-142 | CID |
| 555 | 893.078 | 3 | -18.87 | -0.0168 | 23.83 | 128.05 | 3 | 0.0 | 1 | K.VQWKVDNALQSGNSQESVTEQDSK.D | | 146-169 | CID |
| 348 | 1068.541 | 2 | 49.60 | 0.0530 | 19.55 | 170.02 | 73 | 0.0 | 0 | K.VDNALQSGNSQESVTEQDSK.D | | 150-169 | CID |
| 718 | 1207.477 | 3 | 195.24 | 0.2357 | 27.56 | 153.68 | 21 | 0.0 | 1 | K.VDNALQSGNSQESVTEQDSKDSTYSLSSTLTLS K.A | | 150-183 | CID |
| 744 | 751.941 | 2 | 77.33 | 0.0581 | 28.07 | 96.92 | 7 | 0.0 | 0 | K.DSTYSLSSTLTLSK.A | | 170-183 | CID |
| 6 | 445.774 | 2 | 116.74 | 0.0520 | 11.31 | 15.82 | 2 | 0.0 | 1 | K.ADYEKHK.V | | 184-190 | CID |
| 338 | 536.321 | 4 | 550.99 | 0.2953 | 19.35 | 67.26 | 8 | 0.0 | 1 | K.HKVYACEVTHQGLSSPVTK.S | Carbamidomethyl: 6 | 189-207 | CID |
| 456 | 626.151 | 3 | 272.39 | 0.1705 | 21.74 | 115.13 | 11 | 0.0 | 0 | K.VYACEVTHQGLSSPVTK.S | Carbamidomethyl: 4 | 191-207 | CID |
| 108 | 435.178 | 2 | -9.54 | -0.0042 | 14.06 | 48.19 | 6 | 0.0 | 1 | K.SFNRGEC.- | Carbamidomethyl: 7 | 208-214 | CID |

BRUKER

#### Protein 3: Immunoglobulin heavy constant gamma 1 OS=Homo sapiens GN=IGHG1 PE=1 SV=1

| | | | |
|---|---|---|---|
| **Accession:** | IGHG1_HUMAN | **Score:** | 1006.1 |
| **Database:** | SwissProt | **Seq. Coverage [%]:** | 54.8 |
| **MW [kDa] / pI:** | 36.1 / 8.5 | **No. of Peptides:** | 15 |
| **Modification(s):** | Carbamidomethyl, Oxidation | | |

| 10 | 20 | 30 | 40 | 50 | 60 | 70 | 80 | 90 | 100 |
|---|---|---|---|---|---|---|---|---|---|
| ASTKGPSVFP | LAPSSKSTSG | GTAALGCLVK | DYFPEPVTVS | WNSGALTSGV | HTFPAVLQSS | GLYSLSSVVT | VPSSSLGTQT | YICNVNHKPS | NTKVDKKVEP |
| **110** | **120** | **130** | **140** | **150** | **160** | **170** | **180** | **190** | **200** |
| KSCDKTHTCP | PCPAPELLGG | PSVFLFPPKP | KDTLMISRTP | EVTCVVVDVS | HEDPEVKFNW | YVDGVEVHNA | KTKPREEQYN | STYRVVSVLT | VLHQDWLNGK |
| **210** | **220** | **230** | **240** | **250** | **260** | **270** | **280** | **290** | **300** |
| EYKCKVSNKA | LPAPIEKTIS | KAKGQPREPQ | VYTLPPSRDE | LTKNQVSLTC | LVKGFYPSDI | AVEWESNGQP | ENNYKTTPPV | LDSDGSFFLY | SKLTVDKSRW |
| **310** | **320** | **330** | **340** | | | | | | |
| QQGNVFSCSV | MHEALHNHYT | QKSLSLSPGK | | | | | | | |

##### MS/MS Peptide Matches

| Cmpd. | m/z meas. | z | Δ m/z [ppm] | Δ m/z [Da] | Rt [min] | Score | No. of Cmpds. | Site [%] | P | Sequence | Modification | Range | Type |
|---|---|---|---|---|---|---|---|---|---|---|---|---|---|
| 725 | 593.841 | 2 | 23.64 | 0.0140 | 27.72 | 51.48 | 15 | 0.0 | 0 | K.GPSVFPLAPSSK.S | | 5-16 | CID |
| 625 | 661.39 | 2 | 71.59 | 0.0473 | 25.31 | 127.88 | 17 | 0.0 | 0 | K.STSGGTAALGCLVK.D | Carbamidomethyl: 11 | 17-30 | CID |
| 1013 | 712.209 | 4 | 476.43 | 0.3392 | 33.62 | 45.7 | 11 | 0.0 | 0 | K.THTCPPCPAPELLGGPSVFLFPPKPK.D | Carbamidomethyl: 4, 7 | 106-131 | CID |
| 478 | 418.208 | 2 | -30.48 | -0.0127 | 22.18 | 47.59 | 14 | 0.0 | 0 | K.DTLMISR.T | | 132-138 | CID |
| 364 | 426.255 | 2 | 86.33 | 0.0368 | 19.90 | 47.25 | 8 | 0.0 | 0 | K.DTLMISR.T | Oxidation: 4 | 132-138 | CID |
| 715 | 713.691 | 3 | 14.49 | 0.0103 | 27.52 | 102.58 | 2 | 0.0 | 0 | R.TPEVTCVVVDVSHEDPEVK.F | Carbamidomethyl: 6 | 139-157 | CID |
| 1340 | 950.545 | 4 | 354.29 | 0.3367 | 40.59 | 34.43 | 3 | 0.0 | 1 | R.TPEVTCVVVDVSHEDPEVKFNWYVDGVEVHNAK.T | Carbamidomethyl: 6 | 139-171 | CID |
| 722 | 559.993 | 3 | 96.72 | 0.0542 | 27.65 | 77.22 | 10 | 0.0 | 0 | K.FNWYVDGVEVHNAK.T | | 158-171 | CID |

| Cmpd. | m/z meas. | z | Δ m/z [ppm] | Δ m/z [Da] | Rt [min] | Score | No. of Cmpds. | Site [%] | P | Sequence | Modification | Range | Type |
|---|---|---|---|---|---|---|---|---|---|---|---|---|---|
| 1061 | 603.708 | 3 | 609.36 | 0.3677 | 34.73 | 61.24 | 10 | 0.0 | 0 | R.VVSVLTVLHQDWLNGK.E | | 185-200 | CID |
| 447 | 419.81 | 2 | 130.37 | 0.0547 | 21.48 | 23.37 | 4 | 0.0 | 0 | K.ALPAPIEK.T | | 210-217 | CID |
| 541 | 643.818 | 2 | -35.11 | -0.0226 | 23.55 | 59.61 | 9 | 0.0 | 0 | R.EPQVYTLPPSR.D | | 228-238 | CID |
| 592 | 625.036 | 3 | 65.75 | 0.0411 | 24.61 | 31.97 | 11 | 0.0 | 1 | R.EPQVYTLPPSRDELTK.N | | 228-243 | CID |
| 682 | 581.356 | 2 | 64.60 | 0.0376 | 26.78 | 74.13 | 13 | 0.0 | 0 | K.NQVSLTCLVK.G | Carbamidomethyl: 7 | 244-253 | CID |
| 891 | 1272.495 | 2 | -58.41 | -0.0743 | 31.07 | 130.22 | 46 | 0.0 | 0 | K.GFYPSDIAVEWESNGQPENNYK.T | | 254-275 | CID |
| 962 | 937.911 | 2 | 476.23 | 0.4465 | 32.49 | 91.44 | 15 | 0.0 | 0 | K.TTPPVLDSDGSFFLYSK.L | | 276-292 | CID |

#### Protein 4: Immunoglobulin heavy constant gamma 2 OS=Homo sapiens GN=IGHG2 PE=1 SV=2

**Accession:** IGHG2_HUMAN
**Database:** SwissProt
**MW [kDa] / pI:** 35.9 / 7.7
**Modification(s):** Carbamidomethyl, Oxidation

**Score:** 846.9
**Seq. Coverage [%]:** 49.1
**No. of Peptides:** 14

| 10 | 20 | 30 | 40 | 50 | 60 | 70 | 80 | 90 | 100 |
|---|---|---|---|---|---|---|---|---|---|
| ASTKGPSVFP | LAPCSRSTSE | STAALGCLVK | DYFPEPVTVS | WNSGALTSGV | HTFPAVLQSS | GLYSLSSVVT | VPSSNFGTQT | YTCNVDHKPS | NTKVDKTVER |
| **110** | **120** | **130** | **140** | **150** | **160** | **170** | **180** | **190** | **200** |
| KCCVECPPCP | APPVAGPSVF | LFPPKPKDTL | MISRTPEVTC | VVVDVSHEDP | EVQFNWYVDG | VEVHNAKTKP | REEQFNSTFR | VVSVLTVVHQ | DWLNGKEYKC |
| **210** | **220** | **230** | **240** | **250** | **260** | **270** | **280** | **290** | **300** |
| KVSNKGLPAP | IEKTISKTKG | QPREPQVYTL | PPSREEMTKN | QVSLTCLVKG | FYPSDISVEW | ESNGQPENNY | KTTPPMLDSD | GSFFLYSKLT | VDKSRWQQGN |
| **310** | **320** | **330** | | | | | | | |
| VFSCSVMHEA | LHNHYTQKSL | SLSPGK | | | | | | | |

MS/MS Peptide Matches

| Cmpd. | m/z meas. | z | Δ m/z [ppm] | Δ m/z [Da] | Rt [min] | Score | No. of Cmpds. | Site [%] | P | Sequence | Modification | Range | Type |
|---|---|---|---|---|---|---|---|---|---|---|---|---|---|
| 758 | 644.367 | 2 | 58.44 | 0.0377 | 28.38 | 99.42 | 14 | 0.0 | 0 | K.GPSVFPLAPCSR.S | Carbamidomethyl: 10 | 5-16 | CID |
| 645 | 712.42 | 2 | 86.34 | 0.0615 | 25.82 | 94.97 | 15 | 0.0 | 0 | R.STSESTAALGCLVK.D | Carbamidomethyl: 11 | 17-30 | CID |
| 886 | 760.222 | 4 | 450.57 | 0.3424 | 30.96 | 42.2 | 3 | 0.0 | 1 | R.KCCVECPPCPAPPVAGPSVFLFPPKPK.D | Carbamidomethyl: 2, 3, 6, 9 | 101-127 | CID |
| 965 | 970.449 | 3 | 319.81 | 0.3103 | 32.57 | 87.52 | 8 | 0.0 | 0 | K.CCVECPPCPAPPVAGPSVFLFPPKPK.D | Carbamidomethyl: 1, 2, 5, 8 | 102-127 | CID |
| 478 | 418.208 | 2 | -30.48 | -0.0127 | 22.18 | 47.59 | 14 | 0.0 | 0 | K.DTLMISR.T | | 128-134 | CID |
| 364 | 426.255 | 2 | 86.33 | 0.0368 | 19.90 | 47.25 | 8 | 0.0 | 0 | K.DTLMISR.T | Oxidation: 4 | 128-134 | CID |
| 1340 | 950.545 | 4 | 363.87 | 0.3457 | 40.59 | 34.43 | 3 | 0.0 | 0 | R.TPEVTCVVVDVSHEDPEVQFNWYVDGVEVHNAK.T | Carbamidomethyl: 6 | 135-167 | CID |
| 964 | 599.033 | 3 | 608.91 | 0.3645 | 32.55 | 48.76 | 5 | 0.0 | 0 | R.VVSVLTVVHQDWLNGK.E | | 181-196 | CID |
| 468 | 412.767 | 2 | 47.36 | 0.0195 | 21.96 | 20.16 | 1 | 0.0 | 0 | K.GLPAPIEK.T | | 206-213 | CID |
| 541 | 643.818 | 2 | -35.11 | -0.0226 | 23.55 | 59.61 | 9 | 0.0 | 0 | R.EPQVYTLPPSR.E | | 224-234 | CID |
| 514 | 635.643 | 3 | -14.57 | -0.0093 | 22.91 | 42.28 | 25 | 0.0 | 1 | R.EPQVYTLPPSREEMTK.N | | 224-239 | CID |
| 682 | 581.356 | 2 | 64.60 | 0.0376 | 26.78 | 74.13 | 13 | 0.0 | 0 | K.NQVSLTCLVK.G | Carbamidomethyl: 7 | 240-249 | CID |
| 973 | 953.511 | 2 | 63.37 | 0.0604 | 32.76 | 97.93 | 11 | 0.0 | 0 | K.TTPPMLDSDGSFFLYSK.L | | 272-288 | CID |
| 890 | 961.431 | 2 | -17.73 | -0.0170 | 31.04 | 50.67 | 3 | 0.0 | 0 | K.TTPPMLDSDGSFFLYSK.L | Oxidation: 5 | 272-288 | CID |

#### Protein 5: Immunoglobulin heavy constant gamma 3 OS=Homo sapiens GN=IGHG3 PE=1 SV=2

**Accession:** IGHG3_HUMAN
**Database:** SwissProt
**MW [kDa] / pI:** 41.3 / 8.2
**Modification(s):** Carbamidomethyl, Oxidation

**Score:** 846.2
**Seq. Coverage [%]:** 36.3
**No. of Peptides:** 13

| 10 | 20 | 30 | 40 | 50 | 60 | 70 | 80 | 90 | 100 |
|---|---|---|---|---|---|---|---|---|---|
| ASTKGPSVFP | LAPCSRSTSG | GTAALGCLVK | DYFPEPVTVS | WNSGALTSGV | HTFPAVLQSS | GLYSLSSVVT | VPSSSLGTQT | YTCNVNHKPS | NTKVDKRVEL |
| **110** | **120** | **130** | **140** | **150** | **160** | **170** | **180** | **190** | **200** |
| KTPLGDTTHT | CPRCPEPKSC | DTPPPCPRCP | EPKSCDTPPP | CPRCPEPKSC | DTPPPCPRCP | APELLGGPSV | FLFPPKPKDT | LMISRTPEVT | CVVVDVSHED |
| **210** | **220** | **230** | **240** | **250** | **260** | **270** | **280** | **290** | **300** |
| PEVQFKWYVD | GVEVHNAKTK | PREEQYNSTF | RVVSVLTVLH | QDWLNGKEYK | CKVSNKALPA | PIEKTISKTK | GQPREPQVYT | LPPSREEMTK | NQVSLTCLVK |
| **310** | **320** | **330** | **340** | **350** | **360** | **370** | **380** | | |
| GFYPSDIAVE | WESSGQPENN | YNTTPPMLDS | DGSFFLYSKL | TVDKSRWQQG | NIFSCSVMHE | ALHNRFTQKS | LSLSPGK | | |

##### MS/MS Peptide Matches

| Cmpd. | m/z meas. | z | Δ m/z [ppm] | Δ m/z [Da] | Rt [min] | Score | No. of Cmpds. | Site [%] | P | Sequence | Modification | Range | Type |
|---|---|---|---|---|---|---|---|---|---|---|---|---|---|
| 758 | 644.367 | 2 | 58.44 | 0.0377 | 28.38 | 99.42 | 14 | 0.0 | 0 | K.GPSVFPLAPCSR.S | Carbamidomethyl: 10 | 5-16 | CID |
| 625 | 661.39 | 2 | 71.59 | 0.0473 | 25.31 | 127.88 | 17 | 0.0 | 0 | R.STSGGTAALGCLVK.D | Carbamidomethyl: 11 | 17-30 | CID |
| 231 | 452.545 | 3 | -12.36 | -0.0056 | 16.93 | 59.45 | 3 | 0.0 | 0 | K.TPLGDTTHTCPR.C | Carbamidomethyl: 10 | 102-113 | CID |
| 195 | 593.768 | 2 | 25.91 | 0.0154 | 16.10 | 63.76 | 11 | 0.0 | 0 | K.SCDTPPPCPR.C | Carbamidomethyl: 2, 8 | 119-128 | CID |
| 1056 | 717.799 | 3 | 100.02 | 0.0718 | 34.64 | 101.14 | 3 | 0.0 | 0 | R.CPAPELLGGPSVFLFPPKPK.D | Carbamidomethyl: 1 | 159-178 | CID |
| 478 | 418.208 | 2 | -30.48 | -0.0127 | 22.18 | 47.59 | 14 | 0.0 | 0 | K.DTLMISR.T | | 179-185 | CID |
| 364 | 426.255 | 2 | 86.33 | 0.0368 | 19.90 | 47.25 | 8 | 0.0 | 0 | K.DTLMISR.T | Oxidation: 4 | 179-185 | CID |
| 556 | 472.956 | 3 | 114.76 | 0.0543 | 23.87 | 39.11 | 3 | 0.0 | 0 | K.WYVDGVEVHNAK.T | | 207-218 | CID |
| 1061 | 603.708 | 3 | 609.36 | 0.3677 | 34.73 | 61.24 | 10 | 0.0 | 0 | R.VVSVLTVLHQDWLNGK.E | | 232-247 | CID |
| 447 | 419.81 | 2 | 130.37 | 0.0547 | 21.48 | 23.37 | 4 | 0.0 | 0 | K.ALPAPIEK.T | | 257-264 | CID |
| 541 | 643.818 | 2 | -35.11 | -0.0226 | 23.55 | 59.61 | 9 | 0.0 | 0 | R.EPQVYTLPPSR.E | | 275-285 | CID |
| 514 | 635.643 | 3 | -14.57 | -0.0093 | 22.91 | 42.28 | 25 | 0.0 | 1 | R.EPQVYTLPPSREEMTK.N | | 275-290 | CID |
| 682 | 581.356 | 2 | 64.60 | 0.0376 | 26.78 | 74.13 | 13 | 0.0 | 0 | K.NQVSLTCLVK.G | Carbamidomethyl: 7 | 291-300 | CID |

##### Protein 6: Immunoglobulin heavy constant gamma 4 OS=Homo sapiens GN=IGHG4 PE=1 SV=1

| | | | |
|---|---|---|---|
| **Accession:** | IGHG4_HUMAN | **Score:** | 689.6 |
| **Database:** | SwissProt | **Seq. Coverage [%]:** | 42.2 |
| **MW [kDa] / pI:** | 35.9 / 7.2 | **No. of Peptides:** | 10 |
| **Modification(s):** | Carbamidomethyl, Oxidation | | |

| 10 | 20 | 30 | 40 | 50 | 60 | 70 | 80 | 90 | 100 |
|---|---|---|---|---|---|---|---|---|---|
| ASTKGPSVFP | LAPCSRSTSE | STAALGCLVK | DYFPEPVTVS | WNSGALTSGV | HTFPAVLQSS | GLYSLSSVVT | VPSSSLGTKT | YTCNVDHKPS | NTKVDKRVES |
| **110** | **120** | **130** | **140** | **150** | **160** | **170** | **180** | **190** | **200** |
| KYGPPCPSCP | APEFLGGPSV | FLFPPKPKDT | LMISRTPEVT | CVVVDVSQED | PEVQFNWYVD | GVEVHNAKTK | PREEQFNSTY | RVVSVLTVLH | QDWLNGKEYK |
| **210** | **220** | **230** | **240** | **250** | **260** | **270** | **280** | **290** | **300** |
| CKVSNKGLPS | SIEKTISKAK | GQPREPQVYT | LPPSQEEMTK | NQVSLTCLVK | GFYPSDIAVE | WESNGQPENN | YKTTPPVLDS | DGSFFLYSRL | TVDKSRWQEG |
| **310** | **320** | **330** | | | | | | | |
| NVFSCSVMHE | ALHNHYTQKS | LSLSLGK | | | | | | | |

MS/MS Peptide Matches

| Cmpd. | m/z meas. | z | Δ m/z [ppm] | Δ m/z [Da] | Rt [min] | Score | No. of Cmpds. | Site [%] | P | Sequence | Modification | Range | Type |
|---|---|---|---|---|---|---|---|---|---|---|---|---|---|
| 758 | 644.367 | 2 | 58.44 | 0.0377 | 28.38 | 99.42 | 14 | 0.0 | 0 | K.GPSVFPLAPCSR.S | Carbamidomethyl: 10 | 5-16 | CID |
| 645 | 712.42 | 2 | 86.34 | 0.0615 | 25.82 | 94.97 | 15 | 0.0 | 0 | R.STSESTAALGCLVK.D | Carbamidomethyl: 11 | 17-30 | CID |
| 107 | 416.968 | 4 | 48.25 | 0.0201 | 14.06 | 30.26 | 3 | 0.0 | 0 | K.TYTCNVDHKPSNTK.V | Carbamidomethyl: 4 | 80-93 | CID |
| 1090 | 982.213 | 3 | 396.28 | 0.3891 | 35.33 | 82.02 | 4 | 0.0 | 0 | K.YGPPCPSCPAPEFLGGPSVFLFPPKPK.D | Carbamidomethyl: 5, 8 | 102-128 | CID |
| 478 | 418.208 | 2 | -30.48 | -0.0127 | 22.18 | 47.59 | 14 | 0.0 | 0 | K.DTLMISR.T | | 129-135 | CID |
| 364 | 426.255 | 2 | 86.33 | 0.0368 | 19.90 | 47.25 | 8 | 0.0 | 0 | K.DTLMISR.T | Oxidation: 4 | 129-135 | CID |
| 1061 | 603.708 | 3 | 609.36 | 0.3677 | 34.73 | 61.24 | 10 | 0.0 | 0 | R.VVSVLTVLHQDWLNGK.E | | 182-197 | CID |
| 603 | 938.938 | 2 | -16.50 | -0.0155 | 24.86 | 22.47 | 1 | 0.0 | 0 | R.EPQVYTLPPSQEEMTK.N | | 225-240 | CID |
| 682 | 581.356 | 2 | 64.60 | 0.0376 | 26.78 | 74.13 | 13 | 0.0 | 0 | K.NQVSLTCLVK.G | Carbamidomethyl: 7 | 241-250 | CID |

| Cmpd. | m/z meas. | z | Δ m/z [ppm] | Δ m/z [Da] | Rt [min] | Score | No. of Cmpds. | Site [%] | P | Sequence | Modification | Range | Type |
|---|---|---|---|---|---|---|---|---|---|---|---|---|---|
| 891 | 1272.495 | 2 | -58.41 | -0.0743 | 31.07 | 130.22 | 46 | 0.0 | 0 | K.GFYPSDIAVEWESNGQPENNYK.T | | 251-272 | CID |

#### Protein 7: Immunoglobulin lambda constant 3 OS=Homo sapiens GN=IGLC3 PE=1 SV=1

**Accession:** IGLC3_HUMAN
**Database:** SwissProt
**MW [kDa] / pI:** 11.3 / 6.9
**Modification(s):** Carbamidomethyl

**Score:** 615.8
**Seq. Coverage [%]:** 93.4
**No. of Peptides:** 8

```
10         20         30         40         50         60         70         80         90         100
GQPKAAPSVT LFPPSSEELQ ANKATLVCLI SDFYPGAVTV AWKADSSPVK AGVETTTPSK QSNNKYAASS YLSLTPEQWK SHKSYSCQVT HEGSTVEKTV
110
APTECS
```

MS/MS Peptide Matches

| Cmpd. | m/z meas. | z | Δ m/z [ppm] | Δ m/z [Da] | Rt [min] | Score | No. of Cmpds. | Site [%] | P | Sequence | Modification | Range | Type |
|---|---|---|---|---|---|---|---|---|---|---|---|---|---|
| 830 | 993.596 | 2 | 83.98 | 0.0834 | 29.76 | 87.6 | 18 | 0.0 | 0 | K.AAPSVTLFPPSSEELQANK.A | | 5-23 | CID |
| 1339 | 1106.091 | 2 | 10.35 | 0.0114 | 40.58 | 120.32 | 3 | 0.0 | 0 | K.ATLVCLISDFYPGAVTVAWK.A | Carbamidomethyl: 5 | 24-43 | CID |
| 270 | 558.97 | 3 | 24.44 | 0.0137 | 17.76 | 50.84 | 2 | 0.0 | 1 | K.ADSSPVKAGVETTTPSK.Q | | 44-60 | CID |
| 150 | 495.791 | 2 | 65.06 | 0.0323 | 15.02 | 73.02 | 17 | 0.0 | 0 | K.AGVETTTPSK.Q | | 51-60 | CID |
| 85 | 521.321 | 3 | 106.67 | 0.0556 | 13.50 | 64.7 | 3 | 0.0 | 1 | K.AGVETTTPSKQSNNK.Y | | 51-65 | CID |
| 879 | 872.563 | 2 | 148.95 | 0.1299 | 30.77 | 92.78 | 19 | 0.0 | 0 | K.YAASSYLSLTPEQWK.S | | 66-80 | CID |
| 224 | 571.445 | 3 | 327.50 | 0.1871 | 16.74 | 67.62 | 14 | 0.0 | 0 | K.SYSCQVTHEGSTVEK.T | Carbamidomethyl: 4 | 84-98 | CID |
| 435 | 853.058 | 3 | 16.15 | 0.0138 | 21.24 | 58.94 | 3 | 0.0 | 1 | K.SYSCQVTHEGSTVEKTVAPTECS.- | Carbamidomethyl: 4, 22 | 84-106 | CID |

#### Protein 8: Immunoglobulin lambda-like polypeptide 5 OS=Homo sapiens GN=IGLL5 PE=2 SV=2

**Accession:** IGLL5_HUMAN
**Database:** SwissProt
**MW [kDa] / pI:** 23.0 / 9.1
**Modification(s):** Carbamidomethyl

**Score:** 504.8
**Seq. Coverage [%]:** 43.9
**No. of Peptides:** 6

| 10 | 20 | 30 | 40 | 50 | 60 | 70 | 80 | 90 | 100 |
|---|---|---|---|---|---|---|---|---|---|
| MRPKTGQVGC | ETPEELGPGP | RQRWPLLLLG | LAMVAHGLLR | PMVAPQSGDP | DPGASVGSSR | SSLRSLWGRL | LLQPSPQRAD | PRCWPRGFWS | EPQSLCYVFG |
| **110** | **120** | **130** | **140** | **150** | **160** | **170** | **180** | **190** | **200** |
| TGTKVTVLGQ | PKANPTVTLF | PPSSEELQAN | KATLVCLISD | FYPGAVTVAW | KADGSPVKAG | VETTKPSKQS | NNKYAASSYL | SLTPEQWKSH | RSYSCQVTHE |
| **210** | **220** | | | | | | | | |
| GSTVEKTVAP | TECS | | | | | | | | |

##### MS/MS Peptide Matches

| Cmpd. | m/z meas. | z | Δ m/z [ppm] | Δ m/z [Da] | Rt [min] | Score | No. of Cmpds. | Site [%] | P | Sequence | Modification | Range | Type |
|---|---|---|---|---|---|---|---|---|---|---|---|---|---|
| 356 | 421.309 | 2 | 114.59 | 0.0483 | 19.75 | 57.51 | 6 | 0.0 | 0 | K.VTVLGQPK.A | | 105-112 | CID |
| 836 | 1022.007 | 2 | -15.94 | -0.0163 | 29.85 | 90.25 | 16 | 0.0 | 0 | K.ANPTVTLFPPSSEELQANK.A | | 113-131 | CID |
| 1339 | 1106.091 | 2 | 10.35 | 0.0114 | 40.58 | 120.32 | 3 | 0.0 | 0 | K.ATLVCLISDFYPGAVTVAWK.A | Carbamidomethyl: 5 | 132-151 | CID |
| 879 | 872.563 | 2 | 148.95 | 0.1299 | 30.77 | 92.78 | 19 | 0.0 | 0 | K.YAASSYLSLTPEQWK.S | | 174-188 | CID |
| 224 | 571.445 | 3 | 327.50 | 0.1871 | 16.74 | 67.62 | 14 | 0.0 | 0 | R.SYSCQVTHEGSTVEK.T | Carbamidomethyl: 4 | 192-206 | CID |
| 435 | 853.058 | 3 | 16.15 | 0.0138 | 21.24 | 58.94 | 3 | 0.0 | 1 | R.SYSCQVTHEGSTVEKTVAPTECS.- | Carbamidomethyl: 4, 22 | 192-214 | CID |

#### Protein 9: Immunoglobulin kappa variable 2D-28 OS=Homo sapiens GN=IGKV2D-28 PE=1 SV=2

**Accession:** KVD28_HUMAN
**Database:** SwissProt
**MW [kDa] / pI:** 12.9 / 5.6

**Score:** 160.1
**Seq. Coverage [%]:** 16.7
**No. of Peptides:** 2

| 10 | 20 | 30 | 40 | 50 | 60 | 70 | 80 | 90 | 100 |
|---|---|---|---|---|---|---|---|---|---|
| MRLPAQLLGL | LMLWVSGSSG | DIVMTQSPLS | LPVTPGEPAS | ISCRSSQSLL | HSNGYNYLDW | YLQKPGQSPQ | LLIYLGSNRA | SGVPDRFSGS | GSGTDFTLKI |
| 110 | 120 | 130 | | | | | | | |
| SRVEAEDVGV | YYCMQALQTP | | | | | | | | |

MS/MS Peptide Matches

| Cmpd. | m/z meas. | z | Δ m/z [ppm] | Δ m/z [Da] | Rt [min] | Score | No. of Cmpds. | Site [%] | P | Sequence | Modification | Range | Type |
|---|---|---|---|---|---|---|---|---|---|---|---|---|---|
| 614 | 662.682 | 3 | 37.80 | 0.0250 | 25.09 | 60.16 | 3 | 0.0 | 1 | R.ASGVPDRFSGSGSGTDFTLK.I | | 80-99 | CID |
| 561 | 652.287 | 2 | -38.13 | -0.0249 | 23.95 | 99.91 | 3 | 0.0 | 0 | R.FSGSGSGTDFTLK.I | | 87-99 | CID |

#### Protein 10: Serum albumin OS=Homo sapiens GN=ALB PE=1 SV=2

**Accession:** ALBU_HUMAN
**Database:** SwissProt
**MW [kDa] / pI:** 69.3 / 5.9
**Modification(s):** Carbamidomethyl

**Score:** 124.0
**Seq. Coverage [%]:** 6.6
**No. of Peptides:** 2

| 10 | 20 | 30 | 40 | 50 | 60 | 70 | 80 | 90 | 100 |
|---|---|---|---|---|---|---|---|---|---|
| MKWVTFISLL | FLFSSAYSRG | VFRRDAHKSE | VAHRFKDLGE | ENFKALVLIA | FAQYLQQCPF | EDHVKLVNEV | TEFAKTCVAD | ESAENCDKSL | HTLFGDKLCT |
| **110** | **120** | **130** | **140** | **150** | **160** | **170** | **180** | **190** | **200** |
| VATLRETYGE | MADCCAKQEP | ERNECFLQHK | DDNPNLPRLV | RPEVDVMCTA | FHDNEETFLK | KYLYEIARRH | PYFYAPELLF | FAKRYKAAFT | ECCQAADKAA |
| **210** | **220** | **230** | **240** | **250** | **260** | **270** | **280** | **290** | **300** |
| CLLPKLDELR | DEGKASSAKQ | RLKCASLQKF | GERAFKAWAV | ARLSQRFPKA | EFAEVSKLVT | DLTKVHTECC | HGDLLECADD | RADLAKYICE | NQDSISSKLK |
| **310** | **320** | **330** | **340** | **350** | **360** | **370** | **380** | **390** | **400** |
| ECCEKPLLEK | SHCIAEVEND | EMPADLPSLA | ADFVESKDVC | KNYAEAKDVF | LGMFLYEYAR | RHPDYSVVLL | LRLAKTYETT | LEKCCAAADP | HECYAKVFDE |
| **410** | **420** | **430** | **440** | **450** | **460** | **470** | **480** | **490** | **500** |
| FKPLVEEPQN | LIKQNCELFE | QLGEYKFQNA | LLVRYTKKVP | QVSTPTLVEV | SRNLGKVGSK | CCKHPEAKRM | PCAEDYLSVV | LNQLCVLHEK | TPVSDRVTKC |
| **510** | **520** | **530** | **540** | **550** | **560** | **570** | **580** | **590** | **600** |
| CTESLVNRRP | CFSALEVDET | YVPKEFNAET | FTFHADICTL | SEKERQIKKQ | TALVELVKHK | PKATKEQLKA | VMDDFAAFVE | KCCKADDKET | CFAEEGKKLV |
| **610** | | | | | | | | | |
| AASQAALGL | | | | | | | | | |

MS/MS Peptide Matches

| Cmpd. | m/z meas. | z | Δ m/z [ppm] | Δ m/z [Da] | Rt [min] | Score | No. of Cmpds. | Site [%] | P | Sequence | Modification | Range | Type |
|---|---|---|---|---|---|---|---|---|---|---|---|---|---|
| 1171 | 992.114 | 3 | -5.72 | -0.0057 | 37.07 | 36.94 | 3 | 0.0 | 0 | K.SHCIAEVENDEMPADLPSLAADFVESK.D | Carbamidomethyl: 3 | 311-337 | CID |
| 1300 | 812.461 | 2 | 78.25 | 0.0636 | 39.70 | 87.04 | 3 | 0.0 | 0 | K.DVFLGMFLYEYAR.R | | 348-360 | CID |

#### Protein 11: Alpha-1-antitrypsin OS=Homo sapiens GN=SERPINA1 PE=1 SV=3

| | | | |
|---|---|---|---|
| **Accession:** | A1AT_HUMAN | **Score:** | 114.4 |
| **Database:** | SwissProt | **Seq. Coverage [%]:** | 13.6 |
| **MW [kDa] / pI:** | 46.7 / 5.4 | **No. of Peptides:** | 3 |

| 10 | 20 | 30 | 40 | 50 | 60 | 70 | 80 | 90 | 100 |
|---|---|---|---|---|---|---|---|---|---|
| MPSSVSWGIL | LLAGLCCLVP | VSLAEDPQGD | AAQKTDTSHH | DQDHPTFNKI | TPNLAEFAFS | LYRQLAHQSN | STNIFFSPVS | IATAFAMLSL | GTKADTHDEI |
| 110 | 120 | 130 | 140 | 150 | 160 | 170 | 180 | 190 | 200 |
| LEGLNFNLTE | IPEAQIHEGF | QELLRTLNQP | DSQLQLTTGN | GLFLSEGLKL | VDKFLEDVKK | LYHSEAFTVN | FGDTEEAKKQ | INDYVEKGTQ | GKIVDLVKEL |
| 210 | 220 | 230 | 240 | 250 | 260 | 270 | 280 | 290 | 300 |
| DRDTVFALVN | YIFFKGKWER | PFEVKDTEEE | DFHVDQVTTV | KVPMMKRLGM | FNIQHCKKLS | SWVLLMKYLG | NATAIFFLPD | EGKLQHLENE | LTHDIITKFL |
| 310 | 320 | 330 | 340 | 350 | 360 | 370 | 380 | 390 | 400 |
| ENEDRRSASL | HLPKLSITGT | YDLKSVLGQL | GITKVFSNGA | DLSGVTEEAP | LKLSKAVHKA | VLTIDEKGTE | AAGAMFLEAI | PMSIPPEVKF | NKPFVFLMIE |
| 410 | 420 | | | | | | | | |
| QNTKSPLFMG | KVVNPTQK | | | | | | | | |

MS/MS Peptide Matches

| Cmpd. | m/z meas. | z | Δ m/z [ppm] | Δ m/z [Da] | Rt [min] | Score | No. of Cmpds. | Site [%] | P | Sequence | Modification | Range | Type |
|---|---|---|---|---|---|---|---|---|---|---|---|---|---|
| 1407 | 697.355 | 3 | -21.63 | -0.0151 | 41.92 | 58.04 | 1 | 0.0 | 1 | K.ELDRDTVFALVNYIFFK.G | | 199-215 | CID |
| 1633 | 508.324 | 2 | 25.68 | 0.0131 | 47.48 | 27.38 | 1 | 0.0 | 0 | K.SVLGQLGITK.V | | 325-334 | CID |
| 1235 | 1043.884 | 3 | 323.15 | 0.3372 | 38.32 | 28.93 | 3 | 0.0 | 1 | K.AVLTIDEKGTEAAGAMFLEAIPMSIPPEVK.F | | 360-389 | CID |

#### Protein 12: Immunoglobulin heavy variable 3-30-3 OS=Homo sapiens GN=IGHV3-30-3 PE=3 SV=1

**Accession:** HVC33_HUMAN
**Database:** SwissProt
**MW [kDa] / pI:** 13.0 / 9.1
**Modification(s):** Carbamidomethyl

**Score:** 108.4
**Seq. Coverage [%]:** 15.4
**No. of Peptides:** 2

| 10 | 20 | 30 | 40 | 50 | 60 | 70 | 80 | 90 | 100 |
|---|---|---|---|---|---|---|---|---|---|
| MEFGLSWVFL | VALLRGVQCQ | VQLVESGGGV | VQPGRSLRLS | CAASGFTFSS | YAMHWVRQAP | GKGLEWVAVI | SYDGSNKYYA | DSVKGRFTIS | RDNSKNTLYL |
| 110 | 120 | | | | | | | | |
| QMNSLRAEDT | AVYYCAR | | | | | | | | |

MS/MS Peptide Matches

| Cmpd. | m/z meas. | z | Δ m/z [ppm] | Δ m/z [Da] | Rt [min] | Score | No. of Cmpds. | Site [%] | P | Sequence | Modification | Range | Type |
|---|---|---|---|---|---|---|---|---|---|---|---|---|---|
| 214 | 423.262 | 2 | 133.23 | 0.0564 | 16.50 | 28.74 | 4 | 0.0 | 0 | K.YYADSVK.G | | 78-84 | CID |
| 391 | 659.82 | 2 | 45.09 | 0.0297 | 20.37 | 79.62 | 3 | 0.0 | 0 | R.AEDTAVYYCAR.- | Carbamidomethyl: 9 | 107-117 | CID |

#### Protein 13: Immunoglobulin heavy variable 1-69 OS=Homo sapiens GN=IGHV1-69 PE=1 SV=2

**Accession:** HV169_HUMAN
**Database:** SwissProt
**MW [kDa] / pI:** 12.7 / 9.3
**Modification(s):** Carbamidomethyl

**Score:** 91.5
**Seq. Coverage [%]:** 9.4
**No. of Peptides:** 1

| 10 | 20 | 30 | 40 | 50 | 60 | 70 | 80 | 90 | 100 |
|---|---|---|---|---|---|---|---|---|---|
| MDWTWRFLFV | VAAATGVQSQ | VQLVQSGAEV | KKPGSSVKVS | CKASGGTFSS | YAISWVRQAP | GQGLEWMGGI | IPIFGTANYA | QKFQGRVTIT | ADKSTSTAYM |

| 110 | 120 |
|---|---|
| ELSSLRSEDT | AVYYCAR |

MS/MS Peptide Matches

| Cmpd. | m/z meas. | z | Δ m/z [ppm] | Δ m/z [Da] | Rt [min] | Score | No. of Cmpds. | Site [%] | P | Sequence | Modification | Range | Type |
|---|---|---|---|---|---|---|---|---|---|---|---|---|---|
| 392 | 667.79 | 2 | 3.43 | 0.0023 | 20.38 | 91.45 | 4 | 0.0 | 0 | R.SEDTAVYYCAR.- | Carbamidomethyl: 9 | 107-117 | CID |

#### Protein 14: Immunoglobulin kappa variable 3-20 OS=Homo sapiens GN=IGKV3-20 PE=1 SV=2

**Accession:** KV320_HUMAN
**Database:** SwissProt
**MW [kDa] / pI:** 12.5 / 4.9

**Score:** 69.7
**Seq. Coverage [%]:** 7.8
**No. of Peptides:** 1

| 10 | 20 | 30 | 40 | 50 | 60 | 70 | 80 | 90 | 100 |
|---|---|---|---|---|---|---|---|---|---|
| METPAQLLFL | LLLWLPDTTG | EIVLTQSPGT | LSLSPGERAT | LSCRASQSVS | SSYLAWYQQK | PGQAPRLLIY | GASSRATGIP | DRFSGSGSGT | DFTLTISRLE |
| 110 | 120 | | | | | | | | |
| PEDFAVYYCQ | QYGSSP | | | | | | | | |

MS/MS Peptide Matches

| Cmpd. | m/z meas. | z | Δ m/z [ppm] | Δ m/z [Da] | Rt [min] | Score | No. of Cmpds. | Site [%] | P | Sequence | Modification | Range | Type |
|---|---|---|---|---|---|---|---|---|---|---|---|---|---|
| 570 | 490.337 | 2 | 111.79 | 0.0548 | 24.14 | 69.71 | 3 | 0.0 | 0 | R.LLIYGASSR.A | | 67-75 | CID |

#### Protein 15: Immunoglobulin kappa variable 3-11 OS=Homo sapiens GN=IGKV3-11 PE=1 SV=1

**Accession:** KV311_HUMAN
**Database:** SwissProt
**MW [kDa] / pI:** 12.6 / 4.9
**Score:** 69.0
**Seq. Coverage [%]:** 7.8
**No. of Peptides:** 1

| 10 | 20 | 30 | 40 | 50 | 60 | 70 | 80 | 90 | 100 |
|---|---|---|---|---|---|---|---|---|---|
| MEAPAQLLFL | LLLWLPDTTG | EIVLTQSPAT | LSLSPGERAT | LSCRASQSVS | SYLAWYQQKP | GQAPRLLIYD | ASNRATGIPA | RFSGSGSGTD | FTLTISSLEP |
| 110 | 120 | | | | | | | | |
| EDFAVYYCQQ | RSNWP | | | | | | | | |

MS/MS Peptide Matches

| Cmpd. | m/z meas. | z | Δ m/z [ppm] | Δ m/z [Da] | Rt [min] | Score | No. of Cmpds. | Site [%] | P | Sequence | Modification | Range | Type |
|---|---|---|---|---|---|---|---|---|---|---|---|---|---|
| 580 | 532.831 | 2 | 76.24 | 0.0406 | 24.36 | 69.03 | 5 | 0.0 | 0 | R.LLIYDASNR.A | | 66-74 | CID |

#### Protein 16: Immunoglobulin heavy constant alpha 1 OS=Homo sapiens GN=IGHA1 PE=1 SV=2

**Accession:** IGHA1_HUMAN
**Database:** SwissProt
**MW [kDa] / pI:** 37.6 / 6.1
**Score:** 58.9
**Seq. Coverage [%]:** 3.4
**No. of Peptides:** 1

**Modification(s):** Carbamidomethyl

| 10 | 20 | 30 | 40 | 50 | 60 | 70 | 80 | 90 | 100 |
|---|---|---|---|---|---|---|---|---|---|
| ASPTSPKVFP | LSLCSTQPDG | NVVIACLVQG | FFPQEPLSVT | WSESGQGVTA | RNFPPSQDAS | GDLYTTSSQL | TLPATQCLAG | KSVTCHVKHY | TNPSQDVTVP |
| **110** | **120** | **130** | **140** | **150** | **160** | **170** | **180** | **190** | **200** |
| CPVPSTPPTP | SPSTPPTPSP | SCCHPRLSLH | RPALEDLLLG | SEANLTCTLT | GLRDASGVTF | TWTPSSGKSA | VQGPPERDLC | GCYSVSSVLP | GCAEPWNHGK |
| **210** | **220** | **230** | **240** | **250** | **260** | **270** | **280** | **290** | **300** |
| TFTCTAAYPE | SKTPLTATLS | KSGNTFRPEV | HLLPPPSEEL | ALNELVTLTC | LARGFSPKDV | LVRWLQGSQE | LPREKYLTWA | SRQEPSQGTT | TFAVTSILRV |
| **310** | **320** | **330** | **340** | **350** | **360** | | | | |
| AAEDWKKGDT | FSCMVGHEAL | PLAFTQKTID | RLAGKPTHVN | VSVVMAEVDG | TCY | | | | |

MS/MS Peptide Matches

| Cmpd. | m/z meas. | z | Δ m/z [ppm] | Δ m/z [Da] | Rt [min] | Score | No. of Cmpds. | Site [%] | P | Sequence | Modification | Range | Type |
|---|---|---|---|---|---|---|---|---|---|---|---|---|---|
| 404 | 688.337 | 2 | 34.05 | 0.0234 | 20.60 | 58.94 | 3 | 0.0 | 0 | K.TFTCTAAYPESK.T | Carbamidomethyl: 4 | 201-212 | CID |

#### Protein 17: Immunoglobulin kappa variable 3-15 OS=Homo sapiens GN=IGKV3-15 PE=1 SV=2

**Accession:** KV315_HUMAN
**Database:** SwissProt
**MW [kDa] / pI:** 12.5 / 5.1
**Score:** 56.3
**Seq. Coverage [%]:** 7.8
**No. of Peptides:** 1

| 10 | 20 | 30 | 40 | 50 | 60 | 70 | 80 | 90 | 100 |
|---|---|---|---|---|---|---|---|---|---|
| MEAPAQLLFL | LLLWLPDTTG | EIVMTQSPAT | LSVSPGERAT | LSCRASQSVS | SNLAWYQQKP | GQAPRLLIYG | ASTRATGIPA | RFSGSGSGTE | FTLTISSLQS |
| **110** | **120** | | | | | | | | |
| EDFAVYYCQQ | YNNWP | | | | | | | | |

MS/MS Peptide Matches

| Cmpd. | m/z meas. | z | Δ m/z [ppm] | Δ m/z [Da] | Rt [min] | Score | No. of Cmpds. | Site [%] | P | Sequence | Modification | Range | Type |
|---|---|---|---|---|---|---|---|---|---|---|---|---|---|
| 589 | 497.326 | 2 | 72.36 | 0.0360 | 24.54 | 56.25 | 3 | 0.0 | 0 | R.LLIYGASTR.A | | 66-74 | CID |

#### Protein 18: Immunoglobulin lambda variable 5-45 OS=Homo sapiens GN=IGLV5-45 PE=3 SV=1

**Accession:** LV545_HUMAN
**Database:** SwissProt
**MW [kDa] / pI:** 13.2 / 6.7
**Score:** 42.1
**Seq. Coverage [%]:** 11.4
**No. of Peptides:** 1

| 10 | 20 | 30 | 40 | 50 | 60 | 70 | 80 | 90 | 100 |
|---|---|---|---|---|---|---|---|---|---|
| MAWTPLLLLF | LSHCTGSLSQ | AVLTQPSSLS | ASPGASASLT | CTLCSGINVG | TYRIYWYQQK | PGSPPQYLLR | YKSDSDKQQG | SGVPSRFSGS | KDASANAGIL |
| **110** | **120** | **130** | | | | | | | |
| LISGLQSEDE | ADYYCMIWHS | SAS | | | | | | | |

MS/MS Peptide Matches

| Cmpd. | m/z meas. | z | Δ m/z [ppm] | Δ m/z [Da] | Rt [min] | Score | No. of Cmpds. | Site [%] | P | Sequence | Modification | Range | Type |
|---|---|---|---|---|---|---|---|---|---|---|---|---|---|
| 70 | 483.276 | 3 | 94.00 | 0.0454 | 13.17 | 42.06 | 1 | 0.0 | 1 | K.SDSDKQQGSGVPSR.F | | 73-86 | CID |

#### Protein 19: Immunoglobulin heavy variable 3-73 OS=Homo sapiens GN=IGHV3-73 PE=3 SV=1

**Accession:** HV373_HUMAN
**Database:** SwissProt
**MW [kDa] / pI:** 12.8 / 9.3
**Score:** 40.8
**Seq. Coverage [%]:** 5.9
**No. of Peptides:** 1

| 10 | 20 | 30 | 40 | 50 | 60 | 70 | 80 | 90 | 100 |
|---|---|---|---|---|---|---|---|---|---|
| MEFGLSWVFL | VAILKGVQCE | VQLVESGGGL | VQPGGSLKLS | CAASGFTFSG | SAMHWVRQAS | GKGLEWVGRI | RSKANSYATA | YAASVKGRFT | ISRDDSKNTA |
| **110** | **120** | | | | | | | | |
| YLQMNSLKTE | DTAVYYCTR | | | | | | | | |

MS/MS Peptide Matches

| Cmpd. | m/z meas. | z | Δ m/z [ppm] | Δ m/z [Da] | Rt [min] | Score | No. of Cmpds. | Site [%] | P | Sequence | Modification | Range | Type |
|---|---|---|---|---|---|---|---|---|---|---|---|---|---|
| 593 | 408.762 | 2 | 98.43 | 0.0402 | 24.62 | 40.8 | 3 | 0.0 | 0 | K.GLEWVGR.I | | 63-69 | CID |

#### Project Info

| | | | |
|---|---|---|---|
| **Name:** | Alagesan | **Date:** | Mar 5, 2019 |
| **Note:** | Nanobooster | | |

#### Sample Info & Protocols

| | |
|---|---|
| **Name:** | Nanobooster - IgG |
| **Date:** | Mar 5, 2019 |

#### Search Result Info

| | | | |
|---|---|---|---|
| **Search Result:** | IonTrap_Human_Swiss_CID_2019-03-05 17:45:09 | | |
| **Location:** | /Alagesan/Nanobooster - IgG/Ka_IgG Trypsin NoEnrich NB ACN_BA5_01_8750.mgf | | |
| **Search Method:** | IonTrap_Human_Swiss_CID | **Version:** | 1.2 |
| **Search Engine(s):** | Mascot, 2.6.0 | **Ident. Compound(s):** | 63/2396 |
| **Database(s):** | SwissProt, SwissProt_2017_04.fasta | | |
| **Note:** | | | |

BRUKER

#### Results

##### Protein 1: Immunoglobulin gamma-1 heavy chain OS=Homo sapiens PE=1 SV=1

| | | | |
|---|---|---|---|
| **Accession:** | IGG1_HUMAN | **Score:** | 931.3 |
| **Database:** | SwissProt | **Seq. Coverage [%]:** | 40.3 |
| **MW [kDa] / pI:** | 49.3 / 8.9 | **No. of Peptides:** | 14 |
| **Modification(s):** | Carbamidomethyl, Oxidation | | |

```
        10         20         30         40         50         60         70         80         90        100
QVQLVQSGGG VVQPGRSLRL SCAASGFTFS RYTIHWVRQA PGKGLEWVAV MSYBGBBKHY ADSVNGRFTI SRNDSKNTLY LNMNSLRPED TAVYYCARIR
       110        120        130        140        150        160        170        180        190        200
DTAMFFAHWG QGTLVTVSSA STKGPSVFPL APSSKSTSGG TAALGCLVKD YFPEPVTVSW NSGALTSGVH TFPAVLQSSG LYSLSSVVTV PSSSLGTQTY
       210        220        230        240        250        260        270        280        290        300
ICNVNHKPSN TKVDKKVEPK SCDKTHTCPP CPAPELLGGP SVFLFPPKPK DTLMISRTPE VTCVVVDVSH EDPEVKFNWY VDGVEVHNAK TKPREEQYNS
       310        320        330        340        350        360        370        380        390        400
TYRVVSVLTV LHQDWLNGKE YKCKVSNKAL PAPIEKTISK AKGQPREPQV YTLPPSRDEL TKNQVSLTCL VKGFYPSDIA VEWESNGQPE NNYKTTPPVL
       410        420        430        440        450
DSDGSFFLYS KLTVDKSRWQ QGNVFSCSVM HEALHNHYTQ KSLSLSPGK
```

MS/MS Peptide Matches

| Cmpd. | m/z meas. | z | Δ m/z [ppm] | Δ m/z [Da] | Rt [min] | Score | No. of Cmpds. | Site [%] | P | Sequence | Modification | Range | Type |
|---|---|---|---|---|---|---|---|---|---|---|---|---|---|
| 906 | 593.848 | 2 | 35.43 | 0.0210 | 27.24 | 72.97 | 12 | 0.0 | 0 | K.GPSVFPLAPSSK.S | | 124-135 | CID |
| 813 | 661.373 | 2 | 45.89 | 0.0303 | 25.15 | 119.54 | 14 | 0.0 | 0 | K.STSGGTAALGCLVK.D | Carbamidomethyl: 11 | 136-149 | CID |
| 1202 | 712.199 | 4 | 462.39 | 0.3292 | 32.79 | 46.13 | 9 | 0.0 | 0 | K.THTCPPCPAPELLGGPSVFLFPPKPK.D | Carbamidomethyl: 4, 7 | 225-250 | CID |
| 662 | 418.227 | 2 | 14.96 | 0.0063 | 22.01 | 57.64 | 11 | 0.0 | 0 | K.DTLMISR.T | | 251-257 | CID |
| 551 | 426.243 | 2 | 58.18 | 0.0248 | 19.76 | 47.37 | 4 | 0.0 | 0 | K.DTLMISR.T | Oxidation: 4 | 251-257 | CID |
| 901 | 713.719 | 3 | 53.72 | 0.0383 | 27.15 | 98.69 | 2 | 0.0 | 0 | R.TPEVTCVVVDVSHEDPEVK.F | Carbamidomethyl: 6 | 258-276 | CID |

| Cmpd. | m/z meas. | z | Δ m/z [ppm] | Δ m/z [Da] | Rt [min] | Score | No. of Cmpds. | Site [%] | P | Sequence | Modification | Range | Type |
|---|---|---|---|---|---|---|---|---|---|---|---|---|---|
| 910 | 560.012 | 3 | 130.65 | 0.0732 | 27.31 | 64.64 | 10 | 0.0 | 0 | K.FNWYVDGVEVHNAK.T | | 277-290 | CID |
| 1276 | 603.426 | 3 | 141.96 | 0.0857 | 34.33 | 63.26 | 13 | 0.0 | 0 | R.VVSVLTVLHQDWLNGK.E | | 304-319 | CID |
| 632 | 419.801 | 2 | 108.92 | 0.0457 | 21.36 | 25.66 | 5 | 0.0 | 0 | K.ALPAPIEK.T | | 329-336 | CID |
| 736 | 643.793 | 2 | -73.94 | -0.0476 | 23.57 | 53.73 | 11 | 0.0 | 0 | R.EPQVYTLPPSR.D | | 347-357 | CID |
| 767 | 625.042 | 3 | 75.35 | 0.0471 | 24.21 | 32.56 | 11 | 0.0 | 1 | R.EPQVYTLPPSRDELTK.N | | 347-362 | CID |
| 873 | 581.349 | 2 | 52.56 | 0.0306 | 26.52 | 71.57 | 11 | 0.0 | 0 | K.NQVSLTCLVK.G | Carbamidomethyl: 7 | 363-372 | CID |
| 1106 | 849.069 | 3 | 416.74 | 0.3537 | 31.02 | 79.75 | 32 | 0.0 | 0 | K.GFYPSDIAVEWESNGQPENNYK.T | | 373-394 | CID |
| 1168 | 937.662 | 2 | 210.62 | 0.1975 | 32.13 | 97.82 | 9 | 0.0 | 0 | K.TTPPVLDSDGSFFLYSK.L | | 395-411 | CID |

#### Protein 2: Immunoglobulin heavy constant gamma 3 OS=Homo sapiens GN=IGHG3 PE=1 SV=2

**Accession:** IGHG3_HUMAN
**Database:** SwissProt
**MW [kDa] / pI:** 41.3 / 8.2
**Modification(s):** Carbamidomethyl, Oxidation

**Score:** 840.2
**Seq. Coverage [%]:** 36.3
**No. of Peptides:** 13

```
        10         20         30         40         50         60         70         80         90        100
ASTKGPSVFP LAPCSRSTSG GTAALGCLVK DYFPEPVTVS WNSGALTSGV HTFPAVLQSS GLYSLSSVVT VPSSSLGTQT YTCNVNHKPS NTKVDKRVEL
       110        120        130        140        150        160        170        180        190        200
KTPLGDTTHT CPRCPEPKSC DTPPPCPRCP EPKSCDTPPP CPRCPEPKSC DTPPPCPRCP APELLGGPSV FLFPPKPKDT LMISRTPEVT CVVVDVSHED
       210        220        230        240        250        260        270        280        290        300
PEVQFKWYVD GVEVHNAKTK PREEQYNSTF RVVSVLTVLH QDWLNGKEYK CKVSNKALPA PIEKTISKTK GQPREPQVYT LPPSREEMTK NQVSLTCLVK
       310        320        330        340        350        360        370        380
GFYPSDIAVE WESSGQPENN YNTTPPMLDS DGSFFLYSKL TVDKSRWQQG NIFSCSVMHE ALHNRFTQKS LSLSPGK
```

MS/MS Peptide Matches

| Cmpd. | m/z meas. | z | Δ m/z [ppm] | Δ m/z [Da] | Rt [min] | Score | No. of Cmpds. | Site [%] | P | Sequence | Modification | Range | Type |
|---|---|---|---|---|---|---|---|---|---|---|---|---|---|
| 950 | 644.342 | 2 | 19.64 | 0.0127 | 28.10 | 67.75 | 15 | 0.0 | 0 | K.GPSVFPLAPCSR.S | Carbamidomethyl: 10 | 5-16 | CID |
| 813 | 661.373 | 2 | 45.89 | 0.0303 | 25.15 | 119.54 | 14 | 0.0 | 0 | R.STSGGTAALGCLVK.D | Carbamidomethyl: 11 | 17-30 | CID |
| 420 | 452.579 | 3 | 62.77 | 0.0284 | 17.10 | 65.76 | 6 | 0.0 | 0 | K.TPLGDTTHTCPR.C | Carbamidomethyl: 10 | 102-113 | CID |
| 383 | 593.75 | 2 | -4.41 | -0.0026 | 16.25 | 64.32 | 10 | 0.0 | 0 | K.SCDTPPPCPR.C | Carbamidomethyl: 2, 8 | 119-128 | CID |
| 1265 | 717.772 | 3 | 62.40 | 0.0448 | 34.13 | 117.39 | 3 | 0.0 | 0 | R.CPAPELLGGPSVFLFPPKPK.D | Carbamidomethyl: 1 | 159-178 | CID |
| 662 | 418.227 | 2 | 14.96 | 0.0063 | 22.01 | 57.64 | 11 | 0.0 | 0 | K.DTLMISR.T | | 179-185 | CID |
| 551 | 426.243 | 2 | 58.18 | 0.0248 | 19.76 | 47.37 | 4 | 0.0 | 0 | K.DTLMISR.T | Oxidation: 4 | 179-185 | CID |
| 741 | 472.953 | 3 | 108.41 | 0.0513 | 23.69 | 37.37 | 5 | 0.0 | 0 | K.WYVDGVEVHNAK.T | | 207-218 | CID |
| 1276 | 603.426 | 3 | 141.96 | 0.0857 | 34.33 | 63.26 | 13 | 0.0 | 0 | R.VVSVLTVLHQDWLNGK.E | | 232-247 | CID |
| 632 | 419.801 | 2 | 108.92 | 0.0457 | 21.36 | 25.66 | 5 | 0.0 | 0 | K.ALPAPIEK.T | | 257-264 | CID |
| 736 | 643.793 | 2 | -73.94 | -0.0476 | 23.57 | 53.73 | 11 | 0.0 | 0 | R.EPQVYTLPPSR.E | | 275-285 | CID |
| 718 | 635.757 | 3 | 164.77 | 0.1047 | 23.20 | 48.79 | 30 | 0.0 | 1 | R.EPQVYTLPPSREEMTK.N | | 275-290 | CID |
| 873 | 581.349 | 2 | 52.56 | 0.0306 | 26.52 | 71.57 | 11 | 0.0 | 0 | K.NQVSLTCLVK.G | Carbamidomethyl: 7 | 291-300 | CID |

##### Protein 3: Immunoglobulin kappa constant OS=Homo sapiens GN=IGKC PE=1 SV=2

**Accession:** IGKC_HUMAN
**Database:** SwissProt
**MW [kDa] / pI:** 11.8 / 6.1
**Modification(s):** Carbamidomethyl

**Score:** 766.3
**Seq. Coverage [%]:** 86.9
**No. of Peptides:** 8

```
        10         20         30         40         50         60         70         80         90         100
RTVAAPSVFI FPPSDEQLKS GTASVVCLLN NFYPREAKVQ WKVDNALQSG NSQESVTEQD SKDSTYSLSS TLTLSKADYE KHKVYACEVT HQGLSSPVTK
       110
SFNRGEC
```

MS/MS Peptide Matches

| Cmpd. | m/z meas. | z | Δ m/z [ppm] | Δ m/z [Da] | Rt [min] | Score | No. of Cmpds. | Site [%] | P | Sequence | Modification | Range | Type |
|---|---|---|---|---|---|---|---|---|---|---|---|---|---|
| 1104 | 701.424 | 3 | 61.49 | 0.0431 | 30.96 | 45.44 | 10 | 0.0 | 1 | -.RTVAAPSVFIFPPSDEQLK.S | | 1-19 | CID |
| 1243 | 973.609 | 2 | 94.38 | 0.0919 | 33.72 | 95.81 | 18 | 0.0 | 0 | R.TVAAPSVFIFPPSDEQLK.S | | 2-19 | CID |
| 1323 | 600.047 | 3 | 128.46 | 0.0771 | 35.27 | 71.3 | 14 | 0.0 | 0 | K.SGTASVVCLLNNFYPR.E | Carbamidomethyl: 8 | 20-35 | CID |
| 489 | 1068.463 | 2 | -23.40 | -0.0250 | 18.50 | 159.92 | 16 | 0.0 | 0 | K.VDNALQSGNSQESVTEQDSK.D | | 43-62 | CID |
| 893 | 1207.55 | 3 | 255.71 | 0.3087 | 26.97 | 138.2 | 4 | 0.0 | 1 | K.VDNALQSGNSQESVTEQDSKDSTYSLSSTLTLSK.A | | 43-76 | CID |
| 944 | 751.902 | 2 | 25.46 | 0.0191 | 27.96 | 97.34 | 8 | 0.0 | 0 | K.DSTYSLSSTLTLSK.A | | 63-76 | CID |
| 640 | 626.253 | 3 | 435.33 | 0.2725 | 21.57 | 103.98 | 11 | 0.0 | 0 | K.VYACEVTHQGLSSPVTK.S | Carbamidomethyl: 4 | 84-100 | CID |
| 275 | 435.17 | 2 | -27.92 | -0.0122 | 13.88 | 54.34 | 5 | 0.0 | 1 | K.SFNRGEC.- | Carbamidomethyl: 7 | 101-107 | CID |

#### Protein 4: Immunoglobulin heavy constant gamma 2 OS=Homo sapiens GN=IGHG2 PE=1 SV=2

**Accession:** IGHG2_HUMAN
**Database:** SwissProt
**MW [kDa] / pI:** 35.9 / 7.7
**Modification(s):** Carbamidomethyl, Oxidation

**Score:** 735.5
**Seq. Coverage [%]:** 37.4
**No. of Peptides:** 13

```
        10         20         30         40         50         60         70         80         90        100
ASTKGPSVFP LAPCSRSTSE STAALGCLVK DYFPEPVTVS WNSGALTSGV HTFPAVLQSS GLYSLSSVVT VPSSNFGTQT YTCNVDHKPS NTKVDKTVER
       110        120        130        140        150        160        170        180        190        200
KCCVECPPCP APPVAGPSVF LFPPKPKDTL MISRTPEVTC VVVDVSHEDP EVQFNWYVDG VEVHNAKTKP REEQFNSTFR VVSVLTVVHQ DWLNGKEYKC
       210        220        230        240        250        260        270        280        290        300
KVSNKGLPAP IEKTISKTKG QPREPQVYTL PPSREEMTKN QVSLTCLVKG FYPSDISVEW ESNGQPENNY KTTPPMLDSD GSFFLYSKLT VDKSRWQQGN
       310        320        330
VFSCSVMHEA LHNHYTQKSL SLSPGK
```

MS/MS Peptide Matches

| Cmpd. | m/z meas. | z | Δ m/z [ppm] | Δ m/z [Da] | Rt [min] | Score | No. of Cmpds. | Site [%] | P | Sequence | Modification | Range | Type |
|---|---|---|---|---|---|---|---|---|---|---|---|---|---|
| 950 | 644.342 | 2 | 19.64 | 0.0127 | 28.10 | 67.75 | 15 | 0.0 | 0 | K.GPSVFPLAPCSR.S | Carbamidomethyl: 10 | 5-16 | CID |
| 825 | 712.373 | 2 | 20.36 | 0.0145 | 25.44 | 85.03 | 10 | 0.0 | 0 | R.STSESTAALGCLVK.D | Carbamidomethyl: 11 | 17-30 | CID |
| 1087 | 760.167 | 4 | 378.19 | 0.2874 | 30.64 | 28.8 | 3 | 0.0 | 1 | R.KCCVECPPCPAPPVAGPSVFLFPPKPK.D | Carbamidomethyl: 2, 3, 6, 9 | 101-127 | CID |
| 1173 | 970.481 | 3 | 352.79 | 0.3423 | 32.25 | 91.15 | 5 | 0.0 | 0 | K.CCVECPPCPAPPVAGPSVFLFPPKPK.D | Carbamidomethyl: 1, 2, 5, 8 | 102-127 | CID |
| 662 | 418.227 | 2 | 14.96 | 0.0063 | 22.01 | 57.64 | 11 | 0.0 | 0 | K.DTLMISR.T | | 128-134 | CID |
| 551 | 426.243 | 2 | 58.18 | 0.0248 | 19.76 | 47.37 | 4 | 0.0 | 0 | K.DTLMISR.T | Oxidation: 4 | 128-134 | CID |
| 1166 | 599.084 | 3 | 694.10 | 0.4155 | 32.09 | 40.48 | 3 | 0.0 | 0 | R.VVSVLTVVHQDWLNGK.E | | 181-196 | CID |
| 1090 | 554.628 | 4 | 585.62 | 0.3246 | 30.69 | 21.3 | 2 | 0.0 | 1 | R.VVSVLTVVHQDWLNGKEYK.C | | 181-199 | CID |
| 736 | 643.793 | 2 | -73.94 | -0.0476 | 23.57 | 53.73 | 11 | 0.0 | 0 | R.EPQVYTLPPSR.E | | 224-234 | CID |
| 718 | 635.757 | 3 | 164.77 | 0.1047 | 23.20 | 48.79 | 30 | 0.0 | 1 | R.EPQVYTLPPSREEMTK.N | | 224-239 | CID |
| 873 | 581.349 | 2 | 52.56 | 0.0306 | 26.52 | 71.57 | 11 | 0.0 | 0 | K.NQVSLTCLVK.G | Carbamidomethyl: 7 | 240-249 | CID |
| 1186 | 953.586 | 2 | 142.03 | 0.1354 | 32.51 | 87.12 | 6 | 0.0 | 0 | K.TTPPMLDSDGSFFLYSK.L | | 272-288 | CID |
| 1092 | 961.459 | 2 | 11.40 | 0.0110 | 30.70 | 34.74 | 2 | 0.0 | 0 | K.TTPPMLDSDGSFFLYSK.L | Oxidation: 5 | 272-288 | CID |

##### Protein 5: Immunoglobulin lambda constant 2 OS=Homo sapiens GN=IGLC2 PE=1 SV=1

**Accession:** IGLC2_HUMAN
**Database:** SwissProt
**MW [kDa] / pI:** 11.3 / 6.9
**Modification(s):** Carbamidomethyl

**Score:** 641.8
**Seq. Coverage [%]:** 93.4
**No. of Peptides:** 9

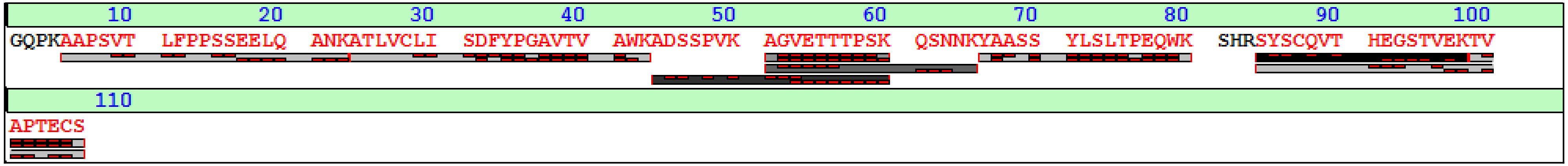


MS/MS Peptide Matches

| Cmpd. | m/z meas. | z | Δ m/z [ppm] | Δ m/z [Da] | Rt [min] | Score | No. of Cmpds. | Site [%] | P | Sequence | Modification | Range | Type |
|---|---|---|---|---|---|---|---|---|---|---|---|---|---|
| 1025 | 662.731 | 3 | 80.78 | 0.0535 | 29.51 | 80.23 | 9 | 0.0 | 0 | K.AAPSVTLFPPSSEELQANK.A | | 5-23 | CID |
| 1589 | 1106.083 | 2 | 3.11 | 0.0034 | 40.25 | 73.3 | 1 | 0.0 | 0 | K.ATLVCLISDFYPGAVTVAWK.A | Carbamidomethyl: 5 | 24-43 | CID |
| 449 | 558.964 | 3 | 13.70 | 0.0077 | 17.68 | 76.61 | 3 | 0.0 | 1 | K.ADSSPVKAGVETTTPSK.Q | | 44-60 | CID |
| 300 | 495.784 | 2 | 50.94 | 0.0253 | 14.47 | 75.51 | 16 | 0.0 | 0 | K.AGVETTTPSK.Q | | 51-60 | CID |
| 254 | 521.339 | 3 | 141.20 | 0.0736 | 13.34 | 57.13 | 4 | 0.0 | 1 | K.AGVETTTPSKQSNNK.Y | | 51-65 | CID |
| 1065 | 872.476 | 2 | 49.23 | 0.0429 | 30.25 | 91.18 | 8 | 0.0 | 0 | K.YAASSYLSLTPEQWK.S | | 66-80 | CID |
| 414 | 571.264 | 3 | 10.65 | 0.0061 | 16.99 | 78.36 | 17 | 0.0 | 0 | R.SYSCQVTHEGSTVEK.T | Carbamidomethyl: 4 | 84-98 | CID |
| 616 | 853.066 | 3 | 25.53 | 0.0218 | 21.04 | 59.88 | 3 | 0.0 | 1 | R.SYSCQVTHEGSTVEKTVAPTECS.- | Carbamidomethyl: 4, 22 | 84-106 | CID |
| 353 | 432.759 | 2 | 154.80 | 0.0670 | 15.63 | 49.57 | 1 | 0.0 | 0 | K.TVAPTECS.- | Carbamidomethyl: 7 | 99-106 | CID |

#### Protein 6: Immunoglobulin heavy constant gamma 4 OS=Homo sapiens GN=IGHG4 PE=1 SV=1

**Accession:** IGHG4_HUMAN
**Database:** SwissProt
**MW [kDa] / pI:** 35.9 / 7.2
**Modification(s):** Carbamidomethyl, Oxidation

**Score:** 576.8
**Seq. Coverage [%]:** 37.3
**No. of Peptides:** 9

| 10 | 20 | 30 | 40 | 50 | 60 | 70 | 80 | 90 | 100 |
|---|---|---|---|---|---|---|---|---|---|
| ASTKGPSVFP | LAPCSRSTSE | STAALGCLVK | DYFPEPVTVS | WNSGALTSGV | HTFPAVLQSS | GLYSLSSVVT | VPSSSLGTKT | YTCNVDHKPS | NTKVDKRVES |
| **110** | **120** | **130** | **140** | **150** | **160** | **170** | **180** | **190** | **200** |
| KYGPPCPSCP | APEFLGGPSV | FLFPPKPKDT | LMISRTPEVT | CVVVDVSQED | PEVQFNWYVD | GVEVHNAKTK | PREEQFNSTY | RVVSVLTVLH | QDWLNGKEYK |
| **210** | **220** | **230** | **240** | **250** | **260** | **270** | **280** | **290** | **300** |
| CKVSNKGLPS | SIEKTISKAK | GQPREPQVYT | LPPSQEEMTK | NQVSLTCLVK | GFYPSDIAVE | WESNGQPENN | YKTTPPVLDS | DGSFFLYSRL | TVDKSRWQEG |
| **310** | **320** | **330** | | | | | | | |
| NVFSCSVMHE | ALHNHYTQKS | LSLSLGK | | | | | | | |

**MS/MS Peptide Matches**

| Cmpd. | m/z meas. | z | Δ m/z [ppm] | Δ m/z [Da] | Rt [min] | Score | No. of Cmpds. | Site [%] | P | Sequence | Modification | Range | Type |
|---|---|---|---|---|---|---|---|---|---|---|---|---|---|
| 950 | 644.342 | 2 | 19.64 | 0.0127 | 28.10 | 67.75 | 15 | 0.0 | 0 | K.GPSVFPLAPCSR.S | Carbamidomethyl: 10 | 5-16 | CID |
| 825 | 712.373 | 2 | 20.36 | 0.0145 | 25.44 | 85.03 | 10 | 0.0 | 0 | R.STSESTAALGCLVK.D | Carbamidomethyl: 11 | 17-30 | CID |
| 282 | 416.972 | 4 | 57.85 | 0.0241 | 14.05 | 17.58 | 2 | 0.0 | 0 | K.TYTCNVDHKPSNTK.V | Carbamidomethyl: 4 | 80-93 | CID |
| 1308 | 982.198 | 3 | 381.01 | 0.3741 | 35.00 | 86.86 | 6 | 0.0 | 0 | K.YGPPCPSCPAPEFLGGPSVFLFPPKPK.D | Carbamidomethyl: 5, 8 | 102-128 | CID |
| 662 | 418.227 | 2 | 14.96 | 0.0063 | 22.01 | 57.64 | 11 | 0.0 | 0 | K.DTLMISR.T | | 129-135 | CID |
| 551 | 426.243 | 2 | 58.18 | 0.0248 | 19.76 | 47.37 | 4 | 0.0 | 0 | K.DTLMISR.T | Oxidation: 4 | 129-135 | CID |
| 1276 | 603.426 | 3 | 141.96 | 0.0857 | 34.33 | 63.26 | 13 | 0.0 | 0 | R.VVSVLTVLHQDWLNGK.E | | 182-197 | CID |
| 873 | 581.349 | 2 | 52.56 | 0.0306 | 26.52 | 71.57 | 11 | 0.0 | 0 | K.NQVSLTCLVK.G | Carbamidomethyl: 7 | 241-250 | CID |
| 1106 | 849.069 | 3 | 416.74 | 0.3537 | 31.02 | 79.75 | 32 | 0.0 | 0 | K.GFYPSDIAVEWESNGQPENNYK.T | | 251-272 | CID |

#### Protein 7: Immunoglobulin lambda-1 light chain OS=Homo sapiens PE=1 SV=1

| | | | |
|---|---|---|---|
| **Accession:** | IGL1_HUMAN | **Score:** | 443.1 |
| **Database:** | SwissProt | **Seq. Coverage [%]:** | 39.4 |
| **MW [kDa] / pI:** | 22.8 / 6.3 | **No. of Peptides:** | 7 |
| **Modification(s):** | Carbamidomethyl | | |

```
        10         20         30         40         50         60         70         80         90        100
QSALTQPPSA SGSLGQSVTI SCTGTSSDVG GYNYVSWYQQ HAGKAPKVII YEVNKRPSGV PDRFSGSKSG NTASLTVSGL QAEDEADYYC SSYEGSDNFV
       110        120        130        140        150        160        170        180        190        200
FGTGTKVTVL GQPKANPTVT LFPPSSEELQ ANKATLVCLI SDFYPGAVTV AWKADGSPVK AGVETTKPSK QSNNKYAASS YLSLTPEQWK SHRSYSCQVT
       210        220
HEGSTVEKTV APTECS
```

MS/MS Peptide Matches

| Cmpd. | m/z meas. | z | Δ m/z [ppm] | Δ m/z [Da] | Rt [min] | Score | No. of Cmpds. | Site [%] | P | Sequence | Modification | Range | Type |
|---|---|---|---|---|---|---|---|---|---|---|---|---|---|
| 561 | 421.322 | 2 | 145.45 | 0.0613 | 19.94 | 39.73 | 4 | 0.0 | 0 | K.VTVLGQPK.A | | 107-114 | CID |
| 1019 | 681.698 | 3 | 19.63 | 0.0134 | 29.35 | 51.11 | 6 | 0.0 | 0 | K.ANPTVTLFPPSSEELQANK.A | | 115-133 | CID |
| 1589 | 1106.083 | 2 | 3.11 | 0.0034 | 40.25 | 73.3 | 1 | 0.0 | 0 | K.ATLVCLISDFYPGAVTVAWK.A | Carbamidomethyl: 5 | 134-153 | CID |
| 1065 | 872.476 | 2 | 49.23 | 0.0429 | 30.25 | 91.18 | 8 | 0.0 | 0 | K.YAASSYLSLTPEQWK.S | | 176-190 | CID |
| 414 | 571.264 | 3 | 10.65 | 0.0061 | 16.99 | 78.36 | 17 | 0.0 | 0 | R.SYSCQVTHEGSTVEK.T | Carbamidomethyl: 4 | 194-208 | CID |
| 616 | 853.066 | 3 | 25.53 | 0.0218 | 21.04 | 59.88 | 3 | 0.0 | 1 | R.SYSCQVTHEGSTVEKTVAPTECS.- | Carbamidomethyl: 4, 22 | 194-216 | CID |
| 353 | 432.759 | 2 | 154.80 | 0.0670 | 15.63 | 49.57 | 1 | 0.0 | 0 | K.TVAPTECS.- | Carbamidomethyl: 7 | 209-216 | CID |

#### Protein 8: Immunoglobulin heavy variable 3-7 OS=Homo sapiens GN=IGHV3-7 PE=1 SV=2

**Accession:** HV307_HUMAN
**Database:** SwissProt
**MW [kDa] / pI:** 12.9 / 6.2
**Modification(s):** Carbamidomethyl

**Score:** 170.0
**Seq. Coverage [%]:** 23.1
**No. of Peptides:** 3

| 10 | 20 | 30 | 40 | 50 | 60 | 70 | 80 | 90 | 100 |
|---|---|---|---|---|---|---|---|---|---|
| MELGLSWVFL | VAILEGVQCE | VQLVESGGGL | VQPGGSLRLS | CAASGFTFSS | YWMSWVRQAP | GKGLEWVANI | KQDGSEKYYV | DSVKGRFTIS | RDNAKNSLYL |
| 110 | 120 | | | | | | | | |
| QMNSLRAEDT | AVYYCAR | | | | | | | | |

MS/MS Peptide Matches

| Cmpd. | m/z meas. | z | Δ m/z [ppm] | Δ m/z [Da] | Rt [min] | Score | No. of Cmpds. | Site [%] | P | Sequence | Modification | Range | Type |
|---|---|---|---|---|---|---|---|---|---|---|---|---|---|
| 993 | 515.346 | 2 | 108.64 | 0.0560 | 28.86 | 63.25 | 2 | 0.0 | 0 | K.GLEWVANIK.Q | | 63-71 | CID |
| 513 | 437.227 | 2 | 13.11 | 0.0057 | 18.98 | 28.92 | 2 | 0.0 | 0 | K.YYVDSVK.G | | 78-84 | CID |
| 580 | 659.809 | 2 | 28.42 | 0.0187 | 20.33 | 77.8 | 3 | 0.0 | 0 | R.AEDTAVYYCAR.- | Carbamidomethyl: 9 | 107-117 | CID |

#### Protein 9: Immunoglobulin kappa variable 2D-28 OS=Homo sapiens GN=IGKV2D-28 PE=1 SV=2

**Accession:** KVD28_HUMAN
**Database:** SwissProt
**MW [kDa] / pI:** 12.9 / 5.6

**Score:** 166.4
**Seq. Coverage [%]:** 16.7
**No. of Peptides:** 2

| 10 | 20 | 30 | 40 | 50 | 60 | 70 | 80 | 90 | 100 |
|---|---|---|---|---|---|---|---|---|---|
| MRLPAQLLGL | LMLWVSGSSG | DIVMTQSPLS | LPVTPGEPAS | ISCRSSQSLL | HSNGYNYLDW | YLQKPGQSPQ | LLIYLGSNRA | SGVPDRFSGS | GSGTDFTLKI |
| 110 | 120 | 130 | | | | | | | |
| SRVEAEDVGV | YYCMQALQTP | | | | | | | | |

MS/MS Peptide Matches

| Cmpd. | m/z meas. | z | Δ m/z [ppm] | Δ m/z [Da] | Rt [min] | Score | No. of Cmpds. | Site [%] | P | Sequence | Modification | Range | Type |
|---|---|---|---|---|---|---|---|---|---|---|---|---|---|
| 799 | 662.661 | 3 | 6.11 | 0.0040 | 24.85 | 68.98 | 3 | 0.0 | 1 | R.ASGVPDRFSGSGSGTDFTLK.I | | 80-99 | CID |
| 747 | 652.281 | 2 | -47.33 | -0.0309 | 23.81 | 97.38 | 3 | 0.0 | 0 | R.FSGSGSGTDFTLK.I | | 87-99 | CID |

#### Protein 10: Serum albumin OS=Homo sapiens GN=ALB PE=1 SV=2

| | | | |
|---|---|---|---|
| **Accession:** | ALBU_HUMAN | **Score:** | 90.2 |
| **Database:** | SwissProt | **Seq. Coverage [%]:** | 6.6 |
| **MW [kDa] / pI:** | 69.3 / 5.9 | **No. of Peptides:** | 2 |
| **Modification(s):** | Carbamidomethyl | | |

| 10 | 20 | 30 | 40 | 50 | 60 | 70 | 80 | 90 | 100 |
|---|---|---|---|---|---|---|---|---|---|
| MKWVTFISLL | FLFSSAYSRG | VFRRDAHKSE | VAHRFKDLGE | ENFKALVLIA | FAQYLQQCPF | EDHVKLVNEV | TEFAKTCVAD | ESAENCDKSL | HTLFGDKLCT |
| **110** | **120** | **130** | **140** | **150** | **160** | **170** | **180** | **190** | **200** |
| VATLRETYGE | MADCCAKQEP | ERNECFLQHK | DDNPNLPRLV | RPEVDVMCTA | FHDNEETFLK | KYLYEIARRH | PYFYAPELLF | FAKRYKAAFT | ECCQAADKAA |
| **210** | **220** | **230** | **240** | **250** | **260** | **270** | **280** | **290** | **300** |
| CLLPKLDELR | DEGKASSAKQ | RLKCASLQKF | GERAFKAWAV | ARLSQRFPKA | EFAEVSKLVT | DLTKVHTECC | HGDLLECADD | RADLAKYICE | NQDSISSKLK |
| **310** | **320** | **330** | **340** | **350** | **360** | **370** | **380** | **390** | **400** |
| ECCEKPLLEK | SHCIAEVEND | EMPADLPSLA | ADFVESKDVC | KNYAEAKDVF | LGMFLYEYAR | RHPDYSVVLL | LRLAKTYETT | LEKCCAADPH | ECYAKVFDE |
| **410** | **420** | **430** | **440** | **450** | **460** | **470** | **480** | **490** | **500** |
| FKPLVEEPQN | LIKQNCELFE | QLGEYKFQNA | LLVRYTKKVP | QVSTPTLVEV | SRNLGKVGSK | CCKHPEAKRM | PCAEDYLSVV | LNQLCVLHEK | TPVSDRVTKC |
| **510** | **520** | **530** | **540** | **550** | **560** | **570** | **580** | **590** | **600** |
| CTESLVNRRP | CFSALEVDET | YVPKEFNAET | FTFHADICTL | SEKERQIKKQ | TALVELVKHK | PKATKEQLKA | VMDDFAAFVE | KCCKADDKET | CFAEEGKKLV |
| **610** | | | | | | | | | |
| AASQAALGL | | | | | | | | | |

MS/MS Peptide Matches

| Cmpd. | m/z meas. | z | Δ m/z [ppm] | Δ m/z [Da] | Rt [min] | Score | No. of Cmpds. | Site [%] | P | Sequence | Modification | Range | Type |
|---|---|---|---|---|---|---|---|---|---|---|---|---|---|
| 2310 | 992.087 | 3 | -32.94 | -0.0327 | 57.11 | 24.92 | 1 | 0.0 | 0 | K.SHCIAEVENDEMPADLPSLAADFVESK.D | Carbamidomethyl: 3 | 311-337 | CID |
| 1543 | 812.5 | 2 | 126.26 | 0.1026 | 39.30 | 65.25 | 2 | 0.0 | 0 | K.DVFLGMFLYEYAR.R | | 348-360 | CID |

#### Protein 11: Alpha-1-antitrypsin OS=Homo sapiens GN=SERPINA1 PE=1 SV=3

**Accession:** A1AT_HUMAN
**Database:** SwissProt
**MW [kDa] / pI:** 46.7 / 5.4
**Score:** 80.2
**Seq. Coverage [%]:** 4.1
**No. of Peptides:** 1

| 10 | 20 | 30 | 40 | 50 | 60 | 70 | 80 | 90 | 100 |
|---|---|---|---|---|---|---|---|---|---|
| MPSSVSWGIL | LLAGLCCLVP | VSLAEDPQGD | AAQKTDTSHH | DQDHPTFNKI | TPNLAEFAFS | LYRQLAHQSN | STNIFFSPVS | IATAFAMLSL | GTKADTHDEI |
| 110 | 120 | 130 | 140 | 150 | 160 | 170 | 180 | 190 | 200 |
| LEGLNFNLTE | IPEAQIHEGF | QELLRTLNQP | DSQLQLTTGN | GLFLSEGLKL | VDKFLEDVKK | LYHSEAFTVN | FGDTEEAKKQ | INDYVEKGTQ | GKIVDLVKEL |
| 210 | 220 | 230 | 240 | 250 | 260 | 270 | 280 | 290 | 300 |
| DRDTVFALVN | YIFFKGKWER | PFEVKDTEEE | DFHVDQVTTV | KVPMMKRLGM | FNIQHCKKLS | SWVLLMKYLG | NATAIFFLPD | EGKLQHLENE | LTHDIITKFL |
| 310 | 320 | 330 | 340 | 350 | 360 | 370 | 380 | 390 | 400 |
| ENEDRRSASL | HLPKLSITGT | YDLKSVLGQL | GITKVFSNGA | DLSGVTEEAP | LKLSKAVHKA | VLTIDEKGTE | AAGAMFLEAI | PMSIPPEVKF | NKPFVFLMIE |
| 410 | 420 | | | | | | | | |
| QNTKSPLFMG | KVVNPTQK | | | | | | | | |

MS/MS Peptide Matches

| Cmpd. | m/z meas. | z | Δ m/z [ppm] | Δ m/z [Da] | Rt [min] | Score | No. of Cmpds. | Site [%] | P | Sequence | Modification | Range | Type |
|---|---|---|---|---|---|---|---|---|---|---|---|---|---|
| 1672 | 697.313 | 3 | -81.86 | -0.0571 | 41.92 | 80.21 | 1 | 0.0 | 1 | K.ELDRDTVFALVNYIFFK.G | | 199-215 | CID |

#### Protein 12: Immunoglobulin kappa variable 3-20 OS=Homo sapiens GN=IGKV3-20 PE=1 SV=2

**Accession:** KV320_HUMAN
**Database:** SwissProt
**MW [kDa] / pI:** 12.5 / 4.9
**Score:** 65.5
**Seq. Coverage [%]:** 7.8
**No. of Peptides:** 1

| 10 | 20 | 30 | 40 | 50 | 60 | 70 | 80 | 90 | 100 |
|---|---|---|---|---|---|---|---|---|---|
| METPAQLLFL | LLLWLPDTTG | EIVLTQSPGT | LSLSPGERAT | LSCRASQSVS | SSYLAWYQQK | PGQAPRLLIY | GASSRATGIP | DRFSGSGSGT | DFTLTISRLE |
| 110 | 120 | | | | | | | | |
| PEDFAVYYCQ | QYGSSP | | | | | | | | |

MS/MS Peptide Matches

| Cmpd. | m/z meas. | z | Δ m/z [ppm] | Δ m/z [Da] | Rt [min] | Score | No. of Cmpds. | Site [%] | P | Sequence | Modification | Range | Type |
|---|---|---|---|---|---|---|---|---|---|---|---|---|---|
| 756 | 490.342 | 2 | 121.99 | 0.0598 | 23.96 | 65.51 | 3 | 0.0 | 0 | R.LLIYGASSR.A | | 67-75 | CID |

#### Protein 13: Immunoglobulin kappa variable 3-15 OS=Homo sapiens GN=IGKV3-15 PE=1 SV=2

**Accession:** KV315_HUMAN
**Database:** SwissProt
**MW [kDa] / pI:** 12.5 / 5.1
**Score:** 65.2
**Seq. Coverage [%]:** 7.8
**No. of Peptides:** 1

| 10 | 20 | 30 | 40 | 50 | 60 | 70 | 80 | 90 | 100 |
|---|---|---|---|---|---|---|---|---|---|
| MEAPAQLLFL | LLLWLPDTTG | EIVMTQSPAT | LSVSPGERAT | LSCRASQSVS | SNLAWYQQKP | GQAPRLLIYG | ASTRATGIPA | RFSGSGSGTE | FTLTISSLQS |
| 110 | 120 | | | | | | | | |
| EDFAVYYCQQ | YNNWP | | | | | | | | |

MS/MS Peptide Matches

| Cmpd. | m/z meas. | z | Δ m/z [ppm] | Δ m/z [Da] | Rt [min] | Score | No. of Cmpds. | Site [%] | P | Sequence | Modification | Range | Type |
|---|---|---|---|---|---|---|---|---|---|---|---|---|---|
| 772 | 497.336 | 2 | 92.47 | 0.0460 | 24.31 | 65.18 | 4 | 0.0 | 0 | R.LLIYGASTR.A | | 66-74 | CID |

#### Protein 14: Immunoglobulin kappa variable 3-11 OS=Homo sapiens GN=IGKV3-11 PE=1 SV=1

**Accession:** KV311_HUMAN
**Database:** SwissProt
**MW [kDa] / pI:** 12.6 / 4.9
**Score:** 45.6
**Seq. Coverage [%]:** 26.1
**No. of Peptides:** 1

**Modification(s):** Carbamidomethyl

```
        10         20         30         40         50         60         70         80         90        100
MEAPAQLLFL LLLWLPDTTG EIVLTQSPAT LSLSPGERAT LSCRASQSVS SYLAWYQQKP GQAPRLLIYD ASNRATGIPA RFSGSGSGTD FTLTISSLEP
       110        120
EDFAVYYCQQ RSNWP
```

MS/MS Peptide Matches

| Cmpd. | m/z meas. | z | Δ m/z [ppm] | Δ m/z [Da] | Rt [min] | Score | No. of Cmpds. | Site [%] | P | Sequence | Modification | Range | Type |
|---|---|---|---|---|---|---|---|---|---|---|---|---|---|
| 1369 | 1121.801 | 3 | 259.37 | 0.2909 | 36.13 | 45.57 | 4 | 0.0 | 0 | R.FSGSGSGTDFTLTISSLEPEDFAVYYCQQR.S | Carbamidomethyl: 27 | 82-111 | CID |

#### Protein 15: Lysosomal-trafficking regulator OS=Homo sapiens GN=LYST PE=1 SV=3

**Accession:** LYST_HUMAN
**Database:** SwissProt
**MW [kDa] / pI:** 428.9 / 6.2

**Score:** 41.4
**Seq. Coverage [%]:** 0.2
**No. of Peptides:** 1

| | | | | | | | | | |
|---|---|---|---|---|---|---|---|---|---|
| 10 | 20 | 30 | 40 | 50 | 60 | 70 | 80 | 90 | 100 |
| MSTDSNSLAR | EFLTDVNRLC | NAVVQRVEAR | EEEEEETHMA | TLGQYLVHGR | GFLLLTKLNS | IIDQALTCRE | ELLTLLLSLL | PLVWKIPVQE | EKATDFNLPL |
| 110 | 120 | 130 | 140 | 150 | 160 | 170 | 180 | 190 | 200 |
| SADIILTKEK | NSSSQRSTQE | KLHLEGSALS | SQVSAKVNVF | RKSRRQRKIT | HRYSVRDARK | TQLSTSDSEA | NSDEKGIAMN | KHRRPHLLHH | FLTSFPKQDH |
| 210 | 220 | 230 | 240 | 250 | 260 | 270 | 280 | 290 | 300 |
| PKAKLDRLAT | KEQTPPDAMA | LENSREIIPR | QGSNTDILSE | PAALSVISNM | NNSPFDLCHV | LLSLLEKVCK | FDVTLNHNSP | LAASVVPTLT | EFLAGFGDCC |
| 310 | 320 | 330 | 340 | 350 | 360 | 370 | 380 | 390 | 400 |
| SLSDNLESRV | VSAGWTEEPV | ALIQRMLFRT | VLHLLSVDVS | TAEMMPENLR | KNLTELLRAA | LKIRICLEKQ | PDPFAPRQKK | TLQEVQEDFV | FSKYRHRALL |
| 410 | 420 | 430 | 440 | 450 | 460 | 470 | 480 | 490 | 500 |
| LPELLEGVLQ | ILICCLQSAA | SNPFYFSQAM | DLVQEFIQHH | GFNLFETAVL | QMEWLVLRDG | VPPEASEHLK | ALINSVMKIM | STVKKVKSEQ | LHHSMCTRKR |
| 510 | 520 | 530 | 540 | 550 | 560 | 570 | 580 | 590 | 600 |
| HRRCEYSHFM | HHHRDLSGLL | VSAFKNQVSK | NPFEETADGD | VYYPERCCCI | AVCAHQCLRL | LQQASLSSTC | VQILSGVHNI | GICCCMDPKS | VIIPLLHAFK |
| 610 | 620 | 630 | 640 | 650 | 660 | 670 | 680 | 690 | 700 |
| LPALKNFQQH | ILNILNKLIL | DQLGGAEISP | KIKKAACNIC | TVDSDQLAQL | EETLQGNLCD | AELSSSLSSP | SYRFQGILPS | SGSEDLLWKW | DALKAYQNFV |
| 710 | 720 | 730 | 740 | 750 | 760 | 770 | 780 | 790 | 800 |
| FEEDRLHSIQ | IANHICNLIQ | KGNIVVQWKL | YNYIFNPVLQ | RGVELAHHCQ | HLSVTSAQSH | VCSHHNQCLP | QDVLQIYVKT | LPILLKSRVI | RDLFLSCNGV |
| 810 | 820 | 830 | 840 | 850 | 860 | 870 | 880 | 890 | 900 |
| SQIIELNCLN | GIRSHSLKAF | ETLIISLGEQ | QKDASVPDID | GIDIEQKELS | SVHVGTSFHH | QQAYSDSPQS | LSKFYAGLKE | AYPKRRKTVN | QDVHINTINL |
| 910 | 920 | 930 | 940 | 950 | 960 | 970 | 980 | 990 | 1000 |
| FLCVAFLCVS | KEAESDRESA | NDSEDTSGYD | STASEPLSHM | LPCISLESLV | LPSPEHMHQA | ADIWSMCRWI | YMLSSVFQKQ | FYRLGGFRVC | HKLIFMIIQK |
| 1010 | 1020 | 1030 | 1040 | 1050 | 1060 | 1070 | 1080 | 1090 | 1100 |
| LFRSHKEEQG | KKEGDTSVNE | NQDLNRISQP | KRTMKEDLLS | LAIKSDPIPS | ELGSLKKSAD | SLGKLELQHI | SSINVEEVSA | TEAAPEEAKL | FTSQESETSL |
| 1110 | 1120 | 1130 | 1140 | 1150 | 1160 | 1170 | 1180 | 1190 | 1200 |
| QSIRLLEALL | AICLHGARTS | QQKMELELPN | QNLSVESILF | EMRDHLSQSK | VIETQLAKPL | FDALLRVALG | NYSADFEHND | AMTEKSHQSA | EELSSQPGDF |
| 1210 | 1220 | 1230 | 1240 | 1250 | 1260 | 1270 | 1280 | 1290 | 1300 |
| SEEAEDSQCC | SFKLLVEEEG | YEADSESNPE | DGETQDDGVD | LKSETEGFSA | SSSPNDLLEN | LTQGEIIYPE | ICMLELNLLS | ASKAKLDVLA | HVFESFLKII |
| [illegible] | [illegible] | [illegible] | [illegible] | [illegible] | [illegible] | [illegible] | [illegible] | [illegible] | [illegible] |

MS/MS Peptide Matches

| Cmpd. | m/z meas. | z | Δ m/z [ppm] | Δ m/z [Da] | Rt [min] | Score | No. of Cmpds. | Site [%] | P | Sequence | Modification | Range | Type |
|---|---|---|---|---|---|---|---|---|---|---|---|---|---|
| 625 | 443.309 | 2 | 66.66 | 0.0295 | 21.20 | 41.35 | 3 | 0.0 | 0 | R.LTVLAVNR.I | | 2646-2653 | CID |

#### Protein 16: Immunoglobulin heavy constant alpha 1 OS=Homo sapiens GN=IGHA1 PE=1 SV=2

| | | | |
|---|---|---|---|
| **Accession:** | IGHA1_HUMAN | **Score:** | 40.2 |
| **Database:** | SwissProt | **Seq. Coverage [%]:** | 3.4 |
| **MW [kDa] / pI:** | 37.6 / 6.1 | **No. of Peptides:** | 1 |
| **Modification(s):** | Carbamidomethyl | | |

| 10 | 20 | 30 | 40 | 50 | 60 | 70 | 80 | 90 | 100 |
|---|---|---|---|---|---|---|---|---|---|
| ASPTSPKVFP | LSLCSTQPDG | NVVIACLVQG | FFPQEPLSVT | WSESGQGVTA | RNFPPSQDAS | GDLYTTSSQL | TLPATQCLAG | KSVTCHVKHY | TNPSQDVTVP |
| 110 | 120 | 130 | 140 | 150 | 160 | 170 | 180 | 190 | 200 |
| CPVPSTPPTP | SPSTPPTPSP | SCCHPRLSLH | RPALEDLLLG | SEANLTCTLT | GLRDASGVTF | TWTPSSGKSA | VQGPPERDLC | GCYSVSSVLP | GCAEPWNHGK |
| 210 | 220 | 230 | 240 | 250 | 260 | 270 | 280 | 290 | 300 |
| TFTCTAAYPE | SKTPLTATLS | KSGNTFRPEV | HLLPPPSEEL | ALNELVTLTC | LARGFSPKDV | LVRWLQGSQE | LPREKYLTWA | SRQEPSQGTT | TFAVTSILRV |
| 310 | 320 | 330 | 340 | 350 | 360 | | | | |
| AAEDWKKGDT | FSCMVGHEAL | PLAFTQKTID | RLAGKPTHVN | VSVVMAEVDG | TCY | | | | |

MS/MS Peptide Matches

| Cmpd. | m/z meas. | z | Δ m/z [ppm] | Δ m/z [Da] | Rt [min] | Score | No. of Cmpds. | Site [%] | P | Sequence | Modification | Range | Type |
|---|---|---|---|---|---|---|---|---|---|---|---|---|---|
| 590 | 688.332 | 2 | 26.79 | 0.0184 | 20.51 | 40.2 | 1 | 0.0 | 0 | K.TFTCTAAYPESK.T | Carbamidomethyl: 4 | 201-212 | CID |

#### Protein 17: Immunoglobulin heavy variable 3-73 OS=Homo sapiens GN=IGHV3-73 PE=3 SV=1

| | | | |
|---|---|---|---|
| **Accession:** | HV373_HUMAN | **Score:** | 37.6 |
| **Database:** | SwissProt | **Seq. Coverage [%]:** | 5.9 |
| **MW [kDa] / pI:** | 12.8 / 9.3 | **No. of Peptides:** | 1 |

| 10 | 20 | 30 | 40 | 50 | 60 | 70 | 80 | 90 | 100 |
|---|---|---|---|---|---|---|---|---|---|
| MEFGLSWVFL | VAILKGVQCE | VQLVESGGGL | VQPGGSLKLS | CAASGFTFSG | SAMHWVRQAS | GKGLEWVGRI | RSKANSYATA | YAASVKGRFT | ISRDDSKNTA |
| 110 | 120 | | | | | | | | |
| YLQMNSLKTE | DTAVYYCTR | | | | | | | | |

MS/MS Peptide Matches

| Cmpd. | m/z meas. | z | Δ m/z [ppm] | Δ m/z [Da] | Rt [min] | Score | No. of Cmpds. | Site [%] | P | Sequence | Modification | Range | Type |
|---|---|---|---|---|---|---|---|---|---|---|---|---|---|
| 777 | 408.751 | 2 | 71.52 | 0.0292 | 24.42 | 37.57 | 3 | 0.0 | 0 | K.GLEWVGR.I | | 63-69 | CID |

#### Project Info

| | | | |
|---|---|---|---|
| **Name:** | Alagesan | **Date:** | Mar 5, 2019 |
| **Note:** | Nanobooster | | |

#### Sample Info & Protocols

| | |
|---|---|
| **Name:** | Nanobooster - IgG |
| **Date:** | Mar 5, 2019 |

#### Search Result Info

| | | | |
|---|---|---|---|
| **Search Result:** | IonTrap_Human_Swiss_CID_2019-03-05 17:55:08 | | |
| **Location:** | /Alagesan/Nanobooster - IgG/Ka_IgG Trypsin NoEnrich NB ACN_BA5_01_8740.mgf | | |
| **Search Method:** | IonTrap_Human_Swiss_CID | **Version:** | 1.2 |
| **Search Engine(s):** | Mascot, 2.6.0 | **Ident. Compound(s):** | 0/2189 |
| **Database(s):** | SwissProt, SwissProt_2017_04.fasta | | |
| **Note:** | | | |

#### Results

##### Protein 1: Immunoglobulin heavy constant gamma 1 OS=Homo sapiens GN=IGHG1 PE=1 SV=1

| | | | |
|---|---|---|---|
| **Accession:** | IGHG1_HUMAN | **Score:** | 968.1 |
| **Database:** | SwissProt | **Seq. Coverage [%]:** | 54.8 |
| **MW [kDa] / pI:** | 36.1 / 8.5 | **No. of Peptides:** | 14 |
| **Modification(s):** | Carbamidomethyl, Oxidation | | |

| 10 | 20 | 30 | 40 | 50 | 60 | 70 | 80 | 90 | 100 |
|---|---|---|---|---|---|---|---|---|---|
| ASTKGPSVFP | LAPSSKSTSG | GTAALGCLVK | DYFPEPVTVS | WNSGALTSGV | HTFPAVLQSS | GLYSLSSVVT | VPSSSLGTQT | YICNVNHKPS | NTKVDKKVEP |
| **110** | **120** | **130** | **140** | **150** | **160** | **170** | **180** | **190** | **200** |
| KSCDKTHTCP | PCPAPELLGG | PSVFLFPPKP | KDTLMISRTP | EVTCVVVDVS | HEDPEVKFNW | YVDGVEVHNA | KTKPREEQYN | STYRVVSVLT | VLHQDWLNGK |
| **210** | **220** | **230** | **240** | **250** | **260** | **270** | **280** | **290** | **300** |
| EYKCKVSNKA | LPAPIEKTIS | KAKGQPREPQ | VYTLPPSRDE | LTKNQVSLTC | LVKGFYPSDI | AVEWESNGQP | ENNYKTTPPV | LDSDGSFFLY | SKLTVDKSRW |
| **310** | **320** | **330** | **340** | | | | | | |
| QQGNVFSCSV | MHEALHNHYT | QKSLSLSPGK | | | | | | | |

**MS/MS Peptide Matches**

| Cmpd. | m/z meas. | z | Δ m/z [ppm] | Δ m/z [Da] | Rt [min] | Score | No. of Cmpds. | Site [%] | P | Sequence | Modification | Range | Type |
|---|---|---|---|---|---|---|---|---|---|---|---|---|---|
| 761 | 593.836 | 2 | 15.22 | 0.0090 | 27.79 | 59.99 | 13 | 0.0 | 0 | K.GPSVFPLAPSSK.S | | 5-16 | CID |
| 657 | 661.384 | 2 | 62.52 | 0.0413 | 25.48 | 112.72 | 13 | 0.0 | 0 | K.STSGGTAALGCLVK.D | Carbamidomethyl: 11 | 17-30 | CID |
| 1046 | 712.176 | 4 | 430.08 | 0.3062 | 33.18 | 63.81 | 13 | 0.0 | 0 | K.THTCPPCPAPELLGGPSVFLFPPKPK.D | Carbamidomethyl: 4, 7 | 106-131 | CID |
| 534 | 418.267 | 2 | 110.60 | 0.0463 | 22.88 | 50.66 | 13 | 0.0 | 0 | K.DTLMISR.T | | 132-138 | CID |
| 389 | 426.263 | 2 | 105.10 | 0.0448 | 20.00 | 53.61 | 3 | 0.0 | 0 | K.DTLMISR.T | Oxidation: 4 | 132-138 | CID |
| 745 | 713.726 | 3 | 63.53 | 0.0453 | 27.48 | 105.07 | 3 | 0.0 | 0 | R.TPEVTCVVVDVSHEDPEVK.F | Carbamidomethyl: 6 | 139-157 | CID |
| 747 | 560.012 | 3 | 130.65 | 0.0732 | 27.53 | 72.51 | 17 | 0.0 | 0 | K.FNWYVDGVEVHNAK.T | | 158-171 | CID |
| 1118 | 603.398 | 3 | 95.55 | 0.0577 | 34.67 | 63.61 | 12 | 0.0 | 0 | R.VVSVLTVLHQDWLNGK.E | | 185-200 | CID |

| Cmpd. | m/z meas. | z | Δ m/z [ppm] | Δ m/z [Da] | Rt [min] | Score | No. of Cmpds. | Site [%] | P | Sequence | Modification | Range | Type |
|---|---|---|---|---|---|---|---|---|---|---|---|---|---|
| 477 | 419.805 | 2 | 118.45 | 0.0497 | 21.72 | 21.74 | 3 | 0.0 | 0 | K.ALPAPIEK.T | | 210-217 | CID |
| 582 | 643.797 | 2 | -67.72 | -0.0436 | 23.91 | 52.57 | 14 | 0.0 | 0 | R.EPQVYTLPPSR.D | | 228-238 | CID |
| 594 | 625.019 | 3 | 38.55 | 0.0241 | 24.16 | 28.95 | 14 | 0.0 | 1 | R.EPQVYTLPPSRDELTK.N | | 228-243 | CID |
| 721 | 581.365 | 2 | 80.08 | 0.0466 | 26.99 | 76.52 | 10 | 0.0 | 0 | K.NQVSLTCLVK.G | Carbamidomethyl: 7 | 244-253 | CID |
| 947 | 848.794 | 3 | 92.72 | 0.0787 | 31.31 | 98.77 | 26 | 0.0 | 0 | K.GFYPSDIAVEWESNGQPENNYK.T | | 254-275 | CID |
| 1036 | 937.507 | 2 | 45.28 | 0.0425 | 32.97 | 107.61 | 11 | 0.0 | 0 | K.TTPPVLDSDGSFFLYSK.L | | 276-292 | CID |

#### Protein 2: Immunoglobulin heavy constant gamma 3 OS=Homo sapiens GN=IGHG3 PE=1 SV=2

**Accession:** IGHG3_HUMAN
**Database:** SwissProt
**MW [kDa] / pI:** 41.3 / 8.2
**Modification(s):** Carbamidomethyl, Oxidation

**Score:** 792.9
**Seq. Coverage [%]:** 36.3
**No. of Peptides:** 13

| 10 | 20 | 30 | 40 | 50 | 60 | 70 | 80 | 90 | 100 |
|---|---|---|---|---|---|---|---|---|---|
| ASTKGPSVFP | LAPCSRSTSG | GTAALGCLVK | DYFPEPVTVS | WNSGALTSGV | HTFPAVLQSS | GLYSLSSVVT | VPSSSLGTQT | YTCNVNHKPS | NTKVDKRVEL |
| **110** | **120** | **130** | **140** | **150** | **160** | **170** | **180** | **190** | **200** |
| KTPLGDTTHT | CPRCPEPKSC | DTPPPCPRCP | EPKSCDTPPP | CPRCPEPKSC | DTPPPCPRCP | APELLGGPSV | FLFPPKPKDT | LMISRTPEVT | CVVVDVSHED |
| **210** | **220** | **230** | **240** | **250** | **260** | **270** | **280** | **290** | **300** |
| PEVQFKWYVD | GVEVHNAKTK | PREEQYNSTF | RVVSVLTVLH | QDWLNGKEYK | CKVSNKALPA | PIEKTISKTK | GQPREPQVYT | LPPSREEMTK | NQVSLTCLVK |
| **310** | **320** | **330** | **340** | **350** | **360** | **370** | **380** | | |
| GFYPSDIAVE | WESSGQPENN | YNTTPPMLDS | DGSFFLYSKL | TVDKSRWQQG | NIFSCSVMHE | ALHNRFTQKS | LSLSPGK | | |

MS/MS Peptide Matches

| Cmpd. | m/z meas. | z | Δ m/z [ppm] | Δ m/z [Da] | Rt [min] | Score | No. of Cmpds. | Site [%] | P | Sequence | Modification | Range | Type |
|---|---|---|---|---|---|---|---|---|---|---|---|---|---|
| 796 | 644.374 | 2 | 69.30 | 0.0447 | 28.51 | 69.42 | 13 | 0.0 | 0 | K.GPSVFPLAPCSR.S | Carbamidomethyl: 10 | 5-16 | CID |

| Cmpd. | m/z meas. | z | Δ m/z [ppm] | Δ m/z [Da] | Rt [min] | Score | No. of Cmpds. | Site [%] | P | Sequence | Modification | Range | Type |
|---|---|---|---|---|---|---|---|---|---|---|---|---|---|
| 657 | 661.384 | 2 | 62.52 | 0.0413 | 25.48 | 112.72 | 13 | 0.0 | 0 | R.STSGGTAALGCLVK.D | Carbamidomethyl: 11 | 17-30 | CID |
| 243 | 452.558 | 3 | 16.37 | 0.0074 | 17.11 | 52.33 | 6 | 0.0 | 0 | K.TPLGDTTHTCPR.C | Carbamidomethyl: 10 | 102-113 | CID |
| 214 | 593.778 | 2 | 42.75 | 0.0254 | 16.45 | 59.68 | 9 | 0.0 | 0 | K.SCDTPPPCPR.C | Carbamidomethyl: 2, 8 | 119-128 | CID |
| 1115 | 717.777 | 3 | 69.36 | 0.0498 | 34.62 | 78.23 | 3 | 0.0 | 0 | R.CPAPELLGGPSVFLFPPKPK.D | Carbamidomethyl: 1 | 159-178 | CID |
| 534 | 418.267 | 2 | 110.60 | 0.0463 | 22.88 | 50.66 | 13 | 0.0 | 0 | K.DTLMISR.T | | 179-185 | CID |
| 389 | 426.263 | 2 | 105.10 | 0.0448 | 20.00 | 53.61 | 3 | 0.0 | 0 | K.DTLMISR.T | Oxidation: 4 | 179-185 | CID |
| 590 | 472.966 | 3 | 135.90 | 0.0643 | 24.08 | 54.04 | 5 | 0.0 | 0 | K.WYVDGVEVHNAK.T | | 207-218 | CID |
| 1118 | 603.398 | 3 | 95.55 | 0.0577 | 34.67 | 63.61 | 12 | 0.0 | 0 | R.VVSVLTVLHQDWLNGK.E | | 232-247 | CID |
| 477 | 419.805 | 2 | 118.45 | 0.0497 | 21.72 | 21.74 | 3 | 0.0 | 0 | K.ALPAPIEK.T | | 257-264 | CID |
| 582 | 643.797 | 2 | -67.72 | -0.0436 | 23.91 | 52.57 | 14 | 0.0 | 0 | R.EPQVYTLPPSR.E | | 275-285 | CID |
| 608 | 635.667 | 3 | 23.18 | 0.0147 | 24.44 | 47.78 | 28 | 0.0 | 1 | R.EPQVYTLPPSREEMTK.N | | 275-290 | CID |
| 721 | 581.365 | 2 | 80.08 | 0.0466 | 26.99 | 76.52 | 10 | 0.0 | 0 | K.NQVSLTCLVK.G | Carbamidomethyl: 7 | 291-300 | CID |

#### Protein 3: Immunoglobulin kappa constant OS=Homo sapiens GN=IGKC PE=1 SV=2

**Accession:** IGKC_HUMAN
**Database:** SwissProt
**MW [kDa] / pI:** 11.8 / 6.1
**Modification(s):** Carbamidomethyl

**Score:** 766.0
**Seq. Coverage [%]:** 86.9
**No. of Peptides:** 8

```
        10         20         30         40         50         60         70         80         90        100
RTVAAPSVFI FPPSDEQLKS GTASVVCLLN NFYPREAKVQ WKVDNALQSG NSQESVTEQD SKDSTYSLSS TLTLSKADYE KHKVYACEVT HQGLSSPVTK
       110
SFNRGEC
```

MS/MS Peptide Matches

| Cmpd. | m/z meas. | z | Δ m/z [ppm] | Δ m/z [Da] | Rt [min] | Score | No. of Cmpds. | Site [%] | P | Sequence | Modification | Range | Type |
|---|---|---|---|---|---|---|---|---|---|---|---|---|---|
| 951 | 701.404 | 3 | 32.97 | 0.0231 | 31.36 | 34.2 | 8 | 0.0 | 1 | -.RTVAAPSVFIFPPSDEQLK.S | | 1-19 | CID |
| 1088 | 973.606 | 2 | 91.30 | 0.0889 | 34.10 | 89.45 | 19 | 0.0 | 0 | R.TVAAPSVFIFPPSDEQLK.S | | 2-19 | CID |
| 1172 | 600.046 | 3 | 126.79 | 0.0761 | 35.76 | 90.43 | 10 | 0.0 | 0 | K.SGTASVVCLLNNFYPR.E | Carbamidomethyl: 8 | 20-35 | CID |
| 348 | 1068.474 | 2 | -13.10 | -0.0140 | 19.14 | 155.06 | 14 | 0.0 | 0 | K.VDNALQSGNSQESVTEQDSK.D | | 43-62 | CID |
| 737 | 1207.537 | 3 | 244.94 | 0.2957 | 27.32 | 138.1 | 4 | 0.0 | 1 | K.VDNALQSGNSQESVTEQDSKDSTYSLSSTLTLSK.A | | 43-76 | CID |
| 779 | 751.961 | 2 | 103.93 | 0.0781 | 28.13 | 95.16 | 9 | 0.0 | 0 | K.DSTYSLSSTLTLSK.A | | 63-76 | CID |
| 482 | 626.223 | 3 | 387.40 | 0.2425 | 21.86 | 109.37 | 11 | 0.0 | 0 | K.VYACEVTHQGLSSPVTK.S | Carbamidomethyl: 4 | 84-100 | CID |
| 106 | 435.168 | 2 | -32.52 | -0.0142 | 14.09 | 54.21 | 8 | 0.0 | 1 | K.SFNRGEC.- | Carbamidomethyl: 7 | 101-107 | CID |

#### Protein 4: Immunoglobulin heavy constant gamma 2 OS=Homo sapiens GN=IGHG2 PE=1 SV=2

**Accession:** IGHG2_HUMAN
**Database:** SwissProt
**MW [kDa] / pI:** 35.9 / 7.7
**Modification(s):** Carbamidomethyl, Oxidation

**Score:** 742.4
**Seq. Coverage [%]:** 39.0
**No. of Peptides:** 13

| 10 | 20 | 30 | 40 | 50 | 60 | 70 | 80 | 90 | 100 |
|---|---|---|---|---|---|---|---|---|---|
| ASTKGPSVFP | LAPCSRSTSE | STAALGCLVK | DYFPEPVTVS | WNSGALTSGV | HTFPAVLQSS | GLYSLSSVVT | VPSSNFGTQT | YTCNVDHKPS | NTKVDKTVER |
| **110** | **120** | **130** | **140** | **150** | **160** | **170** | **180** | **190** | **200** |
| KCCVECPPCP | APPVAGPSVF | LFPPKPKDTL | MISRTPEVTC | VVVDVSHEDP | EVQFNWYVDG | VEVHNAKTKP | REEQFNSTFR | VVSVLTVVHQ | DWLNGKEYKC |
| **210** | **220** | **230** | **240** | **250** | **260** | **270** | **280** | **290** | **300** |
| KVSNKGLPAP | IEKTISKTKG | QPREPQVYTL | PPSREEMTKN | QVSLTCLVKG | FYPSDISVEW | ESNGQPENNY | KTTPPMLDSD | GSFFLYSKLT | VDKSRWQQGN |
| **310** | **320** | **330** | | | | | | | |
| VFSCSVMHEA | LHNHYTQKSL | SLPGK | | | | | | | |

MS/MS Peptide Matches

| Cmpd. | m/z meas. | z | Δ m/z [ppm] | Δ m/z [Da] | Rt [min] | Score | No. of Cmpds. | Site [%] | P | Sequence | Modification | Range | Type |
|---|---|---|---|---|---|---|---|---|---|---|---|---|---|
| 796 | 644.374 | 2 | 69.30 | 0.0447 | 28.51 | 69.42 | 13 | 0.0 | 0 | K.GPSVFPLAPCSR.S | Carbamidomethyl: 10 | 5-16 | CID |
| 677 | 712.393 | 2 | 48.43 | 0.0345 | 25.96 | 91.56 | 10 | 0.0 | 0 | R.STSESTAALGCLVK.D | Carbamidomethyl: 11 | 17-30 | CID |
| 935 | 760.2 | 4 | 421.62 | 0.3204 | 31.10 | 46.73 | 2 | 0.0 | 1 | R.KCCVECPPCPAPPVAGPSVFLFPPKPK.D | Carbamidomethyl: 2, 3, 6, 9 | 101-127 | CID |
| 1023 | 970.51 | 3 | 382.68 | 0.3713 | 32.74 | 60.63 | 6 | 0.0 | 0 | K.CCVECPPCPAPPVAGPSVFLFPPKPK.D | Carbamidomethyl: 1, 2, 5, 8 | 102-127 | CID |
| 534 | 418.267 | 2 | 110.60 | 0.0463 | 22.88 | 50.66 | 13 | 0.0 | 0 | K.DTLMISR.T | | 128-134 | CID |
| 389 | 426.263 | 2 | 105.10 | 0.0448 | 20.00 | 53.61 | 3 | 0.0 | 0 | K.DTLMISR.T | Oxidation: 4 | 128-134 | CID |
| 1007 | 599.054 | 3 | 643.99 | 0.3855 | 32.42 | 37.52 | 3 | 0.0 | 0 | R.VVSVLTVVHQDWLNGK.E | | 181-196 | CID |
| 494 | 412.769 | 2 | 52.20 | 0.0215 | 22.08 | 19.78 | 1 | 0.0 | 0 | K.GLPAPIEK.T | | 206-213 | CID |
| 582 | 643.797 | 2 | -67.72 | -0.0436 | 23.91 | 52.57 | 14 | 0.0 | 0 | R.EPQVYTLPPSR.E | | 224-234 | CID |
| 608 | 635.667 | 3 | 23.18 | 0.0147 | 24.44 | 47.78 | 28 | 0.0 | 1 | R.EPQVYTLPPSREEMTK.N | | 224-239 | CID |
| 721 | 581.365 | 2 | 80.08 | 0.0466 | 26.99 | 76.52 | 10 | 0.0 | 0 | K.NQVSLTCLVK.G | Carbamidomethyl: 7 | 240-249 | CID |
| 1034 | 953.576 | 2 | 131.54 | 0.1254 | 32.96 | 98.4 | 8 | 0.0 | 0 | K.TTPPMLDSDGSFFLYSK.L | | 272-288 | CID |
| 934 | 961.448 | 2 | -0.04 | 0.0000 | 31.06 | 37.19 | 3 | 0.0 | 0 | K.TTPPMLDSDGSFFLYSK.L | Oxidation: 5 | 272-288 | CID |

#### Protein 5: Immunoglobulin kappa light chain OS=Homo sapiens PE=1 SV=1

**Accession:** IGK_HUMAN
**Database:** SwissProt
**MW [kDa] / pI:** 23.4 / 6.9
**Modification(s):** Carbamidomethyl

**Score:** 696.5
**Seq. Coverage [%]:** 39.7
**No. of Peptides:** 7

| 10 | 20 | 30 | 40 | 50 | 60 | 70 | 80 | 90 | 100 |
|---|---|---|---|---|---|---|---|---|---|
| DIQMTQSPST | LSASVGDRVT | ITCRASQSIN | TWLAWYQQKP | GKAPKLLMYK | ASSLESGVPS | RFIGSGSGTE | FTLTISSLQP | DDFATYYCQQ | YNSDSKMFGQ |
| **110** | **120** | **130** | **140** | **150** | **160** | **170** | **180** | **190** | **200** |
| GTKVEVKGTV | AAPSVFIFPP | SDEQLKSGTA | SVVCLLNNFY | PREAKVQWKV | DNALQSGNSQ | ESVTEQDSKD | STYSLSSTLT | LSKADYEKHK | VYACEVTHQG |
| **210** | **220** | | | | | | | | |
| LSSPVTKSFN | RGEC | | | | | | | | |

MS/MS Peptide Matches

| Cmpd. | m/z meas. | z | Δ m/z [ppm] | Δ m/z [Da] | Rt [min] | Score | No. of Cmpds. | Site [%] | P | Sequence | Modification | Range | Type |
|---|---|---|---|---|---|---|---|---|---|---|---|---|---|
| 320 | 545.311 | 2 | 56.16 | 0.0306 | 18.63 | 54.16 | 3 | 0.0 | 0 | K.ASSLESGVPSR.F | | 51-61 | CID |
| 1172 | 600.046 | 3 | 126.79 | 0.0761 | 35.76 | 90.43 | 10 | 0.0 | 0 | K.SGTASVVCLLNNFYPR.E | Carbamidomethyl: 8 | 127-142 | CID |
| 348 | 1068.474 | 2 | -13.10 | -0.0140 | 19.14 | 155.06 | 14 | 0.0 | 0 | K.VDNALQSGNSQESVTEQDSK.D | | 150-169 | CID |
| 737 | 1207.537 | 3 | 244.94 | 0.2957 | 27.32 | 138.1 | 4 | 0.0 | 1 | K.VDNALQSGNSQESVTEQDSKDSTYSLSSTLTLSK.A | | 150-183 | CID |
| 779 | 751.961 | 2 | 103.93 | 0.0781 | 28.13 | 95.16 | 9 | 0.0 | 0 | K.DSTYSLSSTLTLSK.A | | 170-183 | CID |
| 482 | 626.223 | 3 | 387.40 | 0.2425 | 21.86 | 109.37 | 11 | 0.0 | 0 | K.VYACEVTHQGLSSPVTK.S | Carbamidomethyl: 4 | 191-207 | CID |
| 106 | 435.168 | 2 | -32.52 | -0.0142 | 14.09 | 54.21 | 8 | 0.0 | 1 | K.SFNRGEC.- | Carbamidomethyl: 7 | 208-214 | CID |

##### Protein 6: Immunoglobulin heavy constant gamma 4 OS=Homo sapiens GN=IGHG4 PE=1 SV=1

**Accession:** IGHG4_HUMAN
**Database:** SwissProt
**MW [kDa] / pI:** 35.9 / 7.2
**Modification(s):** Carbamidomethyl, Oxidation

**Score:** 639.8
**Seq. Coverage [%]:** 37.3
**No. of Peptides:** 9

| 10 | 20 | 30 | 40 | 50 | 60 | 70 | 80 | 90 | 100 |
|---|---|---|---|---|---|---|---|---|---|
| ASTKGPSVFP | LAPCSRSTSE | STAALGCLVK | DYFPEPVTVS | WNSGALTSGV | HTFPAVLQSS | GLYSLSSVVT | VPSSSLGTKT | YTCNVDHKPS | NTKVDKRVES |
| 110 | 120 | 130 | 140 | 150 | 160 | 170 | 180 | 190 | 200 |
| KYGPPCPSCP | APEFLGGPSV | FLFPPKPKDT | LMISRTPEVT | CVVVDVSQED | PEVQFNWYVD | GVEVHNAKTK | PREEQFNSTY | RVVSVLTVLH | QDWLNGKEYK |
| 210 | 220 | 230 | 240 | 250 | 260 | 270 | 280 | 290 | 300 |
| CKVSNKGLPS | SIEKTISKAK | GQPREPQVYT | LPPSQEEMTK | NQVSLTCLVK | GFYPSDIAVE | WESNGQPENN | YKTTPPVLDS | DGSFFLYSRL | TVDKSRWQEG |
| 310 | 320 | 330 | | | | | | | |
| NVFSCSVMHE | ALHNHYTQKS | LSLSLGK | | | | | | | |

MS/MS Peptide Matches

| Cmpd. | m/z meas. | z | Δ m/z [ppm] | Δ m/z [Da] | Rt [min] | Score | No. of Cmpds. | Site [%] | P | Sequence | Modification | Range | Type |
|---|---|---|---|---|---|---|---|---|---|---|---|---|---|
| 796 | 644.374 | 2 | 69.30 | 0.0447 | 28.51 | 69.42 | 13 | 0.0 | 0 | K.GPSVFPLAPCSR.S | Carbamidomethyl: 10 | 5-16 | CID |
| 677 | 712.393 | 2 | 48.43 | 0.0345 | 25.96 | 91.56 | 10 | 0.0 | 0 | R.STSESTAALGCLVK.D | Carbamidomethyl: 11 | 17-30 | CID |
| 98 | 417.041 | 4 | 223.33 | 0.0931 | 13.94 | 28.5 | 4 | 0.0 | 0 | K.TYTCNVDHKPSNTK.V | Carbamidomethyl: 4 | 80-93 | CID |
| 1149 | 982.176 | 3 | 358.60 | 0.3521 | 35.33 | 107.1 | 7 | 0.0 | 0 | K.YGPPCPSCPAPEFLGGPSVFLFPPKPK.D | Carbamidomethyl: 5, 8 | 102-128 | CID |
| 534 | 418.267 | 2 | 110.60 | 0.0463 | 22.88 | 50.66 | 13 | 0.0 | 0 | K.DTLMISR.T | | 129-135 | CID |
| 389 | 426.263 | 2 | 105.10 | 0.0448 | 20.00 | 53.61 | 3 | 0.0 | 0 | K.DTLMISR.T | Oxidation: 4 | 129-135 | CID |
| 1118 | 603.398 | 3 | 95.55 | 0.0577 | 34.67 | 63.61 | 12 | 0.0 | 0 | R.VVSVLTVLHQDWLNGK.E | | 182-197 | CID |
| 721 | 581.365 | 2 | 80.08 | 0.0466 | 26.99 | 76.52 | 10 | 0.0 | 0 | K.NQVSLTCLVK.G | Carbamidomethyl: 7 | 241-250 | CID |
| 947 | 848.794 | 3 | 92.72 | 0.0787 | 31.31 | 98.77 | 26 | 0.0 | 0 | K.GFYPSDIAVEWESNGQPENNYK.T | | 251-272 | CID |

#### Protein 7: Immunoglobulin lambda constant 2 OS=Homo sapiens GN=IGLC2 PE=1 SV=1

**Accession:** IGLC2_HUMAN
**Database:** SwissProt
**MW [kDa] / pI:** 11.3 / 6.9
**Modification(s):** Carbamidomethyl

**Score:** 540.2
**Seq. Coverage [%]:** 93.4
**No. of Peptides:** 8

10 20 30 40 50 60 70 80 90 100

GQPKAAPSVT LFPPSSEELQ ANKATLVCLI SDFYPGAVTV AWKADSSPVK AGVETTTPSK QSNNKYAASS YLSLTPEQWK SHRSYSCQVT HEGSTVEKTV

110

APTECS

**MS/MS Peptide Matches**

| Cmpd. | m/z meas. | z | Δ m/z [ppm] | Δ m/z [Da] | Rt [min] | Score | No. of Cmpds. | Site [%] | P | Sequence | Modification | Range | Type |
|---|---|---|---|---|---|---|---|---|---|---|---|---|---|
| 872 | 993.513 | 2 | 0.44 | 0.0004 | 29.95 | 72.24 | 8 | 0.0 | 0 | K.AAPSVTLFPPSSEELQANK.A | | 5-23 | CID |
| 1422 | 1106.103 | 2 | 21.19 | 0.0234 | 40.55 | 83.56 | 2 | 0.0 | 0 | K.ATLVCLISDFYPGAVTVAWK.A | Carbamidomethyl: 5 | 24-43 | CID |
| 285 | 558.994 | 3 | 67.37 | 0.0377 | 17.94 | 40.49 | 1 | 0.0 | 1 | K.ADSSPVKAGVETTTPSK.Q | | 44-60 | CID |
| 139 | 495.745 | 2 | -27.73 | -0.0137 | 14.84 | 75.28 | 18 | 0.0 | 0 | K.AGVETTTPSK.Q | | 51-60 | CID |
| 80 | 521.321 | 3 | 106.67 | 0.0556 | 13.47 | 53.1 | 3 | 0.0 | 1 | K.AGVETTTPSKQSNNK.Y | | 51-65 | CID |
| 921 | 872.474 | 2 | 46.94 | 0.0409 | 30.85 | 97.03 | 8 | 0.0 | 0 | K.YAASSYLSLTPEQWK.S | | 66-80 | CID |
| 277 | 571.312 | 3 | 94.68 | 0.0541 | 17.80 | 60.33 | 17 | 0.0 | 0 | R.SYSCQVTHEGSTVEK.T | Carbamidomethyl: 4 | 84-98 | CID |
| 457 | 853.051 | 3 | 7.95 | 0.0068 | 21.29 | 58.12 | 3 | 0.0 | 1 | R.SYSCQVTHEGSTVEKTVAPTECS.- | Carbamidomethyl: 4, 22 | 84-106 | CID |

#### Protein 8: Immunoglobulin lambda-like polypeptide 5 OS=Homo sapiens GN=IGLL5 PE=2 SV=2

| | | | |
|---|---|---|---|
| **Accession:** | IGLL5_HUMAN | **Score:** | 414.6 |
| **Database:** | SwissProt | **Seq. Coverage [%]:** | 43.9 |
| **MW [kDa] / pI:** | 23.0 / 9.1 | **No. of Peptides:** | 6 |
| **Modification(s):** | Carbamidomethyl | | |

```
        10         20         30         40         50         60         70         80         90        100
MRPKTGQVGC ETPEELGPGP RQRWPLLLLG LAMVAHGLLR PMVAPQSGDP DPGASVGSSR SSLRSLWGRL LLQPSPQRAD PRCWPRGFWS EPQSLCYVFG
       110        120        130        140        150        160        170        180        190        200
TGTKVTVLGQ PKANPTVTLF PPSSEELQAN KATLVCLISD FYPGAVTVAW KADGSPVKAG VETTKPSKQS NNKYAASSYL SLTPEQWKSH RSYSCQVTHE
       210        220
GSTVEKTVAP TECS
```

MS/MS Peptide Matches

| Cmpd. | m/z meas. | z | Δ m/z [ppm] | Δ m/z [Da] | Rt [min] | Score | No. of Cmpds. | Site [%] | P | Sequence | Modification | Range | Type |
|---|---|---|---|---|---|---|---|---|---|---|---|---|---|
| 385 | 421.297 | 2 | 86.10 | 0.0363 | 19.94 | 60.32 | 7 | 0.0 | 0 | K.VTVLGQPK.A | | 105-112 | CID |
| 862 | 681.71 | 3 | 37.23 | 0.0254 | 29.72 | 40.07 | 4 | 0.0 | 0 | K.ANPTVTLFPPSSEELQANK.A | | 113-131 | CID |
| 1422 | 1106.103 | 2 | 21.19 | 0.0234 | 40.55 | 83.56 | 2 | 0.0 | 0 | K.ATLVCLISDFYPGAVTVAWK.A | Carbamidomethyl: 5 | 132-151 | CID |
| 921 | 872.474 | 2 | 46.94 | 0.0409 | 30.85 | 97.03 | 8 | 0.0 | 0 | K.YAASSYLSLTPEQWK.S | | 174-188 | CID |
| 277 | 571.312 | 3 | 94.68 | 0.0541 | 17.80 | 60.33 | 17 | 0.0 | 0 | R.SYSCQVTHEGSTVEK.T | Carbamidomethyl: 4 | 192-206 | CID |
| 457 | 853.051 | 3 | 7.95 | 0.0068 | 21.29 | 58.12 | 3 | 0.0 | 1 | R.SYSCQVTHEGSTVEKTVAPTECS.- | Carbamidomethyl: 4, 22 | 192-214 | CID |

#### Protein 9: Immunoglobulin kappa variable 2D-28 OS=Homo sapiens GN=IGKV2D-28 PE=1 SV=2

**Accession:** KVD28_HUMAN
**Database:** SwissProt
**MW [kDa] / pI:** 12.9 / 5.6

**Score:** 171.1
**Seq. Coverage [%]:** 16.7
**No. of Peptides:** 2

```
        10         20         30         40         50         60         70         80         90        100
MRLPAQLLGL LMLWVSGSSG DIVMTQSPLS LPVTPGEPAS ISCRSSQSLL HSNGYNYLDW YLQKPGQSPQ LLIYLGSNRA SGVPDRFSGS GSGTDFTLKI
       110        120        130
SRVEAEDVGV YYCMQALQTP
```

MS/MS Peptide Matches

| Cmpd. | m/z meas. | z | Δ m/z [ppm] | Δ m/z [Da] | Rt [min] | Score | No. of Cmpds. | Site [%] | P | Sequence | Modification | Range | Type |
|---|---|---|---|---|---|---|---|---|---|---|---|---|---|
| 640 | 662.668 | 3 | 16.67 | 0.0110 | 25.12 | 68.6 | 3 | 0.0 | 1 | R.ASGVPDRFSGSGSGTDFTLK.I | | 80-99 | CID |
| 593 | 652.284 | 2 | -42.73 | -0.0279 | 24.16 | 102.51 | 3 | 0.0 | 0 | R.FSGSGSGTDFTLK.I | | 87-99 | CID |

#### Protein 10: Immunoglobulin heavy variable 3-30-3 OS=Homo sapiens GN=IGHV3-30-3 PE=3 SV=1

**Accession:** HVC33_HUMAN
**Database:** SwissProt
**MW [kDa] / pI:** 13.0 / 9.1
**Modification(s):** Carbamidomethyl

**Score:** 97.2
**Seq. Coverage [%]:** 15.4
**No. of Peptides:** 2

```
        10         20         30         40         50         60         70         80         90        100
MEFGLSWVFL VALLRGVQCQ VQLVESGGGV VQPGRSLRLS CAASGFTFSS YAMHWVRQAP GKGLEWVAVI SYDGSNKYYA DSVKGRFTIS RDNSKNTLYL
       110        120
QMNSLRAEDT AVYYCAR
```

MS/MS Peptide Matches

| Cmpd. | m/z meas. | z | Δ m/z [ppm] | Δ m/z [Da] | Rt [min] | Score | No. of Cmpds. | Site [%] | P | Sequence | Modification | Range | Type |
|---|---|---|---|---|---|---|---|---|---|---|---|---|---|
| 223 | 423.24 | 2 | 81.24 | 0.0344 | 16.63 | 30.23 | 4 | 0.0 | 0 | K.YYADSVK.G | | 78-84 | CID |
| 435 | 659.832 | 2 | 63.28 | 0.0417 | 20.87 | 67.0 | 3 | 0.0 | 0 | R.AEDTAVYYCAR.- | Carbamidomethyl: 9 | 107-117 | CID |

#### Protein 11: Alpha-1-antitrypsin OS=Homo sapiens GN=SERPINA1 PE=1 SV=3

**Accession:** A1AT_HUMAN
**Database:** SwissProt
**MW [kDa] / pI:** 46.7 / 5.4

**Score:** 66.2
**Seq. Coverage [%]:** 4.1
**No. of Peptides:** 1

| 10 | 20 | 30 | 40 | 50 | 60 | 70 | 80 | 90 | 100 |
|---|---|---|---|---|---|---|---|---|---|
| MPSSVSWGIL | LLAGLCCLVP | VSLAEDPQGD | AAQKTDTSHH | DQDHPTFNKI | TPNLAEFAFS | LYRQLAHQSN | STNIFFSPVS | IATAFAMLSL | GTKADTHDEI |
| 110 | 120 | 130 | 140 | 150 | 160 | 170 | 180 | 190 | 200 |
| LEGLNFNLTE | IPEAQIHEGF | QELLRTLNQP | DSQLQLTTGN | GLFLSEGLKL | VDKFLEDVKK | LYHSEAFTVN | FGDTEEAKKQ | INDYVEKGTQ | GKIVDLVKEL |
| 210 | 220 | 230 | 240 | 250 | 260 | 270 | 280 | 290 | 300 |
| DRDTVFALVN | YIFFKGKWER | PFEVKDTEEE | DFHVDQVTTV | KVPMMKRLGM | FNIQHCKKLS | SWVLLMKYLG | NATAIFFLPD | EGKLQHLENE | LTHDIITKFL |
| 310 | 320 | 330 | 340 | 350 | 360 | 370 | 380 | 390 | 400 |
| ENEDRRSASL | HLPKLSITGT | YDLKSVLGQL | GITKVFSNGA | DLSGVTEEAP | LKLSKAVHKA | VLTIDEKGTE | AAGAMFLEAI | PMSIPPEVKF | NKPFVFLMIE |
| 410 | 420 | | | | | | | | |
| QNTKSPLFMG | KVVNPTQK | | | | | | | | |

MS/MS Peptide Matches

| Cmpd. | m/z meas. | z | Δ m/z [ppm] | Δ m/z [Da] | Rt [min] | Score | No. of Cmpds. | Site [%] | P | Sequence | Modification | Range | Type |
|---|---|---|---|---|---|---|---|---|---|---|---|---|---|
| 1484 | 697.528 | 3 | 226.44 | 0.1579 | 41.92 | 66.23 | 1 | 0.0 | 1 | K.ELDRDTVFALVNYIFFK.G | | 199-215 | CID |

#### Protein 12: Immunoglobulin kappa variable 3-15 OS=Homo sapiens GN=IGKV3-15 PE=1 SV=2

**Accession:** KV315_HUMAN
**Database:** SwissProt
**MW [kDa] / pI:** 12.5 / 5.1

**Score:** 65.0
**Seq. Coverage [%]:** 7.8
**No. of Peptides:** 1

| 10 | 20 | 30 | 40 | 50 | 60 | 70 | 80 | 90 | 100 |
|---|---|---|---|---|---|---|---|---|---|
| MEAPAQLLFL | LLLWLPDTTG | EIVMTQSPAT | LSVSPGERAT | LSCRASQSVS | SNLAWYQQKP | GQAPRLLIYG | ASTRATGIPA | RFSGSGSGTE | FTLTISSLQS |
| 110 | 120 | | | | | | | | |
| EDFAVYYCQQ | YNNWP | | | | | | | | |

MS/MS Peptide Matches

| Cmpd. | m/z meas. | z | Δ m/z [ppm] | Δ m/z [Da] | Rt [min] | Score | No. of Cmpds. | Site [%] | P | Sequence | Modification | Range | Type |
|---|---|---|---|---|---|---|---|---|---|---|---|---|---|
| 618 | 497.327 | 2 | 74.37 | 0.0370 | 24.65 | 64.98 | 5 | 0.0 | 0 | R.LLIYGASTR.A | | 66-74 | CID |

#### Protein 13: Immunoglobulin kappa variable 3-20 OS=Homo sapiens GN=IGKV3-20 PE=1 SV=2

| | | | |
|---|---|---|---|
| **Accession:** | KV320_HUMAN | **Score:** | 57.7 |
| **Database:** | SwissProt | **Seq. Coverage [%]:** | 7.8 |
| **MW [kDa] / pI:** | 12.5 / 4.9 | **No. of Peptides:** | 1 |

```
        10         20         30         40         50         60         70         80         90        100
METPAQLLFL LLLWLPDTTG EIVLTQSPGT LSLSPGERAT LSCRASQSVS SSYLAWYQQK PGQAPRLLIY GASSRATGIP DRFSGSGSGT DFTLTISRLE
       110        120
PEDFAVYYCQ QYGSSP
```

MS/MS Peptide Matches

| Cmpd. | m/z meas. | z | Δ m/z [ppm] | Δ m/z [Da] | Rt [min] | Score | No. of Cmpds. | Site [%] | P | Sequence | Modification | Range | Type |
|---|---|---|---|---|---|---|---|---|---|---|---|---|---|
| 606 | 490.317 | 2 | 71.00 | 0.0348 | 24.38 | 57.71 | 4 | 0.0 | 0 | R.LLIYGASSR.A | | 67-75 | CID |

#### Protein 14: Immunoglobulin heavy variable 3-73 OS=Homo sapiens GN=IGHV3-73 PE=3 SV=1

| | | | |
|---|---|---|---|
| **Accession:** | HV373_HUMAN | **Score:** | 50.9 |
| **Database:** | SwissProt | **Seq. Coverage [%]:** | 5.9 |
| **MW [kDa] / pI:** | 12.8 / 9.3 | **No. of Peptides:** | 1 |

```
        10         20         30         40         50         60         70         80         90        100
MEFGLSWVFL VAILKGVQCE VQLVESGGGL VQPGGSLKLS CAASGFTFSG SAMHWVRQAS GKGLEWVGRI RSKANSYATA YAASVKGRFT ISRDDSKNTA
       110        120
YLQMNSLKTE DTAVYYCTR
```

MS/MS Peptide Matches

| Cmpd. | m/z meas. | z | Δ m/z [ppm] | Δ m/z [Da] | Rt [min] | Score | No. of Cmpds. | Site [%] | P | Sequence | Modification | Range | Type |
|---|---|---|---|---|---|---|---|---|---|---|---|---|---|
| 628 | 408.762 | 2 | 98.43 | 0.0402 | 24.85 | 50.89 | 3 | 0.0 | 0 | K.GLEWVGR.I | | 63-69 | CID |

#### Project Info

| | | | |
|---|---|---|---|
| **Name:** | Alagesan | **Date:** | Mar 5, 2019 |
| **Note:** | Nanobooster | | |

#### Sample Info & Protocols

| | |
|---|---|
| **Name:** | Nanobooster - IgG |
| **Date:** | Mar 5, 2019 |

#### Search Result Info

| | | | |
|---|---|---|---|
| **Search Result:** | IonTrap_Human_Swiss_CID_2019-03-05 17:55:08 | | |
| **Location:** | /Alagesan/Nanobooster - IgG/Ka_IgG Trypsin NoEnrich NB ACN_BA5_01_8745.mgf | | |
| **Search Method:** | IonTrap_Human_Swiss_CID | **Version:** | 1.2 |
| **Search Engine(s):** | Mascot, 2.6.0 | **Ident. Compound(s):** | 0/2433 |
| **Database(s):** | SwissProt, SwissProt_2017_04.fasta | | |
| **Note:** | | | |

#### Results

##### Protein 1: Immunoglobulin heavy constant gamma 1 OS=Homo sapiens GN=IGHG1 PE=1 SV=1

| | | | |
|---|---|---|---|
| **Accession:** | IGHG1_HUMAN | **Score:** | 961.2 |
| **Database:** | SwissProt | **Seq. Coverage [%]:** | 54.8 |
| **MW [kDa] / pI:** | 36.1 / 8.5 | **No. of Peptides:** | 14 |
| **Modification(s):** | Carbamidomethyl, Oxidation | | |

```
        10         20         30         40         50         60         70         80         90        100
ASTKGPSVFP LAPSSKSTSG GTAALGCLVK DYFPEPVTVS WNSGALTSGV HTFPAVLQSS GLYSLSSVVT VPSSSLGTQT YICNVNHKPS NTKVDKKVEP
       110        120        130        140        150        160        170        180        190        200
KSCDKTHTCP PCPAPELLGG PSVFLFPPKP KDTLMISRTP EVTCVVVDVS HEDPEVKFNW YVDGVEVHNA KTKPREEQYN STYRVVSVLT VLHQDWLNGK
       210        220        230        240        250        260        270        280        290        300
EYKCKVSNKA LPAPIEKTIS KAKGQPREPQ VYTLPPSRDE LTKNQVSLTC LVKGFYPSDI AVEWESNGQP ENNYKTTPPV LDSDGSFFLY SKLTVDKSRW
       310        320        330        340
QQGNVFSCSV MHEALHNHYT QKSLSLSPGK
```

**MS/MS Peptide Matches**

| Cmpd. | m/z meas. | z | Δ m/z [ppm] | Δ m/z [Da] | Rt [min] | Score | No. of Cmpds. | Site [%] | P | Sequence | Modification | Range | Type |
|---|---|---|---|---|---|---|---|---|---|---|---|---|---|
| 924 | 593.847 | 2 | 33.74 | 0.0200 | 27.41 | 54.45 | 15 | 0.0 | 0 | K.GPSVFPLAPSSK.S | | 5-16 | CID |
| 818 | 661.364 | 2 | 32.28 | 0.0213 | 25.08 | 132.45 | 14 | 0.0 | 0 | K.STSGGTAALGCLVK.D | Carbamidomethyl: 11 | 17-30 | CID |
| 1231 | 711.946 | 4 | 106.98 | 0.0762 | 33.18 | 54.83 | 5 | 0.0 | 0 | K.THTCPPCPAPELLGGPSVFLFPPKPK.D | Carbamidomethyl: 4, 7 | 106-131 | CID |
| 694 | 418.268 | 2 | 112.99 | 0.0473 | 22.50 | 50.57 | 11 | 0.0 | 0 | K.DTLMISR.T | | 132-138 | CID |
| 555 | 426.269 | 2 | 119.18 | 0.0508 | 19.74 | 47.24 | 3 | 0.0 | 0 | K.DTLMISR.T | Oxidation: 4 | 132-138 | CID |
| 912 | 713.703 | 3 | 31.30 | 0.0223 | 27.17 | 93.98 | 3 | 0.0 | 0 | R.TPEVTCVVVDVSHEDPEVK.F | Carbamidomethyl: 6 | 139-157 | CID |
| 915 | 560.028 | 3 | 159.22 | 0.0892 | 27.24 | 76.88 | 15 | 0.0 | 0 | K.FNWYVDGVEVHNAK.T | | 158-171 | CID |
| 1285 | 603.389 | 3 | 80.64 | 0.0487 | 34.28 | 67.94 | 13 | 0.0 | 0 | R.VVSVLTVLHQDWLNGK.E | | 185-200 | CID |

| Cmpd. | m/z meas. | z | Δ m/z [ppm] | Δ m/z [Da] | Rt [min] | Score | No. of Cmpds. | Site [%] | P | Sequence | Modification | Range | Type |
|---|---|---|---|---|---|---|---|---|---|---|---|---|---|
| 646 | 419.795 | 2 | 94.63 | 0.0397 | 21.52 | 25.34 | 5 | 0.0 | 0 | K.ALPAPIEK.T | | 210-217 | CID |
| 727 | 643.82 | 2 | -32.00 | -0.0206 | 23.24 | 61.44 | 13 | 0.0 | 0 | R.EPQVYTLPPSR.D | | 228-238 | CID |
| 773 | 625.062 | 3 | 107.35 | 0.0671 | 24.17 | 28.9 | 13 | 0.0 | 1 | R.EPQVYTLPPSRDELTK.N | | 228-243 | CID |
| 883 | 581.347 | 2 | 49.12 | 0.0286 | 26.58 | 69.4 | 9 | 0.0 | 0 | K.NQVSLTCLVK.G | Carbamidomethyl: 7 | 244-253 | CID |
| 1108 | 848.796 | 3 | 95.07 | 0.0807 | 30.82 | 91.26 | 32 | 0.0 | 0 | K.GFYPSDIAVEWESNGQPENNYK.T | | 254-275 | CID |
| 1200 | 937.515 | 2 | 53.82 | 0.0505 | 32.52 | 106.54 | 9 | 0.0 | 0 | K.TTPPVLDSDGSFFLYSK.L | | 276-292 | CID |

#### Protein 2: Immunoglobulin heavy constant gamma 3 OS=Homo sapiens GN=IGHG3 PE=1 SV=2

**Accession:** IGHG3_HUMAN
**Database:** SwissProt
**MW [kDa] / pI:** 41.3 / 8.2
**Modification(s):** Carbamidomethyl, Oxidation

**Score:** 839.0
**Seq. Coverage [%]:** 36.3
**No. of Peptides:** 13

| 10 | 20 | 30 | 40 | 50 | 60 | 70 | 80 | 90 | 100 |
|---|---|---|---|---|---|---|---|---|---|
| ASTKGPSVFP | LAPCSRSTSG | GTAALGCLVK | DYFPEPVTVS | WNSGALTSGV | HTFPAVLQSS | GLYSLSSVVT | VPSSSLGTQT | YTCNVNHKPS | NTKVDKRVEL |
| **110** | **120** | **130** | **140** | **150** | **160** | **170** | **180** | **190** | **200** |
| KTPLGDTTHT | CPRCPEPKSC | DTPPPCPRCP | EPKSCDTPPP | CPRCPEPKSC | DTPPPCPRCP | APELLGGPSV | FLFPPKPKDT | LMISRTPEVT | CVVVDVSHED |
| **210** | **220** | **230** | **240** | **250** | **260** | **270** | **280** | **290** | **300** |
| PEVQFKWYVD | GVEVHNAKTK | PREEQYNSTF | RVVSVLTVLH | QDWLNGKEYK | CKVSNKALPA | PIEKTISKTK | GQPREPQVYT | LPPSREEMTK | NQVSLTCLVK |
| **310** | **320** | **330** | **340** | **350** | **360** | **370** | **380** | | |
| GFYPSDIAVE | WESSGQPENN | YNTTPPMLDS | DGSFFLYSKL | TVDKSRWQQG | NIFSCSVMHE | ALHNRFTQKS | LSLSPGK | | |

MS/MS Peptide Matches

| Cmpd. | m/z meas. | z | Δ m/z [ppm] | Δ m/z [Da] | Rt [min] | Score | No. of Cmpds. | Site [%] | P | Sequence | Modification | Range | Type |
|---|---|---|---|---|---|---|---|---|---|---|---|---|---|
| 959 | 644.385 | 2 | 86.37 | 0.0557 | 28.09 | 67.53 | 13 | 0.0 | 0 | K.GPSVFPLAPCSR.S | Carbamidomethyl: 10 | 5-16 | CID |

| Cmpd. | m/z meas. | z | Δ m/z [ppm] | Δ m/z [Da] | Rt [min] | Score | No. of Cmpds. | Site [%] | P | Sequence | Modification | Range | Type |
|---|---|---|---|---|---|---|---|---|---|---|---|---|---|
| 818 | 661.364 | 2 | 32.28 | 0.0213 | 25.08 | 132.45 | 14 | 0.0 | 0 | R.STSGGTAALGCLVK.D | Carbamidomethyl: 11 | 17-30 | CID |
| 417 | 452.561 | 3 | 23.00 | 0.0104 | 16.98 | 63.47 | 6 | 0.0 | 0 | K.TPLGDTTHTCPR.C | Carbamidomethyl: 10 | 102-113 | CID |
| 364 | 593.772 | 2 | 32.65 | 0.0194 | 15.80 | 65.29 | 9 | 0.0 | 0 | K.SCDTPPPCPR.C | Carbamidomethyl: 2, 8 | 119-128 | CID |
| 1277 | 717.801 | 3 | 102.80 | 0.0738 | 34.13 | 91.28 | 3 | 0.0 | 0 | R.CPAPELLGGPSVFLFPPKPK.D | Carbamidomethyl: 1 | 159-178 | CID |
| 694 | 418.268 | 2 | 112.99 | 0.0473 | 22.50 | 50.57 | 11 | 0.0 | 0 | K.DTLMISR.T | | 179-185 | CID |
| 555 | 426.269 | 2 | 119.18 | 0.0508 | 19.74 | 47.24 | 3 | 0.0 | 0 | K.DTLMISR.T | Oxidation: 4 | 179-185 | CID |
| 745 | 472.965 | 3 | 133.79 | 0.0633 | 23.60 | 42.89 | 4 | 0.0 | 0 | K.WYVDGVEVHNAK.T | | 207-218 | CID |
| 1285 | 603.389 | 3 | 80.64 | 0.0487 | 34.28 | 67.94 | 13 | 0.0 | 0 | R.VVSVLTVLHQDWLNGK.E | | 232-247 | CID |
| 646 | 419.795 | 2 | 94.63 | 0.0397 | 21.52 | 25.34 | 5 | 0.0 | 0 | K.ALPAPIEK.T | | 257-264 | CID |
| 727 | 643.82 | 2 | -32.00 | -0.0206 | 23.24 | 61.44 | 13 | 0.0 | 0 | R.EPQVYTLPPSR.E | | 275-285 | CID |
| 747 | 635.73 | 3 | 122.29 | 0.0777 | 23.65 | 54.18 | 30 | 0.0 | 1 | R.EPQVYTLPPSREEMTK.N | | 275-290 | CID |
| 883 | 581.347 | 2 | 49.12 | 0.0286 | 26.58 | 69.4 | 9 | 0.0 | 0 | K.NQVSLTCLVK.G | Carbamidomethyl: 7 | 291-300 | CID |

##### Protein 3: Immunoglobulin kappa constant OS=Homo sapiens GN=IGKC PE=1 SV=2

**Accession:** IGKC_HUMAN
**Database:** SwissProt
**MW [kDa] / pI:** 11.8 / 6.1
**Modification(s):** Carbamidomethyl

**Score:** 811.9
**Seq. Coverage [%]:** 86.9
**No. of Peptides:** 8

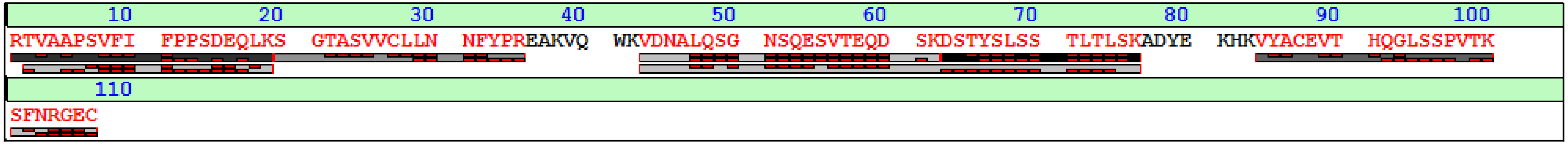


MS/MS Peptide Matches

| Cmpd. | m/z meas. | z | Δ m/z [ppm] | Δ m/z [Da] | Rt [min] | Score | No. of Cmpds. | Site [%] | P | Sequence | Modification | Range | Type |
|---|---|---|---|---|---|---|---|---|---|---|---|---|---|
| 1113 | 701.418 | 3 | 52.93 | 0.0371 | 30.94 | 31.22 | 9 | 0.0 | 1 | -.RTVAAPSVFIFPPSDEQLK.S | | 1-19 | CID |
| 1260 | 973.555 | 2 | 38.91 | 0.0379 | 33.86 | 91.9 | 20 | 0.0 | 0 | R.TVAAPSVFIFPPSDEQLK.S | | 2-19 | CID |
| 1350 | 600.018 | 3 | 80.12 | 0.0481 | 35.57 | 81.11 | 12 | 0.0 | 0 | K.SGTASVVCLLNNFYPR.E | Carbamidomethyl: 8 | 20-35 | CID |
| 546 | 1068.465 | 2 | -21.53 | -0.0230 | 19.57 | 169.91 | 16 | 0.0 | 0 | K.VDNALQSGNSQESVTEQDSK.D | | 43-62 | CID |
| 901 | 1207.52 | 3 | 230.86 | 0.2787 | 26.94 | 176.3 | 3 | 0.0 | 1 | K.VDNALQSGNSQESVTEQDSKDSTYSLSSTLTLSK.A | | 43-76 | CID |
| 932 | 751.919 | 2 | 48.07 | 0.0361 | 27.57 | 106.75 | 10 | 0.0 | 0 | K.DSTYSLSSTLTLSK.A | | 63-76 | CID |
| 650 | 626.063 | 3 | 131.81 | 0.0825 | 21.62 | 108.92 | 12 | 0.0 | 0 | K.VYACEVTHQGLSSPVTK.S | Carbamidomethyl: 4 | 84-100 | CID |
| 285 | 435.144 | 2 | -87.67 | -0.0382 | 14.05 | 45.75 | 8 | 0.0 | 1 | K.SFNRGEC.- | Carbamidomethyl: 7 | 101-107 | CID |

#### Protein 4: Immunoglobulin heavy constant gamma 2 OS=Homo sapiens GN=IGHG2 PE=1 SV=2

**Accession:** IGHG2_HUMAN
**Database:** SwissProt
**MW [kDa] / pI:** 35.9 / 7.7
**Modification(s):** Carbamidomethyl, Oxidation

**Score:** 755.5
**Seq. Coverage [%]:** 37.4
**No. of Peptides:** 13

| 10 | 20 | 30 | 40 | 50 | 60 | 70 | 80 | 90 | 100 |
|---|---|---|---|---|---|---|---|---|---|
| ASTKGPSVFP | LAPCSRSTSE | STAALGCLVK | DYFPEPVTVS | WNSGALTSGV | HTFPAVLQSS | GLYSLSSVVT | VPSSNFGTQT | YTCNVDHKPS | NTKVDKTVER |
| **110** | **120** | **130** | **140** | **150** | **160** | **170** | **180** | **190** | **200** |
| KCCVECPPCP | APPVAGPSVF | LFPPKPKDTL | MISRTPEVTC | VVVDVSHEDP | EVQFNWYVDG | VEVHNAKTKP | REEQFNSTFR | VVSVLTVVHQ | DWLNGKEYKC |
| **210** | **220** | **230** | **240** | **250** | **260** | **270** | **280** | **290** | **300** |
| KVSNKGLPAP | IEKTISKTKG | QPREPQVYTL | PPSREEMTKN | QVSLTCLVKG | FYPSDISVEW | ESNGQPENNY | KTTPPMLDSD | GSFFLYSKLT | VDKSRWQQGN |
| **310** | **320** | **330** | | | | | | | |
| VFSCSVMHEA | LHNHYTQKSL | SLSPGK | | | | | | | |

MS/MS Peptide Matches

| Cmpd. | m/z meas. | z | Δ m/z [ppm] | Δ m/z [Da] | Rt [min] | Score | No. of Cmpds. | Site [%] | P | Sequence | Modification | Range | Type |
|---|---|---|---|---|---|---|---|---|---|---|---|---|---|
| 959 | 644.385 | 2 | 86.37 | 0.0557 | 28.09 | 67.53 | 13 | 0.0 | 0 | K.GPSVFPLAPCSR.S | Carbamidomethyl: 10 | 5-16 | CID |
| 845 | 712.381 | 2 | 31.59 | 0.0225 | 25.70 | 83.88 | 10 | 0.0 | 0 | R.STSESTAALGCLVK.D | Carbamidomethyl: 11 | 17-30 | CID |
| 1093 | 760.155 | 4 | 362.40 | 0.2754 | 30.56 | 35.56 | 3 | 0.0 | 1 | R.KCCVECPPCPAPPVAGPSVFLFPPKPK.D | Carbamidomethyl: 2, 3, 6, 9 | 101-127 | CID |
| 1189 | 970.483 | 3 | 354.85 | 0.3443 | 32.31 | 76.82 | 5 | 0.0 | 0 | K.CCVECPPCPAPPVAGPSVFLFPPKPK.D | Carbamidomethyl: 1, 2, 5, 8 | 102-127 | CID |
| 694 | 418.268 | 2 | 112.99 | 0.0473 | 22.50 | 50.57 | 11 | 0.0 | 0 | K.DTLMISR.T | | 128-134 | CID |
| 555 | 426.269 | 2 | 119.18 | 0.0508 | 19.74 | 47.24 | 3 | 0.0 | 0 | K.DTLMISR.T | Oxidation: 4 | 128-134 | CID |
| 1177 | 599.07 | 3 | 670.71 | 0.4015 | 32.09 | 39.68 | 6 | 0.0 | 0 | R.VVSVLTVVHQDWLNGK.E | | 181-196 | CID |
| 1098 | 554.398 | 4 | 170.68 | 0.0946 | 30.63 | 15.09 | 1 | 0.0 | 1 | R.VVSVLTVVHQDWLNGKEYK.C | | 181-199 | CID |
| 727 | 643.82 | 2 | -32.00 | -0.0206 | 23.24 | 61.44 | 13 | 0.0 | 0 | R.EPQVYTLPPSR.E | | 224-234 | CID |
| 747 | 635.73 | 3 | 122.29 | 0.0777 | 23.65 | 54.18 | 30 | 0.0 | 1 | R.EPQVYTLPPSREEMTK.N | | 224-239 | CID |
| 883 | 581.347 | 2 | 49.12 | 0.0286 | 26.58 | 69.4 | 9 | 0.0 | 0 | K.NQVSLTCLVK.G | Carbamidomethyl: 7 | 240-249 | CID |
| 1195 | 953.55 | 2 | 104.27 | 0.0994 | 32.44 | 96.07 | 6 | 0.0 | 0 | K.TTPPMLDSDGSFFLYSK.L | | 272-288 | CID |
| 1101 | 961.446 | 2 | -2.12 | -0.0020 | 30.68 | 58.07 | 2 | 0.0 | 0 | K.TTPPMLDSDGSFFLYSK.L | Oxidation: 5 | 272-288 | CID |

##### Protein 5: Immunoglobulin heavy constant gamma 4 OS=Homo sapiens GN=IGHG4 PE=1 SV=1

**Accession:** IGHG4_HUMAN
**Database:** SwissProt
**MW [kDa] / pI:** 35.9 / 7.2
**Modification(s):** Carbamidomethyl, Oxidation

**Score:** 650.5
**Seq. Coverage [%]:** 42.5
**No. of Peptides:** 10

| 10 | 20 | 30 | 40 | 50 | 60 | 70 | 80 | 90 | 100 |
|---|---|---|---|---|---|---|---|---|---|
| ASTKGPSVFP | LAPCSRSTSE | STAALGCLVK | DYFPEPVTVS | WNSGALTSGV | HTFPAVLQSS | GLYSLSSVVT | VPSSSLGTKT | YTCNVDHKPS | NTKVDKRVES |
| **110** | **120** | **130** | **140** | **150** | **160** | **170** | **180** | **190** | **200** |
| KYGPPCPSCP | APEFLGGPSV | FLFPPKPKDT | LMISRTPEVT | CVVVDVSQED | PEVQFNWYVD | GVEVHNAKTK | PREEQFNSTY | RVVSVLTVLH | QDWLNGKEYK |
| **210** | **220** | **230** | **240** | **250** | **260** | **270** | **280** | **290** | **300** |
| CKVSNKGLPS | SIEKTISKAK | GQPREPQVYT | LPPSQEEMTK | NQVSLTCLVK | GFYPSDIAVE | WESNGQPENN | YKTTPPVLDS | DGSFFLYSRL | TVDKSRWQEG |
| **310** | **320** | **330** | | | | | | | |
| NVFSCSVMHE | ALHNHYTQKS | LSLSLGK | | | | | | | |

**MS/MS Peptide Matches**

| Cmpd. | m/z meas. | z | Δ m/z [ppm] | Δ m/z [Da] | Rt [min] | Score | No. of Cmpds. | Site [%] | P | Sequence | Modification | Range | Type |
|---|---|---|---|---|---|---|---|---|---|---|---|---|---|
| 959 | 644.385 | 2 | 86.37 | 0.0557 | 28.09 | 67.53 | 13 | 0.0 | 0 | K.GPSVFPLAPCSR.S | Carbamidomethyl: 10 | 5-16 | CID |
| 845 | 712.381 | 2 | 31.59 | 0.0225 | 25.70 | 83.88 | 10 | 0.0 | 0 | R.STSESTAALGCLVK.D | Carbamidomethyl: 11 | 17-30 | CID |
| 273 | 416.992 | 4 | 105.81 | 0.0441 | 13.78 | 24.24 | 2 | 0.0 | 0 | K.TYTCNVDHKPSNTK.V | Carbamidomethyl: 4 | 80-93 | CID |
| 1322 | 982.205 | 3 | 388.14 | 0.3811 | 35.06 | 76.09 | 6 | 0.0 | 0 | K.YGPPCPSCPAPEFLGGPSVFLFPPKPK.D | Carbamidomethyl: 5, 8 | 102-128 | CID |
| 694 | 418.268 | 2 | 112.99 | 0.0473 | 22.50 | 50.57 | 11 | 0.0 | 0 | K.DTLMISR.T | | 129-135 | CID |
| 555 | 426.269 | 2 | 119.18 | 0.0508 | 19.74 | 47.24 | 3 | 0.0 | 0 | K.DTLMISR.T | Oxidation: 4 | 129-135 | CID |
| 1285 | 603.389 | 3 | 80.64 | 0.0487 | 34.28 | 67.94 | 13 | 0.0 | 0 | R.VVSVLTVLHQDWLNGK.E | | 182-197 | CID |
| 883 | 581.347 | 2 | 49.12 | 0.0286 | 26.58 | 69.4 | 9 | 0.0 | 0 | K.NQVSLTCLVK.G | Carbamidomethyl: 7 | 241-250 | CID |
| 1108 | 848.796 | 3 | 95.07 | 0.0807 | 30.82 | 91.26 | 32 | 0.0 | 0 | K.GFYPSDIAVEWESNGQPENNYK.T | | 251-272 | CID |
| 1194 | 951.519 | 2 | 54.00 | 0.0514 | 32.40 | 72.31 | 1 | 0.0 | 0 | K.TTPPVLDSDGSFFLYSR.L | | 273-289 | CID |

#### Protein 6: Immunoglobulin lambda constant 3 OS=Homo sapiens GN=IGLC3 PE=1 SV=1

| | | | |
|---|---|---|---|
| **Accession:** | IGLC3_HUMAN | **Score:** | 566.2 |
| **Database:** | SwissProt | **Seq. Coverage [%]:** | 93.4 |
| **MW [kDa] / pI:** | 11.3 / 6.9 | **No. of Peptides:** | 8 |
| **Modification(s):** | Carbamidomethyl | | |

```
        10         20         30         40         50         60         70         80         90        100
GQPKAAPSVT LFPPSSEELQ ANKATLVCLI SDFYPGAVTV AWKADSSPVK AGVETTTPSK QSNNKYAASS YLSLTPEQWK SHKSYSCQVT HEGSTVEKTV

       110
APTECS
```

**MS/MS Peptide Matches**

| Cmpd. | m/z meas. | z | Δ m/z [ppm] | Δ m/z [Da] | Rt [min] | Score | No. of Cmpds. | Site [%] | P | Sequence | Modification | Range | Type |
|---|---|---|---|---|---|---|---|---|---|---|---|---|---|
| 1028 | 993.493 | 2 | -19.69 | -0.0196 | 29.38 | 76.51 | 9 | 0.0 | 0 | K.AAPSVTLFPPSSEELQANK.A | | 5-23 | CID |
| 1601 | 738.099 | 3 | 510.86 | 0.3769 | 40.24 | 101.69 | 3 | 0.0 | 0 | K.ATLVCLISDFYPGAVTVAWK.A | Carbamidomethyl: 5 | 24-43 | CID |
| 453 | 558.977 | 3 | 36.96 | 0.0207 | 17.71 | 51.64 | 3 | 0.0 | 1 | K.ADSSPVKAGVETTTPSK.Q | | 44-60 | CID |
| 306 | 495.809 | 2 | 101.37 | 0.0503 | 14.51 | 78.86 | 19 | 0.0 | 0 | K.AGVETTTPSK.Q | | 51-60 | CID |
| 258 | 521.327 | 3 | 118.18 | 0.0616 | 13.42 | 58.19 | 3 | 0.0 | 1 | K.AGVETTTPSKQSNNK.Y | | 51-65 | CID |
| 1071 | 872.447 | 2 | 15.99 | 0.0139 | 30.15 | 81.13 | 8 | 0.0 | 0 | K.YAASSYLSLTPEQWK.S | | 66-80 | CID |
| 408 | 571.281 | 3 | 40.41 | 0.0231 | 16.73 | 76.5 | 14 | 0.0 | 0 | K.SYSCQVTHEGSTVEK.T | Carbamidomethyl: 4 | 84-98 | CID |
| 624 | 853.027 | 3 | -20.19 | -0.0172 | 21.06 | 41.65 | 3 | 0.0 | 1 | K.SYSCQVTHEGSTVEKTVAPTECS.- | Carbamidomethyl: 4, 22 | 84-106 | CID |

#### Protein 7: Immunoglobulin lambda-like polypeptide 5 OS=Homo sapiens GN=IGLL5 PE=2 SV=2

| | | | |
|---|---|---|---|
| **Accession:** | IGLL5_HUMAN | **Score:** | 391.1 |
| **Database:** | SwissProt | **Seq. Coverage [%]:** | 39.7 |
| **MW [kDa] / pI:** | 23.0 / 9.1 | **No. of Peptides:** | 6 |
| **Modification(s):** | Carbamidomethyl | | |

| 10 | 20 | 30 | 40 | 50 | 60 | 70 | 80 | 90 | 100 |
|---|---|---|---|---|---|---|---|---|---|
| MRPKTGQVGC | ETPEELGPGP | RQRWPLLLLG | LAMVAHGLLR | PMVAPQSGDP | DPGASVGSSR | SSLRSLWGRL | LLQPSPQRAD | PRCWPRGFWS | EPQSLCYVFG |
| **110** | **120** | **130** | **140** | **150** | **160** | **170** | **180** | **190** | **200** |
| TGTKVTVLGQ | PKANPTVTLF | PPSSEELQAN | KATLVCLISD | FYPGAVTVAW | KADGSPVKAG | VETTKPSKQS | NNKYAASSYL | SLTPEQWKSH | RSYSCQVTHE |
| **210** | **220** | | | | | | | | |
| GSTVEKTVAP | TECS | | | | | | | | |

MS/MS Peptide Matches

| Cmpd. | m/z meas. | z | Δ m/z [ppm] | Δ m/z [Da] | Rt [min] | Score | No. of Cmpds. | Site [%] | P | Sequence | Modification | Range | Type |
|---|---|---|---|---|---|---|---|---|---|---|---|---|---|
| 550 | 421.316 | 2 | 131.21 | 0.0553 | 19.66 | 41.68 | 5 | 0.0 | 0 | K.VTVLGQPK.A | | 105-112 | CID |
| 1037 | 681.72 | 3 | 51.90 | 0.0354 | 29.56 | 48.4 | 9 | 0.0 | 0 | K.ANPTVTLFPPSSEELQANK.A | | 113-131 | CID |
| 1601 | 738.099 | 3 | 510.86 | 0.3769 | 40.24 | 101.69 | 3 | 0.0 | 0 | K.ATLVCLISDFYPGAVTVAWK.A | Carbamidomethyl: 5 | 132-151 | CID |
| 1071 | 872.447 | 2 | 15.99 | 0.0139 | 30.15 | 81.13 | 8 | 0.0 | 0 | K.YAASSYLSLTPEQWK.S | | 174-188 | CID |
| 408 | 571.281 | 3 | 40.41 | 0.0231 | 16.73 | 76.5 | 14 | 0.0 | 0 | R.SYSCQVTHEGSTVEK.T | Carbamidomethyl: 4 | 192-206 | CID |
| 624 | 853.027 | 3 | -20.19 | -0.0172 | 21.06 | 41.65 | 3 | 0.0 | 1 | R.SYSCQVTHEGSTVEKTVAPTECS.- | Carbamidomethyl: 4, 22 | 192-214 | CID |

#### Protein 8: Immunoglobulin kappa variable 2D-28 OS=Homo sapiens GN=IGKV2D-28 PE=1 SV=2

| | | | |
|---|---|---|---|
| **Accession:** | KVD28_HUMAN | **Score:** | 167.9 |
| **Database:** | SwissProt | **Seq. Coverage [%]:** | 16.7 |
| **MW [kDa] / pI:** | 12.9 / 5.6 | **No. of Peptides:** | 2 |

| 10 | 20 | 30 | 40 | 50 | 60 | 70 | 80 | 90 | 100 |
|---|---|---|---|---|---|---|---|---|---|
| MRLPAQLLGL | LMLWVSGSSG | DIVMTQSPLS | LPVTPGEPAS | ISCRSSQSLL | HSNGYNYLDW | YLQKPGQSPQ | LLIYLGSNRA | SGVPDRFSGS | GSGTDFTLKI |
| **110** | **120** | **130** | | | | | | | |
| SRVEAEDVGV | YYCMQALQTP | | | | | | | | |

MS/MS Peptide Matches

| Cmpd. | m/z meas. | z | Δ m/z [ppm] | Δ m/z [Da] | Rt [min] | Score | No. of Cmpds. | Site [%] | P | Sequence | Modification | Range | Type |
|---|---|---|---|---|---|---|---|---|---|---|---|---|---|
| 804 | 662.644 | 3 | -19.55 | -0.0130 | 24.79 | 65.71 | 3 | 0.0 | 1 | R.ASGVPDRFSGSGSGTDFTLK.I | | 80-99 | CID |
| 759 | 652.29 | 2 | -33.53 | -0.0219 | 23.86 | 102.23 | 3 | 0.0 | 0 | R.FSGSGSGTDFTLK.I | | 87-99 | CID |

#### Protein 9: Immunoglobulin heavy variable 3-30-3 OS=Homo sapiens GN=IGHV3-30-3 PE=3 SV=1

**Accession:** HVC33_HUMAN
**Database:** SwissProt
**MW [kDa] / pI:** 13.0 / 9.1
**Modification(s):** Carbamidomethyl

**Score:** 126.4
**Seq. Coverage [%]:** 15.4
**No. of Peptides:** 2

```
        10         20         30         40         50         60         70         80         90        100
MEFGLSWVFL VALLRGVQCQ VQLVESGGGV VQPGRSLRLS CAASGFTFSS YAMHWVRQAP GKGLEWVAVI SYDGSNKYYA DSVKGRFTIS RDNSKNTLYL
       110        120
QMNSLRAEDT AVYYCAR
```

MS/MS Peptide Matches

| Cmpd. | m/z meas. | z | Δ m/z [ppm] | Δ m/z [Da] | Rt [min] | Score | No. of Cmpds. | Site [%] | P | Sequence | Modification | Range | Type |
|---|---|---|---|---|---|---|---|---|---|---|---|---|---|
| 389 | 423.238 | 2 | 76.52 | 0.0324 | 16.33 | 36.31 | 4 | 0.0 | 0 | K.YYADSVK.G | | 78-84 | CID |
| 584 | 659.832 | 2 | 63.28 | 0.0417 | 20.29 | 90.1 | 3 | 0.0 | 0 | R.AEDTAVYYCAR.- | Carbamidomethyl: 9 | 107-117 | CID |

#### Protein 10: Immunoglobulin kappa variable 3-15 OS=Homo sapiens GN=IGKV3-15 PE=1 SV=2

**Accession:** KV315_HUMAN
**Database:** SwissProt
**MW [kDa] / pI:** 12.5 / 5.1

**Score:** 67.3
**Seq. Coverage [%]:** 7.8
**No. of Peptides:** 1

| 10 | 20 | 30 | 40 | 50 | 60 | 70 | 80 | 90 | 100 |
|---|---|---|---|---|---|---|---|---|---|
| MEAPAQLLFL | LLLWLPDTTG | EIVMTQSPAT | LSVSPGERAT | LSCRASQSVS | SNLAWYQQKP | GQAPRLLIYG | ASTRATGIPA | RFSGSGSGTE | FTLTISSLQS |
| 110 | 120 | | | | | | | | |
| EDFAVYYCQQ | YNNWP | | | | | | | | |

MS/MS Peptide Matches

| Cmpd. | m/z meas. | z | Δ m/z [ppm] | Δ m/z [Da] | Rt [min] | Score | No. of Cmpds. | Site [%] | P | Sequence | Modification | Range | Type |
|---|---|---|---|---|---|---|---|---|---|---|---|---|---|
| 779 | 497.327 | 2 | 74.37 | 0.0370 | 24.30 | 67.3 | 5 | 0.0 | 0 | R.LLIYGASTR.A | | 66-74 | CID |

#### Protein 11: Immunoglobulin kappa variable 3-20 OS=Homo sapiens GN=IGKV3-20 PE=1 SV=2

| | | | |
|---|---|---|---|
| **Accession:** | KV320_HUMAN | **Score:** | 65.5 |
| **Database:** | SwissProt | **Seq. Coverage [%]:** | 7.8 |
| **MW [kDa] / pI:** | 12.5 / 4.9 | **No. of Peptides:** | 1 |

| 10 | 20 | 30 | 40 | 50 | 60 | 70 | 80 | 90 | 100 |
|---|---|---|---|---|---|---|---|---|---|
| METPAQLLFL | LLLWLPDTTG | EIVLTQSPGT | LSLSPGERAT | LSCRASQSVS | SSYLAWYQQK | PGQAPRLLIY | GASSRATGIP | DRFSGSGSGT | DFTLTISRLE |
| 110 | 120 | | | | | | | | |
| PEDFAVYYCQ | QYGSSP | | | | | | | | |

MS/MS Peptide Matches

| Cmpd. | m/z meas. | z | Δ m/z [ppm] | Δ m/z [Da] | Rt [min] | Score | No. of Cmpds. | Site [%] | P | Sequence | Modification | Range | Type |
|---|---|---|---|---|---|---|---|---|---|---|---|---|---|
| 762 | 490.327 | 2 | 91.39 | 0.0448 | 23.93 | 65.49 | 4 | 0.0 | 0 | R.LLIYGASSR.A | | 67-75 | CID |

#### Protein 12: Immunoglobulin heavy constant alpha 1 OS=Homo sapiens GN=IGHA1 PE=1 SV=2

| | | | |
|---|---|---|---|
| **Accession:** | IGHA1_HUMAN | **Score:** | 43.9 |
| **Database:** | SwissProt | **Seq. Coverage [%]:** | 3.4 |
| **MW [kDa] / pI:** | 37.6 / 6.1 | **No. of Peptides:** | 1 |

**Modification(s):** Carbamidomethyl

| 10 | 20 | 30 | 40 | 50 | 60 | 70 | 80 | 90 | 100 |
|---|---|---|---|---|---|---|---|---|---|
| ASPTSPKVFP | LSLCSTQPDG | NVVIACLVQG | FFPQEPLSVT | WSESGQGVTA | RNFPPSQDAS | GDLYTTSSQL | TLPATQCLAG | KSVTCHVKHY | TNPSQDVTVP |
| **110** | **120** | **130** | **140** | **150** | **160** | **170** | **180** | **190** | **200** |
| CPVPSTPPTP | SPSTPPTPSP | SCCHPRLSLH | RPALEDLLLG | SEANLTCTLT | GLRDASGVTF | TWTPSSGKSA | VQGPPERDLC | GCYSVSSVLP | GCAEPWNHGK |
| **210** | **220** | **230** | **240** | **250** | **260** | **270** | **280** | **290** | **300** |
| TFTCTAAYPE | SKTPLTATLS | KSGNTFRPEV | HLLPPPSEEL | ALNELVTLTC | LARGFSPKDV | LVRWLQGSQE | LPREKYLTWA | SRQEPSQGTT | TFAVTSILRV |
| **310** | **320** | **330** | **340** | **350** | **360** | | | | |
| AAEDWKKGDT | FSCMVGHEAL | PLAFTQKTID | RLAGKPTHVN | VSVVMAEVDG | TCY | | | | |

MS/MS Peptide Matches

| Cmpd. | m/z meas. | z | Δ m/z [ppm] | Δ m/z [Da] | Rt [min] | Score | No. of Cmpds. | Site [%] | P | Sequence | Modification | Range | Type |
|---|---|---|---|---|---|---|---|---|---|---|---|---|---|
| 594 | 688.306 | 2 | -10.98 | -0.0076 | 20.46 | 43.85 | 1 | 0.0 | 0 | K.TFTCTAAYPESK.T | Carbamidomethyl: 4 | 201-212 | CID |

BRUKER

#### Project Info

| | | | |
|---|---|---|---|
| **Name:** | Alagesan | **Date:** | Mar 5, 2019 |
| **Note:** | Nanobooster | | |

#### Sample Info & Protocols

| | |
|---|---|
| **Name:** | Nanobooster - IgG |
| **Date:** | Mar 5, 2019 |

#### Search Result Info

| | | | |
|---|---|---|---|
| **Search Result:** | IonTrap_Human_Swiss_CID_2019-03-05 17:55:08 | | |
| **Location:** | /Alagesan/Nanobooster - IgG/KA_IgG1 HD NB ACN CID_GA2_01_9077.mgf | | |
| **Search Method:** | IonTrap_Human_Swiss_CID | **Version:** | 1.2 |
| **Search Engine(s):** | Mascot, 2.6.0 | **Ident. Compound(s):** | 64/2248 |
| **Database(s):** | SwissProt, SwissProt_2017_04.fasta | | |
| **Note:** | | | |

#### Results

##### Protein 1: Immunoglobulin kappa constant OS=Homo sapiens GN=IGKC PE=1 SV=2

**Accession:** IGKC_HUMAN
**Database:** SwissProt
**MW [kDa] / pI:** 11.8 / 6.1
**Modification(s):** Carbamidomethyl

**Score:** 1084.1
**Seq. Coverage [%]:** 86.0
**No. of Peptides:** 10

10 20 30 40 50 60 70 80 90 100
RTVAAPSVFI FPPSDEQLKS GTASVVCLLN NFYPREAKVQ WKVDNALQSG NSQESVTEQD SKDSTYSLSS TLTLSKADYE KHKVYACEVT HQGLSSPVTK
110
SFNRGEC

MS/MS Peptide Matches

| Cmpd. | m/z meas. | z | Δ m/z [ppm] | Δ m/z [Da] | Rt [min] | Score | No. of Cmpds. | Site [%] | P | Sequence | Modification | Range | Type |
|---|---|---|---|---|---|---|---|---|---|---|---|---|---|
| 929 | 701.449 | 3 | 97.13 | 0.0681 | 28.76 | 46.58 | 5 | 0.0 | 1 | -.RTVAAPSVFIFPPSDEQLK.S | | 1-19 | CID |
| 1033 | 973.588 | 2 | 72.81 | 0.0709 | 30.58 | 81.11 | 15 | 0.0 | 0 | R.TVAAPSVFIFPPSDEQLK.S | | 2-19 | CID |
| 1479 | 1242.789 | 3 | 388.55 | 0.4827 | 38.76 | 93.78 | 11 | 0.0 | 1 | R.TVAAPSVFIFPPSDEQLKSGTASVVCLLNNFYPR.E | Carbamidomethyl: 26 | 2-35 | CID |
| 1185 | 899.59 | 2 | 154.26 | 0.1387 | 33.12 | 67.59 | 17 | 0.0 | 0 | K.SGTASVVCLLNNFYPR.E | Carbamidomethyl: 8 | 20-35 | CID |
| 498 | 893.518 | 3 | 473.80 | 0.4232 | 21.12 | 128.16 | 9 | 0.0 | 1 | K.VQWKVDNALQSGNSQESVTEQDSK.D | | 39-62 | CID |
| 261 | 1068.956 | 2 | 438.00 | 0.4680 | 15.73 | 155.44 | 29 | 0.0 | 0 | K.VDNALQSGNSQESVTEQDSK.D | | 43-62 | CID |
| 686 | 1207.591 | 3 | 289.67 | 0.3497 | 24.57 | 152.96 | 2 | 0.0 | 1 | K.VDNALQSGNSQESVTEQDSKDSTYSLSSTLTLSK.A | | 43-76 | CID |
| 699 | 751.976 | 2 | 123.88 | 0.0931 | 24.82 | 111.01 | 3 | 0.0 | 0 | K.DSTYSLSSTLTLSK.A | | 63-76 | CID |
| 315 | 714.461 | 3 | 134.22 | 0.0959 | 17.05 | 147.37 | 13 | 0.0 | 1 | K.HKVYACEVTHQGLSSPVTK.S | Carbamidomethyl: 6 | 82-100 | CID |
| 398 | 939.001 | 2 | 568.91 | 0.5339 | 19.16 | 100.08 | 12 | 0.0 | 0 | K.VYACEVTHQGLSSPVTK.S | Carbamidomethyl: 4 | 84-100 | CID |

#### Protein 2: Immunoglobulin heavy constant gamma 1 OS=Homo sapiens GN=IGHG1 PE=1 SV=1

| | | | |
|---|---|---|---|
| **Accession:** | IGHG1_HUMAN | **Score:** | 1032.9 |
| **Database:** | SwissProt | **Seq. Coverage [%]:** | 58.2 |
| **MW [kDa] / pI:** | 36.1 / 8.5 | **No. of Peptides:** | 14 |
| **Modification(s):** | Carbamidomethyl | | |

| 10 | 20 | 30 | 40 | 50 | 60 | 70 | 80 | 90 | 100 |
|---|---|---|---|---|---|---|---|---|---|
| ASTKGPSVFP | LAPSSKSTSG | GTAALGCLVK | DYFPEPVTVS | WNSGALTSGV | HTFPAVLQSS | GLYSLSSVVT | VPSSSLGTQT | YICNVNHKPS | NTKVDKKVEP |
| **110** | **120** | **130** | **140** | **150** | **160** | **170** | **180** | **190** | **200** |
| KSCDKTHTCP | PCPAPELLGG | PSVFLFPPKP | KDTLMISRTP | EVTCVVVDVS | HEDPEVKFNW | YVDGVEVHNA | KTKPREEQYN | STYRVVSVLT | VLHQDWLNGK |
| **210** | **220** | **230** | **240** | **250** | **260** | **270** | **280** | **290** | **300** |
| EYKCKVSNKA | LPAPIEKTIS | KAKGQPREPQ | VYTLPPSRDE | LTKNQVSLTC | LVKGFYPSDI | AVEWESNGQP | ENNYKTTPPV | LDSDGSFFLY | SKLTVDKSRW |
| **310** | **320** | **330** | **340** | | | | | | |
| QQGNVFSCSV | MHEALHNHYT | QKSLSLSPGK | | | | | | | |

##### MS/MS Peptide Matches

| Cmpd. | m/z meas. | z | Δ m/z [ppm] | Δ m/z [Da] | Rt [min] | Score | No. of Cmpds. | Site [%] | P | Sequence | Modification | Range | Type |
|---|---|---|---|---|---|---|---|---|---|---|---|---|---|
| 898 | 830.533 | 3 | 111.43 | 0.0925 | 28.16 | 71.97 | 3 | 0.0 | 1 | K.GPSVFPLAPSSKSTSGGTAALGCLVK.D | Carbamidomethyl: 23 | 5-30 | CID |
| 1020 | 1112.582 | 3 | 326.47 | 0.3631 | 30.33 | 91.58 | 11 | 0.0 | 1 | K.SCDKTHTCPPCPAPELLGGPSVFLFPPKPK.D | Carbamidomethyl: 2, 8, 11 | 102-131 | CID |
| 1070 | 949.302 | 3 | 503.75 | 0.4780 | 31.22 | 88.4 | 7 | 0.0 | 0 | K.THTCPPCPAPELLGGPSVFLFPPKPK.D | Carbamidomethyl: 4, 7 | 106-131 | CID |
| 1095 | 916.336 | 4 | 395.28 | 0.3621 | 31.60 | 38.63 | 3 | 0.0 | 1 | K.THTCPPCPAPELLGGPSVFLFPPKPKDTLMIS R.T | Carbamidomethyl: 4, 7 | 106-138 | CID |
| 903 | 986.252 | 3 | 438.77 | 0.4325 | 28.22 | 25.17 | 1 | 0.0 | 1 | K.DTLMISRTPEVTCVVVDVSHEDPEVK.F | Carbamidomethyl: 13 | 132-157 | CID |
| 707 | 1070.07 | 2 | 49.20 | 0.0526 | 24.94 | 114.78 | 1 | 0.0 | 0 | R.TPEVTCVVVDVSHEDPEVK.F | Carbamidomethyl: 6 | 139-157 | CID |
| 990 | 1266.949 | 3 | 268.67 | 0.3403 | 29.82 | 50.02 | 3 | 0.0 | 1 | R.TPEVTCVVVDVSHEDPEVKFNWYVDGVEVHNA K.T | Carbamidomethyl: 6 | 139-171 | CID |

| Cmpd. | m/z meas. | z | Δ m/z [ppm] | Δ m/z [Da] | Rt [min] | Score | No. of Cmpds. | Site [%] | P | Sequence | Modification | Range | Type |
|---|---|---|---|---|---|---|---|---|---|---|---|---|---|
| 696 | 839.498 | 2 | 111.24 | 0.0934 | 24.75 | 92.26 | 4 | 0.0 | 0 | K.FNWYVDGVEVHNAK.T | | 158-171 | CID |
| 1139 | 905.11 | 2 | 666.79 | 0.6031 | 32.35 | 91.17 | 5 | 0.0 | 0 | R.VVSVLTVLHQDWLNGK.E | | 185-200 | CID |
| 515 | 937.07 | 2 | 86.74 | 0.0813 | 21.43 | 59.44 | 6 | 0.0 | 1 | R.EPQVYTLPPSRDELTK.N | | 228-243 | CID |
| 914 | 1273.22 | 2 | 511.31 | 0.6507 | 28.44 | 112.51 | 10 | 0.0 | 0 | K.GFYPSDIAVEWESNGQPENNYK.T | | 254-275 | CID |
| 979 | 937.628 | 2 | 174.35 | 0.1635 | 29.60 | 84.59 | 5 | 0.0 | 0 | K.TTPPVLDSDGSFFLYSK.L | | 276-292 | CID |
| 606 | 762.198 | 4 | 449.53 | 0.3425 | 23.22 | 38.07 | 3 | 0.0 | 1 | K.SRWQQGNVFSCSVMHEALHNHYTQK.S | Carbamidomethyl: 11 | 298-322 | CID |
| 650 | 934.545 | 3 | 126.04 | 0.1178 | 24.00 | 74.28 | 4 | 0.0 | 0 | R.WQQGNVFSCSVMHEALHNHYTQK.S | Carbamidomethyl: 9 | 300-322 | CID |

#### Protein 3: Immunoglobulin kappa light chain OS=Homo sapiens PE=1 SV=1

**Accession:** IGK_HUMAN
**Database:** SwissProt
**MW [kDa] / pI:** 23.4 / 6.9
**Modification(s):** Carbamidomethyl

**Score:** 985.8
**Seq. Coverage [%]:** 42.5
**No. of Peptides:** 8

```
        10         20         30         40         50         60         70         80         90        100
DIQMTQSPST LSASVGDRVT ITCRASQSIN TWLAWYQQKP GKAPKLLMYK ASSLESGVPS RFIGSGSGTE FTLTISSLQP DDFATYYCQQ YNSDSKMFGQ
       110        120        130        140        150        160        170        180        190        200
GTKVEVKGTV AAPSVFIFPP SDEQLKSGTA SVVCLLNNFY PREAKVQWKV DNALQSGNSQ ESVTEQDSKD STYSLSSTLT LSKADYEKHK VYACEVTHQG
       210        220
LSSPVTKSFN RGEC
```

MS/MS Peptide Matches

| Cmpd. | m/z meas. | z | Δ m/z [ppm] | Δ m/z [Da] | Rt [min] | Score | No. of Cmpds. | Site [%] | P | Sequence | Modification | Range | Type |
|---|---|---|---|---|---|---|---|---|---|---|---|---|---|
| 556 | 947.03 | 2 | 79.67 | 0.0754 | 22.30 | 123.2 | 2 | 0.0 | 0 | -.DIQMTQSPSTLSASVGDR.V | | 1-18 | CID |
| 1185 | 899.59 | 2 | 154.26 | 0.1387 | 33.12 | 67.59 | 17 | 0.0 | 0 | K.SGTASVVCLLNNFYPR.E | Carbamidomethyl: 8 | 127-142 | CID |

| Cmpd. | m/z meas. | z | Δ m/z [ppm] | Δ m/z [Da] | Rt [min] | Score | No. of Cmpds. | Site [%] | P | Sequence | Modification | Range | Type |
|---|---|---|---|---|---|---|---|---|---|---|---|---|---|
| 498 | 893.518 | 3 | 473.80 | 0.4232 | 21.12 | 128.16 | 9 | 0.0 | 1 | K.VQWKVDNALQSGNSQESVTEQDSK.D | | 146-169 | CID |
| 261 | 1068.956 | 2 | 438.00 | 0.4680 | 15.73 | 155.44 | 29 | 0.0 | 0 | K.VDNALQSGNSQESVTEQDSK.D | | 150-169 | CID |
| 686 | 1207.591 | 3 | 289.67 | 0.3497 | 24.57 | 152.96 | 2 | 0.0 | 1 | K.VDNALQSGNSQESVTEQDSKDSTYSLSSTLTLSK.A | | 150-183 | CID |
| 699 | 751.976 | 2 | 123.88 | 0.0931 | 24.82 | 111.01 | 3 | 0.0 | 0 | K.DSTYSLSSTLTLSK.A | | 170-183 | CID |
| 315 | 714.461 | 3 | 134.22 | 0.0959 | 17.05 | 147.37 | 13 | 0.0 | 1 | K.HKVYACEVTHQGLSSPVTK.S | Carbamidomethyl: 6 | 189-207 | CID |
| 398 | 939.001 | 2 | 568.91 | 0.5339 | 19.16 | 100.08 | 12 | 0.0 | 0 | K.VYACEVTHQGLSSPVTK.S | Carbamidomethyl: 4 | 191-207 | CID |

##### Protein 4: Immunoglobulin lambda constant 3 OS=Homo sapiens GN=IGLC3 PE=1 SV=1

**Accession:** IGLC3_HUMAN
**Database:** SwissProt
**MW [kDa] / pI:** 11.3 / 6.9
**Modification(s):** Carbamidomethyl

**Score:** 658.2
**Seq. Coverage [%]:** 96.2
**No. of Peptides:** 9

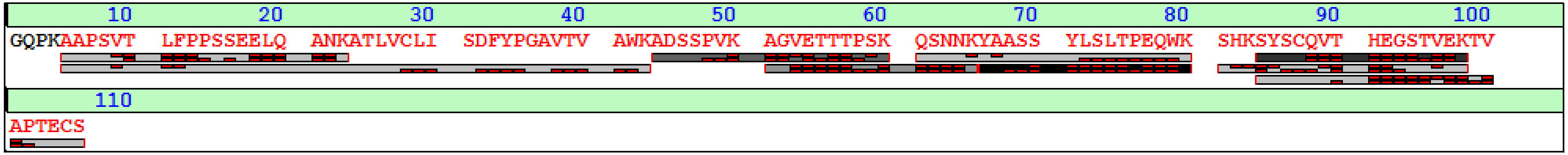


MS/MS Peptide Matches

| Cmpd. | m/z meas. | z | Δ m/z [ppm] | Δ m/z [Da] | Rt [min] | Score | No. of Cmpds. | Site [%] | P | Sequence | Modification | Range | Type |
|---|---|---|---|---|---|---|---|---|---|---|---|---|---|
| 803 | 993.58 | 2 | 67.88 | 0.0674 | 26.53 | 74.91 | 9 | 0.0 | 0 | K.AAPSVTLFPPSSEELQANK.A | | 5-23 | CID |
| 1538 | 1393.645 | 3 | 183.87 | 0.2562 | 39.94 | 24.96 | 2 | 0.0 | 1 | K.AAPSVTLFPPSSEELQANKATLVCLISDFYPGAVTVAWK.A | Carbamidomethyl: 24 | 5-43 | CID |
| 232 | 838.045 | 2 | 136.20 | 0.1141 | 15.07 | 57.48 | 3 | 0.0 | 1 | K.ADSSPVKAGVETTTPSK.Q | | 44-60 | CID |
| 58 | 781.514 | 2 | 152.99 | 0.1195 | 10.91 | 80.96 | 15 | 0.0 | 1 | K.AGVETTTPSKQSNNK.Y | | 51-65 | CID |

| Cmpd. | m/z meas. | z | Δ m/z [ppm] | Δ m/z [Da] | Rt [min] | Score | No. of Cmpds. | Site [%] | P | Sequence | Modification | Range | Type |
|---|---|---|---|---|---|---|---|---|---|---|---|---|---|
| 798 | 772.812 | 3 | 557.24 | 0.4304 | 26.47 | 57.3 | 6 | 0.0 | 1 | K.QSNNKYAASSYLSLTPEQWK.S | | 61-80 | CID |
| 873 | 872.54 | 2 | 122.59 | 0.1069 | 27.70 | 97.03 | 10 | 0.0 | 0 | K.YAASSYLSLTPEQWK.S | | 66-80 | CID |
| 122 | 1032.585 | 2 | 105.39 | 0.1088 | 12.44 | 61.36 | 4 | 0.0 | 1 | K.SHKSYSCQVTHEGSTVEK.T | Carbamidomethyl: 7 | 81-98 | CID |
| 196 | 856.533 | 2 | 174.88 | 0.1498 | 14.24 | 111.23 | 14 | 0.0 | 0 | K.SYSCQVTHEGSTVEK.T | Carbamidomethyl: 4 | 84-98 | CID |
| 389 | 1279.56 | 2 | 388.81 | 0.4973 | 18.97 | 93.01 | 10 | 0.0 | 1 | K.SYSCQVTHEGSTVEKTVAPTECS.- | Carbamidomethyl: 4, 22 | 84-106 | CID |

#### Protein 5: Keratin, type II cytoskeletal 1 OS=Homo sapiens GN=KRT1 PE=1 SV=6

**Accession:** K2C1_HUMAN
**Database:** SwissProt
**MW [kDa] / pI:** 66.0 / 8.2

**Score:** 605.6
**Seq. Coverage [%]:** 18.3
**No. of Peptides:** 6

| | | | | | | | | | |
|---|---|---|---|---|---|---|---|---|---|
| 10 | 20 | 30 | 40 | 50 | 60 | 70 | 80 | 90 | 100 |
| MSRQFSSRSG | YRSGGGFSSG | SAGIINYQRR | TTSSSTRRSG | GGGGRFSSCG | GGGGSFGAGG | GFGSRSLVNL | GGSKSISISV | ARGGGRGSGF | GGGYGGGGFG |
| 110 | 120 | 130 | 140 | 150 | 160 | 170 | 180 | 190 | 200 |
| GGGFGGGGFG | GGGIGGGGFG | GFGSGGGGFG | GGGFGGGGYG | GGYGPVCPPG | GIQEVTINQS | LLQPLNVEID | PEIQKVKSRE | REQIKSLNNQ | FASFIDKVRF |
| 210 | 220 | 230 | 240 | 250 | 260 | 270 | 280 | 290 | 300 |
| LEQQNQVLQT | KWELLQQVDT | STRTHNLEPY | FESFINNLRR | RVDQLKSDQS | RLDSELKNMQ | DMVEDYRNKY | EDEINKRTNA | ENEFVTIKKD | VDGAYMTKVD |
| 310 | 320 | 330 | 340 | 350 | 360 | 370 | 380 | 390 | 400 |
| LQAKLDNLQQ | EIDFLTALYQ | AELSQMQTQI | SETNVILSMD | NNRSLDLDSI | IAEVKAQYED | IAQKSKAEAE | SLYQSKYEEL | QITAGRHGDS | VRNSKIEISE |
| 410 | 420 | 430 | 440 | 450 | 460 | 470 | 480 | 490 | 500 |
| LNRVIQRLRS | EIDNVKKQIS | NLQQSISDAE | QRGENALKDA | KNKLNDLEDA | LQQAKEDLAR | LLRDYQELMN | TKLALDLEIA | TYRTLLEGEE | SRMSGECAPN |
| 510 | 520 | 530 | 540 | 550 | 560 | 570 | 580 | 590 | 600 |
| VSVSVSTSHT | TISGGGSRGG | GGGGYGSGGS | SYGSGGGSYG | SGGGGGGGRG | SYGSGGSSYG | SGGGSYGSGG | GGGGHGSYGS | GSSSGGYRGG | SGGGGGGSSG |
| 610 | 620 | 630 | 640 | 650 | | | | | |
| GRGSGGGSSG | GSIGGRGSSS | GGVKSSGGSS | SVKFVSTTYS | GVTR | | | | | |

MS/MS Peptide Matches

| Cmpd. | m/z meas. | z | Δ m/z [ppm] | Δ m/z [Da] | Rt [min] | Score | No. of Cmpds. | Site [%] | P | Sequence | Modification | Range | Type |
|---|---|---|---|---|---|---|---|---|---|---|---|---|---|
| 536 | 829.484 | 2 | 101.19 | 0.0839 | 21.85 | 49.13 | 1 | 0.0 | 0 | R.SGGGFSSGSAGIINYQR.R | | 13-29 | CID |
| 444 | 738.501 | 2 | 141.83 | 0.1047 | 20.11 | 88.21 | 6 | 0.0 | 0 | R.FLEQQNQVLQTK.W | | 200-211 | CID |
| 1428 | 783.83 | 3 | 109.33 | 0.0857 | 37.75 | 151.05 | 4 | 0.0 | 1 | R.SLDLDSIIAEVKAQYEDIAQK.S | | 344-364 | CID |
| 524 | 858.997 | 2 | 78.94 | 0.0678 | 21.60 | 107.51 | 3 | 0.0 | 0 | K.QISNLQQSISDAEQR.G | | 418-432 | CID |
| 201 | 1192.583 | 2 | 86.74 | 0.1034 | 14.36 | 189.29 | 5 | 0.0 | 0 | R.GGGGGGYGSGGSSYGSGGGSYGSGGGGGG GR.G | | 519-549 | CID |
| 69 | 876.54 | 2 | 140.36 | 0.1230 | 11.25 | 20.44 | 1 | 0.0 | 1 | R.GSGGGSSGGSIGGRGSSSGGVK.S | | 603-624 | CID |

#### Protein 6: Keratin, type I cytoskeletal 9 OS=Homo sapiens GN=KRT9 PE=1 SV=3

**Accession:** K1C9_HUMAN
**Database:** SwissProt
**MW [kDa] / pI:** 62.0 / 5.1

**Score:** 513.5
**Seq. Coverage [%]:** 23.0
**No. of Peptides:** 6

| 10 | 20 | 30 | 40 | 50 | 60 | 70 | 80 | 90 | 100 |
|---|---|---|---|---|---|---|---|---|---|
| MSCRQFSSSY | LSRSGGGGGG | GLGSGGSIRS | SYSRFSSSGG | GGGGGRFSSS | SGYGGGSSRV | CGRGGGGSFG | YSYGGGSGGG | FSASSLGGGF | GGGSRGFGGA |
| 110 | 120 | 130 | 140 | 150 | 160 | 170 | 180 | 190 | 200 |
| SGGGYSSSGG | FGGGFGGGSG | GGFGGGYGSG | FGGFGGFGGG | AGGGDGGILT | ANEKSTMQEL | NSRLASYLDK | VQALEEANND | LENKIQDWYD | KKGPAAIQKN |
| 210 | 220 | 230 | 240 | 250 | 260 | 270 | 280 | 290 | 300 |
| YSPYYNTIDD | LKDQIVDLTV | GNNKTLLDID | NTRMTLDDFR | IKFEMEQNLR | QGVDADINGL | RQVLDNLTME | KSDLEMQYET | LQEELMALKK | NHKEEMSQLT |
| 310 | 320 | 330 | 340 | 350 | 360 | 370 | 380 | 390 | 400 |
| GQNSGDVNVE | INVAPGKDLT | KTLNDMRQEY | EQLIAKNRKD | IENQYETQIT | QIEHEVSSSG | QEVQSSAKEV | TQLRHGVQEL | EIELQSQLSK | KAALEKSLED |
| 410 | 420 | 430 | 440 | 450 | 460 | 470 | 480 | 490 | 500 |
| TKNRYCGQLQ | MIQEQISNLE | AQITDVRQEI | ECQNQEYSLL | LSIKMRLEKE | IETYHNLLEG | GQEDFESSGA | GKIGLGGRGG | SGGSYGRGSR | GGSGGSYGGG |
| 510 | 520 | 530 | 540 | 550 | 560 | 570 | 580 | 590 | 600 |
| GSGGGYGGGS | GSRGGSGGSY | GGGSGSGGGS | GGGYGGGSGG | GHSGGSGGGH | SGGSGGNYGG | GSGSGGGSGG | GYGGGSGSRG | GSGGSHGGGS | GFGGESGGSY |
| 610 | 620 | 630 | | | | | | | |
| GGGEEASGSG | GGYGGGSGKS | SHS | | | | | | | |

**MS/MS Peptide Matches**

| Cmpd. | m/z meas. | z | Δ m/z [ppm] | Δ m/z [Da] | Rt [min] | Score | No. of Cmpds. | Site [%] | P | Sequence | Modification | Range | Type |
|---|---|---|---|---|---|---|---|---|---|---|---|---|---|
| 1229 | 968.234 | 3 | 95.35 | 0.0923 | 33.93 | 65.55 | 2 | 0.0 | 1 | K.NYSPYYNTIDDLKDQIVDLTVGNNK.T | | 200-224 | CID |
| 996 | 1089.212 | 3 | 339.07 | 0.3692 | 29.92 | 107.67 | 1 | 0.0 | 0 | K.DIENQYETQITQIEHEVSSSGQEVQSSAK.E | | 340-368 | CID |
| 1123 | 1331.267 | 3 | 217.92 | 0.2900 | 32.06 | 25.06 | 1 | 0.0 | 1 | K.DIENQYETQITQIEHEVSSSGQEVQSSAKEVTQLR.H | | 340-374 | CID |
| 875 | 920.089 | 2 | 655.42 | 0.6027 | 27.72 | 110.0 | 1 | 0.0 | 0 | R.HGVQELEIELQSQLSK.K | | 375-390 | CID |
| 113 | 896.458 | 2 | 100.97 | 0.0905 | 12.29 | 167.08 | 3 | 0.0 | 0 | R.GGSGGSYGGGGSGGGYGGGSGSR.G | | 491-513 | CID |
| 231 | 906.486 | 4 | 409.88 | 0.3714 | 15.04 | 38.15 | 1 | 0.0 | 1 | R.GGSGGSHGGGSGFGGESGGSYGGGEEASGSGGGYGGGSGKSSHS.- | | 580-623 | CID |

#### Protein 7: Immunoglobulin heavy constant gamma 4 OS=Homo sapiens GN=IGHG4 PE=1 SV=1

| | | | |
|---|---|---|---|
| **Accession:** | IGHG4_HUMAN | **Score:** | 394.2 |
| **Database:** | SwissProt | **Seq. Coverage [%]:** | 35.5 |
| **MW [kDa] / pI:** | 35.9 / 7.2 | **No. of Peptides:** | 5 |
| **Modification(s):** | Carbamidomethyl | | |

| 10 | 20 | 30 | 40 | 50 | 60 | 70 | 80 | 90 | 100 |
|---|---|---|---|---|---|---|---|---|---|
| ASTKGPSVFP | LAPCSRSTSE | STAALGCLVK | DYFPEPVTVS | WNSGALTSGV | HTFPAVLQSS | GLYSLSSVVT | VPSSSLGTKT | YTCNVDHKPS | NTKVDKRVES |
| **110** | **120** | **130** | **140** | **150** | **160** | **170** | **180** | **190** | **200** |
| KYGPPCPSCP | APEFLGGPSV | FLFPPKPKDT | LMISRTPEVT | CVVVDVSQED | PEVQFNWYVD | GVEVHNAKTK | PREEQFNSTY | RVVSVLTVLH | QDWLNGKEYK |
| **210** | **220** | **230** | **240** | **250** | **260** | **270** | **280** | **290** | **300** |
| CKVSNKGLPS | SIEKTISKAK | GQPREPQVYT | LPPSQEEMTK | NQVSLTCLVK | GFYPSDIAVE | WESNGQPENN | YKTTPPVLDS | DGSFFLYSRL | TVDKSRWQEG |
| **310** | **320** | **330** | | | | | | | |
| NVFSCSVMHE | ALHNHYTQKS | LSLSLGK | | | | | | | |

MS/MS Peptide Matches

| Cmpd. | m/z meas. | z | Δ m/z [ppm] | Δ m/z [Da] | Rt [min] | Score | No. of Cmpds. | Site [%] | P | Sequence | Modification | Range | Type |
|---|---|---|---|---|---|---|---|---|---|---|---|---|---|
| 574 | 712.43 | 2 | 100.37 | 0.0715 | 22.63 | 78.96 | 3 | 0.0 | 0 | R.STSESTAALGCLVK.D | Carbamidomethyl: 11 | 17-30 | CID |
| 72 | 833.034 | 2 | 174.71 | 0.1455 | 11.34 | 52.27 | 6 | 0.0 | 0 | K.TYTCNVDHKPSNTK.V | Carbamidomethyl: 4 | 80-93 | CID |
| 1182 | 982.259 | 3 | 443.14 | 0.4351 | 33.04 | 32.22 | 1 | 0.0 | 0 | K.YGPPCPSCPAPEFLGGPSVFLFPPKPK.D | Carbamidomethyl: 5, 8 | 102-128 | CID |
| 1139 | 905.11 | 2 | 666.79 | 0.6031 | 32.35 | 91.17 | 5 | 0.0 | 0 | R.VVSVLTVLHQDWLNGK.E | | 182-197 | CID |
| 914 | 1273.22 | 2 | 511.31 | 0.6507 | 28.44 | 112.51 | 10 | 0.0 | 0 | K.GFYPSDIAVEWESNGQPENNYK.T | | 251-272 | CID |

#### Protein 8: Immunoglobulin heavy constant gamma 2 OS=Homo sapiens GN=IGHG2 PE=1 SV=2

| | | | |
|---|---|---|---|
| **Accession:** | IGHG2_HUMAN | **Score:** | 369.0 |
| **Database:** | SwissProt | **Seq. Coverage [%]:** | 37.4 |
| **MW [kDa] / pI:** | 35.9 / 7.7 | **No. of Peptides:** | 6 |

**Modification(s):** Carbamidomethyl

```
        10         20         30         40         50         60         70         80         90        100
ASTKGPSVFP LAPCSRSTSE STAALGCLVK DYFPEPVTVS WNSGALTSGV HTFPAVLQSS GLYSLSSVVT VPSSNFGTQT YTCNVDHKPS NTKVDKTVER
       110        120        130        140        150        160        170        180        190        200
KCCVECPPCP APPVAGPSVF LFPPKPKDTL MISRTPEVTC VVVDVSHEDP EVQFNWYVDG VEVHNAKTKP REEQFNSTFR VVSVLTVVHQ DWLNGKEYKC
       210        220        230        240        250        260        270        280        290        300
KVSNKGLPAP IEKTISKTKG QPREPQVYTL PPSREEMTKN QVSLTCLVKG FYPSDISVEW ESNGQPENNY KTTPPMLDSD GSFFLYSKLT VDKSRWQQGN
       310        320        330
VFSCSVMHEA LHNHYTQKSL SLSPGK
```

**MS/MS Peptide Matches**

| Cmpd. | m/z meas. | z | Δ m/z [ppm] | Δ m/z [Da] | Rt [min] | Score | No. of Cmpds. | Site [%] | P | Sequence | Modification | Range | Type |
|---|---|---|---|---|---|---|---|---|---|---|---|---|---|
| 574 | 712.43 | 2 | 100.37 | 0.0715 | 22.63 | 78.96 | 3 | 0.0 | 0 | R.STSESTAALGCLVK.D | Carbamidomethyl: 11 | 17-30 | CID |
| 936 | 760.213 | 4 | 438.73 | 0.3334 | 28.91 | 18.38 | 1 | 0.0 | 1 | R.KCCVECPPCPAPPVAGPSVFLFPPKPK.D | Carbamidomethyl: 2, 3, 6, 9 | 101-127 | CID |
| 1078 | 1154.604 | 4 | 260.46 | 0.3007 | 31.32 | 50.12 | 1 | 0.0 | 1 | K.DTLMISRTPEVTCVVVDVSHEDPEVQFNWYVDGVEVHNAK.T | Carbamidomethyl: 13 | 128-167 | CID |
| 456 | 953.029 | 2 | 56.92 | 0.0542 | 20.34 | 62.51 | 12 | 0.0 | 1 | R.EPQVYTLPPSREEMTK.N | | 224-239 | CID |
| 606 | 762.198 | 4 | 449.53 | 0.3425 | 23.22 | 38.07 | 3 | 0.0 | 1 | K.SRWQQGNVFSCSVMHEALHNHYTQK.S | Carbamidomethyl: 11 | 294-318 | CID |
| 650 | 934.545 | 3 | 126.04 | 0.1178 | 24.00 | 74.28 | 4 | 0.0 | 0 | R.WQQGNVFSCSVMHEALHNHYTQK.S | Carbamidomethyl: 9 | 296-318 | CID |

#### Protein 9: Keratin, type II cytoskeletal 2 epidermal OS=Homo sapiens GN=KRT2 PE=1 SV=2

| | | | |
|---|---|---|---|
| **Accession:** | K22E_HUMAN | **Score:** | 342.7 |
| **Database:** | SwissProt | **Seq. Coverage [%]:** | 8.6 |
| **MW [kDa] / pI:** | 65.4 / 8.1 | **No. of Peptides:** | 3 |

| 10 | 20 | 30 | 40 | 50 | 60 | 70 | 80 | 90 | 100 |
|---|---|---|---|---|---|---|---|---|---|
| MSCQISCKSR | GRGGGGGGFR | GFSSGSAVVS | GGSRRSTSSF | SCLSRHGGGG | GGFGGGGFGS | RSLVGLGGTK | SISISVAGGG | GGFGAAGGFG | GRGGGFGGGS |
| **110** | **120** | **130** | **140** | **150** | **160** | **170** | **180** | **190** | **200** |
| SFGGGSGFSG | GGFGGGGFGG | GRFGGFGGPG | GVGGLGGPGG | FGPGGYPGGI | HEVSVNQSLL | QPLNVKVDPE | IQNVKAQERE | QIKTLNNKFA | SFIDKVRFLE |
| **210** | **220** | **230** | **240** | **250** | **260** | **270** | **280** | **290** | **300** |
| QQNQVLQTKW | ELLQQMNVGT | RPINLEPIFQ | GYIDSLKRYL | DGLTAERTSQ | NSELNNMQDL | VEDYKKKYED | EINKRTAAEN | DFVTLKKDVD | NAYMIKVELQ |
| **310** | **320** | **330** | **340** | **350** | **360** | **370** | **380** | **390** | **400** |
| SKVDLLNQEI | EFLKVLYDAE | ISQIHQSVTD | TNVILSMDNS | RNLDLDSIIA | EVKAQYEEIA | QRSKEEAEAL | YHSKYEELQV | TVGRHGDSLK | EIKIEISELN |
| **410** | **420** | **430** | **440** | **450** | **460** | **470** | **480** | **490** | **500** |
| RVIQRLQGEI | AHVKKQCKNV | QDAIADAEQR | GEHALKDARN | KLNDLEEALQ | QAKEDLARLL | RDYQELMNVK | LALDVEIATY | RKLLEGEECR | MSGDLSSNVT |
| **510** | **520** | **530** | **540** | **550** | **560** | **570** | **580** | **590** | **600** |
| VSVTSSTISS | NVASKAAFGG | SGGRGSSSGG | GYSSGSSSYG | SGGRQSGSRG | GSGGGGSISG | GGYGSGGGSG | GRYGSGGGSK | GGSISGGGYG | SGGGKHSSGG |
| **610** | **620** | **630** | **640** | | | | | | |
| GSRGGSSSGG | GYGSGGGSS | SVKGSSGEAF | GSSVTFSFR | | | | | | |

**MS/MS Peptide Matches**

| Cmpd. | m/z meas. | z | Δ m/z [ppm] | Δ m/z [Da] | Rt [min] | Score | No. of Cmpds. | Site [%] | P | Sequence | Modification | Range | Type |
|---|---|---|---|---|---|---|---|---|---|---|---|---|---|
| 444 | 738.501 | 2 | 141.83 | 0.1047 | 20.11 | 88.21 | 6 | 0.0 | 0 | R.FLEQQNQVLQTK.W | | 198-209 | CID |
| 123 | 870.943 | 2 | 99.41 | 0.0866 | 12.49 | 99.22 | 3 | 0.0 | 0 | R.GSSSGGGYSSGSSSYGSGGR.Q | | 525-544 | CID |
| 154 | 871.49 | 2 | 128.69 | 0.1121 | 13.14 | 155.25 | 3 | 0.0 | 0 | R.GGSGGGGSISGGGYGSGGGSGGR.Y | | 550-572 | CID |

#### Protein 10: Keratin, type I cytoskeletal 10 OS=Homo sapiens GN=KRT10 PE=1 SV=6

**Accession:** K1C10_HUMAN
**Database:** SwissProt
**MW [kDa] / pI:** 58.8 / 5.1

**Score:** 290.0
**Seq. Coverage [%]:** 9.9
**No. of Peptides:** 3

| 10 | 20 | 30 | 40 | 50 | 60 | 70 | 80 | 90 | 100 |
|---|---|---|---|---|---|---|---|---|---|
| MSVRYSSSKH | YSSSRSGGGG | GGGGCGGGGG | VSSLRISSSK | GSLGGGFSSG | GFSGGSFSRG | SSGGGCFGGS | SGGYGGLGGF | GGGSFRGSYG | SSSFGGSYGG |
| 110 | 120 | 130 | 140 | 150 | 160 | 170 | 180 | 190 | 200 |
| IFGGGSFGGG | SFGGGSFGGG | GFGGGGFGGG | FGGGFGGDGG | LLSGNEKVTM | QNLNDRLASY | LDKVRALEES | NYELEGKIKE | WYEKHGNSHQ | GEPRDYSKYY |
| 210 | 220 | 230 | 240 | 250 | 260 | 270 | 280 | 290 | 300 |
| KTIDDLKNQI | LNLTTDNANI | LLQIDNARLA | ADDFRLKYEN | EVALRQSVEA | DINGLRRVLD | ELTLTKADLE | MQIESLTEEL | AYLKKNHEEE | MKDLRNVSTG |
| 310 | 320 | 330 | 340 | 350 | 360 | 370 | 380 | 390 | 400 |
| DVNVEMNAAP | GVDLTQLLNN | MRSQYEQLAE | QNRKDAEAWF | NEKSKELTTE | IDNNIEQISS | YKSEITELRR | NVQALEIELQ | SQLALKQSLE | ASLAETEGRY |
| 410 | 420 | 430 | 440 | 450 | 460 | 470 | 480 | 490 | 500 |
| CVQLSQIQAQ | ISALEEQLQQ | IRAETECQNT | EYQQLLDIKI | RLENEIQTYR | SLLEGEGSSG | GGGRGGGSFG | GGYGGGSSGG | GSSGGGHGGG | HGGSSGGGYG |
| 510 | 520 | 530 | 540 | 550 | 560 | 570 | 580 | 590 | |
| GGSSGGGSSG | GGYGGGSSSG | GHGGSSSGGY | GGGSSGGGGG | GYGGGSSGGG | SSSGGGYGGG | SSSGGHKSSS | SGSVGESSSK | GPRY | |

MS/MS Peptide Matches

| Cmpd. | m/z meas. | z | Δ m/z [ppm] | Δ m/z [Da] | Rt [min] | Score | No. of Cmpds. | Site [%] | P | Sequence | Modification | Range | Type |
|---|---|---|---|---|---|---|---|---|---|---|---|---|---|
| 638 | 854.494 | 2 | 122.07 | 0.1043 | 23.81 | 135.06 | 3 | 0.0 | 0 | K.GSLGGGFSSGGFSGGSFSR.G | | 41-59 | CID |
| 1274 | 1437.125 | 2 | 295.80 | 0.4250 | 34.80 | 125.89 | 3 | 0.0 | 0 | R.NVSTGDVNVEMNAAPGVDLTQLLNNMR.S | | 296-322 | CID |
| 227 | 747.426 | 2 | 73.88 | 0.0552 | 14.95 | 29.02 | 4 | 0.0 | 1 | R.SQYEQLAEQNRK.D | | 323-334 | CID |

#### Protein 11: Hornerin OS=Homo sapiens GN=HRNR PE=1 SV=2

| | | | |
|---|---|---|---|
| **Accession:** | HORN_HUMAN | **Score:** | 127.9 |
| **Database:** | SwissProt | **Seq. Coverage [%]:** | 2.6 |
| **MW [kDa] / pI:** | 282.2 / 10.0 | **No. of Peptides:** | 3 |

| | | | | | | | | | |
|---|---|---|---|---|---|---|---|---|---|
| 10 | 20 | 30 | 40 | 50 | 60 | 70 | 80 | 90 | 100 |
| MPKLLQGVIT | VIDVFYQYAT | QHGEYDTLNK | AELKELLENE | FHQILKNPND | PDTVDIILQS | LDRDHNKKVD | FTEYLLMIFK | LVQARNKIIG | KDYCQVSGSK |
| 110 | 120 | 130 | 140 | 150 | 160 | 170 | 180 | 190 | 200 |
| LRDDTHQHQE | EQEETEKEEN | KRQESSFSHS | SWSAGENDSY | SRNVRGSLKP | GTESISRRLS | FQRDFSGQHN | SYSGQSSSYG | EQNSDSHQSS | GRGQCGSGSG |
| 210 | 220 | 230 | 240 | 250 | 260 | 270 | 280 | 290 | 300 |
| QSPNYGQHGS | GSGQSSSNDT | HGSGSGQSSG | FSQHKSSSGQ | SSGYSQHGSG | SGHSSGYGQH | GSRSGQSSRG | ERHRSSSGSS | SSYGQHGSGS | RQSLGHGRQG |
| 310 | 320 | 330 | 340 | 350 | 360 | 370 | 380 | 390 | 400 |
| SGSRQSPSHV | RHGSGSGHSS | SHGQHGSGSS | YSYSRGHYES | GSGQTSGFGQ | HESGSGQSSG | YSKHGSGSGH | SSSQGQHGST | SGQASSSGQH | GSSSRQSSSY |
| 410 | 420 | 430 | 440 | 450 | 460 | 470 | 480 | 490 | 500 |
| GQHESASRHS | SGRGQHSSGS | GQSPGHGQRG | SGSGQSPSSG | QHGTGFGRSS | SSGPYVSGSG | YSSGFGHHES | SSEHSSGYTQ | HGSGSGHSSG | HGQHGSRSGQ |
| 510 | 520 | 530 | 540 | 550 | 560 | 570 | 580 | 590 | 600 |
| SSRGERQGSS | AGSSSSYGQH | GSGSRQSLGH | SRHGSGSGQS | PSPSRGRHES | GSRQSSSYGP | HGYGSGRSSS | RGPYESGSGH | SSGLGHQESR | SGQSSGYGQH |
| 610 | 620 | 630 | 640 | 650 | 660 | 670 | 680 | 690 | 700 |
| GSSSGHSSTH | GQHGSTSGQS | SSCGQHGATS | GQSSSHGQHG | SGSSQSSRYG | QQGSGSGQSP | SRGRHGSDFG | HSSSYGQHGS | GSGWSSSNGP | HGSVSGQSSG |
| 710 | 720 | 730 | 740 | 750 | 760 | 770 | 780 | 790 | 800 |
| FGHKSGSGQS | SGYSQHGSGS | SHSSGYRKHG | SRSGQSSRSE | QHGSSSGLSS | SYGQHGSGSH | QSSGHGRQGS | GSGHSPSRVR | HGSSSGHSSS | HGQHGSGTSC |
| 810 | 820 | 830 | 840 | 850 | 860 | 870 | 880 | 890 | 900 |
| SSSCGHYESG | SGQASGFGQH | ESGSGQGYSQ | HGSASGHFSS | QGRHGSTSGQ | SSSSGQHDSS | SGQSSSYGQH | ESASHHASGR | GRHGSGSGQS | PGHGQRGSGS |
| 910 | 920 | 930 | 940 | 950 | 960 | 970 | 980 | 990 | 1000 |
| GQSPSYGRHG | SGSGRSSSSG | RHGSGSGQSS | GFGHKSSSGQ | SSGYTQHGSG | SGHSSSYEQH | GSRSGQSSRS | EQHGSSSGSS | SSYGQHGSGS | RQSLGHGQHG |
| 1010 | 1020 | 1030 | 1040 | 1050 | 1060 | 1070 | 1080 | 1090 | 1100 |
| SGSGQSPSPS | RGRHGSGSGQ | SSSYGPYRSG | SGWSSSRGPY | ESGSGHSSGL | GHRESRSGQS | SGYGQHGSSS | GHSSTHGQHG | STSGQSSSCG | QHGASSGQSS |
| 1110 | 1120 | 1130 | 1140 | 1150 | 1160 | 1170 | 1180 | 1190 | 1200 |
| SHGQHGSGSS | QSSGYGRQGS | GSGQSPGHGQ | RGSGSRQSPS | YGRHGSGSGR | SSSSGQHGSG | LGESSGFGHH | ESSSGQSSSY | SQHGSGSGHS | SGYGQHGSRS |
| 1210 | 1220 | 1230 | 1240 | 1250 | 1260 | 1270 | 1280 | 1290 | 1300 |
| GQSSRGERHG | SSSGSSSHYG | QHGSGSRQSS | GHGRQGSGSG | HSPSRGRHGS | GLGHSSSHGQ | HGSGSGRSSS | RGPYESRSGH | SSVFGQHESG | SGHSSAYSQH |
| [illegible] | [illegible] | [illegible] | [illegible] | [illegible] | [illegible] | [illegible] | [illegible] | [illegible] | [illegible] |

MS/MS Peptide Matches

| Cmpd. | m/z meas. | z | Δ m/z [ppm] | Δ m/z [Da] | Rt [min] | Score | No. of Cmpds. | Site [%] | P | Sequence | Modification | Range | Type |
|---|---|---|---|---|---|---|---|---|---|---|---|---|---|
| 47 | 974.558 | 2 | 106.28 | 0.1036 | 10.61 | 28.22 | 1 | 0.0 | 0 | R.QSLGHGQHGSGSGQSPSPSR.G | | 992-1011 | CID |
| 88 | 826.772 | 3 | 516.54 | 0.4268 | 11.83 | 65.66 | 1 | 0.0 | 0 | R.HGSGSGHSSSYGQHGSGSGWSSSSGR.H | | 1365-1390 | CID |
| 33 | 909.458 | 3 | 441.97 | 0.4018 | 9.43 | 34.0 | 1 | 0.0 | 1 | R.SGQSSRGEQHGSSSGSSSSYGQHGSGSR.Q | | 1904-1931 | CID |

#### Protein 12: Immunoglobulin lambda variable 3-21 OS=Homo sapiens GN=IGLV3-21 PE=1 SV=2

**Accession:** LV321_HUMAN
**Database:** SwissProt
**MW [kDa] / pI:** 12.4 / 5.0

**Score:** 92.0
**Seq. Coverage [%]:** 13.7
**No. of Peptides:** 1

| 10 | 20 | 30 | 40 | 50 | 60 | 70 | 80 | 90 | 100 |
|---|---|---|---|---|---|---|---|---|---|
| MAWTVLLLGL | LSHCTGSVTS | YVLTQPPSVS | VAPGQTARIT | CGGNNIGSKS | VHWYQQKPGQ | APVLVVYDDS | DRPSGIPERF | SGSNSGNTAT | LTISRVEAGD |
| 110 | 120 | | | | | | | | |
| EADYYCQVWD | SSSDHPT | | | | | | | | |

MS/MS Peptide Matches

| Cmpd. | m/z meas. | z | Δ m/z [ppm] | Δ m/z [Da] | Rt [min] | Score | No. of Cmpds. | Site [%] | P | Sequence | Modification | Range | Type |
|---|---|---|---|---|---|---|---|---|---|---|---|---|---|
| 405 | 807.008 | 2 | 133.96 | 0.1081 | 19.31 | 91.95 | 2 | 0.0 | 0 | R.FSGSNSGNTATLTISR.V | | 80-95 | CID |

#### Protein 13: Immunoglobulin kappa variable 3-15 OS=Homo sapiens GN=IGKV3-15 PE=1 SV=2

**Accession:** KV315_HUMAN
**Database:** SwissProt
**MW [kDa] / pI:** 12.5 / 5.1

**Score:** 86.0
**Seq. Coverage [%]:** 18.3
**No. of Peptides:** 1

| 10 | 20 | 30 | 40 | 50 | 60 | 70 | 80 | 90 | 100 |
|---|---|---|---|---|---|---|---|---|---|
| MEAPAQLLFL | LLLWLPDTTG | EIVMTQSPAT | LSVSPGERAT | LSCRASQSVS | SNLAWYQQKP | GQAPRLLIYG | ASTRATGIPA | RFSGSGSGTE | FTLTISSLQS |
| 110 | 120 | | | | | | | | |
| EDFAVYYCQQ | YNNWP | | | | | | | | |

MS/MS Peptide Matches

| Cmpd. | m/z meas. | z | Δ m/z [ppm] | Δ m/z [Da] | Rt [min] | Score | No. of Cmpds. | Site [%] | P | Sequence | Modification | Range | Type |
|---|---|---|---|---|---|---|---|---|---|---|---|---|---|
| 512 | 768.468 | 3 | 102.70 | 0.0789 | 21.38 | 86.01 | 3 | 0.0 | 0 | R.ASQSVSSNLAWYQQKPGQAPR.L | | 45-65 | CID |

#### Protein 14: Immunoglobulin heavy variable 4-4 OS=Homo sapiens GN=IGHV4-4 PE=3 SV=2

**Accession:** HV404_HUMAN
**Database:** SwissProt
**MW [kDa] / pI:** 12.8 / 9.6
**Modification(s):** Carbamidomethyl

**Score:** 80.6
**Seq. Coverage [%]:** 13.7
**No. of Peptides:** 1

| 10 | 20 | 30 | 40 | 50 | 60 | 70 | 80 | 90 | 100 |
|---|---|---|---|---|---|---|---|---|---|
| MKHLWFFLLL | VAAPRWVLSQ | VQLQESGPGL | VKPSGTLSLT | CAVSGGSISS | SNWWSWVRQP | PGKGLEWIGE | IYHSGSTNYN | PSLKSRVTIS | VDKSKNQFSL |
| 110 | 120 | | | | | | | | |
| KLSSVTAADT | AVYYCAR | | | | | | | | |

MS/MS Peptide Matches

| Cmpd. | m/z meas. | z | Δ m/z [ppm] | Δ m/z [Da] | Rt [min] | Score | No. of Cmpds. | Site [%] | P | Sequence | Modification | Range | Type |
|---|---|---|---|---|---|---|---|---|---|---|---|---|---|
| 561 | 874.533 | 2 | 129.66 | 0.1134 | 22.41 | 80.64 | 3 | 0.0 | 0 | K.LSSVTAADTAVYYCAR.- | Carbamidomethyl: 14 | 102-117 | CID |

#### Project Info

| | | | |
|---|---|---|---|
| **Name:** | Alagesan | **Date:** | Mar 5, 2019 |
| **Note:** | Nanobooster | | |

#### Sample Info & Protocols

| | |
|---|---|
| **Name:** | Nanobooster - IgG |
| **Date:** | Mar 5, 2019 |

#### Search Result Info

| | | | |
|---|---|---|---|
| **Search Result:** | IonTrap_Human_Swiss_CID_2019-03-05 17:55:08 | | |
| **Location:** | /Alagesan/Nanobooster - IgG/KA_IgG1 HD NB ACN CID_GA2_01_9078.mgf | | |
| **Search Method:** | IonTrap_Human_Swiss_CID | **Version:** | 1.2 |
| **Search Engine(s):** | Mascot, 2.6.0 | **Ident. Compound(s):** | 65/2214 |
| **Database(s):** | SwissProt, SwissProt_2017_04.fasta | | |
| **Note:** | | | |

#### Results

##### Protein 1: Immunoglobulin kappa constant OS=Homo sapiens GN=IGKC PE=1 SV=2

**Accession:** IGKC_HUMAN
**Database:** SwissProt
**MW [kDa] / pI:** 11.8 / 6.1
**Modification(s):** Carbamidomethyl

**Score:** 1157.0
**Seq. Coverage [%]:** 88.8
**No. of Peptides:** 11

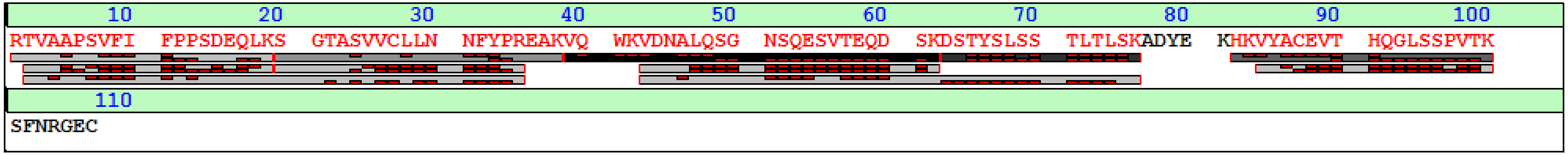


MS/MS Peptide Matches

| Cmpd. | m/z meas. | z | Δ m/z [ppm] | Δ m/z [Da] | Rt [min] | Score | No. of Cmpds. | Site [%] | P | Sequence | Modification | Range | Type |
|---|---|---|---|---|---|---|---|---|---|---|---|---|---|
| 901 | 701.449 | 3 | 97.13 | 0.0681 | 28.65 | 59.76 | 5 | 0.0 | 1 | -.RTVAAPSVFIFPPSDEQLK.S | | 1-19 | CID |
| 1012 | 973.604 | 2 | 89.25 | 0.0869 | 30.59 | 93.19 | 15 | 0.0 | 0 | R.TVAAPSVFIFPPSDEQLK.S | | 2-19 | CID |
| 1446 | 1242.769 | 3 | 372.45 | 0.4627 | 38.73 | 82.91 | 9 | 0.0 | 1 | R.TVAAPSVFIFPPSDEQLKSGTASVVCLLNNFYPR.E | Carbamidomethyl: 26 | 2-35 | CID |
| 1166 | 899.592 | 2 | 156.48 | 0.1407 | 33.20 | 80.76 | 15 | 0.0 | 0 | K.SGTASVVCLLNNFYPR.E | Carbamidomethyl: 8 | 20-35 | CID |
| 1055 | 709.462 | 3 | 141.70 | 0.1005 | 31.33 | 41.56 | 1 | 0.0 | 1 | K.SGTASVVCLLNNFYPREAK.V | Carbamidomethyl: 8 | 20-38 | CID |
| 485 | 893.495 | 3 | 448.05 | 0.4002 | 21.15 | 116.92 | 9 | 0.0 | 1 | K.VQWKVDNALQSGNSQESVTEQDSK.D | | 39-62 | CID |
| 256 | 1068.685 | 2 | 184.37 | 0.1970 | 15.91 | 169.94 | 29 | 0.0 | 0 | K.VDNALQSGNSQESVTEQDSK.D | | 43-62 | CID |
| 668 | 1207.567 | 3 | 269.79 | 0.3257 | 24.60 | 129.02 | 1 | 0.0 | 1 | K.VDNALQSGNSQESVTEQDSKDSTYSLSSTLTLSK.A | | 43-76 | CID |
| 684 | 751.978 | 2 | 126.54 | 0.0951 | 24.89 | 107.57 | 3 | 0.0 | 0 | K.DSTYSLSSTLTLSK.A | | 63-76 | CID |
| 299 | 714.461 | 3 | 134.22 | 0.0959 | 17.00 | 144.85 | 13 | 0.0 | 1 | K.HKVYACEVTHQGLSSPVTK.S | Carbamidomethyl: 6 | 82-100 | CID |
| 380 | 938.596 | 2 | 137.35 | 0.1289 | 19.04 | 130.5 | 10 | 0.0 | 0 | K.VYACEVTHQGLSSPVTK.S | Carbamidomethyl: 4 | 84-100 | CID |

| Cmpd. | m/z meas. | z | Δ m/z [ppm] | Δ m/z [Da] | Rt [min] | Score | No. of Cmpds. | Site [%] | P | Sequence | Modification | Range | Type |
|---|---|---|---|---|---|---|---|---|---|---|---|---|---|

**Protein 2: Immunoglobulin gamma-1 heavy chain OS=Homo sapiens PE=1 SV=1**

| | | | |
|---|---|---|---|
| **Accession:** | IGG1_HUMAN | **Score:** | 1125.2 |
| **Database:** | SwissProt | **Seq. Coverage [%]:** | 43.4 |
| **MW [kDa] / pI:** | 49.3 / 8.9 | **No. of Peptides:** | 14 |
| **Modification(s):** | Carbamidomethyl | | |

| 10 | 20 | 30 | 40 | 50 | 60 | 70 | 80 | 90 | 100 |
|---|---|---|---|---|---|---|---|---|---|
| QVQLVQSGGG | VVQPGRSLRL | SCAASGFTFS | RYTIHWVRQA | PGKGLEWVAV | MSYBGBBKHY | ADSVNGRFTI | SRNDSKNTLY | LNMNSLRPED | TAVYYCARIR |
| **110** | **120** | **130** | **140** | **150** | **160** | **170** | **180** | **190** | **200** |
| DTAMFFAHWG | QGTLVTVSSA | STKGPSVFPL | APSSKSTSGG | TAALGCLVKD | YFPEPVTVSW | NSGALTSGVH | TFPAVLQSSG | LYSLSSVVTV | PSSSLGTQTY |
| **210** | **220** | **230** | **240** | **250** | **260** | **270** | **280** | **290** | **300** |
| ICNVNHKPSN | TKVDKKVEPK | SCDKTHTCPP | CPAPELLGGP | SVFLFPPKPK | DTLMISRTPE | VTCVVVDVSH | EDPEVKFNWY | VDGVEVHNAK | TKPREEQYNS |
| **310** | **320** | **330** | **340** | **350** | **360** | **370** | **380** | **390** | **400** |
| TYRVVSVLTV | LHQDWLNGKE | YKCKVSNKAL | PAPIEKTISK | AKGQPREPQV | YTLPPSRDEL | TKNQVSLTCL | VKGFYPSDIA | VEWESNGQPE | NNYKTTPPVL |
| **410** | **420** | **430** | **440** | **450** | | | | | |
| DSDGSFFLYS | KLTVDKSRWQ | QGNVFSCSVM | HEALHNHYTQ | KSLSLSPGK | | | | | |

MS/MS Peptide Matches

| Cmpd. | m/z meas. | z | Δ m/z [ppm] | Δ m/z [Da] | Rt [min] | Score | No. of Cmpds. | Site [%] | P | Sequence | Modification | Range | Type |
|---|---|---|---|---|---|---|---|---|---|---|---|---|---|
| 874 | 830.551 | 3 | 133.11 | 0.1105 | 28.13 | 77.47 | 3 | 0.0 | 1 | K.GPSVFPLAPSSKSTSGGTAALGCLVK.D | Carbamidomethyl: 23 | 124-149 | CID |
| 1003 | 1112.608 | 3 | 349.85 | 0.3891 | 30.41 | 98.33 | 12 | 0.0 | 1 | K.SCDKTHTCPPCPAPELLGGPSVFLFPPKPK.D | Carbamidomethyl: 2, 8, 11 | 221-250 | CID |
| 1031 | 949.251 | 3 | 450.00 | 0.4270 | 30.93 | 115.61 | 9 | 0.0 | 0 | K.THTCPPCPAPELLGGPSVFLFPPKPK.D | Carbamidomethyl: 4, 7 | 225-250 | CID |

| Cmpd. | m/z meas. | z | Δ m/z [ppm] | Δ m/z [Da] | Rt [min] | Score | No. of Cmpds. | Site [%] | P | Sequence | Modification | Range | Type |
|---|---|---|---|---|---|---|---|---|---|---|---|---|---|
| 1064 | 916.339 | 4 | 398.55 | 0.3651 | 31.47 | 49.33 | 3 | 0.0 | 1 | K.THTCPPCPAPELLGGPSVFLFPPKPKDTLMIS R.T | Carbamidomethyl: 4, 7 | 225-257 | CID |
| 691 | 1070.082 | 2 | 60.42 | 0.0646 | 24.98 | 105.05 | 2 | 0.0 | 0 | R.TPEVTCVVVDVSHEDPEVK.F | Carbamidomethyl: 6 | 258-276 | CID |
| 966 | 950.548 | 4 | 357.45 | 0.3397 | 29.78 | 32.78 | 3 | 0.0 | 1 | R.TPEVTCVVVDVSHEDPEVKFNWYVDGVEVHNA K.T | Carbamidomethyl: 6 | 258-290 | CID |
| 677 | 839.518 | 2 | 135.06 | 0.1134 | 24.77 | 85.56 | 5 | 0.0 | 0 | K.FNWYVDGVEVHNAK.T | | 277-290 | CID |
| 1113 | 904.627 | 2 | 132.80 | 0.1201 | 32.31 | 116.28 | 3 | 0.0 | 0 | R.VVSVLTVLHQDWLNGK.E | | 304-319 | CID |
| 1023 | 743.492 | 3 | 113.92 | 0.0847 | 30.78 | 70.33 | 2 | 0.0 | 1 | R.VVSVLTVLHQDWLNGKEYK.C | | 304-322 | CID |
| 499 | 937.056 | 2 | 71.80 | 0.0673 | 21.40 | 55.11 | 8 | 0.0 | 1 | R.EPQVYTLPPSRDELTK.N | | 347-362 | CID |
| 894 | 1273.231 | 2 | 519.95 | 0.6617 | 28.49 | 119.24 | 12 | 0.0 | 0 | K.GFYPSDIAVEWESNGQPENNYK.T | | 373-394 | CID |
| 952 | 937.563 | 2 | 105.02 | 0.0985 | 29.53 | 92.91 | 7 | 0.0 | 0 | K.TTPPVLDSDGSFFLYSK.L | | 395-411 | CID |
| 592 | 762.194 | 4 | 444.28 | 0.3385 | 23.28 | 34.97 | 2 | 0.0 | 1 | K.SRWQQGNVFSCSVMHEALHNHYTQK.S | Carbamidomethyl: 11 | 417-441 | CID |
| 634 | 934.851 | 3 | 453.52 | 0.4238 | 24.03 | 72.27 | 3 | 0.0 | 0 | R.WQQGNVFSCSVMHEALHNHYTQK.S | Carbamidomethyl: 9 | 419-441 | CID |

#### Protein 3: Immunoglobulin lambda constant 3 OS=Homo sapiens GN=IGLC3 PE=1 SV=1

**Accession:** IGLC3_HUMAN
**Database:** SwissProt
**MW [kDa] / pI:** 11.3 / 6.9
**Modification(s):** Carbamidomethyl

**Score:** 678.3
**Seq. Coverage [%]:** 77.4
**No. of Peptides:** 8

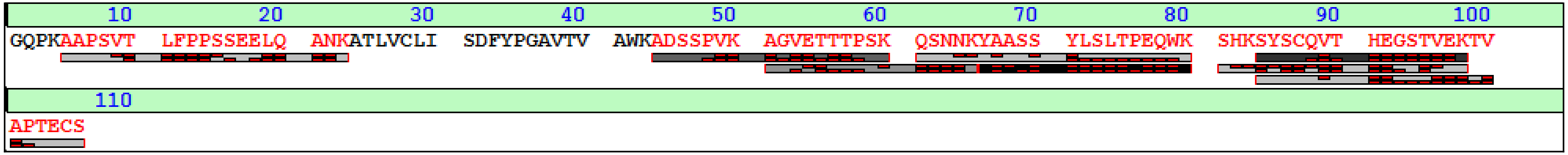

MS/MS Peptide Matches

| Cmpd. | m/z meas. | z | Δ m/z [ppm] | Δ m/z [Da] | Rt [min] | Score | No. of Cmpds. | Site [%] | P | Sequence | Modification | Range | Type |
|---|---|---|---|---|---|---|---|---|---|---|---|---|---|
| 789 | 993.575 | 2 | 62.85 | 0.0624 | 26.65 | 74.2 | 11 | 0.0 | 0 | K.AAPSVTLFPPSSEELQANK.A | | 5-23 | CID |
| 217 | 838.037 | 2 | 126.65 | 0.1061 | 15.01 | 58.87 | 3 | 0.0 | 1 | K.ADSSPVKAGVETTTPSK.Q | | 44-60 | CID |
| 41 | 781.521 | 2 | 161.94 | 0.1265 | 10.87 | 82.16 | 13 | 0.0 | 1 | K.AGVETTTPSKQSNNK.Y | | 51-65 | CID |
| 763 | 772.512 | 3 | 168.83 | 0.1304 | 26.20 | 83.07 | 6 | 0.0 | 1 | K.QSNNKYAASSYLSLTPEQWK.S | | 61-80 | CID |
| 859 | 872.604 | 2 | 195.94 | 0.1709 | 27.86 | 91.15 | 10 | 0.0 | 0 | K.YAASSYLSLTPEQWK.S | | 66-80 | CID |
| 100 | 1032.584 | 2 | 104.42 | 0.1078 | 12.34 | 90.02 | 3 | 0.0 | 1 | K.SHKSYSCQVTHEGSTVEK.T | Carbamidomethyl: 7 | 81-98 | CID |
| 181 | 856.559 | 2 | 205.24 | 0.1758 | 14.20 | 108.41 | 15 | 0.0 | 0 | K.SYSCQVTHEGSTVEK.T | Carbamidomethyl: 4 | 84-98 | CID |
| 377 | 1279.546 | 2 | 377.86 | 0.4833 | 18.98 | 90.45 | 10 | 0.0 | 1 | K.SYSCQVTHEGSTVEKTVAPTECS.- | Carbamidomethyl: 4, 22 | 84-106 | CID |

#### Protein 4: Keratin, type II cytoskeletal 1 OS=Homo sapiens GN=KRT1 PE=1 SV=6

| | | | |
|---|---|---|---|
| **Accession:** | K2C1_HUMAN | **Score:** | 670.8 |
| **Database:** | SwissProt | **Seq. Coverage [%]:** | 17.5 |
| **MW [kDa] / pI:** | 66.0 / 8.2 | **No. of Peptides:** | 6 |

| 10 | 20 | 30 | 40 | 50 | 60 | 70 | 80 | 90 | 100 |
|---|---|---|---|---|---|---|---|---|---|
| MSRQFSSRSG | YRSGGGFSSG | SAGIINYQRR | TTSSSTRRSG | GGGGRFSSCG | GGGGSFGAGG | GFGSRSLVNL | GGSKSISISV | ARGGGRGSGF | GGGYGGGGFG |
| 110 | 120 | 130 | 140 | 150 | 160 | 170 | 180 | 190 | 200 |
| GGGFGGGGFG | GGGIGGGGFG | GFGSGGGGFG | GGGFGGGGYG | GGYGPVCPPG | GIQEVTINQS | LLQPLNVEID | PEIQKVKSRE | REQIKSLNNQ | FASFIDKVRF |
| 210 | 220 | 230 | 240 | 250 | 260 | 270 | 280 | 290 | 300 |
| LEQQNQVLQT | KWELLQQVDT | STRTHNLEPY | FESFINNLRR | RVDQLKSDQS | RLDSELKNMQ | DMVEDYRNKY | EDEINKRTNA | ENEFVTIKKD | VDGAYMTKVD |
| 310 | 320 | 330 | 340 | 350 | 360 | 370 | 380 | 390 | 400 |
| LQAKLDNLQQ | EIDFLTALYQ | AELSQMQTQI | SETNVILSMD | NNRSLDLDSI | IAEVKAQYED | IAQKSKAEAE | SLYQSKYEEL | QITAGRHGDS | VRNSKIEISE |
| 410 | 420 | 430 | 440 | 450 | 460 | 470 | 480 | 490 | 500 |
| LNRVIQRLRS | EIDNVKKQIS | NLQQSISDAE | QRGENALKDA | KNKLNDLEDA | LQQAKEDLAR | LLRDYQELMN | TKLALDLEIA | TYRTLLEGEE | SRMSGECAPN |
| 510 | 520 | 530 | 540 | 550 | 560 | 570 | 580 | 590 | 600 |
| VSVSVSTSHT | TISGGGSRGG | GGGGYGSGGS | SYGSGGGSYG | SGGGGGGGRG | SYGSGGSSYG | SGGGSYGSGG | GGGGHGSYGS | GSSSGGYRGG | SGGGGGGSSG |
| 610 | 620 | 630 | 640 | 650 | | | | | |
| GRGSGGGSSG | GSIGGRGSSS | GGVKSSGGSS | SVKFVSTTYS | GVTR | | | | | |

**MS/MS Peptide Matches**

| Cmpd. | m/z meas. | z | Δ m/z [ppm] | Δ m/z [Da] | Rt [min] | Score | No. of Cmpds. | Site [%] | P | Sequence | Modification | Range | Type |
|---|---|---|---|---|---|---|---|---|---|---|---|---|---|
| 428 | 738.512 | 2 | 156.73 | 0.1157 | 20.05 | 98.3 | 5 | 0.0 | 0 | R.FLEQQNQVLQTK.W | | 200-211 | CID |
| 950 | 978.538 | 3 | 369.10 | 0.3610 | 29.49 | 85.46 | 1 | 0.0 | 1 | R.FLEQQNQVLQTKWELLQQVDTSTR.T | | 200-223 | CID |
| 1399 | 783.843 | 3 | 125.91 | 0.0987 | 37.74 | 163.41 | 3 | 0.0 | 1 | R.SLDLDSIIAEVKAQYEDIAQK.S | | 344-364 | CID |
| 514 | 859.031 | 2 | 118.52 | 0.1018 | 21.71 | 100.38 | 1 | 0.0 | 0 | K.QISNLQQSISDAEQR.G | | 418-432 | CID |
| 186 | 1192.54 | 2 | 50.68 | 0.0604 | 14.32 | 207.31 | 5 | 0.0 | 0 | R.GGGGGGYGSGGSSYGSGGGSYGSGGGGGG GR.G | | 519-549 | CID |
| 50 | 876.528 | 2 | 126.67 | 0.1110 | 11.15 | 15.94 | 2 | 0.0 | 1 | R.GSGGGSSGGSIGGRGSSSGGVK.S | | 603-624 | CID |

##### Protein 5: Immunoglobulin heavy constant gamma 4 OS=Homo sapiens GN=IGHG4 PE=1 SV=1

| | | | |
|---|---|---|---|
| **Accession:** | IGHG4_HUMAN | **Score:** | 594.6 |
| **Database:** | SwissProt | **Seq. Coverage [%]:** | 37.0 |
| **MW [kDa] / pI:** | 35.9 / 7.2 | **No. of Peptides:** | 7 |
| **Modification(s):** | Carbamidomethyl | | |

| 10 | 20 | 30 | 40 | 50 | 60 | 70 | 80 | 90 | 100 |
|---|---|---|---|---|---|---|---|---|---|
| ASTKGPSVFP | LAPCSRSTSE | STAALGCLVK | DYFPEPVTVS | WNSGALTSGV | HTFPAVLQSS | GLYSLSSVVT | VPSSSLGTKT | YTCNVDHKPS | NTKVDKRVES |
| **110** | **120** | **130** | **140** | **150** | **160** | **170** | **180** | **190** | **200** |
| KYGPPCPSCP | APEFLGGPSV | FLFPPKPKDT | LMISRTPEVT | CVVVDVSQED | PEVQFNWYVD | GVEVHNAKTK | PREEQFNSTY | RVVSVLTVLH | QDWLNGKEYK |
| **210** | **220** | **230** | **240** | **250** | **260** | **270** | **280** | **290** | **300** |
| CKVSNKGLPS | SIEKTISKAK | GQPREPQVYT | LPPSQEEMTK | NQVSLTCLVK | GFYPSDIAVE | WESNGQPENN | YKTTPPVLDS | DGSFFLYSRL | TVDKSRWQEG |
| **310** | **320** | **330** | | | | | | | |
| NVFSCSVMHE | ALHNHYTQKS | LSLSLGK | | | | | | | |

MS/MS Peptide Matches

| Cmpd. | m/z meas. | z | Δ m/z [ppm] | Δ m/z [Da] | Rt [min] | Score | No. of Cmpds. | Site [%] | P | Sequence | Modification | Range | Type |
|---|---|---|---|---|---|---|---|---|---|---|---|---|---|
| 557 | 712.451 | 2 | 129.85 | 0.0925 | 22.63 | 79.15 | 3 | 0.0 | 0 | R.STSESTAALGCLVK.D | Carbamidomethyl: 11 | 17-30 | CID |
| 58 | 833.035 | 2 | 175.91 | 0.1465 | 11.36 | 49.87 | 6 | 0.0 | 0 | K.TYTCNVDHKPSNTK.V | Carbamidomethyl: 4 | 80-93 | CID |
| 1159 | 982.251 | 3 | 434.99 | 0.4271 | 33.04 | 65.83 | 1 | 0.0 | 0 | K.YGPPCPSCPAPEFLGGPSVFLFPPKPK.D | Carbamidomethyl: 5, 8 | 102-128 | CID |
| 1113 | 904.627 | 2 | 132.80 | 0.1201 | 32.31 | 116.28 | 3 | 0.0 | 0 | R.VVSVLTVLHQDWLNGK.E | | 182-197 | CID |
| 1023 | 743.492 | 3 | 113.92 | 0.0847 | 30.78 | 70.33 | 2 | 0.0 | 1 | R.VVSVLTVLHQDWLNGKEYK.C | | 182-200 | CID |
| 894 | 1273.231 | 2 | 519.95 | 0.6617 | 28.49 | 119.24 | 12 | 0.0 | 0 | K.GFYPSDIAVEWESNGQPENNYK.T | | 251-272 | CID |
| 588 | 762.26 | 4 | 207.95 | 0.1585 | 23.19 | 47.65 | 2 | 0.0 | 1 | K.SRWQEGNVFSCSVMHEALHNHYTQK.S | Carbamidomethyl: 11 | 295-319 | CID |

BRUKER

#### Protein 6: Immunoglobulin lambda constant 1 OS=Homo sapiens GN=IGLC1 PE=1 SV=1

| | | | |
|---|---|---|---|
| **Accession:** | IGLC1_HUMAN | **Score:** | 437.4 |
| **Database:** | SwissProt | **Seq. Coverage [%]:** | 59.4 |
| **MW [kDa] / pI:** | 11.3 / 7.9 | **No. of Peptides:** | 6 |
| **Modification(s):** | Carbamidomethyl | | |

```
        10         20         30         40         50         60         70         80         90        100
GQPKANPTVT LFPPSSEELQ ANKATLVCLI SDFYPGAVTV AWKADGSPVK AGVETTKPSK QSNNKYAASS YLSLTPEQWK SHRSYSCQVT HEGSTVEKTV
       110
APTECS
```

MS/MS Peptide Matches

| Cmpd. | m/z meas. | z | Δ m/z [ppm] | Δ m/z [Da] | Rt [min] | Score | No. of Cmpds. | Site [%] | P | Sequence | Modification | Range | Type |
|---|---|---|---|---|---|---|---|---|---|---|---|---|---|
| 153 | 836.564 | 2 | 137.21 | 0.1148 | 13.53 | 40.83 | 2 | 0.0 | 1 | K.ADGSPVKAGVETTKPSK.Q | | 44-60 | CID |
| 763 | 772.512 | 3 | 168.83 | 0.1304 | 26.20 | 83.07 | 6 | 0.0 | 1 | K.QSNNKYAASSYLSLTPEQWK.S | | 61-80 | CID |
| 859 | 872.604 | 2 | 195.94 | 0.1709 | 27.86 | 91.15 | 10 | 0.0 | 0 | K.YAASSYLSLTPEQWK.S | | 66-80 | CID |
| 110 | 1046.536 | 2 | 54.22 | 0.0567 | 12.53 | 23.45 | 1 | 0.0 | 1 | K.SHRSYSCQVTHEGSTVEK.T | Carbamidomethyl: 7 | 81-98 | CID |
| 181 | 856.559 | 2 | 205.24 | 0.1758 | 14.20 | 108.41 | 15 | 0.0 | 0 | R.SYSCQVTHEGSTVEK.T | Carbamidomethyl: 4 | 84-98 | CID |
| 377 | 1279.546 | 2 | 377.86 | 0.4833 | 18.98 | 90.45 | 10 | 0.0 | 1 | R.SYSCQVTHEGSTVEKTVAPTECS.- | Carbamidomethyl: 4, 22 | 84-106 | CID |

#### Protein 7: Keratin, type II cytoskeletal 2 epidermal OS=Homo sapiens GN=KRT2 PE=1 SV=2

| | | | |
|---|---|---|---|
| **Accession:** | K22E_HUMAN | **Score:** | 408.1 |
| **Database:** | SwissProt | **Seq. Coverage [%]:** | 8.6 |
| **MW [kDa] / pI:** | 65.4 / 8.1 | **No. of Peptides:** | 3 |

| | | | | | | | | | |
|---|---|---|---|---|---|---|---|---|---|
| 10 | 20 | 30 | 40 | 50 | 60 | 70 | 80 | 90 | 100 |
| MSCQISCKSR | GRGGGGGGFR | GFSSGSAVVS | GGSRRSTSSF | SCLSRHGGGG | GGFGGGGFGS | RSLVGLGGTK | SISISVAGGG | GGFGAAGGFG | GRGGGFGGGS |
| 110 | 120 | 130 | 140 | 150 | 160 | 170 | 180 | 190 | 200 |
| SFGGGSGFSG | GGFGGGGFGG | GRFGGFGGPG | GVGGLGGPGG | FGPGGYPGGI | HEVSVNQSLL | QPLNVKVDPE | IQNVKAQERE | QIKTLNNKFA | SFIDKVRFLE |
| 210 | 220 | 230 | 240 | 250 | 260 | 270 | 280 | 290 | 300 |
| QQNQVLQTKW | ELLQQMNVGT | RPINLEPIFQ | GYIDSLKRYL | DGLTAERTSQ | NSELNNMQDL | VEDYKKKYED | EINKRTAAEN | DFVTLKKDVD | NAYMIKVELQ |
| 310 | 320 | 330 | 340 | 350 | 360 | 370 | 380 | 390 | 400 |
| SKVDLLNQEI | EFLKVLYDAE | ISQIHQSVTD | TNVILSMDNS | RNLDLDSIIA | EVKAQYEEIA | QRSKEEAEAL | YHSKYEELQV | TVGRHGDSLK | EIKIEISELN |
| 410 | 420 | 430 | 440 | 450 | 460 | 470 | 480 | 490 | 500 |
| RVIQRLQGEI | AHVKKQCKNV | QDAIADAEQR | GEHALKDARN | KLNDLEEALQ | QAKEDLARLL | RDYQELMNVK | LALDVEIATY | RKLLEGEECR | MSGDLSSNVT |
| 510 | 520 | 530 | 540 | 550 | 560 | 570 | 580 | 590 | 600 |
| VSVTSSTISS | NVASKAAFGG | SGGRGSSSGG | GYSSGSSSYG | SGGRQSGSRG | GSGGGGSISG | GGYGSGGGSG | GRYGSGGGSK | GGSISGGGYG | SGGGKHSSGG |
| 610 | 620 | 630 | 640 | | | | | | |
| GSRGGSSSGG | GYGSGGGGSS | SVKGSSGEAF | GSSVTFSFR | | | | | | |

MS/MS Peptide Matches

| Cmpd. | m/z meas. | z | Δ m/z [ppm] | Δ m/z [Da] | Rt [min] | Score | No. of Cmpds. | Site [%] | P | Sequence | Modification | Range | Type |
|---|---|---|---|---|---|---|---|---|---|---|---|---|---|
| 428 | 738.512 | 2 | 156.73 | 0.1157 | 20.05 | 98.3 | 5 | 0.0 | 0 | R.FLEQQNQVLQTK.W | | 198-209 | CID |
| 104 | 870.973 | 2 | 133.86 | 0.1166 | 12.40 | 148.49 | 3 | 0.0 | 0 | R.GSSSGGGYSSGSSSYGSGGR.Q | | 525-544 | CID |
| 139 | 871.478 | 2 | 114.92 | 0.1001 | 13.17 | 161.35 | 3 | 0.0 | 0 | R.GGSGGGGSISGGGYGSGGGSGGR.Y | | 550-572 | CID |

##### Protein 8: Keratin, type I cytoskeletal 9 OS=Homo sapiens GN=KRT9 PE=1 SV=3

| | | | |
|---|---|---|---|
| **Accession:** | K1C9_HUMAN | **Score:** | 367.0 |
| **Database:** | SwissProt | **Seq. Coverage [%]:** | 24.1 |
| **MW [kDa] / pI:** | 62.0 / 5.1 | **No. of Peptides:** | 6 |

| 10 | 20 | 30 | 40 | 50 | 60 | 70 | 80 | 90 | 100 |
|---|---|---|---|---|---|---|---|---|---|
| MSCRQFSSSY | LSRSGGGGGG | GLGSGGSIRS | SYSRFSSSGG | GGGGGRFSSS | SGYGGGSSRV | CGRGGGGSFG | YSYGGGSGGG | FSASSLGGGF | GGGSRGFGGA |
| **110** | **120** | **130** | **140** | **150** | **160** | **170** | **180** | **190** | **200** |
| SGGGYSSSGG | FGGGFGGGSG | GGFGGGYGSG | FGGFGGFGGG | AGGGDGGILT | ANEKSTMQEL | NSRLASYLDK | VQALEEANND | LENKIQDWYD | KKGPAAIQKN |
| **210** | **220** | **230** | **240** | **250** | **260** | **270** | **280** | **290** | **300** |
| YSPYYNTIDD | LKDQIVDLTV | GNNKTLLDID | NTRMTLDDFR | IKFEMEQNLR | QGVDADINGL | RQVLDNLTME | KSDLEMQYET | LQEELMALKK | NHKEEMSQLT |
| **310** | **320** | **330** | **340** | **350** | **360** | **370** | **380** | **390** | **400** |
| GQNSGDVNVE | INVAPGKDLT | KTLNDMRQEY | EQLIAKNRKD | IENQYETQIT | QIEHEVSSSG | QEVQSSAKEV | TQLRHGVQEL | EIELQSQLSK | KAALEKSLED |
| **410** | **420** | **430** | **440** | **450** | **460** | **470** | **480** | **490** | **500** |
| TKNRYCGQLQ | MIQEQISNLE | AQITDVRQEI | ECQNQEYSLL | LSIKMRLEKE | IETYHNLLEG | GQEDFESSGA | GKIGLGGRGG | SGGSYGRGSR | GGSGGSYGGG |
| **510** | **520** | **530** | **540** | **550** | **560** | **570** | **580** | **590** | **600** |
| GSGGGYGGGS | GSRGGSGGSY | GGGSGSGGGS | GGGYGGGSGG | GHSGGSGGGH | SGGSGGNYGG | GSGSGGGSGG | GYGGGSGSRG | GSGGSHGGGS | GFGGESGGSY |
| **610** | **620** | **630** | | | | | | | |
| GGGEEASGSG | GGYGGGSGKS | SHS | | | | | | | |

**MS/MS Peptide Matches**

| Cmpd. | m/z meas. | z | Δ m/z [ppm] | Δ m/z [Da] | Rt [min] | Score | No. of Cmpds. | Site [%] | P | Sequence | Modification | Range | Type |
|---|---|---|---|---|---|---|---|---|---|---|---|---|---|
| 1204 | 968.214 | 3 | 74.69 | 0.0723 | 33.89 | 60.25 | 5 | 0.0 | 1 | K.NYSPYYNTIDDLKDQIVDLTVGNNK.T | | 200-224 | CID |
| 977 | 1089.191 | 3 | 319.78 | 0.3482 | 29.96 | 87.55 | 2 | 0.0 | 0 | K.DIENQYETQITQIEHEVSSSGQEVQSSAK.E | | 340-368 | CID |
| 1098 | 998.819 | 4 | 334.98 | 0.3345 | 32.05 | 15.35 | 1 | 0.0 | 1 | K.DIENQYETQITQIEHEVSSSGQEVQSSAKEVTQL R.H | | 340-374 | CID |
| 661 | 837.464 | 3 | 97.80 | 0.0819 | 24.47 | 18.42 | 1 | 0.0 | 0 | K.EIETYHNLLEGGQEDFESSGAGK.I | | 450-472 | CID |
| 91 | 896.465 | 2 | 108.78 | 0.0975 | 12.15 | 127.1 | 3 | 0.0 | 0 | R.GGSGGSYGGGGSGGGYGGGSGSR.G | | 491-513 | CID |
| 216 | 906.464 | 4 | 385.60 | 0.3494 | 14.98 | 58.3 | 1 | 0.0 | 1 | R.GGSGGSHGGGSGFGGESGGSYGGGEEASGS GGGYGGGSGKSSHS.- | | 580-623 | CID |

#### Protein 9: Immunoglobulin lambda constant 7 OS=Homo sapiens GN=IGLC7 PE=1 SV=3

| | | | |
|---|---|---|---|
| **Accession:** | IGLC7_HUMAN | **Score:** | 294.9 |
| **Database:** | SwissProt | **Seq. Coverage [%]:** | 53.8 |
| **MW [kDa] / pI:** | 11.2 / 8.5 | **No. of Peptides:** | 4 |
| **Modification(s):** | Carbamidomethyl | | |

```
10         20         30         40         50         60         70         80         90         100
GQPKAAPSVT LFPPSSEELQ ANKATLVCLV SDFNPGAVTV AWKADGSPVK VGVETTKPSK QSNNKYAASS YLSLTPEQWK SHRSYSCRVT HEGSTVEKTV
110
APAECS
```

**MS/MS Peptide Matches**

| Cmpd. | m/z meas. | z | Δ m/z [ppm] | Δ m/z [Da] | Rt [min] | Score | No. of Cmpds. | Site [%] | P | Sequence | Modification | Range | Type |
|---|---|---|---|---|---|---|---|---|---|---|---|---|---|
| 789 | 993.575 | 2 | 62.85 | 0.0624 | 26.65 | 74.2 | 11 | 0.0 | 0 | K.AAPSVTLFPPSSEELQANK.A | | 5-23 | CID |
| 763 | 772.512 | 3 | 168.83 | 0.1304 | 26.20 | 83.07 | 6 | 0.0 | 1 | K.QSNNKYAASSYLSLTPEQWK.S | | 61-80 | CID |
| 859 | 872.604 | 2 | 195.94 | 0.1709 | 27.86 | 91.15 | 10 | 0.0 | 0 | K.YAASSYLSLTPEQWK.S | | 66-80 | CID |
| 230 | 951.536 | 2 | 91.33 | 0.0869 | 15.34 | 46.44 | 1 | 0.0 | 1 | R.VTHEGSTVEKTVAPAECS.- | Carbamidomethyl: 17 | 89-106 | CID |

#### Protein 10: Immunoglobulin heavy constant gamma 2 OS=Homo sapiens GN=IGHG2 PE=1 SV=2

| | | | |
|---|---|---|---|
| **Accession:** | IGHG2_HUMAN | **Score:** | 285.5 |
| **Database:** | SwissProt | **Seq. Coverage [%]:** | 35.3 |
| **MW [kDa] / pI:** | 35.9 / 7.7 | **No. of Peptides:** | 6 |
| **Modification(s):** | Carbamidomethyl | | |

```
        10         20         30         40         50         60         70         80         90        100
ASTKGPSVFP LAPCSRSTSE STAALGCLVK DYFPEPVTVS WNSGALTSGV HTFPAVLQSS GLYSLSSVVT VPSSNFGTQT YTCNVDHKPS NTKVDKTVER
       110        120        130        140        150        160        170        180        190        200
KCCVECPPCP APPVAGPSVF LFPPKPKDTL MISRTPEVTC VVVDVSHEDP EVQFNWYVDG VEVHNAKTKP REEQFNSTFR VVSVLTVVHQ DWLNGKEYKC
       210        220        230        240        250        260        270        280        290        300
KVSNKGLPAP IEKTISKTKG QPREPQVYTL PPSREEMTKN QVSLTCLVKG FYPSDISVEW ESNGQPENNY KTTPPMLDSD GSFFLYSKLT VDKSRWQQGN
       310        320        330
VFSCSVMHEA LHNHYTQKSL SLSPGK
```

MS/MS Peptide Matches

| Cmpd. | m/z meas. | z | Δ m/z [ppm] | Δ m/z [Da] | Rt [min] | Score | No. of Cmpds. | Site [%] | P | Sequence | Modification | Range | Type |
|---|---|---|---|---|---|---|---|---|---|---|---|---|---|
| 557 | 712.451 | 2 | 129.85 | 0.0925 | 22.63 | 79.15 | 3 | 0.0 | 0 | R.STSESTAALGCLVK.D | Carbamidomethyl: 11 | 17-30 | CID |
| 910 | 759.993 | 4 | 149.21 | 0.1134 | 28.83 | 20.23 | 2 | 0.0 | 1 | R.KCCVECPPCPAPPVAGPSVFLFPPKPK.D | Carbamidomethyl: 2, 3, 6, 9 | 101-127 | CID |
| 966 | 950.548 | 4 | 367.03 | 0.3487 | 29.78 | 32.78 | 3 | 0.0 | 0 | R.TPEVTCVVVDVSHEDPEVQFNWYVDGVEVHNAK.T | Carbamidomethyl: 6 | 135-167 | CID |
| 467 | 953.053 | 2 | 82.10 | 0.0782 | 20.79 | 46.06 | 12 | 0.0 | 1 | R.EPQVYTLPPSREEMTK.N | | 224-239 | CID |
| 592 | 762.194 | 4 | 444.28 | 0.3385 | 23.28 | 34.97 | 2 | 0.0 | 1 | K.SRWQQGNVFSCSVMHEALHNHYTQK.S | Carbamidomethyl: 11 | 294-318 | CID |
| 634 | 934.851 | 3 | 453.52 | 0.4238 | 24.03 | 72.27 | 3 | 0.0 | 0 | R.WQQGNVFSCSVMHEALHNHYTQK.S | Carbamidomethyl: 9 | 296-318 | CID |

#### Protein 11: Keratin, type I cytoskeletal 10 OS=Homo sapiens GN=KRT10 PE=1 SV=6

| | | | |
|---|---|---|---|
| **Accession:** | K1C10_HUMAN | **Score:** | 252.3 |
| **Database:** | SwissProt | **Seq. Coverage [%]:** | 9.9 |
| **MW [kDa] / pI:** | 58.8 / 5.1 | **No. of Peptides:** | 3 |

| 10 | 20 | 30 | 40 | 50 | 60 | 70 | 80 | 90 | 100 |
|---|---|---|---|---|---|---|---|---|---|
| MSVRYSSSKH | YSSSRSGGGG | GGGGCGGGGG | VSSLRISSSK | GSLGGGFSSG | GFSGGSFSRG | SSGGGCFGGS | SGGYGGLGGF | GGGSFRGSYG | SSSFGGSYGG |
| 110 | 120 | 130 | 140 | 150 | 160 | 170 | 180 | 190 | 200 |
| IFGGGSFGGG | SFGGGSFGGG | GFGGGGFGGG | FGGGFGGDGG | LLSGNEKVTM | QNLNDRLASY | LDKVRALEES | NYELEGKIKE | WYEKHGNSHQ | GEPRDYSKYY |
| 210 | 220 | 230 | 240 | 250 | 260 | 270 | 280 | 290 | 300 |
| KTIDDLKNQI | LNLTTDNANI | LLQIDNARLA | ADDFRLKYEN | EVALRQSVEA | DINGLRRVLD | ELTLTKADLE | MQIESLTEEL | AYLKKNHEEE | MKDLRNVSTG |
| 310 | 320 | 330 | 340 | 350 | 360 | 370 | 380 | 390 | 400 |
| DVNVEMNAAP | GVDLTQLLNN | MRSQYEQLAE | QNRKDAEAWF | NEKSKELTTE | IDNNIEQISS | YKSEITELRR | NVQALEIELQ | SQLALKQSLE | ASLAETEGRY |
| 410 | 420 | 430 | 440 | 450 | 460 | 470 | 480 | 490 | 500 |
| CVQLSQIQAQ | ISALEEQLQQ | IRAETECQNT | EYQQLLDIKI | RLENEIQTYR | SLLEGEGSSG | GGGRGGGSFG | GGYGGGSSGG | GSSGGGHGGG | HGGSSGGGYG |
| 510 | 520 | 530 | 540 | 550 | 560 | 570 | 580 | 590 | |
| GGSSGGGSSG | GGYGGGSSSG | GHGGSSSGGY | GGGSSGGGGG | GYGGGSSGGG | SSSGGGYGGG | SSSGGHKSSS | SGSVGESSSK | GPRY | |

MS/MS Peptide Matches

| Cmpd. | m/z meas. | z | Δ m/z [ppm] | Δ m/z [Da] | Rt [min] | Score | No. of Cmpds. | Site [%] | P | Sequence | Modification | Range | Type |
|---|---|---|---|---|---|---|---|---|---|---|---|---|---|
| 620 | 854.495 | 2 | 123.24 | 0.1053 | 23.79 | 147.3 | 3 | 0.0 | 0 | K.GSLGGGFSSGGFSGGSFSR.G | | 41-59 | CID |
| 1255 | 1436.615 | 2 | -59.18 | -0.0850 | 34.90 | 66.05 | 2 | 0.0 | 0 | R.NVSTGDVNVEMNAAPGVDLTQLLNNMR.S | | 296-322 | CID |
| 212 | 747.412 | 2 | 55.15 | 0.0412 | 14.90 | 38.92 | 3 | 0.0 | 1 | R.SQYEQLAEQNRK.D | | 323-334 | CID |

#### Protein 12: Hornerin OS=Homo sapiens GN=HRNR PE=1 SV=2

**Accession:** HORN_HUMAN
**Database:** SwissProt
**MW [kDa] / pI:** 282.2 / 10.0

**Score:** 119.7
**Seq. Coverage [%]:** 1.6
**No. of Peptides:** 2

| | | | | | | | | | |
|---|---|---|---|---|---|---|---|---|---|
| 10 | 20 | 30 | 40 | 50 | 60 | 70 | 80 | 90 | 100 |
| MPKLLQGVIT | VIDVFYQYAT | QHGEYDTLNK | AELKELLENE | FHQILKNPND | PDTVDIILQS | LDRDHNKKVD | FTEYLLMIFK | LVQARNKIIG | KDYCQVSGSK |
| 110 | 120 | 130 | 140 | 150 | 160 | 170 | 180 | 190 | 200 |
| LRDDTHQHQE | EQEETEKEEN | KRQESSFSHS | SWSAGENDSY | SRNVRGSLKP | GTESISRRLS | FQRDFSGQHN | SYSGQSSSYG | EQNSDSHQSS | GRGQCGSGSG |
| 210 | 220 | 230 | 240 | 250 | 260 | 270 | 280 | 290 | 300 |
| QSPNYGQHGS | GSGQSSSNDT | HGSGSGQSSG | FSQHKSSSGQ | SSGYSQHGSG | SGHSSGYGQH | GSRSGQSSRG | ERHRSSSGSS | SSYGQHGSGS | RQSLGHGRQG |
| 310 | 320 | 330 | 340 | 350 | 360 | 370 | 380 | 390 | 400 |
| SGSRQSPSHV | RHGSGSGHSS | SHGQHGSGSS | YSYSRGHYES | GSGQTSGFGQ | HESGSGQSSG | YSKHGSGSGH | SSSQGQHGST | SGQASSSGQH | GSSSRQSSSY |
| 410 | 420 | 430 | 440 | 450 | 460 | 470 | 480 | 490 | 500 |
| GQHESASRHS | SGRGQHSSGS | GQSPGHGQRG | SGSGQSPSSG | QHGTGFGRSS | SSGPYVSGSG | YSSGFGHHES | SSEHSSGYTQ | HGSGSGHSSG | HGQHGSRSGQ |
| 510 | 520 | 530 | 540 | 550 | 560 | 570 | 580 | 590 | 600 |
| SSRGERQGSS | AGSSSSYGQH | GSGSRQSLGH | SRHGSGSGQS | PSPSRGRHES | GSRQSSSYGP | HGYGSGRSSS | RGPYESGSGH | SSGLGHQESR | SGQSSGYGQH |
| 610 | 620 | 630 | 640 | 650 | 660 | 670 | 680 | 690 | 700 |
| GSSSGHSSTH | GQHGSTSGQS | SSCGQHGATS | GQSSSHGQHG | SGSSQSSRYG | QQGSGSGQSP | SRGRHGSDFG | HSSSYGQHGS | GSGWSSSNGP | HGSVSGQSSG |
| 710 | 720 | 730 | 740 | 750 | 760 | 770 | 780 | 790 | 800 |
| FGHKSGSGQS | SGYSQHGSGS | SHSSGYRKHG | SRSGQSSRSE | QHGSSSGLSS | SYGQHGSGSH | QSSGHGRQGS | GSGHSPSRVR | HGSSSGHSSS | HGQHGSGTSC |
| 810 | 820 | 830 | 840 | 850 | 860 | 870 | 880 | 890 | 900 |
| SSSCGHYESG | SGQASGFGQH | ESGSGQGYSQ | HGSASGHFSS | QGRHGSTSGQ | SSSSGQHDSS | SGQSSSYGQH | ESASHHASGR | GRHGSGSGQS | PGHGQRGSGS |
| 910 | 920 | 930 | 940 | 950 | 960 | 970 | 980 | 990 | 1000 |
| GQSPSYGRHG | SGSGRSSSSG | RHGSGSGQSS | GFGHKSSSGQ | SSGYTQHGSG | SGHSSSYEQH | GSRSGQSSRS | EQHGSSSGSS | SSYGQHGSGS | RQSLGHGQHG |
| 1010 | 1020 | 1030 | 1040 | 1050 | 1060 | 1070 | 1080 | 1090 | 1100 |
| SGSGQSPSPS | RGRHGSGSGQ | SSSYGPYRSG | SGWSSSRGPY | ESGSGHSSGL | GHRESRSGQS | SGYGQHGSSS | GHSSTHGQHG | STSGQSSSCG | QHGASSGQSS |
| 1110 | 1120 | 1130 | 1140 | 1150 | 1160 | 1170 | 1180 | 1190 | 1200 |
| SHGQHGSGSS | QSSGYGRQGS | GSGQSPGHGQ | RGSGSRQSPS | YGRHGSGSGR | SSSSGQHGSG | LGESSGFGHH | ESSSGQSSSY | SQHGSGSGHS | SGYGQHGSRS |
| 1210 | 1220 | 1230 | 1240 | 1250 | 1260 | 1270 | 1280 | 1290 | 1300 |
| GQSSRGERHG | SSSGSSSHYG | QHGSGSRQSS | GHGRQGSGSG | HSPSRGRHGS | GLGHSSSHGQ | HGSGSGRSSS | RGPYESRSGH | SSVFGQHESG | SGHSSAYSQH |
| [illegible] | [illegible] | [illegible] | [illegible] | [illegible] | [illegible] | [illegible] | [illegible] | [illegible] | [illegible] |

MS/MS Peptide Matches

| Cmpd. | m/z meas. | z | Δ m/z [ppm] | Δ m/z [Da] | Rt [min] | Score | No. of Cmpds. | Site [%] | P | Sequence | Modification | Range | Type |
|---|---|---|---|---|---|---|---|---|---|---|---|---|---|
| 32 | 974.536 | 2 | 83.71 | 0.0816 | 10.64 | 31.12 | 2 | 0.0 | 0 | R.QSLGHGQHGSGSGQSPSPSR.G | | 992-1011 | CID |
| 71 | 826.434 | 3 | 107.51 | 0.0888 | 11.80 | 88.53 | 2 | 0.0 | 0 | R.HGSGSGHSSSYGQHGSGSGWSSSSGR.H | | 1365-1390 | CID |

#### Protein 13: Immunoglobulin heavy variable 4-4 OS=Homo sapiens GN=IGHV4-4 PE=3 SV=2

**Accession:** HV404_HUMAN
**Database:** SwissProt
**MW [kDa] / pI:** 12.8 / 9.6
**Modification(s):** Carbamidomethyl

**Score:** 92.8
**Seq. Coverage [%]:** 13.7
**No. of Peptides:** 1

```
        10         20         30         40         50         60         70         80         90        100
MKHLWFFLLL VAAPRWVLSQ VQLQESGPGL VKPSGTLSLT CAVSGGSISS SNWWSWVRQP PGKGLEWIGE IYHSGSTNYN PSLKSRVTIS VDKSKNQFSL
       110        120
KLSSVTAADT AVYYCAR
```

MS/MS Peptide Matches

| Cmpd. | m/z meas. | z | Δ m/z [ppm] | Δ m/z [Da] | Rt [min] | Score | No. of Cmpds. | Site [%] | P | Sequence | Modification | Range | Type |
|---|---|---|---|---|---|---|---|---|---|---|---|---|---|
| 542 | 874.51 | 2 | 103.36 | 0.0904 | 22.37 | 92.77 | 3 | 0.0 | 0 | K.LSSVTAADTAVYYCAR.- | Carbamidomethyl: 14 | 102-117 | CID |

#### Protein 14: Immunoglobulin lambda variable 3-21 OS=Homo sapiens GN=IGLV3-21 PE=1 SV=2

**Accession:** LV321_HUMAN
**Database:** SwissProt
**MW [kDa] / pI:** 12.4 / 5.0

**Score:** 91.0
**Seq. Coverage [%]:** 13.7
**No. of Peptides:** 1

| 10 | 20 | 30 | 40 | 50 | 60 | 70 | 80 | 90 | 100 |
|---|---|---|---|---|---|---|---|---|---|
| MAWTVLLLGL | LSHCTGSVTS | YVLTQPPSVS | VAPGQTARIT | CGGNNIGSKS | VHWYQQKPGQ | APVLVVYDDS | DRPSGIPERF | SGSNSGNTAT | LTISRVEAGD |
| 110 | 120 | | | | | | | | |
| EADYYCQVWD | SSSDHPT | | | | | | | | |

MS/MS Peptide Matches

| Cmpd. | m/z meas. | z | Δ m/z [ppm] | Δ m/z [Da] | Rt [min] | Score | No. of Cmpds. | Site [%] | P | Sequence | Modification | Range | Type |
|---|---|---|---|---|---|---|---|---|---|---|---|---|---|
| 392 | 806.983 | 2 | 102.98 | 0.0831 | 19.32 | 90.97 | 3 | 0.0 | 0 | R.FSGSNSGNTATLTISR.V | | 80-95 | CID |

#### Protein 15: Immunoglobulin kappa variable 3-15 OS=Homo sapiens GN=IGKV3-15 PE=1 SV=2

**Accession:** KV315_HUMAN
**Database:** SwissProt
**MW [kDa] / pI:** 12.5 / 5.1

**Score:** 76.1
**Seq. Coverage [%]:** 18.3
**No. of Peptides:** 1

| 10 | 20 | 30 | 40 | 50 | 60 | 70 | 80 | 90 | 100 |
|---|---|---|---|---|---|---|---|---|---|
| MEAPAQLLFL | LLLWLPDTTG | EIVMTQSPAT | LSVSPGERAT | LSCRASQSVS | SNLAWYQQKP | GQAPRLLIYG | ASTRATGIPA | RFSGSGSGTE | FTLTISSLQS |
| 110 | 120 | | | | | | | | |
| EDFAVYYCQQ | YNNWP | | | | | | | | |

MS/MS Peptide Matches

| Cmpd. | m/z meas. | z | Δ m/z [ppm] | Δ m/z [Da] | Rt [min] | Score | No. of Cmpds. | Site [%] | P | Sequence | Modification | Range | Type |
|---|---|---|---|---|---|---|---|---|---|---|---|---|---|
| 500 | 768.484 | 3 | 123.52 | 0.0949 | 21.44 | 76.13 | 3 | 0.0 | 0 | R.ASQSVSSNLAWYQQKPGQAPR.L | | 45-65 | CID |

#### Protein 16: Immunoglobulin kappa variable 2D-28 OS=Homo sapiens GN=IGKV2D-28 PE=1 SV=2

**Accession:** KVD28_HUMAN
**Database:** SwissProt
**MW [kDa] / pI:** 12.9 / 5.6

**Score:** 40.2
**Seq. Coverage [%]:** 16.7
**No. of Peptides:** 1

| 10 | 20 | 30 | 40 | 50 | 60 | 70 | 80 | 90 | 100 |
|---|---|---|---|---|---|---|---|---|---|
| MRLPAQLLGL | LMLWVSGSSG | DIVMTQSPLS | LPVTPGEPAS | ISCRSSQSLL | HSNGYNYLDW | YLQKPGQSPQ | LLIYLGSNRA | SGVPDRFSGS | GSGTDFTLKI |
| **110** | **120** | **130** | | | | | | | |
| SRVEAEDVGV | YYCMQALQTP | | | | | | | | |

MS/MS Peptide Matches

| Cmpd. | m/z meas. | z | Δ m/z [ppm] | Δ m/z [Da] | Rt [min] | Score | No. of Cmpds. | Site [%] | P | Sequence | Modification | Range | Type |
|---|---|---|---|---|---|---|---|---|---|---|---|---|---|
| 548 | 993.546 | 2 | 64.63 | 0.0642 | 22.48 | 40.16 | 3 | 0.0 | 1 | R.ASGVPDRFSGSGSGTDFTLK.I | | 80-99 | CID |

#### Protein Report
#### IonTrap_Human_Swiss_CID_2019-03-05 17:55:08

##### Project Info

| | | | |
|---|---|---|---|
| **Name:** | Alagesan | **Date:** | Mar 5, 2019 |
| **Note:** | Nanobooster | | |

##### Sample Info & Protocols

| | |
|---|---|
| **Name:** | Nanobooster - IgG |
| **Date:** | Mar 5, 2019 |

##### Search Result Info

| | | | |
|---|---|---|---|
| **Search Result:** | IonTrap_Human_Swiss_CID_2019-03-05 17:55:08 | | |
| **Location:** | /Alagesan/Nanobooster - IgG/KA_IgG1 HD NB ACN CID_GA2_01_9079.mgf | | |
| **Search Method:** | IonTrap_Human_Swiss_CID | **Version:** | 1.2 |
| **Search Engine(s):** | Mascot, 2.6.0 | **Ident. Compound(s):** | 0/2209 |
| **Database(s):** | SwissProt, SwissProt_2017_04.fasta | | |
| **Note:** | | | |

#### Results

##### Protein 1: Immunoglobulin kappa constant OS=Homo sapiens GN=IGKC PE=1 SV=2

| | | | |
|---|---|---|---|
| **Accession:** | IGKC_HUMAN | **Score:** | 1109.5 |
| **Database:** | SwissProt | **Seq. Coverage [%]:** | 88.8 |
| **MW [kDa] / pI:** | 11.8 / 6.1 | **No. of Peptides:** | 11 |
| **Modification(s):** | Carbamidomethyl | | |

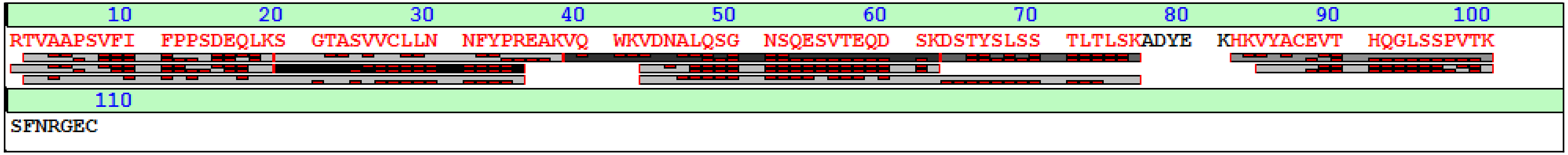


MS/MS Peptide Matches

| Cmpd. | m/z meas. | z | Δ m/z [ppm] | Δ m/z [Da] | Rt [min] | Score | No. of Cmpds. | Site [%] | P | Sequence | Modification | Range | Type |
|---|---|---|---|---|---|---|---|---|---|---|---|---|---|
| 911 | 1051.644 | 2 | 72.59 | 0.0763 | 28.84 | 48.44 | 4 | 0.0 | 1 | -.RTVAAPSVFIFPPSDEQLK.S | | 1-19 | CID |
| 1019 | 973.668 | 2 | 154.99 | 0.1509 | 30.73 | 92.52 | 14 | 0.0 | 0 | R.TVAAPSVFIFPPSDEQLK.S | | 2-19 | CID |
| 1415 | 1242.652 | 3 | 278.27 | 0.3457 | 38.12 | 94.15 | 14 | 0.0 | 1 | R.TVAAPSVFIFPPSDEQLKSGTASVVCLLNNFYPR.E | Carbamidomethyl: 26 | 2-35 | CID |
| 1152 | 899.561 | 2 | 122.02 | 0.1097 | 33.00 | 66.59 | 16 | 0.0 | 0 | K.SGTASVVCLLNNFYPR.E | Carbamidomethyl: 8 | 20-35 | CID |
| 1049 | 709.426 | 3 | 90.95 | 0.0645 | 31.28 | 48.47 | 1 | 0.0 | 1 | K.SGTASVVCLLNNFYPREAK.V | Carbamidomethyl: 8 | 20-38 | CID |
| 477 | 893.502 | 3 | 455.89 | 0.4072 | 21.02 | 121.75 | 10 | 0.0 | 1 | K.VQWKVDNALQSGNSQESVTEQDSK.D | | 39-62 | CID |
| 252 | 1068.673 | 2 | 173.14 | 0.1850 | 15.89 | 165.46 | 33 | 0.0 | 0 | K.VDNALQSGNSQESVTEQDSK.D | | 43-62 | CID |
| 666 | 1207.57 | 3 | 272.28 | 0.3287 | 24.55 | 117.41 | 3 | 0.0 | 1 | K.VDNALQSGNSQESVTEQDSKDSTYSLSSTLTLSK.A | | 43-76 | CID |
| 683 | 751.97 | 2 | 115.90 | 0.0871 | 24.87 | 102.79 | 3 | 0.0 | 0 | K.DSTYSLSSTLTLSK.A | | 63-76 | CID |
| 292 | 714.408 | 3 | 60.03 | 0.0429 | 16.90 | 139.0 | 14 | 0.0 | 1 | K.HKVYACEVTHQGLSSPVTK.S | Carbamidomethyl: 6 | 82-100 | CID |
| 383 | 938.963 | 2 | 528.41 | 0.4959 | 19.14 | 112.87 | 11 | 0.0 | 0 | K.VYACEVTHQGLSSPVTK.S | Carbamidomethyl: 4 | 84-100 | CID |

| Cmpd. | m/z meas. | z | Δ m/z [ppm] | Δ m/z [Da] | Rt [min] | Score | No. of Cmpds. | Site [%] | P | Sequence | Modification | Range | Type |
|---|---|---|---|---|---|---|---|---|---|---|---|---|---|

#### Protein 2: Immunoglobulin heavy constant gamma 1 OS=Homo sapiens GN=IGHG1 PE=1 SV=1

**Accession:** IGHG1_HUMAN
**Database:** SwissProt
**MW [kDa] / pI:** 36.1 / 8.5
**Modification(s):** Carbamidomethyl

**Score:** 1001.6
**Seq. Coverage [%]:** 57.0
**No. of Peptides:** 13

```
        10         20         30         40         50         60         70         80         90        100
ASTKGPSVFP LAPSSKSTSG GTAALGCLVK DYFPEPVTVS WNSGALTSGV HTFPAVLQSS GLYSLSSVVT VPSSSLGTQT YICNVNHKPS NTKVDKKVEP
       110        120        130        140        150        160        170        180        190        200
KSCDKTHTCP PCPAPELLGG PSVFLFPPKP KDTLMISRTP EVTCVVVDVS HEDPEVKFNW YVDGVEVHNA KTKPREEQYN STYRVVSVLT VLHQDWLNGK
       210        220        230        240        250        260        270        280        290        300
EYKCKVSNKA LPAPIEKTIS KAKGQPREPQ VYTLPPSRDE LTKNQVSLTC LVKGFYPSDI AVEWESNGQP ENNYKTTPPV LDSDGSFFLY SKLTVDKSRW
       310        320        330        340
QQGNVFSCSV MHEALHNHYT QKSLSLSPGK
```

**MS/MS Peptide Matches**

| Cmpd. | m/z meas. | z | Δ m/z [ppm] | Δ m/z [Da] | Rt [min] | Score | No. of Cmpds. | Site [%] | P | Sequence | Modification | Range | Type |
|---|---|---|---|---|---|---|---|---|---|---|---|---|---|
| 874 | 830.526 | 3 | 103.00 | 0.0855 | 28.11 | 91.3 | 4 | 0.0 | 1 | K.GPSVFPLAPSSKSTSGGTAALGCLVK.D | Carbamidomethyl: 23 | 5-30 | CID |
| 992 | 1112.594 | 3 | 337.26 | 0.3751 | 30.22 | 78.17 | 14 | 0.0 | 1 | K.SCDKTHTCPPCPAPELLGGPSVFLFPPKPK.D | Carbamidomethyl: 2, 8, 11 | 102-131 | CID |
| 1029 | 949.261 | 3 | 460.54 | 0.4370 | 30.93 | 82.07 | 8 | 0.0 | 0 | K.THTCPPCPAPELLGGPSVFLFPPKPK.D | Carbamidomethyl: 4, 7 | 106-131 | CID |
| 687 | 1070.085 | 2 | 63.22 | 0.0676 | 24.92 | 97.55 | 1 | 0.0 | 0 | R.TPEVTCVVVDVSHEDPEVK.F | Carbamidomethyl: 6 | 139-157 | CID |
| 966 | 950.584 | 4 | 395.34 | 0.3757 | 29.76 | 44.2 | 2 | 0.0 | 1 | R.TPEVTCVVVDVSHEDPEVKFNWYVDGVEVHNAK.T | Carbamidomethyl: 6 | 139-171 | CID |
| 676 | 839.517 | 2 | 133.87 | 0.1124 | 24.73 | 95.85 | 3 | 0.0 | 0 | K.FNWYVDGVEVHNAK.T | | 158-171 | CID |

| Cmpd. | m/z meas. | z | Δ m/z [ppm] | Δ m/z [Da] | Rt [min] | Score | No. of Cmpds. | Site [%] | P | Sequence | Modification | Range | Type |
|---|---|---|---|---|---|---|---|---|---|---|---|---|---|
| 1107 | 905.12 | 2 | 677.84 | 0.6131 | 32.27 | 106.6 | 3 | 0.0 | 0 | R.VVSVLTVLHQDWLNGK.E | | 185-200 | CID |
| 1034 | 743.839 | 3 | 580.69 | 0.4317 | 31.01 | 26.88 | 1 | 0.0 | 1 | R.VVSVLTVLHQDWLNGKEYK.C | | 185-203 | CID |
| 499 | 937.075 | 2 | 92.08 | 0.0863 | 21.44 | 60.17 | 8 | 0.0 | 1 | R.EPQVYTLPPSRDELTK.N | | 228-243 | CID |
| 892 | 1273.152 | 2 | 457.87 | 0.5827 | 28.44 | 111.4 | 11 | 0.0 | 0 | K.GFYPSDIAVEWESNGQPENNYK.T | | 254-275 | CID |
| 956 | 937.596 | 2 | 140.22 | 0.1315 | 29.58 | 95.45 | 6 | 0.0 | 0 | K.TTPPVLDSDGSFFLYSK.L | | 276-292 | CID |
| 587 | 762.202 | 4 | 454.78 | 0.3465 | 23.21 | 41.17 | 3 | 0.0 | 1 | K.SRWQQGNVFSCSVMHEALHNHYTQK.S | Carbamidomethyl: 11 | 298-322 | CID |
| 635 | 934.869 | 3 | 472.78 | 0.4418 | 24.06 | 70.81 | 4 | 0.0 | 0 | R.WQQGNVFSCSVMHEALHNHYTQK.S | Carbamidomethyl: 9 | 300-322 | CID |

#### Protein 3: Keratin, type II cytoskeletal 1 OS=Homo sapiens GN=KRT1 PE=1 SV=6

| | | | |
|---|---|---|---|
| **Accession:** | K2C1_HUMAN | **Score:** | 674.2 |
| **Database:** | SwissProt | **Seq. Coverage [%]:** | 23.4 |
| **MW [kDa] / pI:** | 66.0 / 8.2 | **No. of Peptides:** | 8 |

| 10 | 20 | 30 | 40 | 50 | 60 | 70 | 80 | 90 | 100 |
|---|---|---|---|---|---|---|---|---|---|
| MSRQFSSRSG | YRSGGGFSSG | SAGIINYQRR | TTSSSTRRSG | GGGGRFSSCG | GGGGSFGAGG | GFGSRSLVNL | GGSKSISISV | ARGGGRGSGF | GGGYGGGGFG |
| 110 | 120 | 130 | 140 | 150 | 160 | 170 | 180 | 190 | 200 |
| GGGFGGGGFG | GGGIGGGGFG | GFGSGGGGFG | GGGFGGGGYG | GGYGPVCPPG | GIQEVTINQS | LLQPLNVEID | PEIQKVKSRE | REQIKSLNNQ | FASFIDKVRF |
| 210 | 220 | 230 | 240 | 250 | 260 | 270 | 280 | 290 | 300 |
| LEQQNQVLQT | KWELLQQVDT | STRTHNLEPY | FESFINNLRR | RVDQLKSDQS | RLDSELKNMQ | DMVEDYRNKY | EDEINKRTNA | ENEFVTIKKD | VDGAYMTKVD |
| 310 | 320 | 330 | 340 | 350 | 360 | 370 | 380 | 390 | 400 |
| LQAKLDNLQQ | EIDFLTALYQ | AELSQMQTQI | SETNVILSMD | NNRSLDLDSI | IAEVKAQYED | IAQKSKAEAE | SLYQSKYEEL | QITAGRHGDS | VRNSKIEISE |
| 410 | 420 | 430 | 440 | 450 | 460 | 470 | 480 | 490 | 500 |
| LNRVIQRLRS | EIDNVKKQIS | NLQQSISDAE | QRGENALKDA | KNKLNDLEDA | LQQAKEDLAR | LLRDYQELMN | TKLALDLEIA | TYRTLLEGEE | SRMSGECAPN |
| 510 | 520 | 530 | 540 | 550 | 560 | 570 | 580 | 590 | 600 |
| VSVSVSTSHT | TISGGGSRGG | GGGGYGSGGS | SYGSGGGSYG | SGGGGGGGRG | SYGSGGSSYG | SGGGSYGSGG | GGGGHGSYGS | GSSSGGYRGG | SGGGGGGSSG |
| 610 | 620 | 630 | 640 | 650 | | | | | |
| GRGSGGGSSG | GSIGGRGSSS | GGVKSSGGSS | SVKFVSTTYS | GVTR | | | | | |

**MS/MS Peptide Matches**

| Cmpd. | m/z meas. | z | Δ m/z [ppm] | Δ m/z [Da] | Rt [min] | Score | No. of Cmpds. | Site [%] | P | Sequence | Modification | Range | Type |
|---|---|---|---|---|---|---|---|---|---|---|---|---|---|
| 516 | 829.473 | 2 | 87.93 | 0.0729 | 21.79 | 97.24 | 1 | 0.0 | 0 | R.SGGGFSSGSAGIINYQR.R | | 13-29 | CID |
| 688 | 941.553 | 2 | 68.09 | 0.0641 | 24.93 | 15.02 | 1 | 0.0 | 1 | R.EQIKSLNNQFASFIDK.V | | 182-197 | CID |
| 425 | 738.5 | 2 | 140.48 | 0.1037 | 20.01 | 88.75 | 6 | 0.0 | 0 | R.FLEQQNQVLQTK.W | | 200-211 | CID |
| 1073 | 717.421 | 3 | 79.31 | 0.0569 | 31.67 | 26.22 | 1 | 0.0 | 1 | R.THNLEPYFESFINNLRR.R | | 224-240 | CID |
| 1397 | 783.841 | 3 | 123.36 | 0.0967 | 37.76 | 144.01 | 4 | 0.0 | 1 | R.SLDLDSIIAEVKAQYEDIAQK.S | | 344-364 | CID |
| 507 | 858.999 | 2 | 81.27 | 0.0698 | 21.61 | 98.21 | 3 | 0.0 | 0 | K.QISNLQQSISDAEQR.G | | 418-432 | CID |
| 184 | 1192.587 | 2 | 90.10 | 0.1074 | 14.36 | 183.5 | 5 | 0.0 | 0 | R.GGGGGGYGSGGSSYGSGGGSYGSGGGGGG GR.G | | 519-549 | CID |
| 50 | 876.557 | 2 | 159.76 | 0.1400 | 11.24 | 21.2 | 3 | 0.0 | 1 | R.GSGGGSSGGSIGGRGSSSGGVK.S | | 603-624 | CID |

#### Protein 4: Immunoglobulin lambda constant 3 OS=Homo sapiens GN=IGLC3 PE=1 SV=1

| | | | |
|---|---|---|---|
| **Accession:** | IGLC3_HUMAN | **Score:** | 651.3 |
| **Database:** | SwissProt | **Seq. Coverage [%]:** | 77.4 |
| **MW [kDa] / pI:** | 11.3 / 6.9 | **No. of Peptides:** | 8 |
| **Modification(s):** | Carbamidomethyl | | |

```
        10         20         30         40         50         60         70         80         90        100
GQPKAAPSVT LFPPSSEELQ ANKATLVCLI SDFYPGAVTV AWKADSSPVK AGVETTTPSK QSNNKYAASS YLSLTPEQWK SHKSYSCQVT HEGSTVEKTV
       110
APTECS
```

**MS/MS Peptide Matches**

| Cmpd. | m/z meas. | z | Δ m/z [ppm] | Δ m/z [Da] | Rt [min] | Score | No. of Cmpds. | Site [%] | P | Sequence | Modification | Range | Type |
|---|---|---|---|---|---|---|---|---|---|---|---|---|---|
| 778 | 993.559 | 2 | 46.74 | 0.0464 | 26.44 | 71.58 | 8 | 0.0 | 0 | K.AAPSVTLFPPSSEELQANK.A | | 5-23 | CID |
| 214 | 838.026 | 2 | 113.53 | 0.0951 | 15.04 | 57.08 | 3 | 0.0 | 1 | K.ADSSPVKAGVETTTPSK.Q | | 44-60 | CID |
| 39 | 781.511 | 2 | 149.15 | 0.1165 | 10.90 | 82.31 | 15 | 0.0 | 1 | K.AGVETTTPSKQSNNK.Y | | 51-65 | CID |
| 759 | 772.494 | 3 | 145.52 | 0.1124 | 26.13 | 57.15 | 7 | 0.0 | 1 | K.QSNNKYAASSYLSLTPEQWK.S | | 61-80 | CID |
| 861 | 872.574 | 2 | 161.56 | 0.1409 | 27.89 | 80.62 | 11 | 0.0 | 0 | K.YAASSYLSLTPEQWK.S | | 66-80 | CID |
| 101 | 1032.573 | 2 | 93.77 | 0.0968 | 12.40 | 85.42 | 3 | 0.0 | 1 | K.SHKSYSCQVTHEGSTVEK.T | Carbamidomethyl: 7 | 81-98 | CID |
| 171 | 856.493 | 2 | 128.18 | 0.1098 | 14.04 | 111.39 | 14 | 0.0 | 0 | K.SYSCQVTHEGSTVEK.T | Carbamidomethyl: 4 | 84-98 | CID |
| 373 | 1279.561 | 2 | 389.59 | 0.4983 | 18.94 | 105.79 | 10 | 0.0 | 1 | K.SYSCQVTHEGSTVEKTVAPTECS.- | Carbamidomethyl: 4, 22 | 84-106 | CID |

#### Protein 5: Immunoglobulin heavy constant gamma 4 OS=Homo sapiens GN=IGHG4 PE=1 SV=1

| | | | |
|---|---|---|---|
| **Accession:** | IGHG4_HUMAN | **Score:** | 496.9 |
| **Database:** | SwissProt | **Seq. Coverage [%]:** | 37.0 |
| **MW [kDa] / pI:** | 35.9 / 7.2 | **No. of Peptides:** | 6 |

**Modification(s):** Carbamidomethyl

| | | | | | | | | | |
|---|---|---|---|---|---|---|---|---|---|
| 10 | 20 | 30 | 40 | 50 | 60 | 70 | 80 | 90 | 100 |
| ASTKGPSVFP | LAPCSRSTSE | STAALGCLVK | DYFPEPVTVS | WNSGALTSGV | HTFPAVLQSS | GLYSLSSVVT | VPSSSLGTKT | YTCNVDHKPS | NTKVDKRVES |
| 110 | 120 | 130 | 140 | 150 | 160 | 170 | 180 | 190 | 200 |
| KYGPPCPSCP | APEFLGGPSV | FLFPPKPKDT | LMISRTPEVT | CVVVDVSQED | PEVQFNWYVD | GVEVHNAKTK | PREEQFNSTY | RVVSVLTVLH | QDWLNGKEYK |
| 210 | 220 | 230 | 240 | 250 | 260 | 270 | 280 | 290 | 300 |
| CKVSNKGLPS | SIEKTISKAK | GQPREPQVYT | LPPSQEEMTK | NQVSLTCLVK | GFYPSDIAVE | WESNGQPENN | YKTTPPVLDS | DGSFFLYSRL | TVDKSRWQEG |
| 310 | 320 | 330 | | | | | | | |
| NVFSCSVMHE | ALHNHYTQKS | LSLSLGK | | | | | | | |

**MS/MS Peptide Matches**

| Cmpd. | m/z meas. | z | Δ m/z [ppm] | Δ m/z [Da] | Rt [min] | Score | No. of Cmpds. | Site [%] | P | Sequence | Modification | Range | Type |
|---|---|---|---|---|---|---|---|---|---|---|---|---|---|
| 558 | 712.466 | 2 | 150.91 | 0.1075 | 22.69 | 81.99 | 3 | 0.0 | 0 | R.STSESTAALGCLVK.D | Carbamidomethyl: 11 | 17-30 | CID |
| 53 | 833.003 | 2 | 137.49 | 0.1145 | 11.34 | 50.21 | 6 | 0.0 | 0 | K.TYTCNVDHKPSNTK.V | Carbamidomethyl: 4 | 80-93 | CID |
| 1153 | 982.234 | 3 | 417.67 | 0.4101 | 33.01 | 52.94 | 1 | 0.0 | 0 | K.YGPPCPSCPAPEFLGGPSVFLFPPKPK.D | Carbamidomethyl: 5, 8 | 102-128 | CID |
| 1107 | 905.12 | 2 | 677.84 | 0.6131 | 32.27 | 106.6 | 3 | 0.0 | 0 | R.VVSVLTVLHQDWLNGK.E | | 182-197 | CID |
| 1034 | 743.839 | 3 | 580.69 | 0.4317 | 31.01 | 26.88 | 1 | 0.0 | 1 | R.VVSVLTVLHQDWLNGKEYK.C | | 182-200 | CID |
| 892 | 1273.152 | 2 | 457.87 | 0.5827 | 28.44 | 111.4 | 11 | 0.0 | 0 | K.GFYPSDIAVEWESNGQPENNYK.T | | 251-272 | CID |

#### Protein 6: Keratin, type I cytoskeletal 9 OS=Homo sapiens GN=KRT9 PE=1 SV=3

**Accession:** K1C9_HUMAN
**Database:** SwissProt
**MW [kDa] / pI:** 62.0 / 5.1

**Score:** 454.4
**Seq. Coverage [%]:** 20.4
**No. of Peptides:** 5

| 10 | 20 | 30 | 40 | 50 | 60 | 70 | 80 | 90 | 100 |
|---|---|---|---|---|---|---|---|---|---|
| MSCRQFSSSY | LSRSGGGGGG | GLGSGGSIRS | SYSRFSSSGG | GGGGGRFSSS | SGYGGGSSRV | CGRGGGGSFG | YSYGGGSGGG | FSASSLGGGF | GGGSRGFGGA |
| 110 | 120 | 130 | 140 | 150 | 160 | 170 | 180 | 190 | 200 |
| SGGGYSSSGG | FGGGFGGGSG | GGFGGGYGSG | FGGFGGFGGG | AGGGDGGILT | ANEKSTMQEL | NSRLASYLDK | VQALEEANND | LENKIQDWYD | KKGPAAIQKN |
| 210 | 220 | 230 | 240 | 250 | 260 | 270 | 280 | 290 | 300 |
| YSPYYNTIDD | LKDQIVDLTV | GNNKTLLDID | NTRMTLDDFR | IKFEMEQNLR | QGVDADINGL | RQVLDNLTME | KSDLEMQYET | LQEELMALKK | NHKEEMSQLT |
| 310 | 320 | 330 | 340 | 350 | 360 | 370 | 380 | 390 | 400 |
| GQNSGDVNVE | INVAPGKDLT | KTLNDMRQEY | EQLIAKNRKD | IENQYETQIT | QIEHEVSSSG | QEVQSSAKEV | TQLRHGVQEL | EIELQSQLSK | KAALEKSLED |
| 410 | 420 | 430 | 440 | 450 | 460 | 470 | 480 | 490 | 500 |
| TKNRYCGQLQ | MIQEQISNLE | AQITDVRQEI | ECQNQEYSLL | LSIKMRLEKE | IETYHNLLEG | GQEDFESSGA | GKIGLGGRGG | SGGSYGRGSR | GGSGGSYGGG |
| 510 | 520 | 530 | 540 | 550 | 560 | 570 | 580 | 590 | 600 |
| GSGGGYGGGS | GSRGGSGGSY | GGGSGSGGGS | GGGYGGGSGG | GHSGGSGGGH | SGGSGGNYGG | GSGSGGGSGG | GYGGGSGSRG | GSGGSHGGGS | GFGGESGGSY |
| 610 | 620 | 630 | | | | | | | |
| GGGEEASGSG | GGYGGGSGKS | SHS | | | | | | | |

##### MS/MS Peptide Matches

| Cmpd. | m/z meas. | z | Δ m/z [ppm] | Δ m/z [Da] | Rt [min] | Score | No. of Cmpds. | Site [%] | P | Sequence | Modification | Range | Type |
|---|---|---|---|---|---|---|---|---|---|---|---|---|---|
| 1201 | 968.578 | 3 | 450.67 | 0.4363 | 33.91 | 76.86 | 2 | 0.0 | 1 | K.NYSPYYNTIDDLKDQIVDLTVGNNK.T | | 200-224 | CID |
| 969 | 1089.224 | 3 | 350.09 | 0.3812 | 29.83 | 153.12 | 3 | 0.0 | 0 | K.DIENQYETQITQIEHEVSSSGQEVQSSAK.E | | 340-368 | CID |
| 1089 | 998.576 | 4 | 91.61 | 0.0915 | 31.96 | 30.85 | 3 | 0.0 | 1 | K.DIENQYETQITQIEHEVSSSGQEVQSSAKEVTQLR.H | | 340-374 | CID |
| 95 | 896.474 | 2 | 118.82 | 0.1065 | 12.29 | 153.07 | 3 | 0.0 | 0 | R.GGSGGSYGGGGSGGGYGGGSGSR.G | | 491-513 | CID |
| 213 | 906.468 | 4 | 390.02 | 0.3534 | 15.01 | 40.45 | 1 | 0.0 | 1 | R.GGSGGSHGGGSGFGGESGGSYGGGEEASGSGGGYGGGSGKSSHS.- | | 580-623 | CID |

#### Protein 7: Immunoglobulin lambda constant 1 OS=Homo sapiens GN=IGLC1 PE=1 SV=1

| | | | |
|---|---|---|---|
| **Accession:** | IGLC1_HUMAN | **Score:** | 396.3 |
| **Database:** | SwissProt | **Seq. Coverage [%]:** | 56.6 |
| **MW [kDa] / pI:** | 11.3 / 7.9 | **No. of Peptides:** | 5 |
| **Modification(s):** | Carbamidomethyl | | |

| 10 | 20 | 30 | 40 | 50 | 60 | 70 | 80 | 90 | 100 |
|---|---|---|---|---|---|---|---|---|---|
| GQPKANPTVT | LFPPSSEELQ | ANKATLVCLI | SDFYPGAVTV | AWKADGSPVK | AGVETTKPSK | QSNNKYAASS | YLSLTPEQWK | SHRSYSCQVT | HEGSTVEKTV |
| 110 | | | | | | | | | |
| APTECS | | | | | | | | | |

MS/MS Peptide Matches

| Cmpd. | m/z meas. | z | Δ m/z [ppm] | Δ m/z [Da] | Rt [min] | Score | No. of Cmpds. | Site [%] | P | Sequence | Modification | Range | Type |
|---|---|---|---|---|---|---|---|---|---|---|---|---|---|
| 149 | 836.552 | 2 | 122.86 | 0.1028 | 13.53 | 41.36 | 1 | 0.0 | 1 | K.ADGSPVKAGVETTKPSK.Q | | 44-60 | CID |
| 759 | 772.494 | 3 | 145.52 | 0.1124 | 26.13 | 57.15 | 7 | 0.0 | 1 | K.QSNNKYAASSYLSLTPEQWK.S | | 61-80 | CID |
| 861 | 872.574 | 2 | 161.56 | 0.1409 | 27.89 | 80.62 | 11 | 0.0 | 0 | K.YAASSYLSLTPEQWK.S | | 66-80 | CID |
| 171 | 856.493 | 2 | 128.18 | 0.1098 | 14.04 | 111.39 | 14 | 0.0 | 0 | R.SYSCQVTHEGSTVEK.T | Carbamidomethyl: 4 | 84-98 | CID |
| 373 | 1279.561 | 2 | 389.59 | 0.4983 | 18.94 | 105.79 | 10 | 0.0 | 1 | R.SYSCQVTHEGSTVEKTVAPTECS.- | Carbamidomethyl: 4, 22 | 84-106 | CID |

#### Protein 8: Keratin, type II cytoskeletal 2 epidermal OS=Homo sapiens GN=KRT2 PE=1 SV=2

| | | | |
|---|---|---|---|
| **Accession:** | K22E_HUMAN | **Score:** | 367.7 |
| **Database:** | SwissProt | **Seq. Coverage [%]:** | 8.6 |
| **MW [kDa] / pI:** | 65.4 / 8.1 | **No. of Peptides:** | 3 |

| 10 | 20 | 30 | 40 | 50 | 60 | 70 | 80 | 90 | 100 |
|---|---|---|---|---|---|---|---|---|---|
| MSCQISCKSR | GRGGGGGGFR | GFSSGSAVVS | GGSRRSTSSF | SCLSRHGGGG | GGFGGGGFGS | RSLVGLGGTK | SISISVAGGG | GGFGAAGGFG | GRGGGFGGGS |
| **110** | **120** | **130** | **140** | **150** | **160** | **170** | **180** | **190** | **200** |
| SFGGGSGFSG | GGFGGGGFGG | GRFGGFGGPG | GVGGLGGPGG | FGPGGYPGGI | HEVSVNQSLL | QPLNVKVDPE | IQNVKAQERE | QIKTLNNKFA | SFIDKVRFLE |
| **210** | **220** | **230** | **240** | **250** | **260** | **270** | **280** | **290** | **300** |
| QQNQVLQTKW | ELLQQMNVGT | RPINLEPIFQ | GYIDSLKRYL | DGLTAERTSQ | NSELNNMQDL | VEDYKKKYED | EINKRTAAEN | DFVTLKKDVD | NAYMIKVELQ |
| **310** | **320** | **330** | **340** | **350** | **360** | **370** | **380** | **390** | **400** |
| SKVDLLNQEI | EFLKVLYDAE | ISQIHQSVTD | TNVILSMDNS | RNLDLDSIIA | EVKAQYEEIA | QRSKEEAEAL | YHSKYEELQV | TVGRHGDSLK | EIKIEISELN |
| **410** | **420** | **430** | **440** | **450** | **460** | **470** | **480** | **490** | **500** |
| RVIQRLQGEI | AHVKKQCKNV | QDAIADAEQR | GEHALKDARN | KLNDLEEALQ | QAKEDLARLL | RDYQELMNVK | LALDVEIATY | RKLLEGEECR | MSGDLSSNVT |
| **510** | **520** | **530** | **540** | **550** | **560** | **570** | **580** | **590** | **600** |
| VSVTSSTISS | NVASKAAFGG | SGGRGSSSGG | GYSSGSSSYG | SGGRQSGSRG | GSGGGGSISG | GGYGSGGGSG | GRYGSGGGSK | GGSISGGGYG | SGGGKHSSGG |
| **610** | **620** | **630** | **640** | | | | | | |
| GSRGGSSSGG | GYGSGGGGSS | SVKGSSGEAF | GSSVTFSFR | | | | | | |

**MS/MS Peptide Matches**

| Cmpd. | m/z meas. | z | Δ m/z [ppm] | Δ m/z [Da] | Rt [min] | Score | No. of Cmpds. | Site [%] | P | Sequence | Modification | Range | Type |
|---|---|---|---|---|---|---|---|---|---|---|---|---|---|
| 425 | 738.5 | 2 | 140.48 | 0.1037 | 20.01 | 88.75 | 6 | 0.0 | 0 | R.FLEQQNQVLQTK.W | | 198-209 | CID |
| 107 | 870.943 | 2 | 99.41 | 0.0866 | 12.53 | 113.29 | 3 | 0.0 | 0 | R.GSSSGGGYSSGSSSYGSGGR.Q | | 525-544 | CID |
| 138 | 871.486 | 2 | 124.10 | 0.1081 | 13.20 | 165.63 | 3 | 0.0 | 0 | R.GGSGGGGSISGGGYGSGGGSGGR.Y | | 550-572 | CID |

#### Protein 9: Immunoglobulin heavy constant gamma 2 OS=Homo sapiens GN=IGHG2 PE=1 SV=2

| | | | |
|---|---|---|---|
| **Accession:** | IGHG2_HUMAN | **Score:** | 309.4 |
| **Database:** | SwissProt | **Seq. Coverage [%]:** | 35.3 |
| **MW [kDa] / pI:** | 35.9 / 7.7 | **No. of Peptides:** | 6 |
| **Modification(s):** | Carbamidomethyl | | |

| 10 | 20 | 30 | 40 | 50 | 60 | 70 | 80 | 90 | 100 |
|---|---|---|---|---|---|---|---|---|---|
| ASTKGPSVFP | LAPCSRSTSE | STAALGCLVK | DYFPEPVTVS | WNSGALTSGV | HTFPAVLQSS | GLYSLSSVVT | VPSSNFGTQT | YTCNVDHKPS | NTKVDKTVER |
| 110 | 120 | 130 | 140 | 150 | 160 | 170 | 180 | 190 | 200 |
| KCCVECPPCP | APPVAGPSVF | LFPPKPKDTL | MISRTPEVTC | VVVDVSHEDP | EVQFNWYVDG | VEVHNAKTKP | REEQFNSTFR | VVSVLTVVHQ | DWLNGKEYKC |
| 210 | 220 | 230 | 240 | 250 | 260 | 270 | 280 | 290 | 300 |
| KVSNKGLPAP | IEKTISKTKG | QPREPQVYTL | PPSREEMTKN | QVSLTCLVKG | FYPSDISVEW | ESNGQPENNY | KTTPPMLDSD | GSFFLYSKLT | VDKSRWQQGN |
| 310 | 320 | 330 | | | | | | | |
| VFSCSVMHEA | LHNHYTQKSL | SLSPGK | | | | | | | |

MS/MS Peptide Matches

| Cmpd. | m/z meas. | z | Δ m/z [ppm] | Δ m/z [Da] | Rt [min] | Score | No. of Cmpds. | Site [%] | P | Sequence | Modification | Range | Type |
|---|---|---|---|---|---|---|---|---|---|---|---|---|---|
| 558 | 712.466 | 2 | 150.91 | 0.1075 | 22.69 | 81.99 | 3 | 0.0 | 0 | R.STSESTAALGCLVK.D | Carbamidomethyl: 11 | 17-30 | CID |
| 913 | 760.213 | 4 | 438.73 | 0.3334 | 28.88 | 37.39 | 2 | 0.0 | 1 | R.KCCVECPPCPAPPVAGPSVFLFPPKPK.D | Carbamidomethyl: 2, 3, 6, 9 | 101-127 | CID |
| 967 | 950.573 | 4 | 393.34 | 0.3737 | 29.78 | 15.08 | 1 | 0.0 | 0 | R.TPEVTCVVVDVSHEDPEVQFNWYVDGVEVHNAK.T | Carbamidomethyl: 6 | 135-167 | CID |
| 445 | 953.049 | 2 | 77.91 | 0.0742 | 20.43 | 62.92 | 8 | 0.0 | 1 | R.EPQVYTLPPSREEMTK.N | | 224-239 | CID |
| 587 | 762.202 | 4 | 454.78 | 0.3465 | 23.21 | 41.17 | 3 | 0.0 | 1 | K.SRWQQGNVFSCSVMHEALHNHYTQK.S | Carbamidomethyl: 11 | 294-318 | CID |
| 635 | 934.869 | 3 | 472.78 | 0.4418 | 24.06 | 70.81 | 4 | 0.0 | 0 | R.WQQGNVFSCSVMHEALHNHYTQK.S | Carbamidomethyl: 9 | 296-318 | CID |

#### Protein 10: Keratin, type I cytoskeletal 10 OS=Homo sapiens GN=KRT10 PE=1 SV=6

| | | | |
|---|---|---|---|
| **Accession:** | K1C10_HUMAN | **Score:** | 276.4 |
| **Database:** | SwissProt | **Seq. Coverage [%]:** | 9.9 |
| **MW [kDa] / pI:** | 58.8 / 5.1 | **No. of Peptides:** | 3 |

| 10 | 20 | 30 | 40 | 50 | 60 | 70 | 80 | 90 | 100 |
|---|---|---|---|---|---|---|---|---|---|
| MSVRYSSSKH | YSSSRSGGGG | GGGGCGGGGG | VSSLRISSSK | GSLGGGFSSG | GFSGGSFSRG | SSGGGCFGGS | SGGYGGLGGF | GGGSFRGSYG | SSSFGGSYGG |
| **110** | **120** | **130** | **140** | **150** | **160** | **170** | **180** | **190** | **200** |
| IFGGGSFGGG | SFGGGSFGGG | GFGGGGFGGG | FGGGFGGDGG | LLSGNEKVTM | QNLNDRLASY | LDKVRALEES | NYELEGKIKE | WYEKHGNSHQ | GEPRDYSKYY |
| **210** | **220** | **230** | **240** | **250** | **260** | **270** | **280** | **290** | **300** |
| KTIDDLKNQI | LNLTTDNANI | LLQIDNARLA | ADDFRLKYEN | EVALRQSVEA | DINGLRRVLD | ELTLTKADLE | MQIESLTEEL | AYLKKNHEEE | MKDLRNVSTG |
| **310** | **320** | **330** | **340** | **350** | **360** | **370** | **380** | **390** | **400** |
| DVNVEMNAAP | GVDLTQLLNN | MRSQYEQLAE | QNRKDAEAWF | NEKSKELTTE | IDNNIEQISS | YKSEITELRR | NVQALEIELQ | SQLALKQSLE | ASLAETEGRY |
| **410** | **420** | **430** | **440** | **450** | **460** | **470** | **480** | **490** | **500** |
| CVQLSQIQAQ | ISALEEQLQQ | IRAETECQNT | EYQQLLDIKI | RLENEIQTYR | SLLEGEGSSG | GGGRGGGSFG | GGYGGGSSGG | GSSGGGHGGG | HGGSSGGGYG |
| **510** | **520** | **530** | **540** | **550** | **560** | **570** | **580** | **590** | |
| GGSSGGGSSG | GGYGGGSSSG | GHGGSSSGGY | GGGSSGGGGG | GYGGGSSGGG | SSSGGGYGGG | SSSGGHKSSS | SGSVGESSSK | GPRY | |

MS/MS Peptide Matches

| Cmpd. | m/z meas. | z | Δ m/z [ppm] | Δ m/z [Da] | Rt [min] | Score | No. of Cmpds. | Site [%] | P | Sequence | Modification | Range | Type |
|---|---|---|---|---|---|---|---|---|---|---|---|---|---|
| 621 | 854.47 | 2 | 93.98 | 0.0803 | 23.83 | 152.32 | 4 | 0.0 | 0 | K.GSLGGGFSSGGFSGGSFSR.G | | 41-59 | CID |
| 1248 | 1437.121 | 2 | 293.02 | 0.4210 | 34.86 | 91.82 | 2 | 0.0 | 0 | R.NVSTGDVNVEMNAAPGVDLTQLLNNMR.S | | 296-322 | CID |
| 210 | 747.428 | 2 | 76.55 | 0.0572 | 14.96 | 32.27 | 4 | 0.0 | 1 | R.SQYEQLAEQNRK.D | | 323-334 | CID |

#### Protein 11: Hornerin OS=Homo sapiens GN=HRNR PE=1 SV=2

**Accession:** HORN_HUMAN
**Database:** SwissProt
**MW [kDa] / pI:** 282.2 / 10.0

**Score:** 109.3
**Seq. Coverage [%]:** 0.9
**No. of Peptides:** 1

| 10 | 20 | 30 | 40 | 50 | 60 | 70 | 80 | 90 | 100 |
|---|---|---|---|---|---|---|---|---|---|
| MPKLLQGVIT | VIDVFYQYAT | QHGEYDTLNK | AELKELLENE | FHQILKNPND | PDTVDIILQS | LDRDHNKKVD | FTEYLLMIFK | LVQARNKIIG | KDYCQVSGSK |
| 110 | 120 | 130 | 140 | 150 | 160 | 170 | 180 | 190 | 200 |
| LRDDTHQHQE | EQEETEKEEN | KRQESSFSHS | SWSAGENDSY | SRNVRGSLKP | GTESISRRLS | FQRDFSGQHN | SYSGQSSSYG | EQNSDSHQSS | GRGQCGSGSG |
| 210 | 220 | 230 | 240 | 250 | 260 | 270 | 280 | 290 | 300 |
| QSPNYGQHGS | GSGQSSSNDT | HGSGSGQSSG | FSQHKSSSGQ | SSGYSQHGSG | SGHSSGYGQH | GSRSGQSSRG | ERHRSSSGSS | SSYGQHGSGS | RQSLGHGRQG |
| 310 | 320 | 330 | 340 | 350 | 360 | 370 | 380 | 390 | 400 |
| SGSRQSPSHV | RHGSGSGHSS | SHGQHGSGSS | YSYSRGHYES | GSGQTSGFGQ | HESGSGQSSG | YSKHGSGSGH | SSSQGQHGST | SGQASSSGQH | GSSSRQSSSY |
| 410 | 420 | 430 | 440 | 450 | 460 | 470 | 480 | 490 | 500 |
| GQHESASRHS | SGRGQHSSGS | GQSPGHGQRG | SGSGQSPSSG | QHGTGFGRSS | SSGPYVSGSG | YSSGFGHHES | SSEHSSGYTQ | HGSGSGHSSG | HGQHGSRSGQ |
| 510 | 520 | 530 | 540 | 550 | 560 | 570 | 580 | 590 | 600 |
| SSRGERQGSS | AGSSSSYGQH | GSGSRQSLGH | SRHGSGSGQS | PSPSRGRHES | GSRQSSSYGP | HGYGSGRSSS | RGPYESGSGH | SSGLGHQESR | SGQSSGYGQH |
| 610 | 620 | 630 | 640 | 650 | 660 | 670 | 680 | 690 | 700 |
| GSSSGHSSTH | GQHGSTSGQS | SSCGQHGATS | GQSSSHGQHG | SGSSQSSRYG | QQGSGSGQSP | SRGRHGSDFG | HSSSYGQHGS | GSGWSSSNGP | HGSVSGQSSG |
| 710 | 720 | 730 | 740 | 750 | 760 | 770 | 780 | 790 | 800 |
| FGHKSGSGQS | SGYSQHGSGS | SHSSGYRKHG | SRSGQSSRSE | QHGSSSGLSS | SYGQHGSGSH | QSSGHGRQGS | GSGHSPSRVR | HGSSSGHSSS | HGQHGSGTSC |
| 810 | 820 | 830 | 840 | 850 | 860 | 870 | 880 | 890 | 900 |
| SSSCGHYESG | SGQASGFGQH | ESGSGQGYSQ | HGSASGHFSS | QGRHGSTSGQ | SSSSGQHDSS | SGQSSSYGQH | ESASHHASGR | GRHGSGSGQS | PGHGQRGSGS |
| 910 | 920 | 930 | 940 | 950 | 960 | 970 | 980 | 990 | 1000 |
| GQSPSYGRHG | SGSGRSSSSG | RHGSGSGQSS | GFGHKSSSGQ | SSGYTQHGSG | SGHSSSYEQH | GSRSGQSSRS | EQHGSSSGSS | SSYGQHGSGS | RQSLGHGQHG |
| 1010 | 1020 | 1030 | 1040 | 1050 | 1060 | 1070 | 1080 | 1090 | 1100 |
| SGSGQSPSPS | RGRHGSGSGQ | SSSYGPYRSG | SGWSSSRGPY | ESGSGHSSGL | GHRESRSGQS | SGYGQHGSSS | GHSSTHGQHG | STSGQSSSCG | QHGASSGQSS |
| 1110 | 1120 | 1130 | 1140 | 1150 | 1160 | 1170 | 1180 | 1190 | 1200 |
| SHGQHGSGSS | QSSGYGRQGS | GSGQSPGHGQ | RGSGSRQSPS | YGRHGSGSGR | SSSSGQHGSG | LGESSGFGHH | ESSSGQSSSY | SQHGSGSGHS | SGYGQHGSRS |
| 1210 | 1220 | 1230 | 1240 | 1250 | 1260 | 1270 | 1280 | 1290 | 1300 |
| GQSSRGERHG | SSSGSSSHYG | QHGSGSRQSS | GHGRQGSGSG | HSPSRGRHGS | GLGHSSSHGQ | HGSGSGRSSS | RGPYESRSGH | SSVFGQHESG | SGHSSAYSQH |
| [illegible] | [illegible] | [illegible] | [illegible] | [illegible] | [illegible] | [illegible] | [illegible] | [illegible] | [illegible] |

MS/MS Peptide Matches

| Cmpd. | m/z meas. | z | Δ m/z [ppm] | Δ m/z [Da] | Rt [min] | Score | No. of Cmpds. | Site [%] | P | Sequence | Modification | Range | Type |
|---|---|---|---|---|---|---|---|---|---|---|---|---|---|
| 70 | 826.808 | 3 | 560.10 | 0.4628 | 11.82 | 109.31 | 2 | 0.0 | 0 | R.HGSGSGHSSSYGQHGSGSGWSSSSGR.H | | 1365-1390 | CID |

#### Protein 12: Immunoglobulin lambda variable 3-21 OS=Homo sapiens GN=IGLV3-21 PE=1 SV=2

| | | | |
|---|---|---|---|
| **Accession:** | LV321_HUMAN | **Score:** | 84.9 |
| **Database:** | SwissProt | **Seq. Coverage [%]:** | 13.7 |
| **MW [kDa] / pI:** | 12.4 / 5.0 | **No. of Peptides:** | 1 |

```
        10         20         30         40         50         60         70         80         90        100
MAWTVLLLGL LSHCTGSVTS YVLTQPPSVS VAPGQTARIT CGGNNIGSKS VHWYQQKPGQ APVLVVYDDS DRPSGIPERF SGSNSGNTAT LTISRVEAGD
       110        120
EADYYCQVWD SSSDHPT
```

MS/MS Peptide Matches

| Cmpd. | m/z meas. | z | Δ m/z [ppm] | Δ m/z [Da] | Rt [min] | Score | No. of Cmpds. | Site [%] | P | Sequence | Modification | Range | Type |
|---|---|---|---|---|---|---|---|---|---|---|---|---|---|
| 391 | 806.962 | 2 | 76.95 | 0.0621 | 19.34 | 84.86 | 2 | 0.0 | 0 | R.FSGSNSGNTATLTISR.V | | 80-95 | CID |

#### Protein 13: Immunoglobulin heavy variable 4-4 OS=Homo sapiens GN=IGHV4-4 PE=3 SV=2

| | | | |
|---|---|---|---|
| **Accession:** | HV404_HUMAN | **Score:** | 78.9 |
| **Database:** | SwissProt | **Seq. Coverage [%]:** | 13.7 |
| **MW [kDa] / pI:** | 12.8 / 9.6 | **No. of Peptides:** | 1 |
| **Modification(s):** | Carbamidomethyl | | |

| 10 | 20 | 30 | 40 | 50 | 60 | 70 | 80 | 90 | 100 |
|---|---|---|---|---|---|---|---|---|---|
| MKHLWFFLLL | VAAPRWVLSQ | VQLQESGPGL | VKPSGTLSLT | CAVSGGSISS | SNWWSWVRQP | PGKGLEWIGE | IYHSGSTNYN | PSLKSRVTIS | VDKSKNQFSL |
| 110 | 120 | | | | | | | | |
| KLSSVTAADT | AVYYCAR | | | | | | | | |

MS/MS Peptide Matches

| Cmpd. | m/z meas. | z | Δ m/z [ppm] | Δ m/z [Da] | Rt [min] | Score | No. of Cmpds. | Site [%] | P | Sequence | Modification | Range | Type |
|---|---|---|---|---|---|---|---|---|---|---|---|---|---|
| 540 | 874.512 | 2 | 105.65 | 0.0924 | 22.36 | 78.93 | 3 | 0.0 | 0 | K.LSSVTAADTAVYYCAR.- | Carbamidomethyl: 14 | 102-117 | CID |

#### Protein 14: Immunoglobulin kappa variable 3-15 OS=Homo sapiens GN=IGKV3-15 PE=1 SV=2

| | | | |
|---|---|---|---|
| **Accession:** | KV315_HUMAN | **Score:** | 75.4 |
| **Database:** | SwissProt | **Seq. Coverage [%]:** | 18.3 |
| **MW [kDa] / pI:** | 12.5 / 5.1 | **No. of Peptides:** | 1 |

| 10 | 20 | 30 | 40 | 50 | 60 | 70 | 80 | 90 | 100 |
|---|---|---|---|---|---|---|---|---|---|
| MEAPAQLLFL | LLLWLPDTTG | EIVMTQSPAT | LSVSPGERAT | LSCRASQSVS | SNLAWYQQKP | GQAPRLLIYG | ASTRATGIPA | RFSGSGSGTE | FTLTISSLQS |
| 110 | 120 | | | | | | | | |
| EDFAVYYCQQ | YNNWP | | | | | | | | |

MS/MS Peptide Matches

| Cmpd. | m/z meas. | z | Δ m/z [ppm] | Δ m/z [Da] | Rt [min] | Score | No. of Cmpds. | Site [%] | P | Sequence | Modification | Range | Type |
|---|---|---|---|---|---|---|---|---|---|---|---|---|---|
| 501 | 768.489 | 3 | 130.03 | 0.0999 | 21.49 | 75.39 | 3 | 0.0 | 0 | R.ASQSVSSNLAWYQQKPGQAPR.L | | 45-65 | CID |

#### Protein 15: Immunoglobulin kappa variable 2D-28 OS=Homo sapiens GN=IGKV2D-28 PE=1 SV=2

| | | | |
|---|---|---|---|
| **Accession:** | KVD28_HUMAN | **Score:** | 50.6 |
| **Database:** | SwissProt | **Seq. Coverage [%]:** | 16.7 |
| **MW [kDa] / pI:** | 12.9 / 5.6 | **No. of Peptides:** | 1 |

| 10 | 20 | 30 | 40 | 50 | 60 | 70 | 80 | 90 | 100 |
|---|---|---|---|---|---|---|---|---|---|
| MRLPAQLLGL | LMLWVSGSSG | DIVMTQSPLS | LPVTPGEPAS | ISCRSSQSLL | HSNGYNYLDW | YLQKPGQSPQ | LLIYLGSNRA | SGVPDRFSGS | GSGTDFTLKI |
| 110 | 120 | 130 | | | | | | | |
| SRVEAEDVGV | YYCMQALQTP | | | | | | | | |

MS/MS Peptide Matches

| Cmpd. | m/z meas. | z | Δ m/z [ppm] | Δ m/z [Da] | Rt [min] | Score | No. of Cmpds. | Site [%] | P | Sequence | Modification | Range | Type |
|---|---|---|---|---|---|---|---|---|---|---|---|---|---|
| 544 | 993.528 | 2 | 46.51 | 0.0462 | 22.41 | 50.59 | 3 | 0.0 | 1 | R.ASGVPDRFSGSGSGTDFTLK.I | | 80-99 | CID |

BRUKER

#### Project Info

| | | | |
|---|---|---|---|
| **Name:** | Alagesan | **Date:** | Mar 5, 2019 |
| **Note:** | Nanobooster | | |

#### Sample Info & Protocols

| | |
|---|---|
| **Name:** | Nanobooster - IgG |
| **Date:** | Mar 5, 2019 |

#### Search Result Info

| | | | |
|---|---|---|---|
| **Search Result:** | IonTrap_Human_Swiss_CID_2019-03-05 17:55:08 | | |
| **Location:** | /Alagesan/Nanobooster - IgG/KA_IgG2 HD NB ACN CID_GA3_01_9081.mgf | | |
| **Search Method:** | IonTrap_Human_Swiss_CID | **Version:** | 1.2 |
| **Search Engine(s):** | Mascot, 2.6.0 | **Ident. Compound(s):** | 0/2323 |
| **Database(s):** | SwissProt, SwissProt_2017_04.fasta | | |
| **Note:** | | | |

#### Results

##### Protein 1: Immunoglobulin kappa constant OS=Homo sapiens GN=IGKC PE=1 SV=2

| | | | |
|---|---|---|---|
| **Accession:** | IGKC_HUMAN | **Score:** | 1023.4 |
| **Database:** | SwissProt | **Seq. Coverage [%]:** | 88.8 |
| **MW [kDa] / pI:** | 11.8 / 6.1 | **No. of Peptides:** | 10 |
| **Modification(s):** | Carbamidomethyl | | |

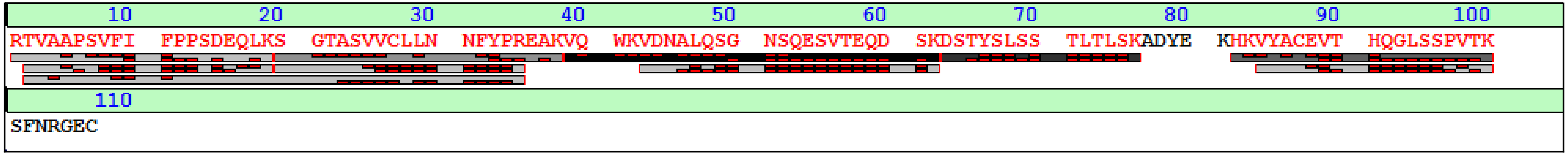


MS/MS Peptide Matches

| Cmpd. | m/z meas. | z | Δ m/z [ppm] | Δ m/z [Da] | Rt [min] | Score | No. of Cmpds. | Site [%] | P | Sequence | Modification | Range | Type |
|---|---|---|---|---|---|---|---|---|---|---|---|---|---|
| 924 | 701.774 | 3 | 560.50 | 0.3931 | 28.73 | 45.91 | 3 | 0.0 | 1 | -.RTVAAPSVFIFPPSDEQLK.S | | 1-19 | CID |
| 1049 | 973.619 | 2 | 104.66 | 0.1019 | 30.86 | 78.47 | 10 | 0.0 | 0 | R.TVAAPSVFIFPPSDEQLK.S | | 2-19 | CID |
| 1483 | 1242.632 | 3 | 262.17 | 0.3257 | 38.10 | 79.62 | 18 | 0.0 | 1 | R.TVAAPSVFIFPPSDEQLKSGTASVVCLLNNFYPR.E | Carbamidomethyl: 26 | 2-35 | CID |
| 1182 | 899.569 | 2 | 130.91 | 0.1177 | 33.05 | 68.19 | 14 | 0.0 | 0 | K.SGTASVVCLLNNFYPR.E | Carbamidomethyl: 8 | 20-35 | CID |
| 1064 | 709.411 | 3 | 69.80 | 0.0495 | 31.12 | 101.87 | 3 | 0.0 | 1 | K.SGTASVVCLLNNFYPREAK.V | Carbamidomethyl: 8 | 20-38 | CID |
| 483 | 893.501 | 3 | 454.77 | 0.4062 | 21.01 | 139.59 | 13 | 0.0 | 1 | K.VQWKVDNALQSGNSQESVTEQDSK.D | | 39-62 | CID |
| 249 | 1069.056 | 2 | 531.59 | 0.5680 | 16.10 | 157.72 | 18 | 0.0 | 0 | K.VDNALQSGNSQESVTEQDSK.D | | 43-62 | CID |
| 696 | 751.974 | 2 | 121.22 | 0.0911 | 24.90 | 102.54 | 3 | 0.0 | 0 | K.DSTYSLSSTLTLSK.A | | 63-76 | CID |
| 290 | 714.457 | 3 | 128.62 | 0.0919 | 16.97 | 140.74 | 9 | 0.0 | 1 | K.HKVYACEVTHQGLSSPVTK.S | Carbamidomethyl: 6 | 82-100 | CID |
| 380 | 938.759 | 2 | 311.04 | 0.2919 | 19.06 | 108.73 | 12 | 0.0 | 0 | K.VYACEVTHQGLSSPVTK.S | Carbamidomethyl: 4 | 84-100 | CID |

#### Protein 2: Immunoglobulin kappa light chain OS=Homo sapiens PE=1 SV=1

| | | | |
|---|---|---|---|
| **Accession:** | IGK_HUMAN | **Score:** | 926.8 |
| **Database:** | SwissProt | **Seq. Coverage [%]:** | 43.9 |
| **MW [kDa] / pI:** | 23.4 / 6.9 | **No. of Peptides:** | 8 |
| **Modification(s):** | Carbamidomethyl | | |

| 10 | 20 | 30 | 40 | 50 | 60 | 70 | 80 | 90 | 100 |
|---|---|---|---|---|---|---|---|---|---|
| DIQMTQSPST | LSASVGDRVT | ITCRASQSIN | TWLAWYQQKP | GKAPKLLMYK | ASSLESGVPS | RFIGSGSGTE | FTLTISSLQP | DDFATYYCQQ | YNSDSKMFGQ |
| **110** | **120** | **130** | **140** | **150** | **160** | **170** | **180** | **190** | **200** |
| GTKVEVKGTV | AAPSVFIFPP | SDEQLKSGTA | SVVCLLNNFY | PREAKVQWKV | DNALQSGNSQ | ESVTEQDSKD | STYSLSSTLT | LSKADYEKHK | VYACEVTHQG |
| **210** | **220** | | | | | | | | |
| LSSPVTKSFN | RGEC | | | | | | | | |

##### MS/MS Peptide Matches

| Cmpd. | m/z meas. | z | Δ m/z [ppm] | Δ m/z [Da] | Rt [min] | Score | No. of Cmpds. | Site [%] | P | Sequence | Modification | Range | Type |
|---|---|---|---|---|---|---|---|---|---|---|---|---|---|
| 549 | 947.01 | 2 | 58.55 | 0.0554 | 22.27 | 107.37 | 3 | 0.0 | 0 | -.DIQMTQSPSTLSASVGDR.V | | 1-18 | CID |
| 1182 | 899.569 | 2 | 130.91 | 0.1177 | 33.05 | 68.19 | 14 | 0.0 | 0 | K.SGTASVVCLLNNFYPR.E | Carbamidomethyl: 8 | 127-142 | CID |
| 1064 | 709.411 | 3 | 69.80 | 0.0495 | 31.12 | 101.87 | 3 | 0.0 | 1 | K.SGTASVVCLLNNFYPREAK.V | Carbamidomethyl: 8 | 127-145 | CID |
| 483 | 893.501 | 3 | 454.77 | 0.4062 | 21.01 | 139.59 | 13 | 0.0 | 1 | K.VQWKVDNALQSGNSQESVTEQDSK.D | | 146-169 | CID |
| 249 | 1069.056 | 2 | 531.59 | 0.5680 | 16.10 | 157.72 | 18 | 0.0 | 0 | K.VDNALQSGNSQESVTEQDSK.D | | 150-169 | CID |
| 696 | 751.974 | 2 | 121.22 | 0.0911 | 24.90 | 102.54 | 3 | 0.0 | 0 | K.DSTYSLSSTLTLSK.A | | 170-183 | CID |
| 290 | 714.457 | 3 | 128.62 | 0.0919 | 16.97 | 140.74 | 9 | 0.0 | 1 | K.HKVYACEVTHQGLSSPVTK.S | Carbamidomethyl: 6 | 189-207 | CID |
| 380 | 938.759 | 2 | 311.04 | 0.2919 | 19.06 | 108.73 | 12 | 0.0 | 0 | K.VYACEVTHQGLSSPVTK.S | Carbamidomethyl: 4 | 191-207 | CID |

#### Protein 3: Immunoglobulin heavy constant gamma 2 OS=Homo sapiens GN=IGHG2 PE=1 SV=2

| | | | |
|---|---|---|---|
| **Accession:** | IGHG2_HUMAN | **Score:** | 870.4 |
| **Database:** | SwissProt | **Seq. Coverage [%]:** | 55.2 |
| **MW [kDa] / pI:** | 35.9 / 7.7 | **No. of Peptides:** | 14 |
| **Modification(s):** | Carbamidomethyl, Oxidation | | |

| 10 | 20 | 30 | 40 | 50 | 60 | 70 | 80 | 90 | 100 |
|---|---|---|---|---|---|---|---|---|---|
| ASTKGPSVFP | LAPCSRSTSE | STAALGCLVK | DYFPEPVTVS | WNSGALTSGV | HTFPAVLQSS | GLYSLSSVVT | VPSSNFGTQT | YTCNVDHKPS | NTKVDKTVER |
| **110** | **120** | **130** | **140** | **150** | **160** | **170** | **180** | **190** | **200** |
| KCCVECPPCP | APPVAGPSVF | LFPPKPKDTL | MISRTPEVTC | VVVDVSHEDP | EVQFNWYVDG | VEVHNAKTKP | REEQFNSTFR | VVSVLTVVHQ | DWLNGKEYKC |
| **210** | **220** | **230** | **240** | **250** | **260** | **270** | **280** | **290** | **300** |
| KVSNKGLPAP | IEKTISKTKG | QPREPQVYTL | PPSREEMTKN | QVSLTCLVKG | FYPSDISVEW | ESNGQPENNY | KTTPPMLDSD | GSFFLYSKLT | VDKSRWQQGN |
| **310** | **320** | **330** | | | | | | | |
| VFSCSVMHEA | LHNHYTQKSL | SLSPGK | | | | | | | |

**MS/MS Peptide Matches**

| Cmpd. | m/z meas. | z | Δ m/z [ppm] | Δ m/z [Da] | Rt [min] | Score | No. of Cmpds. | Site [%] | P | Sequence | Modification | Range | Type |
|---|---|---|---|---|---|---|---|---|---|---|---|---|---|
| 558 | 712.456 | 2 | 136.87 | 0.0975 | 22.44 | 88.23 | 23 | 0.0 | 0 | R.STSESTAALGCLVK.D | Carbamidomethyl: 11 | 17-30 | CID |
| 916 | 1013.245 | 3 | 402.76 | 0.4079 | 28.60 | 80.12 | 5 | 0.0 | 1 | R.KCCVECPPCPAPPVAGPSVFLFPPKPK.D | Carbamidomethyl: 2, 3, 6, 9 | 101-127 | CID |
| 1000 | 970.554 | 3 | 428.04 | 0.4153 | 29.97 | 68.15 | 14 | 0.0 | 0 | K.CCVECPPCPAPPVAGPSVFLFPPKPK.D | Carbamidomethyl: 1, 2, 5, 8 | 102-127 | CID |
| 1043 | 1242.628 | 3 | 282.11 | 0.3505 | 30.75 | 50.55 | 6 | 0.0 | 1 | K.CCVECPPCPAPPVAGPSVFLFPPKPKDTLMIS R.T | Carbamidomethyl: 1, 2, 5, 8 | 102-134 | CID |
| 1151 | 1154.342 | 4 | 33.49 | 0.0387 | 32.54 | 39.61 | 1 | 0.0 | 1 | K.DTLMISRTPEVTCVVVDVSHEDPEVQFNWYVD GVEVHNAK.T | Carbamidomethyl: 13 | 128-167 | CID |

| Cmpd. | m/z meas. | z | Δ m/z [ppm] | Δ m/z [Da] | Rt [min] | Score | No. of Cmpds. | Site [%] | P | Sequence | Modification | Range | Type |
|---|---|---|---|---|---|---|---|---|---|---|---|---|---|
| 1072 | 1266.91 | 3 | 247.45 | 0.3134 | 31.24 | 50.81 | 4 | 0.0 | 0 | R.TPEVTCVVVDVSHEDPEVQFNWYVDGVEVHNAK.T | Carbamidomethyl: 6 | 135-167 | CID |
| 917 | 738.83 | 3 | 128.02 | 0.0946 | 28.60 | 75.69 | 4 | 0.0 | 1 | R.VVSVLTVVHQDWLNGKEYK.C | | 181-199 | CID |
| 500 | 953.002 | 2 | 28.59 | 0.0272 | 21.35 | 48.48 | 9 | 0.0 | 1 | R.EPQVYTLPPSREEMTK.N | | 224-239 | CID |
| 847 | 1280.546 | 2 | -16.23 | -0.0208 | 27.35 | 70.89 | 3 | 0.0 | 0 | K.GFYPSDISVEWESNGQPENNYK.T | | 250-271 | CID |
| 999 | 953.58 | 2 | 135.73 | 0.1294 | 29.97 | 82.41 | 6 | 0.0 | 0 | K.TTPPMLDSDGSFFLYSK.L | | 272-288 | CID |
| 888 | 961.524 | 2 | 79.00 | 0.0760 | 28.09 | 67.22 | 6 | 0.0 | 0 | K.TTPPMLDSDGSFFLYSK.L | Oxidation: 5 | 272-288 | CID |
| 598 | 762.205 | 4 | 458.72 | 0.3495 | 23.23 | 56.59 | 3 | 0.0 | 1 | K.SRWQQGNVFSCSVMHEALHNHYTQK.S | Carbamidomethyl: 11 | 294-318 | CID |
| 641 | 934.849 | 3 | 451.38 | 0.4218 | 23.97 | 61.73 | 4 | 0.0 | 0 | R.WQQGNVFSCSVMHEALHNHYTQK.S | Carbamidomethyl: 9 | 296-318 | CID |
| 486 | 940.162 | 3 | 428.98 | 0.4031 | 21.05 | 29.89 | 1 | 0.0 | 0 | R.WQQGNVFSCSVMHEALHNHYTQK.S | Carbamidomethyl: 9; Oxidation: 12 | 296-318 | CID |

#### Protein 4: Immunoglobulin lambda constant 3 OS=Homo sapiens GN=IGLC3 PE=1 SV=1

**Accession:** IGLC3_HUMAN
**Database:** SwissProt
**MW [kDa] / pI:** 11.3 / 6.9
**Modification(s):** Carbamidomethyl

**Score:** 686.1
**Seq. Coverage [%]:** 77.4
**No. of Peptides:** 8

```
         10         20         30         40         50         60         70         80         90        100
GQPKAAPSVT LFPPSSEELQ ANKATLVCLI SDFYPGAVTV AWKADSSPVK AGVETTTPSK QSNNKYAASS YLSLTPEQWK SHKSYSCQVT HEGSTVEKTV
        110
APTECS
```

MS/MS Peptide Matches

| Cmpd. | m/z meas. | z | Δ m/z [ppm] | Δ m/z [Da] | Rt [min] | Score | No. of Cmpds. | Site [%] | P | Sequence | Modification | Range | Type |
|---|---|---|---|---|---|---|---|---|---|---|---|---|---|
| 813 | 994.052 | 2 | 542.96 | 0.5394 | 26.77 | 37.34 | 2 | 0.0 | 0 | K.AAPSVTLFPPSSEELQANK.A | | 5-23 | CID |

| Cmpd. | m/z meas. | z | Δ m/z [ppm] | Δ m/z [Da] | Rt [min] | Score | No. of Cmpds. | Site [%] | P | Sequence | Modification | Range | Type |
|---|---|---|---|---|---|---|---|---|---|---|---|---|---|
| 193 | 838.02 | 2 | 106.37 | 0.0891 | 15.01 | 76.64 | 6 | 0.0 | 1 | K.ADSSPVKAGVETTTPSK.Q | | 44-60 | CID |
| 36 | 781.526 | 2 | 168.34 | 0.1315 | 10.71 | 81.65 | 15 | 0.0 | 1 | K.AGVETTTPSKQSNNK.Y | | 51-65 | CID |
| 778 | 772.471 | 3 | 115.74 | 0.0894 | 26.22 | 73.52 | 4 | 0.0 | 1 | K.QSNNKYAASSYLSLTPEQWK.S | | 61-80 | CID |
| 877 | 872.532 | 2 | 113.42 | 0.0989 | 27.91 | 94.2 | 10 | 0.0 | 0 | K.YAASSYLSLTPEQWK.S | | 66-80 | CID |
| 97 | 1032.615 | 2 | 134.45 | 0.1388 | 12.34 | 115.88 | 9 | 0.0 | 1 | K.SHKSYSCQVTHEGSTVEK.T | Carbamidomethyl: 7 | 81-98 | CID |
| 155 | 856.498 | 2 | 134.01 | 0.1148 | 14.07 | 99.37 | 15 | 0.0 | 0 | K.SYSCQVTHEGSTVEK.T | Carbamidomethyl: 4 | 84-98 | CID |
| 359 | 1279.565 | 2 | 392.72 | 0.5023 | 18.68 | 107.49 | 8 | 0.0 | 1 | K.SYSCQVTHEGSTVEKTVAPTECS.- | Carbamidomethyl: 4, 22 | 84-106 | CID |

#### Protein 5: Immunoglobulin lambda constant 2 OS=Homo sapiens GN=IGLC2 PE=1 SV=1

**Accession:** IGLC2_HUMAN
**Database:** SwissProt
**MW [kDa] / pI:** 11.3 / 6.9
**Modification(s):** Carbamidomethyl

**Score:** 607.3
**Seq. Coverage [%]:** 77.4
**No. of Peptides:** 8

```
         10         20         30         40         50         60         70         80         90        100
GQPKAAPSVT LFPPSSEELQ ANKATLVCLI SDFYPGAVTV AWKADSSPVK AGVETTTPSK QSNNKYAASS YLSLTPEQWK SHRSYSCQVT HEGSTVEKTV
       110
APTECS
```

MS/MS Peptide Matches

| Cmpd. | m/z meas. | z | Δ m/z [ppm] | Δ m/z [Da] | Rt [min] | Score | No. of Cmpds. | Site [%] | P | Sequence | Modification | Range | Type |
|---|---|---|---|---|---|---|---|---|---|---|---|---|---|
| 813 | 994.052 | 2 | 542.96 | 0.5394 | 26.77 | 37.34 | 2 | 0.0 | 0 | K.AAPSVTLFPPSSEELQANK.A | | 5-23 | CID |
| 193 | 838.02 | 2 | 106.37 | 0.0891 | 15.01 | 76.64 | 6 | 0.0 | 1 | K.ADSSPVKAGVETTTPSK.Q | | 44-60 | CID |
| 36 | 781.526 | 2 | 168.34 | 0.1315 | 10.71 | 81.65 | 15 | 0.0 | 1 | K.AGVETTTPSKQSNNK.Y | | 51-65 | CID |
| 778 | 772.471 | 3 | 115.74 | 0.0894 | 26.22 | 73.52 | 4 | 0.0 | 1 | K.QSNNKYAASSYLSLTPEQWK.S | | 61-80 | CID |

| Cmpd. | m/z meas. | z | Δ m/z [ppm] | Δ m/z [Da] | Rt [min] | Score | No. of Cmpds. | Site [%] | P | Sequence | Modification | Range | Type |
|---|---|---|---|---|---|---|---|---|---|---|---|---|---|
| 877 | 872.532 | 2 | 113.42 | 0.0989 | 27.91 | 94.2 | 10 | 0.0 | 0 | K.YAASSYLSLTPEQWK.S | | 66-80 | CID |
| 102 | 1047.046 | 2 | 541.57 | 0.5667 | 12.45 | 37.1 | 4 | 0.0 | 1 | K.SHRSYSCQVTHEGSTVEK.T | Carbamidomethyl: 7 | 81-98 | CID |
| 155 | 856.498 | 2 | 134.01 | 0.1148 | 14.07 | 99.37 | 15 | 0.0 | 0 | R.SYSCQVTHEGSTVEK.T | Carbamidomethyl: 4 | 84-98 | CID |
| 359 | 1279.565 | 2 | 392.72 | 0.5023 | 18.68 | 107.49 | 8 | 0.0 | 1 | R.SYSCQVTHEGSTVEKTVAPTECS.- | Carbamidomethyl: 4, 22 | 84-106 | CID |

#### Protein 6: Immunoglobulin lambda constant 1 OS=Homo sapiens GN=IGLC1 PE=1 SV=1

**Accession:** IGLC1_HUMAN
**Database:** SwissProt
**MW [kDa] / pI:** 11.3 / 7.9
**Modification(s):** Carbamidomethyl
**Score:** 498.6
**Seq. Coverage [%]:** 59.4
**No. of Peptides:** 6

```
         10         20         30         40         50         60         70         80         90        100
GQPKANPTVT LFPPSSEELQ ANKATLVCLI SDFYPGAVTV AWKADGSPVK AGVETTKPSK QSNNKYAASS YLSLTPEQWK SHRSYSCQVT HEGSTVEKTV

        110
APTECS
```

MS/MS Peptide Matches

| Cmpd. | m/z meas. | z | Δ m/z [ppm] | Δ m/z [Da] | Rt [min] | Score | No. of Cmpds. | Site [%] | P | Sequence | Modification | Range | Type |
|---|---|---|---|---|---|---|---|---|---|---|---|---|---|
| 129 | 836.554 | 2 | 125.25 | 0.1048 | 13.45 | 86.91 | 3 | 0.0 | 1 | K.ADGSPVKAGVETTKPSK.Q | | 44-60 | CID |
| 778 | 772.471 | 3 | 115.74 | 0.0894 | 26.22 | 73.52 | 4 | 0.0 | 1 | K.QSNNKYAASSYLSLTPEQWK.S | | 61-80 | CID |
| 877 | 872.532 | 2 | 113.42 | 0.0989 | 27.91 | 94.2 | 10 | 0.0 | 0 | K.YAASSYLSLTPEQWK.S | | 66-80 | CID |
| 102 | 1047.046 | 2 | 541.57 | 0.5667 | 12.45 | 37.1 | 4 | 0.0 | 1 | K.SHRSYSCQVTHEGSTVEK.T | Carbamidomethyl: 7 | 81-98 | CID |
| 155 | 856.498 | 2 | 134.01 | 0.1148 | 14.07 | 99.37 | 15 | 0.0 | 0 | R.SYSCQVTHEGSTVEK.T | Carbamidomethyl: 4 | 84-98 | CID |
| 359 | 1279.565 | 2 | 392.72 | 0.5023 | 18.68 | 107.49 | 8 | 0.0 | 1 | R.SYSCQVTHEGSTVEKTVAPTECS.- | Carbamidomethyl: 4, 22 | 84-106 | CID |

#### Protein 7: Immunoglobulin heavy constant gamma 4 OS=Homo sapiens GN=IGHG4 PE=1 SV=1

**Accession:** IGHG4_HUMAN
**Database:** SwissProt
**MW [kDa] / pI:** 35.9 / 7.2
**Modification(s):** Carbamidomethyl, Oxidation

**Score:** 374.4
**Seq. Coverage [%]:** 31.2
**No. of Peptides:** 4

| 10 | 20 | 30 | 40 | 50 | 60 | 70 | 80 | 90 | 100 |
|---|---|---|---|---|---|---|---|---|---|
| ASTKGPSVFP | LAPCSRSTSE | STAALGCLVK | DYFPEPVTVS | WNSGALTSGV | HTFPAVLQSS | GLYSLSSVVT | VPSSSLGTKT | YTCNVDHKPS | NTKVDKRVES |
| 110 | 120 | 130 | 140 | 150 | 160 | 170 | 180 | 190 | 200 |
| KYGPPCPSCP | APEFLGGPSV | FLFPPKPKDT | LMISRTPEVT | CVVVDVSQED | PEVQFNWYVD | GVEVHNAKTK | PREEQFNSTY | RVVSVLTVLH | QDWLNGKEYK |
| 210 | 220 | 230 | 240 | 250 | 260 | 270 | 280 | 290 | 300 |
| CKVSNKGLPS | SIEKTISKAK | GQPREPQVYT | LPPSQEEMTK | NQVSLTCLVK | GFYPSDIAVE | WESNGQPENN | YKTTPPVLDS | DGSFFLYSRL | TVDKSRWQEG |
| 310 | 320 | 330 | | | | | | | |
| NVFSCSVMHE | ALHNHYTQKS | LSLSLGK | | | | | | | |

MS/MS Peptide Matches

| Cmpd. | m/z meas. | z | Δ m/z [ppm] | Δ m/z [Da] | Rt [min] | Score | No. of Cmpds. | Site [%] | P | Sequence | Modification | Range | Type |
|---|---|---|---|---|---|---|---|---|---|---|---|---|---|
| 558 | 712.456 | 2 | 136.87 | 0.0975 | 22.44 | 88.23 | 23 | 0.0 | 0 | R.STSESTAALGCLVK.D | Carbamidomethyl: 11 | 17-30 | CID |
| 58 | 833.002 | 2 | 136.29 | 0.1135 | 11.34 | 51.93 | 5 | 0.0 | 0 | K.TYTCNVDHKPSNTK.V | Carbamidomethyl: 4 | 80-93 | CID |
| 1180 | 982.252 | 3 | 436.01 | 0.4281 | 33.00 | 58.59 | 2 | 0.0 | 0 | K.YGPPCPSCPAPEFLGGPSVFLFPPKPK.D | Carbamidomethyl: 5, 8 | 102-128 | CID |
| 906 | 1273.135 | 2 | 444.51 | 0.5657 | 28.41 | 90.76 | 4 | 0.0 | 0 | K.GFYPSDIAVEWESNGQPENNYK.T | | 251-272 | CID |

#### Protein 8: Immunoglobulin gamma-1 heavy chain OS=Homo sapiens PE=1 SV=1

**Accession:** IGG1_HUMAN
**Database:** SwissProt
**MW [kDa] / pI:** 49.3 / 8.9
**Modification(s):** Carbamidomethyl, Oxidation

**Score:** 352.6
**Seq. Coverage [%]:** 21.6
**No. of Peptides:** 6

| 10 | 20 | 30 | 40 | 50 | 60 | 70 | 80 | 90 | 100 |
|---|---|---|---|---|---|---|---|---|---|
| QVQLVQSGGG | VVQPGRSLRL | SCAASGFTFS | RYTIHWVRQA | PGKGLEWVAV | MSYBGBBKHY | ADSVNGRFTI | SRNDSKNTLY | LNMNSLRPED | TAVYYCARIR |
| **110** | **120** | **130** | **140** | **150** | **160** | **170** | **180** | **190** | **200** |
| DTAMFFAHWG | QGTLVTVSSA | STKGPSVFPL | APSSKSTSGG | TAALGCLVKD | YFPEPVTVSW | NSGALTSGVH | TFPAVLQSSG | LYSLSSVVTV | PSSSLGTQTY |
| **210** | **220** | **230** | **240** | **250** | **260** | **270** | **280** | **290** | **300** |
| ICNVNHKPSN | TKVDKKVEPK | SCDKTHTCPP | CPAPELLGGP | SVFLFPPKPK | DTLMISRTPE | VTCVVVDVSH | EDPEVKFNWY | VDGVEVHNAK | TKPREEQYNS |
| **310** | **320** | **330** | **340** | **350** | **360** | **370** | **380** | **390** | **400** |
| TYRVVSVLTV | LHQDWLNGKE | YKCKVSNKAL | PAPIEKTISK | AKGQPREPQV | YTLPPSRDEL | TKNQVSLTCL | VKGFYPSDIA | VEWESNGQPE | NNYKTTPPVL |
| **410** | **420** | **430** | **440** | **450** | | | | | |
| DSDGSFFLYS | KLTVDKSRWQ | QGNVFSCSVM | HEALHNHYTQ | KSLSLSPGK | | | | | |

**MS/MS Peptide Matches**

| Cmpd. | m/z meas. | z | Δ m/z [ppm] | Δ m/z [Da] | Rt [min] | Score | No. of Cmpds. | Site [%] | P | Sequence | Modification | Range | Type |
|---|---|---|---|---|---|---|---|---|---|---|---|---|---|
| 1072 | 1266.91 | 3 | 237.87 | 0.3013 | 31.24 | 50.81 | 4 | 0.0 | 1 | R.TPEVTCVVVDVSHEDPEVKFNWYVDGVEVHNA K.T | Carbamidomethyl: 6 | 258-290 | CID |
| 906 | 1273.135 | 2 | 444.51 | 0.5657 | 28.41 | 90.76 | 4 | 0.0 | 0 | K.GFYPSDIAVEWESNGQPENNYK.T | | 373-394 | CID |
| 983 | 937.538 | 2 | 78.35 | 0.0735 | 29.69 | 62.77 | 3 | 0.0 | 0 | K.TTPPVLDSDGSFFLYSK.L | | 395-411 | CID |
| 598 | 762.205 | 4 | 458.72 | 0.3495 | 23.23 | 56.59 | 3 | 0.0 | 1 | K.SRWQQGNVFSCSVMHEALHNHYTQK.S | Carbamidomethyl: 11 | 417-441 | CID |
| 641 | 934.849 | 3 | 451.38 | 0.4218 | 23.97 | 61.73 | 4 | 0.0 | 0 | R.WQQGNVFSCSVMHEALHNHYTQK.S | Carbamidomethyl: 9 | 419-441 | CID |
| 486 | 940.162 | 3 | 428.98 | 0.4031 | 21.05 | 29.89 | 1 | 0.0 | 0 | R.WQQGNVFSCSVMHEALHNHYTQK.S | Carbamidomethyl: 9; Oxidation: 12 | 419-441 | CID |

#### Protein 9: Keratin, type II cytoskeletal 1 OS=Homo sapiens GN=KRT1 PE=1 SV=6

**Accession:** K2C1_HUMAN
**Database:** SwissProt
**MW [kDa] / pI:** 66.0 / 8.2

**Score:** 204.7
**Seq. Coverage [%]:** 4.8
**No. of Peptides:** 1

| 10 | 20 | 30 | 40 | 50 | 60 | 70 | 80 | 90 | 100 |
|---|---|---|---|---|---|---|---|---|---|
| MSRQFSSRSG | YRSGGGFSSG | SAGIINYQRR | TTSSSTRRSG | GGGGRFSSCG | GGGGSFGAGG | GFGSRSLVNL | GGSKSISISV | ARGGGRGSGF | GGGYGGGGFG |
| 110 | 120 | 130 | 140 | 150 | 160 | 170 | 180 | 190 | 200 |
| GGGFGGGGFG | GGGIGGGGFG | GFGSGGGGFG | GGGFGGGGYG | GGYGPVCPPG | GIQEVTINQS | LLQPLNVEID | PEIQKVKSRE | REQIKSLNNQ | FASFIDKVRF |
| 210 | 220 | 230 | 240 | 250 | 260 | 270 | 280 | 290 | 300 |
| LEQQNQVLQT | KWELLQQVDT | STRTHNLEPY | FESFINNLRR | RVDQLKSDQS | RLDSELKNMQ | DMVEDYRNKY | EDEINKRTNA | ENEFVTIKKD | VDGAYMTKVD |
| 310 | 320 | 330 | 340 | 350 | 360 | 370 | 380 | 390 | 400 |
| LQAKLDNLQQ | EIDFLTALYQ | AELSQMQTQI | SETNVILSMD | NNRSLDLDSI | IAEVKAQYED | IAQKSKAEAE | SLYQSKYEEL | QITAGRHGDS | VRNSKIEISE |
| 410 | 420 | 430 | 440 | 450 | 460 | 470 | 480 | 490 | 500 |
| LNRVIQRLRS | EIDNVKKQIS | NLQQSISDAE | QRGENALKDA | KNKLNDLEDA | LQQAKEDLAR | LLRDYQELMN | TKLALDLEIA | TYRTLLEGEE | SRMSGECAPN |
| 510 | 520 | 530 | 540 | 550 | 560 | 570 | 580 | 590 | 600 |
| VSVSVSTSHT | TISGGGSRGG | GGGGYGSGGS | SYGSGGGSYG | SGGGGGGGRG | SYGSGGSSYG | SGGGSYGSGG | GGGGHGSYGS | GSSSGGYRGG | SGGGGGGSSG |
| 610 | 620 | 630 | 640 | 650 | | | | | |
| GRGSGGGSSG | GSIGGRGSSS | GGVKSSGGSS | SVKFVSTTYS | GVTR | | | | | |

MS/MS Peptide Matches

| Cmpd. | m/z meas. | z | Δ m/z [ppm] | Δ m/z [Da] | Rt [min] | Score | No. of Cmpds. | Site [%] | P | Sequence | Modification | Range | Type |
|---|---|---|---|---|---|---|---|---|---|---|---|---|---|
| 166 | 1192.494 | 2 | 12.11 | 0.0144 | 14.36 | 204.67 | 4 | 0.0 | 0 | R.GGGGGGYGSGGSSYGSGGGSYGSGGGGGG GR.G | | 519-549 | CID |

#### Protein 10: Keratin, type I cytoskeletal 9 OS=Homo sapiens GN=KRT9 PE=1 SV=3

| | | | |
|---|---|---|---|
| **Accession:** | K1C9_HUMAN | **Score:** | 132.6 |
| **Database:** | SwissProt | **Seq. Coverage [%]:** | 3.7 |
| **MW [kDa] / pI:** | 62.0 / 5.1 | **No. of Peptides:** | 1 |

| 10 | 20 | 30 | 40 | 50 | 60 | 70 | 80 | 90 | 100 |
|---|---|---|---|---|---|---|---|---|---|
| MSCRQFSSSY | LSRSGGGGGG | GLGSGGSIRS | SYSRFSSSGG | GGGGGRFSSS | SGYGGGSSRV | CGRGGGGSFG | YSYGGGSGGG | FSASSLGGGF | GGGSRGFGGA |
| 110 | 120 | 130 | 140 | 150 | 160 | 170 | 180 | 190 | 200 |
| SGGGYSSSGG | FGGGFGGGSG | GGFGGGYGSG | FGGFGGFGGG | AGGGDGGILT | ANEKSTMQEL | NSRLASYLDK | VQALEEANND | LENKIQDWYD | KKGPAAIQKN |
| 210 | 220 | 230 | 240 | 250 | 260 | 270 | 280 | 290 | 300 |
| YSPYYNTIDD | LKDQIVDLTV | GNNKTLLDID | NTRMTLDDFR | IKFEMEQNLR | QGVDADINGL | RQVLDNLTME | KSDLEMQYET | LQEELMALKK | NHKEEMSQLT |
| 310 | 320 | 330 | 340 | 350 | 360 | 370 | 380 | 390 | 400 |
| GQNSGDVNVE | INVAPGKDLT | KTLNDMRQEY | EQLIAKNRKD | IENQYETQIT | QIEHEVSSSG | QEVQSSAKEV | TQLRHGVQEL | EIELQSQLSK | KAALEKSLED |
| 410 | 420 | 430 | 440 | 450 | 460 | 470 | 480 | 490 | 500 |
| TKNRYCGQLQ | MIQEQISNLE | AQITDVRQEI | ECQNQEYSLL | LSIKMRLEKE | IETYHNLLEG | GQEDFESSGA | GKIGLGGRGG | SGGSYGRGSR | GGSGGSYGGG |
| 510 | 520 | 530 | 540 | 550 | 560 | 570 | 580 | 590 | 600 |
| GSGGGYGGGS | GSRGGSGGSY | GGGSGSGGGS | GGGYGGGSGG | GHSGGSGGGH | SGGSGGNYGG | GSGSGGGSGG | GYGGGSGSRG | GSGGSHGGGS | GFGGESGGSY |
| 610 | 620 | 630 | | | | | | | |
| GGGEEASGSG | GGYGGGSGKS | SHS | | | | | | | |

MS/MS Peptide Matches

| Cmpd. | m/z meas. | z | Δ m/z [ppm] | Δ m/z [Da] | Rt [min] | Score | No. of Cmpds. | Site [%] | P | Sequence | Modification | Range | Type |
|---|---|---|---|---|---|---|---|---|---|---|---|---|---|
| 93 | 896.459 | 2 | 102.09 | 0.0915 | 12.24 | 132.56 | 3 | 0.0 | 0 | R.GGSGGSYGGGGSGGGYGGGSGSR.G | | 491-513 | CID |

#### Protein 11: Keratin, type II cytoskeletal 2 epidermal OS=Homo sapiens GN=KRT2 PE=1 SV=2

| | | | |
|---|---|---|---|
| **Accession:** | K22E_HUMAN | **Score:** | 122.6 |
| **Database:** | SwissProt | **Seq. Coverage [%]:** | 3.6 |
| **MW [kDa] / pI:** | 65.4 / 8.1 | **No. of Peptides:** | 1 |

| 10 | 20 | 30 | 40 | 50 | 60 | 70 | 80 | 90 | 100 |
|---|---|---|---|---|---|---|---|---|---|
| MSCQISCKSR | GRGGGGGGFR | GFSSGSAVVS | GGSRRSTSSF | SCLSRHGGGG | GGFGGGGFGS | RSLVGLGGTK | SISISVAGGG | GGFGAAGGFG | GRGGGFGGGS |
| 110 | 120 | 130 | 140 | 150 | 160 | 170 | 180 | 190 | 200 |
| SFGGGSGFSG | GGFGGGGFGG | GRFGGFGGPG | GVGGLGGPGG | FGPGGYPGGI | HEVSVNQSLL | QPLNVKVDPE | IQNVKAQERE | QIKTLNNKFA | SFIDKVRFLE |
| 210 | 220 | 230 | 240 | 250 | 260 | 270 | 280 | 290 | 300 |
| QQNQVLQTKW | ELLQQMNVGT | RPINLEPIFQ | GYIDSLKRYL | DGLTAERTSQ | NSELNNMQDL | VEDYKKKYED | EINKRTAAEN | DFVTLKKDVD | NAYMIKVELQ |
| 310 | 320 | 330 | 340 | 350 | 360 | 370 | 380 | 390 | 400 |
| SKVDLLNQEI | EFLKVLYDAE | ISQIHQSVTD | TNVILSMDNS | RNLDLDSIIA | EVKAQYEEIA | QRSKEEAEAL | YHSKYEELQV | TVGRHGDSLK | EIKIEISELN |
| 410 | 420 | 430 | 440 | 450 | 460 | 470 | 480 | 490 | 500 |
| RVIQRLQGEI | AHVKKQCKNV | QDAIADAEQR | GEHALKDARN | KLNDLEEALQ | QAKEDLARLL | RDYQELMNVK | LALDVEIATY | RKLLEGEECR | MSGDLSSNVT |
| 510 | 520 | 530 | 540 | 550 | 560 | 570 | 580 | 590 | 600 |
| VSVTSSTISS | NVASKAAFGG | SGGRGSSSGG | GYSSGSSSYG | SGGRQSGSRG | GSGGGGSISG | GGYGSGGGSG | GRYGSGGGSK | GGSISGGGYG | SGGGKHSSGG |
| 610 | 620 | 630 | 640 | | | | | | |
| GSRGGSSSGG | GYGSGGGGSS | SVKGSSGEAF | GSSVTFSFR | | | | | | |

MS/MS Peptide Matches

| Cmpd. | m/z meas. | z | Δ m/z [ppm] | Δ m/z [Da] | Rt [min] | Score | No. of Cmpds. | Site [%] | P | Sequence | Modification | Range | Type |
|---|---|---|---|---|---|---|---|---|---|---|---|---|---|
| 121 | 871.452 | 2 | 85.08 | 0.0741 | 13.13 | 122.64 | 3 | 0.0 | 0 | R.GGSGGGGSISGGGYGSGGGSGGR.Y | | 550-572 | CID |

#### Protein 12: Keratin, type I cytoskeletal 10 OS=Homo sapiens GN=KRT10 PE=1 SV=6

**Accession:** K1C10_HUMAN
**Database:** SwissProt
**MW [kDa] / pI:** 58.8 / 5.1

**Score:** 110.4
**Seq. Coverage [%]:** 3.3
**No. of Peptides:** 1

| 10 | 20 | 30 | 40 | 50 | 60 | 70 | 80 | 90 | 100 |
|---|---|---|---|---|---|---|---|---|---|
| MSVRYSSSKH | YSSSRSGGGG | GGGGCGGGGG | VSSLRISSSK | GSLGGGFSSG | GFSGGSFSRG | SSGGGCFGGS | SGGYGGLGGF | GGGSFRGSYG | SSSFGGSYGG |
| **110** | **120** | **130** | **140** | **150** | **160** | **170** | **180** | **190** | **200** |
| IFGGGSFGGG | SFGGGSFGGG | GFGGGGFGGG | FGGGFGGDGG | LLSGNEKVTM | QNLNDRLASY | LDKVRALEES | NYELEGKIKE | WYEKHGNSHQ | GEPRDYSKYY |
| **210** | **220** | **230** | **240** | **250** | **260** | **270** | **280** | **290** | **300** |
| KTIDDLKNQI | LNLTTDNANI | LLQIDNARLA | ADDFRLKYEN | EVALRQSVEA | DINGLRRVLD | ELTLTKADLE | MQIESLTEEL | AYLKKNHEEE | MKDLRNVSTG |
| **310** | **320** | **330** | **340** | **350** | **360** | **370** | **380** | **390** | **400** |
| DVNVEMNAAP | GVDLTQLLNN | MRSQYEQLAE | QNRKDAEAWF | NEKSKELTTE | IDNNIEQISS | YKSEITELRR | NVQALEIELQ | SQLALKQSLE | ASLAETEGRY |
| **410** | **420** | **430** | **440** | **450** | **460** | **470** | **480** | **490** | **500** |
| CVQLSQIQAQ | ISALEEQLQQ | IRAETECQNT | EYQQLLDIKI | RLENEIQTYR | SLLEGEGSSG | GGGRGGGSFG | GGYGGGSSGG | GSSGGGHGGG | HGGSSGGGYG |
| **510** | **520** | **530** | **540** | **550** | **560** | **570** | **580** | **590** | |
| GGSSGGGSSG | GGYGGGSSSG | GHGGSSSGGY | GGGSSGGGGG | GYGGGSSGGG | SSSGGGYGGG | SSSGGHKSSS | SGSVGESSSK | GPRY | |

MS/MS Peptide Matches

| Cmpd. | m/z meas. | z | Δ m/z [ppm] | Δ m/z [Da] | Rt [min] | Score | No. of Cmpds. | Site [%] | P | Sequence | Modification | Range | Type |
|---|---|---|---|---|---|---|---|---|---|---|---|---|---|
| 632 | 854.471 | 2 | 95.15 | 0.0813 | 23.82 | 110.4 | 3 | 0.0 | 0 | K.GSLGGGFSSGGFSGGSFSR.G | | 41-59 | CID |

#### Protein 13: Immunoglobulin kappa variable 3-15 OS=Homo sapiens GN=IGKV3-15 PE=1 SV=2

**Accession:** KV315_HUMAN
**Database:** SwissProt
**MW [kDa] / pI:** 12.5 / 5.1
**Score:** 89.7
**Seq. Coverage [%]:** 18.3
**No. of Peptides:** 1

| 10 | 20 | 30 | 40 | 50 | 60 | 70 | 80 | 90 | 100 |
|---|---|---|---|---|---|---|---|---|---|
| MEAPAQLLFL | LLLWLPDTTG | EIVMTQSPAT | LSVSPGERAT | LSCRASQSVS | SNLAWYQQKP | GQAPRLLIYG | ASTRATGIPA | RFSGSGSGTE | FTLTISSLQS |
| **110** | **120** | | | | | | | | |
| EDFAVYYCQQ | YNNWP | | | | | | | | |

MS/MS Peptide Matches

| Cmpd. | m/z meas. | z | Δ m/z [ppm] | Δ m/z [Da] | Rt [min] | Score | No. of Cmpds. | Site [%] | P | Sequence | Modification | Range | Type |
|---|---|---|---|---|---|---|---|---|---|---|---|---|---|
| 504 | 768.5 | 3 | 144.34 | 0.1109 | 21.43 | 89.65 | 3 | 0.0 | 0 | R.ASQSVSSNLAWYQQKPGQAPR.L | | 45-65 | CID |

#### Protein 14: Immunoglobulin heavy variable 4-4 OS=Homo sapiens GN=IGHV4-4 PE=3 SV=2

| | | | |
|---|---|---|---|
| **Accession:** | HV404_HUMAN | **Score:** | 86.5 |
| **Database:** | SwissProt | **Seq. Coverage [%]:** | 13.7 |
| **MW [kDa] / pI:** | 12.8 / 9.6 | **No. of Peptides:** | 1 |
| **Modification(s):** | Carbamidomethyl | | |

| 10 | 20 | 30 | 40 | 50 | 60 | 70 | 80 | 90 | 100 |
|---|---|---|---|---|---|---|---|---|---|
| MKHLWFFLLL | VAAPRWVLSQ | VQLQESGPGL | VKPSGTLSLT | CAVSGGSISS | SNWWSWVRQP | PGKGLEWIGE | IYHSGSTNYN | PSLKSRVTIS | VDKSKNQFSL |

| 110 | 120 |
|---|---|
| KLSSVTAADT | AVYYCAR |

MS/MS Peptide Matches

| Cmpd. | m/z meas. | z | Δ m/z [ppm] | Δ m/z [Da] | Rt [min] | Score | No. of Cmpds. | Site [%] | P | Sequence | Modification | Range | Type |
|---|---|---|---|---|---|---|---|---|---|---|---|---|---|
| 555 | 874.482 | 2 | 71.34 | 0.0624 | 22.38 | 86.5 | 3 | 0.0 | 0 | K.LSSVTAADTAVYYCAR.- | Carbamidomethyl: 14 | 102-117 | CID |

#### Project Info

| | | | |
|---|---|---|---|
| **Name:** | Alagesan | **Date:** | Mar 5, 2019 |
| **Note:** | Nanobooster | | |

#### Sample Info & Protocols

| | |
|---|---|
| **Name:** | Nanobooster - IgG |
| **Date:** | Mar 5, 2019 |

#### Search Result Info

| | | | |
|---|---|---|---|
| **Search Result:** | IonTrap_Human_Swiss_CID_2019-03-05 17:55:08 | | |
| **Location:** | /Alagesan/Nanobooster - IgG/KA_IgG2 HD NB ACN CID_GA3_01_9080.mgf | | |
| **Search Method:** | IonTrap_Human_Swiss_CID | **Version:** | 1.2 |
| **Search Engine(s):** | Mascot, 2.6.0 | **Ident. Compound(s):** | 0/2310 |
| **Database(s):** | SwissProt, SwissProt_2017_04.fasta | | |
| **Note:** | | | |

#### Results

##### Protein 1: Immunoglobulin kappa constant OS=Homo sapiens GN=IGKC PE=1 SV=2

| | | | |
|---|---|---|---|
| **Accession:** | IGKC_HUMAN | **Score:** | 1143.4 |
| **Database:** | SwissProt | **Seq. Coverage [%]:** | 88.8 |
| **MW [kDa] / pI:** | 11.8 / 6.1 | **No. of Peptides:** | 11 |
| **Modification(s):** | Carbamidomethyl | | |

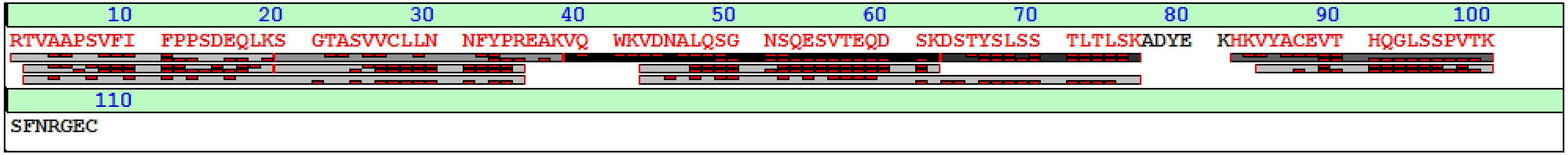


**MS/MS Peptide Matches**

| Cmpd. | m/z meas. | z | Δ m/z [ppm] | Δ m/z [Da] | Rt [min] | Score | No. of Cmpds. | Site [%] | P | Sequence | Modification | Range | Type |
|---|---|---|---|---|---|---|---|---|---|---|---|---|---|
| 908 | 701.455 | 3 | 105.69 | 0.0741 | 28.70 | 50.28 | 3 | 0.0 | 1 | -.RTVAAPSVFIFPPSDEQLK.S | | 1-19 | CID |
| 1035 | 973.637 | 2 | 123.15 | 0.1199 | 30.84 | 90.33 | 10 | 0.0 | 0 | R.TVAAPSVFIFPPSDEQLK.S | | 2-19 | CID |
| 1495 | 1242.706 | 3 | 321.74 | 0.3997 | 38.54 | 97.95 | 17 | 0.0 | 1 | R.TVAAPSVFIFPPSDEQLKSGTASVVCLLNNFYPR.E | Carbamidomethyl: 26 | 2-35 | CID |
| 1176 | 899.568 | 2 | 129.80 | 0.1167 | 33.17 | 75.61 | 16 | 0.0 | 0 | K.SGTASVVCLLNNFYPR.E | Carbamidomethyl: 8 | 20-35 | CID |
| 1054 | 709.455 | 3 | 131.83 | 0.0935 | 31.15 | 101.39 | 3 | 0.0 | 1 | K.SGTASVVCLLNNFYPREAK.V | Carbamidomethyl: 8 | 20-38 | CID |
| 481 | 893.508 | 3 | 462.61 | 0.4132 | 21.20 | 136.87 | 13 | 0.0 | 1 | K.VQWKVDNALQSGNSQESVTEQDSK.D | | 39-62 | CID |
| 220 | 1068.644 | 2 | 146.00 | 0.1560 | 15.82 | 155.63 | 17 | 0.0 | 0 | K.VDNALQSGNSQESVTEQDSK.D | | 43-62 | CID |
| 684 | 1207.595 | 3 | 292.99 | 0.3537 | 24.91 | 102.28 | 3 | 0.0 | 1 | K.VDNALQSGNSQESVTEQDSKDSTYSLSSTLTLSK.A | | 43-76 | CID |
| 682 | 751.97 | 2 | 115.90 | 0.0871 | 24.88 | 88.92 | 4 | 0.0 | 0 | K.DSTYSLSSTLTLSK.A | | 63-76 | CID |
| 276 | 714.466 | 3 | 141.22 | 0.1009 | 16.99 | 137.54 | 9 | 0.0 | 1 | K.HKVYACEVTHQGLSSPVTK.S | Carbamidomethyl: 6 | 82-100 | CID |
| 368 | 938.598 | 2 | 139.48 | 0.1309 | 19.04 | 106.62 | 13 | 0.0 | 0 | K.VYACEVTHQGLSSPVTK.S | Carbamidomethyl: 4 | 84-100 | CID |

| Cmpd. | m/z meas. | z | Δ m/z [ppm] | Δ m/z [Da] | Rt [min] | Score | No. of Cmpds. | Site [%] | P | Sequence | Modification | Range | Type |
|---|---|---|---|---|---|---|---|---|---|---|---|---|---|

#### Protein 2: Immunoglobulin kappa light chain OS=Homo sapiens PE=1 SV=1

**Accession:** IGK_HUMAN
**Database:** SwissProt
**MW [kDa] / pI:** 23.4 / 6.9
**Modification(s):** Carbamidomethyl

**Score:** 1002.1
**Seq. Coverage [%]:** 43.9
**No. of Peptides:** 9

```
        10         20         30         40         50         60         70         80         90        100
DIQMTQSPST LSASVGDRVT ITCRASQSIN TWLAWYQQKP GKAPKLLMYK ASSLESGVPS RFIGSGSGTE FTLTISSLQP DDFATYYCQQ YNSDSKMFGQ
       110        120        130        140        150        160        170        180        190        200
GTKVEVKGTV AAPSVFIFPP SDEQLKSGTA SVVCLLNNFY PREAKVQWKV DNALQSGNSQ ESVTEQDSKD STYSLSSTLT LSKADYEKHK VYACEVTHQG
       210        220
LSSPVTKSFN RGEC
```

##### MS/MS Peptide Matches

| Cmpd. | m/z meas. | z | Δ m/z [ppm] | Δ m/z [Da] | Rt [min] | Score | No. of Cmpds. | Site [%] | P | Sequence | Modification | Range | Type |
|---|---|---|---|---|---|---|---|---|---|---|---|---|---|
| 536 | 947.002 | 2 | 50.10 | 0.0474 | 22.25 | 97.22 | 3 | 0.0 | 0 | -.DIQMTQSPSTLSASVGDR.V | | 1-18 | CID |
| 1176 | 899.568 | 2 | 129.80 | 0.1167 | 33.17 | 75.61 | 16 | 0.0 | 0 | K.SGTASVVCLLNNFYPR.E | Carbamidomethyl: 8 | 127-142 | CID |
| 1054 | 709.455 | 3 | 131.83 | 0.0935 | 31.15 | 101.39 | 3 | 0.0 | 1 | K.SGTASVVCLLNNFYPREAK.V | Carbamidomethyl: 8 | 127-145 | CID |
| 481 | 893.508 | 3 | 462.61 | 0.4132 | 21.20 | 136.87 | 13 | 0.0 | 1 | K.VQWKVDNALQSGNSQESVTEQDSK.D | | 146-169 | CID |
| 220 | 1068.644 | 2 | 146.00 | 0.1560 | 15.82 | 155.63 | 17 | 0.0 | 0 | K.VDNALQSGNSQESVTEQDSK.D | | 150-169 | CID |
| 684 | 1207.595 | 3 | 292.99 | 0.3537 | 24.91 | 102.28 | 3 | 0.0 | 1 | K.VDNALQSGNSQESVTEQDSKDSTYSLSSTLTLSK.A | | 150-183 | CID |
| 682 | 751.97 | 2 | 115.90 | 0.0871 | 24.88 | 88.92 | 4 | 0.0 | 0 | K.DSTYSLSSTLTLSK.A | | 170-183 | CID |
| 276 | 714.466 | 3 | 141.22 | 0.1009 | 16.99 | 137.54 | 9 | 0.0 | 1 | K.HKVYACEVTHQGLSSPVTK.S | Carbamidomethyl: 6 | 189-207 | CID |

| Cmpd. | m/z meas. | z | Δ m/z [ppm] | Δ m/z [Da] | Rt [min] | Score | No. of Cmpds. | Site [%] | P | Sequence | Modification | Range | Type |
|---|---|---|---|---|---|---|---|---|---|---|---|---|---|
| 368 | 938.598 | 2 | 139.48 | 0.1309 | 19.04 | 106.62 | 13 | 0.0 | 0 | K.VYACEVTHQGLSSPVTK.S | Carbamidomethyl: 4 | 191-207 | CID |

#### Protein 3: Immunoglobulin heavy constant gamma 2 OS=Homo sapiens GN=IGHG2 PE=1 SV=2

| | | | |
|---|---|---|---|
| **Accession:** | IGHG2_HUMAN | **Score:** | 851.0 |
| **Database:** | SwissProt | **Seq. Coverage [%]:** | 55.2 |
| **MW [kDa] / pI:** | 35.9 / 7.7 | **No. of Peptides:** | 11 |
| **Modification(s):** | Carbamidomethyl, Oxidation | | |

```
        10         20         30         40         50         60         70         80         90        100
ASTKGPSVFP LAPCSRSTSE STAALGCLVK DYFPEPVTVS WNSGALTSGV HTFPAVLQSS GLYSLSSVVT VPSSNFGTQT YTCNVDHKPS NTKVDKTVER
       110        120        130        140        150        160        170        180        190        200
KCCVECPPCP APPVAGPSVF LFPPKPKDTL MISRTPEVTC VVVDVSHEDP EVQFNWYVDG VEVHNAKTKP REEQFNSTFR VVSVLTVVHQ DWLNGKEYKC
       210        220        230        240        250        260        270        280        290        300
KVSNKGLPAP IEKTISKTKG QPREPQVYTL PPSREEMTKN QVSLTCLVKG FYPSDISVEW ESNGQPENNY KTTPPMLDSD GSFFLYSKLT VDKSRWQQGN
       310        320        330
VFSCSVMHEA LHNHYTQKSL SLSPGK
```

##### MS/MS Peptide Matches

| Cmpd. | m/z meas. | z | Δ m/z [ppm] | Δ m/z [Da] | Rt [min] | Score | No. of Cmpds. | Site [%] | P | Sequence | Modification | Range | Type |
|---|---|---|---|---|---|---|---|---|---|---|---|---|---|
| 554 | 712.479 | 2 | 169.16 | 0.1205 | 22.61 | 81.7 | 21 | 0.0 | 0 | R.STSESTAALGCLVK.D | Carbamidomethyl: 11 | 17-30 | CID |
| 900 | 1013.265 | 3 | 422.51 | 0.4279 | 28.56 | 94.3 | 6 | 0.0 | 1 | R.KCCVECPPCPAPPVAGPSVFLFPPKPK.D | Carbamidomethyl: 2, 3, 6, 9 | 101-127 | CID |
| 1014 | 970.582 | 3 | 456.90 | 0.4433 | 30.45 | 76.7 | 10 | 0.0 | 0 | K.CCVECPPCPAPPVAGPSVFLFPPKPK.D | Carbamidomethyl: 1, 2, 5, 8 | 102-127 | CID |

| Cmpd. | m/z meas. | z | Δ m/z [ppm] | Δ m/z [Da] | Rt [min] | Score | No. of Cmpds. | Site [%] | P | Sequence | Modification | Range | Type |
|---|---|---|---|---|---|---|---|---|---|---|---|---|---|
| 1037 | 1242.628 | 3 | 282.11 | 0.3505 | 30.85 | 64.23 | 5 | 0.0 | 1 | K.CCVECPPCPAPPVAGPSVFLFPPKPKDTLMIS R.T | Carbamidomethyl: 1, 2, 5, 8 | 102-134 | CID |
| 904 | 738.801 | 3 | 88.76 | 0.0656 | 28.61 | 62.89 | 3 | 0.0 | 1 | R.VVSVLTVVHQDWLNGKEYK.C | | 181-199 | CID |
| 439 | 953.06 | 2 | 89.45 | 0.0852 | 20.42 | 72.96 | 7 | 0.0 | 1 | R.EPQVYTLPPSREEMTK.N | | 224-239 | CID |
| 836 | 1281.02 | 2 | 353.92 | 0.4532 | 27.42 | 53.05 | 3 | 0.0 | 0 | K.GFYPSDISVEWESNGQPENNYK.T | | 250-271 | CID |
| 986 | 953.583 | 2 | 138.88 | 0.1324 | 29.97 | 100.63 | 7 | 0.0 | 0 | K.TTPPMLDSDGSFFLYSK.L | | 272-288 | CID |
| 878 | 961.527 | 2 | 82.12 | 0.0790 | 28.16 | 59.49 | 5 | 0.0 | 0 | K.TTPPMLDSDGSFFLYSK.L | Oxidation: 5 | 272-288 | CID |
| 580 | 762.182 | 4 | 428.53 | 0.3265 | 23.14 | 49.62 | 3 | 0.0 | 1 | K.SRWQQGNVFSCSVMHEALHNHYTQK.S | Carbamidomethyl: 11 | 294-318 | CID |
| 629 | 934.543 | 3 | 123.90 | 0.1158 | 23.99 | 82.73 | 4 | 0.0 | 0 | R.WQQGNVFSCSVMHEALHNHYTQK.S | Carbamidomethyl: 9 | 296-318 | CID |

#### Protein 4: Immunoglobulin lambda constant 3 OS=Homo sapiens GN=IGLC3 PE=1 SV=1

**Accession:** IGLC3_HUMAN
**Database:** SwissProt
**MW [kDa] / pI:** 11.3 / 6.9
**Modification(s):** Carbamidomethyl

**Score:** 627.0
**Seq. Coverage [%]:** 59.4
**No. of Peptides:** 7

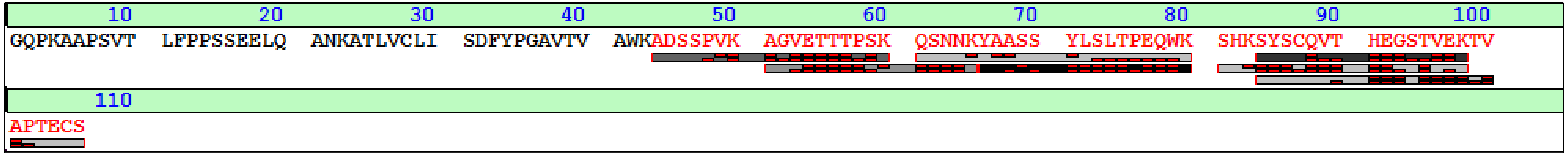


MS/MS Peptide Matches

| Cmpd. | m/z meas. | z | Δ m/z [ppm] | Δ m/z [Da] | Rt [min] | Score | No. of Cmpds. | Site [%] | P | Sequence | Modification | Range | Type |
|---|---|---|---|---|---|---|---|---|---|---|---|---|---|
| 180 | 838.027 | 2 | 114.72 | 0.0961 | 15.05 | 76.47 | 5 | 0.0 | 1 | K.ADSSPVKAGVETTTPSK.Q | | 44-60 | CID |
| 22 | 781.531 | 2 | 174.74 | 0.1365 | 10.74 | 76.54 | 15 | 0.0 | 1 | K.AGVETTTPSKQSNNK.Y | | 51-65 | CID |
| 761 | 772.49 | 3 | 140.34 | 0.1084 | 26.19 | 72.4 | 5 | 0.0 | 1 | K.QSNNKYAASSYLSLTPEQWK.S | | 61-80 | CID |

| Cmpd. | m/z meas. | z | Δ m/z [ppm] | Δ m/z [Da] | Rt [min] | Score | No. of Cmpds. | Site [%] | P | Sequence | Modification | Range | Type |
|---|---|---|---|---|---|---|---|---|---|---|---|---|---|
| 863 | 872.537 | 2 | 119.15 | 0.1039 | 27.89 | 87.34 | 9 | 0.0 | 0 | K.YAASSYLSLTPEQWK.S | | 66-80 | CID |
| 82 | 1032.619 | 2 | 138.32 | 0.1428 | 12.35 | 91.77 | 8 | 0.0 | 1 | K.SHKSYSCQVTHEGSTVEK.T | Carbamidomethyl: 7 | 81-98 | CID |
| 145 | 856.543 | 2 | 186.56 | 0.1598 | 14.21 | 114.36 | 16 | 0.0 | 0 | K.SYSCQVTHEGSTVEK.T | Carbamidomethyl: 4 | 84-98 | CID |
| 361 | 1279.557 | 2 | 386.46 | 0.4943 | 18.90 | 108.08 | 9 | 0.0 | 1 | K.SYSCQVTHEGSTVEKTVAPTECS.- | Carbamidomethyl: 4, 22 | 84-106 | CID |

#### Protein 5: Immunoglobulin lambda-like polypeptide 5 OS=Homo sapiens GN=IGLL5 PE=2 SV=2

**Accession:** IGLL5_HUMAN
**Database:** SwissProt
**MW [kDa] / pI:** 23.0 / 9.1
**Modification(s):** Carbamidomethyl
**Score:** 478.7
**Seq. Coverage [%]:** 29.4
**No. of Peptides:** 6

```
        10         20         30         40         50         60         70         80         90        100
MRPKTGQVGC ETPEELGPGP RQRWPLLLLG LAMVAHGLLR PMVAPQSGDP DPGASVGSSR SSLRSLWGRL LLQPSPQRAD PRCWPRGFWS EPQSLCYVFG
       110        120        130        140        150        160        170        180        190        200
TGTKVTVLGQ PKANPTVTLF PPSSEELQAN KATLVCLISD FYPGAVTVAW KADGSPVKAG VETTKPSKQS NNKYAASSYL SLTPEQWKSH RSYSCQVTHE
       210        220
GSTVEKTVAP TECS
```

MS/MS Peptide Matches

| Cmpd. | m/z meas. | z | Δ m/z [ppm] | Δ m/z [Da] | Rt [min] | Score | No. of Cmpds. | Site [%] | P | Sequence | Modification | Range | Type |
|---|---|---|---|---|---|---|---|---|---|---|---|---|---|
| 112 | 836.543 | 2 | 112.10 | 0.0938 | 13.40 | 68.25 | 3 | 0.0 | 1 | K.ADGSPVKAGVETTKPSK.Q | | 152-168 | CID |
| 761 | 772.49 | 3 | 140.34 | 0.1084 | 26.19 | 72.4 | 5 | 0.0 | 1 | K.QSNNKYAASSYLSLTPEQWK.S | | 169-188 | CID |
| 863 | 872.537 | 2 | 119.15 | 0.1039 | 27.89 | 87.34 | 9 | 0.0 | 0 | K.YAASSYLSLTPEQWK.S | | 174-188 | CID |
| 91 | 1046.572 | 2 | 88.62 | 0.0927 | 12.56 | 28.27 | 2 | 0.0 | 1 | K.SHRSYSCQVTHEGSTVEK.T | Carbamidomethyl: 7 | 189-206 | CID |

| Cmpd. | m/z meas. | z | Δ m/z [ppm] | Δ m/z [Da] | Rt [min] | Score | No. of Cmpds. | Site [%] | P | Sequence | Modification | Range | Type |
|---|---|---|---|---|---|---|---|---|---|---|---|---|---|
| 145 | 856.543 | 2 | 186.56 | 0.1598 | 14.21 | 114.36 | 16 | 0.0 | 0 | R.SYSCQVTHEGSTVEK.T | Carbamidomethyl: 4 | 192-206 | CID |
| 361 | 1279.557 | 2 | 386.46 | 0.4943 | 18.90 | 108.08 | 9 | 0.0 | 1 | R.SYSCQVTHEGSTVEKTVAPTECS.- | Carbamidomethyl: 4, 22 | 192-214 | CID |

#### Protein 6: Immunoglobulin heavy constant gamma 4 OS=Homo sapiens GN=IGHG4 PE=1 SV=1

**Accession:** IGHG4_HUMAN
**Database:** SwissProt
**MW [kDa] / pI:** 35.9 / 7.2
**Modification(s):** Carbamidomethyl

**Score:** 328.4
**Seq. Coverage [%]:** 32.1
**No. of Peptides:** 5

| 10 | 20 | 30 | 40 | 50 | 60 | 70 | 80 | 90 | 100 |
|---|---|---|---|---|---|---|---|---|---|
| ASTKGPSVFP | LAPCSRSTSE | STAALGCLVK | DYFPEPVTVS | WNSGALTSGV | HTFPAVLQSS | GLYSLSSVVT | VPSSSLGTKT | YTCNVDHKPS | NTKVDKRVES |
| 110 | 120 | 130 | 140 | 150 | 160 | 170 | 180 | 190 | 200 |
| KYGPPCPSCP | APEFLGGPSV | FLFPPKPKDT | LMISRTPEVT | CVVVDVSQED | PEVQFNWYVD | GVEVHNAKTK | PREEQFNSTY | RVVSVLTVLH | QDWLNGKEYK |
| 210 | 220 | 230 | 240 | 250 | 260 | 270 | 280 | 290 | 300 |
| CKVSNKGLPS | SIEKTISKAK | GQPREPQVYT | LPPSQEEMTK | NQVSLTCLVK | GFYPSDIAVE | WESNGQPENN | YKTTPPVLDS | DGSFFLYSRL | TVDKSRWQEG |
| 310 | 320 | 330 | | | | | | | |
| NVFSCSVMHE | ALHNHYTQKS | LSLSLGK | | | | | | | |

MS/MS Peptide Matches

| Cmpd. | m/z meas. | z | Δ m/z [ppm] | Δ m/z [Da] | Rt [min] | Score | No. of Cmpds. | Site [%] | P | Sequence | Modification | Range | Type |
|---|---|---|---|---|---|---|---|---|---|---|---|---|---|
| 554 | 712.479 | 2 | 169.16 | 0.1205 | 22.61 | 81.7 | 21 | 0.0 | 0 | R.STSESTAALGCLVK.D | Carbamidomethyl: 11 | 17-30 | CID |
| 44 | 833.02 | 2 | 157.90 | 0.1315 | 11.37 | 40.39 | 6 | 0.0 | 0 | K.TYTCNVDHKPSNTK.V | Carbamidomethyl: 4 | 80-93 | CID |
| 75 | 1004.46 | 2 | 474.46 | 0.4764 | 12.15 | 19.24 | 1 | 0.0 | 1 | K.TYTCNVDHKPSNTKVDK.R | Carbamidomethyl: 4 | 80-96 | CID |
| 1168 | 981.9 | 3 | 77.49 | 0.0761 | 32.97 | 53.0 | 1 | 0.0 | 0 | K.YGPPCPSCPAPEFLGGPSVFLFPPKPK.D | Carbamidomethyl: 5, 8 | 102-128 | CID |
| 893 | 1273.143 | 2 | 450.80 | 0.5737 | 28.43 | 86.06 | 5 | 0.0 | 0 | K.GFYPSDIAVEWESNGQPENNYK.T | | 251-272 | CID |

#### Protein 7: Immunoglobulin heavy constant gamma 1 OS=Homo sapiens GN=IGHG1 PE=1 SV=1

**Accession:** IGHG1_HUMAN
**Database:** SwissProt
**MW [kDa] / pI:** 36.1 / 8.5
**Modification(s):** Carbamidomethyl

**Score:** 322.5
**Seq. Coverage [%]:** 29.4
**No. of Peptides:** 5

| 10 | 20 | 30 | 40 | 50 | 60 | 70 | 80 | 90 | 100 |
|---|---|---|---|---|---|---|---|---|---|
| ASTKGPSVFP | LAPSSKSTSG | GTAALGCLVK | DYFPEPVTVS | WNSGALTSGV | HTFPAVLQSS | GLYSLSSVVT | VPSSSLGTQT | YICNVNHKPS | NTKVDKKVEP |
| **110** | **120** | **130** | **140** | **150** | **160** | **170** | **180** | **190** | **200** |
| KSCDKTHTCP | PCPAPELLGG | PSVFLFPPKP | KDTLMISRTP | EVTCVVVDVS | HEDPEVKFNW | YVDGVEVHNA | KTKPREEQYN | STYRVVSVLT | VLHQDWLNGK |
| **210** | **220** | **230** | **240** | **250** | **260** | **270** | **280** | **290** | **300** |
| EYKCKVSNKA | LPAPIEKTIS | KAKGQPREPQ | VYTLPPSRDE | LTKNQVSLTC | LVKGFYPSDI | AVEWESNGQP | ENNYKTTPPV | LDSDGSFFLY | SKLTVDKSRW |
| **310** | **320** | **330** | **340** | | | | | | |
| QQGNVFSCSV | MHEALHNHYT | QKSLSLSPGK | | | | | | | |

MS/MS Peptide Matches

| Cmpd. | m/z meas. | z | Δ m/z [ppm] | Δ m/z [Da] | Rt [min] | Score | No. of Cmpds. | Site [%] | P | Sequence | Modification | Range | Type |
|---|---|---|---|---|---|---|---|---|---|---|---|---|---|
| 1063 | 950.567 | 4 | 377.45 | 0.3587 | 31.29 | 54.13 | 3 | 0.0 | 1 | R.TPEVTCVVVDVSHEDPEVKFNWYVDGVEVHNA K.T | Carbamidomethyl: 6 | 139-171 | CID |
| 893 | 1273.143 | 2 | 450.80 | 0.5737 | 28.43 | 86.06 | 5 | 0.0 | 0 | K.GFYPSDIAVEWESNGQPENNYK.T | | 254-275 | CID |
| 968 | 937.521 | 2 | 60.22 | 0.0565 | 29.66 | 49.93 | 2 | 0.0 | 0 | K.TTPPVLDSDGSFFLYSK.L | | 276-292 | CID |
| 580 | 762.182 | 4 | 428.53 | 0.3265 | 23.14 | 49.62 | 3 | 0.0 | 1 | K.SRWQQGNVFSCSVMHEALHNHYTQK.S | Carbamidomethyl: 11 | 298-322 | CID |
| 629 | 934.543 | 3 | 123.90 | 0.1158 | 23.99 | 82.73 | 4 | 0.0 | 0 | R.WQQGNVFSCSVMHEALHNHYTQK.S | Carbamidomethyl: 9 | 300-322 | CID |

#### Protein 8: Keratin, type I cytoskeletal 9 OS=Homo sapiens GN=KRT9 PE=1 SV=3

| | | | |
|---|---|---|---|
| **Accession:** | K1C9_HUMAN | **Score:** | 249.9 |
| **Database:** | SwissProt | **Seq. Coverage [%]:** | 14.3 |
| **MW [kDa] / pI:** | 62.0 / 5.1 | **No. of Peptides:** | 2 |

| 10 | 20 | 30 | 40 | 50 | 60 | 70 | 80 | 90 | 100 |
|---|---|---|---|---|---|---|---|---|---|
| MSCRQFSSSY | LSRSGGGGGG | GLGSGGSIRS | SYSRFSSSGG | GGGGGRFSSS | SGYGGGSSRV | CGRGGGGSFG | YSYGGGSGGG | FSASSLGGGF | GGGSRGFGGA |
| 110 | 120 | 130 | 140 | 150 | 160 | 170 | 180 | 190 | 200 |
| SGGGYSSSGG | FGGGFGGGSG | GGFGGGYGSG | FGGFGGFGGG | AGGGDGGILT | ANEKSTMQEL | NSRLASYLDK | VQALEEANND | LENKIQDWYD | KKGPAAIQKN |
| 210 | 220 | 230 | 240 | 250 | 260 | 270 | 280 | 290 | 300 |
| YSPYYNTIDD | LKDQIVDLTV | GNNKTLLDID | NTRMTLDDFR | IKFEMEQNLR | QGVDADINGL | RQVLDNLTME | KSDLEMQYET | LQEELMALKK | NHKEEMSQLT |
| 310 | 320 | 330 | 340 | 350 | 360 | 370 | 380 | 390 | 400 |
| GQNSGDVNVE | INVAPGKDLT | KTLNDMRQEY | EQLIAKNRKD | IENQYETQIT | QIEHEVSSSG | QEVQSSAKEV | TQLRHGVQEL | EIELQSQLSK | KAALEKSLED |
| 410 | 420 | 430 | 440 | 450 | 460 | 470 | 480 | 490 | 500 |
| TKNRYCGQLQ | MIQEQISNLE | AQITDVRQEI | ECQNQEYSLL | LSIKMRLEKE | IETYHNLLEG | GQEDFESSGA | GKIGLGGRGG | SGGSYGRGSR | GGSGGSYGGG |
| 510 | 520 | 530 | 540 | 550 | 560 | 570 | 580 | 590 | 600 |
| GSGGGYGGGS | GSRGGSGGSY | GGGSGSGGGS | GGGYGGGSGG | GHSGGSGGGH | SGGSGGNYGG | GSGSGGGSGG | GYGGGSGSRG | GSGGSHGGGS | GFGGESGGSY |
| 610 | 620 | 630 | | | | | | | |
| GGGEEASGSG | GGYGGGSGKS | SHS | | | | | | | |

##### MS/MS Peptide Matches

| Cmpd. | m/z meas. | z | Δ m/z [ppm] | Δ m/z [Da] | Rt [min] | Score | No. of Cmpds. | Site [%] | P | Sequence | Modification | Range | Type |
|---|---|---|---|---|---|---|---|---|---|---|---|---|---|
| 74 | 896.476 | 2 | 121.05 | 0.1085 | 12.14 | 209.72 | 5 | 0.0 | 0 | R.GGSGGSYGGGGSGGGYGGGSGSR.G | | 491-513 | CID |
| 105 | 1244.219 | 4 | 184.44 | 0.2294 | 13.09 | 40.13 | 1 | 0.0 | 0 | R.GGSGGSYGGGSGSGGGSGGGYGGGSGGGH SGGSGGGHSGGSGGNYGGGSGSGGGSGGGYG GGSGSR.G | | 514-579 | CID |

#### Protein 9: Keratin, type II cytoskeletal 1 OS=Homo sapiens GN=KRT1 PE=1 SV=6

| | | | |
|---|---|---|---|
| **Accession:** | K2C1_HUMAN | **Score:** | 199.8 |
| **Database:** | SwissProt | **Seq. Coverage [%]:** | 4.8 |
| **MW [kDa] / pI:** | 66.0 / 8.2 | **No. of Peptides:** | 1 |

| 10 | 20 | 30 | 40 | 50 | 60 | 70 | 80 | 90 | 100 |
|---|---|---|---|---|---|---|---|---|---|
| MSRQFSSRSG | YRSGGGFSSG | SAGIINYQRR | TTSSSTRRSG | GGGGRFSSCG | GGGGSFGAGG | GFGSRSLVNL | GGSKSISISV | ARGGGRGSGF | GGGYGGGGFG |
| 110 | 120 | 130 | 140 | 150 | 160 | 170 | 180 | 190 | 200 |
| GGGFGGGGFG | GGGIGGGGFG | GFGSGGGGFG | GGGFGGGGYG | GGYGPVCPPG | GIQEVTINQS | LLQPLNVEID | PEIQKVKSRE | REQIKSLNNQ | FASFIDKVRF |
| 210 | 220 | 230 | 240 | 250 | 260 | 270 | 280 | 290 | 300 |
| LEQQNQVLQT | KWELLQQVDT | STRTHNLEPY | FESFINNLRR | RVDQLKSDQS | RLDSELKNMQ | DMVEDYRNKY | EDEINKRTNA | ENEFVTIKKD | VDGAYMTKVD |
| 310 | 320 | 330 | 340 | 350 | 360 | 370 | 380 | 390 | 400 |
| LQAKLDNLQQ | EIDFLTALYQ | AELSQMQTQI | SETNVILSMD | NNRSLDLDSI | IAEVKAQYED | IAQKSKAEAE | SLYQSKYEEL | QITAGRHGDS | VRNSKIEISE |
| 410 | 420 | 430 | 440 | 450 | 460 | 470 | 480 | 490 | 500 |
| LNRVIQRLRS | EIDNVKKQIS | NLQQSISDAE | QRGENALKDA | KNKLNDLEDA | LQQAKEDLAR | LLRDYQELMN | TKLALDLEIA | TYRTLLEGEE | SRMSGECAPN |
| 510 | 520 | 530 | 540 | 550 | 560 | 570 | 580 | 590 | 600 |
| VSVSVSTSHT | TISGGGSRGG | GGGGYGSGGS | SYGSGGGSYG | SGGGGGGGRG | SYGSGGSSYG | SGGGSYGSGG | GGGGHGSYGS | GSSSGGYRGG | SGGGGGGSSG |
| 610 | 620 | 630 | 640 | 650 | | | | | |
| GRGSGGGSSG | GSIGGRGSSS | GGVKSSGGSS | SVKFVSTTYS | GVTR | | | | | |

MS/MS Peptide Matches

| Cmpd. | m/z meas. | z | Δ m/z [ppm] | Δ m/z [Da] | Rt [min] | Score | No. of Cmpds. | Site [%] | P | Sequence | Modification | Range | Type |
|---|---|---|---|---|---|---|---|---|---|---|---|---|---|
| 152 | 1192.525 | 2 | 38.10 | 0.0454 | 14.38 | 199.78 | 5 | 0.0 | 0 | R.GGGGGGYGSGGSSYGSGGGSYGSGGGGGG GR.G | | 519-549 | CID |

##### Protein 10: Keratin, type II cytoskeletal 2 epidermal OS=Homo sapiens GN=KRT2 PE=1 SV=2

| | | | |
|---|---|---|---|
| **Accession:** | K22E_HUMAN | **Score:** | 165.0 |
| **Database:** | SwissProt | **Seq. Coverage [%]:** | 3.6 |
| **MW [kDa] / pI:** | 65.4 / 8.1 | **No. of Peptides:** | 1 |

| 10 | 20 | 30 | 40 | 50 | 60 | 70 | 80 | 90 | 100 |
|---|---|---|---|---|---|---|---|---|---|
| MSCQISCKSR | GRGGGGGGFR | GFSSGSAVVS | GGSRRSTSSF | SCLSRHGGGG | GGFGGGGFGS | RSLVGLGGTK | SISISVAGGG | GGFGAAGGFG | GRGGGFGGGS |
| 110 | 120 | 130 | 140 | 150 | 160 | 170 | 180 | 190 | 200 |
| SFGGGSGFSG | GGFGGGGFGG | GRFGGFGGPG | GVGGLGGPGG | FGPGGYPGGI | HEVSVNQSLL | QPLNVKVDPE | IQNVKAQERE | QIKTLNNKFA | SFIDKVRFLE |
| 210 | 220 | 230 | 240 | 250 | 260 | 270 | 280 | 290 | 300 |
| QQNQVLQTKW | ELLQQMNVGT | RPINLEPIFQ | GYIDSLKRYL | DGLTAERTSQ | NSELNNMQDL | VEDYKKKYED | EINKRTAAEN | DFVTLKKDVD | NAYMIKVELQ |
| 310 | 320 | 330 | 340 | 350 | 360 | 370 | 380 | 390 | 400 |
| SKVDLLNQEI | EFLKVLYDAE | ISQIHQSVTD | TNVILSMDNS | RNLDLDSIIA | EVKAQYEEIA | QRSKEEAEAL | YHSKYEELQV | TVGRHGDSLK | EIKIEISELN |
| 410 | 420 | 430 | 440 | 450 | 460 | 470 | 480 | 490 | 500 |
| RVIQRLQGEI | AHVKKQCKNV | QDAIADAEQR | GEHALKDARN | KLNDLEEALQ | QAKEDLARLL | RDYQELMNVK | LALDVEIATY | RKLLEGEECR | MSGDLSSNVT |
| 510 | 520 | 530 | 540 | 550 | 560 | 570 | 580 | 590 | 600 |
| VSVTSSTISS | NVASKAAFGG | SGGRGSSSGG | GYSSGSSSYG | SGGRQSGSRG | GSGGGGSISG | GGYGSGGGSG | GRYGSGGGSK | GGSISGGGYG | SGGGKHSSGG |
| 610 | 620 | 630 | 640 | | | | | | |
| GSRGGSSSGG | GYGSGGGGSS | SVKGSSGEAF | GSSVTFSFR | | | | | | |

MS/MS Peptide Matches

| Cmpd. | m/z meas. | z | Δ m/z [ppm] | Δ m/z [Da] | Rt [min] | Score | No. of Cmpds. | Site [%] | P | Sequence | Modification | Range | Type |
|---|---|---|---|---|---|---|---|---|---|---|---|---|---|
| 106 | 871.495 | 2 | 134.43 | 0.1171 | 13.10 | 164.96 | 3 | 0.0 | 0 | R.GGSGGGGSISGGGYGSGGGSGGR.Y | | 550-572 | CID |

#### Protein 11: Keratin, type I cytoskeletal 10 OS=Homo sapiens GN=KRT10 PE=1 SV=6

| | | | |
|---|---|---|---|
| **Accession:** | K1C10_HUMAN | **Score:** | 135.0 |
| **Database:** | SwissProt | **Seq. Coverage [%]:** | 3.3 |
| **MW [kDa] / pI:** | 58.8 / 5.1 | **No. of Peptides:** | 1 |

| 10 | 20 | 30 | 40 | 50 | 60 | 70 | 80 | 90 | 100 |
|---|---|---|---|---|---|---|---|---|---|
| MSVRYSSSKH | YSSSRSGGGG | GGGGCGGGGG | VSSLRISSSK | GSLGGGFSSG | GFSGGSFSRG | SSGGGCFGGS | SGGYGGLGGF | GGGSFRGSYG | SSSFGGSYGG |
| **110** | **120** | **130** | **140** | **150** | **160** | **170** | **180** | **190** | **200** |
| IFGGGSFGGG | SFGGGSFGGG | GFGGGGFGGG | FGGGFGGDGG | LLSGNEKVTM | QNLNDRLASY | LDKVRALEES | NYELEGKIKE | WYEKHGNSHQ | GEPRDYSKYY |
| **210** | **220** | **230** | **240** | **250** | **260** | **270** | **280** | **290** | **300** |
| KTIDDLKNQI | LNLTTDNANI | LLQIDNARLA | ADDFRLKYEN | EVALRQSVEA | DINGLRRVLD | ELTLTKADLE | MQIESLTEEL | AYLKKNHEEE | MKDLRNVSTG |
| **310** | **320** | **330** | **340** | **350** | **360** | **370** | **380** | **390** | **400** |
| DVNVEMNAAP | GVDLTQLLNN | MRSQYEQLAE | QNRKDAEAWF | NEKSKELTTE | IDNNIEQISS | YKSEITELRR | NVQALEIELQ | SQLALKQSLE | ASLAETEGRY |
| **410** | **420** | **430** | **440** | **450** | **460** | **470** | **480** | **490** | **500** |
| CVQLSQIQAQ | ISALEEQLQQ | IRAETECQNT | EYQQLLDIKI | RLENEIQTYR | SLLEGEGSSG | GGGRGGGSFG | GGYGGGSSGG | GSSGGGHGGG | HGGSSGGGYG |
| **510** | **520** | **530** | **540** | **550** | **560** | **570** | **580** | **590** | |
| GGSSGGGSSG | GGYGGGSSSG | GHGGSSSGGY | GGGSSGGGGG | GYGGGSSGGG | SSSGGGYGGG | SSSGGHKSSS | SGSVGESSSK | GPRY | |

MS/MS Peptide Matches

| Cmpd. | m/z meas. | z | Δ m/z [ppm] | Δ m/z [Da] | Rt [min] | Score | No. of Cmpds. | Site [%] | P | Sequence | Modification | Range | Type |
|---|---|---|---|---|---|---|---|---|---|---|---|---|---|
| 623 | 854.477 | 2 | 102.17 | 0.0873 | 23.89 | 135.04 | 3 | 0.0 | 0 | K.GSLGGGFSSGGFSGGSFSR.G | | 41-59 | CID |

#### Protein 12: Immunoglobulin heavy variable 4-4 OS=Homo sapiens GN=IGHV4-4 PE=3 SV=2

| | | | |
|---|---|---|---|
| **Accession:** | HV404_HUMAN | **Score:** | 94.4 |
| **Database:** | SwissProt | **Seq. Coverage [%]:** | 13.7 |
| **MW [kDa] / pI:** | 12.8 / 9.6 | **No. of Peptides:** | 1 |

**Modification(s):** Carbamidomethyl

| 10 | 20 | 30 | 40 | 50 | 60 | 70 | 80 | 90 | 100 |
|---|---|---|---|---|---|---|---|---|---|
| MKHLWFFLLL | VAAPRWVLSQ | VQLQESGPGL | VKPSGTLSLT | CAVSGGSISS | SNWWSWVRQP | PGKGLEWIGE | IYHSGSTNYN | PSLKSRVTIS | VDKSKNQFSL |
| **110** | **120** | | | | | | | | |
| KLSSVTAADT | AVYYCAR | | | | | | | | |

MS/MS Peptide Matches

| Cmpd. | m/z meas. | z | Δ m/z [ppm] | Δ m/z [Da] | Rt [min] | Score | No. of Cmpds. | Site [%] | P | Sequence | Modification | Range | Type |
|---|---|---|---|---|---|---|---|---|---|---|---|---|---|
| 541 | 874.465 | 2 | 51.90 | 0.0454 | 22.36 | 94.43 | 3 | 0.0 | 0 | K.LSSVTAADTAVYYCAR.- | Carbamidomethyl: 14 | 102-117 | CID |

#### Protein 13: Immunoglobulin kappa variable 3-15 OS=Homo sapiens GN=IGKV3-15 PE=1 SV=2

**Accession:** KV315_HUMAN
**Database:** SwissProt
**MW [kDa] / pI:** 12.5 / 5.1

**Score:** 68.4
**Seq. Coverage [%]:** 18.3
**No. of Peptides:** 1

| 10 | 20 | 30 | 40 | 50 | 60 | 70 | 80 | 90 | 100 |
|---|---|---|---|---|---|---|---|---|---|
| MEAPAQLLFL | LLLWLPDTTG | EIVMTQSPAT | LSVSPGERAT | LSCRASQSVS | SNLAWYQQKP | GQAPRLLIYG | ASTRATGIPA | RFSGSGSGTE | FTLTISSLQS |
| **110** | **120** | | | | | | | | |
| EDFAVYYCQQ | YNNWP | | | | | | | | |

MS/MS Peptide Matches

| Cmpd. | m/z meas. | z | Δ m/z [ppm] | Δ m/z [Da] | Rt [min] | Score | No. of Cmpds. | Site [%] | P | Sequence | Modification | Range | Type |
|---|---|---|---|---|---|---|---|---|---|---|---|---|---|
| 493 | 768.481 | 3 | 119.61 | 0.0919 | 21.45 | 68.43 | 3 | 0.0 | 0 | R.ASQSVSSNLAWYQQKPGQAPR.L | | 45-65 | CID |

##### Protein 14: Immunoglobulin lambda variable 8-61 OS=Homo sapiens GN=IGLV8-61 PE=3 SV=7

| | | | |
|---|---|---|---|
| **Accession:** | LV861_HUMAN | **Score:** | 35.5 |
| **Database:** | SwissProt | **Seq. Coverage [%]:** | 13.1 |
| **MW [kDa] / pI:** | 12.8 / 4.3 | **No. of Peptides:** | 1 |

```
        10         20         30         40         50         60         70         80         90        100
MSVPTMAWMM LLLGLLAYGS GVDSQTVVTQ EPSFSVSPGG TVTLTCGLSS GSVSTSYYPS WYQQTPGQAP RTLIYSTNTR SSGVPDRFSG SILGNKAALT
       110        120        130
ITGAQADDES DYYCVLYMGS GI
```

MS/MS Peptide Matches

| Cmpd. | m/z meas. | z | Δ m/z [ppm] | Δ m/z [Da] | Rt [min] | Score | No. of Cmpds. | Site [%] | P | Sequence | Modification | Range | Type |
|---|---|---|---|---|---|---|---|---|---|---|---|---|---|
| 539 | 811.011 | 2 | 111.43 | 0.0904 | 22.31 | 35.53 | 1 | 0.0 | 1 | R.SSGVPDRFSGSILGNK.A | | 81-96 | CID |

#### Project Info

| | | | |
|---|---|---|---|
| **Name:** | Alagesan | **Date:** | Mar 5, 2019 |
| **Note:** | Nanobooster | | |

#### Sample Info & Protocols

| | |
|---|---|
| **Name:** | Nanobooster - IgG |
| **Date:** | Mar 5, 2019 |

#### Search Result Info

| | | | |
|---|---|---|---|
| **Search Result:** | IonTrap_Human_Swiss_CID_2019-03-05 17:55:08 | | |
| **Location:** | /Alagesan/Nanobooster - IgG/KA_IgG2 HD NB ACN CID_GA3_01_9082.mgf | | |
| **Search Method:** | IonTrap_Human_Swiss_CID | **Version:** | 1.2 |
| **Search Engine(s):** | Mascot, 2.6.0 | **Ident. Compound(s):** | 0/2318 |
| **Database(s):** | SwissProt, SwissProt_2017_04.fasta | | |
| **Note:** | | | |

#### Results

##### Protein 1: Immunoglobulin kappa constant OS=Homo sapiens GN=IGKC PE=1 SV=2

**Accession:** IGKC_HUMAN
**Database:** SwissProt
**MW [kDa] / pI:** 11.8 / 6.1
**Modification(s):** Carbamidomethyl

**Score:** 1141.0
**Seq. Coverage [%]:** 88.8
**No. of Peptides:** 11

| 10 | 20 | 30 | 40 | 50 | 60 | 70 | 80 | 90 | 100 |
|---|---|---|---|---|---|---|---|---|---|
| RTVAAPSVFI | FPPSDEQLKS | GTASVVCLLN | NFYPREAKVQ | WKVDNALQSG | NSQESVTEQD | SKDSTYSLSS | TLTLSKADYE | KHKVYACEVT | HQGLSSPVTK |

| 110 |
|---|
| SFNRGEC |

MS/MS Peptide Matches

| Cmpd. | m/z meas. | z | Δ m/z [ppm] | Δ m/z [Da] | Rt [min] | Score | No. of Cmpds. | Site [%] | P | Sequence | Modification | Range | Type |
|---|---|---|---|---|---|---|---|---|---|---|---|---|---|
| 910 | 701.443 | 3 | 88.58 | 0.0621 | 28.72 | 44.86 | 3 | 0.0 | 1 | -.RTVAAPSVFIFPPSDEQLK.S | | 1-19 | CID |
| 1040 | 973.615 | 2 | 100.55 | 0.0979 | 30.89 | 86.32 | 14 | 0.0 | 0 | R.TVAAPSVFIFPPSDEQLK.S | | 2-19 | CID |
| 1494 | 1242.731 | 3 | 341.87 | 0.4247 | 38.51 | 78.75 | 18 | 0.0 | 1 | R.TVAAPSVFIFPPSDEQLKSGTASVVCLLNNFYPR.E | Carbamidomethyl: 26 | 2-35 | CID |
| 1174 | 899.555 | 2 | 115.34 | 0.1037 | 33.15 | 72.84 | 12 | 0.0 | 0 | K.SGTASVVCLLNNFYPR.E | Carbamidomethyl: 8 | 20-35 | CID |
| 1056 | 709.435 | 3 | 103.64 | 0.0735 | 31.18 | 88.54 | 4 | 0.0 | 1 | K.SGTASVVCLLNNFYPREAK.V | Carbamidomethyl: 8 | 20-38 | CID |
| 486 | 893.213 | 3 | 132.29 | 0.1182 | 21.22 | 130.48 | 9 | 0.0 | 1 | K.VQWKVDNALQSGNSQESVTEQDSK.D | | 39-62 | CID |
| 237 | 1068.643 | 2 | 145.06 | 0.1550 | 15.85 | 152.79 | 18 | 0.0 | 0 | K.VDNALQSGNSQESVTEQDSK.D | | 43-62 | CID |
| 666 | 1207.545 | 3 | 251.57 | 0.3037 | 24.56 | 134.66 | 2 | 0.0 | 1 | K.VDNALQSGNSQESVTEQDSKDSTYSLSSTLTLSK.A | | 43-76 | CID |
| 678 | 751.944 | 2 | 81.32 | 0.0611 | 24.78 | 94.17 | 4 | 0.0 | 0 | K.DSTYSLSSTLTLSK.A | | 63-76 | CID |
| 286 | 714.463 | 3 | 137.02 | 0.0979 | 16.94 | 147.82 | 9 | 0.0 | 1 | K.HKVYACEVTHQGLSSPVTK.S | Carbamidomethyl: 6 | 82-100 | CID |
| 378 | 938.707 | 2 | 255.63 | 0.2399 | 19.13 | 109.74 | 14 | 0.0 | 0 | K.VYACEVTHQGLSSPVTK.S | Carbamidomethyl: 4 | 84-100 | CID |

| Cmpd. | m/z meas. | z | Δ m/z [ppm] | Δ m/z [Da] | Rt [min] | Score | No. of Cmpds. | Site [%] | P | Sequence | Modification | Range | Type |
|---|---|---|---|---|---|---|---|---|---|---|---|---|---|

##### Protein 2: Immunoglobulin kappa light chain OS=Homo sapiens PE=1 SV=1

| | | | |
|---|---|---|---|
| **Accession:** | IGK_HUMAN | **Score:** | 1055.9 |
| **Database:** | SwissProt | **Seq. Coverage [%]:** | 43.9 |
| **MW [kDa] / pI:** | 23.4 / 6.9 | **No. of Peptides:** | 9 |
| **Modification(s):** | Carbamidomethyl | | |

```
        10         20         30         40         50         60         70         80         90        100
DIQMTQSPST LSASVGDRVT ITCRASQSIN TWLAWYQQKP GKAPKLLMYK ASSLESGVPS RFIGSGSGTE FTLTISSLQP DDFATYYCQQ YNSDSKMFGQ
       110        120        130        140        150        160        170        180        190        200
GTKVEVKGTV AAPSVFIFPP SDEQLKSGTA SVVCLLNNFY PREAKVQWKV DNALQSGNSQ ESVTEQDSKD STYSLSSTLT LSKADYEKHK VYACEVTHQG
       210        220
LSSPVTKSFN RGEC
```

**MS/MS Peptide Matches**

| Cmpd. | m/z meas. | z | Δ m/z [ppm] | Δ m/z [Da] | Rt [min] | Score | No. of Cmpds. | Site [%] | P | Sequence | Modification | Range | Type |
|---|---|---|---|---|---|---|---|---|---|---|---|---|---|
| 537 | 947.003 | 2 | 51.16 | 0.0484 | 22.21 | 124.88 | 3 | 0.0 | 0 | -.DIQMTQSPSTLSASVGDR.V | | 1-18 | CID |
| 1174 | 899.555 | 2 | 115.34 | 0.1037 | 33.15 | 72.84 | 12 | 0.0 | 0 | K.SGTASVVCLLNNFYPR.E | Carbamidomethyl: 8 | 127-142 | CID |
| 1056 | 709.435 | 3 | 103.64 | 0.0735 | 31.18 | 88.54 | 4 | 0.0 | 1 | K.SGTASVVCLLNNFYPREAK.V | Carbamidomethyl: 8 | 127-145 | CID |
| 486 | 893.213 | 3 | 132.29 | 0.1182 | 21.22 | 130.48 | 9 | 0.0 | 1 | K.VQWKVDNALQSGNSQESVTEQDSK.D | | 146-169 | CID |
| 237 | 1068.643 | 2 | 145.06 | 0.1550 | 15.85 | 152.79 | 18 | 0.0 | 0 | K.VDNALQSGNSQESVTEQDSK.D | | 150-169 | CID |
| 666 | 1207.545 | 3 | 251.57 | 0.3037 | 24.56 | 134.66 | 2 | 0.0 | 1 | K.VDNALQSGNSQESVTEQDSKDSTYSLSSTLTLSK.A | | 150-183 | CID |
| 678 | 751.944 | 2 | 81.32 | 0.0611 | 24.78 | 94.17 | 4 | 0.0 | 0 | K.DSTYSLSSTLTLSK.A | | 170-183 | CID |
| 286 | 714.463 | 3 | 137.02 | 0.0979 | 16.94 | 147.82 | 9 | 0.0 | 1 | K.HKVYACEVTHQGLSSPVTK.S | Carbamidomethyl: 6 | 189-207 | CID |

| Cmpd. | m/z meas. | z | Δ m/z [ppm] | Δ m/z [Da] | Rt [min] | Score | No. of Cmpds. | Site [%] | P | Sequence | Modification | Range | Type |
|---|---|---|---|---|---|---|---|---|---|---|---|---|---|
| 378 | 938.707 | 2 | 255.63 | 0.2399 | 19.13 | 109.74 | 14 | 0.0 | 0 | K.VYACEVTHQGLSSPVTK.S | Carbamidomethyl: 4 | 191-207 | CID |

#### Protein 3: Immunoglobulin heavy constant gamma 2 OS=Homo sapiens GN=IGHG2 PE=1 SV=2

**Accession:** IGHG2_HUMAN
**Database:** SwissProt
**MW [kDa] / pI:** 35.9 / 7.7
**Modification(s):** Carbamidomethyl, Oxidation

**Score:** 832.5
**Seq. Coverage [%]:** 55.2
**No. of Peptides:** 14

```
        10         20         30         40         50         60         70         80         90        100
ASTKGPSVFP LAPCSRSTSE STAALGCLVK DYFPEPVTVS WNSGALTSGV HTFPAVLQSS GLYSLSSVVT VPSSNFGTQT YTCNVDHKPS NTKVDKTVER
       110        120        130        140        150        160        170        180        190        200
KCCVECPPCP APPVAGPSVF LFPPKPKDTL MISRTPEVTC VVVDVSHEDP EVQFNWYVDG VEVHNAKTKP REEQFNSTFR VVSVLTVVHQ DWLNGKEYKC
       210        220        230        240        250        260        270        280        290        300
KVSNKGLPAP IEKTISKTKG QPREPQVYTL PPSREEMTKN QVSLTCLVKG FYPSDISVEW ESNGQPENNY KTTPPMLDSD GSFFLYSKLT VDKSRWQQGN
       310        320        330
VFSCSVMHEA LHNHYTQKSL SLSPGK
```

**MS/MS Peptide Matches**

| Cmpd. | m/z meas. | z | Δ m/z [ppm] | Δ m/z [Da] | Rt [min] | Score | No. of Cmpds. | Site [%] | P | Sequence | Modification | Range | Type |
|---|---|---|---|---|---|---|---|---|---|---|---|---|---|
| 551 | 712.453 | 2 | 132.66 | 0.0945 | 22.48 | 95.72 | 19 | 0.0 | 0 | R.STSESTAALGCLVK.D | Carbamidomethyl: 11 | 17-30 | CID |
| 903 | 1013.224 | 3 | 382.03 | 0.3869 | 28.58 | 76.94 | 7 | 0.0 | 1 | R.KCCVECPPCPAPPVAGPSVFLFPPKPK.D | Carbamidomethyl: 2, 3, 6, 9 | 101-127 | CID |
| 1012 | 970.574 | 3 | 448.65 | 0.4353 | 30.39 | 73.88 | 14 | 0.0 | 0 | K.CCVECPPCPAPPVAGPSVFLFPPKPK.D | Carbamidomethyl: 1, 2, 5, 8 | 102-127 | CID |

| Cmpd. | m/z meas. | z | Δ m/z [ppm] | Δ m/z [Da] | Rt [min] | Score | No. of Cmpds. | Site [%] | P | Sequence | Modification | Range | Type |
|---|---|---|---|---|---|---|---|---|---|---|---|---|---|
| 1039 | 1242.605 | 3 | 263.60 | 0.3275 | 30.87 | 41.21 | 5 | 0.0 | 1 | K.CCVECPPCPAPPVAGPSVFLFPPKPKDTLMIS R.T | Carbamidomethyl: 1, 2, 5, 8 | 102-134 | CID |
| 1059 | 950.553 | 4 | 372.29 | 0.3537 | 31.22 | 52.25 | 3 | 0.0 | 0 | R.TPEVTCVVVDVSHEDPEVQFNWYVDGVEVHNA K.T | Carbamidomethyl: 6 | 135-167 | CID |
| 902 | 738.775 | 3 | 53.57 | 0.0396 | 28.57 | 77.02 | 3 | 0.0 | 1 | R.VVSVLTVVHQDWLNGKEYK.C | | 181-199 | CID |
| 448 | 953.075 | 2 | 105.19 | 0.1002 | 20.51 | 62.22 | 8 | 0.0 | 1 | R.EPQVYTLPPSREEMTK.N | | 224-239 | CID |
| 406 | 961.531 | 2 | 581.48 | 0.5588 | 19.67 | 17.05 | 1 | 0.0 | 1 | R.EPQVYTLPPSREEMTK.N | Oxidation: 14 | 224-239 | CID |
| 835 | 1280.998 | 2 | 336.74 | 0.4312 | 27.40 | 68.57 | 2 | 0.0 | 0 | K.GFYPSDISVEWESNGQPENNYK.T | | 250-271 | CID |
| 997 | 953.547 | 2 | 101.12 | 0.0964 | 30.11 | 79.06 | 6 | 0.0 | 0 | K.TTPPMLDSDGSFFLYSK.L | | 272-288 | CID |
| 876 | 961.499 | 2 | 53.00 | 0.0510 | 28.11 | 65.06 | 6 | 0.0 | 0 | K.TTPPMLDSDGSFFLYSK.L | Oxidation: 5 | 272-288 | CID |
| 586 | 762.193 | 4 | 442.97 | 0.3375 | 23.19 | 55.91 | 3 | 0.0 | 1 | K.SRWQQGNVFSCSVMHEALHNHYTQK.S | Carbamidomethyl: 11 | 294-318 | CID |
| 632 | 934.875 | 3 | 479.20 | 0.4478 | 24.00 | 39.36 | 3 | 0.0 | 0 | R.WQQGNVFSCSVMHEALHNHYTQK.S | Carbamidomethyl: 9 | 296-318 | CID |
| 479 | 939.834 | 3 | 79.96 | 0.0751 | 21.05 | 28.27 | 1 | 0.0 | 0 | R.WQQGNVFSCSVMHEALHNHYTQK.S | Carbamidomethyl: 9; Oxidation: 12 | 296-318 | CID |

#### Protein 4: Immunoglobulin lambda constant 3 OS=Homo sapiens GN=IGLC3 PE=1 SV=1

**Accession:** IGLC3_HUMAN
**Database:** SwissProt
**MW [kDa] / pI:** 11.3 / 6.9
**Modification(s):** Carbamidomethyl

**Score:** 591.4
**Seq. Coverage [%]:** 59.4
**No. of Peptides:** 7

10 20 30 40 50 60 70 80 90 100
GQPKAAPSVT LFPPSSEELQ ANKATLVCLI SDFYPGAVTV AWKADSSPVK AGVETTTPSK QSNNKYAASS YLSLTPEQWK SHKSYSCQVT HEGSTVEKTV

110
APTECS

MS/MS Peptide Matches

| Cmpd. | m/z meas. | z | Δ m/z [ppm] | Δ m/z [Da] | Rt [min] | Score | No. of Cmpds. | Site [%] | P | Sequence | Modification | Range | Type |
|---|---|---|---|---|---|---|---|---|---|---|---|---|---|
| 193 | 838.038 | 2 | 127.85 | 0.1071 | 15.00 | 65.37 | 5 | 0.0 | 1 | K.ADSSPVKAGVETTTPSK.Q | | 44-60 | CID |
| 38 | 781.545 | 2 | 192.66 | 0.1505 | 10.75 | 75.64 | 15 | 0.0 | 1 | K.AGVETTTPSKQSNNK.Y | | 51-65 | CID |
| 759 | 772.492 | 3 | 142.93 | 0.1104 | 26.14 | 68.73 | 6 | 0.0 | 1 | K.QSNNKYAASSYLSLTPEQWK.S | | 61-80 | CID |
| 856 | 872.563 | 2 | 148.95 | 0.1299 | 27.77 | 82.57 | 8 | 0.0 | 0 | K.YAASSYLSLTPEQWK.S | | 66-80 | CID |
| 96 | 1032.598 | 2 | 117.98 | 0.1218 | 12.33 | 100.65 | 8 | 0.0 | 1 | K.SHKSYSCQVTHEGSTVEK.T | Carbamidomethyl: 7 | 81-98 | CID |
| 156 | 856.784 | 2 | 467.98 | 0.4008 | 14.12 | 108.0 | 15 | 0.0 | 0 | K.SYSCQVTHEGSTVEK.T | Carbamidomethyl: 4 | 84-98 | CID |
| 364 | 1279.568 | 2 | 395.06 | 0.5053 | 18.85 | 90.47 | 7 | 0.0 | 1 | K.SYSCQVTHEGSTVEKTVAPTECS.- | Carbamidomethyl: 4, 22 | 84-106 | CID |

#### Protein 5: Immunoglobulin lambda constant 2 OS=Homo sapiens GN=IGLC2 PE=1 SV=1

**Accession:** IGLC2_HUMAN
**Database:** SwissProt
**MW [kDa] / pI:** 11.3 / 6.9
**Modification(s):** Carbamidomethyl

**Score:** 535.3
**Seq. Coverage [%]:** 59.4
**No. of Peptides:** 7

```
         10         20         30         40         50         60         70         80         90        100
GQPKAAPSVT LFPPSSEELQ ANKATLVCLI SDFYPGAVTV AWKADSSPVK AGVETTTPSK QSNNKYAASS YLSLTPEQWK SHRSYSCQVT HEGSTVEKTV
        110
APTECS
```

MS/MS Peptide Matches

| Cmpd. | m/z meas. | z | Δ m/z [ppm] | Δ m/z [Da] | Rt [min] | Score | No. of Cmpds. | Site [%] | P | Sequence | Modification | Range | Type |
|---|---|---|---|---|---|---|---|---|---|---|---|---|---|
| 193 | 838.038 | 2 | 127.85 | 0.1071 | 15.00 | 65.37 | 5 | 0.0 | 1 | K.ADSSPVKAGVETTTPSK.Q | | 44-60 | CID |
| 38 | 781.545 | 2 | 192.66 | 0.1505 | 10.75 | 75.64 | 15 | 0.0 | 1 | K.AGVETTTPSKQSNNK.Y | | 51-65 | CID |
| 759 | 772.492 | 3 | 142.93 | 0.1104 | 26.14 | 68.73 | 6 | 0.0 | 1 | K.QSNNKYAASSYLSLTPEQWK.S | | 61-80 | CID |
| 856 | 872.563 | 2 | 148.95 | 0.1299 | 27.77 | 82.57 | 8 | 0.0 | 0 | K.YAASSYLSLTPEQWK.S | | 66-80 | CID |

| Cmpd. | m/z meas. | z | Δ m/z [ppm] | Δ m/z [Da] | Rt [min] | Score | No. of Cmpds. | Site [%] | P | Sequence | Modification | Range | Type |
|---|---|---|---|---|---|---|---|---|---|---|---|---|---|
| 104 | 1046.573 | 2 | 89.58 | 0.0937 | 12.52 | 44.5 | 4 | 0.0 | 1 | K.SHRSYSCQVTHEGSTVEK.T | Carbamidomethyl: 7 | 81-98 | CID |
| 156 | 856.784 | 2 | 467.98 | 0.4008 | 14.12 | 108.0 | 15 | 0.0 | 0 | R.SYSCQVTHEGSTVEK.T | Carbamidomethyl: 4 | 84-98 | CID |
| 364 | 1279.568 | 2 | 395.06 | 0.5053 | 18.85 | 90.47 | 7 | 0.0 | 1 | R.SYSCQVTHEGSTVEKTVAPTECS.- | Carbamidomethyl: 4, 22 | 84-106 | CID |

#### Protein 6: Immunoglobulin lambda-like polypeptide 5 OS=Homo sapiens GN=IGLL5 PE=2 SV=2

**Accession:** IGLL5_HUMAN
**Database:** SwissProt
**MW [kDa] / pI:** 23.0 / 9.1
**Modification(s):** Carbamidomethyl

**Score:** 460.0
**Seq. Coverage [%]:** 29.4
**No. of Peptides:** 6

| 10 | 20 | 30 | 40 | 50 | 60 | 70 | 80 | 90 | 100 |
|---|---|---|---|---|---|---|---|---|---|
| MRPKTGQVGC | ETPEELGPGP | RQRWPLLLLG | LAMVAHGLLR | PMVAPQSGDP | DPGASVGSSR | SSLRSLWGRL | LLQPSPQRAD | PRCWPRGFWS | EPQSLCYVFG |
| **110** | **120** | **130** | **140** | **150** | **160** | **170** | **180** | **190** | **200** |
| TGTKVTVLGQ | PKANPTVTLF | PPSSEELQAN | KATLVCLISD | FYPGAVTVAW | KADGSPVKAG | VETTKPSKQS | NNKYAASSYL | SLTPEQWKSH | RSYSCQVTHE |
| **210** | **220** | | | | | | | | |
| GSTVEKTVAP | TECS | | | | | | | | |

##### MS/MS Peptide Matches

| Cmpd. | m/z meas. | z | Δ m/z [ppm] | Δ m/z [Da] | Rt [min] | Score | No. of Cmpds. | Site [%] | P | Sequence | Modification | Range | Type |
|---|---|---|---|---|---|---|---|---|---|---|---|---|---|
| 128 | 836.566 | 2 | 139.60 | 0.1168 | 13.44 | 65.74 | 3 | 0.0 | 1 | K.ADGSPVKAGVETTKPSK.Q | | 152-168 | CID |
| 759 | 772.492 | 3 | 142.93 | 0.1104 | 26.14 | 68.73 | 6 | 0.0 | 1 | K.QSNNKYAASSYLSLTPEQWK.S | | 169-188 | CID |
| 856 | 872.563 | 2 | 148.95 | 0.1299 | 27.77 | 82.57 | 8 | 0.0 | 0 | K.YAASSYLSLTPEQWK.S | | 174-188 | CID |
| 104 | 1046.573 | 2 | 89.58 | 0.0937 | 12.52 | 44.5 | 4 | 0.0 | 1 | K.SHRSYSCQVTHEGSTVEK.T | Carbamidomethyl: 7 | 189-206 | CID |
| 156 | 856.784 | 2 | 467.98 | 0.4008 | 14.12 | 108.0 | 15 | 0.0 | 0 | R.SYSCQVTHEGSTVEK.T | Carbamidomethyl: 4 | 192-206 | CID |

| Cmpd. | m/z meas. | z | Δ m/z [ppm] | Δ m/z [Da] | Rt [min] | Score | No. of Cmpds. | Site [%] | P | Sequence | Modification | Range | Type |
|---|---|---|---|---|---|---|---|---|---|---|---|---|---|
| 364 | 1279.568 | 2 | 395.06 | 0.5053 | 18.85 | 90.47 | 7 | 0.0 | 1 | R.SYSCQVTHEGSTVEKTVAPTECS.- | Carbamidomethyl: 4, 22 | 192-214 | CID |

##### Protein 7: Immunoglobulin heavy constant gamma 4 OS=Homo sapiens GN=IGHG4 PE=1 SV=1

**Accession:** IGHG4_HUMAN
**Database:** SwissProt
**MW [kDa] / pI:** 35.9 / 7.2
**Modification(s):** Carbamidomethyl

**Score:** 381.4
**Seq. Coverage [%]:** 34.3
**No. of Peptides:** 5

| 10 | 20 | 30 | 40 | 50 | 60 | 70 | 80 | 90 | 100 |
|---|---|---|---|---|---|---|---|---|---|
| ASTKGPSVFP | LAPCSRSTSE | STAALGCLVK | DYFPEPVTVS | WNSGALTSGV | HTFPAVLQSS | GLYSLSSVVT | VPSSSLGTKT | YTCNVDHKPS | NTKVDKRVES |
| **110** | **120** | **130** | **140** | **150** | **160** | **170** | **180** | **190** | **200** |
| KYGPPCPSCP | APEFLGGPSV | FLFPPKPKDT | LMISRTPEVT | CVVVDVSQED | PEVQFNWYVD | GVEVHNAKTK | PREEQFNSTY | RVVSVLTVLH | QDWLNGKEYK |
| **210** | **220** | **230** | **240** | **250** | **260** | **270** | **280** | **290** | **300** |
| CKVSNKGLPS | SIEKTISKAK | GQPREPQVYT | LPPSQEEMTK | NQVSLTCLVK | GFYPSDIAVE | WESNGQPENN | YKTTPPVLDS | DGSFFLYSRL | TVDKSRWQEG |
| **310** | **320** | **330** | | | | | | | |
| NVFSCSVMHE | ALHNHYTQKS | LSLSLGK | | | | | | | |

MS/MS Peptide Matches

| Cmpd. | m/z meas. | z | Δ m/z [ppm] | Δ m/z [Da] | Rt [min] | Score | No. of Cmpds. | Site [%] | P | Sequence | Modification | Range | Type |
|---|---|---|---|---|---|---|---|---|---|---|---|---|---|
| 551 | 712.453 | 2 | 132.66 | 0.0945 | 22.48 | 95.72 | 19 | 0.0 | 0 | R.STSESTAALGCLVK.D | Carbamidomethyl: 11 | 17-30 | CID |
| 59 | 833.004 | 2 | 138.69 | 0.1155 | 11.35 | 60.58 | 6 | 0.0 | 0 | K.TYTCNVDHKPSNTK.V | Carbamidomethyl: 4 | 80-93 | CID |
| 1168 | 982.273 | 3 | 457.39 | 0.4491 | 33.00 | 39.97 | 1 | 0.0 | 0 | K.YGPPCPSCPAPEFLGGPSVFLFPPKPK.D | Carbamidomethyl: 5, 8 | 102-128 | CID |
| 1187 | 1229.913 | 3 | 266.02 | 0.3271 | 33.40 | 21.34 | 1 | 0.0 | 1 | K.NQVSLTCLVKGFYPSDIAVEWESNGQPENNY K.T | Carbamidomethyl: 7 | 241-272 | CID |
| 892 | 1273.142 | 2 | 450.02 | 0.5727 | 28.39 | 101.87 | 7 | 0.0 | 0 | K.GFYPSDIAVEWESNGQPENNYK.T | | 251-272 | CID |

##### Protein 8: Immunoglobulin heavy constant gamma 1 OS=Homo sapiens GN=IGHG1 PE=1 SV=1

| | | | |
|---|---|---|---|
| **Accession:** | IGHG1_HUMAN | **Score:** | 339.5 |
| **Database:** | SwissProt | **Seq. Coverage [%]:** | 32.4 |
| **MW [kDa] / pI:** | 36.1 / 8.5 | **No. of Peptides:** | 7 |
| **Modification(s):** | Carbamidomethyl, Oxidation | | |

| 10 | 20 | 30 | 40 | 50 | 60 | 70 | 80 | 90 | 100 |
|---|---|---|---|---|---|---|---|---|---|
| ASTKGPSVFP | LAPSSKSTSG | GTAALGCLVK | DYFPEPVTVS | WNSGALTSGV | HTFPAVLQSS | GLYSLSSVVT | VPSSSLGTQT | YICNVNHKPS | NTKVDKKVEP |
| **110** | **120** | **130** | **140** | **150** | **160** | **170** | **180** | **190** | **200** |
| KSCDKTHTCP | PCPAPELLGG | PSVFLFPPKP | KDTLMISRTP | EVTCVVVDVS | HEDPEVKFNW | YVDGVEVHNA | KTKPREEQYN | STYRVVSVLT | VLHQDWLNGK |
| **210** | **220** | **230** | **240** | **250** | **260** | **270** | **280** | **290** | **300** |
| EYKCKVSNKA | LPAPIEKTIS | KAKGQPREPQ | VYTLPPSRDE | LTKNQVSLTC | LVKGFYPSDI | AVEWESNGQP | ENNYKTTPPV | LDSDGSFFLY | SKLTVDKSRW |
| **310** | **320** | **330** | **340** | | | | | | |
| QQGNVFSCSV | MHEALHNHYT | QKSLSLSPGK | | | | | | | |

**MS/MS Peptide Matches**

| Cmpd. | m/z meas. | z | Δ m/z [ppm] | Δ m/z [Da] | Rt [min] | Score | No. of Cmpds. | Site [%] | P | Sequence | Modification | Range | Type |
|---|---|---|---|---|---|---|---|---|---|---|---|---|---|
| 1059 | 950.553 | 4 | 362.71 | 0.3447 | 31.22 | 52.25 | 3 | 0.0 | 1 | R.TPEVTCVVVDVSHEDPEVKFNWYVDGVEVHNA K.T | Carbamidomethyl: 6 | 139-171 | CID |
| 1187 | 1229.913 | 3 | 266.02 | 0.3271 | 33.40 | 21.34 | 1 | 0.0 | 1 | K.NQVSLTCLVKGFYPSDIAVEWESNGQPENNY K.T | Carbamidomethyl: 7 | 244-275 | CID |
| 892 | 1273.142 | 2 | 450.02 | 0.5727 | 28.39 | 101.87 | 7 | 0.0 | 0 | K.GFYPSDIAVEWESNGQPENNYK.T | | 254-275 | CID |
| 970 | 937.508 | 2 | 46.35 | 0.0435 | 29.66 | 40.46 | 1 | 0.0 | 0 | K.TTPPVLDSDGSFFLYSK.L | | 276-292 | CID |
| 586 | 762.193 | 4 | 442.97 | 0.3375 | 23.19 | 55.91 | 3 | 0.0 | 1 | K.SRWQQGNVFSCSVMHEALHNHYTQK.S | Carbamidomethyl: 11 | 298-322 | CID |
| 632 | 934.875 | 3 | 479.20 | 0.4478 | 24.00 | 39.36 | 3 | 0.0 | 0 | R.WQQGNVFSCSVMHEALHNHYTQK.S | Carbamidomethyl: 9 | 300-322 | CID |

| Cmpd. | m/z meas. | z | Δ m/z [ppm] | Δ m/z [Da] | Rt [min] | Score | No. of Cmpds. | Site [%] | P | Sequence | Modification | Range | Type |
|---|---|---|---|---|---|---|---|---|---|---|---|---|---|
| 479 | 939.834 | 3 | 79.96 | 0.0751 | 21.05 | 28.27 | 1 | 0.0 | 0 | R.WQQGNVFSCSVMHEALHNHYTQK.S | Carbamidomethyl: 9; Oxidation: 12 | 300-322 | CID |

#### Protein 9: Keratin, type II cytoskeletal 1 OS=Homo sapiens GN=KRT1 PE=1 SV=6

**Accession:** K2C1_HUMAN
**Database:** SwissProt
**MW [kDa] / pI:** 66.0 / 8.2

**Score:** 203.7
**Seq. Coverage [%]:** 4.8
**No. of Peptides:** 1

| 10 | 20 | 30 | 40 | 50 | 60 | 70 | 80 | 90 | 100 |
|---|---|---|---|---|---|---|---|---|---|
| MSRQFSSRSG | YRSGGGFSSG | SAGIINYQRR | TTSSSTRRSG | GGGGRFSSCG | GGGGSFGAGG | GFGSRSLVNL | GGSKSISISV | ARGGGRGSGF | GGGYGGGGFG |
| **110** | **120** | **130** | **140** | **150** | **160** | **170** | **180** | **190** | **200** |
| GGGFGGGGFG | GGGIGGGGFG | GFGSGGGGFG | GGGFGGGGYG | GGYGPVCPPG | GIQEVTINQS | LLQPLNVEID | PEIQKVKSRE | REQIKSLNNQ | FASFIDKVRF |
| **210** | **220** | **230** | **240** | **250** | **260** | **270** | **280** | **290** | **300** |
| LEQQNQVLQT | KWELLQQVDT | STRTHNLEPY | FESFINNLRR | RVDQLKSDQS | RLDSELKNMQ | DMVEDYRNKY | EDEINKRTNA | ENEFVTIKKD | VDGAYMTKVD |
| **310** | **320** | **330** | **340** | **350** | **360** | **370** | **380** | **390** | **400** |
| LQAKLDNLQQ | EIDFLTALYQ | AELSQMQTQI | SETNVILSMD | NNRSLDLDSI | IAEVKAQYED | IAQKSKAEAE | SLYQSKYEEL | QITAGRHGDS | VRNSKIEISE |
| **410** | **420** | **430** | **440** | **450** | **460** | **470** | **480** | **490** | **500** |
| LNRVIQRLRS | EIDNVKKQIS | NLQQSISDAE | QRGENALKDA | KNKLNDLEDA | LQQAKEDLAR | LLRDYQELMN | TKLALDLEIA | TYRTLLEGEE | SRMSGECAPN |
| **510** | **520** | **530** | **540** | **550** | **560** | **570** | **580** | **590** | **600** |
| VSVSVSTSHT | TISGGGSRGG | GGGGYGSGGS | SYGSGGGSYG | SGGGGGGGRG | SYGSGGSSYG | SGGGSYGSGG | GGGGHGSYGS | GSSSGGYRGG | SGGGGGGSSG |
| **610** | **620** | **630** | **640** | **650** | | | | | |
| GRGSGGGSSG | GSIGGRGSSS | GGVKSSGGSS | SVKFVSTTYS | GVTR | | | | | |

MS/MS Peptide Matches

| Cmpd. | m/z meas. | z | Δ m/z [ppm] | Δ m/z [Da] | Rt [min] | Score | No. of Cmpds. | Site [%] | P | Sequence | Modification | Range | Type |
|---|---|---|---|---|---|---|---|---|---|---|---|---|---|
| 165 | 1192.963 | 2 | 405.41 | 0.4834 | 14.34 | 203.69 | 4 | 0.0 | 0 | R.GGGGGGYGSGGSSYGSGGGSYGSGGGGGG GR.G | | 519-549 | CID |

#### Protein 10: Keratin, type I cytoskeletal 9 OS=Homo sapiens GN=KRT9 PE=1 SV=3

| | | | |
|---|---|---|---|
| **Accession:** | K1C9_HUMAN | **Score:** | 159.4 |
| **Database:** | SwissProt | **Seq. Coverage [%]:** | 3.7 |
| **MW [kDa] / pI:** | 62.0 / 5.1 | **No. of Peptides:** | 1 |

| 10 | 20 | 30 | 40 | 50 | 60 | 70 | 80 | 90 | 100 |
|---|---|---|---|---|---|---|---|---|---|
| MSCRQFSSSY | LSRSGGGGGG | GLGSGGSIRS | SYSRFSSSGG | GGGGGRFSSS | SGYGGGSSRV | CGRGGGGSFG | YSYGGGSGGG | FSASSLGGGF | GGGSRGFGGA |
| **110** | **120** | **130** | **140** | **150** | **160** | **170** | **180** | **190** | **200** |
| SGGGYSSSGG | FGGGFGGGSG | GGFGGGYGSG | FGGFGGFGGG | AGGGDGGILT | ANEKSTMQEL | NSRLASYLDK | VQALEEANND | LENKIQDWYD | KKGPAAIQKN |
| **210** | **220** | **230** | **240** | **250** | **260** | **270** | **280** | **290** | **300** |
| YSPYYNTIDD | LKDQIVDLTV | GNNKTLLDID | NTRMTLDDFR | IKFEMEQNLR | QGVDADINGL | RQVLDNLTME | KSDLEMQYET | LQEELMALKK | NHKEEMSQLT |
| **310** | **320** | **330** | **340** | **350** | **360** | **370** | **380** | **390** | **400** |
| GQNSGDVNVE | INVAPGKDLT | KTLNDMRQEY | EQLIAKNRKD | IENQYETQIT | QIEHEVSSSG | QEVQSSAKEV | TQLRHGVQEL | EIELQSQLSK | KAALEKSLED |
| **410** | **420** | **430** | **440** | **450** | **460** | **470** | **480** | **490** | **500** |
| TKNRYCGQLQ | MIQEQISNLE | AQITDVRQEI | ECQNQEYSLL | LSIKMRLEKE | IETYHNLLEG | GQEDFESSGA | GKIGLGGRGG | SGGSYGRGSR | GGSGGSYGGG |
| **510** | **520** | **530** | **540** | **550** | **560** | **570** | **580** | **590** | **600** |
| GSGGGYGGGS | GSRGGSGGSY | GGGSGSGGGS | GGGYGGGSGG | GHSGGSGGGH | SGGSGGNYGG | GSGSGGGSGG | GYGGGSGSRG | GSGGSHGGGS | GFGGESGGSY |
| **610** | **620** | **630** | | | | | | | |
| GGGEEASGSG | GGYGGGSGKS | SHS | | | | | | | |

MS/MS Peptide Matches

| Cmpd. | m/z meas. | z | Δ m/z [ppm] | Δ m/z [Da] | Rt [min] | Score | No. of Cmpds. | Site [%] | P | Sequence | Modification | Range | Type |
|---|---|---|---|---|---|---|---|---|---|---|---|---|---|
| 89 | 896.478 | 2 | 123.28 | 0.1105 | 12.16 | 159.42 | 3 | 0.0 | 0 | R.GGSGGSYGGGGSGGGYGGGSGSR.G | | 491-513 | CID |

#### Protein 11: Immunoglobulin heavy constant gamma 3 OS=Homo sapiens GN=IGHG3 PE=1 SV=2

**Accession:** IGHG3_HUMAN
**Database:** SwissProt
**MW [kDa] / pI:** 41.3 / 8.2
**Modification(s):** Carbamidomethyl, Oxidation

**Score:** 128.2
**Seq. Coverage [%]:** 8.2
**No. of Peptides:** 3

| 10 | 20 | 30 | 40 | 50 | 60 | 70 | 80 | 90 | 100 |
|---|---|---|---|---|---|---|---|---|---|
| ASTKGPSVFP | LAPCSRSTSG | GTAALGCLVK | DYFPEPVTVS | WNSGALTSGV | HTFPAVLQSS | GLYSLSSVVT | VPSSSLGTQT | YTCNVNHKPS | NTKVDKRVEL |
| **110** | **120** | **130** | **140** | **150** | **160** | **170** | **180** | **190** | **200** |
| KTPLGDTTHT | CPRCPEPKSC | DTPPPCPRCP | EPKSCDTPPP | CPRCPEPKSC | DTPPPCPRCP | APELLGGPSV | FLFPPKPKDT | LMISRTPEVT | CVVVDVSHED |
| **210** | **220** | **230** | **240** | **250** | **260** | **270** | **280** | **290** | **300** |
| PEVQFKWYVD | GVEVHNAKTK | PREEQYNSTF | RVVSVLTVLH | QDWLNGKEYK | CKVSNKALPA | PIEKTISKTK | GQPREPQVYT | LPPSREEMTK | NQVSLTCLVK |
| **310** | **320** | **330** | **340** | **350** | **360** | **370** | **380** | | |
| GFYPSDIAVE | WESSGQPENN | YNTTPPMLDS | DGSFFLYSKL | TVDKSRWQQG | NIFSCSVMHE | ALHNRFTQKS | LSLSPGK | | |

MS/MS Peptide Matches

| Cmpd. | m/z meas. | z | Δ m/z [ppm] | Δ m/z [Da] | Rt [min] | Score | No. of Cmpds. | Site [%] | P | Sequence | Modification | Range | Type |
|---|---|---|---|---|---|---|---|---|---|---|---|---|---|
| 154 | 899.459 | 2 | 77.30 | 0.0695 | 14.07 | 48.92 | 2 | 0.0 | 1 | R.CPEPKSCDTPPPCPR.C | Carbamidomethyl: 1, 7, 13 | 114-128 | CID |
| 406 | 961.531 | 2 | 581.48 | 0.5588 | 19.67 | 17.05 | 1 | 0.0 | 1 | R.EPQVYTLPPSREEMTK.N | Oxidation: 14 | 275-290 | CID |
| 448 | 953.075 | 2 | 105.19 | 0.1002 | 20.51 | 62.22 | 8 | 0.0 | 1 | R.EPQVYTLPPSREEMTK.N | | 275-290 | CID |

#### Protein 12: Keratin, type I cytoskeletal 10 OS=Homo sapiens GN=KRT10 PE=1 SV=6

| | | | |
|---|---|---|---|
| **Accession:** | K1C10_HUMAN | **Score:** | 124.4 |
| **Database:** | SwissProt | **Seq. Coverage [%]:** | 3.3 |
| **MW [kDa] / pI:** | 58.8 / 5.1 | **No. of Peptides:** | 1 |

| 10 | 20 | 30 | 40 | 50 | 60 | 70 | 80 | 90 | 100 |
|---|---|---|---|---|---|---|---|---|---|
| MSVRYSSSKH | YSSSRSGGGG | GGGGCGGGGG | VSSLRISSSK | GSLGGGFSSG | GFSGGSFSRG | SSGGGCFGGS | SGGYGGLGGF | GGGSFRGSYG | SSSFGGSYGG |
| **110** | **120** | **130** | **140** | **150** | **160** | **170** | **180** | **190** | **200** |
| IFGGGSFGGG | SFGGGSFGGG | GFGGGGFGGG | FGGGFGGDGG | LLSGNEKVTM | QNLNDRLASY | LDKVRALEES | NYELEGKIKE | WYEKHGNSHQ | GEPRDYSKYY |
| **210** | **220** | **230** | **240** | **250** | **260** | **270** | **280** | **290** | **300** |
| KTIDDLKNQI | LNLTTDNANI | LLQIDNARLA | ADDFRLKYEN | EVALRQSVEA | DINGLRRVLD | ELTLTKADLE | MQIESLTEEL | AYLKKNHEEE | MKDLRNVSTG |
| **310** | **320** | **330** | **340** | **350** | **360** | **370** | **380** | **390** | **400** |
| DVNVEMNAAP | GVDLTQLLNN | MRSQYEQLAE | QNRKDAEAWF | NEKSKELTTE | IDNNIEQISS | YKSEITELRR | NVQALEIELQ | SQLALKQSLE | ASLAETEGRY |
| **410** | **420** | **430** | **440** | **450** | **460** | **470** | **480** | **490** | **500** |
| CVQLSQIQAQ | ISALEEQLQQ | IRAETECQNT | EYQQLLDIKI | RLENEIQTYR | SLLEGEGSSG | GGGRGGGSFG | GGYGGGSSGG | GSSGGGHGGG | HGGSSGGGYG |
| **510** | **520** | **530** | **540** | **550** | **560** | **570** | **580** | **590** | |
| GGSSGGGSSG | GGYGGGSSSG | GHGGSSSGGY | GGGSSGGGGG | GYGGGSSGGG | SSSGGGYGGG | SSSGGHKSSS | SGSVGESSSK | GPRY | |

MS/MS Peptide Matches

| Cmpd. | m/z meas. | z | Δ m/z [ppm] | Δ m/z [Da] | Rt [min] | Score | No. of Cmpds. | Site [%] | P | Sequence | Modification | Range | Type |
|---|---|---|---|---|---|---|---|---|---|---|---|---|---|
| 622 | 854.484 | 2 | 110.36 | 0.0943 | 23.83 | 124.43 | 1 | 0.0 | 0 | K.GSLGGGFSSGGFSGGSFSR.G | | 41-59 | CID |

#### Protein 13: Keratin, type II cytoskeletal 2 epidermal OS=Homo sapiens GN=KRT2 PE=1 SV=2

| | | | |
|---|---|---|---|
| **Accession:** | K22E_HUMAN | **Score:** | 92.0 |
| **Database:** | SwissProt | **Seq. Coverage [%]:** | 3.6 |
| **MW [kDa] / pI:** | 65.4 / 8.1 | **No. of Peptides:** | 1 |

| 10 | 20 | 30 | 40 | 50 | 60 | 70 | 80 | 90 | 100 |
|---|---|---|---|---|---|---|---|---|---|
| MSCQISCKSR | GRGGGGGGFR | GFSSGSAVVS | GGSRRSTSSF | SCLSRHGGGG | GGFGGGGFGS | RSLVGLGGTK | SISISVAGGG | GGFGAAGGFG | GRGGGFGGGS |
| **110** | **120** | **130** | **140** | **150** | **160** | **170** | **180** | **190** | **200** |
| SFGGGSGFSG | GGFGGGGFGG | GRFGGFGGPG | GVGGLGGPGG | FGPGGYPGGI | HEVSVNQSLL | QPLNVKVDPE | IQNVKAQERE | QIKTLNNKFA | SFIDKVRFLE |
| **210** | **220** | **230** | **240** | **250** | **260** | **270** | **280** | **290** | **300** |
| QQNQVLQTKW | ELLQQMNVGT | RPINLEPIFQ | GYIDSLKRYL | DGLTAERTSQ | NSELNNMQDL | VEDYKKKYED | EINKRTAAEN | DFVTLKKDVD | NAYMIKVELQ |
| **310** | **320** | **330** | **340** | **350** | **360** | **370** | **380** | **390** | **400** |
| SKVDLLNQEI | EFLKVLYDAE | ISQIHQSVTD | TNVILSMDNS | RNLDLDSIIA | EVKAQYEEIA | QRSKEEAEAL | YHSKYEELQV | TVGRHGDSLK | EIKIEISELN |
| **410** | **420** | **430** | **440** | **450** | **460** | **470** | **480** | **490** | **500** |
| RVIQRLQGEI | AHVKKQCKNV | QDAIADAEQR | GEHALKDARN | KLNDLEEALQ | QAKEDLARLL | RDYQELMNVK | LALDVEIATY | RKLLEGEECR | MSGDLSSNVT |
| **510** | **520** | **530** | **540** | **550** | **560** | **570** | **580** | **590** | **600** |
| VSVTSSTISS | NVASKAAFGG | SGGRGSSSGG | GYSSGSSSYG | SGGRQSGSRG | GSGGGGSISG | GGYGSGGGSG | GRYGSGGGSK | GGSISGGGYG | SGGGKHSSGG |
| **610** | **620** | **630** | **640** | | | | | | |
| GSRGGSSSGG | GYGSGGGGSS | SVKGSSGEAF | GSSVTFSFR | | | | | | |

MS/MS Peptide Matches

| Cmpd. | m/z meas. | z | Δ m/z [ppm] | Δ m/z [Da] | Rt [min] | Score | No. of Cmpds. | Site [%] | P | Sequence | Modification | Range | Type |
|---|---|---|---|---|---|---|---|---|---|---|---|---|---|
| 122 | 871.448 | 2 | 80.49 | 0.0701 | 13.16 | 92.01 | 2 | 0.0 | 0 | R.GGSGGGGSISGGGYGSGGGSGGR.Y | | 550-572 | CID |

#### Protein 14: Immunoglobulin kappa variable 3-15 OS=Homo sapiens GN=IGKV3-15 PE=1 SV=2

| | | | |
|---|---|---|---|
| **Accession:** | KV315_HUMAN | **Score:** | 63.7 |
| **Database:** | SwissProt | **Seq. Coverage [%]:** | 18.3 |
| **MW [kDa] / pI:** | 12.5 / 5.1 | **No. of Peptides:** | 1 |

| 10 | 20 | 30 | 40 | 50 | 60 | 70 | 80 | 90 | 100 |
|---|---|---|---|---|---|---|---|---|---|
| MEAPAQLLFL | LLLWLPDTTG | EIVMTQSPAT | LSVSPGERAT | LSCRASQSVS | SNLAWYQQKP | GQAPRLLIYG | ASTRATGIPA | RFSGSGSGTE | FTLTISSLQS |
| 110 | 120 | | | | | | | | |
| EDFAVYYCQQ | YNNWP | | | | | | | | |

MS/MS Peptide Matches

| Cmpd. | m/z meas. | z | Δ m/z [ppm] | Δ m/z [Da] | Rt [min] | Score | No. of Cmpds. | Site [%] | P | Sequence | Modification | Range | Type |
|---|---|---|---|---|---|---|---|---|---|---|---|---|---|
| 501 | 768.473 | 3 | 109.20 | 0.0839 | 21.51 | 63.7 | 3 | 0.0 | 0 | R.ASQSVSSNLAWYQQKPGQAPR.L | | 45-65 | CID |

#### Protein 15: Immunoglobulin heavy variable 4-4 OS=Homo sapiens GN=IGHV4-4 PE=3 SV=2

**Accession:** HV404_HUMAN
**Database:** SwissProt
**MW [kDa] / pI:** 12.8 / 9.6
**Modification(s):** Carbamidomethyl

**Score:** 55.2
**Seq. Coverage [%]:** 13.7
**No. of Peptides:** 1

| 10 | 20 | 30 | 40 | 50 | 60 | 70 | 80 | 90 | 100 |
|---|---|---|---|---|---|---|---|---|---|
| MKHLWFFLLL | VAAPRWVLSQ | VQLQESGPGL | VKPSGTLSLT | CAVSGGSISS | SNWWSWVRQP | PGKGLEWIGE | IYHSGSTNYN | PSLKSRVTIS | VDKSKNQFSL |
| 110 | 120 | | | | | | | | |
| KLSSVTAADT | AVYYCAR | | | | | | | | |

MS/MS Peptide Matches

| Cmpd. | m/z meas. | z | Δ m/z [ppm] | Δ m/z [Da] | Rt [min] | Score | No. of Cmpds. | Site [%] | P | Sequence | Modification | Range | Type |
|---|---|---|---|---|---|---|---|---|---|---|---|---|---|
| 547 | 874.485 | 2 | 74.77 | 0.0654 | 22.39 | 55.17 | 3 | 0.0 | 0 | K.LSSVTAADTAVYYCAR.- | Carbamidomethyl: 14 | 102-117 | CID |

#### Protein 16: Immunoglobulin lambda variable 8-61 OS=Homo sapiens GN=IGLV8-61 PE=3 SV=7

| | | | |
|---|---|---|---|
| **Accession:** | LV861_HUMAN | **Score:** | 38.0 |
| **Database:** | SwissProt | **Seq. Coverage [%]:** | 13.1 |
| **MW [kDa] / pI:** | 12.8 / 4.3 | **No. of Peptides:** | 1 |

```
        10         20         30         40         50         60         70         80         90        100
MSVPTMAWMM LLLGLLAYGS GVDSQTVVTQ EPSFSVSPGG TVTLTCGLSS GSVSTSYYPS WYQQTPGQAP RTLIYSTNTR SSGVPDRFSG SILGNKAALT
       110        120        130
ITGAQADDES DYYCVLYMGS GI
```

**MS/MS Peptide Matches**

| Cmpd. | m/z meas. | z | Δ m/z [ppm] | Δ m/z [Da] | Rt [min] | Score | No. of Cmpds. | Site [%] | P | Sequence | Modification | Range | Type |
|---|---|---|---|---|---|---|---|---|---|---|---|---|---|
| 541 | 811 | 2 | 97.86 | 0.0794 | 22.27 | 37.98 | 2 | 0.0 | 1 | R.SSGVPDRFSGSILGNK.A | | 81-96 | CID |

#### Protein 17: Immunoglobulin kappa variable 2D-28 OS=Homo sapiens GN=IGKV2D-28 PE=1 SV=2

| | | | |
|---|---|---|---|
| **Accession:** | KVD28_HUMAN | **Score:** | 34.1 |
| **Database:** | SwissProt | **Seq. Coverage [%]:** | 16.7 |
| **MW [kDa] / pI:** | 12.9 / 5.6 | **No. of Peptides:** | 1 |

```
        10         20         30         40         50         60         70         80         90        100
MRLPAQLLGL LMLWVSGSSG DIVMTQSPLS LPVTPGEPAS ISCRSSQSLL HSNGYNYLDW YLQKPGQSPQ LLIYLGSNRA SGVPDRFSGS GSGTDFTLKI
       110        120        130
SRVEAEDVGV YYCMQALQTP
```

**MS/MS Peptide Matches**

| Cmpd. | m/z meas. | z | Δ m/z [ppm] | Δ m/z [Da] | Rt [min] | Score | No. of Cmpds. | Site [%] | P | Sequence | Modification | Range | Type |
|---|---|---|---|---|---|---|---|---|---|---|---|---|---|
| 552 | 993.552 | 2 | 70.67 | 0.0702 | 22.48 | 34.14 | 2 | 0.0 | 1 | R.ASGVPDRFSGSGSGTDFTLK.I | | 80-99 | CID |

#### Project Info

| | | | |
|---|---|---|---|
| **Name:** | Alagesan | **Date:** | Mar 5, 2019 |
| **Note:** | Nanobooster | | |

#### Sample Info & Protocols

| | |
|---|---|
| **Name:** | Nanobooster - IgG |
| **Date:** | Mar 5, 2019 |

#### Search Result Info

| | | | |
|---|---|---|---|
| **Search Result:** | IonTrap_Human_Swiss_CID_2019-03-05 17:55:08 | | |
| **Location:** | /Alagesan/Nanobooster - IgG/KA_IgG3 HD NB ACN CID_GA4_01_9083.mgf | | |
| **Search Method:** | IonTrap_Human_Swiss_CID | **Version:** | 1.2 |
| **Search Engine(s):** | Mascot, 2.6.0 | **Ident. Compound(s):** | 0/2361 |
| **Database(s):** | SwissProt, SwissProt_2017_04.fasta | | |
| **Note:** | | | |

#### Results

##### Protein 1: Immunoglobulin heavy constant gamma 3 OS=Homo sapiens GN=IGHG3 PE=1 SV=2

| | | | |
|---|---|---|---|
| **Accession:** | IGHG3_HUMAN | **Score:** | 1434.0 |
| **Database:** | SwissProt | **Seq. Coverage [%]:** | 58.6 |
| **MW [kDa] / pI:** | 41.3 / 8.2 | **No. of Peptides:** | 20 |
| **Modification(s):** | Carbamidomethyl, Oxidation | | |

```
        10         20         30         40         50         60         70         80         90        100
ASTKGPSVFP LAPCSRSTSG GTAALGCLVK DYFPEPVTVS WNSGALTSGV HTFPAVLQSS GLYSLSSVVT VPSSSLGTQT YTCNVNHKPS NTKVDKRVEL
       110        120        130        140        150        160        170        180        190        200
KTPLGDTTHT CPRCPEPKSC DTPPPCPRCP EPKSCDTPPP CPRCPEPKSC DTPPPCPRCP APELLGGPSV FLFPPKPKDT LMISRTPEVT CVVVDVSHED
       210        220        230        240        250        260        270        280        290        300
PEVQFKWYVD GVEVHNAKTK PREEQYNSTF RVVSVLTVLH QDWLNGKEYK CKVSNKALPA PIEKTISKTK GQPREPQVYT LPPSREEMTK NQVSLTCLVK
       310        320        330        340        350        360        370        380
GFYPSDIAVE WESSGQPENN YNTTPPMLDS DGSFFLYSKL TVDKSRWQQG NIFSCSVMHE ALHNRFTQKS LSLSPGK
```

**MS/MS Peptide Matches**

| Cmpd. | m/z meas. | z | Δ m/z [ppm] | Δ m/z [Da] | Rt [min] | Score | No. of Cmpds. | Site [%] | P | Sequence | Modification | Range | Type |
|---|---|---|---|---|---|---|---|---|---|---|---|---|---|
| 926 | 864.526 | 3 | 482.90 | 0.4173 | 28.44 | 63.41 | 3 | 0.0 | 1 | K.GPSVFPLAPCSRSTSGGTAALGCLVK.D | Carbamidomethyl: 10, 23 | 5-30 | CID |
| 404 | 913.081 | 2 | 124.57 | 0.1137 | 18.99 | 79.04 | 5 | 0.0 | 1 | R.VELKTPLGDTTHTCPR.C | Carbamidomethyl: 14 | 98-113 | CID |
| 208 | 984.01 | 2 | 51.71 | 0.0509 | 15.14 | 53.79 | 3 | 0.0 | 1 | K.TPLGDTTHTCPRCPEPK.S | Carbamidomethyl: 10, 13 | 102-118 | CID |
| 143 | 899.977 | 2 | 653.24 | 0.5875 | 13.88 | 81.13 | 13 | 0.0 | 1 | R.CPEPKSCDTPPPCPR.C | Carbamidomethyl: 1, 7, 13 | 114-128 | CID |
| 187 | 899.43 | 2 | 45.05 | 0.0405 | 14.76 | 32.25 | 3 | 0.0 | 1 | K.SCDTPPPCPRCPEPK.S | Carbamidomethyl: 2, 8, 11 | 119-133 | CID |
| 1134 | 718.12 | 3 | 547.26 | 0.3928 | 31.97 | 107.77 | 21 | 0.0 | 0 | R.CPAPELLGGPSVFLFPPKPK.D | Carbamidomethyl: 1 | 159-178 | CID |

| Cmpd. | m/z meas. | z | Δ m/z [ppm] | Δ m/z [Da] | Rt [min] | Score | No. of Cmpds. | Site [%] | P | Sequence | Modification | Range | Type |
|---|---|---|---|---|---|---|---|---|---|---|---|---|---|
| 1166 | 742.771 | 4 | 161.15 | 0.1197 | 32.49 | 84.44 | 6 | 0.0 | 1 | R.CPAPELLGGPSVFLFPPKPKDTLMISR.T | Carbamidomethyl: 1 | 159-185 | CID |
| 1115 | 746.766 | 4 | 155.29 | 0.1159 | 31.65 | 93.04 | 3 | 0.0 | 1 | R.CPAPELLGGPSVFLFPPKPKDTLMISR.T | Carbamidomethyl: 1; Oxidation: 24 | 159-185 | CID |
| 1008 | 1077.948 | 3 | 389.37 | 0.4196 | 29.82 | 75.89 | 3 | 0.0 | 1 | K.DTLMISRTPEVTCVVVDVSHEDPEVQFK.W | Carbamidomethyl: 13 | 179-206 | CID |
| 876 | 805.528 | 3 | 171.77 | 0.1383 | 27.36 | 106.95 | 6 | 0.0 | 0 | R.TPEVTCVVVDVSHEDPEVQFK.W | Carbamidomethyl: 6 | 186-206 | CID |
| 1050 | 1271.652 | 3 | 292.15 | 0.3714 | 30.49 | 70.2 | 3 | 0.0 | 1 | R.TPEVTCVVVDVSHEDPEVQFKWYVDGVEVHNAK.T | Carbamidomethyl: 6 | 186-218 | CID |
| 510 | 708.959 | 2 | 155.24 | 0.1100 | 20.93 | 101.23 | 17 | 0.0 | 0 | K.WYVDGVEVHNAK.T | | 207-218 | CID |
| 1158 | 904.578 | 2 | 78.62 | 0.0711 | 32.36 | 92.6 | 3 | 0.0 | 0 | R.VVSVLTVLHQDWLNGK.E | | 232-247 | CID |
| 1062 | 743.527 | 3 | 161.00 | 0.1197 | 30.73 | 90.15 | 7 | 0.0 | 1 | R.VVSVLTVLHQDWLNGKEYK.C | | 232-250 | CID |
| 519 | 953.012 | 2 | 39.08 | 0.0372 | 21.13 | 58.07 | 13 | 0.0 | 1 | R.EPQVYTLPPSREEMTK.N | | 275-290 | CID |
| 457 | 961.021 | 2 | 50.77 | 0.0488 | 19.92 | 28.13 | 1 | 0.0 | 1 | R.EPQVYTLPPSREEMTK.N | Oxidation: 14 | 275-290 | CID |
| 1298 | 1464.236 | 3 | 170.53 | 0.2497 | 34.72 | 60.39 | 2 | 0.0 | 0 | K.GFYPSDIAVEWESSGQPENNYNTTPPMLDSDGSFFLYSK.L | | 301-339 | CID |
| 695 | 853.516 | 3 | 523.56 | 0.4466 | 24.39 | 32.47 | 2 | 0.0 | 1 | K.SRWQQGNIFSCSVMHEALHNR.F | Carbamidomethyl: 11 | 345-365 | CID |
| 770 | 1158.068 | 2 | 461.46 | 0.5342 | 25.64 | 73.63 | 4 | 0.0 | 0 | R.WQQGNIFSCSVMHEALHNR.F | Carbamidomethyl: 9 | 347-365 | CID |
| 823 | 940.528 | 3 | 439.45 | 0.4131 | 26.50 | 49.39 | 3 | 0.0 | 1 | R.WQQGNIFSCSVMHEALHNRFTQK.S | Carbamidomethyl: 9 | 347-369 | CID |

#### Protein 2: Immunoglobulin kappa constant OS=Homo sapiens GN=IGKC PE=1 SV=2

**Accession:** IGKC_HUMAN
**Database:** SwissProt
**MW [kDa] / pI:** 11.8 / 6.1
**Modification(s):** Carbamidomethyl

**Score:** 1002.1
**Seq. Coverage [%]:** 86.9
**No. of Peptides:** 10

| 10 | 20 | 30 | 40 | 50 | 60 | 70 | 80 | 90 | 100 |
|---|---|---|---|---|---|---|---|---|---|
| RTVAAPSVFI | FPPSDEQLKS | GTASVVCLLN | NFYPREAKVQ | WKVDNALQSG | NSQESVTEQD | SKDSTYSLSS | TLTLSKADYE | KHKVYACEVT | HQGLSSPVTK |

| 110 |
|---|
| SFNRGEC |

MS/MS Peptide Matches

| Cmpd. | m/z meas. | z | Δ m/z [ppm] | Δ m/z [Da] | Rt [min] | Score | No. of Cmpds. | Site [%] | P | Sequence | Modification | Range | Type |
|---|---|---|---|---|---|---|---|---|---|---|---|---|---|
| 955 | 701.434 | 3 | 75.75 | 0.0531 | 28.94 | 43.43 | 8 | 0.0 | 1 | -.RTVAAPSVFIFPPSDEQLK.S | | 1-19 | CID |
| 1071 | 973.642 | 2 | 128.28 | 0.1249 | 30.88 | 84.25 | 14 | 0.0 | 0 | R.TVAAPSVFIFPPSDEQLK.S | | 2-19 | CID |
| 1505 | 1242.667 | 3 | 290.35 | 0.3607 | 38.20 | 97.82 | 13 | 0.0 | 1 | R.TVAAPSVFIFPPSDEQLKSGTASVVCLLNNFYPR.E | Carbamidomethyl: 26 | 2-35 | CID |
| 1204 | 899.556 | 2 | 116.46 | 0.1047 | 33.17 | 75.64 | 13 | 0.0 | 0 | K.SGTASVVCLLNNFYPR.E | Carbamidomethyl: 8 | 20-35 | CID |
| 1086 | 709.404 | 3 | 59.94 | 0.0425 | 31.19 | 73.16 | 3 | 0.0 | 1 | K.SGTASVVCLLNNFYPREAK.V | Carbamidomethyl: 8 | 20-38 | CID |
| 515 | 893.48 | 3 | 431.26 | 0.3852 | 21.06 | 121.91 | 6 | 0.0 | 1 | K.VQWKVDNALQSGNSQESVTEQDSK.D | | 39-62 | CID |
| 238 | 1068.809 | 2 | 300.42 | 0.3210 | 15.63 | 170.8 | 7 | 0.0 | 0 | K.VDNALQSGNSQESVTEQDSK.D | | 43-62 | CID |
| 722 | 1207.59 | 3 | 288.85 | 0.3487 | 24.87 | 128.58 | 4 | 0.0 | 1 | K.VDNALQSGNSQESVTEQDSKDSTYSLSSTLTLSK.A | | 43-76 | CID |
| 723 | 751.964 | 2 | 107.92 | 0.0811 | 24.90 | 89.91 | 6 | 0.0 | 0 | K.DSTYSLSSTLTLSK.A | | 63-76 | CID |
| 403 | 938.554 | 2 | 92.60 | 0.0869 | 18.99 | 116.63 | 10 | 0.0 | 0 | K.VYACEVTHQGLSSPVTK.S | Carbamidomethyl: 4 | 84-100 | CID |

#### Protein 3: Immunoglobulin kappa light chain OS=Homo sapiens PE=1 SV=1

**Accession:** IGK_HUMAN
**Database:** SwissProt
**MW [kDa] / pI:** 23.4 / 6.9
**Modification(s):** Carbamidomethyl

**Score:** 874.4
**Seq. Coverage [%]:** 43.0
**No. of Peptides:** 8

| 10 | 20 | 30 | 40 | 50 | 60 | 70 | 80 | 90 | 100 |
|---|---|---|---|---|---|---|---|---|---|
| DIQMTQSPST | LSASVGDRVT | ITCRASQSIN | TWLAWYQQKP | GKAPKLLMYK | ASSLESGVPS | RFIGSGSGTE | FTLTISSLQP | DDFATYYCQQ | YNSDSKMFGQ |
| **110** | **120** | **130** | **140** | **150** | **160** | **170** | **180** | **190** | **200** |
| GTKVEVKGTV | AAPSVFIFPP | SDEQLKSGTA | SVVCLLNNFY | PREAKVQWKV | DNALQSGNSQ | ESVTEQDSKD | STYSLSSTLT | LSKADYEKHK | VYACEVTHQG |
| **210** | **220** | | | | | | | | |
| LSSPVTKSFN | RGEC | | | | | | | | |

MS/MS Peptide Matches

| Cmpd. | m/z meas. | z | Δ m/z [ppm] | Δ m/z [Da] | Rt [min] | Score | No. of Cmpds. | Site [%] | P | Sequence | Modification | Range | Type |
|---|---|---|---|---|---|---|---|---|---|---|---|---|---|
| 580 | 947.032 | 2 | 81.78 | 0.0774 | 22.32 | 97.74 | 2 | 0.0 | 0 | -.DIQMTQSPSTLSASVGDR.V | | 1-18 | CID |
| 1204 | 899.556 | 2 | 116.46 | 0.1047 | 33.17 | 75.64 | 13 | 0.0 | 0 | K.SGTASVVCLLNNFYPR.E | Carbamidomethyl: 8 | 127-142 | CID |
| 1086 | 709.404 | 3 | 59.94 | 0.0425 | 31.19 | 73.16 | 3 | 0.0 | 1 | K.SGTASVVCLLNNFYPREAK.V | Carbamidomethyl: 8 | 127-145 | CID |
| 515 | 893.48 | 3 | 431.26 | 0.3852 | 21.06 | 121.91 | 6 | 0.0 | 1 | K.VQWKVDNALQSGNSQESVTEQDSK.D | | 146-169 | CID |
| 238 | 1068.809 | 2 | 300.42 | 0.3210 | 15.63 | 170.8 | 7 | 0.0 | 0 | K.VDNALQSGNSQESVTEQDSK.D | | 150-169 | CID |
| 722 | 1207.59 | 3 | 288.85 | 0.3487 | 24.87 | 128.58 | 4 | 0.0 | 1 | K.VDNALQSGNSQESVTEQDSKDSTYSLSSTLTLSK.A | | 150-183 | CID |
| 723 | 751.964 | 2 | 107.92 | 0.0811 | 24.90 | 89.91 | 6 | 0.0 | 0 | K.DSTYSLSSTLTLSK.A | | 170-183 | CID |
| 403 | 938.554 | 2 | 92.60 | 0.0869 | 18.99 | 116.63 | 10 | 0.0 | 0 | K.VYACEVTHQGLSSPVTK.S | Carbamidomethyl: 4 | 191-207 | CID |

#### Protein 4: Keratin, type II cytoskeletal 1 OS=Homo sapiens GN=KRT1 PE=1 SV=6

**Accession:** K2C1_HUMAN
**Database:** SwissProt
**MW [kDa] / pI:** 66.0 / 8.2

**Score:** 746.5
**Seq. Coverage [%]:** 18.8
**No. of Peptides:** 7

| 10 | 20 | 30 | 40 | 50 | 60 | 70 | 80 | 90 | 100 |
|---|---|---|---|---|---|---|---|---|---|
| MSRQFSSRSG | YRSGGGFSSG | SAGIINYQRR | TTSSSTRRSG | GGGGRFSSCG | GGGGSFGAGG | GFGSRSLVNL | GGSKSISISV | ARGGGRGSGF | GGGYGGGGFG |
| **110** | **120** | **130** | **140** | **150** | **160** | **170** | **180** | **190** | **200** |
| GGGFGGGGFG | GGGIGGGGFG | GFGSGGGGFG | GGGFGGGGYG | GGYGPVCPPG | GIQEVTINQS | LLQPLNVEID | PEIQKVKSRE | REQIKSLNNQ | FASFIDKVRF |
| **210** | **220** | **230** | **240** | **250** | **260** | **270** | **280** | **290** | **300** |
| LEQQNQVLQT | KWELLQQVDT | STRTHNLEPY | FESFINNLRR | RVDQLKSDQS | RLDSELKNMQ | DMVEDYRNKY | EDEINKRTNA | ENEFVTIKKD | VDGAYMTKVD |
| **310** | **320** | **330** | **340** | **350** | **360** | **370** | **380** | **390** | **400** |
| LQAKLDNLQQ | EIDFLTALYQ | AELSQMQTQI | SETNVILSMD | NNRSLDLDSI | IAEVKAQYED | IAQKSKAEAE | SLYQSKYEEL | QITAGRHGDS | VRNSKIEISE |
| **410** | **420** | **430** | **440** | **450** | **460** | **470** | **480** | **490** | **500** |
| LNRVIQRLRS | EIDNVKKQIS | NLQQSISDAE | QRGENALKDA | KNKLNDLEDA | LQQAKEDLAR | LLRDYQELMN | TKLALDLEIA | TYRTLLEGEE | SRMSGECAPN |
| **510** | **520** | **530** | **540** | **550** | **560** | **570** | **580** | **590** | **600** |
| VSVSVSTSHT | TISGGGSRGG | GGGGYGSGGS | SYGSGGGSYG | SGGGGGGGRG | SYGSGGSSYG | SGGGSYGSGG | GGGGHGSYGS | GSSSGGYRGG | SGGGGGGSSG |
| **610** | **620** | **630** | **640** | **650** | | | | | |
| GRGSGGGSSG | GSIGGRGSSS | GGVKSSGGSS | SVKFVSTTYS | GVTR | | | | | |

##### MS/MS Peptide Matches

| Cmpd. | m/z meas. | z | Δ m/z [ppm] | Δ m/z [Da] | Rt [min] | Score | No. of Cmpds. | Site [%] | P | Sequence | Modification | Range | Type |
|---|---|---|---|---|---|---|---|---|---|---|---|---|---|
| 936 | 820.006 | 2 | 88.36 | 0.0724 | 28.60 | 82.77 | 3 | 0.0 | 1 | K.SLNNQFASFIDKVR.F | | 186-199 | CID |
| 456 | 738.516 | 2 | 162.15 | 0.1197 | 19.91 | 84.81 | 3 | 0.0 | 0 | R.FLEQQNQVLQTK.W | | 200-211 | CID |
| 987 | 978.243 | 3 | 67.52 | 0.0660 | 29.48 | 125.4 | 2 | 0.0 | 1 | R.FLEQQNQVLQTKWELLQQVDTSTR.T | | 200-223 | CID |
| 1182 | 997.93 | 2 | 439.14 | 0.4380 | 32.77 | 38.3 | 1 | 0.0 | 0 | R.THNLEPYFESFINNLR.R | | 224-239 | CID |
| 1483 | 783.812 | 3 | 86.36 | 0.0677 | 37.80 | 143.17 | 3 | 0.0 | 1 | R.SLDLDSIIAEVKAQYEDIAQK.S | | 344-364 | CID |
| 547 | 858.991 | 2 | 71.95 | 0.0618 | 21.61 | 84.52 | 3 | 0.0 | 0 | K.QISNLQQSISDAEQR.G | | 418-432 | CID |
| 171 | 1192.947 | 2 | 391.99 | 0.4674 | 14.42 | 187.51 | 3 | 0.0 | 0 | R.GGGGGGYGSGGSSYGSGGGSYGSGGGGGG GR.G | | 519-549 | CID |

##### Protein 5: Immunoglobulin lambda constant 3 OS=Homo sapiens GN=IGLC3 PE=1 SV=1

| | | | |
|---|---|---|---|
| **Accession:** | IGLC3_HUMAN | **Score:** | 745.1 |
| **Database:** | SwissProt | **Seq. Coverage [%]:** | 96.2 |
| **MW [kDa] / pI:** | 11.3 / 6.9 | **No. of Peptides:** | 9 |
| **Modification(s):** | Carbamidomethyl | | |

```
        10         20         30         40         50         60         70         80         90        100
GQPKAAPSVT LFPPSSEELQ ANKATLVCLI SDFYPGAVTV AWKADSSPVK AGVETTTPSK QSNNKYAASS YLSLTPEQWK SHKSYSCQVT HEGSTVEKTV
       110
APTECS
```

**MS/MS Peptide Matches**

| Cmpd. | m/z meas. | z | Δ m/z [ppm] | Δ m/z [Da] | Rt [min] | Score | No. of Cmpds. | Site [%] | P | Sequence | Modification | Range | Type |
|---|---|---|---|---|---|---|---|---|---|---|---|---|---|
| 832 | 993.577 | 2 | 64.86 | 0.0644 | 26.64 | 101.77 | 13 | 0.0 | 0 | K.AAPSVTLFPPSSEELQANK.A | | 5-23 | CID |
| 1503 | 1105.94 | 2 | -126.17 | -0.1396 | 38.15 | 61.98 | 1 | 0.0 | 0 | K.ATLVCLISDFYPGAVTVAWK.A | Carbamidomethyl: 5 | 24-43 | CID |
| 206 | 838.015 | 2 | 100.40 | 0.0841 | 15.10 | 59.23 | 4 | 0.0 | 1 | K.ADSSPVKAGVETTTPSK.Q | | 44-60 | CID |
| 31 | 781.532 | 2 | 176.02 | 0.1375 | 10.87 | 89.38 | 15 | 0.0 | 1 | K.AGVETTTPSKQSNNK.Y | | 51-65 | CID |
| 809 | 1158.12 | 2 | 44.24 | 0.0512 | 26.28 | 61.82 | 4 | 0.0 | 1 | K.QSNNKYAASSYLSLTPEQWK.S | | 61-80 | CID |
| 904 | 872.567 | 2 | 153.53 | 0.1339 | 27.89 | 86.14 | 13 | 0.0 | 0 | K.YAASSYLSLTPEQWK.S | | 66-80 | CID |
| 74 | 1032.605 | 2 | 124.76 | 0.1288 | 12.32 | 88.24 | 5 | 0.0 | 1 | K.SHKSYSCQVTHEGSTVEK.T | Carbamidomethyl: 7 | 81-98 | CID |
| 152 | 856.529 | 2 | 170.21 | 0.1458 | 14.08 | 114.29 | 8 | 0.0 | 0 | K.SYSCQVTHEGSTVEK.T | Carbamidomethyl: 4 | 84-98 | CID |
| 384 | 1279.554 | 2 | 384.12 | 0.4913 | 18.63 | 82.23 | 6 | 0.0 | 1 | K.SYSCQVTHEGSTVEKTVAPTECS.- | Carbamidomethyl: 4, 22 | 84-106 | CID |

BRUKER

##### Protein 6: Immunoglobulin lambda constant 2 OS=Homo sapiens GN=IGLC2 PE=1 SV=1

**Accession:** IGLC2_HUMAN
**Database:** SwissProt
**MW [kDa] / pI:** 11.3 / 6.9
**Modification(s):** Carbamidomethyl

**Score:** 691.1
**Seq. Coverage [%]:** 96.2
**No. of Peptides:** 9

| 10 | 20 | 30 | 40 | 50 | 60 | 70 | 80 | 90 | 100 |
|---|---|---|---|---|---|---|---|---|---|
| GQPKAAPSVT | LFPPSSEELQ | ANKATLVCLI | SDFYPGAVTV | AWKADSSPVK | AGVETTTPSK | QSNNKYAASS | YLSLTPEQWK | SHRSYSCQVT | HEGSTVEKTV |

| 110 |
|---|
| APTECS |

MS/MS Peptide Matches

| Cmpd. | m/z meas. | z | Δ m/z [ppm] | Δ m/z [Da] | Rt [min] | Score | No. of Cmpds. | Site [%] | P | Sequence | Modification | Range | Type |
|---|---|---|---|---|---|---|---|---|---|---|---|---|---|
| 832 | 993.577 | 2 | 64.86 | 0.0644 | 26.64 | 101.77 | 13 | 0.0 | 0 | K.AAPSVTLFPPSSEELQANK.A | | 5-23 | CID |
| 1503 | 1105.94 | 2 | -126.17 | -0.1396 | 38.15 | 61.98 | 1 | 0.0 | 0 | K.ATLVCLISDFYPGAVTVAWK.A | Carbamidomethyl: 5 | 24-43 | CID |
| 206 | 838.015 | 2 | 100.40 | 0.0841 | 15.10 | 59.23 | 4 | 0.0 | 1 | K.ADSSPVKAGVETTTPSK.Q | | 44-60 | CID |
| 31 | 781.532 | 2 | 176.02 | 0.1375 | 10.87 | 89.38 | 15 | 0.0 | 1 | K.AGVETTTPSKQSNNK.Y | | 51-65 | CID |
| 809 | 1158.12 | 2 | 44.24 | 0.0512 | 26.28 | 61.82 | 4 | 0.0 | 1 | K.QSNNKYAASSYLSLTPEQWK.S | | 61-80 | CID |
| 904 | 872.567 | 2 | 153.53 | 0.1339 | 27.89 | 86.14 | 13 | 0.0 | 0 | K.YAASSYLSLTPEQWK.S | | 66-80 | CID |
| 78 | 1046.575 | 2 | 91.49 | 0.0957 | 12.43 | 34.26 | 4 | 0.0 | 1 | K.SHRSYSCQVTHEGSTVEK.T | Carbamidomethyl: 7 | 81-98 | CID |
| 152 | 856.529 | 2 | 170.21 | 0.1458 | 14.08 | 114.29 | 8 | 0.0 | 0 | R.SYSCQVTHEGSTVEK.T | Carbamidomethyl: 4 | 84-98 | CID |
| 384 | 1279.554 | 2 | 384.12 | 0.4913 | 18.63 | 82.23 | 6 | 0.0 | 1 | R.SYSCQVTHEGSTVEKTVAPTECS.- | Carbamidomethyl: 4, 22 | 84-106 | CID |

**Protein 7: Keratin, type II cytoskeletal 2 epidermal OS=Homo sapiens GN=KRT2 PE=1 SV=2**

| | | | |
|---|---|---|---|
| **Accession:** | K22E_HUMAN | **Score:** | 230.5 |
| **Database:** | SwissProt | **Seq. Coverage [%]:** | 5.0 |
| **MW [kDa] / pI:** | 65.4 / 8.1 | **No. of Peptides:** | 2 |

| 10 | 20 | 30 | 40 | 50 | 60 | 70 | 80 | 90 | 100 |
|---|---|---|---|---|---|---|---|---|---|
| MSCQISCKSR | GRGGGGGGFR | GFSSGSAVVS | GGSRRSTSSF | SCLSRHGGGG | GGFGGGGFGS | RSLVGLGGTK | SISISVAGGG | GGFGAAGGFG | GRGGGFGGGS |
| **110** | **120** | **130** | **140** | **150** | **160** | **170** | **180** | **190** | **200** |
| SFGGGSGFSG | GGFGGGGFGG | GRFGGFGGPG | GVGGLGGPGG | FGPGGYPGGI | HEVSVNQSLL | QPLNVKVDPE | IQNVKAQERE | QIKTLNNKFA | SFIDKVRFLE |
| **210** | **220** | **230** | **240** | **250** | **260** | **270** | **280** | **290** | **300** |
| QQNQVLQTKW | ELLQQMNVGT | RPINLEPIFQ | GYIDSLKRYL | DGLTAERTSQ | NSELNNMQDL | VEDYKKKYED | EINKRTAAEN | DFVTLKKDVD | NAYMIKVELQ |
| **310** | **320** | **330** | **340** | **350** | **360** | **370** | **380** | **390** | **400** |
| SKVDLLNQEI | EFLKVLYDAE | ISQIHQSVTD | TNVILSMDNS | RNLDLDSIIA | EVKAQYEEIA | QRSKEEAEAL | YHSKYEELQV | TVGRHGDSLK | EIKIEISELN |
| **410** | **420** | **430** | **440** | **450** | **460** | **470** | **480** | **490** | **500** |
| RVIQRLQGEI | AHVKKQCKNV | QDAIADAEQR | GEHALKDARN | KLNDLEEALQ | QAKEDLARLL | RDYQELMNVK | LALDVEIATY | RKLLEGEECR | MSGDLSSNVT |
| **510** | **520** | **530** | **540** | **550** | **560** | **570** | **580** | **590** | **600** |
| VSVTSSTISS | NVASKAAFGG | SGGRGSSSGG | GYSSGSSSYG | SGGRQSGSRG | GSGGGGSISG | GGYGSGGGSG | GRYGSGGGSK | GGSISGGGYG | SGGGKHSSGG |
| **610** | **620** | **630** | **640** | | | | | | |
| GSRGGSSSGG | GYGSGGGGSS | SVKGSSGEAF | GSSVTFSFR | | | | | | |

MS/MS Peptide Matches

| Cmpd. | m/z meas. | z | Δ m/z [ppm] | Δ m/z [Da] | Rt [min] | Score | No. of Cmpds. | Site [%] | P | Sequence | Modification | Range | Type |
|---|---|---|---|---|---|---|---|---|---|---|---|---|---|
| 456 | 738.516 | 2 | 162.15 | 0.1197 | 19.91 | 84.81 | 3 | 0.0 | 0 | R.FLEQQNQVLQTK.W | | 198-209 | CID |
| 77 | 870.955 | 2 | 113.19 | 0.0986 | 12.40 | 145.67 | 2 | 0.0 | 0 | R.GSSSGGGYSSGSSSYGSGGR.Q | | 525-544 | CID |

#### Protein 8: Immunoglobulin heavy constant gamma 1 OS=Homo sapiens GN=IGHG1 PE=1 SV=1

| | | | |
|---|---|---|---|
| **Accession:** | IGHG1_HUMAN | **Score:** | 229.5 |
| **Database:** | SwissProt | **Seq. Coverage [%]:** | 10.9 |
| **MW [kDa] / pI:** | 36.1 / 8.5 | **No. of Peptides:** | 3 |

| | | | | | | | | | |
|---|---|---|---|---|---|---|---|---|---|
| 10 | 20 | 30 | 40 | 50 | 60 | 70 | 80 | 90 | 100 |
| ASTKGPSVFP | LAPSSKSTSG | GTAALGCLVK | DYFPEPVTVS | WNSGALTSGV | HTFPAVLQSS | GLYSLSSVVT | VPSSSLGTQT | YICNVNHKPS | NTKVDKKVEP |
| 110 | 120 | 130 | 140 | 150 | 160 | 170 | 180 | 190 | 200 |
| KSCDKTHTCP | PCPAPELLGG | PSVFLFPPKP | KDTLMISRTP | EVTCVVVDVS | HEDPEVKFNW | YVDGVEVHNA | KTKPREEQYN | STYRVVSVLT | VLHQDWLNGK |
| 210 | 220 | 230 | 240 | 250 | 260 | 270 | 280 | 290 | 300 |
| EYKCKVSNKA | LPAPIEKTIS | KAKGQPREPQ | VYTLPPSRDE | LTKNQVSLTC | LVKGFYPSDI | AVEWESNGQP | ENNYKTTPPV | LDSDGSFFLY | SKLTVDKSRW |
| 310 | 320 | 330 | 340 | | | | | | |
| QQGNVFSCSV | MHEALHNHYT | QKSLSLSPGK | | | | | | | |

MS/MS Peptide Matches

| Cmpd. | m/z meas. | z | Δ m/z [ppm] | Δ m/z [Da] | Rt [min] | Score | No. of Cmpds. | Site [%] | P | Sequence | Modification | Range | Type |
|---|---|---|---|---|---|---|---|---|---|---|---|---|---|
| 1158 | 904.578 | 2 | 78.62 | 0.0711 | 32.36 | 92.6 | 3 | 0.0 | 0 | R.VVSVLTVLHQDWLNGK.E | | 185-200 | CID |
| 1062 | 743.527 | 3 | 161.00 | 0.1197 | 30.73 | 90.15 | 7 | 0.0 | 1 | R.VVSVLTVLHQDWLNGKEYK.C | | 185-203 | CID |
| 997 | 937.602 | 2 | 146.62 | 0.1375 | 29.63 | 46.75 | 1 | 0.0 | 0 | K.TTPPVLDSDGSFFLYSK.L | | 276-292 | CID |

#### Protein 9: Immunoglobulin heavy constant gamma 4 OS=Homo sapiens GN=IGHG4 PE=1 SV=1

| | | | |
|---|---|---|---|
| **Accession:** | IGHG4_HUMAN | **Score:** | 221.0 |
| **Database:** | SwissProt | **Seq. Coverage [%]:** | 10.1 |
| **MW [kDa] / pI:** | 35.9 / 7.2 | **No. of Peptides:** | 3 |
| **Modification(s):** | Carbamidomethyl | | |

| | | | | | | | | | |
|---|---|---|---|---|---|---|---|---|---|
| 10 | 20 | 30 | 40 | 50 | 60 | 70 | 80 | 90 | 100 |
| ASTKGPSVFP | LAPCSRSTSE | STAALGCLVK | DYFPEPVTVS | WNSGALTSGV | HTFPAVLQSS | GLYSLSSVVT | VPSSSLGTKT | YTCNVDHKPS | NTKVDKRVES |
| 110 | 120 | 130 | 140 | 150 | 160 | 170 | 180 | 190 | 200 |
| KYGPPCPSCP | APEFLGGPSV | FLFPPKPKDT | LMISRTPEVT | CVVVDVSQED | PEVQFNWYVD | GVEVHNAKTK | PREEQFNSTY | RVVSVLTVLH | QDWLNGKEYK |
| 210 | 220 | 230 | 240 | 250 | 260 | 270 | 280 | 290 | 300 |
| CKVSNKGLPS | SIEKTISKAK | GQPREPQVYT | LPPSQEEMTK | NQVSLTCLVK | GFYPSDIAVE | WESNGQPENN | YKTTPPVLDS | DGSFFLYSRL | TVDKSRWQEG |
| 310 | 320 | 330 | | | | | | | |
| NVFSCSVMHE | ALHNHYTQKS | LSLSLGK | | | | | | | |

MS/MS Peptide Matches

| Cmpd. | m/z meas. | z | Δ m/z [ppm] | Δ m/z [Da] | Rt [min] | Score | No. of Cmpds. | Site [%] | P | Sequence | Modification | Range | Type |
|---|---|---|---|---|---|---|---|---|---|---|---|---|---|
| 50 | 832.987 | 2 | 118.28 | 0.0985 | 11.39 | 38.23 | 4 | 0.0 | 0 | K.TYTCNVDHKPSNTK.V | Carbamidomethyl: 4 | 80-93 | CID |
| 1158 | 904.578 | 2 | 78.62 | 0.0711 | 32.36 | 92.6 | 3 | 0.0 | 0 | R.VVSVLTVLHQDWLNGK.E | | 182-197 | CID |
| 1062 | 743.527 | 3 | 161.00 | 0.1197 | 30.73 | 90.15 | 7 | 0.0 | 1 | R.VVSVLTVLHQDWLNGKEYK.C | | 182-200 | CID |

#### Protein 10: Keratin, type I cytoskeletal 9 OS=Homo sapiens GN=KRT9 PE=1 SV=3

| | | | |
|---|---|---|---|
| **Accession:** | K1C9_HUMAN | **Score:** | 157.5 |
| **Database:** | SwissProt | **Seq. Coverage [%]:** | 3.7 |
| **MW [kDa] / pI:** | 62.0 / 5.1 | **No. of Peptides:** | 1 |

| | | | | | | | | | |
|---|---|---|---|---|---|---|---|---|---|
| 10 | 20 | 30 | 40 | 50 | 60 | 70 | 80 | 90 | 100 |
| MSCRQFSSSY | LSRSGGGGGG | GLGSGGSIRS | SYSRFSSSGG | GGGGGRFSSS | SGYGGGSSRV | CGRGGGGSFG | YSYGGGSGGG | FSASSLGGGF | GGGSRGFGGA |
| 110 | 120 | 130 | 140 | 150 | 160 | 170 | 180 | 190 | 200 |
| SGGGYSSSGG | FGGGFGGGSG | GGFGGGYGSG | FGGFGGFGGG | AGGGDGGILT | ANEKSTMQEL | NSRLASYLDK | VQALEEANND | LENKIQDWYD | KKGPAAIQKN |
| 210 | 220 | 230 | 240 | 250 | 260 | 270 | 280 | 290 | 300 |
| YSPYYNTIDD | LKDQIVDLTV | GNNKTLLDID | NTRMTLDDFR | IKFEMEQNLR | QGVDADINGL | RQVLDNLTME | KSDLEMQYET | LQEELMALKK | NHKEEMSQLT |
| 310 | 320 | 330 | 340 | 350 | 360 | 370 | 380 | 390 | 400 |
| GQNSGDVNVE | INVAPGKDLT | KTLNDMRQEY | EQLIAKNRKD | IENQYETQIT | QIEHEVSSSG | QEVQSSAKEV | TQLRHGVQEL | EIELQSQLSK | KAALEKSLED |
| 410 | 420 | 430 | 440 | 450 | 460 | 470 | 480 | 490 | 500 |
| TKNRYCGQLQ | MIQEQISNLE | AQITDVRQEI | ECQNQEYSLL | LSIKMRLEKE | IETYHNLLEG | GQEDFESSGA | GKIGLGGRGG | SGGSYGRGSR | GGSGGSYGGG |
| 510 | 520 | 530 | 540 | 550 | 560 | 570 | 580 | 590 | 600 |
| GSGGGYGGGS | GSRGGSGGSY | GGGSGSGGGS | GGGYGGGSGG | GHSGGSGGGH | SGGSGGNYGG | GSGSGGGSGG | GYGGGSGSRG | GSGGSHGGGS | GFGGESGGSY |
| 610 | 620 | 630 | | | | | | | |
| GGGEEASGSG | GGYGGGSGKS | SHS | | | | | | | |

MS/MS Peptide Matches

| Cmpd. | m/z meas. | z | Δ m/z [ppm] | Δ m/z [Da] | Rt [min] | Score | No. of Cmpds. | Site [%] | P | Sequence | Modification | Range | Type |
|---|---|---|---|---|---|---|---|---|---|---|---|---|---|
| 65 | 896.445 | 2 | 86.47 | 0.0775 | 12.10 | 157.5 | 7 | 0.0 | 0 | R.GGSGGSYGGGGSGGGYGGGSGSR.G | | 491-513 | CID |

#### Protein 11: Immunoglobulin heavy constant gamma 2 OS=Homo sapiens GN=IGHG2 PE=1 SV=2

**Accession:** IGHG2_HUMAN
**Database:** SwissProt
**MW [kDa] / pI:** 35.9 / 7.7
**Modification(s):** Oxidation

**Score:** 136.8
**Seq. Coverage [%]:** 10.1
**No. of Peptides:** 3

| 10 | 20 | 30 | 40 | 50 | 60 | 70 | 80 | 90 | 100 |
|---|---|---|---|---|---|---|---|---|---|
| ASTKGPSVFP | LAPCSRSTSE | STAALGCLVK | DYFPEPVTVS | WNSGALTSGV | HTFPAVLQSS | GLYSLSSVVT | VPSSNFGTQT | YTCNVDHKPS | NTKVDKTVER |
| 110 | 120 | 130 | 140 | 150 | 160 | 170 | 180 | 190 | 200 |
| KCCVECPPCP | APPVAGPSVF | LFPPKPKDTL | MISRTPEVTC | VVVDVSHEDP | EVQFNWYVDG | VEVHNAKTKP | REEQFNSTFR | VVSVLTVVHQ | DWLNGKEYKC |
| 210 | 220 | 230 | 240 | 250 | 260 | 270 | 280 | 290 | 300 |
| KVSNKGLPAP | IEKTISKTKG | QPREPQVYTL | PPSREEMTKN | QVSLTCLVKG | FYPSDISVEW | ESNGQPENNY | KTTPPMLDSD | GSFFLYSKLT | VDKSRWQQGN |
| 310 | 320 | 330 | | | | | | | |
| VFSCSVMHEA | LHNHYTQKSL | SLSPGK | | | | | | | |

MS/MS Peptide Matches

| Cmpd. | m/z meas. | z | Δ m/z [ppm] | Δ m/z [Da] | Rt [min] | Score | No. of Cmpds. | Site [%] | P | Sequence | Modification | Range | Type |
|---|---|---|---|---|---|---|---|---|---|---|---|---|---|
| 519 | 953.012 | 2 | 39.08 | 0.0372 | 21.13 | 58.07 | 13 | 0.0 | 1 | R.EPQVYTLPPSREEMTK.N | | 224-239 | CID |
| 457 | 961.021 | 2 | 50.77 | 0.0488 | 19.92 | 28.13 | 1 | 0.0 | 1 | R.EPQVYTLPPSREEMTK.N | Oxidation: 14 | 224-239 | CID |
| 1024 | 953.528 | 2 | 81.20 | 0.0774 | 30.07 | 50.62 | 2 | 0.0 | 0 | K.TTPPMLDSDGSFFLYSK.L | | 272-288 | CID |

#### Protein 12: Keratin, type I cytoskeletal 10 OS=Homo sapiens GN=KRT10 PE=1 SV=6

**Accession:** K1C10_HUMAN
**Database:** SwissProt
**MW [kDa] / pI:** 58.8 / 5.1

**Score:** 133.0
**Seq. Coverage [%]:** 3.3
**No. of Peptides:** 1

| 10 | 20 | 30 | 40 | 50 | 60 | 70 | 80 | 90 | 100 |
|---|---|---|---|---|---|---|---|---|---|
| MSVRYSSSKH | YSSSRSGGGG | GGGGCGGGGG | VSSLRISSSK | GSLGGGFSSG | GFSGGSFSRG | SSGGGCFGGS | SGGYGGLGGF | GGGSFRGSYG | SSSFGGSYGG |
| **110** | **120** | **130** | **140** | **150** | **160** | **170** | **180** | **190** | **200** |
| IFGGGSFGGG | SFGGGSFGGG | GFGGGGFGGG | FGGGFGGDGG | LLSGNEKVTM | QNLNDRLASY | LDKVRALEES | NYELEGKIKE | WYEKHGNSHQ | GEPRDYSKYY |
| **210** | **220** | **230** | **240** | **250** | **260** | **270** | **280** | **290** | **300** |
| KTIDDLKNQI | LNLTTDNANI | LLQIDNARLA | ADDFRLKYEN | EVALRQSVEA | DINGLRRVLD | ELTLTKADLE | MQIESLTEEL | AYLKKNHEEE | MKDLRNVSTG |
| **310** | **320** | **330** | **340** | **350** | **360** | **370** | **380** | **390** | **400** |
| DVNVEMNAAP | GVDLTQLLNN | MRSQYEQLAE | QNRKDAEAWF | NEKSKELTTE | IDNNIEQISS | YKSEITELRR | NVQALEIELQ | SQLALKQSLE | ASLAETEGRY |
| **410** | **420** | **430** | **440** | **450** | **460** | **470** | **480** | **490** | **500** |
| CVQLSQIQAQ | ISALEEQLQQ | IRAETECQNT | EYQQLLDIKI | RLENEIQTYR | SLLEGEGSSG | GGGRGGGSFG | GGYGGGSSGG | GSSGGGHGGG | HGGSSGGGYG |
| **510** | **520** | **530** | **540** | **550** | **560** | **570** | **580** | **590** | |
| GGSSGGGSSG | GGYGGGSSSG | GHGGSSSGGY | GGGSSGGGGG | GYGGGSSGGG | SSSGGGYGGG | SSSGGHKSSS | SGSVGESSSK | GPRY | |

MS/MS Peptide Matches

| Cmpd. | m/z meas. | z | Δ m/z [ppm] | Δ m/z [Da] | Rt [min] | Score | No. of Cmpds. | Site [%] | P | Sequence | Modification | Range | Type |
|---|---|---|---|---|---|---|---|---|---|---|---|---|---|
| 660 | 854.5 | 2 | 129.09 | 0.1103 | 23.79 | 133.01 | 3 | 0.0 | 0 | K.GSLGGGFSSGGFSGGSFSR.G | | 41-59 | CID |

#### Protein 13: Immunoglobulin kappa variable 3-20 OS=Homo sapiens GN=IGKV3-20 PE=1 SV=2

**Accession:** KV320_HUMAN
**Database:** SwissProt
**MW [kDa] / pI:** 12.5 / 4.9

**Score:** 117.1
**Seq. Coverage [%]:** 19.0
**No. of Peptides:** 1

| 10 | 20 | 30 | 40 | 50 | 60 | 70 | 80 | 90 | 100 |
|---|---|---|---|---|---|---|---|---|---|
| METPAQLLFL | LLLWLPDTTG | EIVLTQSPGT | LSLSPGERAT | LSCRASQSVS | SSYLAWYQQK | PGQAPRLLIY | GASSRATGIP | DRFSGSGSGT | DFTLTISRLE |
| **110** | **120** | | | | | | | | |
| PEDFAVYYCQ | QYGSSP | | | | | | | | |

MS/MS Peptide Matches

| Cmpd. | m/z meas. | z | Δ m/z [ppm] | Δ m/z [Da] | Rt [min] | Score | No. of Cmpds. | Site [%] | P | Sequence | Modification | Range | Type |
|---|---|---|---|---|---|---|---|---|---|---|---|---|---|
| 671 | 813.834 | 3 | 115.64 | 0.0941 | 23.97 | 117.11 | 3 | 0.0 | 0 | R.ASQSVSSSYLAWYQQKPGQAPR.L | | 45-66 | CID |

##### Protein 14: Immunoglobulin kappa variable 3-15 OS=Homo sapiens GN=IGKV3-15 PE=1 SV=2

**Accession:** KV315_HUMAN
**Database:** SwissProt
**MW [kDa] / pI:** 12.5 / 5.1

**Score:** 88.1
**Seq. Coverage [%]:** 18.3
**No. of Peptides:** 1

```
        10         20         30         40         50         60         70         80         90        100
MEAPAQLLFL LLLWLPDTTG EIVMTQSPAT LSVSPGERAT LSCRASQSVS SNLAWYQQKP GQAPRLLIYG ASTRATGIPA RFSGSGSGTE FTLTISSLQS
       110        120
EDFAVYYCQQ YNNWP
```

MS/MS Peptide Matches

| Cmpd. | m/z meas. | z | Δ m/z [ppm] | Δ m/z [Da] | Rt [min] | Score | No. of Cmpds. | Site [%] | P | Sequence | Modification | Range | Type |
|---|---|---|---|---|---|---|---|---|---|---|---|---|---|
| 537 | 768.477 | 3 | 114.41 | 0.0879 | 21.45 | 88.06 | 3 | 0.0 | 0 | R.ASQSVSSNLAWYQQKPGQAPR.L | | 45-65 | CID |

##### Protein 15: Immunoglobulin heavy variable 4-4 OS=Homo sapiens GN=IGHV4-4 PE=3 SV=2

**Accession:** HV404_HUMAN
**Database:** SwissProt
**MW [kDa] / pI:** 12.8 / 9.6
**Modification(s):** Carbamidomethyl

**Score:** 74.3
**Seq. Coverage [%]:** 13.7
**No. of Peptides:** 1

| 10 | 20 | 30 | 40 | 50 | 60 | 70 | 80 | 90 | 100 |
|---|---|---|---|---|---|---|---|---|---|
| MKHLWFFLLL | VAAPRWVLSQ | VQLQESGPGL | VKPSGTLSLT | CAVSGGSISS | SNWWSWVRQP | PGKGLEWIGE | IYHSGSTNYN | PSLKSRVTIS | VDKSKNQFSL |
| 110 | 120 | | | | | | | | |
| KLSSVTAADT | AVYYCAR | | | | | | | | |

MS/MS Peptide Matches

| Cmpd. | m/z meas. | z | Δ m/z [ppm] | Δ m/z [Da] | Rt [min] | Score | No. of Cmpds. | Site [%] | P | Sequence | Modification | Range | Type |
|---|---|---|---|---|---|---|---|---|---|---|---|---|---|
| 581 | 874.504 | 2 | 96.50 | 0.0844 | 22.36 | 74.25 | 3 | 0.0 | 0 | K.LSSVTAADTAVYYCAR.- | Carbamidomethyl: 14 | 102-117 | CID |

#### Protein 16: Immunoglobulin heavy variable 5-51 OS=Homo sapiens GN=IGHV5-51 PE=3 SV=1

**Accession:** HV551_HUMAN
**Database:** SwissProt
**MW [kDa] / pI:** 12.7 / 8.4
**Score:** 73.8
**Seq. Coverage [%]:** 12.8
**No. of Peptides:** 1

| 10 | 20 | 30 | 40 | 50 | 60 | 70 | 80 | 90 | 100 |
|---|---|---|---|---|---|---|---|---|---|
| MGSTAILALL | LAVLQGVCAE | VQLVQSGAEV | KKPGESLKIS | CKGSGYSFTS | YWIGWVRQMP | GKGLEWMGII | YPGDSDTRYS | PSFQGQVTIS | ADKSISTAYL |
| 110 | 120 | | | | | | | | |
| QWSSLKASDT | AMYYCAR | | | | | | | | |

MS/MS Peptide Matches

| Cmpd. | m/z meas. | z | Δ m/z [ppm] | Δ m/z [Da] | Rt [min] | Score | No. of Cmpds. | Site [%] | P | Sequence | Modification | Range | Type |
|---|---|---|---|---|---|---|---|---|---|---|---|---|---|
| 554 | 814.452 | 2 | 61.70 | 0.0502 | 21.73 | 73.75 | 3 | 0.0 | 0 | R.YSPSFQGQVTISADK.S | | 79-93 | CID |

#### Project Info

| | | | |
|---|---|---|---|
| **Name:** | Alagesan | **Date:** | Mar 5, 2019 |
| **Note:** | Nanobooster | | |

#### Sample Info & Protocols

| | |
|---|---|
| **Name:** | Nanobooster - IgG |
| **Date:** | Mar 5, 2019 |

#### Search Result Info

| | | | |
|---|---|---|---|
| **Search Result:** | IonTrap_Human_Swiss_CID_2019-03-05 17:55:08 | | |
| **Location:** | /Alagesan/Nanobooster - IgG/KA_IgG3 HD NB ACN CID_GA4_01_9084.mgf | | |
| **Search Method:** | IonTrap_Human_Swiss_CID | **Version:** | 1.2 |
| **Search Engine(s):** | Mascot, 2.6.0 | **Ident. Compound(s):** | 0/2370 |
| **Database(s):** | SwissProt, SwissProt_2017_04.fasta | | |
| **Note:** | | | |

#### Results

##### Protein 1: Immunoglobulin heavy constant gamma 3 OS=Homo sapiens GN=IGHG3 PE=1 SV=2

| | | | |
|---|---|---|---|
| **Accession:** | IGHG3_HUMAN | **Score:** | 1323.2 |
| **Database:** | SwissProt | **Seq. Coverage [%]:** | 47.7 |
| **MW [kDa] / pI:** | 41.3 / 8.2 | **No. of Peptides:** | 18 |
| **Modification(s):** | Carbamidomethyl, Oxidation | | |

```
        10         20         30         40         50         60         70         80         90        100
ASTKGPSVFP LAPCSRSTSG GTAALGCLVK DYFPEPVTVS WNSGALTSGV HTFPAVLQSS GLYSLSSVVT VPSSSLGTQT YTCNVNHKPS NTKVDKRVEL
       110        120        130        140        150        160        170        180        190        200
KTPLGDTTHT CPRCPEPKSC DTPPPCPRCP EPKSCDTPPP CPRCPEPKSC DTPPPCPRCP APELLGGPSV FLFPPKPKDT LMISRTPEVT CVVVDVSHED
       210        220        230        240        250        260        270        280        290        300
PEVQFKWYVD GVEVHNAKTK PREEQYNSTF RVVSVLTVLH QDWLNGKEYK CKVSNKALPA PIEKTISKTK GQPREPQVYT LPPSREEMTK NQVSLTCLVK
       310        320        330        340        350        360        370        380
GFYPSDIAVE WESSGQPENN YNTTPPMLDS DGSFFLYSKL TVDKSRWQQG NIFSCSVMHE ALHNRFTQKS LSLSPGK
```

MS/MS Peptide Matches

| Cmpd. | m/z meas. | z | Δ m/z [ppm] | Δ m/z [Da] | Rt [min] | Score | No. of Cmpds. | Site [%] | P | Sequence | Modification | Range | Type |
|---|---|---|---|---|---|---|---|---|---|---|---|---|---|
| 934 | 864.202 | 3 | 107.95 | 0.0933 | 28.44 | 81.06 | 3 | 0.0 | 1 | K.GPSVFPLAPCSRSTSGGTAALGCLVK.D | Carbamidomethyl: 10, 23 | 5-30 | CID |
| 443 | 913.017 | 2 | 54.47 | 0.0497 | 19.58 | 88.04 | 7 | 0.0 | 1 | R.VELKTPLGDTTHTCPR.C | Carbamidomethyl: 14 | 98-113 | CID |
| 215 | 984.025 | 2 | 66.96 | 0.0659 | 15.13 | 50.19 | 3 | 0.0 | 1 | K.TPLGDTTHTCPRCPEPK.S | Carbamidomethyl: 10, 13 | 102-118 | CID |
| 168 | 899.469 | 2 | 88.41 | 0.0795 | 14.24 | 67.85 | 14 | 0.0 | 1 | R.CPEPKSCDTPPPCPR.C | Carbamidomethyl: 1, 7, 13 | 114-128 | CID |
| 198 | 899.434 | 2 | 49.50 | 0.0445 | 14.83 | 24.73 | 2 | 0.0 | 1 | K.SCDTPPPCPRCPEPK.S | Carbamidomethyl: 2, 8, 11 | 119-133 | CID |
| 1139 | 1076.629 | 2 | 503.50 | 0.5418 | 31.92 | 108.43 | 18 | 0.0 | 0 | R.CPAPELLGGPSVFLFPPKPK.D | Carbamidomethyl: 1 | 159-178 | CID |
| 1163 | 742.985 | 4 | 449.30 | 0.3337 | 32.34 | 96.58 | 5 | 0.0 | 1 | R.CPAPELLGGPSVFLFPPKPKDTLMISR.T | Carbamidomethyl: 1 | 159-185 | CID |

| Cmpd. | m/z meas. | z | Δ m/z [ppm] | Δ m/z [Da] | Rt [min] | Score | No. of Cmpds. | Site [%] | P | Sequence | Modification | Range | Type |
|---|---|---|---|---|---|---|---|---|---|---|---|---|---|
| 1125 | 747.01 | 4 | 482.08 | 0.3599 | 31.69 | 71.12 | 3 | 0.0 | 1 | R.CPAPELLGGPSVFLFPPKPKDTLMISR.T | Carbamidomethyl: 1; Oxidation: 24 | 159-185 | CID |
| 1018 | 1077.928 | 3 | 370.80 | 0.3996 | 29.88 | 75.94 | 3 | 0.0 | 1 | K.DTLMISRTPEVTCVVVDVSHEDPEVQFK.W | Carbamidomethyl: 13 | 179-206 | CID |
| 890 | 805.522 | 3 | 164.32 | 0.1323 | 27.47 | 126.98 | 4 | 0.0 | 0 | R.TPEVTCVVVDVSHEDPEVQFK.W | Carbamidomethyl: 6 | 186-206 | CID |
| 1055 | 1271.608 | 3 | 257.54 | 0.3274 | 30.46 | 56.71 | 3 | 0.0 | 1 | R.TPEVTCVVVDVSHEDPEVQFKWYVDGVEVHNA K.T | Carbamidomethyl: 6 | 186-218 | CID |
| 518 | 708.94 | 2 | 128.44 | 0.0910 | 20.99 | 84.47 | 20 | 0.0 | 0 | K.WYVDGVEVHNAK.T | | 207-218 | CID |
| 1167 | 905.096 | 2 | 651.31 | 0.5891 | 32.40 | 74.04 | 3 | 0.0 | 0 | R.VVSVLTVLHQDWLNGK.E | | 232-247 | CID |
| 1067 | 743.505 | 3 | 131.41 | 0.0977 | 30.72 | 92.84 | 3 | 0.0 | 1 | R.VVSVLTVLHQDWLNGKEYK.C | | 232-250 | CID |
| 479 | 953.055 | 2 | 84.20 | 0.0802 | 20.29 | 62.8 | 13 | 0.0 | 1 | R.EPQVYTLPPSREEMTK.N | | 275-290 | CID |
| 444 | 961.022 | 2 | 51.81 | 0.0498 | 19.59 | 34.56 | 3 | 0.0 | 1 | R.EPQVYTLPPSREEMTK.N | Oxidation: 14 | 275-290 | CID |
| 779 | 1158.039 | 2 | 436.41 | 0.5052 | 25.62 | 74.9 | 4 | 0.0 | 0 | R.WQQGNIFSCSVMHEALHNR.F | Carbamidomethyl: 9 | 347-365 | CID |
| 824 | 940.513 | 3 | 423.49 | 0.3981 | 26.40 | 52.0 | 3 | 0.0 | 1 | R.WQQGNIFSCSVMHEALHNRFTQK.S | Carbamidomethyl: 9 | 347-369 | CID |

#### Protein 2: Immunoglobulin kappa constant OS=Homo sapiens GN=IGKC PE=1 SV=2

**Accession:** IGKC_HUMAN
**Database:** SwissProt
**MW [kDa] / pI:** 11.8 / 6.1
**Modification(s):** Carbamidomethyl

**Score:** 1011.7
**Seq. Coverage [%]:** 86.9
**No. of Peptides:** 10

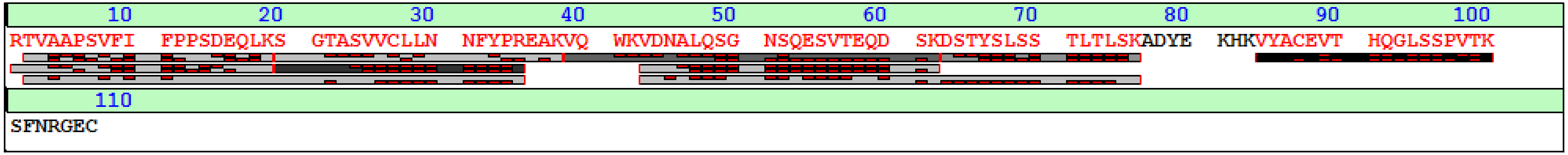


MS/MS Peptide Matches

| Cmpd. | m/z meas. | z | Δ m/z [ppm] | Δ m/z [Da] | Rt [min] | Score | No. of Cmpds. | Site [%] | P | Sequence | Modification | Range | Type |
|---|---|---|---|---|---|---|---|---|---|---|---|---|---|
| 943 | 1051.684 | 2 | 110.62 | 0.1163 | 28.59 | 39.95 | 5 | 0.0 | 1 | -.RTVAAPSVFIFPPSDEQLK.S | | 1-19 | CID |
| 1065 | 973.643 | 2 | 129.31 | 0.1259 | 30.67 | 81.14 | 19 | 0.0 | 0 | R.TVAAPSVFIFPPSDEQLK.S | | 2-19 | CID |
| 1521 | 1242.688 | 3 | 307.25 | 0.3817 | 38.38 | 83.91 | 9 | 0.0 | 1 | R.TVAAPSVFIFPPSDEQLKSGTASVVCLLNNFYPR.E | Carbamidomethyl: 26 | 2-35 | CID |
| 1209 | 899.566 | 2 | 127.57 | 0.1147 | 33.09 | 65.86 | 13 | 0.0 | 0 | K.SGTASVVCLLNNFYPR.E | Carbamidomethyl: 8 | 20-35 | CID |
| 1097 | 709.416 | 3 | 76.85 | 0.0545 | 31.25 | 107.74 | 3 | 0.0 | 1 | K.SGTASVVCLLNNFYPREAK.V | Carbamidomethyl: 8 | 20-38 | CID |
| 522 | 893.195 | 3 | 112.14 | 0.1002 | 21.08 | 127.92 | 6 | 0.0 | 1 | K.VQWKVDNALQSGNSQESVTEQDSK.D | | 39-62 | CID |
| 257 | 1068.569 | 2 | 75.81 | 0.0810 | 15.90 | 169.9 | 10 | 0.0 | 0 | K.VDNALQSGNSQESVTEQDSK.D | | 43-62 | CID |
| 716 | 1207.568 | 3 | 270.62 | 0.3267 | 24.56 | 151.41 | 4 | 0.0 | 1 | K.VDNALQSGNSQESVTEQDSKDSTYSLSSTLTLSK.A | | 43-76 | CID |
| 725 | 751.963 | 2 | 106.59 | 0.0801 | 24.76 | 89.21 | 5 | 0.0 | 0 | K.DSTYSLSSTLTLSK.A | | 63-76 | CID |
| 415 | 938.598 | 2 | 139.48 | 0.1309 | 19.08 | 94.67 | 9 | 0.0 | 0 | K.VYACEVTHQGLSSPVTK.S | Carbamidomethyl: 4 | 84-100 | CID |

#### Protein 3: Immunoglobulin kappa light chain OS=Homo sapiens PE=1 SV=1

**Accession:** IGK_HUMAN
**Database:** SwissProt
**MW [kDa] / pI:** 23.4 / 6.9
**Modification(s):** Carbamidomethyl

**Score:** 905.5
**Seq. Coverage [%]:** 43.0
**No. of Peptides:** 8

| 10 | 20 | 30 | 40 | 50 | 60 | 70 | 80 | 90 | 100 |
|---|---|---|---|---|---|---|---|---|---|
| DIQMTQSPST | LSASVGDRVT | ITCRASQSIN | TWLAWYQQKP | GKAPKLLMYK | ASSLESGVPS | RFIGSGSGTE | FTLTISSLQP | DDFATYYCQQ | YNSDSKMFGQ |
| **110** | **120** | **130** | **140** | **150** | **160** | **170** | **180** | **190** | **200** |
| GTKVEVKGTV | AAPSVFIFPP | SDEQLKSGTA | SVVCLLNNFY | PREAKVQWKV | DNALQSGNSQ | ESVTEQDSKD | STYSLSSTLT | LSKADYEKHK | VYACEVTHQG |
| **210** | **220** | | | | | | | | |
| LSSPVTKSFN | RGEC | | | | | | | | |

MS/MS Peptide Matches

| Cmpd. | m/z meas. | z | Δ m/z [ppm] | Δ m/z [Da] | Rt [min] | Score | No. of Cmpds. | Site [%] | P | Sequence | Modification | Range | Type |
|---|---|---|---|---|---|---|---|---|---|---|---|---|---|
| 586 | 946.994 | 2 | 41.66 | 0.0394 | 22.25 | 98.77 | 3 | 0.0 | 0 | -.DIQMTQSPSTLSASVGDR.V | | 1-18 | CID |
| 1209 | 899.566 | 2 | 127.57 | 0.1147 | 33.09 | 65.86 | 13 | 0.0 | 0 | K.SGTASVVCLLNNFYPR.E | Carbamidomethyl: 8 | 127-142 | CID |
| 1097 | 709.416 | 3 | 76.85 | 0.0545 | 31.25 | 107.74 | 3 | 0.0 | 1 | K.SGTASVVCLLNNFYPREAK.V | Carbamidomethyl: 8 | 127-145 | CID |
| 522 | 893.195 | 3 | 112.14 | 0.1002 | 21.08 | 127.92 | 6 | 0.0 | 1 | K.VQWKVDNALQSGNSQESVTEQDSK.D | | 146-169 | CID |
| 257 | 1068.569 | 2 | 75.81 | 0.0810 | 15.90 | 169.9 | 10 | 0.0 | 0 | K.VDNALQSGNSQESVTEQDSK.D | | 150-169 | CID |
| 716 | 1207.568 | 3 | 270.62 | 0.3267 | 24.56 | 151.41 | 4 | 0.0 | 1 | K.VDNALQSGNSQESVTEQDSKDSTYSLSSTLTLSK.A | | 150-183 | CID |
| 725 | 751.963 | 2 | 106.59 | 0.0801 | 24.76 | 89.21 | 5 | 0.0 | 0 | K.DSTYSLSSTLTLSK.A | | 170-183 | CID |
| 415 | 938.598 | 2 | 139.48 | 0.1309 | 19.08 | 94.67 | 9 | 0.0 | 0 | K.VYACEVTHQGLSSPVTK.S | Carbamidomethyl: 4 | 191-207 | CID |

#### Protein 4: Immunoglobulin lambda constant 2 OS=Homo sapiens GN=IGLC2 PE=1 SV=1

**Accession:** IGLC2_HUMAN
**Database:** SwissProt
**MW [kDa] / pI:** 11.3 / 6.9
**Modification(s):** Carbamidomethyl

**Score:** 694.0
**Seq. Coverage [%]:** 77.4
**No. of Peptides:** 9

10 20 30 40 50 60 70 80 90 100
GQPKAAPSVT LFPPSSEELQ ANKATLVCLI SDFYPGAVTV AWKADSSPVK AGVETTTPSK QSNNKYAASS YLSLTPEQWK SHRSYSCQVT HEGSTVEKTV

110
APTECS

MS/MS Peptide Matches

| Cmpd. | m/z meas. | z | Δ m/z [ppm] | Δ m/z [Da] | Rt [min] | Score | No. of Cmpds. | Site [%] | P | Sequence | Modification | Range | Type |
|---|---|---|---|---|---|---|---|---|---|---|---|---|---|
| 850 | 993.565 | 2 | 52.78 | 0.0524 | 26.81 | 75.06 | 13 | 0.0 | 0 | K.AAPSVTLFPPSSEELQANK.A | | 5-23 | CID |
| 213 | 838.017 | 2 | 102.79 | 0.0861 | 15.08 | 53.7 | 3 | 0.0 | 1 | K.ADSSPVKAGVETTTPSK.Q | | 44-60 | CID |
| 35 | 781.498 | 2 | 132.51 | 0.1035 | 10.90 | 81.24 | 16 | 0.0 | 1 | K.AGVETTTPSKQSNNK.Y | | 51-65 | CID |

| Cmpd. | m/z meas. | z | Δ m/z [ppm] | Δ m/z [Da] | Rt [min] | Score | No. of Cmpds. | Site [%] | P | Sequence | Modification | Range | Type |
|---|---|---|---|---|---|---|---|---|---|---|---|---|---|
| 814 | 772.503 | 3 | 157.17 | 0.1214 | 26.23 | 72.76 | 5 | 0.0 | 1 | K.QSNNKYAASSYLSLTPEQWK.S | | 61-80 | CID |
| 901 | 872.522 | 2 | 101.95 | 0.0889 | 27.69 | 85.72 | 12 | 0.0 | 0 | K.YAASSYLSLTPEQWK.S | | 66-80 | CID |
| 714 | 708.788 | 3 | 140.43 | 0.0995 | 24.52 | 73.19 | 1 | 0.0 | 1 | K.YAASSYLSLTPEQWKSHR.S | | 66-83 | CID |
| 84 | 1046.572 | 2 | 88.62 | 0.0927 | 12.44 | 47.66 | 5 | 0.0 | 1 | K.SHRSYSCQVTHEGSTVEK.T | Carbamidomethyl: 7 | 81-98 | CID |
| 159 | 856.507 | 2 | 144.52 | 0.1238 | 14.06 | 108.39 | 8 | 0.0 | 0 | R.SYSCQVTHEGSTVEK.T | Carbamidomethyl: 4 | 84-98 | CID |
| 399 | 1279.556 | 2 | 385.68 | 0.4933 | 18.78 | 96.24 | 6 | 0.0 | 1 | R.SYSCQVTHEGSTVEKTVAPTECS.- | Carbamidomethyl: 4, 22 | 84-106 | CID |

#### Protein 5: Immunoglobulin lambda constant 3 OS=Homo sapiens GN=IGLC3 PE=1 SV=1

**Accession:** IGLC3_HUMAN
**Database:** SwissProt
**MW [kDa] / pI:** 11.3 / 6.9
**Modification(s):** Carbamidomethyl

**Score:** 668.7
**Seq. Coverage [%]:** 77.4
**No. of Peptides:** 8

| 10 | 20 | 30 | 40 | 50 | 60 | 70 | 80 | 90 | 100 |
|---|---|---|---|---|---|---|---|---|---|
| GQPKAAPSVT | LFPPSSEELQ | ANKATLVCLI | SDFYPGAVTV | AWKADSSPVK | AGVETTTPSK | QSNNKYAASS | YLSLTPEQWK | SHKSYSCQVT | HEGSTVEKTV |

| 110 |
|---|
| APTECS |

MS/MS Peptide Matches

| Cmpd. | m/z meas. | z | Δ m/z [ppm] | Δ m/z [Da] | Rt [min] | Score | No. of Cmpds. | Site [%] | P | Sequence | Modification | Range | Type |
|---|---|---|---|---|---|---|---|---|---|---|---|---|---|
| 850 | 993.565 | 2 | 52.78 | 0.0524 | 26.81 | 75.06 | 13 | 0.0 | 0 | K.AAPSVTLFPPSSEELQANK.A | | 5-23 | CID |
| 213 | 838.017 | 2 | 102.79 | 0.0861 | 15.08 | 53.7 | 3 | 0.0 | 1 | K.ADSSPVKAGVETTTPSK.Q | | 44-60 | CID |
| 35 | 781.498 | 2 | 132.51 | 0.1035 | 10.90 | 81.24 | 16 | 0.0 | 1 | K.AGVETTTPSKQSNNK.Y | | 51-65 | CID |
| 814 | 772.503 | 3 | 157.17 | 0.1214 | 26.23 | 72.76 | 5 | 0.0 | 1 | K.QSNNKYAASSYLSLTPEQWK.S | | 61-80 | CID |
| 901 | 872.522 | 2 | 101.95 | 0.0889 | 27.69 | 85.72 | 12 | 0.0 | 0 | K.YAASSYLSLTPEQWK.S | | 66-80 | CID |

| Cmpd. | m/z meas. | z | Δ m/z [ppm] | Δ m/z [Da] | Rt [min] | Score | No. of Cmpds. | Site [%] | P | Sequence | Modification | Range | Type |
|---|---|---|---|---|---|---|---|---|---|---|---|---|---|
| 77 | 1032.604 | 2 | 123.80 | 0.1278 | 12.27 | 95.57 | 7 | 0.0 | 1 | K.SHKSYSCQVTHEGSTVEK.T | Carbamidomethyl: 7 | 81-98 | CID |
| 159 | 856.507 | 2 | 144.52 | 0.1238 | 14.06 | 108.39 | 8 | 0.0 | 0 | K.SYSCQVTHEGSTVEK.T | Carbamidomethyl: 4 | 84-98 | CID |
| 399 | 1279.556 | 2 | 385.68 | 0.4933 | 18.78 | 96.24 | 6 | 0.0 | 1 | K.SYSCQVTHEGSTVEKTVAPTECS.- | Carbamidomethyl: 4, 22 | 84-106 | CID |

#### Protein 6: Keratin, type II cytoskeletal 1 OS=Homo sapiens GN=KRT1 PE=1 SV=6

**Accession:** K2C1_HUMAN
**Database:** SwissProt
**MW [kDa] / pI:** 66.0 / 8.2

**Score:** 657.9
**Seq. Coverage [%]:** 19.7
**No. of Peptides:** 7

| | | | | | | | | | |
|---|---|---|---|---|---|---|---|---|---|
| 10 | 20 | 30 | 40 | 50 | 60 | 70 | 80 | 90 | 100 |
| MSRQFSSRSG | YRSGGGFSSG | SAGIINYQRR | TTSSSTRRSG | GGGGRFSSCG | GGGGSFGAGG | GFGSRSLVNL | GGSKSISISV | ARGGGRGSGF | GGGYGGGGFG |
| 110 | 120 | 130 | 140 | 150 | 160 | 170 | 180 | 190 | 200 |
| GGGFGGGGFG | GGGIGGGGFG | GFGSGGGGFG | GGGFGGGGYG | GGYGPVCPPG | GIQEVTINQS | LLQPLNVEID | PEIQKVKSRE | REQIKSLNNQ | FASFIDKVRF |
| 210 | 220 | 230 | 240 | 250 | 260 | 270 | 280 | 290 | 300 |
| LEQQNQVLQT | KWELLQQVDT | STRTHNLEPY | FESFINNLRR | RVDQLKSDQS | RLDSELKNMQ | DMVEDYRNKY | EDEINKRTNA | ENEFVTIKKD | VDGAYMTKVD |
| 310 | 320 | 330 | 340 | 350 | 360 | 370 | 380 | 390 | 400 |
| LQAKLDNLQQ | EIDFLTALYQ | AELSQMQTQI | SETNVILSMD | NNRSLDLDSI | IAEVKAQYED | IAQKSKAEAE | SLYQSKYEEL | QITAGRHGDS | VRNSKIEISE |
| 410 | 420 | 430 | 440 | 450 | 460 | 470 | 480 | 490 | 500 |
| LNRVIQRLRS | EIDNVKKQIS | NLQQSISDAE | QRGENALKDA | KNKLNDLEDA | LQQAKEDLAR | LLRDYQELMN | TKLALDLEIA | TYRTLLEGEE | SRMSGECAPN |
| 510 | 520 | 530 | 540 | 550 | 560 | 570 | 580 | 590 | 600 |
| VSVSVSTSHT | TISGGGSRGG | GGGGYGSGGS | SYGSGGGSYG | SGGGGGGGRG | SYGSGGSSYG | SGGGSYGSGG | GGGGHGSYGS | GSSSGGYRGG | SGGGGGGSSG |
| 610 | 620 | 630 | 640 | 650 | | | | | |
| GRGSGGGSSG | GSIGGRGSSS | GGVKSSGGSS | SVKFVSTTYS | GVTR | | | | | |

MS/MS Peptide Matches

| Cmpd. | m/z meas. | z | Δ m/z [ppm] | Δ m/z [Da] | Rt [min] | Score | No. of Cmpds. | Site [%] | P | Sequence | Modification | Range | Type |
|---|---|---|---|---|---|---|---|---|---|---|---|---|---|
| 947 | 820.019 | 2 | 104.21 | 0.0854 | 28.64 | 60.66 | 2 | 0.0 | 1 | K.SLNNQFASFIDKVR.F | | 186-199 | CID |
| 462 | 738.525 | 2 | 174.33 | 0.1287 | 19.91 | 98.21 | 2 | 0.0 | 0 | R.FLEQQNQVLQTK.W | | 200-211 | CID |
| 995 | 978.58 | 3 | 412.04 | 0.4030 | 29.50 | 103.56 | 3 | 0.0 | 1 | R.FLEQQNQVLQTKWELLQQVDTSTR.T | | 200-223 | CID |
| 1492 | 783.77 | 3 | 32.77 | 0.0257 | 37.85 | 124.07 | 1 | 0.0 | 1 | R.SLDLDSIIAEVKAQYEDIAQK.S | | 344-364 | CID |
| 551 | 859.05 | 2 | 140.64 | 0.1208 | 21.57 | 81.75 | 4 | 0.0 | 0 | K.QISNLQQSISDAEQR.G | | 418-432 | CID |
| 178 | 1192.497 | 2 | 14.62 | 0.0174 | 14.43 | 169.91 | 3 | 0.0 | 0 | R.GGGGGGYGSGGSSYGSGGGSYGSGGGGGG GR.G | | 519-549 | CID |
| 46 | 876.504 | 2 | 99.29 | 0.0870 | 11.19 | 19.73 | 2 | 0.0 | 1 | R.GSGGGSSGGSIGGRGSSSGGVK.S | | 603-624 | CID |

#### Protein 7: Keratin, type II cytoskeletal 2 epidermal OS=Homo sapiens GN=KRT2 PE=1 SV=2

| | | | |
|---|---|---|---|
| **Accession:** | K22E_HUMAN | **Score:** | 228.1 |
| **Database:** | SwissProt | **Seq. Coverage [%]:** | 5.0 |
| **MW [kDa] / pI:** | 65.4 / 8.1 | **No. of Peptides:** | 2 |

| 10 | 20 | 30 | 40 | 50 | 60 | 70 | 80 | 90 | 100 |
|---|---|---|---|---|---|---|---|---|---|
| MSCQISCKSR | GRGGGGGGFR | GFSSGSAVVS | GGSRRSTSSF | SCLSRHGGGG | GGFGGGGFGS | RSLVGLGGTK | SISISVAGGG | GGFGAAGGFG | GRGGGFGGGS |
| 110 | 120 | 130 | 140 | 150 | 160 | 170 | 180 | 190 | 200 |
| SFGGGSGFSG | GGFGGGGFGG | GRFGGFGGPG | GVGGLGGPGG | FGPGGYPGGI | HEVSVNQSLL | QPLNVKVDPE | IQNVKAQERE | QIKTLNNKFA | SFIDKVRFLE |
| 210 | 220 | 230 | 240 | 250 | 260 | 270 | 280 | 290 | 300 |
| QQNQVLQTKW | ELLQQMNVGT | RPINLEPIFQ | GYIDSLKRYL | DGLTAERTSQ | NSELNNMQDL | VEDYKKKYED | EINKRTAAEN | DFVTLKKDVD | NAYMIKVELQ |
| 310 | 320 | 330 | 340 | 350 | 360 | 370 | 380 | 390 | 400 |
| SKVDLLNQEI | EFLKVLYDAE | ISQIHQSVTD | TNVILSMDNS | RNLDLDSIIA | EVKAQYEEIA | QRSKEEAEAL | YHSKYEELQV | TVGRHGDSLK | EIKIEISELN |
| 410 | 420 | 430 | 440 | 450 | 460 | 470 | 480 | 490 | 500 |
| RVIQRLQGEI | AHVKKQCKNV | QDAIADAEQR | GEHALKDARN | KLNDLEEALQ | QAKEDLARLL | RDYQELMNVK | LALDVEIATY | RKLLEGEECR | MSGDLSSNVT |
| 510 | 520 | 530 | 540 | 550 | 560 | 570 | 580 | 590 | 600 |
| VSVTSSTISS | NVASKAAFGG | SGGRGSSSGG | GYSSGSSSYG | SGGRQSGSRG | GSGGGGSISG | GGYGSGGGSG | GRYGSGGGSK | GGSISGGGYG | SGGGKHSSGG |
| 610 | 620 | 630 | 640 | | | | | | |
| GSRGGSSSGG | GYGSGGGGSS | SVKGSSGEAF | GSSVTFSFR | | | | | | |

MS/MS Peptide Matches

| Cmpd. | m/z meas. | z | Δ m/z [ppm] | Δ m/z [Da] | Rt [min] | Score | No. of Cmpds. | Site [%] | P | Sequence | Modification | Range | Type |
|---|---|---|---|---|---|---|---|---|---|---|---|---|---|
| 462 | 738.525 | 2 | 174.33 | 0.1287 | 19.91 | 98.21 | 2 | 0.0 | 0 | R.FLEQQNQVLQTK.W | | 198-209 | CID |
| 82 | 870.952 | 2 | 109.75 | 0.0956 | 12.39 | 129.93 | 2 | 0.0 | 0 | R.GSSSGGGYSSGSSSYGSGGR.Q | | 525-544 | CID |

#### Protein 8: Immunoglobulin heavy constant gamma 4 OS=Homo sapiens GN=IGHG4 PE=1 SV=1

**Accession:** IGHG4_HUMAN
**Database:** SwissProt
**MW [kDa] / pI:** 35.9 / 7.2
**Modification(s):** Carbamidomethyl

**Score:** 215.8
**Seq. Coverage [%]:** 10.1
**No. of Peptides:** 3

| 10 | 20 | 30 | 40 | 50 | 60 | 70 | 80 | 90 | 100 |
|---|---|---|---|---|---|---|---|---|---|
| ASTKGPSVFP | LAPCSRSTSE | STAALGCLVK | DYFPEPVTVS | WNSGALTSGV | HTFPAVLQSS | GLYSLSSVVT | VPSSSLGTKT | YTCNVDHKPS | NTKVDKRVES |
| **110** | **120** | **130** | **140** | **150** | **160** | **170** | **180** | **190** | **200** |
| KYGPPCPSCP | APEFLGGPSV | FLFPPKPKDT | LMISRTPEVT | CVVVDVSQED | PEVQFNWYVD | GVEVHNAKTK | PREEQFNSTY | RVVSVLTVLH | QDWLNGKEYK |
| **210** | **220** | **230** | **240** | **250** | **260** | **270** | **280** | **290** | **300** |
| CKVSNKGLPS | SIEKTISKAK | GQPREPQVYT | LPPSQEEMTK | NQVSLTCLVK | GFYPSDIAVE | WESNGQPENN | YKTTPPVLDS | DGSFFLYSRL | TVDKSRWQEG |
| **310** | **320** | **330** | | | | | | | |
| NVFSCSVMHE | ALHNHYTQKS | LSLSLGK | | | | | | | |

MS/MS Peptide Matches

| Cmpd. | m/z meas. | z | Δ m/z [ppm] | Δ m/z [Da] | Rt [min] | Score | No. of Cmpds. | Site [%] | P | Sequence | Modification | Range | Type |
|---|---|---|---|---|---|---|---|---|---|---|---|---|---|
| 54 | 832.993 | 2 | 125.48 | 0.1045 | 11.43 | 48.94 | 4 | 0.0 | 0 | K.TYTCNVDHKPSNTK.V | Carbamidomethyl: 4 | 80-93 | CID |
| 1167 | 905.096 | 2 | 651.31 | 0.5891 | 32.40 | 74.04 | 3 | 0.0 | 0 | R.VVSVLTVLHQDWLNGK.E | | 182-197 | CID |
| 1067 | 743.505 | 3 | 131.41 | 0.0977 | 30.72 | 92.84 | 3 | 0.0 | 1 | R.VVSVLTVLHQDWLNGKEYK.C | | 182-200 | CID |

#### Protein 9: Keratin, type I cytoskeletal 9 OS=Homo sapiens GN=KRT9 PE=1 SV=3

| | | | |
|---|---|---|---|
| **Accession:** | K1C9_HUMAN | **Score:** | 210.6 |
| **Database:** | SwissProt | **Seq. Coverage [%]:** | 8.3 |
| **MW [kDa] / pI:** | 62.0 / 5.1 | **No. of Peptides:** | 2 |

| 10 | 20 | 30 | 40 | 50 | 60 | 70 | 80 | 90 | 100 |
|---|---|---|---|---|---|---|---|---|---|
| MSCRQFSSSY | LSRSGGGGGG | GLGSGGSIRS | SYSRFSSSGG | GGGGGRFSSS | SGYGGGSSRV | CGRGGGGSFG | YSYGGGSGGG | FSASSLGGGF | GGGSRGFGGA |
| 110 | 120 | 130 | 140 | 150 | 160 | 170 | 180 | 190 | 200 |
| SGGGYSSSGG | FGGGFGGGSG | GGFGGGYGSG | FGGFGGFGGG | AGGGDGGILT | ANEKSTMQEL | NSRLASYLDK | VQALEEANND | LENKIQDWYD | KKGPAAIQKN |
| 210 | 220 | 230 | 240 | 250 | 260 | 270 | 280 | 290 | 300 |
| YSPYYNTIDD | LKDQIVDLTV | GNNKTLLDID | NTRMTLDDFR | IKFEMEQNLR | QGVDADINGL | RQVLDNLTME | KSDLEMQYET | LQEELMALKK | NHKEEMSQLT |
| 310 | 320 | 330 | 340 | 350 | 360 | 370 | 380 | 390 | 400 |
| GQNSGDVNVE | INVAPGKDLT | KTLNDMRQEY | EQLIAKNRKD | IENQYETQIT | QIEHEVSSSG | QEVQSSAKEV | TQLRHGVQEL | EIELQSQLSK | KAALEKSLED |
| 410 | 420 | 430 | 440 | 450 | 460 | 470 | 480 | 490 | 500 |
| TKNRYCGQLQ | MIQEQISNLE | AQITDVRQEI | ECQNQEYSLL | LSIKMRLEKE | IETYHNLLEG | GQEDFESSGA | GKIGLGGRGG | SGGSYGRGSR | GGSGGSYGGG |
| 510 | 520 | 530 | 540 | 550 | 560 | 570 | 580 | 590 | 600 |
| GSGGGYGGGS | GSRGGSGGSY | GGGSGSGGGS | GGGYGGGSGG | GHSGGSGGGH | SGGSGGNYGG | GSGSGGGSGG | GYGGGSGSRG | GSGGSHGGGS | GFGGESGGSY |
| 610 | 620 | 630 | | | | | | | |
| GGGEEASGSG | GGYGGGSGKS | SHS | | | | | | | |

MS/MS Peptide Matches

| Cmpd. | m/z meas. | z | Δ m/z [ppm] | Δ m/z [Da] | Rt [min] | Score | No. of Cmpds. | Site [%] | P | Sequence | Modification | Range | Type |
|---|---|---|---|---|---|---|---|---|---|---|---|---|---|
| 1021 | 1089.216 | 3 | 342.74 | 0.3732 | 29.93 | 63.56 | 3 | 0.0 | 0 | K.DIENQYETQITQIEHEVSSSGQEVQSSAK.E | | 340-368 | CID |
| 71 | 896.51 | 2 | 158.98 | 0.1425 | 12.12 | 147.06 | 6 | 0.0 | 0 | R.GGSGGSYGGGGSGGGYGGGSGSR.G | | 491-513 | CID |

#### Protein 10: Keratin, type I cytoskeletal 10 OS=Homo sapiens GN=KRT10 PE=1 SV=6

| | | | |
|---|---|---|---|
| **Accession:** | K1C10_HUMAN | **Score:** | 158.7 |
| **Database:** | SwissProt | **Seq. Coverage [%]:** | 7.9 |
| **MW [kDa] / pI:** | 58.8 / 5.1 | **No. of Peptides:** | 2 |

| 10 | 20 | 30 | 40 | 50 | 60 | 70 | 80 | 90 | 100 |
|---|---|---|---|---|---|---|---|---|---|
| MSVRYSSSKH | YSSSRSGGGG | GGGGCGGGGG | VSSLRISSSK | GSLGGGFSSG | GFSGGSFSRG | SSGGGCFGGS | SGGYGGLGGF | GGGSFRGSYG | SSSFGGSYGG |
| **110** | **120** | **130** | **140** | **150** | **160** | **170** | **180** | **190** | **200** |
| IFGGGSFGGG | SFGGGSFGGG | GFGGGGFGGG | FGGGFGGDGG | LLSGNEKVTM | QNLNDRLASY | LDKVRALEES | NYELEGKIKE | WYEKHGNSHQ | GEPRDYSKYY |
| **210** | **220** | **230** | **240** | **250** | **260** | **270** | **280** | **290** | **300** |
| KTIDDLKNQI | LNLTTDNANI | LLQIDNARLA | ADDFRLKYEN | EVALRQSVEA | DINGLRRVLD | ELTLTKADLE | MQIESLTEEL | AYLKKNHEEE | MKDLRNVSTG |
| **310** | **320** | **330** | **340** | **350** | **360** | **370** | **380** | **390** | **400** |
| DVNVEMNAAP | GVDLTQLLNN | MRSQYEQLAE | QNRKDAEAWF | NEKSKELTTE | IDNNIEQISS | YKSEITELRR | NVQALEIELQ | SQLALKQSLE | ASLAETEGRY |
| **410** | **420** | **430** | **440** | **450** | **460** | **470** | **480** | **490** | **500** |
| CVQLSQIQAQ | ISALEEQLQQ | IRAETECQNT | EYQQLLDIKI | RLENEIQTYR | SLLEGEGSSG | GGGRGGGSFG | GGYGGGSSGG | GSSGGGHGGG | HGGSSGGGYG |
| **510** | **520** | **530** | **540** | **550** | **560** | **570** | **580** | **590** | |
| GGSSGGGSSG | GGYGGGSSSG | GHGGSSSGGY | GGGSSGGGGG | GYGGGSSGGG | SSSGGGYGGG | SSSGGHKSSS | SGSVGESSSK | GPRY | |

**MS/MS Peptide Matches**

| Cmpd. | m/z meas. | z | Δ m/z [ppm] | Δ m/z [Da] | Rt [min] | Score | No. of Cmpds. | Site [%] | P | Sequence | Modification | Range | Type |
|---|---|---|---|---|---|---|---|---|---|---|---|---|---|
| 674 | 854.462 | 2 | 84.61 | 0.0723 | 23.87 | 132.04 | 3 | 0.0 | 0 | K.GSLGGGFSSGGFSGGSFSR.G | | 41-59 | CID |
| 1310 | 958.482 | 3 | 361.35 | 0.3462 | 34.78 | 26.7 | 1 | 0.0 | 0 | R.NVSTGDVNVEMNAAPGVDLTQLLNNMR.S | | 296-322 | CID |

#### Protein 11: Immunoglobulin kappa variable 3-20 OS=Homo sapiens GN=IGKV3-20 PE=1 SV=2

**Accession:** KV320_HUMAN
**Database:** SwissProt
**MW [kDa] / pI:** 12.5 / 4.9

**Score:** 105.6
**Seq. Coverage [%]:** 19.0
**No. of Peptides:** 1

| 10 | 20 | 30 | 40 | 50 | 60 | 70 | 80 | 90 | 100 |
|---|---|---|---|---|---|---|---|---|---|
| METPAQLLFL | LLLWLPDTTG | EIVLTQSPGT | LSLSPGERAT | LSCRASQSVS | SSYLAWYQQK | PGQAPRLLIY | GASSRATGIP | DRFSGSGSGT | DFTLTISRLE |
| **110** | **120** | | | | | | | | |
| PEDFAVYYCQ | QYGSSP | | | | | | | | |

##### MS/MS Peptide Matches

| Cmpd. | m/z meas. | z | Δ m/z [ppm] | Δ m/z [Da] | Rt [min] | Score | No. of Cmpds. | Site [%] | P | Sequence | Modification | Range | Type |
|---|---|---|---|---|---|---|---|---|---|---|---|---|---|
| 685 | 813.825 | 3 | 104.58 | 0.0851 | 24.06 | 105.64 | 4 | 0.0 | 0 | R.ASQSVSSSYLAWYQQKPGQAPR.L | | 45-66 | CID |

#### Protein 12: Immunoglobulin heavy variable 4-4 OS=Homo sapiens GN=IGHV4-4 PE=3 SV=2

**Accession:** HV404_HUMAN
**Database:** SwissProt
**MW [kDa] / pI:** 12.8 / 9.6
**Modification(s):** Carbamidomethyl

**Score:** 90.8
**Seq. Coverage [%]:** 13.7
**No. of Peptides:** 1

| 10 | 20 | 30 | 40 | 50 | 60 | 70 | 80 | 90 | 100 |
|---|---|---|---|---|---|---|---|---|---|
| MKHLWFFLLL | VAAPRWVLSQ | VQLQESGPGL | VKPSGTLSLT | CAVSGGSISS | SNWWSWVRQP | PGKGLEWIGE | IYHSGSTNYN | PSLKSRVTIS | VDKSKNQFSL |
| **110** | **120** | | | | | | | | |
| KLSSVTAADT | AVYYCAR | | | | | | | | |

##### MS/MS Peptide Matches

| Cmpd. | m/z meas. | z | Δ m/z [ppm] | Δ m/z [Da] | Rt [min] | Score | No. of Cmpds. | Site [%] | P | Sequence | Modification | Range | Type |
|---|---|---|---|---|---|---|---|---|---|---|---|---|---|
| 588 | 874.519 | 2 | 113.65 | 0.0994 | 22.33 | 90.78 | 3 | 0.0 | 0 | K.LSSVTAADTAVYYCAR.- | Carbamidomethyl: 14 | 102-117 | CID |

#### Protein 13: Immunoglobulin heavy variable 5-51 OS=Homo sapiens GN=IGHV5-51 PE=3 SV=1

| | | | |
|---|---|---|---|
| **Accession:** | HV551_HUMAN | **Score:** | 58.1 |
| **Database:** | SwissProt | **Seq. Coverage [%]:** | 12.8 |
| **MW [kDa] / pI:** | 12.7 / 8.4 | **No. of Peptides:** | 1 |

```
        10         20         30         40         50         60         70         80         90        100
MGSTAILALL LAVLQGVCAE VQLVQSGAEV KKPGESLKIS CKGSGYSFTS YWIGWVRQMP GKGLEWMGII YPGDSDTRYS PSFQGQVTIS ADKSISTAYL
       110        120
QWSSLKASDT AMYYCAR
```

MS/MS Peptide Matches

| Cmpd. | m/z meas. | z | Δ m/z [ppm] | Δ m/z [Da] | Rt [min] | Score | No. of Cmpds. | Site [%] | P | Sequence | Modification | Range | Type |
|---|---|---|---|---|---|---|---|---|---|---|---|---|---|
| 558 | 814.468 | 2 | 81.35 | 0.0662 | 21.73 | 58.09 | 2 | 0.0 | 0 | R.YSPSFQGQVTISADK.S | | 79-93 | CID |

#### Protein 14: Immunoglobulin kappa variable 3-15 OS=Homo sapiens GN=IGKV3-15 PE=1 SV=2

| | | | |
|---|---|---|---|
| **Accession:** | KV315_HUMAN | **Score:** | 47.3 |
| **Database:** | SwissProt | **Seq. Coverage [%]:** | 18.3 |
| **MW [kDa] / pI:** | 12.5 / 5.1 | **No. of Peptides:** | 1 |

```
        10         20         30         40         50         60         70         80         90        100
MEAPAQLLFL LLLWLPDTTG EIVMTQSPAT LSVSPGERAT LSCRASQSVS SNLAWYQQKP GQAPRLLIYG ASTRATGIPA RFSGSGSGTE FTLTISSLQS
       110        120
EDFAVYYCQQ YNNWP
```

MS/MS Peptide Matches

| Cmpd. | m/z meas. | z | Δ m/z [ppm] | Δ m/z [Da] | Rt [min] | Score | No. of Cmpds. | Site [%] | P | Sequence | Modification | Range | Type |
|---|---|---|---|---|---|---|---|---|---|---|---|---|---|
| 544 | 768.462 | 3 | 94.89 | 0.0729 | 21.46 | 47.28 | 2 | 0.0 | 0 | R.ASQSVSSNLAWYQQKPGQAPR.L | | 45-65 | CID |

BRUKER

#### Project Info

| | | | |
|---|---|---|---|
| **Name:** | Alagesan | **Date:** | Mar 5, 2019 |
| **Note:** | Nanobooster | | |

#### Sample Info & Protocols

| | |
|---|---|
| **Name:** | Nanobooster - IgG |
| **Date:** | Mar 5, 2019 |

#### Search Result Info

| | | | |
|---|---|---|---|
| **Search Result:** | IonTrap_Human_Swiss_CID_2019-03-05 17:55:08 | | |
| **Location:** | /Alagesan/Nanobooster - IgG/KA_IgG3 HD NB ACN CID_GA4_01_9085.mgf | | |
| **Search Method:** | IonTrap_Human_Swiss_CID | **Version:** | 1.2 |
| **Search Engine(s):** | Mascot, 2.6.0 | **Ident. Compound(s):** | 57/2373 |
| **Database(s):** | SwissProt, SwissProt_2017_04.fasta | | |
| **Note:** | | | |

#### Results

##### Protein 1: Immunoglobulin heavy constant gamma 3 OS=Homo sapiens GN=IGHG3 PE=1 SV=2

| | | | |
|---|---|---|---|
| **Accession:** | IGHG3_HUMAN | **Score:** | 1406.0 |
| **Database:** | SwissProt | **Seq. Coverage [%]:** | 58.1 |
| **MW [kDa] / pI:** | 41.3 / 8.2 | **No. of Peptides:** | 19 |
| **Modification(s):** | Carbamidomethyl, Oxidation | | |

```
        10         20         30         40         50         60         70         80         90        100
ASTKGPSVFP LAPCSRSTSG GTAALGCLVK DYFPEPVTVS WNSGALTSGV HTFPAVLQSS GLYSLSSVVT VPSSSLGTQT YTCNVNHKPS NTKVDKRVEL
       110        120        130        140        150        160        170        180        190        200
KTPLGDTTHT CPRCPEPKSC DTPPPCPRCP EPKSCDTPPP CPRCPEPKSC DTPPPCPRCP APELLGGPSV FLFPPKPKDT LMISRTPEVT CVVVDVSHED
       210        220        230        240        250        260        270        280        290        300
PEVQFKWYVD GVEVHNAKTK PREEQYNSTF RVVSVLTVLH QDWLNGKEYK CKVSNKALPA PIEKTISKTK GQPREPQVYT LPPSREEMTK NQVSLTCLVK
       310        320        330        340        350        360        370        380
GFYPSDIAVE WESSGQPENN YNTTPPMLDS DGSFFLYSKL TVDKSRWQQG NIFSCSVMHE ALHNRFTQKS LSLSPGK
```

**MS/MS Peptide Matches**

| Cmpd. | m/z meas. | z | Δ m/z [ppm] | Δ m/z [Da] | Rt [min] | Score | No. of Cmpds. | Site [%] | P | Sequence | Modification | Range | Type |
|---|---|---|---|---|---|---|---|---|---|---|---|---|---|
| 941 | 864.548 | 3 | 508.36 | 0.4393 | 28.47 | 63.76 | 3 | 0.0 | 1 | K.GPSVFPLAPCSRSTSGGTAALGCLVK.D | Carbamidomethyl: 10, 23 | 5-30 | CID |
| 426 | 913.133 | 2 | 181.53 | 0.1657 | 19.11 | 95.69 | 6 | 0.0 | 1 | R.VELKTPLGDTTHTCPR.C | Carbamidomethyl: 14 | 98-113 | CID |
| 224 | 984.011 | 2 | 52.73 | 0.0519 | 15.18 | 45.5 | 3 | 0.0 | 1 | K.TPLGDTTHTCPRCPEPK.S | Carbamidomethyl: 10, 13 | 102-118 | CID |
| 168 | 899.51 | 2 | 134.00 | 0.1205 | 14.15 | 73.57 | 16 | 0.0 | 1 | R.CPEPKSCDTPPPCPR.C | Carbamidomethyl: 1, 7, 13 | 114-128 | CID |
| 207 | 899.452 | 2 | 69.51 | 0.0625 | 14.88 | 29.07 | 2 | 0.0 | 1 | K.SCDTPPPCPRCPEPK.S | Carbamidomethyl: 2, 8, 11 | 119-133 | CID |
| 1149 | 718.052 | 3 | 452.52 | 0.3248 | 32.02 | 121.32 | 23 | 0.0 | 0 | R.CPAPELLGGPSVFLFPPKPK.D | Carbamidomethyl: 1 | 159-178 | CID |

| Cmpd. | m/z meas. | z | Δ m/z [ppm] | Δ m/z [Da] | Rt [min] | Score | No. of Cmpds. | Site [%] | P | Sequence | Modification | Range | Type |
|---|---|---|---|---|---|---|---|---|---|---|---|---|---|
| 1164 | 742.737 | 4 | 115.36 | 0.0857 | 32.27 | 92.14 | 6 | 0.0 | 1 | R.CPAPELLGGPSVFLFPPKPKDTLMISR.T | Carbamidomethyl: 1 | 159-185 | CID |
| 1124 | 747.017 | 4 | 491.46 | 0.3669 | 31.60 | 51.68 | 3 | 0.0 | 1 | R.CPAPELLGGPSVFLFPPKPKDTLMISR.T | Carbamidomethyl: 1; Oxidation: 24 | 159-185 | CID |
| 1021 | 808.496 | 4 | 121.04 | 0.0978 | 29.84 | 82.26 | 3 | 0.0 | 1 | K.DTLMISRTPEVTCVVVDVSHEDPEVQFK.W | Carbamidomethyl: 13 | 179-206 | CID |
| 896 | 805.528 | 3 | 171.77 | 0.1383 | 27.47 | 129.07 | 5 | 0.0 | 0 | R.TPEVTCVVVDVSHEDPEVQFK.W | Carbamidomethyl: 6 | 186-206 | CID |
| 1062 | 1271.629 | 3 | 274.06 | 0.3484 | 30.50 | 58.83 | 3 | 0.0 | 1 | R.TPEVTCVVVDVSHEDPEVQFKWYVDGVEVHNAK.T | Carbamidomethyl: 6 | 186-218 | CID |
| 529 | 708.94 | 2 | 128.44 | 0.0910 | 21.00 | 88.89 | 17 | 0.0 | 0 | K.WYVDGVEVHNAK.T | | 207-218 | CID |
| 1165 | 905.099 | 2 | 654.63 | 0.5921 | 32.28 | 103.91 | 3 | 0.0 | 0 | R.VVSVLTVLHQDWLNGK.E | | 232-247 | CID |
| 1082 | 743.487 | 3 | 107.19 | 0.0797 | 30.84 | 88.75 | 5 | 0.0 | 1 | R.VVSVLTVLHQDWLNGKEYK.C | | 232-250 | CID |
| 492 | 953.036 | 2 | 64.26 | 0.0612 | 20.31 | 55.26 | 9 | 0.0 | 1 | R.EPQVYTLPPSREEMTK.N | | 275-290 | CID |
| 459 | 961.537 | 2 | 587.72 | 0.5648 | 19.68 | 30.86 | 2 | 0.0 | 1 | R.EPQVYTLPPSREEMTK.N | Oxidation: 14 | 275-290 | CID |
| 1309 | 1464.236 | 3 | 170.53 | 0.2497 | 34.72 | 57.69 | 3 | 0.0 | 0 | K.GFYPSDIAVEWESSGQPENNYNTTPPMLDSDGSFFLYSK.L | | 301-339 | CID |
| 793 | 1157.597 | 2 | 54.56 | 0.0632 | 25.76 | 70.58 | 5 | 0.0 | 0 | R.WQQGNIFSCSVMHEALHNR.F | Carbamidomethyl: 9 | 347-365 | CID |
| 830 | 940.503 | 3 | 412.86 | 0.3881 | 26.40 | 67.15 | 3 | 0.0 | 1 | R.WQQGNIFSCSVMHEALHNRFTQK.S | Carbamidomethyl: 9 | 347-369 | CID |

#### Protein 2: Immunoglobulin kappa constant OS=Homo sapiens GN=IGKC PE=1 SV=2

**Accession:** IGKC_HUMAN
**Database:** SwissProt
**MW [kDa] / pI:** 11.8 / 6.1
**Modification(s):** Carbamidomethyl

**Score:** 994.3
**Seq. Coverage [%]:** 86.9
**No. of Peptides:** 10

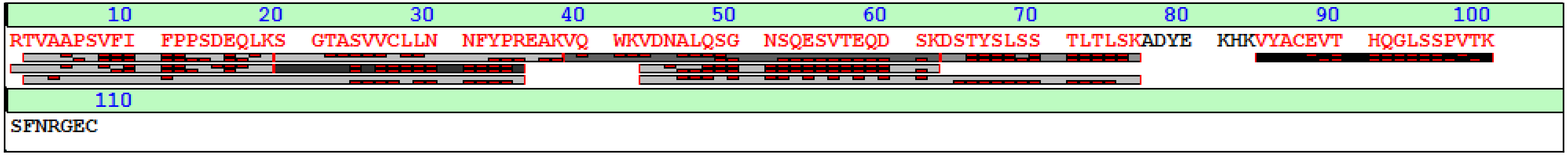


##### MS/MS Peptide Matches

| Cmpd. | m/z meas. | z | Δ m/z [ppm] | Δ m/z [Da] | Rt [min] | Score | No. of Cmpds. | Site [%] | P | Sequence | Modification | Range | Type |
|---|---|---|---|---|---|---|---|---|---|---|---|---|---|
| 967 | 1051.626 | 2 | 55.47 | 0.0583 | 28.92 | 41.87 | 7 | 0.0 | 1 | -.RTVAAPSVFIFPPSDEQLK.S | | 1-19 | CID |
| 1084 | 973.632 | 2 | 118.01 | 0.1149 | 30.90 | 81.33 | 14 | 0.0 | 0 | R.TVAAPSVFIFPPSDEQLK.S | | 2-19 | CID |
| 1518 | 1242.665 | 3 | 288.74 | 0.3587 | 38.24 | 85.74 | 16 | 0.0 | 1 | R.TVAAPSVFIFPPSDEQLKSGTASVVCLLNNFYPR.E | Carbamidomethyl: 26 | 2-35 | CID |
| 1212 | 899.568 | 2 | 129.80 | 0.1167 | 33.07 | 63.02 | 14 | 0.0 | 0 | K.SGTASVVCLLNNFYPR.E | Carbamidomethyl: 8 | 20-35 | CID |
| 1100 | 709.434 | 3 | 102.23 | 0.0725 | 31.21 | 122.74 | 3 | 0.0 | 1 | K.SGTASVVCLLNNFYPREAK.V | Carbamidomethyl: 8 | 20-38 | CID |
| 533 | 893.234 | 3 | 155.81 | 0.1392 | 21.10 | 130.25 | 8 | 0.0 | 1 | K.VQWKVDNALQSGNSQESVTEQDSK.D | | 39-62 | CID |
| 256 | 1068.941 | 2 | 423.96 | 0.4530 | 15.71 | 161.34 | 9 | 0.0 | 0 | K.VDNALQSGNSQESVTEQDSK.D | | 43-62 | CID |
| 724 | 1207.565 | 3 | 268.14 | 0.3237 | 24.59 | 102.87 | 3 | 0.0 | 1 | K.VDNALQSGNSQESVTEQDSKDSTYSLSSTLTLSK.A | | 43-76 | CID |
| 742 | 751.973 | 2 | 119.89 | 0.0901 | 24.94 | 88.83 | 4 | 0.0 | 0 | K.DSTYSLSSTLTLSK.A | | 63-76 | CID |
| 428 | 938.559 | 2 | 97.93 | 0.0919 | 19.14 | 116.33 | 8 | 0.0 | 0 | K.VYACEVTHQGLSSPVTK.S | Carbamidomethyl: 4 | 84-100 | CID |

#### Protein 3: Immunoglobulin kappa light chain OS=Homo sapiens PE=1 SV=1

**Accession:** IGK_HUMAN
**Database:** SwissProt
**MW [kDa] / pI:** 23.4 / 6.9
**Modification(s):** Carbamidomethyl

**Score:** 890.2
**Seq. Coverage [%]:** 43.0
**No. of Peptides:** 8

| 10 | 20 | 30 | 40 | 50 | 60 | 70 | 80 | 90 | 100 |
|---|---|---|---|---|---|---|---|---|---|
| DIQMTQSPST | LSASVGDRVT | ITCRASQSIN | TWLAWYQQKP | GKAPKLLMYK | ASSLESGVPS | RFIGSGSGTE | FTLTISSLQP | DDFATYYCQQ | YNSDSKMFGQ |
| **110** | **120** | **130** | **140** | **150** | **160** | **170** | **180** | **190** | **200** |
| GTKVEVKGTV | AAPSVFIFPP | SDEQLKSGTA | SVVCLLNNFY | PREAKVQWKV | DNALQSGNSQ | ESVTEQDSKD | STYSLSSTLT | LSKADYEKHK | VYACEVTHQG |
| **210** | **220** | | | | | | | | |
| LSSPVTKSFN | RGEC | | | | | | | | |

**MS/MS Peptide Matches**

| Cmpd. | m/z meas. | z | Δ m/z [ppm] | Δ m/z [Da] | Rt [min] | Score | No. of Cmpds. | Site [%] | P | Sequence | Modification | Range | Type |
|---|---|---|---|---|---|---|---|---|---|---|---|---|---|
| 594 | 947.012 | 2 | 60.66 | 0.0574 | 22.29 | 104.86 | 1 | 0.0 | 0 | -.DIQMTQSPSTLSASVGDR.V | | 1-18 | CID |
| 1212 | 899.568 | 2 | 129.80 | 0.1167 | 33.07 | 63.02 | 14 | 0.0 | 0 | K.SGTASVVCLLNNFYPR.E | Carbamidomethyl: 8 | 127-142 | CID |
| 1100 | 709.434 | 3 | 102.23 | 0.0725 | 31.21 | 122.74 | 3 | 0.0 | 1 | K.SGTASVVCLLNNFYPREAK.V | Carbamidomethyl: 8 | 127-145 | CID |
| 533 | 893.234 | 3 | 155.81 | 0.1392 | 21.10 | 130.25 | 8 | 0.0 | 1 | K.VQWKVDNALQSGNSQESVTEQDSK.D | | 146-169 | CID |
| 256 | 1068.941 | 2 | 423.96 | 0.4530 | 15.71 | 161.34 | 9 | 0.0 | 0 | K.VDNALQSGNSQESVTEQDSK.D | | 150-169 | CID |
| 724 | 1207.565 | 3 | 268.14 | 0.3237 | 24.59 | 102.87 | 3 | 0.0 | 1 | K.VDNALQSGNSQESVTEQDSKDSTYSLSSTLTLSK.A | | 150-183 | CID |
| 742 | 751.973 | 2 | 119.89 | 0.0901 | 24.94 | 88.83 | 4 | 0.0 | 0 | K.DSTYSLSSTLTLSK.A | | 170-183 | CID |
| 428 | 938.559 | 2 | 97.93 | 0.0919 | 19.14 | 116.33 | 8 | 0.0 | 0 | K.VYACEVTHQGLSSPVTK.S | Carbamidomethyl: 4 | 191-207 | CID |

##### Protein 4: Keratin, type II cytoskeletal 1 OS=Homo sapiens GN=KRT1 PE=1 SV=6

**Accession:** K2C1_HUMAN
**Database:** SwissProt
**MW [kDa] / pI:** 66.0 / 8.2

**Score:** 688.0
**Seq. Coverage [%]:** 19.7
**No. of Peptides:** 7

| 10 | 20 | 30 | 40 | 50 | 60 | 70 | 80 | 90 | 100 |
|---|---|---|---|---|---|---|---|---|---|
| MSRQFSSRSG | YRSGGGFSSG | SAGIINYQRR | TTSSSTRRSG | GGGGRFSSCG | GGGGSFGAGG | GFGSRSLVNL | GGSKSISISV | ARGGGRGSGF | GGGYGGGGFG |
| **110** | **120** | **130** | **140** | **150** | **160** | **170** | **180** | **190** | **200** |
| GGGFGGGGFG | GGGIGGGGFG | GFGSGGGGFG | GGGFGGGGYG | GGYGPVCPPG | GIQEVTINQS | LLQPLNVEID | PEIQKVKSRE | REQIKSLNNQ | FASFIDKVRF |
| **210** | **220** | **230** | **240** | **250** | **260** | **270** | **280** | **290** | **300** |
| LEQQNQVLQT | KWELLQQVDT | STRTHNLEPY | FESFINNLRR | RVDQLKSDQS | RLDSELKNMQ | DMVEDYRNKY | EDEINKRTNA | ENEFVTIKKD | VDGAYMTKVD |
| **310** | **320** | **330** | **340** | **350** | **360** | **370** | **380** | **390** | **400** |
| LQAKLDNLQQ | EIDFLTALYQ | AELSQMQTQI | SETNVILSMD | NNRSLDLDSI | IAEVKAQYED | IAQKSKAEAE | SLYQSKYEEL | QITAGRHGDS | VRNSKIEISE |
| **410** | **420** | **430** | **440** | **450** | **460** | **470** | **480** | **490** | **500** |
| LNRVIQRLRS | EIDNVKKQIS | NLQQSISDAE | QRGENALKDA | KNKLNDLEDA | LQQAKEDLAR | LLRDYQELMN | TKLALDLEIA | TYRTLLEGEE | SRMSGECAPN |
| **510** | **520** | **530** | **540** | **550** | **560** | **570** | **580** | **590** | **600** |
| VSVSVSTSHT | TISGGGSRGG | GGGGYGSGGS | SYGSGGGSYG | SGGGGGGGRG | SYGSGGSSYG | SGGGSYGSGG | GGGGHGSYGS | GSSSGGYRGG | SGGGGGGSSG |
| **610** | **620** | **630** | **640** | **650** | | | | | |
| GRGSGGGSSG | GSIGGRGSSS | GGVKSSGGSS | SVKFVSTTYS | GVTR | | | | | |

##### MS/MS Peptide Matches

| Cmpd. | m/z meas. | z | Δ m/z [ppm] | Δ m/z [Da] | Rt [min] | Score | No. of Cmpds. | Site [%] | P | Sequence | Modification | Range | Type |
|---|---|---|---|---|---|---|---|---|---|---|---|---|---|
| 956 | 820.029 | 2 | 116.41 | 0.0954 | 28.70 | 59.21 | 1 | 0.0 | 1 | K.SLNNQFASFIDKVR.F | | 186-199 | CID |
| 474 | 738.482 | 2 | 116.10 | 0.0857 | 19.95 | 78.73 | 3 | 0.0 | 0 | R.FLEQQNQVLQTK.W | | 200-211 | CID |
| 999 | 978.559 | 3 | 390.57 | 0.3820 | 29.47 | 89.79 | 3 | 0.0 | 1 | R.FLEQQNQVLQTKWELLQQVDTSTR.T | | 200-223 | CID |
| 1491 | 783.783 | 3 | 49.36 | 0.0387 | 37.78 | 144.96 | 3 | 0.0 | 1 | R.SLDLDSIIAEVKAQYEDIAQK.S | | 344-364 | CID |
| 564 | 859.005 | 2 | 88.25 | 0.0758 | 21.68 | 84.49 | 3 | 0.0 | 0 | K.QISNLQQSISDAEQR.G | | 418-432 | CID |
| 181 | 1192.448 | 2 | -26.47 | -0.0316 | 14.37 | 212.62 | 4 | 0.0 | 0 | R.GGGGGGYGSGGSSYGSGGGSYGSGGGGGG GR.G | | 519-549 | CID |
| 56 | 876.557 | 2 | 159.76 | 0.1400 | 11.20 | 18.2 | 1 | 0.0 | 1 | R.GSGGGSSGGSIGGRGSSSGGVK.S | | 603-624 | CID |

BRUKER

#### Protein 5: Immunoglobulin lambda constant 3 OS=Homo sapiens GN=IGLC3 PE=1 SV=1

| | | | |
|---|---|---|---|
| **Accession:** | IGLC3_HUMAN | **Score:** | 657.3 |
| **Database:** | SwissProt | **Seq. Coverage [%]:** | 77.4 |
| **MW [kDa] / pI:** | 11.3 / 6.9 | **No. of Peptides:** | 8 |
| **Modification(s):** | Carbamidomethyl | | |

| 10 | 20 | 30 | 40 | 50 | 60 | 70 | 80 | 90 | 100 |
|---|---|---|---|---|---|---|---|---|---|
| GQPKAAPSVT | LFPPSSEELQ | ANKATLVCLI | SDFYPGAVTV | AWKADSSPVK | AGVETTTPSK | QSNNKYAASS | YLSLTPEQWK | SHKSYSCQVT | HEGSTVEKTV |

| 110 |
|---|
| APTECS |

MS/MS Peptide Matches

| Cmpd. | m/z meas. | z | Δ m/z [ppm] | Δ m/z [Da] | Rt [min] | Score | No. of Cmpds. | Site [%] | P | Sequence | Modification | Range | Type |
|---|---|---|---|---|---|---|---|---|---|---|---|---|---|
| 831 | 993.559 | 2 | 46.74 | 0.0464 | 26.41 | 77.36 | 14 | 0.0 | 0 | K.AAPSVTLFPPSSEELQANK.A | | 5-23 | CID |
| 219 | 838.024 | 2 | 111.14 | 0.0931 | 15.09 | 54.29 | 3 | 0.0 | 1 | K.ADSSPVKAGVETTTPSK.Q | | 44-60 | CID |
| 55 | 782.029 | 2 | 812.06 | 0.6345 | 11.17 | 92.58 | 20 | 0.0 | 1 | K.AGVETTTPSKQSNNK.Y | | 51-65 | CID |
| 820 | 772.786 | 3 | 523.57 | 0.4044 | 26.20 | 67.85 | 5 | 0.0 | 1 | K.QSNNKYAASSYLSLTPEQWK.S | | 61-80 | CID |
| 912 | 872.619 | 2 | 213.14 | 0.1859 | 27.77 | 92.51 | 14 | 0.0 | 0 | K.YAASSYLSLTPEQWK.S | | 66-80 | CID |
| 84 | 1032.574 | 2 | 94.74 | 0.0978 | 12.27 | 88.92 | 5 | 0.0 | 1 | K.SHKSYSCQVTHEGSTVEK.T | Carbamidomethyl: 7 | 81-98 | CID |
| 164 | 856.495 | 2 | 130.51 | 0.1118 | 14.07 | 107.98 | 9 | 0.0 | 0 | K.SYSCQVTHEGSTVEK.T | Carbamidomethyl: 4 | 84-98 | CID |
| 407 | 1279.574 | 2 | 399.75 | 0.5113 | 18.76 | 75.76 | 6 | 0.0 | 1 | K.SYSCQVTHEGSTVEKTVAPTECS.- | Carbamidomethyl: 4, 22 | 84-106 | CID |

#### Protein 6: Immunoglobulin lambda constant 2 OS=Homo sapiens GN=IGLC2 PE=1 SV=1

| | | | |
|---|---|---|---|
| **Accession:** | IGLC2_HUMAN | **Score:** | 618.0 |
| **Database:** | SwissProt | **Seq. Coverage [%]:** | 77.4 |
| **MW [kDa] / pI:** | 11.3 / 6.9 | **No. of Peptides:** | 8 |

**Modification(s):** Carbamidomethyl

```
         10         20         30         40         50         60         70         80         90        100
GQPKAAPSVT LFPPSSEELQ ANKATLVCLI SDFYPGAVTV AWKADSSPVK AGVETTTPSK QSNNKYAASS YLSLTPEQWK SHRSYSCQVT HEGSTVEKTV
        110
APTECS
```

**MS/MS Peptide Matches**

| Cmpd. | m/z meas. | z | Δ m/z [ppm] | Δ m/z [Da] | Rt [min] | Score | No. of Cmpds. | Site [%] | P | Sequence | Modification | Range | Type |
|---|---|---|---|---|---|---|---|---|---|---|---|---|---|
| 831 | 993.559 | 2 | 46.74 | 0.0464 | 26.41 | 77.36 | 14 | 0.0 | 0 | K.AAPSVTLFPPSSEELQANK.A | | 5-23 | CID |
| 219 | 838.024 | 2 | 111.14 | 0.0931 | 15.09 | 54.29 | 3 | 0.0 | 1 | K.ADSSPVKAGVETTTPSK.Q | | 44-60 | CID |
| 55 | 782.029 | 2 | 812.06 | 0.6345 | 11.17 | 92.58 | 20 | 0.0 | 1 | K.AGVETTTPSKQSNNK.Y | | 51-65 | CID |
| 820 | 772.786 | 3 | 523.57 | 0.4044 | 26.20 | 67.85 | 5 | 0.0 | 1 | K.QSNNKYAASSYLSLTPEQWK.S | | 61-80 | CID |
| 912 | 872.619 | 2 | 213.14 | 0.1859 | 27.77 | 92.51 | 14 | 0.0 | 0 | K.YAASSYLSLTPEQWK.S | | 66-80 | CID |
| 95 | 1046.553 | 2 | 70.47 | 0.0737 | 12.53 | 49.66 | 5 | 0.0 | 1 | K.SHRSYSCQVTHEGSTVEK.T | Carbamidomethyl: 7 | 81-98 | CID |
| 164 | 856.495 | 2 | 130.51 | 0.1118 | 14.07 | 107.98 | 9 | 0.0 | 0 | R.SYSCQVTHEGSTVEK.T | Carbamidomethyl: 4 | 84-98 | CID |
| 407 | 1279.574 | 2 | 399.75 | 0.5113 | 18.76 | 75.76 | 6 | 0.0 | 1 | R.SYSCQVTHEGSTVEKTVAPTECS.- | Carbamidomethyl: 4, 22 | 84-106 | CID |

#### Protein 7: Immunoglobulin heavy constant gamma 4 OS=Homo sapiens GN=IGHG4 PE=1 SV=1

**Accession:** IGHG4_HUMAN
**Database:** SwissProt
**MW [kDa] / pI:** 35.9 / 7.2
**Modification(s):** Carbamidomethyl

**Score:** 241.4
**Seq. Coverage [%]:** 10.1
**No. of Peptides:** 3

| 10 | 20 | 30 | 40 | 50 | 60 | 70 | 80 | 90 | 100 |
|---|---|---|---|---|---|---|---|---|---|
| ASTKGPSVFP | LAPCSRSTSE | STAALGCLVK | DYFPEPVTVS | WNSGALTSGV | HTFPAVLQSS | GLYSLSSVVT | VPSSSLGTKT | YTCNVDHKPS | NTKVDKRVES |
| **110** | **120** | **130** | **140** | **150** | **160** | **170** | **180** | **190** | **200** |
| KYGPPCPSCP | APEFLGGPSV | FLFPPKPKDT | LMISRTPEVT | CVVVDVSQED | PEVQFNWYVD | GVEVHNAKTK | PREEQFNSTY | RVVSVLTVLH | QDWLNGKEYK |
| **210** | **220** | **230** | **240** | **250** | **260** | **270** | **280** | **290** | **300** |
| CKVSNKGLPS | SIEKTISKAK | GQPREPQVYT | LPPSQEEMTK | NQVSLTCLVK | GFYPSDIAVE | WESNGQPENN | YKTTPPVLDS | DGSFFLYSRL | TVDKSRWQEG |
| **310** | **320** | **330** | | | | | | | |
| NVFSCSVMHE | ALHNHYTQKS | LSLSLGK | | | | | | | |

MS/MS Peptide Matches

| Cmpd. | m/z meas. | z | Δ m/z [ppm] | Δ m/z [Da] | Rt [min] | Score | No. of Cmpds. | Site [%] | P | Sequence | Modification | Range | Type |
|---|---|---|---|---|---|---|---|---|---|---|---|---|---|
| 62 | 833.029 | 2 | 168.71 | 0.1405 | 11.37 | 48.7 | 3 | 0.0 | 0 | K.TYTCNVDHKPSNTK.V | Carbamidomethyl: 4 | 80-93 | CID |
| 1165 | 905.099 | 2 | 654.63 | 0.5921 | 32.28 | 103.91 | 3 | 0.0 | 0 | R.VVSVLTVLHQDWLNGK.E | | 182-197 | CID |
| 1082 | 743.487 | 3 | 107.19 | 0.0797 | 30.84 | 88.75 | 5 | 0.0 | 1 | R.VVSVLTVLHQDWLNGKEYK.C | | 182-200 | CID |

#### Protein 8: Keratin, type I cytoskeletal 9 OS=Homo sapiens GN=KRT9 PE=1 SV=3

**Accession:** K1C9_HUMAN
**Database:** SwissProt
**MW [kDa] / pI:** 62.0 / 5.1

**Score:** 214.5
**Seq. Coverage [%]:** 8.3
**No. of Peptides:** 2

| 10 | 20 | 30 | 40 | 50 | 60 | 70 | 80 | 90 | 100 |
|---|---|---|---|---|---|---|---|---|---|
| MSCRQFSSSY | LSRSGGGGGG | GLGSGGSIRS | SYSRFSSSGG | GGGGGRFSSS | SGYGGGSSRV | CGRGGGGSFG | YSYGGGSGGG | FSASSLGGGF | GGGSRGFGGA |
| 110 | 120 | 130 | 140 | 150 | 160 | 170 | 180 | 190 | 200 |
| SGGGYSSSGG | FGGGFGGGSG | GGFGGGYGSG | FGGFGGFGGG | AGGGDGGILT | ANEKSTMQEL | NSRLASYLDK | VQALEEANND | LENKIQDWYD | KKGPAAIQKN |
| 210 | 220 | 230 | 240 | 250 | 260 | 270 | 280 | 290 | 300 |
| YSPYYNTIDD | LKDQIVDLTV | GNNKTLLDID | NTRMTLDDFR | IKFEMEQNLR | QGVDADINGL | RQVLDNLTME | KSDLEMQYET | LQEELMALKK | NHKEEMSQLT |
| 310 | 320 | 330 | 340 | 350 | 360 | 370 | 380 | 390 | 400 |
| GQNSGDVNVE | INVAPGKDLT | KTLNDMRQEY | EQLIAKNRKD | IENQYETQIT | QIEHEVSSSG | QEVQSSAKEV | TQLRHGVQEL | EIELQSQLSK | KAALEKSLED |
| 410 | 420 | 430 | 440 | 450 | 460 | 470 | 480 | 490 | 500 |
| TKNRYCGQLQ | MIQEQISNLE | AQITDVRQEI | ECQNQEYSLL | LSIKMRLEKE | IETYHNLLEG | GQEDFESSGA | GKIGLGGRGG | SGGSYGRGSR | GGSGGSYGGG |
| 510 | 520 | 530 | 540 | 550 | 560 | 570 | 580 | 590 | 600 |
| GSGGGYGGGS | GSRGGSGGSY | GGGSGSGGGS | GGGYGGGSGG | GHSGGSGGGH | SGGSGGNYGG | GSGSGGGSGG | GYGGGSGSRG | GSGGSHGGGS | GFGGESGGSY |
| 610 | 620 | 630 | | | | | | | |
| GGGEEASGSG | GGYGGGSGKS | SHS | | | | | | | |

MS/MS Peptide Matches

| Cmpd. | m/z meas. | z | Δ m/z [ppm] | Δ m/z [Da] | Rt [min] | Score | No. of Cmpds. | Site [%] | P | Sequence | Modification | Range | Type |
|---|---|---|---|---|---|---|---|---|---|---|---|---|---|
| 1028 | 1088.868 | 3 | 23.14 | 0.0252 | 29.94 | 69.08 | 2 | 0.0 | 0 | K.DIENQYETQITQIEHEVSSSGQEVQSSAK.E | | 340-368 | CID |
| 79 | 896.481 | 2 | 126.63 | 0.1135 | 12.14 | 145.45 | 8 | 0.0 | 0 | R.GGSGGSYGGGGSGGGYGGGSGSR.G | | 491-513 | CID |

#### Protein 9: Keratin, type II cytoskeletal 2 epidermal OS=Homo sapiens GN=KRT2 PE=1 SV=2

| | | | |
|---|---|---|---|
| **Accession:** | K22E_HUMAN | **Score:** | 184.8 |
| **Database:** | SwissProt | **Seq. Coverage [%]:** | 5.0 |
| **MW [kDa] / pI:** | 65.4 / 8.1 | **No. of Peptides:** | 2 |

| 10 | 20 | 30 | 40 | 50 | 60 | 70 | 80 | 90 | 100 |
|---|---|---|---|---|---|---|---|---|---|
| MSCQISCKSR | GRGGGGGGFR | GFSSGSAVVS | GGSRRSTSSF | SCLSRHGGGG | GGFGGGGFGS | RSLVGLGGTK | SISISVAGGG | GGFGAAGGFG | GRGGGFGGGS |
| **110** | **120** | **130** | **140** | **150** | **160** | **170** | **180** | **190** | **200** |
| SFGGGSGFSG | GGFGGGGFGG | GRFGGFGGPG | GVGGLGGPGG | FGPGGYPGGI | HEVSVNQSLL | QPLNVKVDPE | IQNVKAQERE | QIKTLNNKFA | SFIDKVRFLE |
| **210** | **220** | **230** | **240** | **250** | **260** | **270** | **280** | **290** | **300** |
| QQNQVLQTKW | ELLQQMNVGT | RPINLEPIFQ | GYIDSLKRYL | DGLTAERTSQ | NSELNNMQDL | VEDYKKKYED | EINKRTAAEN | DFVTLKKDVD | NAYMIKVELQ |
| **310** | **320** | **330** | **340** | **350** | **360** | **370** | **380** | **390** | **400** |
| SKVDLLNQEI | EFLKVLYDAE | ISQIHQSVTD | TNVILSMDNS | RNLDLDSIIA | EVKAQYEEIA | QRSKEEAEAL | YHSKYEELQV | TVGRHGDSLK | EIKIEISELN |
| **410** | **420** | **430** | **440** | **450** | **460** | **470** | **480** | **490** | **500** |
| RVIQRLQGEI | AHVKKQCKNV | QDAIADAEQR | GEHALKDARN | KLNDLEEALQ | QAKEDLARLL | RDYQELMNVK | LALDVEIATY | RKLLEGEECR | MSGDLSSNVT |
| **510** | **520** | **530** | **540** | **550** | **560** | **570** | **580** | **590** | **600** |
| VSVTSSTISS | NVASKAAFGG | SGGRGSSSGG | GYSSGSSSYG | SGGRQSGSRG | GSGGGGSISG | GGYGSGGGSG | GRYGSGGGSK | GGSISGGGYG | SGGGKHSSGG |
| **610** | **620** | **630** | **640** | | | | | | |
| GSRGGSSSGG | GYGSGGGGSS | SVKGSSGEAF | GSSVTFSFR | | | | | | |

MS/MS Peptide Matches

| Cmpd. | m/z meas. | z | Δ m/z [ppm] | Δ m/z [Da] | Rt [min] | Score | No. of Cmpds. | Site [%] | P | Sequence | Modification | Range | Type |
|---|---|---|---|---|---|---|---|---|---|---|---|---|---|
| 474 | 738.482 | 2 | 116.10 | 0.0857 | 19.95 | 78.73 | 3 | 0.0 | 0 | R.FLEQQNQVLQTK.W | | 198-209 | CID |
| 92 | 870.956 | 2 | 114.34 | 0.0996 | 12.45 | 106.03 | 2 | 0.0 | 0 | R.GSSSGGGYSSGSSSYGSGGR.Q | | 525-544 | CID |

#### Protein 10: Keratin, type I cytoskeletal 10 OS=Homo sapiens GN=KRT10 PE=1 SV=6

**Accession:** K1C10_HUMAN
**Database:** SwissProt
**MW [kDa] / pI:** 58.8 / 5.1

**Score:** 141.2
**Seq. Coverage [%]:** 3.3
**No. of Peptides:** 1

| 10 | 20 | 30 | 40 | 50 | 60 | 70 | 80 | 90 | 100 |
|---|---|---|---|---|---|---|---|---|---|
| MSVRYSSSKH | YSSSRSGGGG | GGGGCGGGGG | VSSLRISSSK | GSLGGGFSSG | GFSGGSFSRG | SSGGGCFGGS | SGGYGGLGGF | GGGSFRGSYG | SSSFGGSYGG |
| 110 | 120 | 130 | 140 | 150 | 160 | 170 | 180 | 190 | 200 |
| IFGGGSFGGG | SFGGGSFGGG | GFGGGGFGGG | FGGGFGGDGG | LLSGNEKVTM | QNLNDRLASY | LDKVRALEES | NYELEGKIKE | WYEKHGNSHQ | GEPRDYSKYY |
| 210 | 220 | 230 | 240 | 250 | 260 | 270 | 280 | 290 | 300 |
| KTIDDLKNQI | LNLTTDNANI | LLQIDNARLA | ADDFRLKYEN | EVALRQSVEA | DINGLRRVLD | ELTLTKADLE | MQIESLTEEL | AYLKKNHEEE | MKDLRNVSTG |
| 310 | 320 | 330 | 340 | 350 | 360 | 370 | 380 | 390 | 400 |
| DVNVEMNAAP | GVDLTQLLNN | MRSQYEQLAE | QNRKDAEAWF | NEKSKELTTE | IDNNIEQISS | YKSEITELRR | NVQALEIELQ | SQLALKQSLE | ASLAETEGRY |
| 410 | 420 | 430 | 440 | 450 | 460 | 470 | 480 | 490 | 500 |
| CVQLSQIQAQ | ISALEEQLQQ | IRAETECQNT | EYQQLLDIKI | RLENEIQTYR | SLLEGEGSSG | GGGRGGGSFG | GGYGGGSSGG | GSSGGGHGGG | HGGSSGGGYG |
| 510 | 520 | 530 | 540 | 550 | 560 | 570 | 580 | 590 | |
| GGSSGGGSSG | GGYGGGSSSG | GHGGSSSGGY | GGGSSGGGGG | GYGGGSSGGG | SSSGGGYGGG | SSSGGHKSSS | SGSVGESSSK | GPRY | |

MS/MS Peptide Matches

| Cmpd. | m/z meas. | z | Δ m/z [ppm] | Δ m/z [Da] | Rt [min] | Score | No. of Cmpds. | Site [%] | P | Sequence | Modification | Range | Type |
|---|---|---|---|---|---|---|---|---|---|---|---|---|---|
| 681 | 854.455 | 2 | 76.42 | 0.0653 | 23.88 | 141.21 | 3 | 0.0 | 0 | K.GSLGGGFSSGGFSGGSFSR.G | | 41-59 | CID |

#### Protein 11: Immunoglobulin kappa variable 3-20 OS=Homo sapiens GN=IGKV3-20 PE=1 SV=2

**Accession:** KV320_HUMAN
**Database:** SwissProt
**MW [kDa] / pI:** 12.5 / 4.9
**Score:** 97.1
**Seq. Coverage [%]:** 19.0
**No. of Peptides:** 1

| 10 | 20 | 30 | 40 | 50 | 60 | 70 | 80 | 90 | 100 |
|---|---|---|---|---|---|---|---|---|---|
| METPAQLLFL | LLLWLPDTTG | EIVLTQSPGT | LSLSPGERAT | LSCRASQSVS | SSYLAWYQQK | PGQAPRLLIY | GASSRATGIP | DRFSGSGSGT | DFTLTISRLE |
| 110 | 120 | | | | | | | | |
| PEDFAVYYCQ | QYGSSP | | | | | | | | |

MS/MS Peptide Matches

| Cmpd. | m/z meas. | z | Δ m/z [ppm] | Δ m/z [Da] | Rt [min] | Score | No. of Cmpds. | Site [%] | P | Sequence | Modification | Range | Type |
|---|---|---|---|---|---|---|---|---|---|---|---|---|---|
| 685 | 814.14 | 3 | 491.68 | 0.4001 | 23.95 | 97.05 | 4 | 0.0 | 0 | R.ASQSVSSSYLAWYQQKPGQAPR.L | | 45-66 | CID |

#### Protein 12: Immunoglobulin heavy variable 4-4 OS=Homo sapiens GN=IGHV4-4 PE=3 SV=2

**Accession:** HV404_HUMAN
**Database:** SwissProt
**MW [kDa] / pI:** 12.8 / 9.6
**Modification(s):** Carbamidomethyl

**Score:** 88.4
**Seq. Coverage [%]:** 13.7
**No. of Peptides:** 1

| 10 | 20 | 30 | 40 | 50 | 60 | 70 | 80 | 90 | 100 |
|---|---|---|---|---|---|---|---|---|---|
| MKHLWFFLLL | VAAPRWVLSQ | VQLQESGPGL | VKPSGTLSLT | CAVSGGSISS | SNWWSWVRQP | PGKGLEWIGE | IYHSGSTNYN | PSLKSRVTIS | VDKSKNQFSL |
| 110 | 120 | | | | | | | | |
| KLSSVTAADT | AVYYCAR | | | | | | | | |

MS/MS Peptide Matches

| Cmpd. | m/z meas. | z | Δ m/z [ppm] | Δ m/z [Da] | Rt [min] | Score | No. of Cmpds. | Site [%] | P | Sequence | Modification | Range | Type |
|---|---|---|---|---|---|---|---|---|---|---|---|---|---|
| 596 | 874.509 | 2 | 102.22 | 0.0894 | 22.35 | 88.36 | 3 | 0.0 | 0 | K.LSSVTAADTAVYYCAR.- | Carbamidomethyl: 14 | 102-117 | CID |

#### Protein 13: Immunoglobulin kappa variable 3-15 OS=Homo sapiens GN=IGKV3-15 PE=1 SV=2

**Accession:** KV315_HUMAN
**Database:** SwissProt
**MW [kDa] / pI:** 12.5 / 5.1

**Score:** 82.8
**Seq. Coverage [%]:** 18.3
**No. of Peptides:** 1

| 10 | 20 | 30 | 40 | 50 | 60 | 70 | 80 | 90 | 100 |
|---|---|---|---|---|---|---|---|---|---|
| MEAPAQLLFL | LLLWLPDTTG | EIVMTQSPAT | LSVSPGERAT | LSCRASQSVS | SNLAWYQQKP | GQAPRLLIYG | ASTRATGIPA | RFSGSGSGTE | FTLTISSLQS |
| **110** | **120** | | | | | | | | |
| EDFAVYYCQQ | YNNWP | | | | | | | | |

MS/MS Peptide Matches

| Cmpd. | m/z meas. | z | Δ m/z [ppm] | Δ m/z [Da] | Rt [min] | Score | No. of Cmpds. | Site [%] | P | Sequence | Modification | Range | Type |
|---|---|---|---|---|---|---|---|---|---|---|---|---|---|
| 547 | 768.45 | 3 | 79.27 | 0.0609 | 21.36 | 82.78 | 3 | 0.0 | 0 | R.ASQSVSSNLAWYQQKPGQAPR.L | | 45-65 | CID |

#### Protein 14: Immunoglobulin heavy variable 1-18 OS=Homo sapiens GN=IGHV1-18 PE=3 SV=1

**Accession:** HV118_HUMAN
**Database:** SwissProt
**MW [kDa] / pI:** 12.8 / 9.0
**Score:** 78.1
**Seq. Coverage [%]:** 14.5
**No. of Peptides:** 1

| 10 | 20 | 30 | 40 | 50 | 60 | 70 | 80 | 90 | 100 |
|---|---|---|---|---|---|---|---|---|---|
| MDWTWSILFL | VAAATGAHSQ | VQLVQSGAEV | KKPGASVKVS | CKASGYTFTS | YGISWVRQAP | GQGLEWMGWI | SAYNGNTNYA | QKLQGRVTMT | TDTSTSTAYM |
| **110** | **120** | | | | | | | | |
| ELRSLRSDDT | AVYYCAR | | | | | | | | |

MS/MS Peptide Matches

| Cmpd. | m/z meas. | z | Δ m/z [ppm] | Δ m/z [Da] | Rt [min] | Score | No. of Cmpds. | Site [%] | P | Sequence | Modification | Range | Type |
|---|---|---|---|---|---|---|---|---|---|---|---|---|---|
| 652 | 954.503 | 2 | 66.12 | 0.0631 | 23.34 | 78.13 | 2 | 0.0 | 0 | R.VTMTTDTSTSTAYMELR.S | | 87-103 | CID |

#### Protein 15: Immunoglobulin heavy variable 5-51 OS=Homo sapiens GN=IGHV5-51 PE=3 SV=1

**Accession:** HV551_HUMAN
**Database:** SwissProt
**MW [kDa] / pI:** 12.7 / 8.4
**Score:** 64.0
**Seq. Coverage [%]:** 12.8
**No. of Peptides:** 1

| 10 | 20 | 30 | 40 | 50 | 60 | 70 | 80 | 90 | 100 |
|---|---|---|---|---|---|---|---|---|---|
| MGSTAILALL | LAVLQGVCAE | VQLVQSGAEV | KKPGESLKIS | CKGSGYSFTS | YWIGWVRQMP | GKGLEWMGII | YPGDSDTRYS | PSFQGQVTIS | ADKSISTAYL |
| 110 | 120 | | | | | | | | |
| QWSSLKASDT | AMYYCAR | | | | | | | | |

MS/MS Peptide Matches

| Cmpd. | m/z meas. | z | Δ m/z [ppm] | Δ m/z [Da] | Rt [min] | Score | No. of Cmpds. | Site [%] | P | Sequence | Modification | Range | Type |
|---|---|---|---|---|---|---|---|---|---|---|---|---|---|
| 573 | 814.506 | 2 | 128.01 | 0.1042 | 21.85 | 63.96 | 3 | 0.0 | 0 | R.YSPSFQGQVTISADK.S | | 79-93 | CID |

#### Project Info

| | | | |
|---|---|---|---|
| **Name:** | Alagesan | **Date:** | Mar 5, 2019 |
| **Note:** | Nanobooster | | |

#### Sample Info & Protocols

| | |
|---|---|
| **Name:** | Nanobooster - IgG |
| **Date:** | Mar 5, 2019 |

#### Search Result Info

| | | | |
|---|---|---|---|
| **Search Result:** | IonTrap_Human_Swiss_CID_2019-03-05 17:55:08 | | |
| **Location:** | /Alagesan/Nanobooster - IgG/KA_IgG4 HD NB ACN CID_GA5_01_9086.mgf | | |
| **Search Method:** | IonTrap_Human_Swiss_CID | **Version:** | 1.2 |
| **Search Engine(s):** | Mascot, 2.6.0 | **Ident. Compound(s):** | 0/2367 |
| **Database(s):** | SwissProt, SwissProt_2017_04.fasta | | |
| **Note:** | | | |

#### Results

##### Protein 1: Immunoglobulin heavy constant gamma 4 OS=Homo sapiens GN=IGHG4 PE=1 SV=1

| | | | |
|---|---|---|---|
| **Accession:** | IGHG4_HUMAN | **Score:** | 1254.7 |
| **Database:** | SwissProt | **Seq. Coverage [%]:** | 65.7 |
| **MW [kDa] / pI:** | 35.9 / 7.2 | **No. of Peptides:** | 21 |
| **Modification(s):** | Carbamidomethyl, Oxidation | | |

```
        10         20         30         40         50         60         70         80         90        100
ASTKGPSVFP LAPCSRSTSE STAALGCLVK DYFPEPVTVS WNSGALTSGV HTFPAVLQSS GLYSLSSVVT VPSSSLGTKT YTCNVDHKPS NTKVDKRVES
       110        120        130        140        150        160        170        180        190        200
KYGPPCPSCP APEFLGGPSV FLFPPKPKDT LMISRTPEVT CVVVDVSQED PEVQFNWYVD GVEVHNAKTK PREEQFNSTY RVVSVLTVLH QDWLNGKEYK
       210        220        230        240        250        260        270        280        290        300
CKVSNKGLPS SIEKTISKAK GQPREPQVYT LPPSQEEMTK NQVSLTCLVK GFYPSDIAVE WESNGQPENN YKTTPPVLDS DGSFFLYSRL TVDKSRWQEG
       310        320        330
NVFSCSVMHE ALHNHYTQKS LSLSLGK
```

**MS/MS Peptide Matches**

| Cmpd. | m/z meas. | z | Δ m/z [ppm] | Δ m/z [Da] | Rt [min] | Score | No. of Cmpds. | Site [%] | P | Sequence | Modification | Range | Type |
|---|---|---|---|---|---|---|---|---|---|---|---|---|---|
| 600 | 712.428 | 2 | 97.57 | 0.0695 | 22.84 | 85.07 | 23 | 0.0 | 0 | R.STSESTAALGCLVK.D | Carbamidomethyl: 11 | 17-30 | CID |
| 51 | 833.028 | 2 | 167.51 | 0.1395 | 11.36 | 60.71 | 21 | 0.0 | 0 | K.TYTCNVDHKPSNTK.V | Carbamidomethyl: 4 | 80-93 | CID |
| 82 | 1004.084 | 2 | 99.96 | 0.1004 | 12.03 | 66.02 | 10 | 0.0 | 1 | K.TYTCNVDHKPSNTKVDK.R | Carbamidomethyl: 4 | 80-96 | CID |
| 1115 | 847.789 | 4 | 424.51 | 0.3597 | 31.64 | 18.59 | 1 | 0.0 | 1 | R.VESKYGPPCPSCPAPEFLGGPSVFLFPPKPK.D | Carbamidomethyl: 9, 12 | 98-128 | CID |
| 1208 | 982.273 | 3 | 457.39 | 0.4491 | 33.31 | 75.51 | 36 | 0.0 | 0 | K.YGPPCPSCPAPEFLGGPSVFLFPPKPK.D | Carbamidomethyl: 5, 8 | 102-128 | CID |

| Cmpd. | m/z meas. | z | Δ m/z [ppm] | Δ m/z [Da] | Rt [min] | Score | No. of Cmpds. | Site [%] | P | Sequence | Modification | Range | Type |
|---|---|---|---|---|---|---|---|---|---|---|---|---|---|
| 1199 | 941.088 | 4 | 387.09 | 0.3641 | 33.14 | 67.09 | 5 | 0.0 | 1 | K.YGPPCPSCPAPEFLGGPSVFLFPPKPKDTLMIS R.T | Carbamidomethyl: 5, 8 | 102-135 | CID |
| 1157 | 945.076 | 4 | 374.10 | 0.3534 | 32.32 | 36.23 | 3 | 0.0 | 1 | K.YGPPCPSCPAPEFLGGPSVFLFPPKPKDTLMIS R.T | Carbamidomethyl: 5, 8; Oxidation: 31 | 102-135 | CID |
| 1211 | 1152.342 | 4 | 250.63 | 0.2887 | 33.35 | 61.4 | 3 | 0.0 | 1 | K.DTLMISRTPEVTCVVVDVSQEDPEVQFNWYVD GVEVHNAK.T | Carbamidomethyl: 13 | 129-168 | CID |
| 1150 | 1263.983 | 3 | 305.90 | 0.3865 | 32.21 | 111.23 | 21 | 0.0 | 0 | R.TPEVTCVVVDVSQEDPEVQFNWYVDGVEVHN AK.T | Carbamidomethyl: 6 | 136-168 | CID |
| 1154 | 905.087 | 2 | 641.36 | 0.5801 | 32.28 | 123.35 | 3 | 0.0 | 0 | R.VVSVLTVLHQDWLNGK.E | | 182-197 | CID |
| 1066 | 743.475 | 3 | 91.05 | 0.0677 | 30.79 | 80.99 | 7 | 0.0 | 1 | R.VVSVLTVLHQDWLNGKEYK.C | | 182-200 | CID |
| 522 | 772.472 | 3 | 115.58 | 0.0893 | 21.33 | 28.37 | 4 | 0.0 | 1 | K.GQPREPQVYTLPPSQEEMTK.N | | 221-240 | CID |
| 584 | 939.033 | 2 | 84.68 | 0.0795 | 22.49 | 42.58 | 11 | 0.0 | 0 | R.EPQVYTLPPSQEEMTK.N | | 225-240 | CID |
| 532 | 946.997 | 2 | 48.63 | 0.0461 | 21.49 | 28.16 | 3 | 0.0 | 0 | R.EPQVYTLPPSQEEMTK.N | Oxidation: 14 | 225-240 | CID |
| 911 | 1007.58 | 3 | 401.77 | 0.4047 | 28.10 | 33.82 | 7 | 0.0 | 1 | R.EPQVYTLPPSQEEMTKNQVSLTCLVK.G | Carbamidomethyl: 23 | 225-250 | CID |
| 872 | 1012.875 | 3 | 363.47 | 0.3680 | 27.38 | 20.4 | 2 | 0.0 | 1 | R.EPQVYTLPPSQEEMTKNQVSLTCLVK.G | Carbamidomethyl: 23; Oxidation: 14 | 225-250 | CID |
| 931 | 1273.184 | 2 | 483.02 | 0.6147 | 28.49 | 86.26 | 7 | 0.0 | 0 | K.GFYPSDIAVEWESNGQPENNYK.T | | 251-272 | CID |
| 1016 | 951.638 | 2 | 179.07 | 0.1704 | 29.93 | 96.73 | 32 | 0.0 | 0 | K.TTPPVLDSDGSFFLYSR.L | | 273-289 | CID |
| 646 | 762.42 | 4 | 417.89 | 0.3185 | 23.62 | 43.06 | 3 | 0.0 | 1 | K.SRWQEGNVFSCSVMHEALHNHYTQK.S | Carbamidomethyl: 11 | 295-319 | CID |
| 702 | 935.163 | 3 | 436.24 | 0.4078 | 24.53 | 39.14 | 1 | 0.0 | 0 | R.WQEGNVFSCSVMHEALHNHYTQK.S | Carbamidomethyl: 9 | 297-319 | CID |
| 541 | 940.487 | 3 | 425.64 | 0.4001 | 21.64 | 49.94 | 2 | 0.0 | 0 | R.WQEGNVFSCSVMHEALHNHYTQK.S | Carbamidomethyl: 9; Oxidation: 12 | 297-319 | CID |

#### Protein 2: Immunoglobulin kappa constant OS=Homo sapiens GN=IGKC PE=1 SV=2

**Accession:** IGKC_HUMAN
**Database:** SwissProt
**MW [kDa] / pI:** 11.8 / 6.1
**Modification(s):** Carbamidomethyl

**Score:** 1024.6
**Seq. Coverage [%]:** 88.8
**No. of Peptides:** 11

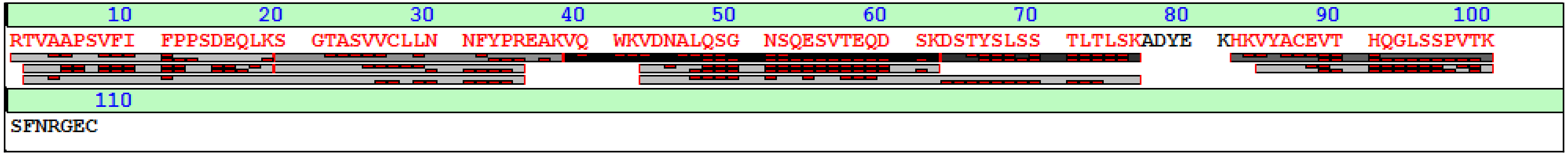


**MS/MS Peptide Matches**

| Cmpd. | m/z meas. | z | Δ m/z [ppm] | Δ m/z [Da] | Rt [min] | Score | No. of Cmpds. | Site [%] | P | Sequence | Modification | Range | Type |
|---|---|---|---|---|---|---|---|---|---|---|---|---|---|
| 955 | 701.442 | 3 | 87.15 | 0.0611 | 28.93 | 44.1 | 7 | 0.0 | 1 | -.RTVAAPSVFIFPPSDEQLK.S | | 1-19 | CID |
| 1071 | 973.662 | 2 | 148.83 | 0.1449 | 30.92 | 80.68 | 20 | 0.0 | 0 | R.TVAAPSVFIFPPSDEQLK.S | | 2-19 | CID |
| 1508 | 1242.721 | 3 | 333.82 | 0.4147 | 38.43 | 73.37 | 13 | 0.0 | 1 | R.TVAAPSVFIFPPSDEQLKSGTASVVCLLNNFYPR.E | Carbamidomethyl: 26 | 2-35 | CID |
| 977 | 899.998 | 2 | 607.87 | 0.5467 | 29.29 | 19.4 | 3 | 0.0 | 0 | K.SGTASVVCLLNNFYPR.E | Carbamidomethyl: 8 | 20-35 | CID |
| 1091 | 709.381 | 3 | 27.51 | 0.0195 | 31.26 | 94.82 | 3 | 0.0 | 1 | K.SGTASVVCLLNNFYPREAK.V | Carbamidomethyl: 8 | 20-38 | CID |
| 508 | 893.499 | 3 | 452.53 | 0.4042 | 21.05 | 118.12 | 10 | 0.0 | 1 | K.VQWKVDNALQSGNSQESVTEQDSK.D | | 39-62 | CID |
| 269 | 1068.666 | 2 | 166.59 | 0.1780 | 15.90 | 161.29 | 27 | 0.0 | 0 | K.VDNALQSGNSQESVTEQDSK.D | | 43-62 | CID |
| 721 | 1207.625 | 3 | 317.84 | 0.3837 | 24.88 | 103.23 | 2 | 0.0 | 1 | K.VDNALQSGNSQESVTEQDSKDSTYSLSSTLTLSK.A | | 43-76 | CID |
| 719 | 751.981 | 2 | 130.53 | 0.0981 | 24.85 | 85.89 | 4 | 0.0 | 0 | K.DSTYSLSSTLTLSK.A | | 63-76 | CID |
| 304 | 714.457 | 3 | 128.62 | 0.0919 | 16.92 | 136.98 | 10 | 0.0 | 1 | K.HKVYACEVTHQGLSSPVTK.S | Carbamidomethyl: 6 | 82-100 | CID |
| 405 | 938.853 | 2 | 411.20 | 0.3859 | 19.09 | 106.76 | 11 | 0.0 | 0 | K.VYACEVTHQGLSSPVTK.S | Carbamidomethyl: 4 | 84-100 | CID |

#### Protein 3: Immunoglobulin kappa light chain OS=Homo sapiens PE=1 SV=1

**Accession:** IGK_HUMAN
**Database:** SwissProt
**MW [kDa] / pI:** 23.4 / 6.9
**Modification(s):** Carbamidomethyl

**Score:** 949.7
**Seq. Coverage [%]:** 43.9
**No. of Peptides:** 9

| 10 | 20 | 30 | 40 | 50 | 60 | 70 | 80 | 90 | 100 |
|---|---|---|---|---|---|---|---|---|---|
| DIQMTQSPST | LSASVGDRVT | ITCRASQSIN | TWLAWYQQKP | GKAPKLLMYK | ASSLESGVPS | RFIGSGSGTE | FTLTISSLQP | DDFATYYCQQ | YNSDSKMFGQ |
| **110** | **120** | **130** | **140** | **150** | **160** | **170** | **180** | **190** | **200** |
| GTKVEVKGTV | AAPSVFIFPP | SDEQLKSGTA | SVVCLLNNFY | PREAKVQWKV | DNALQSGNSQ | ESVTEQDSKD | STYSLSSTLT | LSKADYEKHK | VYACEVTHQG |
| **210** | **220** | | | | | | | | |
| LSSPVTKSFN | RGEC | | | | | | | | |

MS/MS Peptide Matches

| Cmpd. | m/z meas. | z | Δ m/z [ppm] | Δ m/z [Da] | Rt [min] | Score | No. of Cmpds. | Site [%] | P | Sequence | Modification | Range | Type |
|---|---|---|---|---|---|---|---|---|---|---|---|---|---|
| 570 | 946.99 | 2 | 37.43 | 0.0354 | 22.21 | 123.19 | 4 | 0.0 | 0 | -.DIQMTQSPSTLSASVGDR.V | | 1-18 | CID |
| 977 | 899.998 | 2 | 607.87 | 0.5467 | 29.29 | 19.4 | 3 | 0.0 | 0 | K.SGTASVVCLLNNFYPR.E | Carbamidomethyl: 8 | 127-142 | CID |
| 1091 | 709.381 | 3 | 27.51 | 0.0195 | 31.26 | 94.82 | 3 | 0.0 | 1 | K.SGTASVVCLLNNFYPREAK.V | Carbamidomethyl: 8 | 127-145 | CID |
| 508 | 893.499 | 3 | 452.53 | 0.4042 | 21.05 | 118.12 | 10 | 0.0 | 1 | K.VQWKVDNALQSGNSQESVTEQDSK.D | | 146-169 | CID |
| 269 | 1068.666 | 2 | 166.59 | 0.1780 | 15.90 | 161.29 | 27 | 0.0 | 0 | K.VDNALQSGNSQESVTEQDSK.D | | 150-169 | CID |
| 721 | 1207.625 | 3 | 317.84 | 0.3837 | 24.88 | 103.23 | 2 | 0.0 | 1 | K.VDNALQSGNSQESVTEQDSKDSTYSLSSTLTLS K.A | | 150-183 | CID |
| 719 | 751.981 | 2 | 130.53 | 0.0981 | 24.85 | 85.89 | 4 | 0.0 | 0 | K.DSTYSLSSTLTLSK.A | | 170-183 | CID |
| 304 | 714.457 | 3 | 128.62 | 0.0919 | 16.92 | 136.98 | 10 | 0.0 | 1 | K.HKVYACEVTHQGLSSPVTK.S | Carbamidomethyl: 6 | 189-207 | CID |
| 405 | 938.853 | 2 | 411.20 | 0.3859 | 19.09 | 106.76 | 11 | 0.0 | 0 | K.VYACEVTHQGLSSPVTK.S | Carbamidomethyl: 4 | 191-207 | CID |

#### Protein 4: Keratin, type II cytoskeletal 1 OS=Homo sapiens GN=KRT1 PE=1 SV=6

**Accession:** K2C1_HUMAN
**Database:** SwissProt
**MW [kDa] / pI:** 66.0 / 8.2
**Modification(s):** Carbamidomethyl

**Score:** 783.6
**Seq. Coverage [%]:** 17.2
**No. of Peptides:** 6

| 10 | 20 | 30 | 40 | 50 | 60 | 70 | 80 | 90 | 100 |
|---|---|---|---|---|---|---|---|---|---|
| MSRQFSSRSG | YRSGGGFSSG | SAGIINYQRR | TTSSSTRRSG | GGGGRFSSCG | GGGGSFGAGG | GFGSRSLVNL | GGSKSISISV | ARGGGRGSGF | GGGYGGGGFG |
| 110 | 120 | 130 | 140 | 150 | 160 | 170 | 180 | 190 | 200 |
| GGGFGGGGFG | GGGIGGGGFG | GFGSGGGGFG | GGGFGGGGYG | GGYGPVCPPG | GIQEVTINQS | LLQPLNVEID | PEIQKVKSRE | REQIKSLNNQ | FASFIDKVRF |
| 210 | 220 | 230 | 240 | 250 | 260 | 270 | 280 | 290 | 300 |
| LEQQNQVLQT | KWELLQQVDT | STRTHNLEPY | FESFINNLRR | RVDQLKSDQS | RLDSELKNMQ | DMVEDYRNKY | EDEINKRTNA | ENEFVTIKKD | VDGAYMTKVD |
| 310 | 320 | 330 | 340 | 350 | 360 | 370 | 380 | 390 | 400 |
| LQAKLDNLQQ | EIDFLTALYQ | AELSQMQTQI | SETNVILSMD | NNRSLDLDSI | IAEVKAQYED | IAQKSKAEAE | SLYQSKYEEL | QITAGRHGDS | VRNSKIEISE |
| 410 | 420 | 430 | 440 | 450 | 460 | 470 | 480 | 490 | 500 |
| LNRVIQRLRS | EIDNVKKQIS | NLQQSISDAE | QRGENALKDA | KNKLNDLEDA | LQQAKEDLAR | LLRDYQELMN | TKLALDLEIA | TYRTLLEGEE | SRMSGECAPN |
| 510 | 520 | 530 | 540 | 550 | 560 | 570 | 580 | 590 | 600 |
| VSVSVSTSHT | TISGGGSRGG | GGGGYGSGGS | SYGSGGGSYG | SGGGGGGGRG | SYGSGGSSYG | SGGGSYGSGG | GGGGHGSYGS | GSSSGGYRGG | SGGGGGGSSG |
| 610 | 620 | 630 | 640 | 650 | | | | | |
| GRGSGGGSSG | GSIGGRGSSS | GGVKSSGGSS | SVKFVSTTYS | GVTR | | | | | |

**MS/MS Peptide Matches**

| Cmpd. | m/z meas. | z | Δ m/z [ppm] | Δ m/z [Da] | Rt [min] | Score | No. of Cmpds. | Site [%] | P | Sequence | Modification | Range | Type |
|---|---|---|---|---|---|---|---|---|---|---|---|---|---|
| 463 | 883.463 | 2 | 104.16 | 0.0920 | 20.18 | 140.58 | 3 | 0.0 | 0 | R.FSSCGGGGGSFGAGGGFGSR.S | Carbamidomethyl: 4 | 46-65 | CID |
| 453 | 738.519 | 2 | 166.21 | 0.1227 | 20.02 | 81.47 | 5 | 0.0 | 0 | R.FLEQQNQVLQTK.W | | 200-211 | CID |
| 986 | 978.568 | 3 | 399.77 | 0.3910 | 29.43 | 106.59 | 3 | 0.0 | 1 | R.FLEQQNQVLQTKWELLQQVDTSTR.T | | 200-223 | CID |
| 1471 | 783.819 | 3 | 95.29 | 0.0747 | 37.77 | 163.71 | 3 | 0.0 | 1 | R.SLDLDSIIAEVKAQYEDIAQK.S | | 344-364 | CID |
| 539 | 859.021 | 2 | 106.88 | 0.0918 | 21.63 | 110.91 | 3 | 0.0 | 0 | K.QISNLQQSISDAEQR.G | | 418-432 | CID |
| 197 | 1192.543 | 2 | 53.20 | 0.0634 | 14.41 | 180.37 | 3 | 0.0 | 0 | R.GGGGGGYGSGGSSYGSGGGSYGSGGGGGG GR.G | | 519-549 | CID |

BRUKER

##### Protein 5: Keratin, type II cytoskeletal 2 epidermal OS=Homo sapiens GN=KRT2 PE=1 SV=2

| | | | |
|---|---|---|---|
| **Accession:** | K22E_HUMAN | **Score:** | 642.9 |
| **Database:** | SwissProt | **Seq. Coverage [%]:** | 21.3 |
| **MW [kDa] / pI:** | 65.4 / 8.1 | **No. of Peptides:** | 7 |

```
        10         20         30         40         50         60         70         80         90        100
MSCQISCKSR GRGGGGGGFR GFSSGSAVVS GGSRRSTSSF SCLSRHGGGG GGFGGGGFGS RSLVGLGGTK SISISVAGGG GGFGAAGGFG GRGGGFGGGS
       110        120        130        140        150        160        170        180        190        200
SFGGGSGFSG GGFGGGGFGG GRFGGFGGPG GVGGLGGPGG FGPGGYPGGI HEVSVNQSLL QPLNVKVDPE IQNVKAQERE QIKTLNNKFA SFIDKVRFLE
       210        220        230        240        250        260        270        280        290        300
QQNQVLQTKW ELLQQMNVGT RPINLEPIFQ GYIDSLKRYL DGLTAERTSQ NSELNNMQDL VEDYKKKYED EINKRTAAEN DFVTLKKDVD NAYMIKVELQ
       310        320        330        340        350        360        370        380        390        400
SKVDLLNQEI EFLKVLYDAE ISQIHQSVTD TNVILSMDNS RNLDLDSIIA EVKAQYEEIA QRSKEEAEAL YHSKYEELQV TVGRHGDSLK EIKIEISELN
       410        420        430        440        450        460        470        480        490        500
RVIQRLQGEI AHVKKQCKNV QDAIADAEQR GEHALKDARN KLNDLEEALQ QAKEDLARLL RDYQELMNVK LALDVEIATY RKLLEGEECR MSGDLSSNVT
       510        520        530        540        550        560        570        580        590        600
VSVTSSTISS NVASKAAFGG SGGRGSSSGG GYSSGSSSYG SGGRQSGSRG GSGGGGSISG GGYGSGGGSG GRYGSGGGSK GGSISGGGYG SGGGKHSSGG
       610        620        630        640
GSRGGSSSGG GYGSGGGGSS SVKGSSGEAF GSSVTFSFR
```

MS/MS Peptide Matches

| Cmpd. | m/z meas. | z | Δ m/z [ppm] | Δ m/z [Da] | Rt [min] | Score | No. of Cmpds. | Site [%] | P | Sequence | Modification | Range | Type |
|---|---|---|---|---|---|---|---|---|---|---|---|---|---|
| 233 | 705.937 | 2 | 112.14 | 0.0792 | 15.16 | 18.36 | 1 | 0.0 | 1 | R.GFSSGSAVVSGGSRR.S | | 21-35 | CID |
| 808 | 920.04 | 2 | 86.06 | 0.0792 | 26.34 | 156.81 | 2 | 0.0 | 0 | K.SISISVAGGGGGFGAAGGFGGR.G | | 71-92 | CID |
| 453 | 738.519 | 2 | 166.21 | 0.1227 | 20.02 | 81.47 | 5 | 0.0 | 0 | R.FLEQQNQVLQTK.W | | 198-209 | CID |

| Cmpd. | m/z meas. | z | Δ m/z [ppm] | Δ m/z [Da] | Rt [min] | Score | No. of Cmpds. | Site [%] | P | Sequence | Modification | Range | Type |
|---|---|---|---|---|---|---|---|---|---|---|---|---|---|
| 1534 | 806.806 | 3 | 62.95 | 0.0508 | 38.85 | 152.11 | 2 | 0.0 | 1 | R.NLDLDSIIAEVKAQYEEIAQR.S | | 342-362 | CID |
| 135 | 871.472 | 2 | 108.04 | 0.0941 | 13.18 | 138.11 | 2 | 0.0 | 0 | R.GGSGGGGSISGGGYGSGGGSGGR.Y | | 550-572 | CID |
| 34 | 962.005 | 2 | 69.01 | 0.0664 | 10.96 | 24.22 | 1 | 0.0 | 1 | K.GGSISGGGYGSGGGKHSSGGGSR.G | | 581-603 | CID |
| 33 | 772.416 | 3 | 530.90 | 0.4099 | 10.95 | 71.82 | 3 | 0.0 | 1 | K.HSSGGGSRGGSSSGGGYGSGGGGSSSVK.G | | 596-623 | CID |

#### Protein 6: Immunoglobulin lambda constant 3 OS=Homo sapiens GN=IGLC3 PE=1 SV=1

**Accession:** IGLC3_HUMAN
**Database:** SwissProt
**MW [kDa] / pI:** 11.3 / 6.9
**Modification(s):** Carbamidomethyl

**Score:** 580.5
**Seq. Coverage [%]:** 93.4
**No. of Peptides:** 8

10 20 30 40 50 60 70 80 90 100
GQPKAAPSVT LFPPSSEELQ ANKATLVCLI SDFYPGAVTV AWKADSSPVK AGVETTTPSK QSNNKYAASS YLSLTPEQWK SHKSYSCQVT HEGSTVEKTV
110
APTECS

MS/MS Peptide Matches

| Cmpd. | m/z meas. | z | Δ m/z [ppm] | Δ m/z [Da] | Rt [min] | Score | No. of Cmpds. | Site [%] | P | Sequence | Modification | Range | Type |
|---|---|---|---|---|---|---|---|---|---|---|---|---|---|
| 834 | 993.555 | 2 | 42.72 | 0.0424 | 26.76 | 69.84 | 8 | 0.0 | 0 | K.AAPSVTLFPPSSEELQANK.A | | 5-23 | CID |
| 1601 | 1393.626 | 3 | 170.23 | 0.2372 | 39.93 | 18.54 | 1 | 0.0 | 1 | K.AAPSVTLFPPSSEELQANKATLVCLISDFYPGAV TVAWK.A | Carbamidomethyl: 24 | 5-43 | CID |
| 227 | 838 | 2 | 82.50 | 0.0691 | 15.01 | 71.48 | 3 | 0.0 | 1 | K.ADSSPVKAGVETTTPSK.Q | | 44-60 | CID |
| 26 | 781.535 | 2 | 179.86 | 0.1405 | 10.77 | 89.37 | 6 | 0.0 | 1 | K.AGVETTTPSKQSNNK.Y | | 51-65 | CID |
| 798 | 772.479 | 3 | 126.10 | 0.0974 | 26.16 | 61.19 | 8 | 0.0 | 1 | K.QSNNKYAASSYLSLTPEQWK.S | | 61-80 | CID |
| 892 | 872.533 | 2 | 114.56 | 0.0999 | 27.76 | 84.24 | 9 | 0.0 | 0 | K.YAASSYLSLTPEQWK.S | | 66-80 | CID |
| 183 | 856.556 | 2 | 201.74 | 0.1728 | 14.14 | 97.86 | 7 | 0.0 | 0 | K.SYSCQVTHEGSTVEK.T | Carbamidomethyl: 4 | 84-98 | CID |

| Cmpd. | m/z meas. | z | Δ m/z [ppm] | Δ m/z [Da] | Rt [min] | Score | No. of Cmpds. | Site [%] | P | Sequence | Modification | Range | Type |
|---|---|---|---|---|---|---|---|---|---|---|---|---|---|
| 390 | 1279.549 | 2 | 380.21 | 0.4863 | 18.81 | 87.97 | 9 | 0.0 | 1 | K.SYSCQVTHEGSTVEKTVAPTECS.- | Carbamidomethyl: 4, 22 | 84-106 | CID |

#### Protein 7: Immunoglobulin heavy constant gamma 1 OS=Homo sapiens GN=IGHG1 PE=1 SV=1

| | | | |
|---|---|---|---|
| **Accession:** | IGHG1_HUMAN | **Score:** | 366.7 |
| **Database:** | SwissProt | **Seq. Coverage [%]:** | 17.6 |
| **MW [kDa] / pI:** | 36.1 / 8.5 | **No. of Peptides:** | 4 |

| 10 | 20 | 30 | 40 | 50 | 60 | 70 | 80 | 90 | 100 |
|---|---|---|---|---|---|---|---|---|---|
| ASTKGPSVFP | LAPSSKSTSG | GTAALGCLVK | DYFPEPVTVS | WNSGALTSGV | HTFPAVLQSS | GLYSLSSVVT | VPSSSLGTQT | YICNVNHKPS | NTKVDKKVEP |
| **110** | **120** | **130** | **140** | **150** | **160** | **170** | **180** | **190** | **200** |
| KSCDKTHTCP | PCPAPELLGG | PSVFLFPPKP | KDTLMISRTP | EVTCVVVDVS | HEDPEVKFNW | YVDGVEVHNA | KTKPREEQYN | STYRVVSVLT | VLHQDWLNGK |
| **210** | **220** | **230** | **240** | **250** | **260** | **270** | **280** | **290** | **300** |
| EYKCKVSNKA | LPAPIEKTIS | KAKGQPREPQ | VYTLPPSRDE | LTKNQVSLTC | LVKGFYPSDI | AVEWESNGQP | ENNYKTTPPV | LDSDGSFFLY | SKLTVDKSRW |
| **310** | **320** | **330** | **340** | | | | | | |
| QQGNVFSCSV | MHEALHNHYT | QKSLSLSPGK | | | | | | | |

MS/MS Peptide Matches

| Cmpd. | m/z meas. | z | Δ m/z [ppm] | Δ m/z [Da] | Rt [min] | Score | No. of Cmpds. | Site [%] | P | Sequence | Modification | Range | Type |
|---|---|---|---|---|---|---|---|---|---|---|---|---|---|
| 1154 | 905.087 | 2 | 641.36 | 0.5801 | 32.28 | 123.35 | 3 | 0.0 | 0 | R.VVSVLTVLHQDWLNGK.E | | 185-200 | CID |
| 1066 | 743.475 | 3 | 91.05 | 0.0677 | 30.79 | 80.99 | 7 | 0.0 | 1 | R.VVSVLTVLHQDWLNGKEYK.C | | 185-203 | CID |
| 931 | 1273.184 | 2 | 483.02 | 0.6147 | 28.49 | 86.26 | 7 | 0.0 | 0 | K.GFYPSDIAVEWESNGQPENNYK.T | | 254-275 | CID |
| 1001 | 937.567 | 2 | 109.29 | 0.1025 | 29.66 | 76.08 | 3 | 0.0 | 0 | K.TTPPVLDSDGSFFLYSK.L | | 276-292 | CID |

BRUKER

#### Protein 8: Keratin, type I cytoskeletal 10 OS=Homo sapiens GN=KRT10 PE=1 SV=6

| | | | |
|---|---|---|---|
| **Accession:** | K1C10_HUMAN | **Score:** | 345.9 |
| **Database:** | SwissProt | **Seq. Coverage [%]:** | 9.9 |
| **MW [kDa] / pI:** | 58.8 / 5.1 | **No. of Peptides:** | 3 |

| 10 | 20 | 30 | 40 | 50 | 60 | 70 | 80 | 90 | 100 |
|---|---|---|---|---|---|---|---|---|---|
| MSVRYSSSKH | YSSSRSGGGG | GGGGCGGGGG | VSSLRISSSK | GSLGGGFSSG | GFSGGSFSRG | SSGGGCFGGS | SGGYGGLGGF | GGGSFRGSYG | SSSFGGSYGG |
| 110 | 120 | 130 | 140 | 150 | 160 | 170 | 180 | 190 | 200 |
| IFGGGSFGGG | SFGGGSFGGG | GFGGGGFGGG | FGGGFGGDGG | LLSGNEKVTM | QNLNDRLASY | LDKVRALEES | NYELEGKIKE | WYEKHGNSHQ | GEPRDYSKYY |
| 210 | 220 | 230 | 240 | 250 | 260 | 270 | 280 | 290 | 300 |
| KTIDDLKNQI | LNLTTDNANI | LLQIDNARLA | ADDFRLKYEN | EVALRQSVEA | DINGLRRVLD | ELTLTKADLE | MQIESLTEEL | AYLKKNHEEE | MKDLRNVSTG |
| 310 | 320 | 330 | 340 | 350 | 360 | 370 | 380 | 390 | 400 |
| DVNVEMNAAP | GVDLTQLLNN | MRSQYEQLAE | QNRKDAEAWF | NEKSKELTTE | IDNNIEQISS | YKSEITELRR | NVQALEIELQ | SQLALKQSLE | ASLAETEGRY |
| 410 | 420 | 430 | 440 | 450 | 460 | 470 | 480 | 490 | 500 |
| CVQLSQIQAQ | ISALEEQLQQ | IRAETECQNT | EYQQLLDIKI | RLENEIQTYR | SLLEGEGSSG | GGGRGGGSFG | GGYGGGSSGG | GSSGGGHGGG | HGGSSGGGYG |
| 510 | 520 | 530 | 540 | 550 | 560 | 570 | 580 | 590 | |
| GGSSGGGSSG | GGYGGGSSSG | GHGGSSSGGY | GGGSSGGGGG | GYGGGSSGGG | SSSGGGYGGG | SSSGGHKSSS | SGSVGESSSK | GPRY | |

MS/MS Peptide Matches

| Cmpd. | m/z meas. | z | Δ m/z [ppm] | Δ m/z [Da] | Rt [min] | Score | No. of Cmpds. | Site [%] | P | Sequence | Modification | Range | Type |
|---|---|---|---|---|---|---|---|---|---|---|---|---|---|
| 653 | 854.447 | 2 | 67.06 | 0.0573 | 23.76 | 156.11 | 3 | 0.0 | 0 | K.GSLGGGFSSGGFSGGSFSR.G | | 41-59 | CID |
| 1301 | 1437.165 | 2 | 323.64 | 0.4650 | 34.83 | 150.36 | 5 | 0.0 | 0 | R.NVSTGDVNVEMNAAPGVDLTQLLNNMR.S | | 296-322 | CID |
| 221 | 747.398 | 2 | 36.41 | 0.0272 | 14.90 | 39.41 | 4 | 0.0 | 1 | R.SQYEQLAEQNRK.D | | 323-334 | CID |

#### Protein 9: Keratin, type I cytoskeletal 9 OS=Homo sapiens GN=KRT9 PE=1 SV=3

| | | | |
|---|---|---|---|
| **Accession:** | K1C9_HUMAN | **Score:** | 161.0 |
| **Database:** | SwissProt | **Seq. Coverage [%]:** | 3.7 |
| **MW [kDa] / pI:** | 62.0 / 5.1 | **No. of Peptides:** | 1 |

| 10 | 20 | 30 | 40 | 50 | 60 | 70 | 80 | 90 | 100 |
|---|---|---|---|---|---|---|---|---|---|
| MSCRQFSSSY | LSRSGGGGGG | GLGSGGSIRS | SYSRFSSSGG | GGGGGRFSSS | SGYGGGSSRV | CGRGGGGSFG | YSYGGGSGGG | FSASSLGGGF | GGGSRGFGGA |
| 110 | 120 | 130 | 140 | 150 | 160 | 170 | 180 | 190 | 200 |
| SGGGYSSSGG | FGGGFGGGSG | GGFGGGYGSG | FGGFGGFGGG | AGGGDGGILT | ANEKSTMQEL | NSRLASYLDK | VQALEEANND | LENKIQDWYD | KKGPAAIQKN |
| 210 | 220 | 230 | 240 | 250 | 260 | 270 | 280 | 290 | 300 |
| YSPYYNTIDD | LKDQIVDLTV | GNNKTLLDID | NTRMTLDDFR | IKFEMEQNLR | QGVDADINGL | RQVLDNLTME | KSDLEMQYET | LQEELMALKK | NHKEEMSQLT |
| 310 | 320 | 330 | 340 | 350 | 360 | 370 | 380 | 390 | 400 |
| GQNSGDVNVE | INVAPGKDLT | KTLNDMRQEY | EQLIAKNRKD | IENQYETQIT | QIEHEVSSSG | QEVQSSAKEV | TQLRHGVQEL | EIELQSQLSK | KAALEKSLED |
| 410 | 420 | 430 | 440 | 450 | 460 | 470 | 480 | 490 | 500 |
| TKNRYCGQLQ | MIQEQISNLE | AQITDVRQEI | ECQNQEYSLL | LSIKMRLEKE | IETYHNLLEG | GQEDFESSGA | GKIGLGGRGG | SGGSYGRGSR | GGSGGSYGGG |
| 510 | 520 | 530 | 540 | 550 | 560 | 570 | 580 | 590 | 600 |
| GSGGGYGGGS | GSRGGSGGSY | GGGSGSGGGS | GGGYGGGSGG | GHSGGSGGGH | SGGSGGNYGG | GSGSGGGSGG | GYGGGSGSRG | GSGGSHGGGS | GFGGESGGSY |
| 610 | 620 | 630 | | | | | | | |
| GGGEEASGSG | GGYGGGSGKS | SHS | | | | | | | |

MS/MS Peptide Matches

| Cmpd. | m/z meas. | z | Δ m/z [ppm] | Δ m/z [Da] | Rt [min] | Score | No. of Cmpds. | Site [%] | P | Sequence | Modification | Range | Type |
|---|---|---|---|---|---|---|---|---|---|---|---|---|---|
| 91 | 896.426 | 2 | 65.27 | 0.0585 | 12.22 | 160.99 | 4 | 0.0 | 0 | R.GGSGGSYGGGGSGGGYGGGSGSR.G | | 491-513 | CID |

#### Protein 10: Immunoglobulin heavy constant gamma 2 OS=Homo sapiens GN=IGHG2 PE=1 SV=2

| | | | |
|---|---|---|---|
| **Accession:** | IGHG2_HUMAN | **Score:** | 127.4 |
| **Database:** | SwissProt | **Seq. Coverage [%]:** | 10.1 |
| **MW [kDa] / pI:** | 35.9 / 7.7 | **No. of Peptides:** | 2 |
| **Modification(s):** | Carbamidomethyl | | |

| 10 | 20 | 30 | 40 | 50 | 60 | 70 | 80 | 90 | 100 |
|---|---|---|---|---|---|---|---|---|---|
| ASTKGPSVFP | LAPCSRSTSE | STAALGCLVK | DYFPEPVTVS | WNSGALTSGV | HTFPAVLQSS | GLYSLSSVVT | VPSSNFGTQT | YTCNVDHKPS | NTKVDKTVER |
| 110 | 120 | 130 | 140 | 150 | 160 | 170 | 180 | 190 | 200 |
| KCCVECPPCP | APPVAGPSVF | LFPPKPKDTL | MISRTPEVTC | VVVDVSHEDP | EVQFNWYVDG | VEVHNAKTKP | REEQFNSTFR | VVSVLTVVHQ | DWLNGKEYKC |
| 210 | 220 | 230 | 240 | 250 | 260 | 270 | 280 | 290 | 300 |
| KVSNKGLPAP | IEKTISKTKG | QPREPQVYTL | PPSREEMTKN | QVSLTCLVKG | FYPSDISVEW | ESNGQPENNY | KTTPPMLDSD | GSFFLYSKLT | VDKSRWQQGN |
| 310 | 320 | 330 | | | | | | | |
| VFSCSVMHEA | LHNHYTQKSL | SLSPGK | | | | | | | |

MS/MS Peptide Matches

| Cmpd. | m/z meas. | z | Δ m/z [ppm] | Δ m/z [Da] | Rt [min] | Score | No. of Cmpds. | Site [%] | P | Sequence | Modification | Range | Type |
|---|---|---|---|---|---|---|---|---|---|---|---|---|---|
| 600 | 712.428 | 2 | 97.57 | 0.0695 | 22.84 | 85.07 | 23 | 0.0 | 0 | R.STSESTAALGCLVK.D | Carbamidomethyl: 11 | 17-30 | CID |
| 946 | 739.122 | 3 | 523.29 | 0.3866 | 28.77 | 42.31 | 3 | 0.0 | 1 | R.VVSVLTVVHQDWLNGKEYK.C | | 181-199 | CID |

#### Protein 11: Immunoglobulin lambda variable 3-21 OS=Homo sapiens GN=IGLV3-21 PE=1 SV=2

| | | | |
|---|---|---|---|
| **Accession:** | LV321_HUMAN | **Score:** | 100.1 |
| **Database:** | SwissProt | **Seq. Coverage [%]:** | 13.7 |
| **MW [kDa] / pI:** | 12.4 / 5.0 | **No. of Peptides:** | 1 |

| 10 | 20 | 30 | 40 | 50 | 60 | 70 | 80 | 90 | 100 |
|---|---|---|---|---|---|---|---|---|---|
| MAWTVLLLGL | LSHCTGSVTS | YVLTQPPSVS | VAPGQTARIT | CGGNNIGSKS | VHWYQQKPGQ | APVLVVYDDS | DRPSGIPERF | SGSNSGNTAT | LTISRVEAGD |
| 110 | 120 | | | | | | | | |
| EADYYCQVWD | SSSDHPT | | | | | | | | |

MS/MS Peptide Matches

| Cmpd. | m/z meas. | z | Δ m/z [ppm] | Δ m/z [Da] | Rt [min] | Score | No. of Cmpds. | Site [%] | P | Sequence | Modification | Range | Type |
|---|---|---|---|---|---|---|---|---|---|---|---|---|---|
| 414 | 806.979 | 2 | 98.02 | 0.0791 | 19.31 | 100.11 | 3 | 0.0 | 0 | R.FSGSNSGNTATLTISR.V | | 80-95 | CID |

#### Protein 12: Immunoglobulin heavy variable 4-4 OS=Homo sapiens GN=IGHV4-4 PE=3 SV=2

**Accession:** HV404_HUMAN
**Database:** SwissProt
**MW [kDa] / pI:** 12.8 / 9.6
**Modification(s):** Carbamidomethyl

**Score:** 92.6
**Seq. Coverage [%]:** 13.7
**No. of Peptides:** 1

| 10 | 20 | 30 | 40 | 50 | 60 | 70 | 80 | 90 | 100 |
|---|---|---|---|---|---|---|---|---|---|
| MKHLWFFLLL | VAAPRWVLSQ | VQLQESGPGL | VKPSGTLSLT | CAVSGGSISS | SNWWSWVRQP | PGKGLEWIGE | IYHSGSTNYN | PSLKSRVTIS | VDKSKNQFSL |
| 110 | 120 | | | | | | | | |
| KLSSVTAADT | AVYYCAR | | | | | | | | |

MS/MS Peptide Matches

| Cmpd. | m/z meas. | z | Δ m/z [ppm] | Δ m/z [Da] | Rt [min] | Score | No. of Cmpds. | Site [%] | P | Sequence | Modification | Range | Type |
|---|---|---|---|---|---|---|---|---|---|---|---|---|---|
| 578 | 874.511 | 2 | 104.51 | 0.0914 | 22.37 | 92.59 | 1 | 0.0 | 0 | K.LSSVTAADTAVYYCAR.- | Carbamidomethyl: 14 | 102-117 | CID |

BRUKER

#### Protein 13: Immunoglobulin kappa variable 3-15 OS=Homo sapiens GN=IGKV3-15 PE=1 SV=2

| | | | |
|---|---|---|---|
| **Accession:** | KV315_HUMAN | **Score:** | 82.7 |
| **Database:** | SwissProt | **Seq. Coverage [%]:** | 18.3 |
| **MW [kDa] / pI:** | 12.5 / 5.1 | **No. of Peptides:** | 1 |

| 10 | 20 | 30 | 40 | 50 | 60 | 70 | 80 | 90 | 100 |
|---|---|---|---|---|---|---|---|---|---|
| MEAPAQLLFL | LLLWLPDTTG | EIVMTQSPAT | LSVSPGERAT | LSCRASQSVS | SNLAWYQQKP | GQAPRLLIYG | ASTRATGIPA | RFSGSGSGTE | FTLTISSLQS |
| 110 | 120 | | | | | | | | |
| EDFAVYYCQQ | YNNWP | | | | | | | | |

MS/MS Peptide Matches

| Cmpd. | m/z meas. | z | Δ m/z [ppm] | Δ m/z [Da] | Rt [min] | Score | No. of Cmpds. | Site [%] | P | Sequence | Modification | Range | Type |
|---|---|---|---|---|---|---|---|---|---|---|---|---|---|
| 527 | 768.457 | 3 | 88.38 | 0.0679 | 21.43 | 82.69 | 3 | 0.0 | 0 | R.ASQSVSSNLAWYQQKPGQAPR.L | | 45-65 | CID |

#### Protein 14: Keratin, type II cytoskeletal 5 OS=Homo sapiens GN=KRT5 PE=1 SV=3

| | | | |
|---|---|---|---|
| **Accession:** | K2C5_HUMAN | **Score:** | 75.1 |
| **Database:** | SwissProt | **Seq. Coverage [%]:** | 3.6 |
| **MW [kDa] / pI:** | 62.3 / 7.6 | **No. of Peptides:** | 1 |

| | | | | | | | | | |
|---|---|---|---|---|---|---|---|---|---|
| 10 | 20 | 30 | 40 | 50 | 60 | 70 | 80 | 90 | 100 |
| MSRQSSVSFR | SGGSRSFSTA | SAITPSVSRT | SFTSVSRSGG | GGGGGFGRVS | LAGACGVGGY | GSRSLYNLGG | SKRISISTSG | GSFRNRFGAG | AGGGYGFGGG |
| 110 | 120 | 130 | 140 | 150 | 160 | 170 | 180 | 190 | 200 |
| AGSGFGFGGG | AGGGFGLGGG | AGFGGGFGGP | GFPVCPPGGI | QEVTVNQSLL | TPLNLQIDPS | IQRVRTEERE | QIKTLNNKFA | SFIDKVRFLE | QQNKVLDTKW |
| 210 | 220 | 230 | 240 | 250 | 260 | 270 | 280 | 290 | 300 |
| TLLQEQGTKT | VRQNLEPLFE | QYINNLRRQL | DSIVGERGRL | DSELRNMQDL | VEDFKNKYED | EINKRTTAEN | EFVMLKKDVD | AAYMNKVELE | AKVDALMDEI |
| 310 | 320 | 330 | 340 | 350 | 360 | 370 | 380 | 390 | 400 |
| NFMKMFFDAE | LSQMQTHVSD | TSVVLSMDNN | RNLDLDSIIA | EVKAQYEEIA | NRSRTEAESW | YQTKYEELQQ | TAGRHGDDLR | NTKHEISEMN | RMIQRLRAEI |
| 410 | 420 | 430 | 440 | 450 | 460 | 470 | 480 | 490 | 500 |
| DNVKKQCANL | QNAIADAEQR | GELALKDARN | KLAELEEALQ | KAKQDMARLL | REYQELMNTK | LALDVEIATY | RKLLEGEECR | LSGEGVGPVN | ISVVTSSVSS |
| 510 | 520 | 530 | 540 | 550 | 560 | 570 | 580 | 590 | 600 |
| GYGSGSGYGG | GLGGGLGGGL | GGGLAGGSSG | SYYSSSSGGV | GLGGGLSVGG | SGFSASSGRG | LGVGFGSGGG | SSSSVKFVST | TSSSRKSFKS | |

**MS/MS Peptide Matches**

| Cmpd. | m/z meas. | z | Δ m/z [ppm] | Δ m/z [Da] | Rt [min] | Score | No. of Cmpds. | Site [%] | P | Sequence | Modification | Range | Type |
|---|---|---|---|---|---|---|---|---|---|---|---|---|---|
| 1543 | 802.085 | 3 | 2.08 | 0.0017 | 39.00 | 75.13 | 1 | 0.0 | 1 | R.NLDLDSIIAEVKAQYEEIANR.S | | 332-352 | CID |

#### Protein 15: Immunoglobulin kappa variable 2D-28 OS=Homo sapiens GN=IGKV2D-28 PE=1 SV=2

| | | | |
|---|---|---|---|
| **Accession:** | KVD28_HUMAN | **Score:** | 45.1 |
| **Database:** | SwissProt | **Seq. Coverage [%]:** | 16.7 |
| **MW [kDa] / pI:** | 12.9 / 5.6 | **No. of Peptides:** | 1 |

| | | | | | | | | | |
|---|---|---|---|---|---|---|---|---|---|
| 10 | 20 | 30 | 40 | 50 | 60 | 70 | 80 | 90 | 100 |
| MRLPAQLLGL | LMLWVSGSSG | DIVMTQSPLS | LPVTPGEPAS | ISCRSSQSLL | HSNGYNYLDW | YLQKPGQSPQ | LLIYLGSNRA | SGVPDRFSGS | GSGTDFTLKI |
| 110 | 120 | 130 | | | | | | | |
| SRVEAEDVGV | YYCMQALQTP | | | | | | | | |

**MS/MS Peptide Matches**

| Cmpd. | m/z meas. | z | Δ m/z [ppm] | Δ m/z [Da] | Rt [min] | Score | No. of Cmpds. | Site [%] | P | Sequence | Modification | Range | Type |
|---|---|---|---|---|---|---|---|---|---|---|---|---|---|
| 581 | 993.52 | 2 | 38.46 | 0.0382 | 22.43 | 45.05 | 3 | 0.0 | 1 | R.ASGVPDRFSGSGSGTDFTLK.I | | 80-99 | CID |

#### Project Info

| | | | |
|---|---|---|---|
| **Name:** | Alagesan | **Date:** | Mar 5, 2019 |
| **Note:** | Nanobooster | | |

#### Sample Info & Protocols

| | |
|---|---|
| **Name:** | Nanobooster - IgG |
| **Date:** | Mar 5, 2019 |

#### Search Result Info

| | | | |
|---|---|---|---|
| **Search Result:** | IonTrap_Human_Swiss_CID_2019-03-05 17:55:08 | | |
| **Location:** | /Alagesan/Nanobooster - IgG/KA_IgG4 HD NB ACN CID_GA5_01_9087.mgf | | |
| **Search Method:** | IonTrap_Human_Swiss_CID | **Version:** | 1.2 |
| **Search Engine(s):** | Mascot, 2.6.0 | **Ident. Compound(s):** | 0/2424 |
| **Database(s):** | SwissProt, SwissProt_2017_04.fasta | | |
| **Note:** | | | |

#### Results

##### Protein 1: Immunoglobulin heavy constant gamma 4 OS=Homo sapiens GN=IGHG4 PE=1 SV=1

| | | | |
|---|---|---|---|
| **Accession:** | IGHG4_HUMAN | **Score:** | 1315.7 |
| **Database:** | SwissProt | **Seq. Coverage [%]:** | 64.5 |
| **MW [kDa] / pI:** | 35.9 / 7.2 | **No. of Peptides:** | 20 |
| **Modification(s):** | Carbamidomethyl, Oxidation | | |

| 10 | 20 | 30 | 40 | 50 | 60 | 70 | 80 | 90 | 100 |
|---|---|---|---|---|---|---|---|---|---|
| ASTKGPSVFP | LAPCSRSTSE | STAALGCLVK | DYFPEPVTVS | WNSGALTSGV | HTFPAVLQSS | GLYSLSSVVT | VPSSSLGTKT | YTCNVDHKPS | NTKVDKRVES |
| **110** | **120** | **130** | **140** | **150** | **160** | **170** | **180** | **190** | **200** |
| KYGPPCPSCP | APEFLGGPSV | FLFPPKPKDT | LMISRTPEVT | CVVVDVSQED | PEVQFNWYVD | GVEVHNAKTK | PREEQFNSTY | RVVSVLTVLH | QDWLNGKEYK |
| **210** | **220** | **230** | **240** | **250** | **260** | **270** | **280** | **290** | **300** |
| CKVSNKGLPS | SIEKTISKAK | GQPREPQVYT | LPPSQEEMTK | NQVSLTCLVK | GFYPSDIAVE | WESNGQPENN | YKTTPPVLDS | DGSFFLYSRL | TVDKSRWQEG |
| **310** | **320** | **330** | | | | | | | |
| NVFSCSVMHE | ALHNHYTQKS | LSLSLGK | | | | | | | |

**MS/MS Peptide Matches**

| Cmpd. | m/z meas. | z | Δ m/z [ppm] | Δ m/z [Da] | Rt [min] | Score | No. of Cmpds. | Site [%] | P | Sequence | Modification | Range | Type |
|---|---|---|---|---|---|---|---|---|---|---|---|---|---|
| 605 | 712.431 | 2 | 101.78 | 0.0725 | 22.47 | 88.02 | 21 | 0.0 | 0 | R.STSESTAALGCLVK.D | Carbamidomethyl: 11 | 17-30 | CID |
| 91 | 833.41 | 2 | 626.15 | 0.5215 | 11.88 | 70.32 | 18 | 0.0 | 0 | K.TYTCNVDHKPSNTK.V | Carbamidomethyl: 4 | 80-93 | CID |
| 99 | 1004.195 | 2 | 210.52 | 0.2114 | 12.06 | 63.89 | 6 | 0.0 | 1 | K.TYTCNVDHKPSNTKVDK.R | Carbamidomethyl: 4 | 80-96 | CID |
| 1241 | 982.271 | 3 | 455.36 | 0.4471 | 33.51 | 71.21 | 51 | 0.0 | 0 | K.YGPPCPSCPAPEFLGGPSVFLFPPKPK.D | Carbamidomethyl: 5, 8 | 102-128 | CID |

| Cmpd. | m/z meas. | z | Δ m/z [ppm] | Δ m/z [Da] | Rt [min] | Score | No. of Cmpds. | Site [%] | P | Sequence | Modification | Range | Type |
|---|---|---|---|---|---|---|---|---|---|---|---|---|---|
| 1228 | 941.069 | 4 | 366.90 | 0.3451 | 33.27 | 47.68 | 7 | 0.0 | 1 | K.YGPPCPSCPAPEFLGGPSVFLFPPKPKDTLMIS R.T | Carbamidomethyl: 5, 8 | 102-135 | CID |
| 1188 | 1259.636 | 3 | 271.30 | 0.3417 | 32.49 | 49.6 | 2 | 0.0 | 1 | K.YGPPCPSCPAPEFLGGPSVFLFPPKPKDTLMIS R.T | Carbamidomethyl: 5, 8; Oxidation: 31 | 102-135 | CID |
| 1232 | 1152.35 | 4 | 257.57 | 0.2967 | 33.36 | 84.09 | 2 | 0.0 | 1 | K.DTLMISRTPEVTCVVVDVSQEDPEVQFNWYVD GVEVHNAK.T | Carbamidomethyl: 13 | 129-168 | CID |
| 1255 | 1263.908 | 3 | 246.55 | 0.3115 | 33.73 | 124.58 | 14 | 0.0 | 0 | R.TPEVTCVVVDVSQEDPEVQFNWYVDGVEVHN AK.T | Carbamidomethyl: 6 | 136-168 | CID |
| 1176 | 905.088 | 2 | 642.47 | 0.5811 | 32.28 | 127.05 | 3 | 0.0 | 0 | R.VVSVLTVLHQDWLNGK.E | | 182-197 | CID |
| 1087 | 743.507 | 3 | 134.10 | 0.0997 | 30.75 | 85.79 | 4 | 0.0 | 1 | R.VVSVLTVLHQDWLNGKEYK.C | | 182-200 | CID |
| 539 | 772.484 | 3 | 131.12 | 0.1013 | 21.28 | 27.66 | 1 | 0.0 | 1 | K.GQPREPQVYTLPPSQEEMTK.N | | 221-240 | CID |
| 600 | 939.065 | 2 | 118.76 | 0.1115 | 22.38 | 44.92 | 11 | 0.0 | 0 | R.EPQVYTLPPSQEEMTK.N | | 225-240 | CID |
| 545 | 946.978 | 2 | 28.57 | 0.0271 | 21.38 | 30.27 | 9 | 0.0 | 0 | R.EPQVYTLPPSQEEMTK.N | Oxidation: 14 | 225-240 | CID |
| 915 | 1007.632 | 3 | 453.40 | 0.4567 | 27.74 | 41.96 | 10 | 0.0 | 1 | R.EPQVYTLPPSQEEMTKNQVSLTCLVK.G | Carbamidomethyl: 23 | 225-250 | CID |
| 837 | 1012.887 | 3 | 375.32 | 0.3800 | 26.42 | 32.5 | 4 | 0.0 | 1 | R.EPQVYTLPPSQEEMTKNQVSLTCLVK.G | Carbamidomethyl: 23; Oxidation: 14 | 225-250 | CID |
| 949 | 1273.196 | 2 | 492.45 | 0.6267 | 28.40 | 104.52 | 8 | 0.0 | 0 | K.GFYPSDIAVEWESNGQPENNYK.T | | 251-272 | CID |
| 1041 | 951.613 | 2 | 152.79 | 0.1454 | 29.97 | 99.87 | 63 | 0.0 | 0 | K.TTPPVLDSDGSFFLYSR.L | | 273-289 | CID |
| 671 | 1016.158 | 3 | 352.82 | 0.3584 | 23.66 | 33.48 | 1 | 0.0 | 1 | K.SRWQEGNVFSCSVMHEALHNHYTQK.S | Carbamidomethyl: 11 | 295-319 | CID |
| 726 | 935.149 | 3 | 421.26 | 0.3938 | 24.55 | 60.94 | 2 | 0.0 | 0 | R.WQEGNVFSCSVMHEALHNHYTQK.S | Carbamidomethyl: 9 | 297-319 | CID |
| 563 | 940.484 | 3 | 422.44 | 0.3971 | 21.67 | 27.38 | 2 | 0.0 | 0 | R.WQEGNVFSCSVMHEALHNHYTQK.S | Carbamidomethyl: 9; Oxidation: 12 | 297-319 | CID |

#### Protein 2: Immunoglobulin kappa constant OS=Homo sapiens GN=IGKC PE=1 SV=2

**Accession:** IGKC_HUMAN
**Database:** SwissProt
**MW [kDa] / pI:** 11.8 / 6.1
**Modification(s):** Carbamidomethyl

**Score:** 1137.9
**Seq. Coverage [%]:** 88.8
**No. of Peptides:** 11

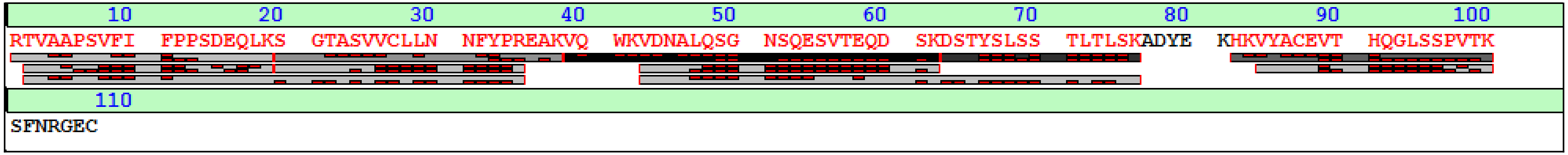


MS/MS Peptide Matches

| Cmpd. | m/z meas. | z | Δ m/z [ppm] | Δ m/z [Da] | Rt [min] | Score | No. of Cmpds. | Site [%] | P | Sequence | Modification | Range | Type |
|---|---|---|---|---|---|---|---|---|---|---|---|---|---|
| 973 | 701.466 | 3 | 121.37 | 0.0851 | 28.83 | 42.1 | 5 | 0.0 | 1 | -.RTVAAPSVFIFPPSDEQLK.S | | 1-19 | CID |
| 1098 | 973.655 | 2 | 141.64 | 0.1379 | 30.98 | 98.64 | 23 | 0.0 | 0 | R.TVAAPSVFIFPPSDEQLK.S | | 2-19 | CID |
| 1552 | 1242.697 | 3 | 314.50 | 0.3907 | 38.72 | 97.86 | 14 | 0.0 | 1 | R.TVAAPSVFIFPPSDEQLKSGTASVVCLLNNFYP R.E | Carbamidomethyl: 26 | 2-35 | CID |
| 1218 | 899.53 | 2 | 87.55 | 0.0787 | 33.08 | 55.01 | 4 | 0.0 | 0 | K.SGTASVVCLLNNFYPR.E | Carbamidomethyl: 8 | 20-35 | CID |
| 1109 | 709.432 | 3 | 99.41 | 0.0705 | 31.19 | 128.18 | 3 | 0.0 | 1 | K.SGTASVVCLLNNFYPREAK.V | Carbamidomethyl: 8 | 20-38 | CID |
| 532 | 893.484 | 3 | 435.73 | 0.3892 | 21.11 | 131.5 | 11 | 0.0 | 1 | K.VQWKVDNALQSGNSQESVTEQDSK.D | | 39-62 | CID |
| 253 | 1068.541 | 2 | 49.60 | 0.0530 | 15.22 | 156.98 | 34 | 0.0 | 0 | K.VDNALQSGNSQESVTEQDSK.D | | 43-62 | CID |
| 728 | 1207.594 | 3 | 292.16 | 0.3527 | 24.59 | 80.38 | 4 | 0.0 | 1 | K.VDNALQSGNSQESVTEQDSKDSTYSLSSTLTLS K.A | | 43-76 | CID |
| 741 | 751.965 | 2 | 109.25 | 0.0821 | 24.82 | 94.05 | 4 | 0.0 | 0 | K.DSTYSLSSTLTLSK.A | | 63-76 | CID |
| 330 | 714.437 | 3 | 100.62 | 0.0719 | 17.06 | 137.3 | 7 | 0.0 | 1 | K.HKVYACEVTHQGLSSPVTK.S | Carbamidomethyl: 6 | 82-100 | CID |
| 447 | 938.517 | 2 | 53.17 | 0.0499 | 19.51 | 115.89 | 13 | 0.0 | 0 | K.VYACEVTHQGLSSPVTK.S | Carbamidomethyl: 4 | 84-100 | CID |

#### Protein 3: Immunoglobulin kappa light chain OS=Homo sapiens PE=1 SV=1

**Accession:** IGK_HUMAN
**Database:** SwissProt
**MW [kDa] / pI:** 23.4 / 6.9
**Modification(s):** Carbamidomethyl

**Score:** 1031.2
**Seq. Coverage [%]:** 43.9
**No. of Peptides:** 9

| 10 | 20 | 30 | 40 | 50 | 60 | 70 | 80 | 90 | 100 |
|---|---|---|---|---|---|---|---|---|---|
| DIQMTQSPST | LSASVGDRVT | ITCRASQSIN | TWLAWYQQKP | GKAPKLLMYK | ASSLESGVPS | RFIGSGSGTE | FTLTISSLQP | DDFATYYCQQ | YNSDSKMFGQ |
| **110** | **120** | **130** | **140** | **150** | **160** | **170** | **180** | **190** | **200** |
| GTKVEVKGTV | AAPSVFIFPP | SDEQLKSGTA | SVVCLLNNFY | PREAKVQWKV | DNALQSGNSQ | ESVTEQDSKD | STYSLSSTLT | LSKADYEKHK | VYACEVTHQG |
| **210** | **220** | | | | | | | | |
| LSSPVTKSFN | RGEC | | | | | | | | |

MS/MS Peptide Matches

| Cmpd. | m/z meas. | z | Δ m/z [ppm] | Δ m/z [Da] | Rt [min] | Score | No. of Cmpds. | Site [%] | P | Sequence | Modification | Range | Type |
|---|---|---|---|---|---|---|---|---|---|---|---|---|---|
| 592 | 947.002 | 2 | 50.10 | 0.0474 | 22.22 | 131.89 | 5 | 0.0 | 0 | -.DIQMTQSPSTLSASVGDR.V | | 1-18 | CID |
| 1218 | 899.53 | 2 | 87.55 | 0.0787 | 33.08 | 55.01 | 4 | 0.0 | 0 | K.SGTASVVCLLNNFYPR.E | Carbamidomethyl: 8 | 127-142 | CID |
| 1109 | 709.432 | 3 | 99.41 | 0.0705 | 31.19 | 128.18 | 3 | 0.0 | 1 | K.SGTASVVCLLNNFYPREAK.V | Carbamidomethyl: 8 | 127-145 | CID |
| 532 | 893.484 | 3 | 435.73 | 0.3892 | 21.11 | 131.5 | 11 | 0.0 | 1 | K.VQWKVDNALQSGNSQESVTEQDSK.D | | 146-169 | CID |
| 253 | 1068.541 | 2 | 49.60 | 0.0530 | 15.22 | 156.98 | 34 | 0.0 | 0 | K.VDNALQSGNSQESVTEQDSK.D | | 150-169 | CID |
| 728 | 1207.594 | 3 | 292.16 | 0.3527 | 24.59 | 80.38 | 4 | 0.0 | 1 | K.VDNALQSGNSQESVTEQDSKDSTYSLSSTLTLSK.A | | 150-183 | CID |
| 741 | 751.965 | 2 | 109.25 | 0.0821 | 24.82 | 94.05 | 4 | 0.0 | 0 | K.DSTYSLSSTLTLSK.A | | 170-183 | CID |
| 330 | 714.437 | 3 | 100.62 | 0.0719 | 17.06 | 137.3 | 7 | 0.0 | 1 | K.HKVYACEVTHQGLSSPVTK.S | Carbamidomethyl: 6 | 189-207 | CID |
| 447 | 938.517 | 2 | 53.17 | 0.0499 | 19.51 | 115.89 | 13 | 0.0 | 0 | K.VYACEVTHQGLSSPVTK.S | Carbamidomethyl: 4 | 191-207 | CID |

#### Protein 4: Keratin, type II cytoskeletal 1 OS=Homo sapiens GN=KRT1 PE=1 SV=6

**Accession:** K2C1_HUMAN
**Database:** SwissProt
**MW [kDa] / pI:** 66.0 / 8.2
**Modification(s):** Carbamidomethyl

**Score:** 763.0
**Seq. Coverage [%]:** 17.2
**No. of Peptides:** 6

| 10 | 20 | 30 | 40 | 50 | 60 | 70 | 80 | 90 | 100 |
|---|---|---|---|---|---|---|---|---|---|
| MSRQFSSRSG | YRSGGGFSSG | SAGIINYQRR | TTSSSTRRSG | GGGGRFSSCG | GGGGSFGAGG | GFGSRSLVNL | GGSKSISISV | ARGGGRGSGF | GGGYGGGGFG |
| 110 | 120 | 130 | 140 | 150 | 160 | 170 | 180 | 190 | 200 |
| GGGFGGGGFG | GGGIGGGGFG | GFGSGGGGFG | GGGFGGGGYG | GGYGPVCPPG | GIQEVTINQS | LLQPLNVEID | PEIQKVKSRE | REQIKSLNNQ | FASFIDKVRF |
| 210 | 220 | 230 | 240 | 250 | 260 | 270 | 280 | 290 | 300 |
| LEQQNQVLQT | KWELLQQVDT | STRTHNLEPY | FESFINNLRR | RVDQLKSDQS | RLDSELKNMQ | DMVEDYRNKY | EDEINKRTNA | ENEFVTIKKD | VDGAYMTKVD |
| 310 | 320 | 330 | 340 | 350 | 360 | 370 | 380 | 390 | 400 |
| LQAKLDNLQQ | EIDFLTALYQ | AELSQMQTQI | SETNVILSMD | NNRSLDLDSI | IAEVKAQYED | IAQKSKAEAE | SLYQSKYEEL | QITAGRHGDS | VRNSKIEISE |
| 410 | 420 | 430 | 440 | 450 | 460 | 470 | 480 | 490 | 500 |
| LNRVIQRLRS | EIDNVKKQIS | NLQQSISDAE | QRGENALKDA | KNKLNDLEDA | LQQAKEDLAR | LLRDYQELMN | TKLALDLEIA | TYRTLLEGEE | SRMSGECAPN |
| 510 | 520 | 530 | 540 | 550 | 560 | 570 | 580 | 590 | 600 |
| VSVSVSTSHT | TISGGGSRGG | GGGGYGSGGS | SYGSGGGSYG | SGGGGGGGRG | SYGSGGSSYG | SGGGSYGSGG | GGGGHGSYGS | GSSSGGYRGG | SGGGGGGSSG |
| 610 | 620 | 630 | 640 | 650 | | | | | |
| GRGSGGGSSG | GSIGGRGSSS | GGVKSSGGSS | SVKFVSTTYS | GVTR | | | | | |

**MS/MS Peptide Matches**

| Cmpd. | m/z meas. | z | Δ m/z [ppm] | Δ m/z [Da] | Rt [min] | Score | No. of Cmpds. | Site [%] | P | Sequence | Modification | Range | Type |
|---|---|---|---|---|---|---|---|---|---|---|---|---|---|
| 482 | 883.458 | 2 | 98.50 | 0.0870 | 20.16 | 133.98 | 3 | 0.0 | 0 | R.FSSCGGGGGSFGAGGGFGSR.S | Carbamidomethyl: 4 | 46-65 | CID |
| 470 | 738.484 | 2 | 118.81 | 0.0877 | 19.96 | 88.6 | 5 | 0.0 | 0 | R.FLEQQNQVLQTK.W | | 200-211 | CID |
| 1012 | 978.269 | 3 | 94.10 | 0.0920 | 29.49 | 96.16 | 3 | 0.0 | 1 | R.FLEQQNQVLQTKWELLQQVDTSTR.T | | 200-223 | CID |
| 1493 | 783.835 | 3 | 115.71 | 0.0907 | 37.74 | 136.81 | 3 | 0.0 | 1 | R.SLDLDSIIAEVKAQYEDIAQK.S | | 344-364 | CID |
| 562 | 859.004 | 2 | 87.09 | 0.0748 | 21.67 | 105.73 | 5 | 0.0 | 0 | K.QISNLQQSISDAEQR.G | | 418-432 | CID |
| 211 | 1192.942 | 2 | 387.80 | 0.4624 | 14.37 | 201.76 | 4 | 0.0 | 0 | R.GGGGGGYGSGGSSYGSGGGSYGSGGGGGG GR.G | | 519-549 | CID |

#### Protein 5: Immunoglobulin lambda constant 3 OS=Homo sapiens GN=IGLC3 PE=1 SV=1

| | | | |
|---|---|---|---|
| **Accession:** | IGLC3_HUMAN | **Score:** | 565.4 |
| **Database:** | SwissProt | **Seq. Coverage [%]:** | 74.5 |
| **MW [kDa] / pI:** | 11.3 / 6.9 | **No. of Peptides:** | 7 |
| **Modification(s):** | Carbamidomethyl | | |

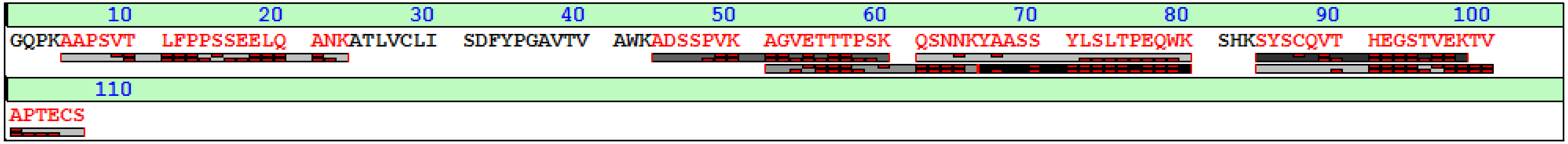


MS/MS Peptide Matches

| Cmpd. | m/z meas. | z | Δ m/z [ppm] | Δ m/z [Da] | Rt [min] | Score | No. of Cmpds. | Site [%] | P | Sequence | Modification | Range | Type |
|---|---|---|---|---|---|---|---|---|---|---|---|---|---|
| 857 | 993.564 | 2 | 51.77 | 0.0514 | 26.78 | 55.7 | 9 | 0.0 | 0 | K.AAPSVTLFPPSSEELQANK.A | | 5-23 | CID |
| 246 | 838.012 | 2 | 96.82 | 0.0811 | 15.08 | 83.06 | 3 | 0.0 | 1 | K.ADSSPVKAGVETTTPSK.Q | | 44-60 | CID |
| 42 | 781.538 | 2 | 183.70 | 0.1435 | 10.76 | 62.48 | 5 | 0.0 | 1 | K.AGVETTTPSKQSNNK.Y | | 51-65 | CID |
| 828 | 772.46 | 3 | 101.50 | 0.0784 | 26.29 | 68.63 | 5 | 0.0 | 1 | K.QSNNKYAASSYLSLTPEQWK.S | | 61-80 | CID |
| 918 | 872.557 | 2 | 142.07 | 0.1239 | 27.81 | 88.21 | 7 | 0.0 | 0 | K.YAASSYLSLTPEQWK.S | | 66-80 | CID |
| 198 | 856.497 | 2 | 132.85 | 0.1138 | 14.15 | 101.16 | 7 | 0.0 | 0 | K.SYSCQVTHEGSTVEK.T | Carbamidomethyl: 4 | 84-98 | CID |
| 408 | 1279.587 | 2 | 409.92 | 0.5243 | 18.78 | 106.15 | 9 | 0.0 | 1 | K.SYSCQVTHEGSTVEKTVAPTECS.- | Carbamidomethyl: 4, 22 | 84-106 | CID |

#### Protein 6: Keratin, type II cytoskeletal 2 epidermal OS=Homo sapiens GN=KRT2 PE=1 SV=2

| | | | |
|---|---|---|---|
| **Accession:** | K22E_HUMAN | **Score:** | 517.4 |
| **Database:** | SwissProt | **Seq. Coverage [%]:** | 16.6 |
| **MW [kDa] / pI:** | 65.4 / 8.1 | **No. of Peptides:** | 5 |

| 10 | 20 | 30 | 40 | 50 | 60 | 70 | 80 | 90 | 100 |
|---|---|---|---|---|---|---|---|---|---|
| MSCQISCKSR | GRGGGGGGFR | GFSSGSAVVS | GGSRRSTSSF | SCLSRHGGGG | GGFGGGGFGS | RSLVGLGGTK | SISISVAGGG | GGFGAAGGFG | GRGGGFGGGS |
| **110** | **120** | **130** | **140** | **150** | **160** | **170** | **180** | **190** | **200** |
| SFGGGSGFSG | GGFGGGGFGG | GRFGGFGGPG | GVGGLGGPGG | FGPGGYPGGI | HEVSVNQSLL | QPLNVKVDPE | IQNVKAQERE | QIKTLNNKFA | SFIDKVRFLE |
| **210** | **220** | **230** | **240** | **250** | **260** | **270** | **280** | **290** | **300** |
| QQNQVLQTKW | ELLQQMNVGT | RPINLEPIFQ | GYIDSLKRYL | DGLTAERTSQ | NSELNNMQDL | VEDYKKKYED | EINKRTAAEN | DFVTLKKDVD | NAYMIKVELQ |
| **310** | **320** | **330** | **340** | **350** | **360** | **370** | **380** | **390** | **400** |
| SKVDLLNQEI | EFLKVLYDAE | ISQIHQSVTD | TNVILSMDNS | RNLDLDSIIA | EVKAQYEEIA | QRSKEEAEAL | YHSKYEELQV | TVGRHGDSLK | EIKIEISELN |
| **410** | **420** | **430** | **440** | **450** | **460** | **470** | **480** | **490** | **500** |
| RVIQRLQGEI | AHVKKQCKNV | QDAIADAEQR | GEHALKDARN | KLNDLEEALQ | QAKEDLARLL | RDYQELMNVK | LALDVEIATY | RKLLEGEECR | MSGDLSSNVT |
| **510** | **520** | **530** | **540** | **550** | **560** | **570** | **580** | **590** | **600** |
| VSVTSSTISS | NVASKAAFGG | SGGRGSSSGG | GYSSGSSSYG | SGGRQSGSRG | GSGGGGSISG | GGYGSGGGSG | GRYGSGGGSK | GGSISGGGYG | SGGGKHSSGG |
| **610** | **620** | **630** | **640** | | | | | | |
| GSRGGSSSGG | GYGSGGGGSS | SVKGSSGEAF | GSSVTFSFR | | | | | | |

**MS/MS Peptide Matches**

| Cmpd. | m/z meas. | z | Δ m/z [ppm] | Δ m/z [Da] | Rt [min] | Score | No. of Cmpds. | Site [%] | P | Sequence | Modification | Range | Type |
|---|---|---|---|---|---|---|---|---|---|---|---|---|---|
| 832 | 920.024 | 2 | 68.67 | 0.0632 | 26.34 | 148.57 | 3 | 0.0 | 0 | K.SISISVAGGGGGFGAAGGFGGR.G | | 71-92 | CID |
| 470 | 738.484 | 2 | 118.81 | 0.0877 | 19.96 | 88.6 | 5 | 0.0 | 0 | R.FLEQQNQVLQTK.W | | 198-209 | CID |
| 1562 | 807.137 | 3 | 473.23 | 0.3818 | 38.87 | 144.21 | 2 | 0.0 | 1 | R.NLDLDSIIAEVKAQYEEIAQR.S | | 342-362 | CID |
| 150 | 871.465 | 2 | 100.00 | 0.0871 | 13.19 | 106.37 | 2 | 0.0 | 0 | R.GGSGGGGSISGGGYGSGGGSGGR.Y | | 550-572 | CID |
| 55 | 772.437 | 3 | 558.10 | 0.4309 | 11.09 | 29.61 | 1 | 0.0 | 1 | K.HSSGGGSRGGSSSGGGYGSGGGGSSSVK.G | | 596-623 | CID |

#### Protein 7: Immunoglobulin heavy constant gamma 1 OS=Homo sapiens GN=IGHG1 PE=1 SV=1

| | | | |
|---|---|---|---|
| **Accession:** | IGHG1_HUMAN | **Score:** | 391.6 |
| **Database:** | SwissProt | **Seq. Coverage [%]:** | 17.6 |
| **MW [kDa] / pI:** | 36.1 / 8.5 | **No. of Peptides:** | 4 |

| 10 | 20 | 30 | 40 | 50 | 60 | 70 | 80 | 90 | 100 |
|---|---|---|---|---|---|---|---|---|---|
| ASTKGPSVFP | LAPSSKSTSG | GTAALGCLVK | DYFPEPVTVS | WNSGALTSGV | HTFPAVLQSS | GLYSLSSVVT | VPSSSLGTQT | YICNVNHKPS | NTKVDKKVEP |
| 110 | 120 | 130 | 140 | 150 | 160 | 170 | 180 | 190 | 200 |
| KSCDKTHTCP | PCPAPELLGG | PSVFLFPPKP | KDTLMISRTP | EVTCVVVDVS | HEDPEVKFNW | YVDGVEVHNA | KTKPREEQYN | STYRVVSVLT | VLHQDWLNGK |
| 210 | 220 | 230 | 240 | 250 | 260 | 270 | 280 | 290 | 300 |
| EYKCKVSNKA | LPAPIEKTIS | KAKGQPREPQ | VYTLPPSRDE | LTKNQVSLTC | LVKGFYPSDI | AVEWESNGQP | ENNYKTTPPV | LDSDGSFFLY | SKLTVDKSRW |
| 310 | 320 | 330 | 340 | | | | | | |
| QQGNVFSCSV | MHEALHNHYT | QKSLSLSPGK | | | | | | | |

**MS/MS Peptide Matches**

| Cmpd. | m/z meas. | z | Δ m/z [ppm] | Δ m/z [Da] | Rt [min] | Score | No. of Cmpds. | Site [%] | P | Sequence | Modification | Range | Type |
|---|---|---|---|---|---|---|---|---|---|---|---|---|---|
| 1176 | 905.088 | 2 | 642.47 | 0.5811 | 32.28 | 127.05 | 3 | 0.0 | 0 | R.VVSVLTVLHQDWLNGK.E | | 185-200 | CID |
| 1087 | 743.507 | 3 | 134.10 | 0.0997 | 30.75 | 85.79 | 4 | 0.0 | 1 | R.VVSVLTVLHQDWLNGKEYK.C | | 185-203 | CID |
| 949 | 1273.196 | 2 | 492.45 | 0.6267 | 28.40 | 104.52 | 8 | 0.0 | 0 | K.GFYPSDIAVEWESNGQPENNYK.T | | 254-275 | CID |
| 1024 | 937.563 | 2 | 105.02 | 0.0985 | 29.67 | 74.21 | 3 | 0.0 | 0 | K.TTPPVLDSDGSFFLYSK.L | | 276-292 | CID |

#### Protein 8: Keratin, type I cytoskeletal 10 OS=Homo sapiens GN=KRT10 PE=1 SV=6

**Accession:** K1C10_HUMAN
**Database:** SwissProt
**MW [kDa] / pI:** 58.8 / 5.1

**Score:** 337.8
**Seq. Coverage [%]:** 9.9
**No. of Peptides:** 3

| 10 | 20 | 30 | 40 | 50 | 60 | 70 | 80 | 90 | 100 |
|---|---|---|---|---|---|---|---|---|---|
| MSVRYSSSKH | YSSSRSGGGG | GGGGCGGGGG | VSSLRISSSK | GSLGGGFSSG | GFSGGSFSRG | SSGGGCFGGS | SGGYGGLGGF | GGGSFRGSYG | SSSFGGSYGG |
| 110 | 120 | 130 | 140 | 150 | 160 | 170 | 180 | 190 | 200 |
| IFGGGSFGGG | SFGGGSFGGG | GFGGGGFGGG | FGGGFGGDGG | LLSGNEKVTM | QNLNDRLASY | LDKVRALEES | NYELEGKIKE | WYEKHGNSHQ | GEPRDYSKYY |
| 210 | 220 | 230 | 240 | 250 | 260 | 270 | 280 | 290 | 300 |
| KTIDDLKNQI | LNLTTDNANI | LLQIDNARLA | ADDFRLKYEN | EVALRQSVEA | DINGLRRVLD | ELTLTKADLE | MQIESLTEEL | AYLKKNHEEE | MKDLRNVSTG |
| 310 | 320 | 330 | 340 | 350 | 360 | 370 | 380 | 390 | 400 |
| DVNVEMNAAP | GVDLTQLLNN | MRSQYEQLAE | QNRKDAEAWF | NEKSKELTTE | IDNNIEQISS | YKSEITELRR | NVQALEIELQ | SQLALKQSLE | ASLAETEGRY |
| 410 | 420 | 430 | 440 | 450 | 460 | 470 | 480 | 490 | 500 |
| CVQLSQIQAQ | ISALEEQLQQ | IRAETECQNT | EYQQLLDIKI | RLENEIQTYR | SLLEGEGSSG | GGGRGGGSFG | GGYGGGSSGG | GSSGGGHGGG | HGGSSGGGYG |
| 510 | 520 | 530 | 540 | 550 | 560 | 570 | 580 | 590 | |
| GGSSGGGSSG | GGYGGGSSSG | GHGGSSSGGY | GGGSSGGGGG | GYGGGSSGGG | SSSGGGYGGG | SSSGGHKSSS | SGSVGESSSK | GPRY | |

**MS/MS Peptide Matches**

| Cmpd. | m/z meas. | z | Δ m/z [ppm] | Δ m/z [Da] | Rt [min] | Score | No. of Cmpds. | Site [%] | P | Sequence | Modification | Range | Type |
|---|---|---|---|---|---|---|---|---|---|---|---|---|---|
| 677 | 854.467 | 2 | 90.47 | 0.0773 | 23.78 | 157.13 | 3 | 0.0 | 0 | K.GSLGGGFSSGGFSGGSFSR.G | | 41-59 | CID |
| 1320 | 1437.117 | 2 | 290.23 | 0.4170 | 34.78 | 145.08 | 4 | 0.0 | 0 | R.NVSTGDVNVEMNAAPGVDLTQLLNNMR.S | | 296-322 | CID |
| 242 | 747.417 | 2 | 61.84 | 0.0462 | 14.99 | 35.54 | 4 | 0.0 | 1 | R.SQYEQLAEQNRK.D | | 323-334 | CID |

#### Protein 9: Immunoglobulin heavy constant gamma 3 OS=Homo sapiens GN=IGHG3 PE=1 SV=2

**Accession:** IGHG3_HUMAN
**Database:** SwissProt
**MW [kDa] / pI:** 41.3 / 8.2
**Modification(s):** Carbamidomethyl

**Score:** 255.5
**Seq. Coverage [%]:** 10.3
**No. of Peptides:** 3

| 10 | 20 | 30 | 40 | 50 | 60 | 70 | 80 | 90 | 100 |
|---|---|---|---|---|---|---|---|---|---|
| ASTKGPSVFP | LAPCSRSTSG | GTAALGCLVK | DYFPEPVTVS | WNSGALTSGV | HTFPAVLQSS | GLYSLSSVVT | VPSSSLGTQT | YTCNVNHKPS | NTKVDKRVEL |
| **110** | **120** | **130** | **140** | **150** | **160** | **170** | **180** | **190** | **200** |
| KTPLGDTTHT | CPRCPEPKSC | DTPPPCPRCP | EPKSCDTPPP | CPRCPEPKSC | DTPPPCPRCP | APELLGGPSV | FLFPPKPKDT | LMISRTPEVT | CVVVDVSHED |
| **210** | **220** | **230** | **240** | **250** | **260** | **270** | **280** | **290** | **300** |
| PEVQFKWYVD | GVEVHNAKTK | PREEQYNSTF | RVVSVLTVLH | QDWLNGKEYK | CKVSNKALPA | PIEKTISKTK | GQPREPQVYT | LPPSREEMTK | NQVSLTCLVK |
| **310** | **320** | **330** | **340** | **350** | **360** | **370** | **380** | | |
| GFYPSDIAVE | WESSGQPENN | YNTTPPMLDS | DGSFFLYSKL | TVDKSRWQQG | NIFSCSVMHE | ALHNRFTQKS | LSLSPGK | | |

**MS/MS Peptide Matches**

| Cmpd. | m/z meas. | z | Δ m/z [ppm] | Δ m/z [Da] | Rt [min] | Score | No. of Cmpds. | Site [%] | P | Sequence | Modification | Range | Type |
|---|---|---|---|---|---|---|---|---|---|---|---|---|---|
| 1144 | 717.771 | 3 | 61.00 | 0.0438 | 31.76 | 42.61 | 1 | 0.0 | 0 | R.CPAPELLGGPSVFLFPPKPK.D | Carbamidomethyl: 1 | 159-178 | CID |
| 1176 | 905.088 | 2 | 642.47 | 0.5811 | 32.28 | 127.05 | 3 | 0.0 | 0 | R.VVSVLTVLHQDWLNGK.E | | 232-247 | CID |
| 1087 | 743.507 | 3 | 134.10 | 0.0997 | 30.75 | 85.79 | 4 | 0.0 | 1 | R.VVSVLTVLHQDWLNGKEYK.C | | 232-250 | CID |

#### Protein 10: Keratin, type I cytoskeletal 9 OS=Homo sapiens GN=KRT9 PE=1 SV=3

**Accession:** K1C9_HUMAN
**Database:** SwissProt
**MW [kDa] / pI:** 62.0 / 5.1

**Score:** 166.5
**Seq. Coverage [%]:** 3.7
**No. of Peptides:** 1

| 10 | 20 | 30 | 40 | 50 | 60 | 70 | 80 | 90 | 100 |
|---|---|---|---|---|---|---|---|---|---|
| MSCRQFSSSY | LSRSGGGGGG | GLGSGGSIRS | SYSRFSSSGG | GGGGGRFSSS | SGYGGGSSRV | CGRGGGGSFG | YSYGGGSGGG | FSASSLGGGF | GGGSRGFGGA |
| 110 | 120 | 130 | 140 | 150 | 160 | 170 | 180 | 190 | 200 |
| SGGGYSSSGG | FGGGFGGGSG | GGFGGGYGSG | FGGFGGFGGG | AGGGDGGILT | ANEKSTMQEL | NSRLASYLDK | VQALEEANND | LENKIQDWYD | KKGPAAIQKN |
| 210 | 220 | 230 | 240 | 250 | 260 | 270 | 280 | 290 | 300 |
| YSPYYNTIDD | LKDQIVDLTV | GNNKTLLDID | NTRMTLDDFR | IKFEMEQNLR | QGVDADINGL | RQVLDNLTME | KSDLEMQYET | LQEELMALKK | NHKEEMSQLT |
| 310 | 320 | 330 | 340 | 350 | 360 | 370 | 380 | 390 | 400 |
| GQNSGDVNVE | INVAPGKDLT | KTLNDMRQEY | EQLIAKNRKD | IENQYETQIT | QIEHEVSSSG | QEVQSSAKEV | TQLRHGVQEL | EIELQSQLSK | KAALEKSLED |
| 410 | 420 | 430 | 440 | 450 | 460 | 470 | 480 | 490 | 500 |
| TKNRYCGQLQ | MIQEQISNLE | AQITDVRQEI | ECQNQEYSLL | LSIKMRLEKE | IETYHNLLEG | GQEDFESSGA | GKIGLGGRGG | SGGSYGRGSR | GGSGGSYGGG |
| 510 | 520 | 530 | 540 | 550 | 560 | 570 | 580 | 590 | 600 |
| GSGGGYGGGS | GSRGGSGGSY | GGGSGSGGGS | GGGYGGGSGG | GHSGGSGGGH | SGGSGGNYGG | GSGSGGGSGG | GYGGGSGSRG | GSGGSHGGGS | GFGGESGGSY |
| 610 | 620 | 630 | | | | | | | |
| GGGEEASGSG | GGYGGGSGKS | SHS | | | | | | | |

MS/MS Peptide Matches

| Cmpd. | m/z meas. | z | Δ m/z [ppm] | Δ m/z [Da] | Rt [min] | Score | No. of Cmpds. | Site [%] | P | Sequence | Modification | Range | Type |
|---|---|---|---|---|---|---|---|---|---|---|---|---|---|
| 105 | 896.444 | 2 | 85.35 | 0.0765 | 12.20 | 166.48 | 5 | 0.0 | 0 | R.GGSGGSYGGGGSGGGYGGGSGSR.G | | 491-513 | CID |

#### Protein 11: Immunoglobulin lambda variable 3-21 OS=Homo sapiens GN=IGLV3-21 PE=1 SV=2

| | | | |
|---|---|---|---|
| **Accession:** | LV321_HUMAN | **Score:** | 96.7 |
| **Database:** | SwissProt | **Seq. Coverage [%]:** | 13.7 |
| **MW [kDa] / pI:** | 12.4 / 5.0 | **No. of Peptides:** | 1 |

| 10 | 20 | 30 | 40 | 50 | 60 | 70 | 80 | 90 | 100 |
|---|---|---|---|---|---|---|---|---|---|
| MAWTVLLLGL | LSHCTGSVTS | YVLTQPPSVS | VAPGQTARIT | CGGNNIGSKS | VHWYQQKPGQ | APVLVVYDDS | DRPSGIPERF | SGSNSGNTAT | LTISRVEAGD |
| 110 | 120 | | | | | | | | |
| EADYYCQVWD | SSSDHPT | | | | | | | | |

MS/MS Peptide Matches

| Cmpd. | m/z meas. | z | Δ m/z [ppm] | Δ m/z [Da] | Rt [min] | Score | No. of Cmpds. | Site [%] | P | Sequence | Modification | Range | Type |
|---|---|---|---|---|---|---|---|---|---|---|---|---|---|
| 436 | 806.959 | 2 | 73.23 | 0.0591 | 19.33 | 96.68 | 2 | 0.0 | 0 | R.FSGSNSGNTATLTISR.V | | 80-95 | CID |

#### Protein 12: Immunoglobulin kappa variable 3-15 OS=Homo sapiens GN=IGKV3-15 PE=1 SV=2

**Accession:** KV315_HUMAN
**Database:** SwissProt
**MW [kDa] / pI:** 12.5 / 5.1
**Score:** 65.5
**Seq. Coverage [%]:** 18.3
**No. of Peptides:** 1

| 10 | 20 | 30 | 40 | 50 | 60 | 70 | 80 | 90 | 100 |
|---|---|---|---|---|---|---|---|---|---|
| MEAPAQLLFL | LLLWLPDTTG | EIVMTQSPAT | LSVSPGERAT | LSCRASQSVS | SNLAWYQQKP | GQAPRLLIYG | ASTRATGIPA | RFSGSGSGTE | FTLTISSLQS |
| 110 | 120 | | | | | | | | |
| EDFAVYYCQQ | YNNWP | | | | | | | | |

MS/MS Peptide Matches

| Cmpd. | m/z meas. | z | Δ m/z [ppm] | Δ m/z [Da] | Rt [min] | Score | No. of Cmpds. | Site [%] | P | Sequence | Modification | Range | Type |
|---|---|---|---|---|---|---|---|---|---|---|---|---|---|
| 546 | 768.44 | 3 | 66.26 | 0.0509 | 21.41 | 65.52 | 3 | 0.0 | 0 | R.ASQSVSSNLAWYQQKPGQAPR.L | | 45-65 | CID |

#### Protein 13: Immunoglobulin kappa variable 4-1 OS=Homo sapiens GN=IGKV4-1 PE=1 SV=1

**Accession:** KV401_HUMAN
**Database:** SwissProt
**MW [kDa] / pI:** 13.4 / 5.1
**Score:** 46.2
**Seq. Coverage [%]:** 12.4
**No. of Peptides:** 1

| 10 | 20 | 30 | 40 | 50 | 60 | 70 | 80 | 90 | 100 |
|---|---|---|---|---|---|---|---|---|---|
| MVLQTQVFIS | LLLWISGAYG | DIVMTQSPDS | LAVSLGERAT | INCKSSQSVL | YSSNNKNYLA | WYQQKPGQPP | KLLIYWASTR | ESGVPDRFSG | SGSGTDFTLT |
| **110** | **120** | **130** | | | | | | | |
| ISSLQAEDVA | VYYCQQYYST | P | | | | | | | |

MS/MS Peptide Matches

| Cmpd. | m/z meas. | z | Δ m/z [ppm] | Δ m/z [Da] | Rt [min] | Score | No. of Cmpds. | Site [%] | P | Sequence | Modification | Range | Type |
|---|---|---|---|---|---|---|---|---|---|---|---|---|---|
| 608 | 909.546 | 2 | 83.23 | 0.0757 | 22.52 | 46.21 | 1 | 0.0 | 0 | K.NYLAWYQQKPGQPPK.L | | 57-71 | CID |

#### Protein 14: Immunoglobulin kappa variable 2D-28 OS=Homo sapiens GN=IGKV2D-28 PE=1 SV=2

**Accession:** KVD28_HUMAN
**Database:** SwissProt
**MW [kDa] / pI:** 12.9 / 5.6

**Score:** 34.8
**Seq. Coverage [%]:** 16.7
**No. of Peptides:** 1

| 10 | 20 | 30 | 40 | 50 | 60 | 70 | 80 | 90 | 100 |
|---|---|---|---|---|---|---|---|---|---|
| MRLPAQLLGL | LMLWVSGSSG | DIVMTQSPLS | LPVTPGEPAS | ISCRSSQSLL | HSNGYNYLDW | YLQKPGQSPQ | LLIYLGSNRA | SGVPDRFSGS | GSGTDFTLKI |
| **110** | **120** | **130** | | | | | | | |
| SRVEAEDVGV | YYCMQALQTP | | | | | | | | |

MS/MS Peptide Matches

| Cmpd. | m/z meas. | z | Δ m/z [ppm] | Δ m/z [Da] | Rt [min] | Score | No. of Cmpds. | Site [%] | P | Sequence | Modification | Range | Type |
|---|---|---|---|---|---|---|---|---|---|---|---|---|---|
| 606 | 993.521 | 2 | 39.46 | 0.0392 | 22.48 | 34.84 | 4 | 0.0 | 1 | R.ASGVPDRFSGSGSGTDFTLK.I | | 80-99 | CID |

#### Project Info

| | | | |
|---|---|---|---|
| **Name:** | Alagesan | **Date:** | Mar 5, 2019 |
| **Note:** | Nanobooster | | |

#### Sample Info & Protocols

| | |
|---|---|
| **Name:** | Nanobooster - IgG |
| **Date:** | Mar 5, 2019 |

#### Search Result Info

| | | | |
|---|---|---|---|
| **Search Result:** | IonTrap_Human_Swiss_CID_2019-03-05 17:55:08 | | |
| **Location:** | /Alagesan/Nanobooster - IgG/KA_IgG4 HD NB ACN CID_GA5_01_9088.mgf | | |
| **Search Method:** | IonTrap_Human_Swiss_CID | **Version:** | 1.2 |
| **Search Engine(s):** | Mascot, 2.6.0 | **Ident. Compound(s):** | 0/2513 |
| **Database(s):** | SwissProt, SwissProt_2017_04.fasta | | |
| **Note:** | | | |

#### Results

##### Protein 1: Immunoglobulin heavy constant gamma 4 OS=Homo sapiens GN=IGHG4 PE=1 SV=1

| | | | |
|---|---|---|---|
| **Accession:** | IGHG4_HUMAN | **Score:** | 1426.1 |
| **Database:** | SwissProt | **Seq. Coverage [%]:** | 65.7 |
| **MW [kDa] / pI:** | 35.9 / 7.2 | **No. of Peptides:** | 23 |
| **Modification(s):** | Carbamidomethyl, Oxidation | | |

```
        10         20         30         40         50         60         70         80         90        100
ASTKGPSVFP LAPCSRSTSE STAALGCLVK DYFPEPVTVS WNSGALTSGV HTFPAVLQSS GLYSLSSVVT VPSSSLGTKT YTCNVDHKPS NTKVDKRVES
       110        120        130        140        150        160        170        180        190        200
KYGPPCPSCP APEFLGGPSV FLFPPKPKDT LMISRTPEVT CVVVDVSQED PEVQFNWYVD GVEVHNAKTK PREEQFNSTY RVVSVLTVLH QDWLNGKEYK
       210        220        230        240        250        260        270        280        290        300
CKVSNKGLPS SIEKTISKAK GQPREPQVYT LPPSQEEMTK NQVSLTCLVK GFYPSDIAVE WESNGQPENN YKTTPPVLDS DGSFFLYSRL TVDKSRWQEG
       310        320        330
NVFSCSVMHE ALHNHYTQKS LSLSLGK
```

MS/MS Peptide Matches

| Cmpd. | m/z meas. | z | Δ m/z [ppm] | Δ m/z [Da] | Rt [min] | Score | No. of Cmpds. | Site [%] | P | Sequence | Modification | Range | Type |
|---|---|---|---|---|---|---|---|---|---|---|---|---|---|
| 679 | 712.41 | 2 | 72.30 | 0.0515 | 22.41 | 92.39 | 23 | 0.0 | 0 | R.STSESTAALGCLVK.D | Carbamidomethyl: 11 | 17-30 | CID |
| 175 | 833.42 | 2 | 638.16 | 0.5315 | 11.97 | 72.87 | 17 | 0.0 | 0 | K.TYTCNVDHKPSNTK.V | Carbamidomethyl: 4 | 80-93 | CID |
| 178 | 1004.075 | 2 | 90.99 | 0.0914 | 12.04 | 66.36 | 9 | 0.0 | 1 | K.TYTCNVDHKPSNTKVDK.R | Carbamidomethyl: 4 | 80-96 | CID |
| 1195 | 847.394 | 4 | -41.61 | -0.0353 | 31.41 | 33.67 | 2 | 0.0 | 1 | R.VESKYGPPCPSCPAPEFLGGPSVFLFPPKPK.D | Carbamidomethyl: 9, 12 | 98-128 | CID |

| Cmpd. | m/z meas. | z | Δ m/z [ppm] | Δ m/z [Da] | Rt [min] | Score | No. of Cmpds. | Site [%] | P | Sequence | Modification | Range | Type |
|---|---|---|---|---|---|---|---|---|---|---|---|---|---|
| 1275 | 982.239 | 3 | 422.77 | 0.4151 | 32.70 | 74.06 | 56 | 0.0 | 0 | K.YGPPCPSCPAPEFLGGPSVFLFPPKPK.D | Carbamidomethyl: 5, 8 | 102-128 | CID |
| 1295 | 941.056 | 4 | 353.08 | 0.3321 | 33.15 | 53.01 | 2 | 0.0 | 1 | K.YGPPCPSCPAPEFLGGPSVFLFPPKPKDTLMISR.T | Carbamidomethyl: 5, 8 | 102-135 | CID |
| 1252 | 945.088 | 4 | 386.80 | 0.3654 | 32.34 | 66.01 | 2 | 0.0 | 1 | K.YGPPCPSCPAPEFLGGPSVFLFPPKPKDTLMISR.T | Carbamidomethyl: 5, 8; Oxidation: 31 | 102-135 | CID |
| 1306 | 1152.374 | 4 | 278.41 | 0.3207 | 33.35 | 54.14 | 2 | 0.0 | 1 | K.DTLMISRTPEVTCVVVDVSQEDPEVQFNWYVDGVEVHNAK.T | Carbamidomethyl: 13 | 129-168 | CID |
| 1274 | 1156.338 | 4 | 247.40 | 0.2860 | 32.67 | 29.8 | 1 | 0.0 | 1 | K.DTLMISRTPEVTCVVVDVSQEDPEVQFNWYVDGVEVHNAK.T | Carbamidomethyl: 13; Oxidation: 4 | 129-168 | CID |
| 1239 | 1263.934 | 3 | 267.12 | 0.3375 | 32.11 | 148.22 | 19 | 0.0 | 0 | R.TPEVTCVVVDVSQEDPEVQFNWYVDGVEVHNAK.T | Carbamidomethyl: 6 | 136-168 | CID |
| 1248 | 905.106 | 2 | 662.37 | 0.5991 | 32.25 | 109.58 | 5 | 0.0 | 0 | R.VVSVLTVLHQDWLNGK.E | | 182-197 | CID |
| 1154 | 743.484 | 3 | 103.16 | 0.0767 | 30.67 | 105.43 | 6 | 0.0 | 1 | R.VVSVLTVLHQDWLNGKEYK.C | | 182-200 | CID |
| 619 | 772.446 | 3 | 81.92 | 0.0633 | 21.29 | 26.87 | 2 | 0.0 | 1 | K.GQPREPQVYTLPPSQEEMTK.N | | 221-240 | CID |
| 540 | 778.085 | 3 | 476.57 | 0.3706 | 19.77 | 18.67 | 1 | 0.0 | 1 | K.GQPREPQVYTLPPSQEEMTK.N | Oxidation: 18 | 221-240 | CID |
| 656 | 939.045 | 2 | 97.46 | 0.0915 | 21.93 | 41.23 | 10 | 0.0 | 0 | R.EPQVYTLPPSQEEMTK.N | | 225-240 | CID |
| 629 | 946.997 | 2 | 48.63 | 0.0461 | 21.46 | 22.15 | 3 | 0.0 | 0 | R.EPQVYTLPPSQEEMTK.N | Oxidation: 14 | 225-240 | CID |
| 1004 | 1007.591 | 3 | 412.69 | 0.4157 | 28.05 | 30.06 | 5 | 0.0 | 1 | R.EPQVYTLPPSQEEMTKNQVSLTCLVK.G | Carbamidomethyl: 23 | 225-250 | CID |
| 965 | 1012.863 | 3 | 351.62 | 0.3560 | 27.31 | 26.87 | 2 | 0.0 | 1 | R.EPQVYTLPPSQEEMTKNQVSLTCLVK.G | Carbamidomethyl: 23; Oxidation: 14 | 225-250 | CID |
| 1025 | 1273.178 | 2 | 478.30 | 0.6087 | 28.46 | 98.83 | 9 | 0.0 | 0 | K.GFYPSDIAVEWESNGQPENNYK.T | | 251-272 | CID |
| 1110 | 951.664 | 2 | 206.39 | 0.1964 | 29.92 | 100.22 | 46 | 0.0 | 0 | K.TTPPVLDSDGSFFLYSR.L | | 273-289 | CID |
| 745 | 762.471 | 4 | 484.81 | 0.3695 | 23.64 | 56.45 | 3 | 0.0 | 1 | K.SRWQEGNVFSCSVMHEALHNHYTQK.S | Carbamidomethyl: 11 | 295-319 | CID |
| 800 | 935.146 | 3 | 418.05 | 0.3908 | 24.57 | 66.68 | 2 | 0.0 | 0 | R.WQEGNVFSCSVMHEALHNHYTQK.S | Carbamidomethyl: 9 | 297-319 | CID |
| 638 | 940.487 | 3 | 425.64 | 0.4001 | 21.61 | 32.49 | 2 | 0.0 | 0 | R.WQEGNVFSCSVMHEALHNHYTQK.S | Carbamidomethyl: 9; Oxidation: 12 | 297-319 | CID |

BRUKER

##### Protein 2: Immunoglobulin kappa constant OS=Homo sapiens GN=IGKC PE=1 SV=2

| | | | |
|---|---|---|---|
| **Accession:** | IGKC_HUMAN | **Score:** | 1046.0 |
| **Database:** | SwissProt | **Seq. Coverage [%]:** | 88.8 |
| **MW [kDa] / pI:** | 11.8 / 6.1 | **No. of Peptides:** | 11 |
| **Modification(s):** | Carbamidomethyl | | |

```
        10         20         30         40         50         60         70         80         90        100
RTVAAPSVFI FPPSDEQLKS GTASVVCLLN NFYPREAKVQ WKVDNALQSG NSQESVTEQD SKDSTYSLSS TLTLSKADYE KHKVYACEVT HQGLSSPVTK
       110
SFNRGEC
```

**MS/MS Peptide Matches**

| Cmpd. | m/z meas. | z | Δ m/z [ppm] | Δ m/z [Da] | Rt [min] | Score | No. of Cmpds. | Site [%] | P | Sequence | Modification | Range | Type |
|---|---|---|---|---|---|---|---|---|---|---|---|---|---|
| 1045 | 1051.627 | 2 | 56.42 | 0.0593 | 28.84 | 49.56 | 3 | 0.0 | 1 | -.RTVAAPSVFIFPPSDEQLK.S | | 1-19 | CID |
| 1164 | 973.656 | 2 | 142.66 | 0.1389 | 30.88 | 85.73 | 24 | 0.0 | 0 | R.TVAAPSVFIFPPSDEQLK.S | | 2-19 | CID |
| 1605 | 1242.702 | 3 | 318.52 | 0.3957 | 38.34 | 77.86 | 14 | 0.0 | 1 | R.TVAAPSVFIFPPSDEQLKSGTASVVCLLNNFYPR.E | Carbamidomethyl: 26 | 2-35 | CID |
| 1347 | 900.021 | 2 | 633.44 | 0.5697 | 34.02 | 26.22 | 4 | 0.0 | 0 | K.SGTASVVCLLNNFYPR.E | Carbamidomethyl: 8 | 20-35 | CID |
| 1183 | 709.411 | 3 | 69.80 | 0.0495 | 31.23 | 114.92 | 3 | 0.0 | 1 | K.SGTASVVCLLNNFYPREAK.V | Carbamidomethyl: 8 | 20-38 | CID |
| 609 | 893.521 | 3 | 477.16 | 0.4262 | 21.07 | 115.26 | 10 | 0.0 | 1 | K.VQWKVDNALQSGNSQESVTEQDSK.D | | 39-62 | CID |
| 393 | 1069.032 | 2 | 509.13 | 0.5440 | 16.39 | 150.54 | 31 | 0.0 | 0 | K.VDNALQSGNSQESVTEQDSK.D | | 43-62 | CID |
| 801 | 1207.533 | 3 | 241.63 | 0.2917 | 24.57 | 94.04 | 2 | 0.0 | 1 | K.VDNALQSGNSQESVTEQDSKDSTYSLSSTLTLSK.A | | 43-76 | CID |
| 811 | 751.968 | 2 | 113.24 | 0.0851 | 24.76 | 79.7 | 4 | 0.0 | 0 | K.DSTYSLSSTLTLSK.A | | 63-76 | CID |
| 407 | 714.45 | 3 | 118.82 | 0.0849 | 16.92 | 135.48 | 9 | 0.0 | 1 | K.HKVYACEVTHQGLSSPVTK.S | Carbamidomethyl: 6 | 82-100 | CID |
| 521 | 938.553 | 2 | 91.53 | 0.0859 | 19.40 | 116.73 | 13 | 0.0 | 0 | K.VYACEVTHQGLSSPVTK.S | Carbamidomethyl: 4 | 84-100 | CID |

#### Protein 3: Immunoglobulin kappa light chain OS=Homo sapiens PE=1 SV=1

| | | | |
|---|---|---|---|
| **Accession:** | IGK_HUMAN | **Score:** | 950.6 |
| **Database:** | SwissProt | **Seq. Coverage [%]:** | 43.9 |
| **MW [kDa] / pI:** | 23.4 / 6.9 | **No. of Peptides:** | 9 |
| **Modification(s):** | Carbamidomethyl | | |

```
        10         20         30         40         50         60         70         80         90        100
DIQMTQSPST LSASVGDRVT ITCRASQSIN TWLAWYQQKP GKAPKLLMYK ASSLESGVPS RFIGSGSGTE FTLTISSLQP DDFATYYCQQ YNSDSKMFGQ
       110        120        130        140        150        160        170        180        190        200
GTKVEVKGTV AAPSVFIFPP SDEQLKSGTA SVVCLLNNFY PREAKVQWKV DNALQSGNSQ ESVTEQDSKD STYSLSSTLT LSKADYEKHK VYACEVTHQG
       210        220
LSSPVTKSFN RGEC
```

**MS/MS Peptide Matches**

| Cmpd. | m/z meas. | z | Δ m/z [ppm] | Δ m/z [Da] | Rt [min] | Score | No. of Cmpds. | Site [%] | P | Sequence | Modification | Range | Type |
|---|---|---|---|---|---|---|---|---|---|---|---|---|---|
| 670 | 946.986 | 2 | 33.21 | 0.0314 | 22.22 | 117.67 | 5 | 0.0 | 0 | -.DIQMTQSPSTLSASVGDR.V | | 1-18 | CID |
| 1347 | 900.021 | 2 | 633.44 | 0.5697 | 34.02 | 26.22 | 4 | 0.0 | 0 | K.SGTASVVCLLNNFYPR.E | Carbamidomethyl: 8 | 127-142 | CID |
| 1183 | 709.411 | 3 | 69.80 | 0.0495 | 31.23 | 114.92 | 3 | 0.0 | 1 | K.SGTASVVCLLNNFYPREAK.V | Carbamidomethyl: 8 | 127-145 | CID |
| 609 | 893.521 | 3 | 477.16 | 0.4262 | 21.07 | 115.26 | 10 | 0.0 | 1 | K.VQWKVDNALQSGNSQESVTEQDSK.D | | 146-169 | CID |
| 393 | 1069.032 | 2 | 509.13 | 0.5440 | 16.39 | 150.54 | 31 | 0.0 | 0 | K.VDNALQSGNSQESVTEQDSK.D | | 150-169 | CID |
| 801 | 1207.533 | 3 | 241.63 | 0.2917 | 24.57 | 94.04 | 2 | 0.0 | 1 | K.VDNALQSGNSQESVTEQDSKDSTYSLSSTLTLSK.A | | 150-183 | CID |
| 811 | 751.968 | 2 | 113.24 | 0.0851 | 24.76 | 79.7 | 4 | 0.0 | 0 | K.DSTYSLSSTLTLSK.A | | 170-183 | CID |
| 407 | 714.45 | 3 | 118.82 | 0.0849 | 16.92 | 135.48 | 9 | 0.0 | 1 | K.HKVYACEVTHQGLSSPVTK.S | Carbamidomethyl: 6 | 189-207 | CID |
| 521 | 938.553 | 2 | 91.53 | 0.0859 | 19.40 | 116.73 | 13 | 0.0 | 0 | K.VYACEVTHQGLSSPVTK.S | Carbamidomethyl: 4 | 191-207 | CID |

BRUKER

#### Protein 4: Keratin, type II cytoskeletal 1 OS=Homo sapiens GN=KRT1 PE=1 SV=6

| | | | |
|---|---|---|---|
| **Accession:** | K2C1_HUMAN | **Score:** | 818.3 |
| **Database:** | SwissProt | **Seq. Coverage [%]:** | 17.2 |
| **MW [kDa] / pI:** | 66.0 / 8.2 | **No. of Peptides:** | 6 |
| **Modification(s):** | Carbamidomethyl | | |

| 10 | 20 | 30 | 40 | 50 | 60 | 70 | 80 | 90 | 100 |
|---|---|---|---|---|---|---|---|---|---|
| MSRQFSSRSG | YRSGGGFSSG | SAGIINYQRR | TTSSSTRRSG | GGGGRFSSCG | GGGGSFGAGG | GFGSRSLVNL | GGSKSISISV | ARGGGRGSGF | GGGYGGGGFG |
| 110 | 120 | 130 | 140 | 150 | 160 | 170 | 180 | 190 | 200 |
| GGGFGGGGFG | GGGIGGGGFG | GFGSGGGGFG | GGGFGGGGYG | GGYGPVCPPG | GIQEVTINQS | LLQPLNVEID | PEIQKVKSRE | REQIKSLNNQ | FASFIDKVRF |
| 210 | 220 | 230 | 240 | 250 | 260 | 270 | 280 | 290 | 300 |
| LEQQNQVLQT | KWELLQQVDT | STRTHNLEPY | FESFINNLRR | RVDQLKSDQS | RLDSELKNMQ | DMVEDYRNKY | EDEINKRTNA | ENEFVTIKKD | VDGAYMTKVD |
| 310 | 320 | 330 | 340 | 350 | 360 | 370 | 380 | 390 | 400 |
| LQAKLDNLQQ | EIDFLTALYQ | AELSQMQTQI | SETNVILSMD | NNRSLDLDSI | IAEVKAQYED | IAQKSKAEAE | SLYQSKYEEL | QITAGRHGDS | VRNSKIEISE |
| 410 | 420 | 430 | 440 | 450 | 460 | 470 | 480 | 490 | 500 |
| LNRVIQRLRS | EIDNVKKQIS | NLQQSISDAE | QRGENALKDA | KNKLNDLEDA | LQQAKEDLAR | LLRDYQELMN | TKLALDLEIA | TYRTLLEGEE | SRMSGECAPN |
| 510 | 520 | 530 | 540 | 550 | 560 | 570 | 580 | 590 | 600 |
| VSVSVSTSHT | TISGGGSRGG | GGGGYGSGGS | SYGSGGGSYG | SGGGGGGGRG | SYGSGGSSYG | SGGGSYGSGG | GGGGHGSYGS | GSSSGGYRGG | SGGGGGGSSG |
| 610 | 620 | 630 | 640 | 650 | | | | | |
| GRGSGGGSSG | GSIGGRGSSS | GGVKSSGGSS | SVKFVSTTYS | GVTR | | | | | |

MS/MS Peptide Matches

| Cmpd. | m/z meas. | z | Δ m/z [ppm] | Δ m/z [Da] | Rt [min] | Score | No. of Cmpds. | Site [%] | P | Sequence | Modification | Range | Type |
|---|---|---|---|---|---|---|---|---|---|---|---|---|---|
| 562 | 883.421 | 2 | 56.61 | 0.0500 | 20.17 | 138.19 | 3 | 0.0 | 0 | R.FSSCGGGGGSFGAGGGFGSR.S | Carbamidomethyl: 4 | 46-65 | CID |
| 550 | 738.487 | 2 | 122.87 | 0.0907 | 19.97 | 98.49 | 4 | 0.0 | 0 | R.FLEQQNQVLQTK.W | | 200-211 | CID |
| 1080 | 978.59 | 3 | 422.26 | 0.4130 | 29.43 | 107.15 | 3 | 0.0 | 1 | R.FLEQQNQVLQTKWELLQQVDTSTR.T | | 200-223 | CID |

| Cmpd. | m/z meas. | z | Δ m/z [ppm] | Δ m/z [Da] | Rt [min] | Score | No. of Cmpds. | Site [%] | P | Sequence | Modification | Range | Type |
|---|---|---|---|---|---|---|---|---|---|---|---|---|---|
| 1569 | 783.852 | 3 | 137.40 | 0.1077 | 37.72 | 160.96 | 3 | 0.0 | 1 | R.SLDLDSIIAEVKAQYEDIAQK.S | | 344-364 | CID |
| 639 | 859.015 | 2 | 99.90 | 0.0858 | 21.64 | 118.4 | 3 | 0.0 | 0 | K.QISNLQQSISDAEQR.G | | 418-432 | CID |
| 293 | 1192.928 | 2 | 376.06 | 0.4484 | 14.37 | 195.16 | 4 | 0.0 | 0 | R.GGGGGGYGSGGSSYGSGGGSYGSGGGGGG GR.G | | 519-549 | CID |

##### Protein 5: Keratin, type II cytoskeletal 2 epidermal OS=Homo sapiens GN=KRT2 PE=1 SV=2

**Accession:** K22E_HUMAN
**Database:** SwissProt
**MW [kDa] / pI:** 65.4 / 8.1
**Score:** 623.0
**Seq. Coverage [%]:** 15.3
**No. of Peptides:** 5

| 10 | 20 | 30 | 40 | 50 | 60 | 70 | 80 | 90 | 100 |
|---|---|---|---|---|---|---|---|---|---|
| MSCQISCKSR | GRGGGGGGFR | GFSSGSAVVS | GGSRRSTSSF | SCLSRHGGGG | GGFGGGGFGS | RSLVGLGGTK | SISISVAGGG | GGFGAAGGFG | GRGGGFGGGS |
| **110** | **120** | **130** | **140** | **150** | **160** | **170** | **180** | **190** | **200** |
| SFGGGSGFSG | GGFGGGGFGG | GRFGGFGGPG | GVGGLGGPGG | FGPGGYPGGI | HEVSVNQSLL | QPLNVKVDPE | IQNVKAQERE | QIKTLNNKFA | SFIDKVRFLE |
| **210** | **220** | **230** | **240** | **250** | **260** | **270** | **280** | **290** | **300** |
| QQNQVLQTKW | ELLQQMNVGT | RPINLEPIFQ | GYIDSLKRYL | DGLTAERTSQ | NSELNNMQDL | VEDYKKKYED | EINKRTAAEN | DFVTLKKDVD | NAYMIKVELQ |
| **310** | **320** | **330** | **340** | **350** | **360** | **370** | **380** | **390** | **400** |
| SKVDLLNQEI | EFLKVLYDAE | ISQIHQSVTD | TNVILSMDNS | RNLDLDSIIA | EVKAQYEEIA | QRSKEEAEAL | YHSKYEELQV | TVGRHGDSLK | EIKIEISELN |
| **410** | **420** | **430** | **440** | **450** | **460** | **470** | **480** | **490** | **500** |
| RVIQRLQGEI | AHVKKQCKNV | QDAIADAEQR | GEHALKDARN | KLNDLEEALQ | QAKEDLARLL | RDYQELMNVK | LALDVEIATY | RKLLEGEECR | MSGDLSSNVT |
| **510** | **520** | **530** | **540** | **550** | **560** | **570** | **580** | **590** | **600** |
| VSVTSSTISS | NVASKAAFGG | SGGRGSSSGG | GYSSGSSSYG | SGGRQSGSRG | GSGGGGSISG | GGYGSGGGSG | GRYGSGGGSK | GGSISGGGYG | SGGGKHSSGG |
| **610** | **620** | **630** | **640** | | | | | | |
| GSRGGSSSGG | GYGSGGGGSS | SVKGSSGEAF | GSSVTFSFR | | | | | | |

MS/MS Peptide Matches

| Cmpd. | m/z meas. | z | Δ m/z [ppm] | Δ m/z [Da] | Rt [min] | Score | No. of Cmpds. | Site [%] | P | Sequence | Modification | Range | Type |
|---|---|---|---|---|---|---|---|---|---|---|---|---|---|
| 905 | 920.029 | 2 | 74.10 | 0.0682 | 26.35 | 198.04 | 2 | 0.0 | 0 | K.SISISVAGGGGGFGAAGGFGGR.G | | 71-92 | CID |
| 550 | 738.487 | 2 | 122.87 | 0.0907 | 19.97 | 98.49 | 4 | 0.0 | 0 | R.FLEQQNQVLQTK.W | | 198-209 | CID |
| 1635 | 806.805 | 3 | 61.71 | 0.0498 | 38.82 | 157.39 | 3 | 0.0 | 1 | R.NLDLDSIIAEVKAQYEEIAQR.S | | 342-362 | CID |
| 204 | 870.871 | 2 | 16.74 | 0.0146 | 12.60 | 26.11 | 2 | 0.0 | 0 | R.GSSSGGGYSSGSSSYGSGGR.Q | | 525-544 | CID |
| 231 | 871.445 | 2 | 77.05 | 0.0671 | 13.18 | 143.01 | 3 | 0.0 | 0 | R.GGSGGGGSISGGGYGSGGGSGGR.Y | | 550-572 | CID |

#### Protein 6: Immunoglobulin lambda constant 3 OS=Homo sapiens GN=IGLC3 PE=1 SV=1

| | | | |
|---|---|---|---|
| **Accession:** | IGLC3_HUMAN | **Score:** | 549.0 |
| **Database:** | SwissProt | **Seq. Coverage [%]:** | 74.5 |
| **MW [kDa] / pI:** | 11.3 / 6.9 | **No. of Peptides:** | 7 |
| **Modification(s):** | Carbamidomethyl | | |

10 20 30 40 50 60 70 80 90 100
GQPKAAPSVT LFPPSSEELQ ANKATLVCLI SDFYPGAVTV AWKADSSPVK AGVETTTPSK QSNNKYAASS YLSLTPEQWK SHKSYSCQVT HEGSTVEKTV
110
APTECS

MS/MS Peptide Matches

| Cmpd. | m/z meas. | z | Δ m/z [ppm] | Δ m/z [Da] | Rt [min] | Score | No. of Cmpds. | Site [%] | P | Sequence | Modification | Range | Type |
|---|---|---|---|---|---|---|---|---|---|---|---|---|---|
| 926 | 993.546 | 2 | 33.66 | 0.0334 | 26.70 | 78.41 | 7 | 0.0 | 0 | K.AAPSVTLFPPSSEELQANK.A | | 5-23 | CID |
| 328 | 838.003 | 2 | 86.08 | 0.0721 | 15.08 | 62.52 | 3 | 0.0 | 1 | K.ADSSPVKAGVETTTPSK.Q | | 44-60 | CID |
| 127 | 781.554 | 2 | 204.18 | 0.1595 | 10.92 | 56.45 | 3 | 0.0 | 1 | K.AGVETTTPSKQSNNK.Y | | 51-65 | CID |
| 892 | 772.461 | 3 | 102.80 | 0.0794 | 26.12 | 65.51 | 5 | 0.0 | 1 | K.QSNNKYAASSYLSLTPEQWK.S | | 61-80 | CID |
| 990 | 872.542 | 2 | 124.88 | 0.1089 | 27.78 | 92.69 | 5 | 0.0 | 0 | K.YAASSYLSLTPEQWK.S | | 66-80 | CID |
| 288 | 856.498 | 2 | 134.01 | 0.1148 | 14.28 | 103.37 | 6 | 0.0 | 0 | K.SYSCQVTHEGSTVEK.T | Carbamidomethyl: 4 | 84-98 | CID |
| 499 | 1279.518 | 2 | 355.97 | 0.4553 | 18.95 | 90.04 | 8 | 0.0 | 1 | K.SYSCQVTHEGSTVEKTVAPTECS.- | Carbamidomethyl: 4, 22 | 84-106 | CID |

| Cmpd. | m/z meas. | z | Δ m/z [ppm] | Δ m/z [Da] | Rt [min] | Score | No. of Cmpds. | Site [%] | P | Sequence | Modification | Range | Type |
|---|---|---|---|---|---|---|---|---|---|---|---|---|---|

**Protein 7: Immunoglobulin heavy constant gamma 1 OS=Homo sapiens GN=IGHG1 PE=1 SV=1**

| | | | |
|---|---|---|---|
| **Accession:** | IGHG1_HUMAN | **Score:** | 383.6 |
| **Database:** | SwissProt | **Seq. Coverage [%]:** | 17.6 |
| **MW [kDa] / pI:** | 36.1 / 8.5 | **No. of Peptides:** | 4 |

| 10 | 20 | 30 | 40 | 50 | 60 | 70 | 80 | 90 | 100 |
|---|---|---|---|---|---|---|---|---|---|
| ASTKGPSVFP | LAPSSKSTSG | GTAALGCLVK | DYFPEPVTVS | WNSGALTSGV | HTFPAVLQSS | GLYSLSSVVT | VPSSSLGTQT | YICNVNHKPS | NTKVDKKVEP |
| **110** | **120** | **130** | **140** | **150** | **160** | **170** | **180** | **190** | **200** |
| KSCDKTHTCP | PCPAPELLGG | PSVFLFPPKP | KDTLMISRTP | EVTCVVVDVS | HEDPEVKFNW | YVDGVEVHNA | KTKPREEQYN | STYRVVSVLT | VLHQDWLNGK |
| **210** | **220** | **230** | **240** | **250** | **260** | **270** | **280** | **290** | **300** |
| EYKCKVSNKA | LPAPIEKTIS | KAKGQPREPQ | VYTLPPSRDE | LTKNQVSLTC | LVKGFYPSDI | AVEWESNGQP | ENNYKTTPPV | LDSDGSFFLY | SKLTVDKSRW |
| **310** | **320** | **330** | **340** | | | | | | |
| QQGNVFSCSV | MHEALHNHYT | QKSLSLSPGK | | | | | | | |

MS/MS Peptide Matches

| Cmpd. | m/z meas. | z | Δ m/z [ppm] | Δ m/z [Da] | Rt [min] | Score | No. of Cmpds. | Site [%] | P | Sequence | Modification | Range | Type |
|---|---|---|---|---|---|---|---|---|---|---|---|---|---|
| 1248 | 905.106 | 2 | 662.37 | 0.5991 | 32.25 | 109.58 | 5 | 0.0 | 0 | R.VVSVLTVLHQDWLNGK.E | | 185-200 | CID |
| 1154 | 743.484 | 3 | 103.16 | 0.0767 | 30.67 | 105.43 | 6 | 0.0 | 1 | R.VVSVLTVLHQDWLNGKEYK.C | | 185-203 | CID |
| 1025 | 1273.178 | 2 | 478.30 | 0.6087 | 28.46 | 98.83 | 9 | 0.0 | 0 | K.GFYPSDIAVEWESNGQPENNYK.T | | 254-275 | CID |
| 1092 | 937.548 | 2 | 89.02 | 0.0835 | 29.62 | 69.72 | 3 | 0.0 | 0 | K.TTPPVLDSDGSFFLYSK.L | | 276-292 | CID |

#### Protein 8: Keratin, type I cytoskeletal 10 OS=Homo sapiens GN=KRT10 PE=1 SV=6

| | | | |
|---|---|---|---|
| **Accession:** | K1C10_HUMAN | **Score:** | 322.4 |
| **Database:** | SwissProt | **Seq. Coverage [%]:** | 9.9 |
| **MW [kDa] / pI:** | 58.8 / 5.1 | **No. of Peptides:** | 3 |

| 10 | 20 | 30 | 40 | 50 | 60 | 70 | 80 | 90 | 100 |
|---|---|---|---|---|---|---|---|---|---|
| MSVRYSSSKH | YSSSRSGGGG | GGGGCGGGGG | VSSLRISSSK | GSLGGGFSSG | GFSGGSFSRG | SSGGGCFGGS | SGGYGGLGGF | GGGSFRGSYG | SSSFGGSYGG |
| **110** | **120** | **130** | **140** | **150** | **160** | **170** | **180** | **190** | **200** |
| IFGGGSFGGG | SFGGGSFGGG | GFGGGGFGGG | FGGGFGGDGG | LLSGNEKVTM | QNLNDRLASY | LDKVRALEES | NYELEGKIKE | WYEKHGNSHQ | GEPRDYSKYY |
| **210** | **220** | **230** | **240** | **250** | **260** | **270** | **280** | **290** | **300** |
| KTIDDLKNQI | LNLTTDNANI | LLQIDNARLA | ADDFRLKYEN | EVALRQSVEA | DINGLRRVLD | ELTLTKADLE | MQIESLTEEL | AYLKKNHEEE | MKDLRNVSTG |
| **310** | **320** | **330** | **340** | **350** | **360** | **370** | **380** | **390** | **400** |
| DVNVEMNAAP | GVDLTQLLNN | MRSQYEQLAE | QNRKDAEAWF | NEKSKELTTE | IDNNIEQISS | YKSEITELRR | NVQALEIELQ | SQLALKQSLE | ASLAETEGRY |
| **410** | **420** | **430** | **440** | **450** | **460** | **470** | **480** | **490** | **500** |
| CVQLSQIQAQ | ISALEEQLQQ | IRAETECQNT | EYQQLLDIKI | RLENEIQTYR | SLLEGEGSSG | GGGRGGGSFG | GGYGGGSSGG | GSSGGGHGGG | HGGSSGGGYG |
| **510** | **520** | **530** | **540** | **550** | **560** | **570** | **580** | **590** | |
| GGSSGGGSSG | GGYGGGSSSG | GHGGSSSGGY | GGGSSGGGGG | GYGGGSSGGG | SSSGGGYGGG | SSSGGHKSSS | SGSVGESSSK | GPRY | |

##### MS/MS Peptide Matches

| Cmpd. | m/z meas. | z | Δ m/z [ppm] | Δ m/z [Da] | Rt [min] | Score | No. of Cmpds. | Site [%] | P | Sequence | Modification | Range | Type |
|---|---|---|---|---|---|---|---|---|---|---|---|---|---|
| 755 | 854.481 | 2 | 106.85 | 0.0913 | 23.84 | 152.96 | 3 | 0.0 | 0 | K.GSLGGGFSSGGFSGGSFSR.G | | 41-59 | CID |
| 1393 | 1436.61 | 2 | -62.66 | -0.0900 | 34.75 | 133.55 | 3 | 0.0 | 0 | R.NVSTGDVNVEMNAAPGVDLTQLLNNMR.S | | 296-322 | CID |
| 322 | 747.427 | 2 | 75.22 | 0.0562 | 14.95 | 35.9 | 4 | 0.0 | 1 | R.SQYEQLAEQNRK.D | | 323-334 | CID |

#### Protein 9: Keratin, type I cytoskeletal 9 OS=Homo sapiens GN=KRT9 PE=1 SV=3

| | | | |
|---|---|---|---|
| **Accession:** | K1C9_HUMAN | **Score:** | 156.5 |
| **Database:** | SwissProt | **Seq. Coverage [%]:** | 3.7 |
| **MW [kDa] / pI:** | 62.0 / 5.1 | **No. of Peptides:** | 1 |

| 10 | 20 | 30 | 40 | 50 | 60 | 70 | 80 | 90 | 100 |
|---|---|---|---|---|---|---|---|---|---|
| MSCRQFSSSY | LSRSGGGGGG | GLGSGGSIRS | SYSRFSSSGG | GGGGGRFSSS | SGYGGGSSRV | CGRGGGGSFG | YSYGGGSGGG | FSASSLGGGF | GGGSRGFGGA |
| **110** | **120** | **130** | **140** | **150** | **160** | **170** | **180** | **190** | **200** |
| SGGGYSSSGG | FGGGFGGGSG | GGFGGGYGSG | FGGFGGFGGG | AGGGDGGILT | ANEKSTMQEL | NSRLASYLDK | VQALEEANND | LENKIQDWYD | KKGPAAIQKN |
| **210** | **220** | **230** | **240** | **250** | **260** | **270** | **280** | **290** | **300** |
| YSPYYNTIDD | LKDQIVDLTV | GNNKTLLDID | NTRMTLDDFR | IKFEMEQNLR | QGVDADINGL | RQVLDNLTME | KSDLEMQYET | LQEELMALKK | NHKEEMSQLT |
| **310** | **320** | **330** | **340** | **350** | **360** | **370** | **380** | **390** | **400** |
| GQNSGDVNVE | INVAPGKDLT | KTLNDMRQEY | EQLIAKNRKD | IENQYETQIT | QIEHEVSSSG | QEVQSSAKEV | TQLRHGVQEL | EIELQSQLSK | KAALEKSLED |
| **410** | **420** | **430** | **440** | **450** | **460** | **470** | **480** | **490** | **500** |
| TKNRYCGQLQ | MIQEQISNLE | AQITDVRQEI | ECQNQEYSLL | LSIKMRLEKE | IETYHNLLEG | GQEDFESSGA | GKIGLGGRGG | SGGSYGRGSR | GGSGGSYGGG |
| **510** | **520** | **530** | **540** | **550** | **560** | **570** | **580** | **590** | **600** |
| GSGGGYGGGS | GSRGGSGGSY | GGGSGSGGGS | GGGYGGGSGG | GHSGGSGGGH | SGGSGGNYGG | GSGSGGGSGG | GYGGGSGSRG | GSGGSHGGGS | GFGGESGGSY |
| **610** | **620** | **630** | | | | | | | |
| GGGEEASGSG | GGYGGGSGKS | SHS | | | | | | | |

MS/MS Peptide Matches

| Cmpd. | m/z meas. | z | Δ m/z [ppm] | Δ m/z [Da] | Rt [min] | Score | No. of Cmpds. | Site [%] | P | Sequence | Modification | Range | Type |
|---|---|---|---|---|---|---|---|---|---|---|---|---|---|
| 191 | 896.434 | 2 | 74.20 | 0.0665 | 12.31 | 156.5 | 5 | 0.0 | 0 | R.GGSGGSYGGGGSGGGYGGGSGSR.G | | 491-513 | CID |

#### Protein 10: Immunoglobulin lambda variable 3-21 OS=Homo sapiens GN=IGLV3-21 PE=1 SV=2

| | | | |
|---|---|---|---|
| **Accession:** | LV321_HUMAN | **Score:** | 114.3 |
| **Database:** | SwissProt | **Seq. Coverage [%]:** | 13.7 |
| **MW [kDa] / pI:** | 12.4 / 5.0 | **No. of Peptides:** | 1 |

```
        10         20         30         40         50         60         70         80         90        100
MAWTVLLLGL LSHCTGSVTS YVLTQPPSVS VAPGQTARIT CGGNNIGSKS VHWYQQKPGQ APVLVVYDDS DRPSGIPERF SGSNSGNTAT LTISRVEAGD
       110        120
EADYYCQVWD SSSDHPT
```

MS/MS Peptide Matches

| Cmpd. | m/z meas. | z | Δ m/z [ppm] | Δ m/z [Da] | Rt [min] | Score | No. of Cmpds. | Site [%] | P | Sequence | Modification | Range | Type |
|---|---|---|---|---|---|---|---|---|---|---|---|---|---|
| 518 | 806.972 | 2 | 89.35 | 0.0721 | 19.35 | 114.28 | 2 | 0.0 | 0 | R.FSGSNSGNTATLTISR.V | | 80-95 | CID |

#### Protein 11: Immunoglobulin kappa variable 3-15 OS=Homo sapiens GN=IGKV3-15 PE=1 SV=2

| | | | |
|---|---|---|---|
| **Accession:** | KV315_HUMAN | **Score:** | 61.3 |
| **Database:** | SwissProt | **Seq. Coverage [%]:** | 18.3 |
| **MW [kDa] / pI:** | 12.5 / 5.1 | **No. of Peptides:** | 1 |

```
        10         20         30         40         50         60         70         80         90        100
MEAPAQLLFL LLLWLPDTTG EIVMTQSPAT LSVSPGERAT LSCRASQSVS SNLAWYQQKP GQAPRLLIYG ASTRATGIPA RFSGSGSGTE FTLTISSLQS
       110        120
EDFAVYYCQQ YNNWP
```

MS/MS Peptide Matches

| Cmpd. | m/z meas. | z | Δ m/z [ppm] | Δ m/z [Da] | Rt [min] | Score | No. of Cmpds. | Site [%] | P | Sequence | Modification | Range | Type |
|---|---|---|---|---|---|---|---|---|---|---|---|---|---|
| 630 | 768.448 | 3 | 76.67 | 0.0589 | 21.49 | 61.3 | 3 | 0.0 | 0 | R.ASQSVSSNLAWYQQKPGQAPR.L | | 45-65 | CID |

#### Protein 12: Immunoglobulin kappa variable 2D-28 OS=Homo sapiens GN=IGKV2D-28 PE=1 SV=2

| | | | |
|---|---|---|---|
| **Accession:** | KVD28_HUMAN | **Score:** | 55.6 |
| **Database:** | SwissProt | **Seq. Coverage [%]:** | 16.7 |
| **MW [kDa] / pI:** | 12.9 / 5.6 | **No. of Peptides:** | 1 |

| 10 | 20 | 30 | 40 | 50 | 60 | 70 | 80 | 90 | 100 |
|---|---|---|---|---|---|---|---|---|---|
| MRLPAQLLGL | LMLWVSGSSG | DIVMTQSPLS | LPVTPGEPAS | ISCRSSQSLL | HSNGYNYLDW | YLQKPGQSPQ | LLIYLGSNRA | SGVPDRFSGS | GSGTDFTLKI |
| **110** | **120** | **130** | | | | | | | |
| SRVEAEDVGV | YYCMQALQTP | | | | | | | | |

MS/MS Peptide Matches

| Cmpd. | m/z meas. | z | Δ m/z [ppm] | Δ m/z [Da] | Rt [min] | Score | No. of Cmpds. | Site [%] | P | Sequence | Modification | Range | Type |
|---|---|---|---|---|---|---|---|---|---|---|---|---|---|
| 678 | 993.497 | 2 | 15.31 | 0.0152 | 22.41 | 55.58 | 3 | 0.0 | 1 | R.ASGVPDRFSGSGSGTDFTLK.I | | 80-99 | CID |

#### Protein 13: Immunoglobulin kappa variable 4-1 OS=Homo sapiens GN=IGKV4-1 PE=1 SV=1

| | | | |
|---|---|---|---|
| **Accession:** | KV401_HUMAN | **Score:** | 40.2 |
| **Database:** | SwissProt | **Seq. Coverage [%]:** | 12.4 |
| **MW [kDa] / pI:** | 13.4 / 5.1 | **No. of Peptides:** | 1 |

| 10 | 20 | 30 | 40 | 50 | 60 | 70 | 80 | 90 | 100 |
|---|---|---|---|---|---|---|---|---|---|
| MVLQTQVFIS | LLLWISGAYG | DIVMTQSPDS | LAVSLGERAT | INCKSSQSVL | YSSNNKNYLA | WYQQKPGQPP | KLLIYWASTR | ESGVPDRFSG | SGSGTDFTLT |
| **110** | **120** | **130** | | | | | | | |
| ISSLQAEDVA | VYYCQQYYST | P | | | | | | | |

MS/MS Peptide Matches

| Cmpd. | m/z meas. | z | Δ m/z [ppm] | Δ m/z [Da] | Rt [min] | Score | No. of Cmpds. | Site [%] | P | Sequence | Modification | Range | Type |
|---|---|---|---|---|---|---|---|---|---|---|---|---|---|
| 684 | 909.548 | 2 | 85.43 | 0.0777 | 22.52 | 40.22 | 1 | 0.0 | 0 | K.NYLAWYQQKPGQPPK.L | | 57-71 | CID |
